# Factors Affecting Response to Recurrent Genomic Selection in Soybeans

**DOI:** 10.1101/2020.02.14.949008

**Authors:** Vishnu Ramasubramanian, William D Beavis

## Abstract

Herein we report the impacts of applying five selection methods across 40 cycles of recurrent selection and identify interactions among factors that affect genetic responses in sets of simulated families of recombinant inbred lines derived from 21 homozygous soybean lines. Our use of recurrence equation to model response from recurrent selection allowed us to estimate the half-lives, asymptotic limits to recurrent selection for purposes of assessing the rates of response and future genetic potential of populations under selection. The simulated factors include selection methods, training sets, and selection intensity that are under the control of the plant breeder as well as genetic architecture and heritability. A factorial design to examine and analyze the main and interaction effects of these factors showed that both the rates of genetic improvement in the early cycles and limits to genetic improvement in the later cycles are significantly affected by interactions among all factors. Some consistent trends are that genomic selection methods provide greater initial rates of genetic improvement (per cycle) than phenotypic selection, but phenotypic selection provides the greatest long term responses in these closed genotypic systems. Model updating with training sets consisting of data from prior cycles of selection significantly improved prediction accuracy and genetic response with three parametric genomic prediction models. Ridge Regression, if updated with training sets consisting of data from prior cycles, achieved better rates of response than BayesB and Bayes LASSO models. A Support Vector Machine method, with a radial basis kernel, had the worst estimated prediction accuracies and the least long term genetic response. Application of genomic selection in a closed breeding population of a self-pollinated crop such as soybean will need to consider the impact of these factors on trade-offs between short term gains and conserving useful genetic diversity in the context of the goals for the breeding program.

## Introduction

Genetic improvement in commercial and public soybean breeding programs include variety development as the evaluation phase of recurrent selection (Orf 2008). Each cycle consists of making crosses among selected lines, creating replicable homozygous lines by self-pollinating progeny for several generations, evaluating the lines in replicated field trials for several years during which poor performing lines are discarded and superior performing lines are retained to cross and begin a new cycle. Pedigrees of modern soybean varieties confirm that genetic improvement of soybeans in Maturity Zones (MZs) II, III and IV has been largely through intra-population recurrent selection (Byrum, personal communication; Hyten et al. 2006; Mikel et al. 2010; Langewisch et al. 2017; Achard et al. 2020). Prior to the Plant Variety Protection Act in 1970 (https://www.ams.usda.gov/rules-regulations/pvpa), a cycle of genetic improvement would require from 10 to 14 years, whereas in the last 40 years commercial organizations have invested in development of continuous nurseries including software that streamlines inventories and logistics of seed transfer, resulting in the capacity to complete a cycle of recurrent selection in five years (Byrum et al. 2017; Anderson et al 2019). Thus, using current best practices, a soybean breeder might experience three to five cycles of genetic improvement.

Despite reducing cycle times to less than ½ of the time required 40 years ago, yield improvements for soybean have not doubled in the corn-soybean agricultural system of the United States (Mikel et al. 2010; Specht et al. 2014). Recognizing the relatively slow yield improvements, farmer-members of the North Central Soybean Research Program (NCSRP) provided funding to public soybean breeders to identify novel QTL in germplasm adapted to MZ’s II, III and IV and subsequently to identify methods to increase soybean yields (https://www.ncsrp.com/NCSRP_research.html#yield). The SoyNAM project provided evidence of novel useful genetic variability for yield among the SoyNAM lines (Song et al. 2017; Diers et al. 2018) and genomic selection (GS) is being investigated for improving yield in pubic germplasm.

After the seminal work by Bernardo and Yu (2007), crop breeders have experimentally demonstrated the relative advantage of GS to phenotypic selection (PS) for a few cycles of recurrent selection in barley, maize, oats, rice and wheat (Bernardo 2008, 2014; Asoro et al. 2011; Heslot et al. 2012, 2015; Nakaya and Isobe 2012; Emily and Bernardo 2013; Crossa et al. 2014; Beyene et al. 2015; Bassi et al. 2016; Marulanda et al. 2016; Jonas and de Koning 2013, 2016; Hickey et al. 2017; Goiffon et al. 2017). Genomic prediction (GP) has been integrated in commercial soybean variety development projects as a tool to identify crosses that are most likely to produce superior progeny and to a lesser extent as a means to reduce the costs of conducting preliminary field trials (Kurek 2018). However, implementation of GS in soybeans has not been based on an understanding of how various factors and their interactions can affect responses to recurrent GS (RGS) in a crop with population structures and genome organizations such as found in soybean.

Simulations have been used to demonstrate that genome organization, population structure, genetic architecture, heritability, selection intensity, GP models and composition of training sets will affect responses to selection (Robertson 1960; Dempfle 1974; Kang 1979; Cockerham & Burrows 1980; Kang and Namkoong 1980; Kang 1987; Kang and Nienstaedt 1987; Podlich and Cooper 1998; Cooper et al. 2002; Habier et al. 2007; Goddard 2009; Zhong et al. 2009; Jannink 2010; Bastiaansen et al. 2012; Bijma 2012; de los Campos et al. 2013; Howard et al. 2014; Liu et al. 2015; Michel et al. 2016). Most of these studies evaluated impacts of subsets of factors on responses across a few cycles of recurrent selection, although a couple of studies investigated limits of responses to RGS across more than 20 cycles (Jannink 2010; Liu et al. 2015).

Comparisons of genomic prediction (GP) models have revealed that simulated genetic architecture and proportion of phenotypic variance due to the genotypic variance affected prediction accuracies and simulated responses to selection (de los Campos et al. 2010, 2013; Wimmer et al. 2013; Howard et al. 2014). Parametric methods such as Ridge Regression (RR) and Bayesian regression in the mixed effects modeling framework provide more accurate predictions and greater gains for traits with additive genetic architecture, whereas non-parametric machine-learning methods such as Neural Networks and Support Vector Machines provide more accurate predictions for traits with epistatic genetic architectures (Howard et al. 2014). Prediction accuracies are essentially the same for all GP models applied to data with additive genetic architectures (Long et al. 2010, 2011; Guo et al. 2012; Howard et al. 2014). Given additive genetic architectures, RR (Endelman 2011) and Bayesian models (Pérez and de los Campos, 2014) provide similar short-term genetic gains, but different responses after ~ 15 cycles of RGS.

Marker densities impact prediction accuracies and short term genetic gains, with a dense marker set performing better than a sparse set, although there are limits to improvements from increased marker densities that depend on linkage disequilibrium (LD) and structure of the breeding population (de Roos et al. 2009; Schulz-Streeck et al. 2012; Hickey et al. 2014; Xavier et al, 2016; Norman et al. 2018). Decreased prediction accuracies of GP models in later cycles of recurrent GS are associated with decay of LD between marker loci (ML) and quantitative trait loci (QTL), loss of relationships between lines in early and later cycles of selection or a combination of both (Habier et al. 2007; Zhong et al. 2009; Hickey et al. 2014; Liu et al. 2015; Müller and Melchinger 2017, 2018). It is possible to offset the loss of relationships and LD, by updating the training sets with each cycle of selection (Jannink 2010; Liu et al. 2015; Müller et al. 2017, 2018). If training data are from only the current cycle of selection, then predictions do not take into account relationships between the current population and the founder population or possible loss of LD. At the other extreme, data from all prior cycles of selection can be included with data from the current cycle in the training set, but practical computation limits will be encountered with large training sets.

With the development of male sterile and insect pollination systems for soybean, it will become possible to conduct one to three cycles of RGS per year (Ortiz-Perez 2008; Davis, personal communication). This will enable increased selection intensities and greater immediate genetic gains from GS because much larger numbers of progeny can be evaluated without the commensurate added expense of increased numbers of field plots (Heslot et al. 2015). However, it is well-established that increased selection intensities will reduce the genetic potential of a breeding population through loss of useful genetic variability and ultimately will limit responses to selection (Robertson 1960; Hill and Robertson 1968; Bulmer 1970).

Emerging technologies, such as genome editing, promise to provide novel useful genetic variability for crops that do not have natural genetic variability for traits of interest (Brooks et al. 2014; Jenko et al. 2015; Hickey et al. 2017; Rodríguez-Leal et al. 2017; Lemmon et al. 2018). However, before assuming that genome editing will enable genetic improvement after loss of useful genetic variance from selection, it is important to recognize that expression of novel genetic modifications within genetic networks depend on the genomic background, even in simple model organisms such as yeast (Peccoud et al. 2004; Forsberg et al. 2017; Hou et al. 2018). Since RGS with one to multiple cycles per year (Gaynor et al. 2017) will become feasible for soybean and because genome editing will need genetic diversity for success, soybean breeders need to determine appropriate trade-offs between short term genetic gains and retention of useful genetic variability in the germplasm that will be inherited by future plant breeders (Rodríguez-Leal et al. 2017; Lemmon et al. 2018).

In anticipation of emerging technologies that will enable soybean breeders to implement a two part strategy for genetic improvement (Gaynor et al. 2017), we investigated the impacts of selection methods (PS and GS), training sets (TS), number of QTL, heritability (H), selection intensity (SI) and their interactions on responses to RGS. We employed a factorial treatment design, where each combination of factor levels was replicated ten times in simulated RGS and RPS of recombinant inbred lines (RILs) across 40 cycles. Outcomes reported herein provide a foundation for comparisons with strategies that use other types of progeny for selection and crossing as well as practical guidelines for understanding trade-offs between genetic gains in early cycles and retention of useful genetic potential for future genetic improvement of soybeans.

## Methods

### Simulations and Treatment Design

The impact of nQTL, SI, h, TS and SM on response to selection across 40 cycles of recurrent selection were evaluated with a factorial design consisting of 306 combinations of factors. In contrast to evaluating one or two variables at a time, the factorial design is widely used to comprehensively determine factors that will optimize processes (Myers 1976; Collins et al. 2014; Dunn 2020). Specifically the treatments consisted of three values for number of simulated QTL, three selection intensities, two values for non-genetic variance, five selection methods and four types of TS used to update four GP models (Table 1). In summary the treatment design consists of 18 combinations of factors for PS plus 288 combinations of factors for GS methods for a total of 306 combinations of factors. Note that TS are irrelevant for PS. Each set of factor combinations was replicated with ten simulated recurrent selection projects across 40 cycles resulting in 3060 simulations with 122400 outcomes.

**TABLE 1:** FACTORIAL DESIGN

| **Factors** | **Number of Levels** | **Values for Levels** |
| --- | --- | --- |
| **Number of QTL** | 3 | 40, 400, 4289 |
| **Heritability** | 2 | 0.7, 0.3 |
| **Selection Intensity** | 3 | 2.67, 2.34, 1.75 |
| **Selection Method** | 5 | i) PS- Phenotypic value<br>ii) GS – GP(RR-REML)<br>iii) GS - GP(Bayes B)<br>iv) GS - GP(Bayes LASSO)<br>v) GS – GP(SVM, Radial basis Function Kernel) |
| **Model Update: number of prior cycles used in training sets** | 4 | i) 0 previous cycles<br>ii) 10 previous cycles<br>iii) 12 previous cycles<br>iv) 14 previous cycles |
| **Total Number of unique combinations** | 360 |  |
| **Total Number of Simulations** | 3600 with 10 reps /condition |  |

Simulated soybean RILs were generated by crossing *in silico* 20 homozygous SoyNAM founder lines with IA3023 to generate 20 distinct F_1_ progeny. The F_1_ progeny from each of the 20 crosses were self-pollinated *in silico* for five generations to generate 100 RILs per family. The resulting 2000 RILS from 20 families were assessed for segregating genotypic information at 4289 SNP loci (Song et al. 2017; Xavier A et al. 2017; Diers B et al. 2018). On average alleles from the common founder occurred at a frequency of 0.9 and alleles at the loci from all other founder lines occurred at a frequency of 0.1. Soybeans has been subjected to phenotypic selection and genetic improvement for several thousand years (Anderson et al. 2019). Thus, the SoyNAM founders represent a realized population structure adapted to maturity zones II, III and IV that is akin to a single coalescent process used in many simulation studies (Daetwyler et al. 2013; Bandillo et al. 2017).

Of the 4289 SNP markers with genotypic scores for the SoyNAM population 3818 were polymorphic among the 20 families used as founders for the simulations. On average, 773 were polymorphic for a family with a standard deviation among families of 34. In the initial founding set of RILs, the average heterozygosity per SNP locus across 20 families was 0.09. ‘Gst’ is a measure of sub-population differentiation estimated as ratio of difference between expected heterozygosity of sub-populations to total expected heterozygosity. The average estimated G_st_ value across the genome for the initial founding set of RILs was 0.32, as determined by the ‘diff_stats’ function in the mmod R package (Jombart 2008; Ryman and Leimar 2009; Jombart and Ahmed 2011). Relative to previous reported founders that used a coalescent process (Woolliams and Corbin 2012; Hickey and Gorjanc 2012; Daetwyler HD et al. 2013), our simulations began with a structure more likely to be found in actual soybean breeding populations and with less, albeit more realistic, genetic diversity of soybeans adapted to maturity zones II, III and IV. We’ve also estimated Pairwise ‘Fst’ using ‘pairwise.fst’ in ‘hierfstat’ R package (Goudet 2005), which is a measure of population differentiation among pairs of populations. It is estimated as the ratio of difference between the average of the expected heterozygosity of the two populations and total expected heterozygosity of the pooled populations to total expected heterozygosity of the pooled populations. Average Fst among the 20 families in simulated SoyNAM data is 0.20. Whereas the average Fst using genotypic data from SoyNAM project among 40 families is 0.09 with a maximum pairwise Fst of 0.15 and a minimum Fst of 0.007. Average Fst among the clusters in USDA soybean germplasm collection is 0.22 - 0.23 (Song et al 2015; Xavier 2018).

Subsets of 40, 400, and 4289 SNP loci were designated as QTL. The QTL were distributed evenly throughout the genome, and each contributed equal additive effects of 5/-5, 0.5/-0.5, or 0.05/-0.05 units respectively to the total genotypic value of the simulated RILs. Thus, all three genetic architectures had the same potential to create genotypic values ranging from +200 to - 200 in the initial founder sets of RILs. Positive and negative allelic effects were simulated to alternate sequentially at QTL that are uniformly distributed across the Soybean genetic map. Because all marker alleles are QTL alleles when there are 4289 QTL, LD between marker alleles and QTL will not deteriorate across cycles of selection and recombination. Phenotypic values were simulated by adding non-genetic variance sampled from an N (0, σ) distribution to the simulated genotypic values, where σ was determined by the heritability on an entry mean basis among the initial sets of founder sets of RILs. Broad sense heritability on an entry mean basis (H) values of 0.7 and 0.3 were simulated for each of the three sets of QTL. After the phenotypic values were simulated in the initial founding RILs, the non-genetic variance was held constant across subsequent cycles of selection.

For each cycle of recurrent selection, 1%, 2.5% or 10% of the most positive phenotypic or predicted phenotypic values among 2000 simulated RILs, corresponding to selection intensities of 2.67, 2.34 and 1.75, were selected as lines to inter-mate for the next cycle. Matings among pairs of selected RILs used a design with greater contributions from the best selected lines (Figure 1) also known as networked families designs (Guo et al. 2013; Guo et al. 2014).

**Figure 1.**
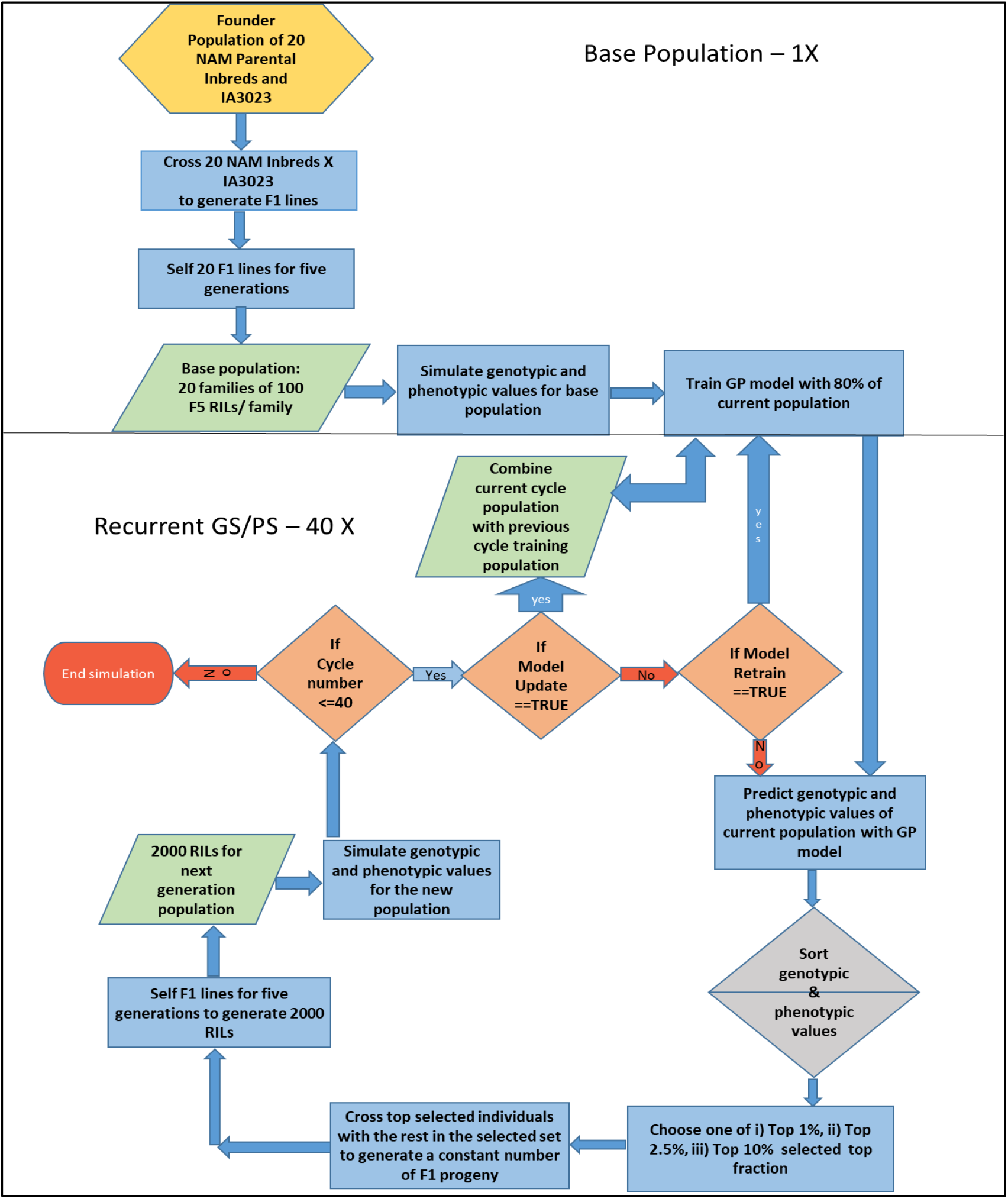
Flow Chart for Simulations of Recurrent Genomic Selection. The upper half panel represents the steps involved in generating the base population of 2000 F5 RILs derived from 20 NAM founder lines crossed, *in silico*, to IA3023. It includes the model training step for genomic prediction models. The lower half panel represents recurrent steps of prediction, sorting, truncation selection, crossing, and generation of F5 RILs for each cycle as well as the decision steps to check if the training set should be updated and if the recurrent process is to be continued for another cycle.

Based on previous results from Howard et al (2014), four GS methods were evaluated. Ridge Regression using restricted maximum likelihood (RR-REML) represented a frequentist parametric model. Bayes-B (BB) and Bayesian LASSO (BL) represented parametric bayesian models and Support Vector Machine with Radial Basis Kernel (SVM-RBF) represented a non-parametric method of machine learning.

Ridge regression was implemented with a method that employs expectation maximization to obtain Restricted Maximum Likelihood estimates of marker effects (Xavier et al. 2019). This computational method is faster than the popular implementation of ridge regression in rrBLUP package (Endelman 2011) and generates values that are highly correlated with the predictions based on the rrBLUP package (Figure S1). The BGLR package (Perez and de los Campos 2014) provided implementations of BB and BL models. The ‘Rgtsvm’ package in R was used for its implementation of the SVM with RBF kernel method (Wang et al. 2017). ‘Rgtsvm’ implements SVM training on GPUs with computing time several hundred times less than that required for the implementation in the ‘caret’ package on high performance computing clusters, and produces similar prediction accuracies and estimates of mean squared errors (Figure S2). The parameters used to train GP models with R packages are provided in Table 2.

**TABLE 2.** PACKAGES IN R FOR PARAMETRIC AND NON-PARAMETRIC MODELS WITH TUNING PARAMETERS

| GS Model | Package (R) | Model Tuning Parameters (BGLR package ) |
| --- | --- | --- |
| Ridge Regression | REML-EM (custom R script Xavier 2019) | EM algorithm for estimation of parameters with REML method without using matrix inversion |
| Bayesian LASSO | BGLR (Perez et al. 2014 ) | Priors for<br>varE (df=3,S=0.25); varU (df=3,S=0.63);<br>lambda(shape=0.53,rate=5e-5)<br>type='random',value=30),<br>nIter=20000,<br>burnIn=2000,<br>thin=1 |
| Bayes B | BGLR (Perez et al. 2014 ) | nIter=41000, burnIn =1000,<br>df0=4, R2=0.7 |
| SVM | Rgtsvm (Wang et al. 2017) | SVM with Radial basis function kernel on GPU |

For purposes of this manuscript we use the phrase ‘model updating’ to refer to retraining GP models with up to 14 previous cycles of training data (Figure S3). A preliminary analysis of TS on genotypic values and prediction accuracies was conducted using RR-REML models trained with data from the current cycle as well as 3, 5, 8, 10, 12, and 14 prior cycles. The results were compared with responses from the RR-REML model updated with TS’s comprised of data from all prior cycles and a RR-REML model with no updating. Training sets for each cycle were obtained by randomly sampling 1600 RILs from the set of 2000 simulated RILs in each cycle. The most accurate prediction and maximum genetic response was obtained with training data that is cumulatively added every cycle (Figure S4 and S5). The results indicate that 3-5 prior cycles of training data did not significantly improve prediction accuracies and responses relative to models that were not updated. Also, the standardized genotypic values and prediction accuracies, obtained using 10 to 14 prior cycles of data in the TS’s, were not significantly different than results based on TS’s consisting of all prior cycles. Based on the results of this preliminary study, we investigated responses to recurrent selection using TS’s consisting of up to 14 prior cycles of selection as well as data from the current cycle. After the 14^th^ cycle, training data consisted of 14 prior cycles of recurrent selection.

### Modeled response to recurrent selection

The averaged genotypic value for each cycle, c, of recurrent selection was modeled with a linear first order recurrence equation:

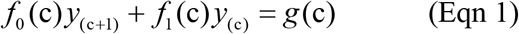

Where c is a sequence of integers from 0 to 39 representing each cycle of recurrent selection from cycle 1 to 40 and *f*_0_, *f*_1_ and *g* are constant functions of c. By rearranging the equation we note that the response in cycle c+1 can be represented as

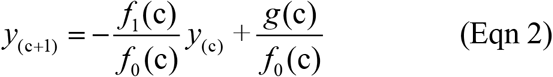

Since the ratios *f*_1_(c)/*f*_0_(c) and *g*(c)/*f*_0_(c) are constants, we can represent the response in cycle c+1 as

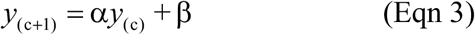

If *y*0 specifies the average genotypic value of the first cycle of RILs derived from the founders, then (3) has a unique solution (Goldberg 1958):

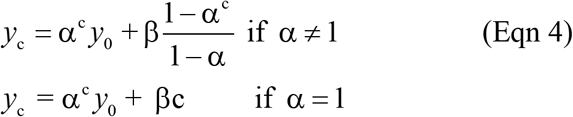

An alternative representation of (4) for the situation of α ≠ 1 is

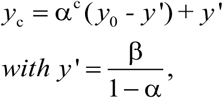

where α is less than 1 for genotypic response to recurrent selection and *y*’ represents the asymptotic limit to selection (Goldberg 1958). To illustrate, values of the sequence of c=0 to 39, with *y*_0_ = 0, for the range of α (0.6-0.9) and β (1.4-38) values are plotted in Figure 2. The curves can be interpreted as response to selection as a function of the frequencies of alleles with additive selective advantage, selection intensity, time and effective population size (Robertson 1960).

The parameters, *y*_o,_ α, and β, were estimated with a non-linear mixed effects method implemented in ‘nlme’ functions in the ‘nlme’ and ‘nlshelper’ packages (Pinheiro and Bates 2000; Baty et al. 2015; Pinheiro et al. 2019).

**Figure 2.**
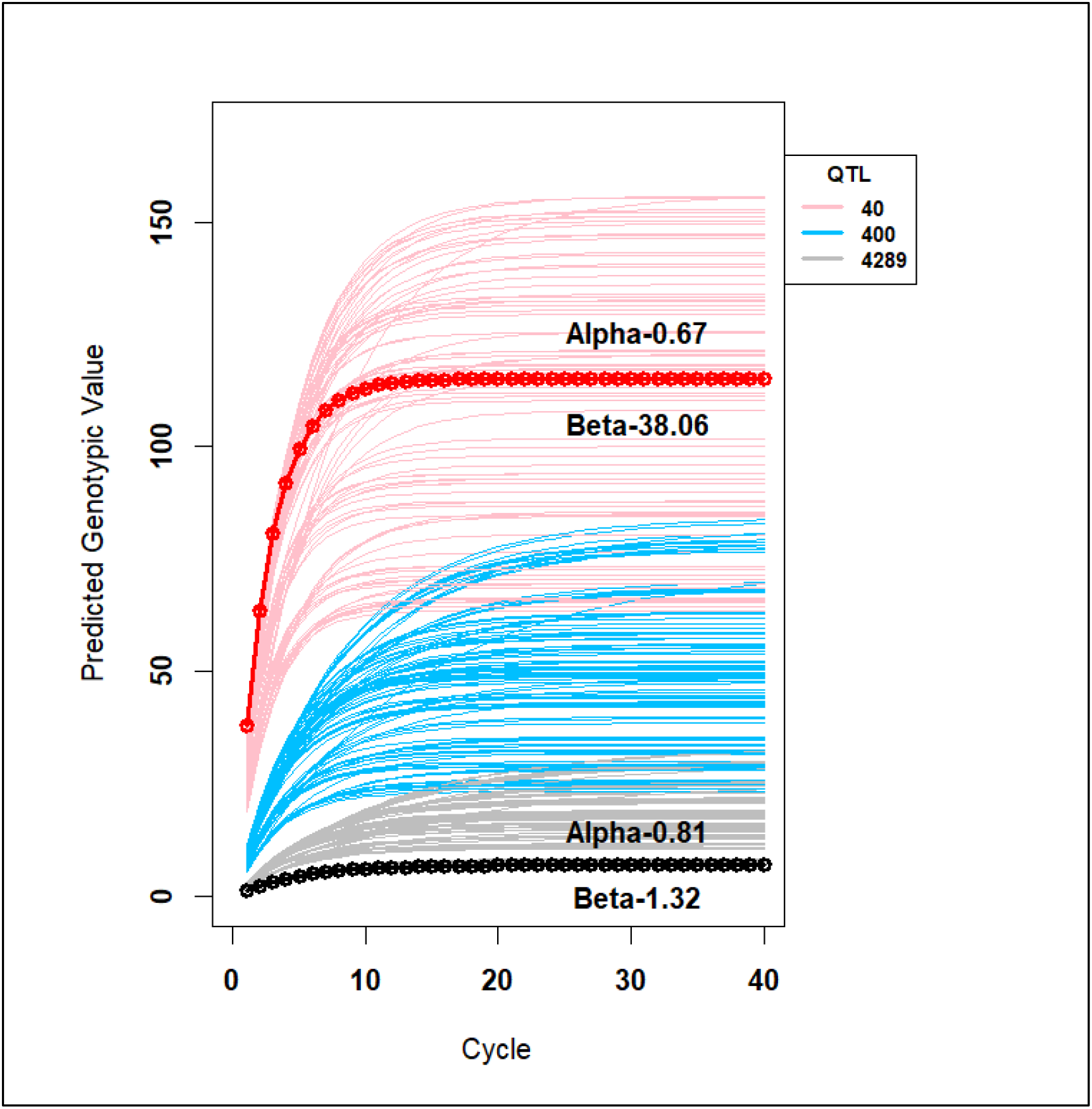
Theoretical Genotypic Values: Theoretical Genotypic Values from 40 cycles of recurrent selection modeled with the recurrence equation, where *y*c represents the genotypic value in cycle c, with c= 1, 2… 40 and values of α and β range from 0.6-0.9 and 1.32-38.06 respectively for 360 combinations of factors across all selection methods, training sets, selection intensities, number of simulated QTL and simulated heritabilities. The bold lines represent curves with the smallest and largest beta values and their corresponding alpha values. The curves are colored corresponding to 40 (pink), 400 (blue) and 4289 (gray) simulated QTL.

Since the limits of responses are asymptotic, the number of cycles before there is no longer response to selection is referred to as the half-life (Robertson 1960; Dempfle 1974; Kang 1979; Cockerham & Burrows 1980; Kang and Namkoong 1980; Kang 1987; Kang and Nienstaedt 1987). From the first order recurrence equation, the half-life is estimated as ln (0.5)/ln (α), when y0 is ‘0’ and the asymptotic limit is estimated as y’.

### *Analyses of variance* (ANOVA) *of modeled response to recurrent selection*

The purpose of the ANOVA is to evaluate the impact of factors and their interactions on the modeled responses to recurrent PS and GS methods. The analyses of variance used single and multi-level nlme models with modeled (eqn 4) responses grouped by treatment factors. The influence of multiple factor treatment combinations on estimated non-linear mixed effect models have not been implemented in standard statistical software packages that report the analysis of variance in terms of sums of squares and traditional ‘F-tests’. For discussions on the challenges of using standard F-test for non-linear mixed effects (nlme) models see (Pinheiro et al. 2000; Baty et al. 2015; Pinheiro et al. 2019). Consequently, we analyzed the variance among modeled responses using AIC, BIC and Likelihood metrics that were grouped based on combinations of factors consisting of selection methods, TS, SI, nQTL and simulated H.

In order to provide a balanced data table for analyses by the non-linear mixed effect model, responses that included PS, which has no TS’s, were assumed constant resulting in a balanced full factorial set of responses for 360 combinations of factors. The process of fitting, selecting and refining mixed effects models closely followed the steps described in Pinheiro et al. 2000; Zuur 2009 and Oddi et al. 2019). The complete process used in the study is provided in File S1.

In the first phase of model fitting, estimates of modeled parameters from nlsList models were retained as starting values for fixed effects. Both alpha and beta were fit only for intercept and deviations from estimated means conditioned on grouping variables were modeled as random effects using the ‘nlme’ R package. Multiple ANOVA of ‘nlme’ objects representing the models were used to identify combinations of factors with significant effects on the non-linear response model. The model with the lowest AIC score was selected as the best model. The best random intercept model in the first phase of model fitting process M31 in Table S1 was further refined by modelling the correlation structure.

### Evaluation of Simulated Response to Recurrent Selection

While the modeled genotypic values are evaluated using half-life and asymptotic limits, we have evaluated the simulated outcomes from recurrent selection using a set of metrics to assess responses, population characteristics, and GP model performance every cycle of selection.

The standardized genotypic value, R_s_ (eqn 5), was estimated every cycle as the change in genotypic value from the average genotypic value of 2000 RILs derived from the initial founders and standardized to the maximum genotypic potential (200 units) among the founders (Meuwissen et al. 2001; Liu et al. 2015).

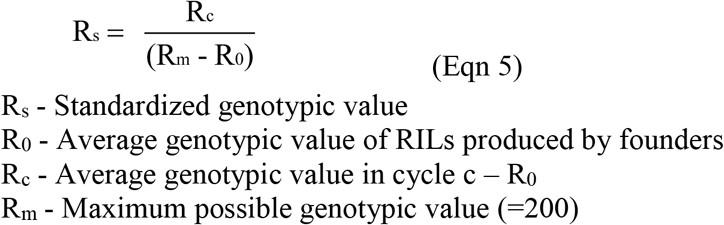

The most positive genotypic value (*M_gv_*) among RIL’s selected in cycle c is a metric used to evaluate the best RIL produced each cycle, while the standardized genotypic variance (*SV_g_*) defined as the estimated genotypic variance divided by the estimated genotypic variance of the initial population, was used to evaluate the loss of genotypic variance. Note that values for the *SV_g_* range from zero to one.

Response standardized to change in standard deviation of genotypic values captures genetic gain with respect to loss of genetic variance (eqn 6). The numerator term represents change in genotypic values of a population in cycle ‘c’ from cycle ‘0’ founder population normalized to standard deviation of genotypic values in cycle ‘0’. The denominator term represents change in standard deviation of genotypic values from cycle ‘0’ to cycle ‘c’ as a fraction of standard deviation of genotypic values in cycle 0. This metric is similar to the metric used to refer to efficiency of converting loss of genetic diversity to genetic gain of a selection method in recurrent selection (Gorjanc et al. 2018). While efficiency is estimated as slope in linear regression model with numerator as ‘y’ term and denominator as ‘x’ term in the linear part of response curve, with Rs_Var it is possible to visualize both linear and non-linear sections of the response curve.

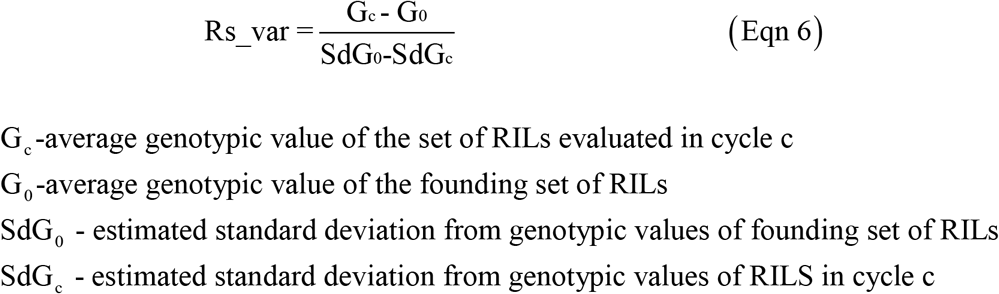

Estimated Linkage disequilibrium (*LD*) among pairs of marker loci on all 20 chromosomes was evaluated as the deviation of observed gametic frequency of alleles at a pair of loci from the product of the individual allele frequencies, assuming independence (Weir 1996). The R function ‘get_PG_LD_StatsGSMethods’ used to estimate pairwise LD between markers is provided in the R package ‘SoyNAMPredictionMethods’. GP models were assessed using the estimated prediction accuracies (r_*ps*_), defined as the estimated linear correlation (Pearson) between predicted and simulated phenotypic values and the estimated Mean Squared Error (MSE), and defined as the mean of the squared deviations of the predicted phenotypic values from the simulated values.

## Analyses and Data Availability

More information on the analyses can be found in the R package ‘SoyNAMPredictionMethods’. Also, simulated data and code are available as part of the package (File S1 found at http://gfspopgen.agron.iastate.edu/SoyNAMPredictionMethods_v2_2020.html). Supplemental material including the R package can be found at https://figshare.com/s/f7d4e515e563beef550b and http://gfspopgen.agron.iastate.edu/SoyNAMPredictionMethods_v2_2020.html describes how to use the package. The complete process for fitting NLME models and ANOVA can be found at http://gfspopgen.agron.iastate.edu/Vishnu_MS1/NLME_Models_Part_I.html .SoyNAM genotypic and phenotypic data are available in SoyBase (Grant et al. 2010).

## Results

### Prediction accuracies in the founding sets of RILs

Estimates of prediction accuracies, r_ps_, of GP models trained with the initial set of 2000 F_5_-derived RILs ranged from 0.75-0.82 for H of 0.7 and ranged from 0.38 - 0.49 for h of 0.3 (Figure 3). The initial r_ps_ for both H values was best with BB and poorest with the SVM-RBF. The nQTL had little effect on r_ps_ within either value of 0.7 or 0.3 for H. RR-REML and BL had smaller magnitude MSE values than BB and SVM RBF for all numbers of simulated QTL and both values for H (Figure 3). Accuracies are lower when GP models are trained without QTL in the training set, but follow a similar pattern as the models with QTL. MSE are greater or comparable for models trained without QTL than for models with QTL (Figure S6-S7). Average within family prediction accuracies are lesser than prediction accuracies from a combined TS comprising of RILs from all the families (Figure S8-S9). However, a combined TS will have (n*population size of family) for ‘n’ families and estimated accuracies will have confounding effects from training set size. Estimated accuracies for models trained with TS generated by random sampling from a combined population to keep the TS size same as family size are lower than average within family accuracies. MSE are lesser for combined TS than for models trained using within family TS and sampled TS (Figure S8-S9).

**Figure 3.**
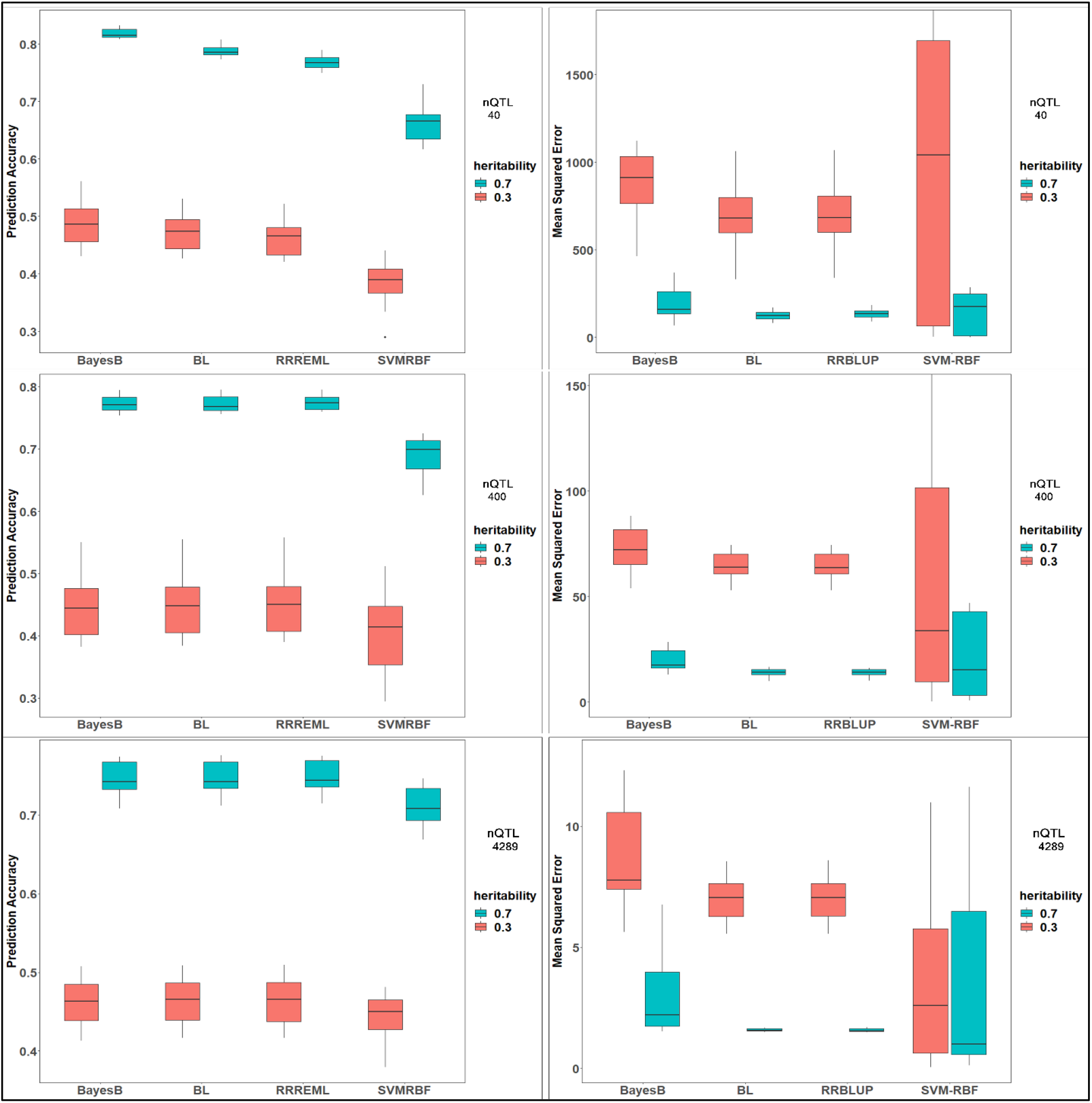
Estimated prediction accuracies and MSE in Founding Set of RILs: Estimated prediction accuracies (left panel) and mean squared errors (right panel) for four genomic prediction (GP) models: BayesB, BL (Bayes LASSO), RRREML (Ridge Regression with REML) and SVMRBF (Support Vector Machines with Radial Basis Function Kernel) trained with F5 RILs derived from crosses of 20 homozygous founder lines with IA3023. Phenotypes used to train the GP models consisted of genetic architectures comprised of 40, 400 and 4289 simulated QTL (top, middle and bottom) that were responsible for 70% (blue) and 30% (red) of phenotypic variability in the initial populations.

### Influence of factors on response metrics

The averaged responses Rs were modeled (eqn 4) and the results are consistent with theory (Figure 2). Averaged across all simulations there was rapid increase of Rs across the first five cycles of selection followed by slower responses from cycles 5 to 10 and no response after cycle 20.

While there are observable general trends for each of the individual factors, response metrics are unique for each combination of all factors (Figure 4–7, S10 - S14). The most parsimonious model requires unique estimates of α, and β (eqn 4) for each of the combinations of factors indicating that interactions among all factors have significant influences on the responses (Table S1). Also analyses of variances on subsets of 10 and 20 cycles of selection demonstrate that the interactions were important from the earliest cycles (File S2: Table S1, S2). Further, the relative importance of factors on interaction effects were consistent (nQTL>SM>SI>H>TS) and significant in analyses of variance conducted on subsets of 10 and 20 selection cycles (Table S1).

**Figure 4.**
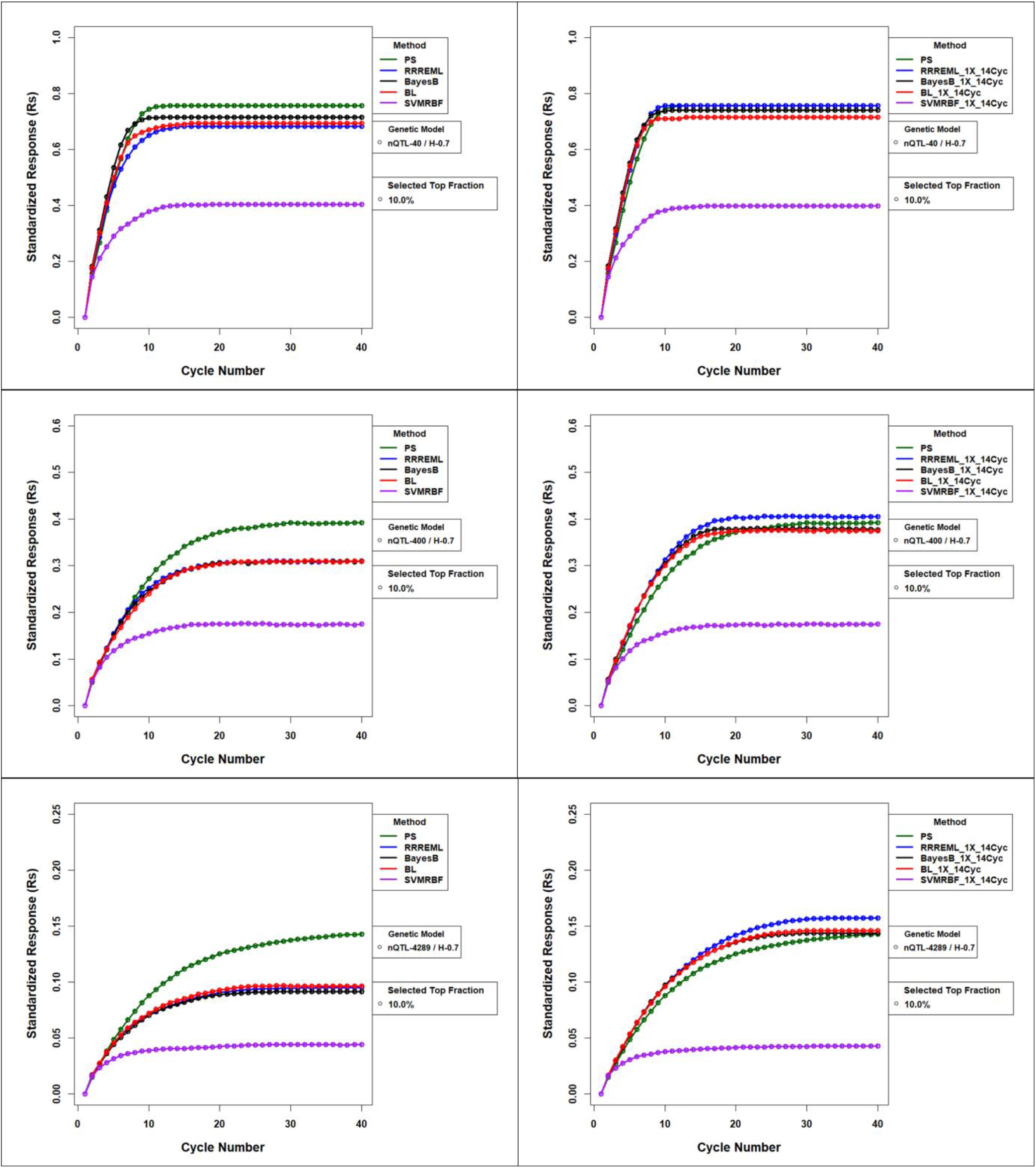
Standardized Responses for Comparison of GS methods with and without Updating for 0.7 H and Top 10% Selected Fraction. Forty cycles of standardized responses to selection of 10% of 2000 soybean RILs per cycle. Standardized responses are plotted by selection methods without (left panels) and with (right panels) model updating using prior cycles as training sets for the four genotypic prediction models. Phenotypic selection (PS) is not updated and hence is the same in the left and right panels. The top panels consist of responses for genetic architectures consisting of 40 simulated QTL. Middle panels consist of responses for genetic architectures consisting of 400 simulated QTL and the bottom panels consist of responses for genetic architectures consisting of 4289 simulated QTL. All 40, 400, and 4289 simulated QTL are responsible for 70% of phenotypic variability in the initial population. PS – Phenotypic Selection, RR-REML-Ridge Regression with Restricted Maximum Likelihood, BL – Bayes LASSO, and SVMRBF-Support Vector Machine with Radial Basis Kernel.

Among the three factors SM, TS and SI that are under the control of the plant breeder, we consider SM as the primary factor of interest, whereas TS and SI are considered secondary factors of interest that significantly modulate the effect of SM on response. In addition, nQTL and H that are not under the control of the plant breeder are noted for their large and significant impacts on response metrics (Table S1).

Relative to PS, the three parametric GP models provided greater initial rates of response, reduced population half-life and faster loss of genetic variance. Whereas, selection using the SVMRBF consistently produced the least effective responses (Figure 4–7). Training sets consisting of data from up to 14 prior cycles of selection, compared to TS’s consisting of data from only the current cycle of selection are observably distinctive for both the short term and long term response metrics (Figure 4–7). Importantly, use of the TS’s significantly improved responses to selection in both the short term and long term for all other combinations of factors (Table S1). Selection intensity is the third most important factor to affect the response metrics (Table S1). Consistent with theory (Robertson 1960; Hill and Robertson 1968; Bulmer 1971,1976), greater SI’s (associated with retention of smaller numbers of RILs to initiate subsequent cycles, were associated with more rapid response rates, shorter half-lives, faster loss of genetic variance and significantly lower Rs values as the population approached it’s limit to respond to selection.

The modeled responses were highly dependent on each of the factors, with nQTL showing the greatest deviations from the average response (Table S1). From the modeled responses the half-life for a responsive population varied from 1.6 to 4.9 cycles for 40 QTL, whereas for 400 and 4289 QTL it ranged from 2.2 to 8.8 and 3 to 9.5 cycles respectively. The modeled responses were highly correlated with the simulated responses (Pearson correlation coefficient: 0.94-0.97).

To illustrate the impact of nQTL, consider Rs values plotted across forty cycles of recurrent selection (Figure 4 and S15). If the genetic architecture of the trait consists of 40 and 400 QTL, responses to selection were limited after 10-15 cycles of selection, whereas for 4289 QTL, limits to selection responses were not realized until 30 to 40 cycles of selection.

Next consider that the maximum realized response for 40 simulated QTL was from 0.32 to 0.78 of maximum genotypic potential among the founders, whereas for 400 and 4289 QTL, the realized response was 0.12 - 0.42 and 0.04-0.17 of the maximum (Figure 4 and S15). Also if there are 40 simulated QTL, the maximum attained values are as high as 80% of the maximum value of 200 in less than ten cycles of recurrent selection (Figure 8 and S16). In contrast, Rs values are no greater than 40% of the maximum value and stop responding to selection in 10-15 cycles if there are 400 simulated QTL. Rs values were never greater than 15% of the maximum value and only begin to approach a limit after 20 cycles if there are 4289 simulated QTL.

As expected, responses to selection reflect declining genetic variances (Figure 9 and S17). The loss of Sgv’s across cycles is much faster with fewer simulated QTL than larger numbers of QTL. Likewise the estimated prediction accuracies (Figure 10 and S18) approach zero as the genotypic variance approaches zero. Average MSE for the GP models increase across cycles of selection (Figure 11 and S19) LD among markers approach zero as the genotypic variance approaches zero although the covariance among these response metrics depend on the other simulated factors (Figure S20-S24).

To interpret the role of nQTL as a factor, it is important to recall that: 1) positive and negative allelic effects were simulated to alternate sequentially at QTL (marker loci) that were distributed across the genome according to the Soybean genetic linkage map. 2) Crossing nearly homozygous lines and subsequently self-pollinating progeny for five generation before genotyping and phenotyping within each cycle creates a limited number of large linkage blocks. Analyses of the number of linkage blocks each generation reveals that regardless of the number of QTL, the number of linkage blocks per cycle ranges from 70-90. It was not the same blocks each cycle, but if the nQTL equals 40, then each linkage block included all segregating QTL each cycle. For 400 and 4289 simulated QTL, each linkage block had a net genetic effect of zero or +/− 0.5 or +/− 0.05 multiplied by the number of QTL in the block respectively. Thus, the nQTL might be better understood as the magnitude of genetic effects associated with segregating linkage blocks.

The observable general trend for H, or perhaps more accurately understood as contributions of non-genetic effects to the phenotypes, was that H values of 0.7 in the initial phenotypic variance resulted in Rs values that were greater than H values of 0.3 of the initial phenotypic variance (Figure 4 and S15). The trend in Rs values is correlated with the other response metrics, in particular prediction accuracies of the GP models. The loss of estimated prediction accuracies are greater with H values of 0.3 than 0.7 with relaxed selection intensities (Figure 10 and S18). Other combinations of SI and H require model updating to provide reasonable GP model prediction accuracies and achieve greater responses across more cycles of selection. As we would expect, for all combinations of SI and nQTL, loss of genotypic variance are greater with H values of 0.7 than 0.3 (Figure 9 and S17).

### Some specific outcomes of interest

For purposes of illustrating interaction effects on SM’s across all 40 cycles of selection, consider the most relaxed SI of 1.75, associated with selecting 10% of the RILs per cycle. When GP models are not updated, BB produced greater R_s_ values than PS in the early cycles for all nQTL and both levels of H, whereas PS resulted in greater responses than all GS methods after the 10^th^ cycle (Figure 6, S15 and S25). SVMRBF did not demonstrate any better responses than PS in either early or late cycles for any nQTL or level of H (Figure 6, S15, and S25; File S3 and S4).

If the parametric GP models are updated with training sets consisting of data from up to 14 prior cycles of recurrent selection, responses to RR-REML demonstrated the greatest responses (Rs) for 40, 400 and 4289 QTL across both levels of H (Figure 7, S15 and S26; File S3 and S4). If the RR-REML model is updated with up to 14 prior cycles of training sets, responses are larger than PS for up to 10-40 cycles depending on the number of QTL, H and SI (Figure 4 and 7; File S3 and S4). When BB and BL GP models are updated, responses are larger than PS for up to 5, 20 and 40 cycles for 40, 400, and 4289 QTL respectively. Similar, albeit distinctive, comparisons among outcomes from GP models with model updating for genetic architectures responsible for 0.3 of the phenotypic variance in the initial sets of RILs (Figure S26) are described in File S4.

Relative to responses without model updating application of RR-REML and Bayesian methods with model updating resulted in greater responses. Model updating with Bayesian methods also resulted in less favorable responses than the RR-REML (Figure S27 and S28; File S4). SVMRBF when updated with TS’s demonstrated no significant improvement relative to SVMRBF without updating for all genetic architectures, levels of H and SI’s (Figure S27 and S28; File S4). If the genetic architecture explains only 30% of the phenotypic variability in the initial sets of RILs, the relative improvements in Rs values across cycles using updated TS’s are better than simulated QTL that explain 70% of the phenotypic variance (Figure S28). Percentage gain in responses for GS with model updating relative to response from GS without model updating are provided in File S5.

In terms of lost genetic potential, every cycle of selection reduced the maximum possible genetic value. When GP models are not updated, the genetic potential is lost at a rapid rate beginning in the early cycles, whereas when GP models are updated, genetic potential is retained in the population and genetic variance decreases at a slower rate. PS had the least loss in genetic potential relative to all four GP models without updating. However, with model updating, the loss of genetic potential using parametric GP models was almost the same as PS. Among the parametric GP models, RR-REML and Bayesian methods showed similar slow losses of genetic potential with and without model updating. SVMRBF GS had the greatest loss of genetic potential beginning with the early cycles (Figure 5).

**Figure 5.**
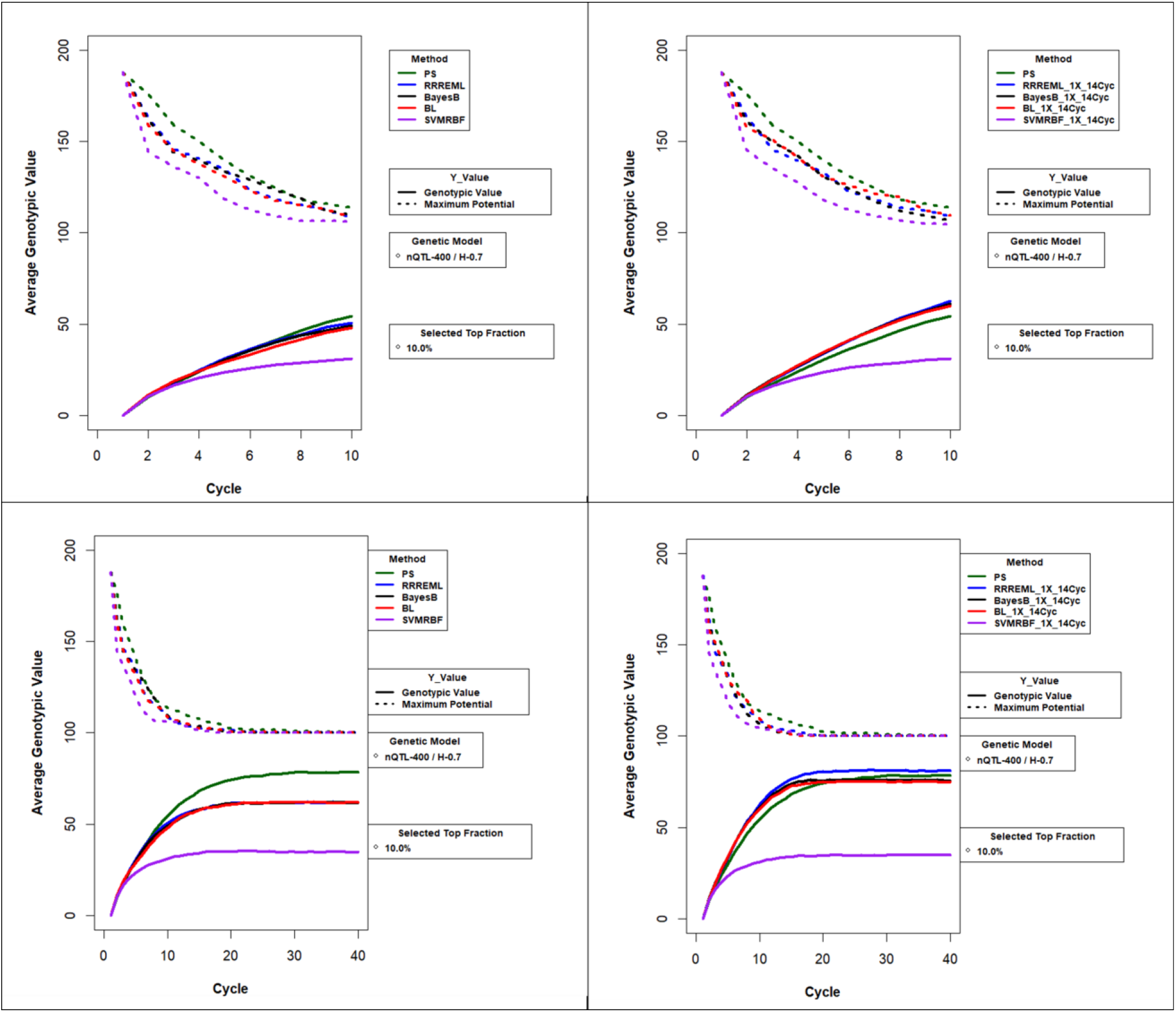
Average Genotypic Value and Maximum Genetic Potential for Comparison of GS methods with and without Updating for 0.7 H and Top 10% Selected Fraction. Average genotypic value and maximum possible genotypic value in recurrent selection of 10% of 2000 soybean RILs per cycle. The values are plotted by selection methods without (left panels) and with (right panels) model updating using prior cycles as training sets for the four genotypic prediction models. Plots demonstrate decrease in maximum possible genotypic value due to loss of favorable alleles and increase in average genotypic value for 10 cycles (upper panel) and 40 cycles (lower panel) of selection. Phenotypic selection (PS) is not updated and hence is the same in the left and right panels. The top and bottom panels represent genotypic values for genetic architectures consisting of 400 simulated QTL responsible for 70% of phenotypic variability in the initial population. PS – Phenotypic Selection, RR-REML-Ridge Regression with Restricted Maximum Likelihood, BL – Bayes LASSO, and SVMRBF-Support Vector Machine with Radial Basis Kernel.

**Figure 6.**
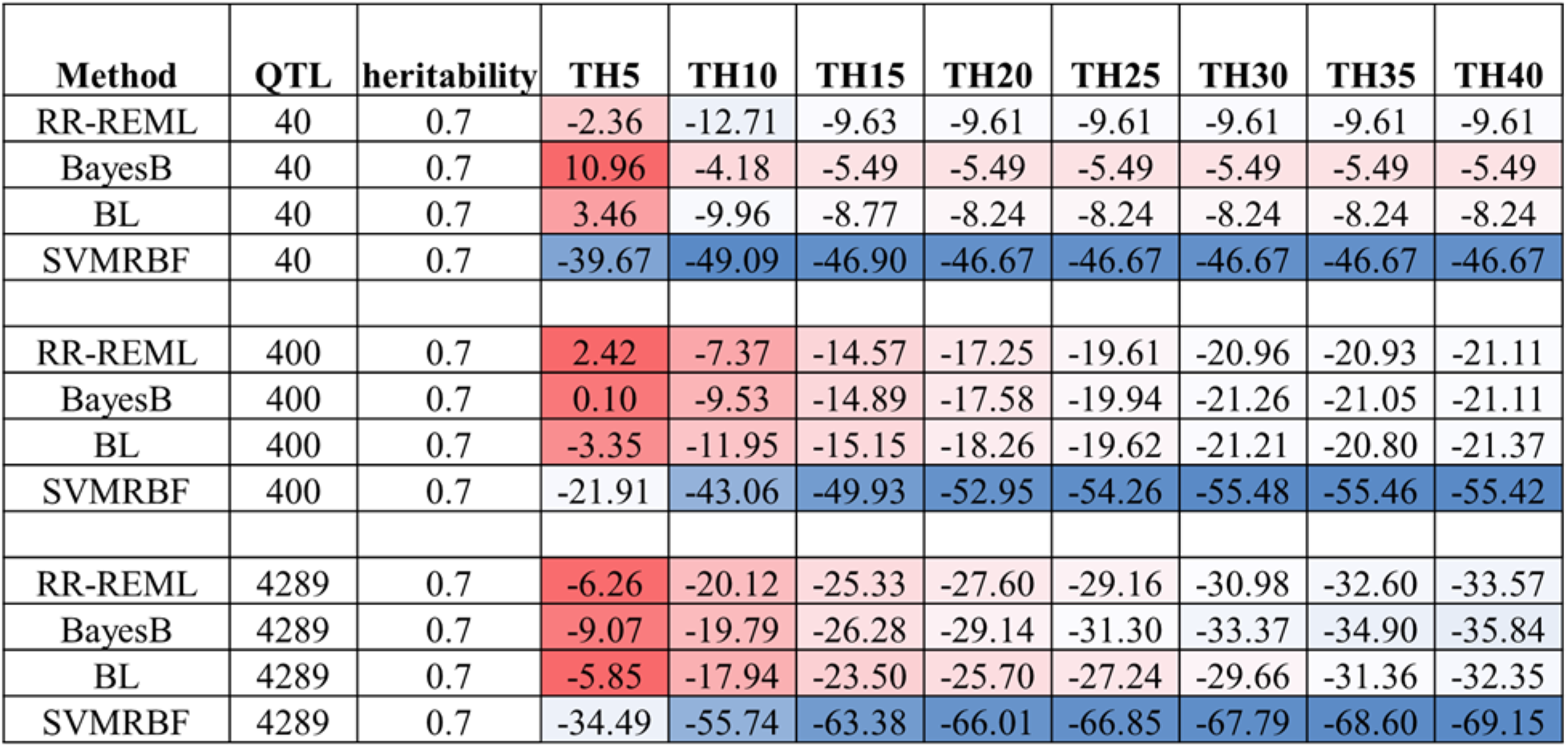
Heat Map for Percent Gain in Rs Relative to PS for 0.7 H for GP Models without Updating: Heat map indicating standardized response relative to PS as percentage gain after 40 cycles of recurrent selection using genomic prediction models without updated training sets for 40, 400, 4289 simulated QTL responsible for 70% of phenotypic variability in the initial population. Blue to red shaded cells represent increasing gain in response relative to PS. RR-REML-Ridge Regression with Restricted Maximum Likelihood, BL – Bayes LASSO, and SVMRBF-Support Vector Machine with Radial Basis Kernel.

**Figure 7.**
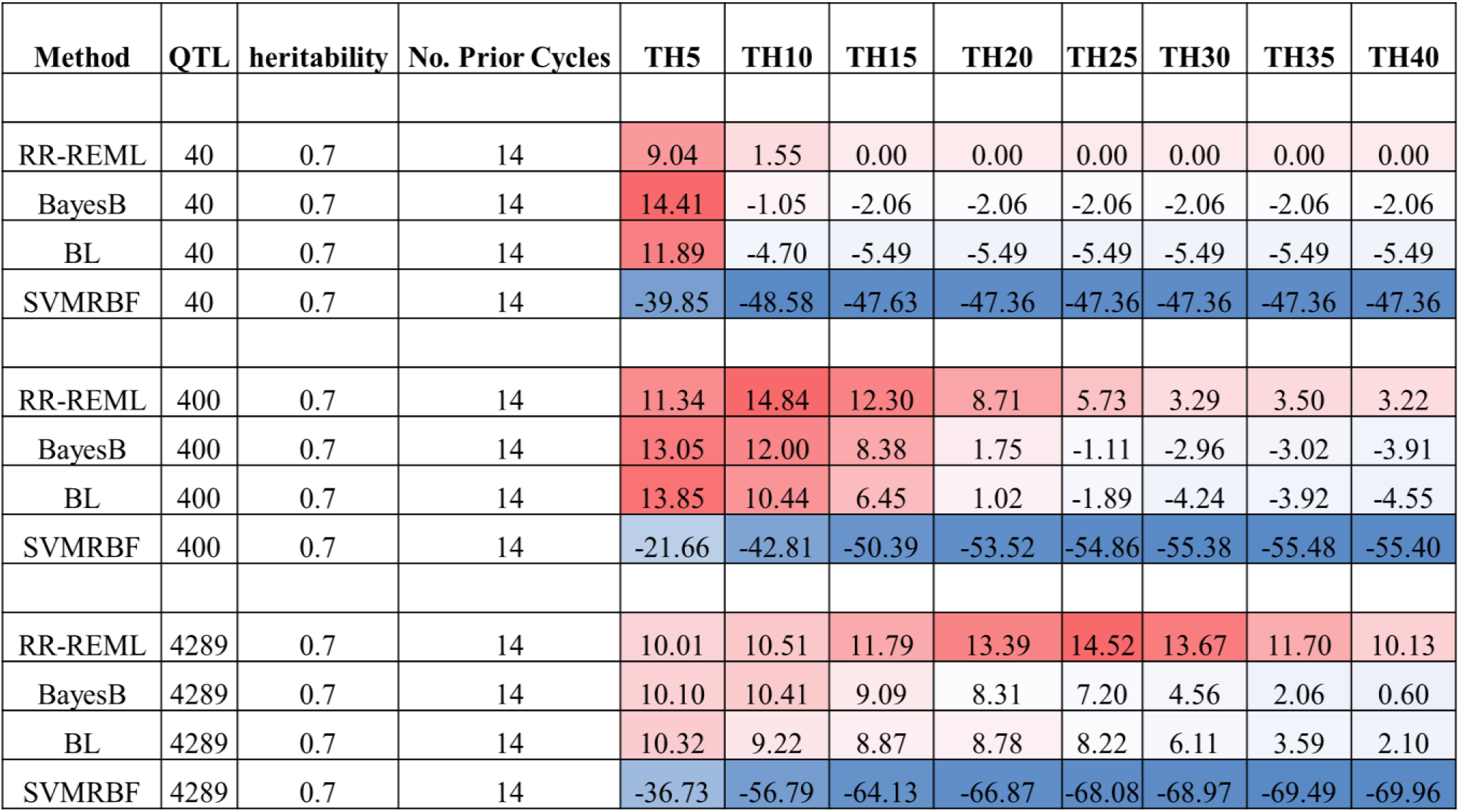
Heat Map for Percent Gain in Rs Relative to PS for 0.7 H for GP Models with Updating: Heat map indicating standardized response relative to PS as percentage gain after 40 cycles of recurrent selection using genomic prediction models with updated training sets from up to 14 prior cycles of selection for 40, 400, and 4289 simulated QTL responsible for 70% of phenotypic variability in the initial population. Blue to red shaded cells represent increasing gain in response relative to PS. RR-REML-Ridge Regression with Restricted Maximum Likelihood, BL – Bayes LASSO, and SVMRBF-Support Vector Machine with Radial Basis Kernel.

**Figure 8.**
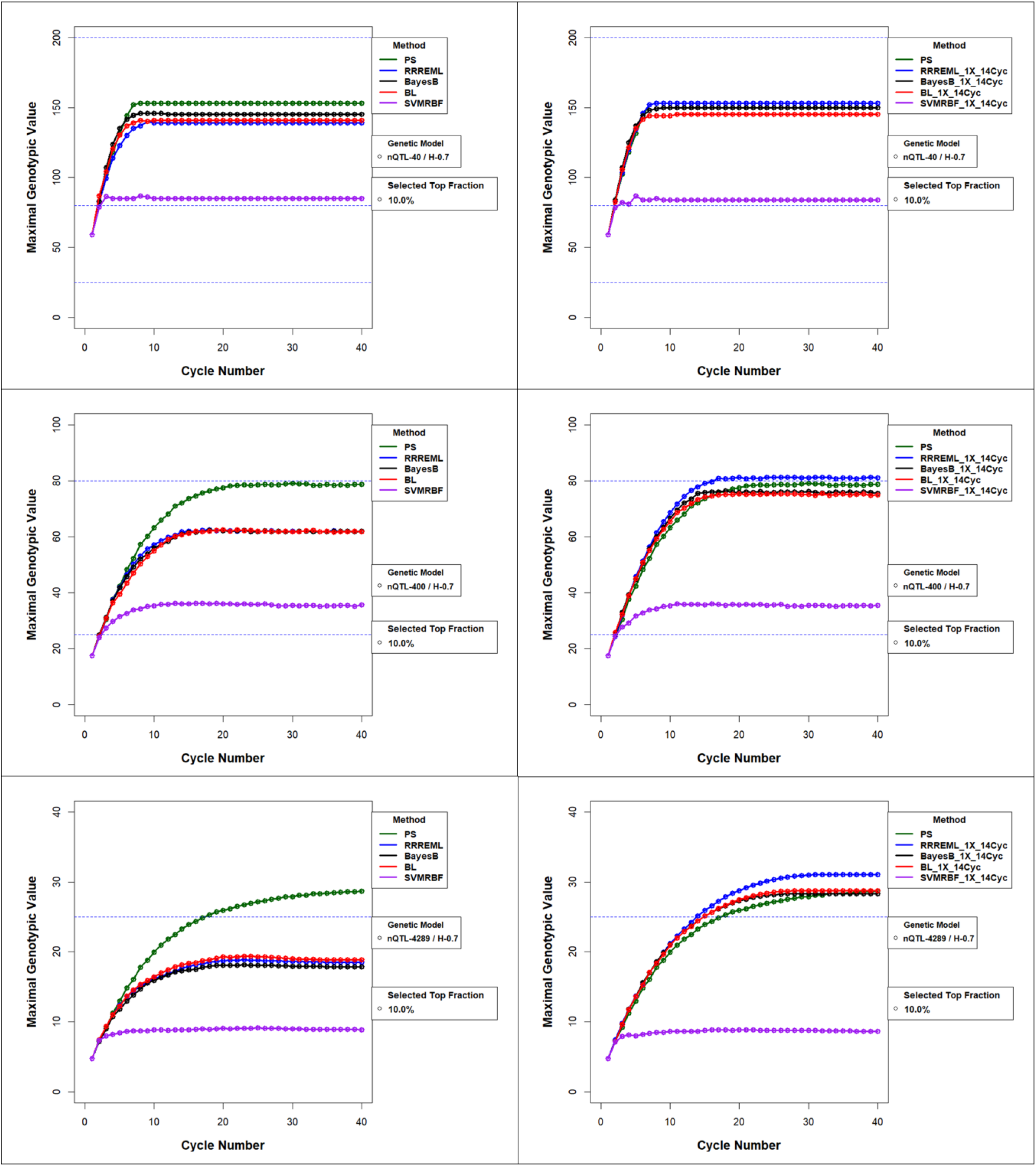
Maximal Genotypic Value for Comparison of GS methods with and without Updating for 0.7 H and Top 10% Selected Fraction: Maximum attained genotypic values (Mgvs) in recurrent genomic selection and phenotypic selection (PS) without updating the training sets in the left panels and with training set updates from up to 14 prior cycles in the right panels. PS has no training sets and hence does not change between the left and right panels. a) 40 QTL (top), b) 400 QTL (middle) and c) 4289 QTL (bottom) responsible for 70% of phenotypic variability in the initial population and selection of 10% of the RILs in each cycle. PS – Phenotypic Selection, RR-REML-Ridge Regression with Restricted Maximum Likelihood, BL – Bayes LASSO, and SVMRBF-Support Vector Machine with Radial Basis Kernel.

**Figure 9.**
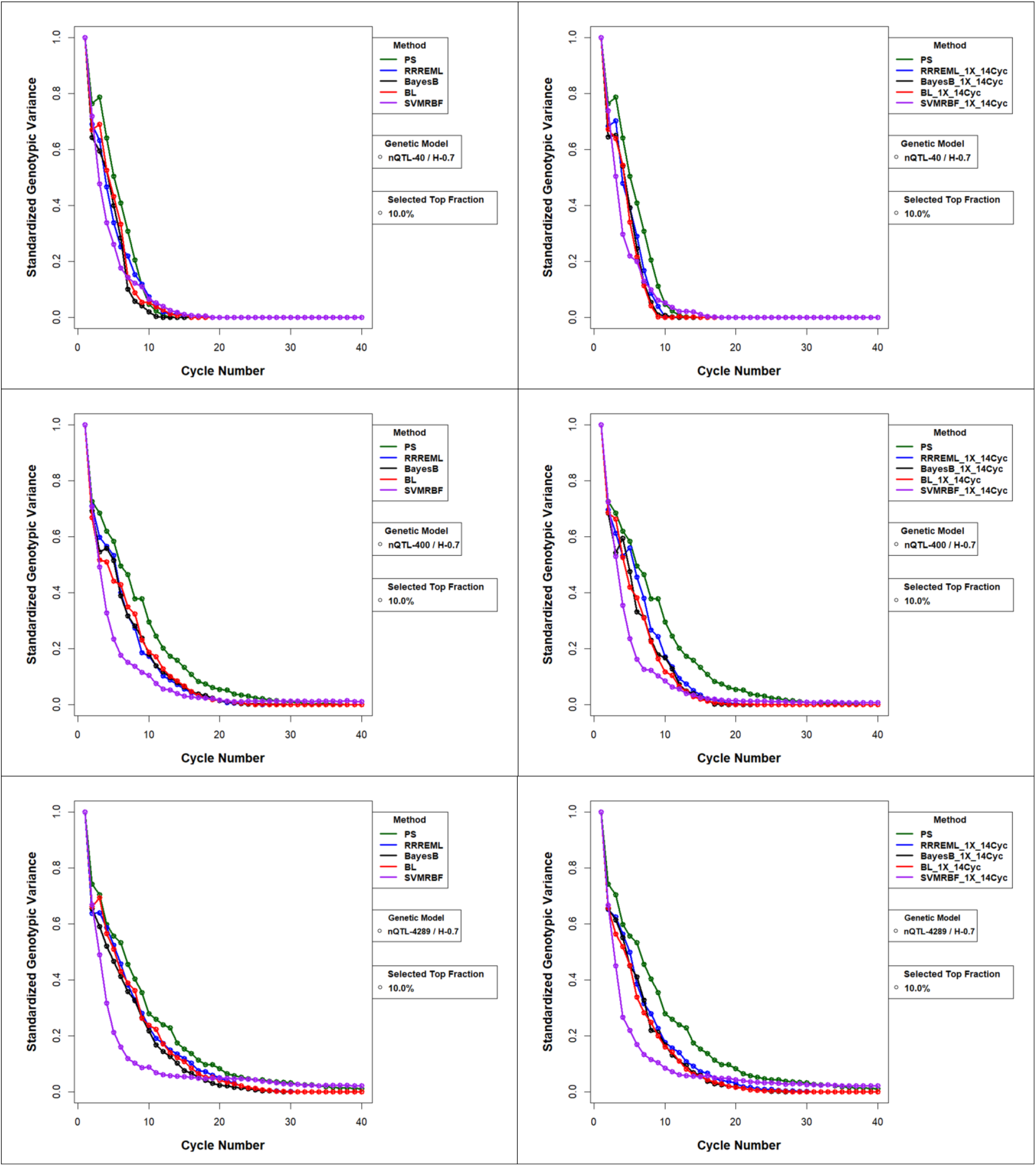
Standardized Genotypic Variance (Sgv) for Comparison of GS methods with and without Updating for 0.7 H and Top 10% Selected Fraction. Standardized genotypic variance without training set updating (left panels) and with training set updating using prior cycle training data (right panels) for the four GP models. PS has no updating and hence is the same in both left and right panels. A) Training data from up to 14 prior cycles for 40 simulated QTL (top), 400 simulated QTL (middle) and 4289 simulated QTL (bottom) responsible for 70% of phenotypic variability in the initial population and top 10% of RILs with the greatest predicted values. PS – Phenotypic Selection, RR-REML-Ridge Regression with Restricted Maximum Likelihood, BL – Bayes LASSO, and SVMRBF-Support Vector Machine with Radial Basis Kernel.

**Figure 10.**
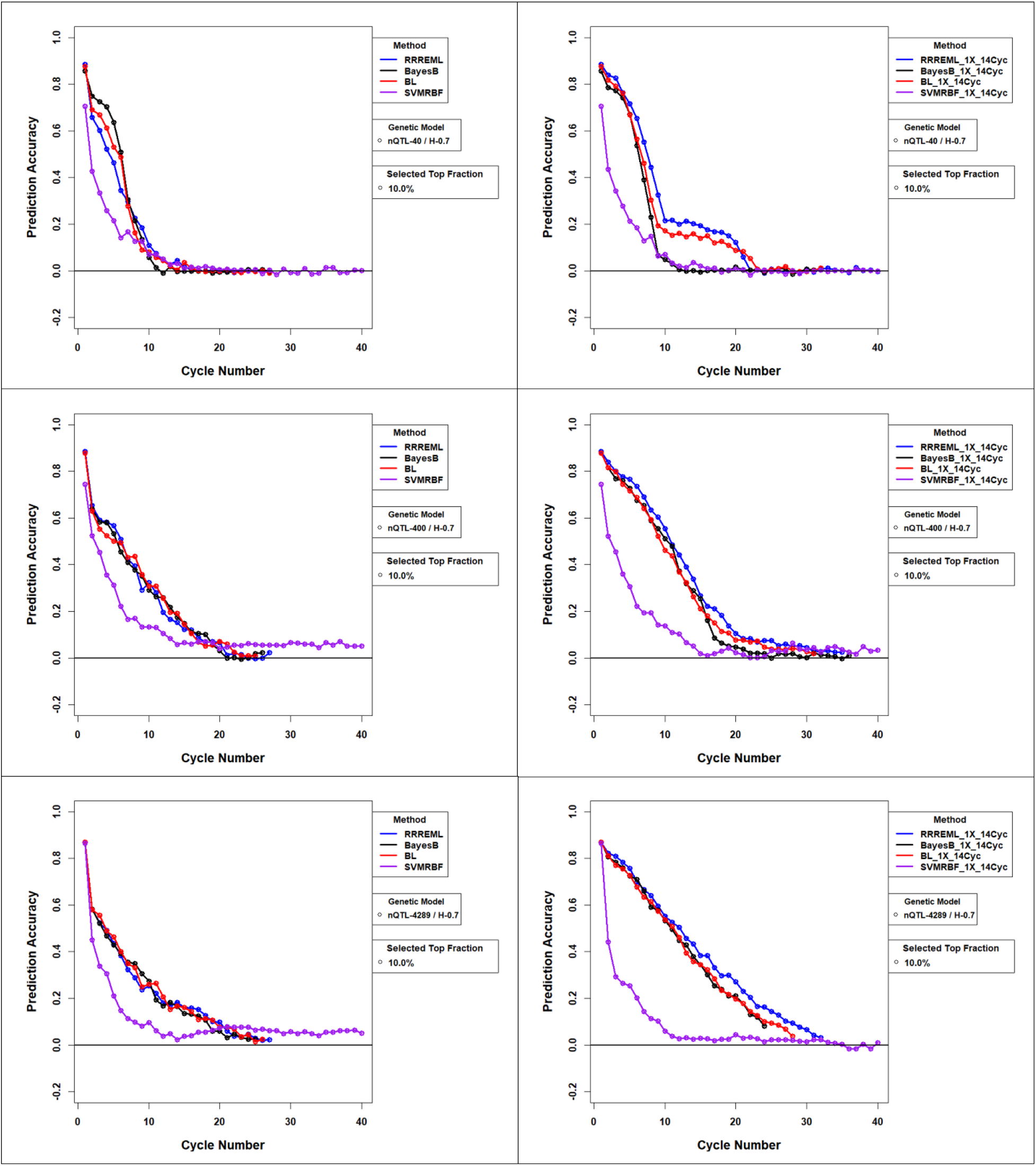
Estimated Prediction Accuracies for Comparison of GS methods with and without Updating for 0.7 H and Top 10% Selected Fraction: Estimated prediction accuracies with updates to the training sets used in genomic prediction (GP) models. Training data from up to 14 prior selection cycles were used to update all four GP models for 40 QTL (top), 400 QTL (middle) and 4289 QTL (bottom) responsible for 70% of phenotypic variability in the initial population and top 10% of RILs with the greatest predicted values. RR-REML-Ridge Regression with Restricted Maximum Likelihood, BL – Bayes LASSO, and SVMRBF-Support Vector Machine with Radial Basis Kernel.

**Figure 11.**
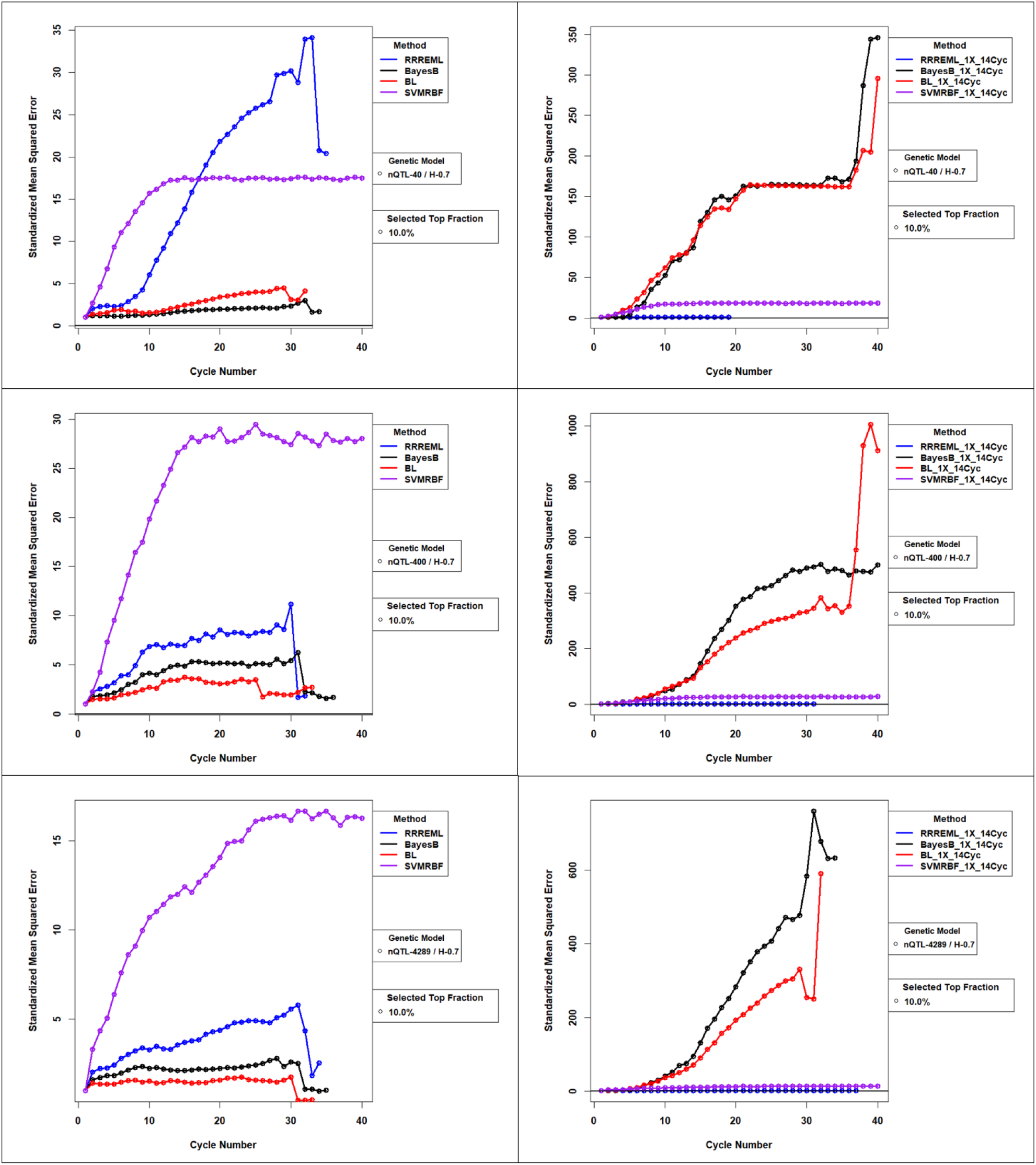
Standardized Mean Squared Error for Comparison of GS methods with and without Updating for 0.7 H and Top 10% Selected Fraction: Mean Squared Error of GP models with updates to the training sets used in genomic prediction (GP) models. Standardized MSE (>=1) is estimated as the ratio of MSE for GP models in cycle ‘c’ to MSE for GP models trained with founder population of RILs. While MSE for RRREML model were lesser with model updating, MSE for bayesian methods increased orders of magnitude in late cycles of selection with model updating. SVMEBF showed relatively constant MSE with and without updating. Training data from up to 14 prior selection cycles were used to update all four GP models for 40 QTL (top), 400 QTL (middle) and 4289 QTL (bottom) responsible for 70% of phenotypic variability in the initial population and top 10% of RILs with the greatest predicted values. RR-REML-Ridge Regression with Restricted Maximum Likelihood, BL – Bayes LASSO, and SVMRBF-Support Vector Machine with Radial

The loss of genetic potential in early cycles determines the limits to selection response in later cycles. For example, with 400 simulated QTL responsible for 70% of the phenotypic variance, the maximum potential was only 50% of the maximum potential (100 units) for PS and parametric GS methods with and without model updating. When parametric GP models are updated, 81% and 75% of the limits of maximum available potential are realized with RR-REML and Bayesian methods respectively. If GP models are not updated, only 62% of the limits of the maximum available potential are realized with RR-REML and Bayesian methods. With PS, 78% of maximum available potential is realized by the cycles in which the population no longer responds to selection. With SVMRBF, only 35 % of the potential is realized with and without updating by the cycles in which the population no longer responds to selection (Figure 5).

If GP models are not updated with data from up to 14 prior cycles, the Mgv’s were consistently greater with PS than the four GS methods. Among GP models without updating, BB provided the best Mgv, while SVM-RBF had the smallest Mgv (Figure 8 and S16). If GP models are updated, the pattern depends mostly on the number of QTL. For initial H values of either 0.7 and 0.3 and 40 simulated QTL, Mgv’s are similar for RR-REML, Bayesian GP models and PS, whereas for 400 QTL, RR-REML produces greater Mgv’s than PS and Bayesian methods. For 4289 QTL, RR-REML and Bayesian methods produce greater Mgv’s than PS. Recurrent GS with SVMRBF produced the least desirable Mgv’s for 40, 400 and 4289 QTL.

If GP models are updated, the standardized genotypic variance (Sgv) decreases at a rate similar to GP models that are not updated (Figure 9 and S17). There is no difference among GS methods in terms of rate at which Sgv decreases. Also, model updating significantly improved estimated prediction accuracies, r_ps,_ for all GP models except SVMRBF. Among RR-REML and Bayesian GP models, model updating has a slightly larger impact on estimated accuracies and MSE using RR-REML than with Bayesian GP models (Figure 10, 11, S18 and S19). MSE were orders of magnitude lesser for RR-REML than bayesian GP models with updates after the first 10-15 cycles of selection (Figure 11 and S19).

If models are updated using data from up to 14 prior cycles, the changes to genetic variance among the RILs selected to be crossed, their average heterozygosity, average rate of inbreeding, and loss of favorable alleles are similar among GS methods (Figure S29-S32). Model updating resulted in faster loss of genotypic variance among genotypes selected to be parental lines for the next cycle of inter-mating. The loss of genotypic variance is similar among parametric GS methods. (Figure S29). When there are 400 simulated QTL responsible for 70% of the phenotypic variance, the average number of favorable alleles that are lost across 40 cycles due to selection and drift are similar among PS and GS methods, but the rate at which they are lost differs among selection methods for the first 20 cycles until they converge at the same limit. SVMRBF GS showed the greatest rate of loss and PS had the least rate of loss while the parametric GS methods had intermediate rates of loss. By the time 40 cycles of selection have completed, model updating didn’t result in any significant difference in rates of loss and the total number of favorable alleles that are lost (Figure S30).

SVMRBF GS showed the greatest loss of average heterozygosity and PS lost heterozygosity at the slowest rate, while RR-REML and bayesian GS methods lost heterozygosity at intermediate rates. Model updating didn’t result in significant changes to rates at which heterozygosity was lost over cycles (Figure S31). PS showed slower rates of inbreeding than GS methods as we would expect from the decay of standardized genotypic variance. Average rates of inbreeding were similar among parametric GP models in the early cycles of recurrent selection, whereas the patterns varied after there is no response to selection (Figure S32).

For all selection methods pairwise LD among markers on the same chromosome decreased across cycles of recurrent selection (Figure S20-S24). LD decreased slowest with PS (Figure S20). Loss of LD in early and late cycles of selection are similar among parametric GP models and SVMRBF with the relaxed selection intensity. By the 20^th^ cycle of recurrent selection, LD approached zero for all selection methods and there was no evidence that selection methods affected linkage disequilibrium (LD) differentially in the earlier cycles. The rates at which LD decays are lower when GP models are updated with training sets compared to GP models without updating (Figure S20-S24).

The limiting values for RsVar (Response standardized to change in genotypic variance) when PS is used to select the best 10% of RILS with genetic architectures consisting of 400 and 4289 QTL are greater than the limiting values using GS methods without model updates (Figure S33 and S34). The parametric GP models, without model updating, resulted in similar changes of RsVar for 40, 400 and 4289 simulated QTL responsible for both 70% and 30% of phenotypic variability in the initial population. Also, if the GP models are not updated, the rates and limits to loss of RsVar are similar among the GS methods for all nQTL and SI.

If GP models are updated with the TS’s, the patterns of RsVar are significantly different among GS methods and are dependent on nQTL, SI and H (Figure S35 and S36). With 0.7 heritability, there are no significant difference in RsVar among GS methods for 40 simulated QTL. If the genetic architecture consists of 400 and 4289 QTL and weaker selection intensities are practiced, the RR-REML GS method maintained genetic variance and RsVar for more cycles than PS and the Bayesian GS methods. Relative Gain in RsVar with RR-REML GS is even larger for 0.3 H treatment with relaxed selection intensities (Figure S35 and S36). SVMRBF demonstrated the least limits of RsVar for treatment combinations with and without model updating (Figure S33 – S36). The plots for all the evaluation metrics discussed above for selection intensities 2.67 and 2.34 are provided in Figure S37 –S60 and discussed in File S6).

## Discussion

Previous publications of *in silico* investigations of factors affecting outcomes from RGS have been conducted using a few factors applied to arbitrary diploid genomes, and an expected population structure from an assumed coalescent process. To our knowledge, the research reported herein is the first designed to reveal interactions among five factors that previously had been shown to affect responses to selection. Our ability to detect and characterize interaction effects is enabled by use of a first order recurrence equation (eqn 4). Hopefully, our explanation of how to implement recurrence equation models in available R packages will encourage others to investigate recurrent genetic improvement designs. Even though our motivation for the use of non-linear modeling in this study was restricted to the systematic investigation of significance of variation in response due to the factors and the relative order of magnitude of interaction effects of factors, non-linear mixed effects models have predictive power that is still relatively unexploited in the study of limits due to recurrent selection. We plan to investigate the use of non-linear recurrence modelling in mixed model framework to estimate the magnitude of interaction effects and optimize the use of genomic information at prediction and selection levels to optimize response in recurrent selection schemes in the future.

Most *in silico* investigations of GS have simulated diploid genomes consisting of ten or fewer linkage groups with evenly spaced molecular markers depending on the organism (Jannink 2010; Liu et al. 2015; Akdemir and Sánchez 2016; Yabe et al. 2016). The organization of the soybean genome is the result of ancient duplication events (Shoemaker et al. 1996; Grant et al. 2000; Cannon and Shoemaker 2012) and consists of many copies of functional genes that are distributed unevenly with high concentrations near the telomeric ends of 20 diploid chromosomes. Also, most prior simulation studies created population structures using an assumed coalescent process (Woolliams and Corbin 2012; Hickey and Gorjanc 2012; Daetwyler et al. 2013). We used a realized structure representing adapted and selected lines by sampling the SoyNAM founders.

Structures of plant populations are highly dependent on the reproductive biology. Indeed, plant breeders design genetic improvement projects based on reproductive biology. Similar to many cereal and pulse crops, the reproductive biology of soybean is primarily through self-pollination (Wilcox et al. 1979; Fehr 1980, 1991). The frequency of natural cross pollination in soybean is only about 0.025 (Garber et al. 1925; Caviness 1966; Carlson and Lersten 1987; Ahrent and Caviness 1994). Because crossing soybean lines is labor intensive and expensive (https://www.youtube.com/watch?v=VnjGijF4KQI) soybean breeder’s use mating designs in which only one or two elite varieties are crossed with a few dozen recently selected RIL’s (Guo et al. 2013; Guo et al. 2014). We refer to this as a hub network design. Soybean breeders subsequently take advantage of natural self-pollination to create RILs with sufficient seed for replicated evaluations across many environments.

Our simulations attempted to emulate cycles of selection, crossing, and self-pollination currently conducted by commercial and academic soybean breeders. Relative to outcrossing species selecting homozygotes to participate in a hub network within each cycle will have smaller effective populations and retain LD for more cycles of recurrent selection. Future studies are needed to determine if the significant interaction effects that we found are due to population structure and genome organization.

### Implications for application of GS for genetic improvement with current soybean breeding

There has been considerable concern expressed about the limited genetic variability among soybean genotypes adapted to specific maturity zones and recurrently selected over 75 years in North America (Carter et al. 2004; Hyten et al. 2006; Mikel et al. 2010; McCouch et al. 2013). The SoyNAM founders represent a sample of current improved breeding germplasm adapted to and used for agricultural production in maturity zones II - IV (Song et al. 2017; Diers et al. 2018).

Despite concerns about limited genetic variability, our assessment of the half-lives of populations suggest that even if only 40 QTL remain in the founders of the SoyNAM population, there is still genetic potential for response to selection for at least five cycles. If there are larger numbers of QTL with smaller additive effects distributed among linkage blocks of SoyNAM founders, then we can expect reasonable half-lives to RGS for 10 cycles (File S2). Because current soybean breeding methods require four to five years before RIL’s are selected to participate in crossing nurseries, soybean breeders can expect reasonable responses to selection using samples from the SoyNAM population for 20 to 60 years. It is possible to obtain reasonable estimates of half-lives and asymptotic limits using the parameters in eqn 4 with sufficient number of simulated cycles of selection. The estimated half-lives and asymptotic limits can be used to assess the impact of selection on genetic potential of populations in the long term. In our simulations, the estimates of half-lives using 10 or 20 simulated cycles, are significantly less accurate than estimates obtained with 40 simulated cycles of recurrent selection (Table S2).

Among the five factors we investigated, a soybean breeder can choose SM’s, SI’s and TS’s. Currently, plant breeders have little control on nQTL and H, because these parameters are determined by the sample of germplasm and environments. While it is possible to adjust the value of H on an entry mean basis by increasing/decreasing the number of replicates (Fehr 1991), estimating the number of segregating QTL and the magnitude of their effects is difficult and usually extremely expensive (Beavis 1994; Goring et al. 2001; Xu 2003). For a fixed budget, the breeder will be faced with a trade-off between numbers of replicates and numbers of RIL’s that can be evaluated. In other words, H on an entry mean basis can be estimated, but not adjusted without adding field plot resources, and nQTL and their contributions to genetic variance can be estimated, but the estimates will be biased.

Although modulated by nQTL and H, PS consistently retained useful genetic variability across many cycles of genetic improvement, Bayesian methods provided the fastest genetic gains in the short term and RR-REML provided a compromise between PS and Bayesian methods. SVM-RBF should not be considered for genetic improvement for the additive genetic architectures in soybean population structures. Previously others have reported that long-term response using GS methods will be more limited than PS in closed populations (Goddard 2009; Zhong et al. 2009; Jannink 2010). We also observed that GS without updated TS’s result in rapid loss of genetic variance in the initial cycles, which results in lower Rs values as responses to selection approach an asymptotic limit. When models were updated with TS’s composed of data from prior cycles of selection, loss of prediction accuracy slowed for all values of nQTL and H (Figure 10).

Given similar rates of decreasing genetic variance among parametric GS methods, the different limits to selection response is possibly due to greater efficiency of retaining genetic diversity with RR-REML for later cycles, although for many combinations of factors the limits of response with RR-REML and Bayesian methods are about the same. Unlike previous studies, we noted in the initial cycles of RGS genetic gains and estimates of accuracy were similar using BL and RR-REML, whereas after 15 cycles genetic responses were not as limited with RR-REML, probably because additive genetic variance was retained for later cycles of selection (Liu et al. 2015).

Replicated responses to high values of SI quickly reach a limit in five to ten cycles of recurrent selection. Replicated responses to lower values of selection intensity consistently result in greater gains over more cycles, indicating that genetic drift is the most likely mechanism for loss of genotypic variance. These constraints on plant breeding programs are well characterized (Brisbane and Gibson 1995; Hayes et al. 2009; Jannink 2010; Hung et al. 2012; Liu et al. 2015; Akdemir and Sánchez 2016; Yabe et al. 2016).

Model updating with TS’s from prior cycles improves the relationship between training sets and validation sets and thus improves responses to GS without updated training sets. While model updating doesn’t significantly change the estimated half-life, model updating did produce greater responses standardized to the rate at which genotypic variance is lost in selected populations with updated RR-REML GP models (File S2; Table S3).

The choice of which combinations of SM, SI and TS depend on the objectives of the breeding program. If the objective is to enter and capture market share in a short time, then maintenance of genetic diversity is not important. Averaged over nQTL and H, the greatest changes in Rs values, i.e., rate of genetic gain, were attained in early cycles using BayesB and SI = 2.84, without model updating. Thus, if a soybean breeding project wants to maximize the rate of genetic gain for a single quantitative trait in a population derived from a sample of the SoyNAM founders, then application of the BayesB GS method and large SI is the best combination to meet the breeding objective. If the breeding project has a long term breeding objective to improve germplasm while maintaining useful genetic diversity for purposes such as providing useful germplasm for future generations or evaluating genome editing, then PS or GS with RR-REML with relaxed SI will be the best combination to meet the breeding objective. The greatest values for Rs were attained using the RR-REML model with model updating and SI = 1.75. Last if the breeding project has multiple objectives for both immediate and longer term goals, then pareto optima among tradeoffs involving responses/cycle and retention of useful genetic variance for multiple traits need to be identified (Akdemir et al. 2019).

### Lessons for future simulation studies

In our simulations we assigned the same amount of time to develop and evaluate RILs for GS as for PS. Application of PS usually requires field trials for three years before lines are selected for use in a crossing nursery. In practice, one of the advantages of GS relative to PS is that only subsets of RILs need to be phenotyped. Commercial soybean breeding projects have used genotypic values obtained from GP models to cull lines in as they are being self-pollinated and prior to the first stage of field trials (Kurek 2018). Also it is possible to train or update GP models with lines derived in earlier filial generations thereby requiring less time per cycle (Bassi et al. 2016). Even if both GS and PS require the same amount of time to develop lines before phenotypic evaluation, GS methods can be conducted as soon as phenotypic information is available from first year field trials (Heffner et al. 2009), while PS usually requires phenotypic evaluations for three years before lines are selected for crossing nurseries to begin a new cycle. Herein we did not investigate the trade-offs between less accurate predictions from models trained with less extensive phenotyping or phenotyping with lines derived in earlier filial generations. These practical adjustments from application of GS methods to current soybean genetic improvement projects still need to be investigated.

Accuracies of GP models in the founding set of RILs were similar to that reported in previous studies (Long et al. 2010, 2011; Guo et al. 2012; Howard et al. 2014). However, our estimates of accuracies are larger as we’ve included QTL in our training sets. Relationship among selected RILs and LD between marker loci (ML) and QTL are considered the two most important sources of prediction accuracy. In previous studies, relationship among RILs had a greater effect on prediction accuracies for RR-REML than BL, whereas accuracy of BL was more dependent on LD and both components showed similar contribution to the accuracy of BayesB GP models (Habier et al. 2007; Zhong et al. 2009; Liu et al. 2015). Similar to our results without model updating, Bayesian GS methods resulted in greater responses as the populations approached asymptotic limits (Meuwissen 1997; Li et al. 2008; Akdemir and Sánchez 2016). However, GP model updating reduced the difference in rate of decrease of prediction accuracies among RR-REML and Bayesian GP models but there was no consistent pattern in relationship among selected RILs and rate of loss for LD to explain the observed estimates of prediction accuracies. However, GP model updating consistently resulted in lesser MSE for RR-REML than BB and BL GP models across all levels of SI, nQTL and H. This pattern is consistent with greater efficiency of converting loss of variance into gain with updated RR-REML GS method. In order to estimate the contribution of LD and linkage blocks to prediction accuracy of GP models will require a design similar to that employed by Müller et al. (2017, 2018) for synthetic populations. Populations with unrelated training and prediction sets with LD and SNP based relationship estimates showed low prediction accuracy and low genetic response in recurrent GS similar to GS without updating in this study. Whereas populations with relationship between training and prediction sets with LD and SNP based relationship estimates showed greater prediction accuracy and greater genetic response similar to GS with model updating (Müller et al. 2017, 2018).

While TS’s had relatively small impacts on response metrics, they were highly significant. Since relatedness of TS’s and validation sets affect estimates of prediction accuracy it has been suggested that model updating needs to be evaluated for accuracies of prediction within and across populations (Crossa et al. 2014; Juliana et al. 2018; Stewart-Brown et al. 2019). Herein, the TS’s were comprised of a combined population of RIL’s derived from all families across cycles of selection. Given a constant number of RILs in the TS’s, within or across family prediction accuracies will depend on genetic differentiation among families (de Roos et al. 2009; Schulz-Streeck et al. 2012; Stewart-Brown et al. 2019). However, actual soybean breeding projects evaluate a few to many dozen RIL’s per family and future simulation studies, especially of two part systems, should consider whether relationships between evaluation units, RIL’s and selection units (Holland et al. 2003), possibly individuals, need to be used in design of the TS’s.

We allowed the size of TS’s to increase every cycle by adding data from prior cycles. Increasing the number of RIL’s per TS requires more computational resources. An alternative strategy is to randomly sample subsets of data from each of the prior cycles to maintain a constant cumulative training population size. It is also possible to assign weights to the samples from prior cycles to place more weight on data from more recent cycles. These possibilities suggest determination of an optimal combination of numbers of RIL’s and weights that will provide maximum prediction accuracy with minimal computational requirements. Some aspects of this optimization problem have been addressed (Lorenz et al. 2013; Hickey et al. 2014; Akdemir et al. 2015; Xavier et al. 2017). For example, Akdemir et al (2015) devised a genetic algorithm for selecting optimal training populations to minimize prediction error variance and Xavier et al (2017) developed sampling methods for training Bayesian GP models. Another consideration is whether TS’s need to be updated every cycle. Instead of updating the model with data from every cycle would it be more effective to retrain GP models every second third, fourth… cycle while maintaining a constant training population size? Modifications to design of TS’s will need to be addressed with simulations before implementation.

Like most simulation studies we fixed levels of SI as constant for all combinations of factors across all cycles of recurrent selection. This is not consistent with actual soybean breeding projects. Just as sampling families result in some with exceptional genotypes and some with poor genotypes, in an actual genetic improvement project there is variability among cycles. The effects of applying a dynamic selection strategy is an alternative and interesting question. We hypothesize that a strategy consisting of applying different SI’s, optimized for each cycle, will achieve improved long-term genetic response by differentially emphasizing genetic variance and genetic gain across multiple cycles of selection.

Unlike actual soybean genetic improvement projects we simulated a closed breeding population derived from a sample of SoyNAM founders in which culled lines were not resampled for discarded favorable alleles, nor did we simulate exchange of lines among breeding projects. In MZ’s II, III and IV there are six public breeding projects and about a dozen commercial soybean breeding projects. Thus, there is potential, depending on material transfer agreements, to exchange genotypes among breeding populations. In our next set of investigations migration among multiple breeding projects will be evaluated for response to recurrent selection within and among breeding projects using island model evolutionary algorithms (Hagan et al. 2012; Yabe et al. 2016). Results reported herein will provide comparators for assessing impacts of migration policies relative to the other factors that affect responses to RGS.

Also, as with prior simulation studies, we simulated truncation selection, but unlike previous investigations we did not randomly mate selected lines each cycle. Rather, we used the hub network design (Guo et al. 2013; Guo et al. 2014). We did not consider relationships among selected RILs nor the trade-offs between genetic gain, genetic variance (inbreeding) when selecting RILs to cross. There exist multi-objective optimization methods such as genomic mating and optimal cross selection (Rutten et al. 2002; Woolliams et al. 2015; Akdemir and Sánchez 2016; Gorjanc et al. 2018; Akdemir et al. 2019) that have been demonstrated to provide both greater rates of genetic gain and assure maintenance of population genetic potential across cycles.

In the future, it is possible that development of male sterile and insect pollination systems for soybean (Ortiz-Perez 2008; Davis 2020) will enable a cycle of RGS to be conducted using two or three mating generations per year. This will enable an alternative genetic improvement system based on decoupling genetic improvement from variety development (Gaynor et al. 2017). By separating the two types of breeding projects, GS can be applied continuously. In the two part system TS’s will need to be composed of genotypic and phenotypic data obtained from annual field trials of RIL’s, although the TS’s will be several selection cycles removed from the cycle used to create the RILs.

Results reported herein suggest that RGS in a two part system will rapidly produce genetic gains and loss of useful genetic variance in very short periods of time. Indeed, 40 cycles of RGS could be completed in 12 to 20 years. To offset the shorter cycle times, application of GP models to select and cross individual progeny instead of RIL’s could result in a larger effective population size per cycle by creating more opportunities for recombination and slow the unintended loss of valuable alleles in discarded linkage blocks. However, before investments in development of male sterile and insect pollination systems for soybean (Ortiz-Perez 2008; Davis, personal communication) research using simulations are needed to understand the trade-offs and whether the investments are justified using approaches from Operations Research (Xu et al. 2011; Cameron et al. 2017, Han et al. 2017).

## Acknowledgements

Funding for this research was provided by the Department of Agronomy, Iowa State University, the North Central Soybean Research Program and an NSF grant (1830478). Supplemental funding for large scale computing was enabled by the Extreme Science and Engineering Discovery Environment (XSEDE), which is supported by National Science Foundation. XSEDE resources consisted of research allocations (DMS190015 & DMS190018) on PSC-Bridges Large Memory nodes for the simulations involving updating of parametric GP models. The Iowa State University-Pronto GPU cluster enabled computation of SVM model update simulations. We want to acknowledge Matheus de Krause for discussions on fitting non-linear models using ‘nlme’ package, Dr. Lizhi Wang for efficient programs to simulate meiosis and Dr. Alencar Xavier for sharing an efficient expectation maximization method for fitting ridge regression GP models. Last, we want to thank anonymous critical reviewers of earlier versions of this manuscript for providing useful suggestions on how to present such a large volume of information from such a comprehensive investigation of expected short and long term outcomes of GS applied to soybeans adapted to MZ’s II to IV.

