## Supplementary material for "Factors Affecting Response to Recurrent Genomic Selection in Soybeans": File S4

**File S4: Comparison of responses from GS Methods when simulated QTL are responsible for 70% (0.7 H) and 30% (0.3 H) of phenotypic variability in the initial population**

**Responses from GS Methods for 0.7 H**

If the BB models are not updated with data from prior cycles, then the Rs from 40 simulated QTL and 0.7 H in the initial population were 10 to 16% greater than they were with PS in the first five cycles. For the same genetic architecture, recurrent selection with BL and RR models resulted in Rs values that were 4 to 13% greater than Rs values from PS in the first five cycles (Figure 6 and File S3). After five cycles, PS resulted in greater responses than all of the GS methods when training sets were not updated. If training sets were not updated after the initial evaluations of RILs and the genetic architectures consisted of 400 and 4289 QTL responsible for 0.7 H in the initial sets of RILs, then RR and Bayesian GS methods provided greater genetic responses than PS only in the first 2-3 cycles and thereafter PS demonstrated 5-50 % greater standardized genetic responses (File S3).

When the RR GP model is updated with data from up to 14 previous cycles of recurrent selection, the Rs values for selection on 40 simulated QTL responsible for 70% of phenotypic variability in the initial sets of RILs was 1.5 to 10% greater than PS for the first 10 cycles of recurrent selection, but after the 10^th^ cycle it was similar to PS (Figure 4 and 7; File S3). For the same genetic architecture the Rs values from BB and BL with model updating were 14.4% and 12% greater than PS respectively for the first five cycles of recurrent selection. After the fifth cycle PS resulted in greater Rs values. If the RR model is updated with data from up to 14 previous cycles of recurrent selection, the responses to selection of RILs with 400 simulated QTL responsible for 70% of phenotypic variability in the initial sets of RILs, then the Rs values were 3 to 15% greater than PS across all 40 cycles. The limits of response to selection using BB and BL GS methods were 1 to 14% greater than PS for up to 20 cycles. If the RR model is updated with data from 14 previous cycles of recurrent selection, the Rs values with 4289 QTL were 10-15% greater than PS for 40 cycles (Figure 4 and 7; File S3). Likewise recurrent selection using BB and BL models resulted in greater responses than PS for 40 cycles (Figure 4 and 7; File S3).

**Responses from GS methods with model updating compared to responses from GS methods without updating**: When GP models are updated, RR models resulted in 10% greater response than RR without updating for 40 simulated QTL responsible for 70% of phenotypic variability among RILs in the initial cycle. Model updating resulted in 30% and 60% greater responses for 400 and 4289 QTL respectively (Figure S27). Recurrent selection with updated BB models resulted in 4% greater Rs values than without updating for 40 QTL and resulted in 22% and 57% greater responses for 400 and 4289 simulated QTL respectively (Figure S27). Recurrent selection with updated BL models resulted in 3% greater responses than without updating for 40 simulated QTL. If the genetic architecture consisted of 400 QTL and 4289 QTL, updated BL models resulted in 21% and 51% greater responses respectively (Figure S27).

**Responses from GS Methods for 0.3 H**

When GP models are updated with up to 14 prior cycles of training sets, parametric GS methods demonstrated 10-60% greater responses in the early cycles of selection relative to PS for genetic architectures consisting of 40, 400 and 4289 simulated QTL and H values of 0.3 (Figure S25 and S26; File S3), whereas selection based on SVM-RBF resulted in lesser responses than all other selection methods across all cycles.

For 40 simulated QTL responsible for 30% of phenotypic variability in the initial population, RR GS when updated with training sets from up to 14 prior cycles resulted in 25 - 47% greater response relative to PS in early cycles whereas PS demonstrated about 5 % greater Rs values as the limits to selection were approached (Figure S15, S25 and S26). For the same genetic architecture, GS using Bayes-B and BL models with training sets from up to 14 prior cycles resulted in 10 to 53 % and 5.4 - 49.2% greater Rs values than PS in the first ten cycles (Figure S15, S25 and S26).

If the genotypic variance was responsible for 30% of phenotypic variability in the initial population, GS with RR demonstrated 12-42% and 22-48% greater standardized responses than PS up to the fortieth cycle for 400 and 4289 QTL (Figure S15, S25 and S26). For 400 QTL, selection using Bayes B provided 0.6-42% greater responses than PS up to the thirtieth cycle. After the thirtieth cycle, PS demonstrated greater responses. For 4289 QTL, selection with Bayes B resulted in 3.7 to 44.4% greater responses than PS up to the fortieth cycle. For 400 QTL, GS with BL models resulted in 5.1-42% greater responses than PS up to 25^th^ cycle. After the 25^th^ cycle, PS demonstrated greater responses. For 4289 QTL, selection with BL resulted in 1.8 to 42.7 % greater responses than PS up to fortieth cycle (S15, S25 and S26; File S3).

PS demonstrated greater responses than recurrent selection using the SVMRBF models for Rs values created by 40, 400 and 4289 simulated QTL responsible for 0.7 and 0.3 of the phenotypic variance in the initial sets of RILs (Figure 4, 6, 7, S15, S25 and S26; File S3). Complete set of percent gain in response relative to response from PS for forty cycles are provided in File S3.
