## Supplementary material for "Factors Affecting Response to Recurrent Genomic Selection in Soybeans": File S6

**File S6: Impact of Selection Intensity on Response for Combinations of Treatment Factors**

When RR models are updated and applied to initial sets of RILs with genotypic architectures consisting of 40 QTL responsible for 70% of the phenotypic variance in the initial population using a stringent selection intensity, the responses were 5.8% greater than responses without updating. In contrast, less stringent selection intensities with updated RR models resulted in 7.1% and 10.6% greater responses than application of RR without updating (Figure S27, S41 and S43).

For 400 QTL, 1% selected fraction with updated RR GS resulted in 22.2% greater responses than RR GS without updating, whereas 2.5 and 10% selected fraction resulted in 22.26% and 30.8% greater responses respectively with model updating. For 4289 QTL, 1% selected fraction with RR GS resulted in 42.96% greater responses with model updating, whereas 2.5 and 10% selected fraction resulted in 52.74 and 65.8% greater responses with model updating respectively (Figure S27, S41 and S43).

With 0.3 heritabilities, 1% top selected fraction and 40 QTL, RR GS when updated demonstrated 5.52% greater response than RR GS without updating. Whereas with 2.5 and 10%, updated RR GS demonstrated 26.73 % and 21.27% greater limits of responses than RR GS without updating for 40 QTL. For 400 QTL, 1% selected fraction with RRGS resulted in 43.6% greater limits of responses than updated RRGS, whereas 2.5 and 10% selected fraction resulted in 46.6% and 64.5% greater limits of responses with model updating respectively. For 4289 QTL, 1% selected fraction with RR GS resulted in 75.46% greater limits of responses with model updating, whereas 2.5 and 10% selected fraction resulted in 88.4% and 132.9% greater responses with model updating respectively (Figure S28, S42, and S44).

BayesB and Bayes LASSO methods have similar patterns of response with small differences among selection intensities, but result in lower limits of responses compared to RR method. For 0.7 heritabilities, with 1% top selected fraction and 40 QTL, BB GS when updated demonstrated 1% greater response than BB GS without updating. Whereas for 40 QTL, selection with 2.5% and 10% selected fraction updated BB GS demonstrated 1% lesser and 3.6% greater limits of response than BB GS without updating. For 400 QTL, 1% selected fraction with BBGS resulted in 10.85% greater responses than updated BB GS, whereas selection with 2.5% and 10% selected fraction resulted in 21.7% and 21.8% greater responses with model updating respectively. For 4289 QTL, 1% selected fraction with BB GS resulted in 37.8% greater responses with model updating, whereas selection with 2.5% and 10% selected fraction resulted in 36.0% and 56.7% greater responses with model updating respectively (Figure S27, S41 and S43).

For 0.3 heritabilities, selection with 1% top selected fraction and 40 QTL, BB GS when updated demonstrated 3.63% greater response than BB GS without updating. Whereas for 40 QTL, selection with 2.5 and 10% selected fraction with updated BB GS demonstrated 14.07% lesser limits of response and 11.43% greater limits of response than BB GS without updating respectively. For 400 QTL, 1% selected fraction with updated BBGS resulted in 27% greater responses than BB GS without updating, whereas 2.5 and 10% selected fraction resulted in 36.9% and 39.8% greater responses with model updating respectively. For 4289 QTL, 1% selected fraction with BB GS resulted in 49.8% greater responses with model updating, whereas selection with 2.5% and 10% selected fraction resulted in 66.4 and 85.7% greater responses with model updating respectively (Figure S28, S42 and S44).

For 0.7 heritabilities, with 1% top selected fraction and 40 QTL, BL GS when updated demonstrated 4.8% greater response than BL GS without updating. Whereas selection with 2.5 and 10% selected fraction with updated BL GS demonstrated 9.85 and 3.0% greater limits of response than BL GS without updating for 40 QTL. For 400 QTL, 1% selected fraction with updated BL GS resulted in 17.2% greater responses than BL GS without updating, whereas 2.5 and 10% selected fraction resulted in 16.9% and 21.4% greater responses with model updating respectively. For 4289 QTL, 1% selected fraction with BL GS resulted in 32.0% greater responses with model updating, whereas 2.5 and 10% selected fraction resulted in 46.21 and 51% greater responses with model updating respectively (Figure S27, S41 and S43).

For 0.3 heritabilities, with 1% top selected fraction and 40 QTL, BL GS when updated demonstrated 2.3% greater response than BL GS without updating. Whereas with 2.5 and 10%, updated BL GS demonstrated 11.3 and 11.0% greater limits of response than BLGS without updating for 40 QTL. For 400 QTL, 1% selected fraction with BLGS resulted in 19.2% greater responses than updated BLGS, whereas 2.5 and 10% selected fraction resulted in 43.6% and 44.4% greater responses with model updating respectively. For 4289 QTL, 1% selected fraction with BL GS resulted in 50.2% greater responses with model updating, whereas 2.5 and 10% selected fraction resulted in 78.9 and 83.6% greater responses with model updating respectively (Figure S28, S42 and S44).

For SVMRBF, model updating significantly improved response only with stringent selection intensity of 2.67, whereas responses to selection intensities of 2.34 and 1.75 using SVMRBF with updated training sets didn’t result in relative improvements (Figure S27, S28, and S41 – S44). Percent gain in response with model updating relative to GS without updating for forty cycles of selection are provided in File S5.
