## Supplementary material for "Factors Affecting Response to Recurrent Genomic Selection in Soybeans": File S1: DESCRIPTION

Package: SoyNAMPredictionMethods
Type: Package
Title: Evaluate genomic prediction model performance in terms of
        long-term genetic response in recurrent selection of SoyNAM
        populations
Version: 0.1.0
Author: Vishnu
Maintainer: Vishnu <>
Description: Data and simulation analysis package to evaluate long-term response in recurrent selection with phenotypic selection and genomic selection of populations with SoyNAM structure
License: What license is it under?
LazyData: TRUE
Imports:
        Rcpp,adegenet,mmod,bigmemory,rrBLUP,foreach,doParallel,BGLR,caret,kernlab,DSL,reshape2
Depends: Rcpp, adegenet, mmod, bigmemory, rrBLUP, foreach, doParallel,
        BGLR, caret, kernlab, DSL, reshape2, R (>= 2.10)
LinkingTo: Rcpp
NeedsCompilation: yes
Packaged: 2020-10-30 04:56:22 UTC; vishnur
