## Supplementary material for "Factors Affecting Response to Recurrent Genomic Selection in Soybeans": File S1: EarlyCycles_GSMethods.r

```
#############
 

getQTLIndices <- function(nLoci){

   nMarkers <- 4289
   nCycles <- 40

  if(nLoci== 40){
      QTL_indices<- seq(50,(nMarkers/2),by=53)
    }


   if(nLoci == 400){
      QTL_indices_1<- seq(8,nMarkers,by=12)
      QTL_indices_2 <- seq(50,nMarkers,by=107)
      n_3 <- 400-(length(QTL_indices_1))-(length(QTL_indices_2))
      QTL_indices_3<- sample(c(1:nMarkers),n_3)

      QTL_indices <- c(QTL_indices_1,QTL_indices_2,QTL_indices_3)
    }

    if(nLoci == nMarkers){
      QTL_indices <- c(1:nMarkers)
    }

    return(QTL_indices) 

 }
 
#############
 
getLostGenoValue <- function(MAF_List_nTS,FavAlleleList,QTL_Indices_400){

 MAF_List <- MAF_List_nTS
 nSelCriteria <- length(MAF_List)
 nRep <- length(MAF_List[[1]][[1]])
 nCycles <- 40 
 FavAllele_List <- list()
 FavAll_Freq_List_Cyc <- rep(list(rep(list(list()),nRep)),nCycles)
 FavAll_Freq_List_Met <- rep(list(rep(list(rep(list(list()),nRep)),nCycles)),nSelCriteria)
 QTL_Freq_List_Cyc <- rep(list(rep(list(list()),nRep)),nCycles)
 QTL_Freq_List_Met <- rep(list(rep(list(rep(list(list()),nRep)),nCycles)),nSelCriteria) 
 
 

 for(nSC in 1:nSelCriteria){

  for(nCyc in 1:nCycles){

   for(nrep in 1:nRep){ 

    FavAllele_List[[nrep]] <- FavAlleleList[[nrep]][[2]] 
 
    FavAllele <-  FavAllele_List[[nrep]]
    
    FavAll_Freq_List <- rep(0,4289)
    QTL_Freq_List <- rep(0,400)
    
	evenIndices<- seq(2,400,by=2)
	oddIndices <- seq(1,399,by=2)

   
	QTL_Freq_List[evenIndices] <- 1-MAF_List[[nSC]][[nCyc]][[nrep]][QTL_Indices_400[evenIndices]]
	QTL_Freq_List[oddIndices] <- MAF_List[[nSC]][[nCyc]][[nrep]][QTL_Indices_400[oddIndices]]
	
    FavAll_Freq_List_Cyc[[nCyc]][[nrep]]<- FavAll_Freq_List
    QTL_Freq_List_Cyc[[nCyc]][[nrep]] <- QTL_Freq_List

   }
  }
    FavAll_Freq_List_Met[[nSC]] <- FavAll_Freq_List_Cyc
    QTL_Freq_List_Met[[nSC]] <- QTL_Freq_List_Cyc
}

############## 
 
 QTL_indices <- QTL_Indices_400
 
 nCycles <- 40
 
 FavAll_Table <- matrix(rep(0,nRep*4289),nrow=nRep,ncol=4289)
 FavAll_List <- rep(list(rep(list(list()),nRep)),nCycles)

 Lost_FavAll_Table <- rep(0,nRep)
 Lost_FavQTLAll_Table <- rep(0,nRep)

 Avg_FavAll_List <- rep(list(rep(list(list()),nCycles)),nSelCriteria)
 Tot_FavAll_List <- rep(list(rep(list(list()),nCycles)),nSelCriteria)
 Tot_FavQTL_List <- rep(list(rep(list(list()),nCycles)),nSelCriteria)

 Tot_LostFavAll_List <- rep(list(rep(list(list()),nCycles)),nSelCriteria)
 Tot_LostFavQTLAll_List <- rep(list(rep(list(list()),nCycles)),nSelCriteria)

Lost_FavQTLAll_Indices <- rep(list(list()),nRep)
Lost_FavQTLAll_Indices_List <- rep(list(rep(list(list()),nCycles)),nSelCriteria)

 Lost_Genetic_Value <- rep(list(list()),nRep)
 Lost_Genetic_Value_List <-rep(list(rep(list(list()),nCycles)),nSelCriteria)
 Lost_FavQTLAll_Freq <- rep(0,nRep)
 Lost_FavQTLAll_Freq_List <- rep(list(rep(list(list()),nCycles)),nSelCriteria)
 
 for(nSC in 1:nSelCriteria){
  for(nCyc in 1:40){
    for(nrep in 1:nRep){

				
		Lost_FavQTLAll_Freq[nrep] <- length(which(QTL_Freq_List_Met[[nSC]][[nCyc]][[nRep]] %in% 0))
		
		Lost_Genetic_Value[[nrep]] <- Lost_FavQTLAll_Freq * 0.5
		
   
    }
	    Lost_FavQTLAll_Freq_List[[nSC]][[nCyc]] <- mean(Lost_FavQTLAll_Freq)
					
		Lost_Genetic_Value_List[[nSC]][[nCyc]] <- mean(unlist(Lost_Genetic_Value))

  }
 }
 
	return(list(Lost_FavQTLAll_Freq_List,Lost_Genetic_Value_List))
	
}

##############	
	
getLostGenoValuePlot <- function(Lost_FavQTLAll_Freq_List_nCyc,TrainCycle,ncycles){
	
  Lost_FavQTLAll_Freq_List <- Lost_FavQTLAll_Freq_List_nCyc[[1]]
  nCycles <- ncycles    
	
  GenoVal_Sim_NX_2k_List_3c_PhenoPred_GP_List2 <- GenoVal_Sim_NX_2k_List_3c_PhenoPred_GP_List_TrainSet_Sel3
		 
  trainCycle <- TrainCycle
	
  Gt_List <- list()
  Tot_MaxPotential_V2_List <- list()

  GenoVal_Sim_NX_2k_List_3c_PhenoPred_GP_List <- GenoVal_Sim_NX_2k_List_3c_PhenoPred_GP_List2 
		
	nSelMethod <- length(GenoVal_Sim_NX_2k_List_3c_PhenoPred_GP_List)
	  
	for(nSel in 1:nSelMethod){

		GenoVal_Sim_NX_2k_List_3c_PhenoPred_GP <- GenoVal_Sim_NX_2k_List_3c_PhenoPred_GP_List[[nSel]][[trainCycle]]
     		
	 	nReps <- 10
		nreps <- nReps
		cycleNo <- c(1:nCycles)
	
		leg_col <- c("dark green","blue","black","red","purple","dark orange","gold3")
			
		nSelSch <- 3
			
		lineType <- 1
		pchar <- 1
						
		yoffset1 <- 100 
		yoffset2 <- 150
		
		ymax <- 200			
				
		Rt <- rep(0,nCycles)
		Rs <- rep(0,nCycles)
		pValue.t <- rep(0,nCycles)

				Gt_Rep <- matrix(rep(0,nreps*nCycles),nrow=nreps,ncol=nCycles)
				GVart_Rep <- matrix(rep(0,nreps*nCycles),nrow=nreps,ncol=nCycles)
				
				
				## mean genotypic value of a population of 2000 individuals every generation for 10 replicates 
				## genotypic variance - variance of genetic values of a ppoulation every generation for 10 replicates 
				## Average of mean genotypic value of 10 reps every generation 
				## Average of genotypic variance of 10 reps every generation 			
				
				
				for(nReps in 1:nreps){ 
				 
				  for (t in 1:nCycles){
					
					Gt_Rep[nReps,t] <- mean(GenoVal_Sim_NX_2k_List_3c_PhenoPred_GP[[nCon]][[nReps]][,t])
					GVart_Rep[nReps,t] <- sd(GenoVal_Sim_NX_2k_List_3c_PhenoPred_GP[[nCon]][[nReps]][,t]) 
				
				  }
				}

			Gt <- apply(Gt_Rep,2,mean)
			Gt_List[[nSel]] <- Gt
					
                   
			Tot_MaxPotential <- (200- (unlist(Lost_FavQTLAll_Freq_List[[nSel]]) * 0.5))
			
			Tot_MaxPotential_V2 <- (Tot_MaxPotential[c(1:(nCycles))])
			
			Tot_MaxPotential_V2_List[[nSel]] <- Tot_MaxPotential_V2

			if(nSel==1){ 
			
				filename <- paste("GSMethods_EarlyCycles_GenoValues","_",nCon,"_",trainCycle,"Cycset",nCycles,"Cyc",".png",sep="")
				png(filename,width=1024,height=768,pointsize=20)
				par(oma=c(0,0,0,6),mar=c(4,8,4,12)) 

				plot(cycleNo,Tot_MaxPotential_V2[1:(nCycles)],type="l",xlim=c(0,nCycles),ylim=c(ymin,ymax),lty=3,lwd=4,col=leg_col[nSel],xlab="Cycle",ylab="Average Genotypic Value",main="",font.lab=2) 
	
				lines(cycleNo,c(Gt[1:nCycles]),type="l",lty=1,lwd=4,col=leg_col[nSel])
	
			}else if(nSel >1){ 
 
				lines(cycleNo,Tot_MaxPotential_V2[1:(nCycles)],type="l",lty=3,lwd=4,col=leg_col[nSel])
				lines(cycleNo,c(Gt[1:nCycles]),type="l",lty=1,lwd=4,col=leg_col[nSel])
			}
}

 

    
    nSelSch <- 3		
	index <- 1
	Sellabel <- c("1.0%","2.5%","10.0%")
	SelCondition <- c("nQTL-40 / H-0.7","nQTL-40 / H-0.3","nQTL-400 / H-0.7","nQTL-400 / H-0.3","nQTL-4289 / H-0.7","nQTL-4289 / H-0.3")
	if(trainCycle ==1){
		SelMethod <- c("PS","RRREML","BayesB","BL","SVMRBF")
		
		}
	if(trainCycle ==4){
		SelMethod <- c("PS","RRREML_1X_14Cyc","BayesB_1X_14Cyc","BL_1X_14Cyc","SVMRBF_1X_14Cyc")
	}	
	x_c <- nCycles+1.5
	
	y_Value <- c("Genotypic Value","Maximum Potential")
		
	par(font=2)
	legend(x_c,ymax,legend=SelMethod[1:nSelMethod],xpd=TRUE,lty=c(1,1),lwd=c(3,3),col=leg_col,cex=0.7,title="Method") 
	legend(x_c,ymax-65,legend=y_Value[1:2],xpd=TRUE,col="black",lty=c(1,3),lwd=c(3,3),title="Y_Value",text.width=strwidth("Maximum Potential"),cex=0.70)
	
	legend(x_c,ymax-yoffset1,legend=SelCondition[nCon],xpd=TRUE,col="black",pch=c(1,3,5),title="Genetic Model",text.width=strwidth("Genetic Model")+0.05,cex=0.7)
	legend(x_c,ymax-yoffset2,legend=Sellabel[nSelSch],xpd=TRUE,col="black",pch=c(1,3,5),title=" Selected Top Fraction",text.width=strwidth("Selected Top Fraction"),cex=0.7) 
		
	dev.off()
	
	return(list(Tot_MaxPotential_V2_List,Gt_List))
	
}

##############
```
