## Supplementary material for "Factors Affecting Response to Recurrent Genomic Selection in Soybeans": File S1: functionBayesAModel.R

```
buildBayesAModel <- function(trainGenoTable,trainPhenoTable,testGenoTable,testPhenoTable,heritability){

	
	 trainPheno <- trainPhenoTable[,1]
   trainGeno <- trainGenoTable
	 h2 <- heritability
	 nIndividuals <- 2000


	 predBayesA <- BGLR(trainPheno,response_type="gaussian",ETA=list(list(X=trainGeno,model="BayesA")),nIter=41000,burnIn=1000,df0=4,R2=h2)

	 #nIter=41000,burnIn =1000, df0=4, R2=0.7


###########################################################################################################################################3
############################################################################################################################################

  testPheno <- testPhenoTable[,1]
  testGeno <- testGenoTable

  train_nIndividuals <- (0.8*nIndividuals)
  test_nIndividuals <-  (0.2*nIndividuals)

  # pred.valid <- predBayesA$ETA[[1]]$b
  Pred.pheno.train <- (trainGeno %*% predBayesA$ETA[[1]]$b)
  Pred.pheno.valid <- (testGeno %*% predBayesA$ETA[[1]]$b)

  # Pred.pheno.valid <- predict(testGeno,predBayesA)

  # Pred.pheno.valid <- predBayesA$ETA[[1]] + (as.vector(pred.valid))

  Pred.Accuracy <- cor(Pred.pheno.valid,testPheno)

  train_MSE <- (sum((Pred.pheno.train-trainPheno))^2)/(train_nIndividuals)
  test_MSE <- (sum((Pred.pheno.valid-testPheno))^2)/(test_nIndividuals)

  sol <- list(Pred.Accuracy,predBayesA,predBayesA$ETA[[1]]$b,train_MSE,test_MSE)

  return(sol)


}
```
