## Supplementary material for "Factors Affecting Response to Recurrent Genomic Selection in Soybeans": File S1: functionBayesBModel.R

```
###Function for BayesB model

buildBayesBModel <- function(trainGenoTable,trainPhenoTable,testGenoTable,testPhenoTable,heritability){ 

	
	 trainPheno <- trainPhenoTable
     trainGeno <- trainGenoTable
	 h2 <- heritability
	 nIndividuals <- 2000
	 
 	 train_nIndividuals <- (0.8*nIndividuals)
     test_nIndividuals <-  (0.2*nIndividuals)
   
	 
	 predBayesB <- BGLR(trainPheno,response_type="gaussian",ETA=list(list(X=trainGeno,model="BayesB")),nIter=41000,burnIn=1000,df0=4,R2=h2,verbose=FALSE)
	 
	 	
	
###############################################################################3
 
  testPheno <- testPhenoTable
  testGeno <- testGenoTable 
  
  Pred.pheno.train <- (trainGeno %*% predBayesB$ETA[[1]]$b) 
  Pred.pheno.valid <- (testGeno %*% predBayesB$ETA[[1]]$b) 
  
  # Pred.pheno.valid <- predBL_BRR$mu + (as.vector(pred.valid))
  
  Pred.Accuracy <- cor(Pred.pheno.valid,testPheno)
  
  train_MSE <- (sum((Pred.pheno.train-trainPheno))^2)/(train_nIndividuals)
  test_MSE <- (sum((Pred.pheno.valid-testPheno))^2)/(test_nIndividuals) 
  
  sol <- list(Pred.Accuracy,predBayesB,predBayesB$ETA[[1]]$b,train_MSE,test_MSE)
  
  return(sol)


}
```
