## Supplementary material for "Factors Affecting Response to Recurrent Genomic Selection in Soybeans": File S1: functionBLModel.R

```
buildBLModel <- function(trainGenoTable,trainPhenoTable,testGenoTable,testPhenoTable,heritability){ 

	
	 trainPheno <- trainPhenoTable
         trainGeno <- trainGenoTable
	 h2 <- heritability
	 nIndividuals <- 2000
	 train_nIndividuals <- (0.8*nIndividuals)
         test_nIndividuals <-  (0.2*nIndividuals)
	     
     
	 predBL_BLR <-BLR(y=trainPheno,XL=trainGeno,GF=NULL,
           prior=list(varE=list(df=3,S=0.25),
		   varU=list(df=3,S=0.63),
		   lambda=list(shape=0.53,rate=5e-5,
		   type='random',value=30)),
		   nIter=20000,burnIn=2000,thin=1)
		   
	
	  
	 	 
################################################################################3
	  
  estimated.Marker.Effects <- predBL_BLR$bL 
  
  pred.train <- trainGeno %*% predBL_BLR$bL 
  
  Pred.pheno.train <-  predBL_BLR$yHat
  
  # cor(Pred.pheno.train,trainPheno)
  
#################################################################################################################
  
  testPheno <- testPhenoTable 
  testGeno <- testGenoTable 
  
  
  
  
  pred.valid <- (testGeno %*% predBL_BLR$bL) 
  
  Pred.pheno.valid <- predBL_BLR$mu + (as.vector(pred.valid))
  
  Pred.Accuracy <- cor(Pred.pheno.valid,testPheno)
  
  train_MSE <- (sum((Pred.pheno.train-trainPheno))^2)/(train_nIndividuals)
  test_MSE <- (sum((Pred.pheno.valid-testPheno))^2)/(test_nIndividuals) 
  
  sol <- list(Pred.Accuracy,predBL_BLR,estimated.Marker.Effects,train_MSE,test_MSE)
  
  return(sol)
  
} 

 
```
