## Supplementary material for "Factors Affecting Response to Recurrent Genomic Selection in Soybeans": File S1: functionBRRModel.R

```
### function Bayesian Ridge regression 

buildBRRModel <- function(trainGenoTable,trainPhenoTable,testGenoTable,testPhenoTable,heritability){ 

	
	 trainPheno <- trainPhenoTable[,1]
         trainGeno <- trainGenoTable
	 h2 <- heritability
	 nIndividuals <- 2000
	 
################################################################################3
	
  predBL_BRR <- BLR(y=trainPheno,XR=trainGeno,GF=NULL,
           prior=list(varE=list(df=4,S=1),varU=list(df=5,S=0.01)),
		   nIter=20000,burnIn=2000,thin=1)  
		   


	
  estimated.Marker.Effects <- predBL_BRR$bR 
  
  pred.train <- trainGeno %*% predBL_BRR$bR 
  
  Pred.pheno.train <-  predBL_BRR$mu + predBL_BRR$bR
  
#################################################################################################################
  
  testPheno <- testPhenoTable[,1] 
  testGeno <- testGenoTable 
  
  train_nIndividuals <- (0.8*nIndividuals)
  test_nIndividuals <-  (0.2*nIndividuals)
  
  
  pred.valid <- (testGeno %*% predBL_BRR$bR) 
  
  Pred.pheno.valid <- predBL_BRR$mu + (as.vector(pred.valid))
  
  Pred.Accuracy <- cor(Pred.pheno.valid,testPheno)
  
  train_MSE <- (sum((Pred.pheno.train-trainPheno))^2)/(train_nIndividuals)
  test_MSE <- (sum((Pred.pheno.valid-testPheno))^2)/(test_nIndividuals) 
  
  sol <- list(Pred.Accuracy,predBL_BRR,estimated.Marker.Effects,train_MSE,test_MSE)
  
  return(sol)


}
```
