## Supplementary material for "Factors Affecting Response to Recurrent Genomic Selection in Soybeans": File S1: functionElasticNetModel.R

```
buildElasticNetModel <- function(trainGenoTable,trainPhenoTable,testGenoTable,testPhenoTable,heritability){ 

	 library(glmnet)
	 trainPheno <- trainPhenoTable[,1]
         trainGeno <- trainGenoTable
	 testPheno <- testPhenoTable[,1]
         testGeno <- testGenoTable
         h2 <- heritability
	 nIndividuals <- 2000
	 
################################################################################3
   
 
 pred_ENet=glmnet(y=trainPheno,x=trainGeno,alpha=0.2)

 Cor.ENet <- rep(0,100)
 # evaluating correlation in TST set
 for(i in 1:100){
   
   Cor.ENet[i]=cor(testPheno,testGeno%*%pred_ENet$beta[,i])
 }
 
  
  selindex <- which.max(Cor.ENet)
	
  estimated.Marker.Effects <-  pred_ENet$beta[,selindex]
  
  pred.train <- trainGeno %*%  pred_ENet$beta[,selindex]
  
  Pred.pheno.train <-  pred_ENet$a0[selindex] + as.vector(pred.train)
  
###########################################################################################################
  
  
  
  train_nIndividuals <- (0.8*nIndividuals)
  test_nIndividuals <-  (0.2*nIndividuals)
  
  
  pred.valid <- (testGeno %*% pred_ENet$beta[,selindex]) 
  
  Pred.pheno.valid <- pred_ENet$a0[selindex] + (as.vector(pred.valid))
  
  Pred.Accuracy <- Cor.ENet[selindex]
  
  train_MSE <- (sum((Pred.pheno.train-trainPheno))^2)/(train_nIndividuals)
  test_MSE <- (sum((Pred.pheno.valid-testPheno))^2)/(test_nIndividuals) 
  
  sol <- list(Pred.Accuracy,pred_ENet,estimated.Marker.Effects,train_MSE,test_MSE)
  
  return(sol)


}
```
