## Supplementary material for "Factors Affecting Response to Recurrent Genomic Selection in Soybeans": File S1: functionLASSOModel.R

```
 
 

buildLASSOModel <- function(trainGenoTable,trainPhenoTable,testGenoTable,testPhenoTable,heritability){ 

	 library(glmnet)
	 trainPheno <- trainPhenoTable[,1]
         trainGeno <- trainGenoTable
	 testPheno <- testPhenoTable[,1] 
         testGeno <- testGenoTable 
	 h2 <- heritability
	 nIndividuals <- 2000
	 
################################################################################3
	
         pred_LASSO =glmnet(y=trainPheno,x=trainGeno, alpha=1)	  
 
 

 # evaluating correlation in TST set
	 Cor.LASSO <- rep(0,100)
	 for(i in 1:100){
	   
	   Cor.LASSO[i]=cor(testPheno,testGeno%*%pred_LASSO$beta[,i])
	   
	 }
 

  selindex <- which.max(Cor.LASSO)
	
  estimated.Marker.Effects <-  pred_LASSO$beta[,selindex]
  
  pred.train <- trainGeno %*%  pred_LASSO$beta[,selindex]
  
  Pred.pheno.train <-  pred_LASSO$a0[selindex] + as.vector(pred.train)
  
###########################################################################################################
  
  
  
  train_nIndividuals <- (0.8*nIndividuals)
  test_nIndividuals <-  (0.2*nIndividuals)
  
  
  pred.valid <- (testGeno %*% pred_LASSO$beta[,selindex]) 
  
  Pred.pheno.valid <- pred_LASSO$a0[selindex] + (as.vector(pred.valid))
  
  Pred.Accuracy <- Cor.LASSO[selindex]
  
  train_MSE <- (sum((Pred.pheno.train-trainPheno))^2)/(train_nIndividuals)
  test_MSE <- (sum((Pred.pheno.valid-testPheno))^2)/(test_nIndividuals) 
  
  sol <- list(Pred.Accuracy,pred_LASSO,estimated.Marker.Effects,train_MSE,test_MSE)
  
  return(sol)


}


  # trainPheno <- trainSimPhenoTableListReps[[1]][[1]][,1]
  # trainGeno <- trainGenoNewTableListReps[[1]][[1]]
  # testPheno <- testSimPhenoTableListReps[[1]][[1]][,1]
  # testGeno <- testGenoNewTableListReps[[1]][[1]]

  # h2 <- 0.7
  # nIndividuals <- 2000
	
  # pred_LASSO =glmnet(y=trainPheno,x=trainGeno, alpha=1)	
  
  # heritability <- h2
  

# PredictionModel_Pheno <-(buildLASSOModel(trainGenoNewTableListReps[[1]][[1]],trainSimPhenoTableListReps[[1]][[1]],testGenoNewTableListReps[[1]][[1]],testSimPhenoTableListReps[[1]][[1]],heritability))
```
