## Supplementary material for "Factors Affecting Response to Recurrent Genomic Selection in Soybeans": File S1: functionRRBLUP_EM_Model.R

```
#### Function to build RRBLUP model
#### rrBLUP to build genomic prediction model
### rrReml(NumericVector y, NumericMatrix gen, int maxit = 500, double tol = 10e-6)


buildRRBLUPModel_REML <- function(trainGenoTable,trainPhenoTable,testGenoTable,testPhenoTable){
  
  options(warn=-1) 
  
  nIndividuals_train <- nrow(trainGenoTable)
  nIndividuals_test <- nrow(testGenoTable)

  trainPheno <- trainPhenoTable
  trainGeno <- trainGenoTable
 
 
  pred <-  (rrReml(trainPheno,trainGeno,500,10e-6))
  
  estimated.Marker.Effects <- pred$b 
  
  pred.train <- trainGeno %*% pred$b
  
  Pred.pheno.train <- pred.train[,1] + (pred$mu)
  
  
#################################################################################################################
  
  testPheno <- testPhenoTable
  testGeno <- testGenoTable 
  
  
  pred.valid <- (testGeno %*% pred$b) 
  
  Pred.pheno.valid <- pred.valid[,1] + (pred$mu) 
  
  Pred.Accuracy <- suppressWarnings(cor(Pred.pheno.valid,testPheno))
  
  train_MSE <- (sum((Pred.pheno.train-trainPheno))^2)/(nIndividuals_train)
  test_MSE <- (sum((Pred.pheno.valid-testPheno))^2)/(nIndividuals_test) 
  
  sol <- list(estimated.Marker.Effects,pred$mu,Pred.Accuracy,train_MSE,test_MSE)
  
  return(sol)
  
} 

################################################################
## Test PredictionModel _Geno
# PredictionModel_Geno <- buildRRBLUPModel_REML(trainGenoNewTable,trainSimPhenoTable,testGenoNewTable,testSimPhenoTable)


 # trainGeno <- trainGenoNewTable
  # trainPheno <- trainSimPhenoTable[,1]
  # testGeno <- testGenoNewTable 
  # testPheno <- testSimPhenoTable[,1]
  
```
