## Supplementary material for "Factors Affecting Response to Recurrent Genomic Selection in Soybeans": File S1: functionRRBLUPModel.R

```
#### Function to build RRBLUP model
#### rrBLUP to build genomic prediction model

buildRRBLUPModel <- function(trainGenoTable,trainPhenoTable,testGenoTable,testPhenoTable){
  
 
  nIndividuals_train <- nrow(trainGenoTable)
  nIndividuals_test <- nrow(testGenoTable)

  trainPheno <- trainPhenoTable
  trainGeno <- trainGenoTable
  
  pred <- mixed.solve(trainPheno,Z=trainGeno,K=NULL,SE=FALSE,return.Hinv =FALSE) 
  
  estimated.Marker.Effects <- pred$u 
  
  pred.train <- trainGeno %*% pred$u
  
  Pred.pheno.train <- pred.train[,1] + (pred$beta)
  
  
  #cor(Pred.pheno.train,trainPheno)
  
#################################################################################################################
  
  testPheno <- testPhenoTable
  testGeno <- testGenoTable 
  
  
  pred.valid <- (testGeno %*% pred$u) 
  
  Pred.pheno.valid <- pred.valid[,1] + (pred$beta) 
  
  Pred.Accuracy <- cor(Pred.pheno.valid,testPheno)
  
  train_MSE <- (sum((Pred.pheno.train-trainPheno))^2)/(nIndividuals_train)
  test_MSE <- (sum((Pred.pheno.valid-testPheno))^2)/(nIndividuals_test) 
  
  sol <- list(estimated.Marker.Effects,pred$beta,Pred.Accuracy,train_MSE,test_MSE)
  
  return(sol)
  
} 

################################################################
```
