## Supplementary material for "Factors Affecting Response to Recurrent Genomic Selection in Soybeans": File S1: functionRunSimulations_WGSV4.R

```
## Module 4: Functions to run the simulation
## Apply selection to choose parents for the next generation ##

### Extract Genotype data in long array format ########################################################
### Meant for geno data with 20 crosses and 100 progeny wih 2 columns of markers and 4000 loci

# RF_Vector <- as.vector(NAM_LinkMap_New[,3])
# nMarkers <- length(RF_Vector)

## Fn 1

extractGenoData <- function(GenoTableIndices1,ProgenyGenoData,no_selected){

  GenoTableIndices <- GenoTableIndices1
  nFamilies <- dim(ProgenyGenoData)[1]
  nMarkers <- 4289

  if(nFamilies == 20){

	n<- length(GenoTableIndices)

	genoData <- array(0,c(n,nMarkers,2))

	for(j in 1:n){

	 if(GenoTableIndices[j]<100){
		i<-1
		}

		if(GenoTableIndices[j]>=100 && GenoTableIndices[j]!=5000 ){
			i <- (GenoTableIndices[j] %/% 100)+1
		}

		if(GenoTableIndices[j]>=100 && GenoTableIndices[j]!=2000 ){
			i <- (GenoTableIndices[j] %/% 100)+1
		}


		else if(GenoTableIndices[j]==5000 ){
			i<-50
			l<-100
		}

		else if(GenoTableIndices[j]==2000 ){
			i<-20
			l<-100
		}

		l <- GenoTableIndices[j] %% 100

		if(l==0){

			l=100
		}

		genoData[j,,] <- ProgenyGenoData[i,,,l]
    }
  }

  if(nFamilies ==50){

	n<- length(GenoTableIndices)

	genoData <- array(0,c(n,nMarkers,2))

    for(j in 1:n){

		if(GenoTableIndices[j]<40){
			i<-1
		}

		if(GenoTableIndices[j]>=40 && GenoTableIndices[j]!= 5000 ){
			i <- (GenoTableIndices[j] %/% 40)+1
		}

		if(GenoTableIndices[j]>=40 && GenoTableIndices[j]!= 2000 ){
			i <- (GenoTableIndices[j] %/% 40)+1
		}


		# else if(GenoTableIndices[j]==5000 ){
			# i<-50
			# l<-40
		# }

		else if(GenoTableIndices[j]==2000 ){
			i<-50
			l<-40
		}

		l <- GenoTableIndices[j] %% 40

		if(l==0){

			l=40
		}

		genoData[j,,] <- ProgenyGenoData[i,,,l]

	}
  }

  if(nFamilies == 200){

	n<- length(GenoTableIndices)

	genoData <- array(0,c(n,nMarkers,2))

    for(j in 1:n){

		if(GenoTableIndices[j]< 10){
			i<-1
		}

		if(GenoTableIndices[j]>=10 && GenoTableIndices[j]!= 5000 ){
			i <- (GenoTableIndices[j] %/% 10)+1
		}

		if(GenoTableIndices[j]>=10 && GenoTableIndices[j]!= 2000 ){
			i <- (GenoTableIndices[j] %/% 10)+1
		}


		# else if(GenoTableIndices[j]==5000 ){
			# i<-50
			# l<-40
		# }

		else if(GenoTableIndices[j]==2000 ){
			i<-200
			l<-10
		}

		l <- GenoTableIndices[j] %% 10

		if(l==0){

			l=10
		}

		genoData[j,,] <- ProgenyGenoData[i,,,l]

	}
  }


  if(nFamilies == 400){

	n<- length(GenoTableIndices)

	genoData <- array(0,c(n,nMarkers,2))

    for(j in 1:n){

		if(GenoTableIndices[j]< 5){
			i<-1
		}

		if(GenoTableIndices[j]>=5 && GenoTableIndices[j]!= 5000 ){
			i <- (GenoTableIndices[j] %/% 10)+1
		}

		if(GenoTableIndices[j]>=5 && GenoTableIndices[j]!= 2000 ){
			i <- (GenoTableIndices[j] %/% 5)+1
		}


		# else if(GenoTableIndices[j]==5000 ){
			# i<-50
			# l<-40
		# }

		else if(GenoTableIndices[j]==2000 ){
			i<-400
			l<-5
		}

		l <- GenoTableIndices[j] %% 5

		if(l==0){

			l=5
		}

		genoData[j,,] <- ProgenyGenoData[i,,,l]

	}
  }

 	return(genoData)
}

## Function Test #######################################################################
# gen1_top20_genoData <- extractGenoData(GenoTableIndices_top20,F5_Progeny)
# gen1_top50_genoData <- extractGenoData(selectedGenoIndividualIndices,Cycle_Progeny)

########### Generate Doubled Haploid ####################################################

## Fn 2

generateDoubledHaploid <- function(selectedGenoData){

  no_of_lines <- dim(selectedGenoData)[1]
  nMarkers <- dim(selectedGenoData)[2]
  nChr <- dim(selectedGenoData)[3]

  DH_Lines<- array(0,c(2*no_of_lines,nMarkers,nChr))

  j <-1

  for( i in 1: no_of_lines){

    DH_Lines[j,,] <- cbind(selectedGenoData[i,,1],selectedGenoData[i,,1])
    DH_Lines[j+1,,]<-  cbind(selectedGenoData[i,,2],selectedGenoData[i,,2])
    j<- j+2

  }

  return(DH_Lines)
}


### test generateDoubledHaploid
# DH_test <- generateDoubledHaploid(gen1_top20_genoData)

## Fn 3
########### Generate Doubled Haploid for (2a) ####################################################################


generateDoubledHaploid_2a <- function(selectedGenoData){

  no_of_lines <- dim(selectedGenoData)[1]
  nMarkers <- dim(selectedGenoData)[2]
  nChr <- dim(selectedGenoData)[3]
  nProgeny <- dim(selectedGenoData)[4]

  DH_Lines<- array(0,c(no_of_lines,nMarkers,nChr,nProgeny,2))
  DH_Lines_New <- array(0,c(no_of_lines,nMarkers,nChr,nProgeny))


  for( i in 1: no_of_lines){
    for( k in 1: nProgeny){

      index <-sample(c(1,2),1)

      DH_Lines_New[i,,,k] <- cbind(selectedGenoData[i,,index,k],selectedGenoData[i,,index,k])
      #DH_Lines[j,,,k,2] <-  cbind(selectedGenoData[i,,2,k],selectedGenoData[i,,2,k])


      }
  }

     return(DH_Lines_New)
  }

##  test generateDoubledHaploid_2a

#DH_test <- generateDoubledHaploid_2a(Cycle_Progeny)


#################################################################################################################

### Function to generate progeny given two parents and the number of Progeny
### generateProgeny(Parent1,Parent2,Number_of_progeny)

## Fn4

#generateProgeny <- function(Parent1,Parent2,Number_Progeny){

#  RF_Vector <- as.vector(NAM_LinkMap_New[,3])
#  rr <- as.vector(NAM_LinkMap_New[,3])


#  x1<-Parent1
#  x2<- Parent2


#  a<-Number_Progeny
#  n <- dim(x1)[1]

#  y<-array(0,dim=c(n,2,a))


#  for (i in 1:a) {

#    p1<-cumsum(runif(n)<=rr)%%2
#    p2<-cumsum(runif(n)<=rr)%%2 #creating gamete from p2


#    y[p1==0,1,i]=x1[p1==0,1]

#    y[p1==1,1,i] = x1[p1==1,2]

#    y[p2==0,2,i] = x2[p2==0,1]

#    y[p2==1,2,i] = x2[p2==1,2]
#  }

#  progeny <- y

#  return(progeny)

#}


## Fn5
######### Function to generate GenoTable in 012 format ########################

generate012GenoFormat <- function(Cycle_Progeny_Data,number_of_Crosses,n_Progeny){

  Cycle_Progeny <- Cycle_Progeny_Data


  nCrosses <- number_of_Crosses
  nProgeny<- n_Progeny
  nMarkers <- 4289


  Cycle_Progeny_table <-c()
  cycle_family_table<- array(0,c(1,nMarkers,nProgeny))

  for(i in 1:nCrosses){

    for(k in 1:nProgeny){


      cycle_family_table[1,,k] <- (Cycle_Progeny[i,,1,k] + Cycle_Progeny[i,,2,k])

    }

    Cycle_Progeny_table <-cbind(Cycle_Progeny_table,cycle_family_table[1,,1:nProgeny])

  }

  IndividualNames <-rep(0,(nCrosses*nProgeny))

  for(i in 1:(nCrosses*nProgeny)){
    IndividualNames[i]<- paste("Ind",i,sep="")
  }

# IndividualNames <- c("Animal_ID",IndividualNames)
#### RowNames

  markerNames <-rep(0,nMarkers)


  for(i in 1:(nMarkers)){
    markerNames[i]<- paste("m",i,sep="")
  }


  colnames(Cycle_Progeny_table)<- IndividualNames
  row.names(Cycle_Progeny_table)<- markerNames

  genoTable<- t(Cycle_Progeny_table)


  return(genoTable)
}

## Test Function generate012GenoFormat
#Cycle_test <- generate012GenoFormat(Cycle_Progeny,50)


## Fn6
###############################################################################################################

generateMinus101GenoFormat <- function(Cycle_Progeny_Data,number_of_Crosses,n_Progeny){

  Cycle_Progeny <- Cycle_Progeny_Data

  nCrosses <- number_of_Crosses
  nProgeny <- n_Progeny
  nMarkers <- 4289

  Cycle_Progeny_table <-c()
  cycle_family_table<- array(0,c(1,nMarkers,nProgeny))

  for(i in 1:nCrosses){

      cycle_family_table[1,,] <- ((Cycle_Progeny[i,,1,] + Cycle_Progeny[i,,2,])) -1

      Cycle_Progeny_table <-cbind(Cycle_Progeny_table,cycle_family_table[1,,1:nProgeny])

  }

  IndividualNames <-rep(0,(nCrosses*nProgeny))
    
  IndividualNames <- paste("Ind",c(1:(nCrosses*nProgeny)),sep="")
 
  #### RowNames

  markerNames <-rep(0,nMarkers)
 
  markerNames <- paste("m",c(1:nMarkers),sep="")
 
  colnames(Cycle_Progeny_table)<- IndividualNames
  row.names(Cycle_Progeny_table)<- markerNames

  genoTable<- t(Cycle_Progeny_table)

  ########## Translate 0-1-2 code to -1-0-1 code in genotable

  # genoTable_Mod <- apply(genoTable,2,function(x)(gsub("\\b0\\b","minus1",as.matrix(x))))
  # genoTable_Mod <- apply(genoTable_Mod,2,function(x)(gsub("\\b1\\b","0",as.matrix(x))))
  # genoTable_Mod <- apply(genoTable_Mod,2,function(x)(gsub("\\b2\\b","1",as.matrix(x))))
  # genoTable_Mod <- apply(genoTable_Mod,2,function(x)(gsub("\\bminus1\\b","-1",as.matrix(x))))
  
  genoTable_Mod <- apply(genoTable,2,as.numeric)


  return(genoTable_Mod)
}

## Test Function generate012GenoFormat

# Cycle_test <- generateMinus101GenoFormat(F5_Progeny,20)

##################################################################################################

## Fn 9
####### Function 8 get Frequency and state of favorable allele

getFreq_BasePoln<- function(genoTable,MarkerEffects){

    GenoTable <- genoTable
	markerEffects <- MarkerEffects

   	nMarkers <- dim(GenoTable)[2]
	nIndividuals <- dim(GenoTable)[1]
	sign_markerEffects <- sign(markerEffects)
	n_alleles <- 2*nIndividuals

	Freq <- rep(0,nMarkers)
	FavAllele <- rep(0,nMarkers)
	count_1 <- rep(0,nMarkers)
	count_0 <- rep(0,nMarkers)

	for(j in 1:nMarkers){


		p_1_P <- length(which(GenoTable[,j]==1))
		p_0_H <- length(which(GenoTable[,j]==0))
		p_minus1_Q <- length(which(GenoTable[,j]==-1))


		# p_1_P <- length(which(GenoVector==1))/nIndividuals
		# p_0_H <- length(which(GenoVector==0))/nIndividuals
		# p_minus1_Q <- length(which(GenoVector==-1))/nIndividuals

		count_1[j] <- ((2*p_1_P)+p_0_H )
		count_0[j] <- ((2*p_minus1_Q)+p_0_H )

		if((count_1[j] >count_0[j]) && sign_markerEffects[j]==1){
			FavAllele[j] <- 1
			Freq[j] <- count_1[j]/n_alleles
		}else if((count_0[j] > count_1[j]) && sign_markerEffects[j]==1){
			FavAllele[j] <- 0
			Freq[j] <- count_0[j]/n_alleles

		}


	}

	return(list(Freq,FavAllele))

}


# ParentLines - QTL assigned alternating positive and negative allele effects, eg: 400 (
# Favorable allele at assigned QTL loci 
# i)  '1' genotypic state - with positive allele effect
# ii) ,whereas '-1' genotypic state with negative allele effect)

# FavQTL allele using assigned effect values:  

# From parental lines, base population of 2000 F5 RILs are generated. Then genotypic and phenotypic values are simulated from assigned QTL allele effects. 

# Marker effects are estimated using RRREML method with QTL alleles retained. 

# Sign of estimated marker effects 


# GenoTable <- genoTable_Mod
# markerEffects <- TrainModel
#########################################################################################
## Fn 10: get frequency of favorable allele given genotype table and state of favorable allele

getFreq <- function(GenoTable,FavAllele){


	nMarkers <- dim(GenoTable)[2]
	nIndividuals <- dim(GenoTable)[1]
	Freq <- rep(0,nMarkers)
	count_1 <-rep(0,nMarkers)
	count_0 <-rep(0,nMarkers)
	n_alleles <- 2*nIndividuals

	for(j in 1:nMarkers){


		p_1_P <- length(which(GenoTable[,j]==1))
		p_0_H <- length(which(GenoTable[,j]==0))
		p_minus1_Q <- length(which(GenoTable[,j]==-1))


		count_1[j] <- ((2*p_1_P)+p_0_H )
		count_0[j] <- ((2*p_minus1_Q)+p_0_H )

		if(FavAllele[j]==1){
			Freq[j] <- count_1[j]/n_alleles
		}else if(FavAllele[j]==0){
			Freq[j] <- count_0[j]/n_alleles
		}
	}

	return(Freq)

}


## Fn 7
##### Function to Predict Phenotype Values given genotable in 012 format

PredictRRBLUPPhenoValues <- function(GenoTable,TrainModel){

  givenGenoTable<- GenoTable

  nMarkers <- ncol(givenGenoTable)
  markerNames <-rep(0,nMarkers)
  
 
  markerNames <- paste("m",c(1:(nMarkers)),sep="")
  

  beta <- unlist(TrainModel[1])
  names(beta)<- markerNames


  y.hat <- givenGenoTable %*% beta
  Avg <- unlist(TrainModel[2])

  pred.y <- y.hat+ Avg

  return(pred.y)

}

# testRRBLUPPredict<- PredictRRBLUPPhenoValues(genoTable_New2,model)

## Fn 8
#####  Function to predict phenotype values using Bayesian GS models given genotable in 012 format

PredictBayesPhenoValues <- function(GenoTable,TrainModel){

  givenGenoTable <- GenoTable

  # Pred.pheno.valid <- (testGeno %*% predBayesA$ETA[[1]]$b)
  # Pred.Accuracy <- cor(Pred.pheno.valid,testPheno)


  # TrainModel in sol <- list(Pred.Accuracy,predBayesA,predBayesA$ETA[[1]]$b,train_MSE,test_MSE)

  beta.markers <- as.vector(TrainModel[[3]])
  # names(beta)<- markerNames


  y.hat <- givenGenoTable %*% beta.markers

  pred.y <- y.hat
  return(pred.y)

}

# predictBayesPhenoValues(testGeno,sol)

## Fn 9:
##### Function to predict phenotype values using Bayesian GS models with marker effects weighted with ##### Jannink's function given genotable in 012 format

PredictBayesPhenoValues_Wght <- function(GenoTable,TrainModel,FreqVec){

  givenGenoTable <- GenoTable
  Freq <- FreqVec
  beta.markers <- as.vector(TrainModel[[3]])


  weightsVec <- (1/sqrt(Freq))
  wt.beta <- beta.markers* weightsVec

  Neginf.indices<- which(wt.beta == -Inf)
  inf.indices <-  which(wt.beta == Inf)
  indices.unWtd <- c(Neginf.indices,inf.indices)

  if(length(indices.unWtd)!=0) {
	wt.beta[indices.unWtd] <- beta.markers[indices.unWtd]
  }

  y.hat <- givenGenoTable %*% wt.beta

  pred.y <- y.hat
  return(pred.y)

}

## Fn 10: 
##### Function to predict phenotype values using Bayesian GS models with marker effects weighted with ##### dynamic weighting function given genotable in 012 format


PredictBayesPhenoValues_WGS_DWModel<- function(GenoTable,TrainModel,FreqVec,alphaShape,betaShape,cycleNumber,timeHorizon){

	  givenGenoTable<- GenoTable
	  beta.markers <- as.vector(TrainModel[[3]])
	  Freq <- FreqVec

	  alphaS <- alphaShape
	  betaS <- betaShape
	  N <- timeHorizon
	  t <- cycleNumber


	  alphaPar <- (alphaS+ (t*(1-alphaS)/N))
	  betaPar <- betaS

	  weightsVec <- (Freq^(alphaS+ (t*((1-alphaS)/N))))/ beta(alphaPar,betaPar)

	  wt.beta <- beta.markers* weightsVec

	  Neginf.indices<- which(wt.beta == -Inf)
	  inf.indices <-  which(wt.beta == Inf)
	  indices.unWtd <- c(Neginf.indices,inf.indices)

	  if(length(indices.unWtd)!=0) {
		wt.beta[indices.unWtd] <- beta.markers[indices.unWtd]
	  }

	  y.hat <- givenGenoTable %*% wt.beta

	  # Avg <- unlist(TrainModel[2])

	  # Avg <- 0
	  pred.y <- y.hat

	  return(list(pred.y,alphaPar))

}


## Fn 11:
##### Function to predict phenotype values using RidgeRegression GS models with marker effects weighted with Jannink's weighting function given genotable in 012 format

PredictRRBLUPPhenoValues_WGS_Jannink1 <- function(GenoTable,TrainModel,FreqVec){

  givenGenoTable<- GenoTable
  beta.markers <- unlist(TrainModel[1])
  # names(beta)<- markerNames
  Freq <- FreqVec

  weightsVec <- (1/sqrt(Freq))

  wt.beta <- beta.markers* weightsVec

  Neginf.indices<- which(wt.beta == -Inf)
  inf.indices <-  which(wt.beta == Inf)
  indices.unWtd <- c(Neginf.indices,inf.indices)

  if(length(indices.unWtd)!=0) {
	wt.beta[indices.unWtd] <- beta.markers[indices.unWtd]
  }

  y.hat <- givenGenoTable %*% wt.beta

  Avg <- unlist(TrainModel[2])

  pred.y <- y.hat+ Avg

  return(pred.y)

}


## Fn 12:
##### Function to predict phenotype values using RidgeRegression GS models with marker effects weighted with dynamic weighting function given genotable in 012 format

PredictRRBLUPPhenoValues_WGS_DWModel<- function(GenoTable,TrainModel,FreqVec,alphaShape,betaShape,cycleNumber,timeHorizon){

	  givenGenoTable<- GenoTable
	  beta.markers <- unlist(TrainModel[1])
	  Freq <- FreqVec

	  alphaS <- alphaShape
	  betaS <- betaShape
	  N <- timeHorizon
	  t <- cycleNumber


	  alphaPar <- (alphaS+ (t*(1-alphaS)/N))
	  betaPar <- betaS

	  weightsVec <- (Freq^(alphaS+ (t*((1-alphaS)/N))))/ beta(alphaPar,betaPar)

	  wt.beta <- beta.markers* weightsVec

	  Neginf.indices<- which(wt.beta == -Inf)
	  inf.indices <-  which(wt.beta == Inf)
	  indices.unWtd <- c(Neginf.indices,inf.indices)

	  if(length(indices.unWtd)!=0) {
		wt.beta[indices.unWtd] <- beta.markers[indices.unWtd]
	  }

	  y.hat <- givenGenoTable %*% wt.beta

	  Avg <- unlist(TrainModel[2])

	  pred.y <- y.hat+ Avg

	  return(list(pred.y,alphaPar))

}


## Fn 13:
##########################################################################
#### Function to select parents for next gen

ApplySelection <- function(PhenoValues,no_selected){


   nSel<-no_selected

   phenoValues<- PhenoValues
   sortedPhenoValues <- sort(phenoValues,decreasing=TRUE,index.return=TRUE)

   topNPhenoValues<- sortedPhenoValues[[1]][1:nSel]
   GenoTableIndices_topN <-  sortedPhenoValues[[2]][1:nSel]

   selected_PG_table<- cbind(topNPhenoValues,GenoTableIndices_topN)

   return(selected_PG_table)

 }

# testSelection <- ApplySelection(genoValues,20)

######################################################################################################
######################################################################################################

## Fn 14 a:
#### Function to simulate genotypic values give Genotype table from each cycle of crossing
######################################################################################################

simulateGenotypicValues <- function(nextGenGenoTable,noLoci) {

    genoTable <- nextGenGenoTable
    nIndividuals <-nrow(genoTable)
    genotypicValuesSim <-rep(0,nIndividuals)
    nLoci <- noLoci
    nMarkers <- ncol(genoTable)
    ##### Extract QTL table ######

    if(nLoci== 40){
      QTL_indices<- seq(50,(nMarkers/2),by=53)
    }


    if(nLoci == 400){
      QTL_indices_1<- seq(8,nMarkers,by=12)
      QTL_indices_2 <- seq(50,nMarkers,by=107)
      n_3 <- 400-(length(QTL_indices_1))-(length(QTL_indices_2))
      QTL_indices_3<- sample(c(1:nMarkers),n_3)

      QTL_indices <- c(QTL_indices_1,QTL_indices_2,QTL_indices_3)
    }


    if(nLoci == nMarkers){
      QTL_indices<- c(1:nMarkers)
    }


    QTL_table <- genoTable[,c(QTL_indices)]

############### Simulate Genotypic Values ######################

    if(nLoci == 40){

		evenIndices<- seq(2,40,by=2)
		oddIndices <- seq(1,39,by=2)

		for(i in 1:nIndividuals){

			genotypicValuesSim[i] <- sum(QTL_table[i,evenIndices]*5)+sum(QTL_table[i,oddIndices]*-5)

		}
}

	if(nLoci == 400){
		evenIndices<- seq(2,400,by=2)
		oddIndices <- seq(1,399,by=2)

		for(i in 1:nIndividuals){

			genotypicValuesSim[i] <- sum(QTL_table[i,evenIndices]*0.5)+sum(QTL_table[i,oddIndices]*-0.5)

		}
}

	if(nLoci == nMarkers){

		if(nMarkers %% 2 ==1){
			oddIndices <- seq(1,nMarkers,by=2)
			evenIndices<- seq(2,(nMarkers-1),by=2)
		}

		else if(nMarkers %% 2 ==0){
			oddIndices <- seq(1,(nMarkers-1),by=2)
			evenIndices<- seq(2,nMarkers,by=2)
		}

		for(i in 1:nIndividuals){

			genotypicValuesSim[i] <- sum(QTL_table[i,evenIndices]*0.05)+sum(QTL_table[i,oddIndices]*-0.05)

		}
    }

    return(genotypicValuesSim)
}

## Fn14 b:
###########################################################################################
######### Function to simulate Phenotypic Values ##########################################

simulatePhenotypicValues <- function(nextGenGenoTable,varE,noLoci){


    genoTable <- nextGenGenoTable
    nIndividuals <- nrow(genoTable)
    genotypicValuesSim <-rep(0,nIndividuals)
    phenotypicValuesSim <-rep(0,nIndividuals)
    nMarkers <- ncol(genoTable)

    errorVar <- varE
    errorSD <- sqrt(errorVar)
    nLoci <- noLoci
#######################################################################
##### Extract QTL table ###### ###### ###### ###### ###### ###### ######

    if(nLoci== 40){
      QTL_indices<- seq(50,(nMarkers/2),by=53)
    }


    if(nLoci == 400){
      QTL_indices_1<- seq(8,nMarkers,by=12)
      QTL_indices_2 <- seq(50,nMarkers,by=107)
      n_3 <- 400-(length(QTL_indices_1))-(length(QTL_indices_2))
      QTL_indices_3<- sample(c(1:nMarkers),n_3)

      QTL_indices <- c(QTL_indices_1,QTL_indices_2,QTL_indices_3)
    }


    if(nLoci == nMarkers){
      QTL_indices<- c(1:nMarkers)
    }


    QTL_table <- genoTable[,c(QTL_indices)]

############### Simulate Genotypic Values #############################

    if(nLoci == 40){

      evenIndices<- seq(2,40,by=2)
      oddIndices <- seq(1,39,by=2)


      for(i in 1:nIndividuals){

        genotypicValuesSim[i] <- sum(QTL_table[i,evenIndices]*5)+sum(QTL_table[i,oddIndices]*-5)
        phenotypicValuesSim[i] <- genotypicValuesSim[i] + rnorm(1,mean=0,sd=errorSD)
      }
    }

    if(nLoci == 400){

      evenIndices<- seq(2,400,by=2)
      oddIndices <- seq(1,399,by=2)

      for(i in 1:nIndividuals){

        genotypicValuesSim[i] <- sum(QTL_table[i,evenIndices]*0.5)+sum(QTL_table[i,oddIndices]*-0.5)
        phenotypicValuesSim[i] <- genotypicValuesSim[i] + rnorm(1,mean=0,sd=errorSD)

      }
    }

    if(nLoci == nMarkers){

      if(nMarkers %% 2 ==1){
        oddIndices <- seq(1,nMarkers,by=2)
        evenIndices<- seq(2,(nMarkers-1),by=2)
      }

      else if(nMarkers %% 2 ==0){
        oddIndices <- seq(1,(nMarkers-1),by=2)
        evenIndices<- seq(2,nMarkers,by=2)
      }

      for(i in 1:nIndividuals){

        genotypicValuesSim[i] <- sum(QTL_table[i,evenIndices]*0.05)+sum(QTL_table[i,oddIndices]*-0.05)
        phenotypicValuesSim[i] <- genotypicValuesSim[i] + rnorm(1,mean=0,sd=errorSD)
      }
    }

    return(phenotypicValuesSim)
}

### Test Simulated Phenotypic Values
## Fn15:
#########################################################################
#### Generate Weighted Genotypic Table ##################################

generateWeightedGenoTable <- function(nextGenGenoTable,noLoci) {

  genoTable <- nextGenGenoTable
  nLoci <- noLoci
  nMarkers <- ncol(genoTable)

  if(nLoci== 40){
      QTL_indices<- seq(50,(nMarkers/2),by=53)
    }


  if(nLoci == 400){
    QTL_indices_1<- seq(8,nMarkers,by=12)
    QTL_indices_2 <- seq(50,nMarkers,by=107)
    n_3 <- 400-(length(QTL_indices_1))-(length(QTL_indices_2))
    QTL_indices_3<- sample(c(1:nMarkers),n_3)

    QTL_indices <- c(QTL_indices_1,QTL_indices_2,QTL_indices_3)
  }


  if(nLoci == nMarkers){
    QTL_indices<- c(1:nMarkers)
  }


  QTL_table <- genoTable[,c(QTL_indices)]
  QTL_table_New <- QTL_table

############ Weighting QTL based on QTL effect  #########################################


  for(i in 1:nrow(genoTable)){


    if(nLoci==40) {
      evenIndices<- seq(2,40,by=2)
      oddIndices <- seq(1,39,by=2)

      QTL_table_New[i,evenIndices] <- QTL_table[i,evenIndices]*2
      QTL_table_New[i,oddIndices] <- QTL_table[i,oddIndices]*-2

    }

    if(nLoci==400) {
      evenIndices<- seq(2,400,by=2)
      oddIndices <- seq(1,399,by=2)

      QTL_table_New[i,evenIndices] <- QTL_table[i,evenIndices]*2
      QTL_table_New[i,oddIndices] <- QTL_table[i,oddIndices]*-2

    }


    if(nLoci==nMarkers){

      if(nMarkers %% 2 ==1){
        oddIndices <- seq(1,nMarkers,by=2)
        evenIndices<- seq(2,(nMarkers-1),by=2)
      }

      else if(nMarkers %% 2 ==0){
        oddIndices <- seq(1,(nMarkers-1),by=2)
        evenIndices<- seq(2,nMarkers,by=2)
      }

          QTL_table_New[i,evenIndices] <- QTL_table[i,evenIndices]*2
          QTL_table_New[i,oddIndices] <- QTL_table[i,oddIndices]*-2
    }
  }

  genoTable_New <- genoTable
  genoTable_New[,c(QTL_indices)]<- QTL_table_New


  return(genoTable_New)
}

# testweightGeno <- generateWeightedGenoTable(nextGenGenoTable,400)


### Function to estimate average percent heterozygous across all sampled loci...
### GenoTable in minus101 format ################################################################################
### Fn 16:

  getPercentHeterozygous<- function(nextgenoTable){

    genoTable_Mod <- nextgenoTable
    nMarkers <- ncol(genoTable_Mod)

    markerTest <- rep(0,nMarkers)
    no_genotype_1<- rep(0,nMarkers)
    no_genotype_0 <- rep(0,nMarkers)
    no_genotype_minus1 <- rep(0,nMarkers)
    percent_heterozygous_Mod <-rep(0,nMarkers)


    for(i in 1:nMarkers){

      markerTest[i]<- length(levels(factor(genoTable_Mod[,i])))
      no_genotype_1[i] <-length(genoTable_Mod[genoTable_Mod[,i]==1,i])
      no_genotype_0[i] <- length(genoTable_Mod[genoTable_Mod[,i]==0,i])
      no_genotype_minus1[i] <- length(genoTable_Mod[genoTable_Mod[,i]==-1,i])


      percent_heterozygous_Mod[i] <- no_genotype_0[i]/(no_genotype_1[i] + no_genotype_0[i] + no_genotype_minus1[i])

    }

    Avg_percent_heterozygous_Mod <- mean(percent_heterozygous_Mod)


 }

# testpercentHet <- getPercentHeterozygous(nextGenGenoTable)


## Fn 17:
getParentCombnIndices <- function(no_selected) {

	Parent_Combn_indices <- c()

	if(no_selected ==20) {

		combn_ParentVectorIndices <- c(2,3)
		for(l in 2:20){
			combn_ParentVectorIndices <- cbind(combn_ParentVectorIndices,c(1,l))
		}

		Parent_Combn_indices <- as.matrix(combn_ParentVectorIndices)


	}

	if(no_selected ==50) {

		combn_ParentVectorIndices <- c(2,3)

		for(l in 2:25){
			combn_ParentVectorIndices <- cbind(combn_ParentVectorIndices,c(1,l))
		}

		for(m in 25:50){
			combn_ParentVectorIndices <- cbind(combn_ParentVectorIndices,c(2,m))
		}

		Parent_Combn_indices <- as.matrix(combn_ParentVectorIndices)
	}


	if(no_selected ==100) {

		combn_ParentVectorIndices <- c(2,3)
		for(l in 2:20){
			combn_ParentVectorIndices <- cbind(combn_ParentVectorIndices,c(1,l))
		}

		Parent_Combn_indices <- as.matrix(combn_ParentVectorIndices)

	}

	if(no_selected ==200) {

		combn_ParentVectorIndices <- c(9,10)
		init  <- 2
		final  <- 25

		for(k in 1:8){

			for(m in init:final){

				combn_ParentVectorIndices <- cbind(combn_ParentVectorIndices,c(k,m))
			}
			init <- final+1
			final <- final +25
		}

		Parent_Combn_indices <- as.matrix(combn_ParentVectorIndices)

	}

	if(no_selected ==400) {

		combn_ParentVectorIndices <- c(9,10)
		init  <- 2
		final  <- 50

		for(k in 1:8){

			for(m in init:final){

				combn_ParentVectorIndices <- cbind(combn_ParentVectorIndices,c(k,m))
			}
			init <- final+1
			final <- final +50
		}

		Parent_Combn_indices <- as.matrix(combn_ParentVectorIndices)

	}

	return(Parent_Combn_indices)


}

##########################################################################################

## Fn 18:
generateProgenyCpp <- function(nMarkers,nProgeny,RF,Parent1,Parent2){

		library(Rcpp)
		sourceCpp('mei.cpp')
		sourceCpp('mei2.cpp')

		m<- nMarkers
		n<- 2*nProgeny
		rr <- RF
		a1<- Parent1[,1]
		b1<- Parent1[,2]

		Pro <- mei(m,n,rr,a1,b1)

		a2<- Parent2[,1]
		b2<- Parent2[,2]

	    Pro2<- mei2(m,n,rr,a2,b2,Pro)

		return(Pro2)
	}

## Fn 19:
	
### 

getF5RILs_BD <- function(BD,SelectedGenoData,nSelected,nProgeny,nMarkers,NAM_LinkMap){

	NAM_LinkMap_New <- NAM_LinkMap
    selectedGenoData <- SelectedGenoData
	no_selected <- nSelected 
	
    if(BD =="BD2_V2"){ 

		if(no_selected ==2){
			nCrosses <- 1
	    }else{nCrosses <- no_selected}


		Cycle_Progeny_F1 <- array(0,c(nCrosses,nMarkers,2,1))

		Cycle_Progeny_F2 <- array(0,c(nCrosses,nMarkers,2,nProgeny))

		Cycle_Progeny_F3 <- array(0,c(nCrosses,nMarkers,2,nProgeny))

		Cycle_Progeny_F4 <- array(0,c(nCrosses,nMarkers,2,nProgeny))

		Cycle_Progeny_F5 <- array(0,c(nCrosses,nMarkers,2,nProgeny))

###################################################################################

		for( j in 1:(nCrosses)){

## Get parents from selectedGenoData_List 
		
		  if(j ==1 && nCrosses ==1){
			  Parent1<-  selectedGenoData[j,,]
			  Parent2<-  selectedGenoData[j+1,,]
		  }else if ((nCrosses>1)&&(j<=(nCrosses%/%2))) {
		       Parent1<-  selectedGenoData[1,,]
			   Parent2<-  selectedGenoData[j+1,,]
		  }else if ((nCrosses >1)&&(j>(nCrosses %/%2))&&(j<=(nCrosses-1))) {
		       Parent1<-  selectedGenoData[2,,]
			   Parent2<-  selectedGenoData[(j-(nCrosses %/%2)+1),,]
		  }else if(j==nCrosses){
		       Parent1<-  selectedGenoData[j,,]
			   Parent2<-  selectedGenoData[1,,]
		  }
	
## Get F1- F5 RILs
	
		  Cycle_Progeny_F1[j,,,] <- generateProgeny(Parent1,Parent2,1,NAM_LinkMap_New)

		  

		  Parent1<- Cycle_Progeny_F1[j,,,1]
		  Parent2<- Cycle_Progeny_F1[j,,,1]
		  progeny1<- generateProgeny(Parent1,Parent2,nProgeny,NAM_LinkMap_New)
		  Cycle_Progeny_F2[j,,,]  <- progeny1

		 
		  for(m in 1:nProgeny){

			Parent1<- Cycle_Progeny_F2[j,,,m]
			Parent2<- Cycle_Progeny_F2[j,,,m]
			progeny1<- generateProgeny(Parent1,Parent2,1,NAM_LinkMap_New)
			Cycle_Progeny_F3[j,,,m] <- progeny1
		  }


		  for(m in 1:nProgeny){

			Parent1<- Cycle_Progeny_F3[j,,,m]
			Parent2<- Cycle_Progeny_F3[j,,,m]
			progeny1<- generateProgeny(Parent1,Parent2,1,NAM_LinkMap_New)
			Cycle_Progeny_F4[j,,,m] <- progeny1
		  }

		  for(m in 1:nProgeny){

			Parent1<- Cycle_Progeny_F4[j,,,m]
			Parent2<- Cycle_Progeny_F4[j,,,m]
			progeny1<- generateProgeny(Parent1,Parent2,1,NAM_LinkMap_New)
			Cycle_Progeny_F5[j,,,m] <- progeny1
		  }

		}

		
} 


  if (BD =="BD2"){ 
  
    if(no_selected ==2){
			nCrosses <- 1
	}else{nCrosses <- no_selected}

    Parent_Combn_indices <- getParentCombnIndices(no_selected)

    Cycle_Progeny_F1 <- array(0,c(no_selected,nMarkers,2,1))

    Cycle_Progeny_F2 <- array(0,c(no_selected,nMarkers,2,nProgeny))

    Cycle_Progeny_F3 <- array(0,c(no_selected,nMarkers,2,nProgeny))

    Cycle_Progeny_F4 <- array(0,c(no_selected,nMarkers,2,nProgeny))

    Cycle_Progeny_F5 <- array(0,c(no_selected,nMarkers,2,nProgeny))


###################################################################################

    for( j in 1:(nCrosses)){

      parents<- Parent_Combn_indices[,j]

      Parent1<-  selectedGenoData[parents[1],,]
      Parent2<-  selectedGenoData[parents[2],,]

      Cycle_Progeny_F1[j,,,] <- generateProgeny(Parent1,Parent2,1,NAM_LinkMap_New)

      Parent1<- Cycle_Progeny_F1[j,,,1]
      Parent2<- Cycle_Progeny_F1[j,,,1]

      progeny1<- generateProgeny(Parent1,Parent2,nProgeny,NAM_LinkMap_New)
      Cycle_Progeny_F2[j,,,]  <- progeny1


      for(m in 1:nProgeny){

        Parent1<- Cycle_Progeny_F2[j,,,m]
        Parent2<- Cycle_Progeny_F2[j,,,m]
        progeny1<- generateProgeny(Parent1,Parent2,1,NAM_LinkMap_New)
        Cycle_Progeny_F3[j,,,m] <- progeny1
      }


      for(m in 1:nProgeny){

        Parent1<- Cycle_Progeny_F3[j,,,m]
        Parent2<- Cycle_Progeny_F3[j,,,m]
        progeny1<- generateProgeny(Parent1,Parent2,1,NAM_LinkMap_New)
        Cycle_Progeny_F4[j,,,m] <- progeny1
      }

      for(m in 1:nProgeny){

        Parent1<- Cycle_Progeny_F4[j,,,m]
        Parent2<- Cycle_Progeny_F4[j,,,m]
        progeny1<- generateProgeny(Parent1,Parent2,1,NAM_LinkMap_New)
        Cycle_Progeny_F5[j,,,m] <- progeny1
      }

     }
	
   }else if(BD =="Drift"){ 

		if(no_selected ==2){
			nCrosses <- 1
	    }else{nCrosses <- no_selected}


		Cycle_Progeny_F1 <- array(0,c(nCrosses,nMarkers,2,1))

		Cycle_Progeny_F2 <- array(0,c(nCrosses,nMarkers,2,nProgeny))

		Cycle_Progeny_F3 <- array(0,c(nCrosses,nMarkers,2,nProgeny))

		Cycle_Progeny_F4 <- array(0,c(nCrosses,nMarkers,2,nProgeny))

		Cycle_Progeny_F5 <- array(0,c(nCrosses,nMarkers,2,nProgeny))

###################################################################################

	    for( j in 1:(nCrosses)){

## Get parents from selectedGenoData_List 
          parentLineSet <- c(1:nCrosses)
		  parent1Index <- sample(parentLineSet,1)
		  parent2Index <- sample(setdiff(parentLineSet,parent1Index),1)
			
		  Parent1<-  selectedGenoData[parent1Index,,]
		  Parent2<-  selectedGenoData[parent2Index,,]
	
## Get F1- F5 RILs
	
		  Cycle_Progeny_F1[j,,,] <- generateProgeny(Parent1,Parent2,1,NAM_LinkMap_New)
		  

		  Parent1<- Cycle_Progeny_F1[j,,,1]
		  Parent2<- Cycle_Progeny_F1[j,,,1]
		  progeny1<- generateProgeny(Parent1,Parent2,nProgeny,NAM_LinkMap_New)
		  Cycle_Progeny_F2[j,,,]  <- progeny1

		 
		  for(m in 1:nProgeny){

			Parent1<- Cycle_Progeny_F2[j,,,m]
			Parent2<- Cycle_Progeny_F2[j,,,m]
			progeny1<- generateProgeny(Parent1,Parent2,1,NAM_LinkMap_New)
			Cycle_Progeny_F3[j,,,m] <- progeny1
		  }


		  for(m in 1:nProgeny){

			Parent1<- Cycle_Progeny_F3[j,,,m]
			Parent2<- Cycle_Progeny_F3[j,,,m]
			progeny1<- generateProgeny(Parent1,Parent2,1,NAM_LinkMap_New)
			Cycle_Progeny_F4[j,,,m] <- progeny1
		  }

		  for(m in 1:nProgeny){

			Parent1<- Cycle_Progeny_F4[j,,,m]
			Parent2<- Cycle_Progeny_F4[j,,,m]
			progeny1<- generateProgeny(Parent1,Parent2,1,NAM_LinkMap_New)
			Cycle_Progeny_F5[j,,,m] <- progeny1
		  }

	  }
	}else if(BD == "RM"){ 

## RM 
        if(nSelected ==2){
			nCrosses <- 1
	    }else{nCrosses <- nSelected}

       
	    selectedGenoData <- selectedGenoData

		Cycle_Progeny_F1 <- array(0,c(nCrosses,nMarkers,2,1))

		Cycle_Progeny_F2 <- array(0,c(nCrosses,nMarkers,2,nProgeny))

		Cycle_Progeny_F3 <- array(0,c(nCrosses,nMarkers,2,nProgeny))

		Cycle_Progeny_F4 <- array(0,c(nCrosses,nMarkers,2,nProgeny))

		Cycle_Progeny_F5  <- array(0,c(nCrosses,nMarkers,2,nProgeny))

###################################################################################

			    
		for( j in 1:(nCrosses)){
		
		  parentIndices <- c(1:nCrosses)
	      parent1Index <-  sample(parentIndices,1)
		  parent2Index <- sample(parentIndices[-(parent1Index)],1)

## Get parents from selected GenoData List

		  if(j ==1 && nCrosses ==1){
			  Parent1 <-  selectedGenoData[parent1Index,,]
			  Parent2 <-  selectedGenoData[parent1Index,,]
		  }else if ((nCrosses>1)&&(j<=(nCrosses))){
		       Parent1 <-  selectedGenoData[parent1Index,,]
			   Parent2 <-  selectedGenoData[parent2Index,,]
		  }


## Get F1- F5 RILs	  
		  
		  Cycle_Progeny_F1[j,,,] <- generateProgeny(Parent1,Parent2,1,NAM_LinkMap_New)

		  Parent1<- Cycle_Progeny_F1[j,,,1]
		  Parent2<- Cycle_Progeny_F1[j,,,1]
		  progeny1<- generateProgeny(Parent1,Parent2,nProgeny,NAM_LinkMap_New)
		  Cycle_Progeny_F2[j,,,]  <- progeny1


		  for(m in 1:nProgeny){

			Parent1<- Cycle_Progeny_F2[j,,,m]
			Parent2<- Cycle_Progeny_F2[j,,,m]
			progeny1<- generateProgeny(Parent1,Parent2,1,NAM_LinkMap_New)
			Cycle_Progeny_F3[j,,,m] <- progeny1
		  }


		  for(m in 1:nProgeny){

			Parent1<- Cycle_Progeny_F3[j,,,m]
			Parent2<- Cycle_Progeny_F3[j,,,m]
			progeny1<- generateProgeny(Parent1,Parent2,1,NAM_LinkMap_New)
			Cycle_Progeny_F4[j,,,m] <- progeny1
		  }

		  for(m in 1:nProgeny){

			Parent1<- Cycle_Progeny_F4[j,,,m]
			Parent2<- Cycle_Progeny_F4[j,,,m]
			progeny1<- generateProgeny(Parent1,Parent2,1,NAM_LinkMap_New)
			Cycle_Progeny_F5[j,,,m] <- progeny1
		  }

		}

   }	     
   
  return(Cycle_Progeny_F5)

}

#############################################################


generateProgeny <- function(Parent1,Parent2,Number_Progeny,NAM_LinkMap){

  NAM_LinkMap_New <- NAM_LinkMap
  RF_Vector <- as.vector(NAM_LinkMap_New[,3])
  rr <- as.vector(NAM_LinkMap_New[,3])
  number_progeny <- Number_Progeny

  x1<-Parent1
  x2<- Parent2


  a <- Number_Progeny
  n <- dim(x1)[1]

  y<-array(0,dim=c(n,2,a))


  for (i in 1:a) {

    p1<-cumsum(runif(n)<=rr)%%2
    p2<-cumsum(runif(n)<=rr)%%2 #creating gamete from p2


    y[p1==0,1,i]=x1[p1==0,1]

    y[p1==1,1,i] = x1[p1==1,2]

    y[p2==0,2,i] = x2[p2==0,1]

    y[p2==1,2,i] = x2[p2==1,2]
  }

  progeny <- y

  return(progeny)

}


## Fn 31

 getAlleleFormat<- function(GenoTable,AlleleConversionTable_Combined){

	genoTable<- GenoTable
	nIndividuals <- nrow(genoTable)
	IA3023_alleles <- as.character(AlleleConversionTable_Combined[,8])
	Alt_Homozygous_alleles<- as.character(AlleleConversionTable_Combined[,10])
	Het_alleles <- as.character(AlleleConversionTable_Combined[,22])
	nMarkers <- ncol(genoTable)

	for(i in 1:nIndividuals){

		oneIndices <- which(genoTable[i,] ==1)
		zeroIndices <- which(genoTable[i,] ==0)
		minusOneIndices <- which(genoTable[i,] ==-1)

		genoTable[i,oneIndices] <- IA3023_alleles[oneIndices]
		genoTable[i,zeroIndices] <- Het_alleles[zeroIndices]
		genoTable[i,minusOneIndices] <- Alt_Homozygous_alleles[minusOneIndices]
	}

	return(genoTable)


 }

###########################################################
```
