## Supplementary material for "Factors Affecting Response to Recurrent Genomic Selection in Soybeans": File S1: functionRunSimulations_WGSV4_PG.R

```
### 

runSimulations40X_ML_Update <- function(model_Sol_Pheno,model_Sol_Geno,F5_Progeny_List,genoValSimList,phenoValSimList,varEList,noQTLs,H2,numberSelected,noCrosses,no_Progeny,noCycles,selectionOnGenoValues,selectionOnSimulatedValues,ModelRetrain,RetrainFrequency,ModelUpdate,UpdateType,UpdateFrequency,i,k,ModelType,weighted,weightingMethod,AlphaShape,BetaShape,TimeHorizon,trainGenoNewTable,trainSimGenoValTable,trainSimPhenoValTable,TrainTableWindowSize,NAM_LinkMap,gIndStats,alleleConversionTable_Combined){

 options(warn=-1)
 ## Assign Variables

  h2 <- H2
  no_QTL <-noQTLs

  no_selected <- numberSelected
  nCrosses <- noCrosses
  nProgeny <- no_Progeny
  nCycles <- noCycles
  selectionOnGeno <- selectionOnGenoValues
  selectionOnSimulated <- selectionOnSimulatedValues

####################################################

  PredictionModel_Pheno <- model_Sol_Pheno
  PredictionModel_Geno <- model_Sol_Geno

  cycle1_Geno_Data <- F5_Progeny_List
  genoSimValues <- genoValSimList
  phenoSimValues <- phenoValSimList

  varE <- varEList

  cycle1_nProgeny <- 100
  condition <- i
  Rep <- k
  modelRetrain <- ModelRetrain
  retrainFrequency <- RetrainFrequency 

  modelUpdate <- ModelUpdate
  updateType <- UpdateType
  updateFrequency <- UpdateFrequency
  modelType <- ModelType

  nFamilies <- 20
  cycle1_nProgeny <- 100
  nMarkers <- 4289
  nIndividuals <- 2000
  
  alphaShape <- AlphaShape
  betaShape <- BetaShape
  timeHorizon <- TimeHorizon
  alphaPar_Vector <- rep(0,nCycles)

  trainTableWindowSize <- TrainTableWindowSize 
  
  NAM_LinkMap_New <- NAM_LinkMap
  GIndStats <- gIndStats
  AlleleConversionTable_Combined <- alleleConversionTable_Combined 

### Initiate Arrays and Variables

  GenoVal_NX_2k_3c <- matrix(rep(0,(nCrosses*nProgeny)*nCycles),nrow=(nCrosses*nProgeny),ncol=nCycles)
  GenoVal_NX_N_3c <- matrix(rep(0,no_selected*nCycles),nrow=no_selected,ncol=nCycles)
  GenoVal_Sim_NX_2k_3c <- matrix(rep(0,(nCrosses*nProgeny)*nCycles),nrow=(nCrosses*nProgeny),ncol=nCycles)

  PhenoVal_NX_2k_3c <- matrix(rep(0,(nCrosses*nProgeny)*nCycles),nrow=(nCrosses*nProgeny),ncol=nCycles)
  PhenoVal_NX_N_3c <- matrix(rep(0,no_selected*nCycles),nrow=no_selected,ncol=nCycles)
  PhenoVal_Sim_NX_2k_3c <- matrix(rep(0,(nCrosses*nProgeny)*nCycles),nrow=(nCrosses*nProgeny),ncol=nCycles)

  selectedGenoIndividualIndices2 <- rep(0,no_selected)
  attainedGenoValues <- rep(0,nCycles)
  
  selectedGenoIndividualIndices_List <- list()
  selectedPhenoIndividualIndices_List <- list()
 
  trainGenoNewTablePreCycle <- as.big.matrix(trainGenoNewTable)
  trainSimPhenoTablePreCycle <- trainSimPhenoValTable
  trainSimGenoValTablePreCycle <- trainSimGenoValTable
  
  combinedTableCount <- 1
  
#######################################################################################################

	nCyc <- 1

####### Predict geno and pheno values ###################################################################

	cycle1GenoTable <- generateMinus101GenoFormat(cycle1_Geno_Data,nFamilies,cycle1_nProgeny)
	newCycle1GenoTable <- generateWeightedGenoTable(cycle1GenoTable,no_QTL)
    
  
	genoValues <- kernlab::predict(PredictionModel_Geno,newCycle1GenoTable)
    phenoValues <- kernlab::predict(PredictionModel_Pheno,newCycle1GenoTable)
	
	
    if(GIndStats == TRUE){	
###  Get population vector

		populations <- rep(0,20*100)
		init <- 1
		for(pop in 1:20){

			final <- pop*100
			populations[init:final]<- rep(pop,100)
			init <- final+1
		}
		
      cycle1GenoTable_AlleleFormat <- getAlleleFormat(cycle1GenoTable,AlleleConversionTable_Combined)
 	  cycle1GenoTable_GInd <- df2genind(cycle1GenoTable_AlleleFormat,pop=populations,sep="")
		
	  cycle1GenoData_DF <- genind2df(cycle1GenoTable_GInd)
	  write.table(cycle1GenoData_DF,paste("PM_",modelType,"_",no_QTL,"_",h2,"_GenoDataInd_Rep",Rep,"_Cyc_",nCyc,sep=""),sep="\t")
	} 	

######## Functions on Cycle 1 Geno Data ################################################################################
################################################################################

	if(selectionOnSimulated ==FALSE){
	
		if(selectionOnGeno==TRUE){
	
		  sortedGenoValues <- sort(genoValues,decreasing=TRUE,index.return=TRUE)
		  topNGenoValues <- sortedGenoValues[[1]][1:no_selected]
		  GenoTableIndices_topN <- sortedGenoValues[[2]][1:no_selected]
                  topNPhenoValues <- phenoValues[GenoTableIndices_topN] 
		  topNGenoSimValues <- genoSimValues[GenoTableIndices_topN]
		  PhenoTableIndices_topN <- GenoTableIndices_topN
		}else if (selectionOnGeno==FALSE) {
		  	  
		  sortedPhenoValues <- sort(phenoValues,decreasing=TRUE,index.return=TRUE)
		  topNPhenoValues <- sortedPhenoValues[[1]][1:no_selected]
		  PhenoTableIndices_topN <- sortedPhenoValues[[2]][1:no_selected]
		  topNGenoValues <- genoValues[PhenoTableIndices_topN]
		  topNGenoSimValues <- genoSimValues[PhenoTableIndices_topN]
		  GenoTableIndices_topN <- PhenoTableIndices_topN
		} 
  
    }
	if(selectionOnSimulated ==TRUE){
	
		if(selectionOnGeno==TRUE){
		 
   		  sortedGenoValues <- sort(genoSimValues,decreasing=TRUE,index.return=TRUE)
		  topNGenoValues <- sortedGenoValues[[1]][1:no_selected]
		  GenoTableIndices_topN <- sortedGenoValues[[2]][1:no_selected]
		  topNPhenoValues <- phenoValues[GenoTableIndices_topN] 
		  topNGenoSimValues <- genoSimValues[GenoTableIndices_topN]
		  PhenoTableIndices_topN <- GenoTableIndices_topN
		
		}else if(selectionOnGeno==FALSE){

		  sortedPhenoValues <- sort(phenoSimValues,decreasing=TRUE,index.return=TRUE)
		  topNPhenoValues <- sortedPhenoValues[[1]][1:no_selected]
		  PhenoTableIndices_topN <- sortedPhenoValues[[2]][1:no_selected]
		  topNGenoValues <- genoValues[PhenoTableIndices_topN]
		  topNGenoSimValues <- genoSimValues[PhenoTableIndices_topN]
		  GenoTableIndices_topN <- PhenoTableIndices_topN
	    }
	}

##################################################################################################################
## Assign output variables for cycle 1

	  GenoVal_Sim_NX_2k_3c[,1] <- genoSimValues
	  GenoVal_NX_2k_3c[,1] <- genoValues
	  GenoVal_NX_N_3c[,1] <- topNGenoValues
    
	  attainedGenoValues[1] <- max(topNGenoSimValues)
	  
	  selectedGenoIndividualIndices <- GenoTableIndices_topN
      selectedPhenoIndividualIndices <- PhenoTableIndices_topN
	  
	  
	  selectedGenoIndividualIndices_List[[nCyc]] <- selectedGenoIndividualIndices
	  selectedPhenoIndividualIndices_List[[nCyc]] <- selectedPhenoIndividualIndices

	  PhenoVal_Sim_NX_2k_3c[,1] <- phenoSimValues
	  PhenoVal_NX_2k_3c[,1] <- phenoValues
	  PhenoVal_NX_N_3c[,1] <- topNPhenoValues


          rm(cycle1GenoTable)
	  rm(newCycle1GenoTable)

###########################################################################
############### Cycles 2*29 ###################################################################################
### extract genotype data of selected individuals

  for( nCyc in 2:nCycles){

	print(nCyc)
	

    if(nCyc==2){

		if(selectionOnGeno == TRUE){
			selectedGenoData <- extractGenoData(selectedGenoIndividualIndices,cycle1_Geno_Data,no_selected)
		}else if(selectionOnGeno == FALSE){
			selectedGenoData <- extractGenoData(selectedPhenoIndividualIndices,cycle1_Geno_Data,no_selected)
		}

    }else if(nCyc>2){

		if(selectionOnGeno == TRUE){

			selectedGenoData <- extractGenoData(selectedGenoIndividualIndices,Cycle_Progeny_F5,no_selected)
		}else if(selectionOnGeno == FALSE){
			selectedGenoData <- extractGenoData(selectedPhenoIndividualIndices,Cycle_Progeny_F5,no_selected)
		}

    }


    Parent_Combn_indices <- getParentCombnIndices(no_selected)

    Cycle_Progeny_F1 <- array(0,c(no_selected,nMarkers,2,1))

    Cycle_Progeny_F2 <- array(0,c(no_selected,nMarkers,2,nProgeny))

    Cycle_Progeny_F3 <- array(0,c(no_selected,nMarkers,2,nProgeny))

    Cycle_Progeny_F4 <- array(0,c(no_selected,nMarkers,2,nProgeny))

    Cycle_Progeny_F5 <- array(0,c(no_selected,nMarkers,2,nProgeny))


###################################################################################

    for( j in 1:(nCrosses)){

      parents<- Parent_Combn_indices[,j]

      Parent1<-  selectedGenoData[parents[1],,]
      Parent2<-  selectedGenoData[parents[2],,]

      Cycle_Progeny_F1[j,,,] <- generateProgeny(Parent1,Parent2,1,NAM_LinkMap_New)

      Parent1<- Cycle_Progeny_F1[j,,,1]
      Parent2<- Cycle_Progeny_F1[j,,,1]

      progeny1<- generateProgeny(Parent1,Parent2,nProgeny,NAM_LinkMap_New)
      Cycle_Progeny_F2[j,,,]  <- progeny1


      for(m in 1:nProgeny){

        Parent1<- Cycle_Progeny_F2[j,,,m]
        Parent2<- Cycle_Progeny_F2[j,,,m]
        progeny1<- generateProgeny(Parent1,Parent2,1,NAM_LinkMap_New)
        Cycle_Progeny_F3[j,,,m] <- progeny1
      }


      for(m in 1:nProgeny){

        Parent1<- Cycle_Progeny_F3[j,,,m]
        Parent2<- Cycle_Progeny_F3[j,,,m]
        progeny1<- generateProgeny(Parent1,Parent2,1,NAM_LinkMap_New)
        Cycle_Progeny_F4[j,,,m] <- progeny1
      }

      for(m in 1:nProgeny){

        Parent1<- Cycle_Progeny_F4[j,,,m]
        Parent2<- Cycle_Progeny_F4[j,,,m]
        progeny1<- generateProgeny(Parent1,Parent2,1,NAM_LinkMap_New)
        Cycle_Progeny_F5[j,,,m] <- progeny1
      }

    }

########################

    nextGenGenoTable <- generateMinus101GenoFormat(Cycle_Progeny_F5,no_selected,nProgeny)

	
### Simulate genotypic and phenotypic values

    genoSimValues <- simulateGenotypicValues(nextGenGenoTable,no_QTL)
    phenoSimValues <- simulatePhenotypicValues(nextGenGenoTable,varE,no_QTL)

    newNextGenGenoTable <- generateWeightedGenoTable(nextGenGenoTable,no_QTL)


  
 if((modelUpdate==TRUE && nCyc %%updateFrequency ==0) || (modelRetrain==TRUE && nCyc%%updateFrequency ==0)){
### Extract training and test data for model build

	IndividualNames <-rep(0,(nCrosses*nProgeny))
    IndividualNames <- paste("Ind",c(1:(nCrosses*nProgeny)),sep="")

#################################################################################################
    names(phenoSimValues)<- IndividualNames
    Mean_Fixed<- rep(1,nIndividuals)

    phenotypicValuesSimTable<- cbind(phenoSimValues,Mean_Fixed)

#### Table for Simulated Genotypic Values ####################################

    names(genoSimValues)<-IndividualNames
    Mean_Fixed<- rep(1,nIndividuals)

    genotypicValuesSimTable <- cbind(genoSimValues,Mean_Fixed)

#####################

	if(modelRetrain==TRUE){

		indices<-c(1:nIndividuals)

		trainIndices<-sample(c(1:nIndividuals),(0.8*nIndividuals))
		testIndices<- indices[which(!indices %in% trainIndices)]

	#### Training Set table for Genotype and Simulated Phenotype ###################

		# trainGenoTable <- genoTable_Mod[trainIndices,]
		trainGenoNewTable <- newNextGenGenoTable[trainIndices,]
		trainSimPhenoTable <- phenotypicValuesSimTable[trainIndices,]
		trainSimGenoTable <-  genotypicValuesSimTable[trainIndices,]

	#### Validation Set table for Genotype and Simulated Phenotype ###################

		# testGenoTable<- genoTable_Mod[testIndices,]
		testGenoNewTable <- newNextGenGenoTable[testIndices,]
		testSimPhenoTable <- phenotypicValuesSimTable[testIndices,]
		testSimGenoTable <-  genotypicValuesSimTable[testIndices,]

############### Change SVMRBF
	if(!is.null(sd(trainSimGenoTable[,1])) && !is.null(sd(trainSimPhenoTable[,1])) && !is.na(sd(trainSimPhenoTable[,1])) && !is.na(sd(trainSimPhenoTable[,1]))){
	   
	  if(sd(trainSimGenoTable[,1])!=0 && sd(trainSimPhenoTable[,1])!=0 && i%%updateFrequency==0){

	    if(modelType=="SVMRBF"){
	   		PredictionModel_Geno_List<- tryCatch(SoyNAMPredictionMethods::buildSVMRBFModel(trainGenoNewTable,trainSimGenoTable,testGenoNewTable,testSimGenoTable),error=function(e) print(paste("BuildSVMRBF-Error -",e)))
			PredictionModel_Pheno_List <- tryCatch(SoyNAMPredictionMethods::buildSVMRBFModel(trainGenoNewTable,trainSimPhenoTable,testGenoNewTable,testSimPhenoTable),error=function(e) print(paste("BuildSVM-Error -",e)))
			
			PredictionModel_Geno <- PredictionModel_Geno_List[[2]]
		    PredictionModel_Pheno <- PredictionModel_Pheno_List[[2]]
		}
	  }

	 }
	
}

	gc()
	
####################################### 
	if(modelUpdate ==TRUE){
		
			
		PredictionModel_Geno_PreCycle <- PredictionModel_Geno
		PredictionModel_Pheno_PreCycle <- PredictionModel_Pheno
        
	    indices <- c(1:nIndividuals)

        trainIndices <- sample(c(1:nIndividuals),(0.8*nIndividuals))

        testIndices<- indices[which(!indices %in% trainIndices)]
		
		nTrainIndices <- length(trainIndices)

#### Training Set table for Genotype and Simulated Phenotype ###################

        trainGenoNewTable <- newNextGenGenoTable[trainIndices,]

        trainSimPhenoTable <- phenotypicValuesSimTable[trainIndices,]

        trainSimGenoValTable <-  genotypicValuesSimTable[trainIndices,]

#### Validation Set table for Genotype and Simulated Phenotype ###################
        testGenoNewTable <- newNextGenGenoTable[testIndices,]

        testSimPhenoTable <- phenotypicValuesSimTable[testIndices,]

        testSimGenoValTable <-  genotypicValuesSimTable[testIndices,]


##### Build RRBLUP prediction model every cycle


        trainGenoNewTableComb <- as.big.matrix(rbind(bigmemory::as.matrix(trainGenoNewTablePreCycle),as.matrix(trainGenoNewTable)))
        trainSimGenoValTableComb <- rbind((trainSimGenoValTablePreCycle),trainSimGenoValTable)
        trainSimPhenoValTableComb <- rbind((trainSimPhenoTablePreCycle),trainSimPhenoTable)
        print(dim(trainGenoNewTableComb))
		Avg_GenoSim_Prev <- mean(trainSimGenoValTableComb[,1])
		Avg_GenoSim_Current <- mean(trainSimGenoValTable[,1])

		change_GenoSim <- Avg_GenoSim_Prev - Avg_GenoSim_Current
		combinedTableCount <- combinedTableCount+1
        print(change_GenoSim)

        if(!is.null(sd(trainSimGenoValTableComb[,1])) && !is.null(sd(trainSimPhenoValTableComb[,1])) && !is.na(sd(trainSimGenoValTableComb[,1])) && !is.na(sd(trainSimPhenoValTableComb[,1]))){

          if(sd(trainSimGenoValTableComb[,1])!=0 && sd(trainSimPhenoValTableComb[,1])!=0 && change_GenoSim !=0 ){ 

           if(modelType=="SVMRBF"){
				PredictionModel_Geno_List <- tryCatch((SoyNAMPredictionMethods::buildSVMRBFModel(bigmemory::as.matrix(trainGenoNewTableComb),trainSimGenoValTableComb,testGenoNewTable,testSimGenoValTable)),error=function(e) print(paste("Error BigMatrix Cycle No",nCyc,"Cond",condition,"Rep",Rep,e)))
			    
				PredictionModel_Pheno_List <-tryCatch((SoyNAMPredictionMethods::buildSVMRBFModel(bigmemory::as.matrix(trainGenoNewTableComb),trainSimPhenoValTableComb,testGenoNewTable,testSimPhenoTable)),error=function(e) print(paste("Error BigMatrix Cycle No",nCyc,"Cond",condition,"Rep",Rep,e)))
			
				PredictionModel_Geno <- PredictionModel_Geno_List[[2]]
                PredictionModel_Pheno <- PredictionModel_Pheno_List[[2]]

                        
                }

	      } 

           print(sd(PredictionModel_Pheno[[1]])) 
           print(sd(PredictionModel_Geno[[1]])) 
           print(length(levels(factor(PredictionModel_Pheno[[1]]))))
           print(length(levels(factor(PredictionModel_Geno[[1]]))))	 

           genoValues <- (kernlab::predict(PredictionModel_Geno,newNextGenGenoTable))
           phenoValues <-(kernlab::predict(PredictionModel_Pheno,newNextGenGenoTable))
           print(sd(phenoValues))
           print(sd(genoValues))

 
           if(length(levels(factor(PredictionModel_Pheno[[1]]))) ==1 || length(levels(factor(PredictionModel_Geno[[1]]))) ==1 || (is.na(sd(phenoValues)) || sd(phenoValues)==0) || (is.na(sd(genoValues)) || sd(genoValues)==0)){

                         PredictionModel_Pheno <- PredictionModel_Pheno_PreCycle
                         PredictionModel_Geno <- PredictionModel_Geno_PreCycle

           }
	 }
		
		if(combinedTableCount < trainTableWindowSize){
			
			    trainGenoNewTablePreCycle <-  trainGenoNewTableComb

                trainSimPhenoTablePreCycle <-   (trainSimPhenoValTableComb)

                trainSimGenoValTablePreCycle <- (trainSimGenoValTableComb)
					
		 }else if(combinedTableCount >= trainTableWindowSize){
			
			    trainGenoNewTablePreCycle <-  trainGenoNewTableComb[-c(1:nTrainIndices),]

                trainSimPhenoTablePreCycle <-   (trainSimPhenoValTableComb[-c(1:nTrainIndices),])

                trainSimGenoValTablePreCycle <- (trainSimGenoValTableComb[-c(1:nTrainIndices),])
				
				combinedTableCount <- (trainTableWindowSize-1)
					
		  }
		
	
                rm(trainGenoNewTableComb)
                rm(trainSimGenoValTableComb)
                rm(trainSimPhenoValTableComb)
                gc()

       }	
	
 }	

############# 

if(GIndStats == TRUE){
###  Get population vector

		populations <- rep(0,20*100)
		init <- 1
		for(pop in 1:20){

			final <- pop*100
			populations[init:final]<- rep(pop,100)
			init <- final+1
		}
		
### Get GInd objects 	
	
	
	print(length(populations))
	nextGenGenoTable_AlleleFormat <- getAlleleFormat(nextGenGenoTable,AlleleConversionTable_Combined)
    nextGenGenoTable_GInd <- df2genind(as.data.frame(nextGenGenoTable_AlleleFormat),pop=populations,sep="")
    nextGenGenoData_DF <- genind2df(nextGenGenoTable_GInd)
	write.table(nextGenGenoData_DF,paste("PM_",modelType,"_",no_QTL,"_",h2,"_GenoDataInd_Rep",Rep,"_Cyc_",nCyc,sep=""),sep="\t")

}

#################get predicted geno and pheno values with JM and unweighted GP models

    genoValues <- (kernlab::predict(PredictionModel_Geno,newNextGenGenoTable))
    phenoValues <-(kernlab::predict(PredictionModel_Pheno,newNextGenGenoTable))

#############

    GenoVal_Sim_NX_2k_3c[,nCyc] <- genoSimValues
    GenoVal_NX_2k_3c[,nCyc] <- genoValues


    PhenoVal_Sim_NX_2k_3c[,nCyc]<- phenoSimValues
    PhenoVal_NX_2k_3c[,nCyc] <- phenoValues


### Selection based on simulated or predicted values

    if(selectionOnSimulated ==TRUE){

		genoSelectionTable <- ApplySelection(genoSimValues,no_selected)
		phenoSelectionTable <- ApplySelection(phenoSimValues,no_selected)

		if(selectionOnGeno==TRUE){

			selectedGenoIndividualIndices <- as.vector(genoSelectionTable[,2])
			genoSimSelectedValues <- genoSimValues[selectedGenoIndividualIndices]
			phenoSimSelectedValues <- phenoSimValues[selectedGenoIndividualIndices]

		}else if(selectionOnGeno==FALSE){

			selectedPhenoIndividualIndices <- as.vector(phenoSelectionTable[,2])
			genoSimSelectedValues <- genoSimValues[selectedPhenoIndividualIndices]
			phenoSimSelectedValues <- phenoSimValues[selectedPhenoIndividualIndices]
		}

		GenoVal_NX_N_3c[,nCyc] <- genoSimSelectedValues
	    PhenoVal_NX_N_3c[,nCyc] <- phenoSimSelectedValues
		attainedGenoValues[nCyc] <- max(genoSimSelectedValues)
		
		 selectedGenoIndividualIndices_List[[nCyc]] <- selectedGenoIndividualIndices
		selectedPhenoIndividualIndices_List[[nCyc]] <- selectedPhenoIndividualIndices

	}
	if(selectionOnSimulated==FALSE){

		genoSelectionTable <- ApplySelection(genoValues,no_selected)
		phenoSelectionTable <- ApplySelection(phenoValues,no_selected)

		if(selectionOnGeno==TRUE){

			selectedGenoIndividualIndices <- as.vector(genoSelectionTable[,2])
			genoSelectedValues <- genoValues[selectedGenoIndividualIndices]
			phenoSelectedValues <- phenoValues[selectedGenoIndividualIndices]
			genoSimSelectedValues <- genoSimValues[selectedPhenoIndividualIndices]
		
		}else if(selectionOnGeno==FALSE){

			selectedPhenoIndividualIndices <- as.vector(phenoSelectionTable[,2])
			genoSelectedValues <- genoValues[selectedPhenoIndividualIndices]
			phenoSelectedValues <- phenoValues[selectedPhenoIndividualIndices]
			genoSimSelectedValues <- genoSimValues[selectedPhenoIndividualIndices]
		
		}
		GenoVal_NX_N_3c[,nCyc]<- genoSelectedValues
	    PhenoVal_NX_N_3c[,nCyc]<- phenoSelectedValues
		attainedGenoValues[nCyc] <- max(genoSimSelectedValues)
		selectedGenoIndividualIndices_List[[nCyc]] <- selectedGenoIndividualIndices
		selectedPhenoIndividualIndices_List[[nCyc]] <- selectedPhenoIndividualIndices

	}

    
	rm(nextGenGenoTable)
	rm(newNextGenGenoTable)
    gc()
  }

  simResults_List <- list(GenoVal_Sim_NX_2k_3c,GenoVal_NX_2k_3c,GenoVal_NX_N_3c, PhenoVal_Sim_NX_2k_3c,PhenoVal_NX_2k_3c,PhenoVal_NX_N_3c,attainedGenoValues,selectedGenoIndividualIndices_List,selectedPhenoIndividualIndices_List)


  return(simResults_List)

}


###


runSimulations40X_Bayes_WGS_V2 <- function(model_Sol_Pheno,model_Sol_Geno,F5_Progeny_List,genoValSimList,phenoValSimList,varEList,noQTLs,H2,numberSelected,noCrosses,no_Progeny,noCycles,selectionOnGenoValues,selectionOnSimulatedValues,ModelRetrain,RetrainFrequency,ModelUpdate,UpdateType,UpdateFrequency,i,k,ModelType,weighted,weightingMethod,AlphaShape,BetaShape,TimeHorizon,trainGenoNewTable,trainSimGenoValTable,trainSimPhenoValTable,TrainTableWindowSize,NAM_LinkMap,gIndStats,alleleConversionTable_Combined){

 options(warn=-1)

### Assign Variables

  h2 <- H2
  no_QTL <-noQTLs

  no_selected <- numberSelected
  nCrosses <- noCrosses
  nProgeny <- no_Progeny
  nCycles <- noCycles
  selectionOnGeno <- selectionOnGenoValues
  selectionOnSimulated <- selectionOnSimulatedValues

####################################################################################################

  PredictionModel_Pheno <- model_Sol_Pheno
  PredictionModel_Geno <- model_Sol_Geno

  cycle1_Geno_Data <- F5_Progeny_List
  genoSimValues <- genoValSimList
  phenoSimValues <- phenoValSimList

  varE <- varEList

  nFamilies <- 20
  cycle1_nProgeny <- 100

  modelType <- ModelType
  modelRetrain <- ModelRetrain
  retrainFrequency <- RetrainFrequency 

  modelUpdate <- ModelUpdate
  updateFrequency <- UpdateFrequency
  updateType <- UpdateType

  nMarkers <- 4289
  nIndividuals <- 2000

  Weighted <- weighted 
  WghtMethod <- weightingMethod

  
  alphaShape <- AlphaShape
  betaShape <- BetaShape
  timeHorizon <- TimeHorizon
  alphaPar_Vector <- rep(0,nCycles)

  condition <- i
   Rep <- k
  
    
  trainTableWindowSize <- TrainTableWindowSize
#######################################################################################################
   NAM_LinkMap_New <- NAM_LinkMap 
   GIndStats <- gIndStats
   AlleleConversionTable_Combined <- alleleConversionTable_Combined
  
   nCyc <- 1
   print(paste("cycle_No",nCyc))
           
   trainGeno_backingFileName <- paste("trainTable_",no_QTL,"_",h2,"_",condition,"_",Rep,"_Cyc",nCyc,".bin",sep="")
   trainGeno_descriptorFileName <- paste("trainTable_",no_QTL,"_",h2,"_",condition,"_",Rep,"_Cyc",nCyc,".desc",sep="")
 							
   trainGenoNewTablePreCycle <- deepcopy(as.big.matrix(trainGenoNewTable),backingfile=trainGeno_backingFileName,descriptorfile=trainGeno_descriptorFileName)
   
   trainSimPhenoTablePreCycle <- trainSimPhenoValTable  
   trainSimGenoValTablePreCycle <- trainSimGenoValTable

####### Predict geno and pheno values ###################################################################

	cycle1GenoTable <- generateMinus101GenoFormat(cycle1_Geno_Data,nFamilies,cycle1_nProgeny)
	newCycle1GenoTable <- generateWeightedGenoTable(cycle1GenoTable,no_QTL)

    if(selectionOnGeno==TRUE){
		MarkerEffects <- unlist(PredictionModel_Geno[3])
	}else if(selectionOnGeno==FALSE){
	    MarkerEffects <- unlist(PredictionModel_Pheno[3])
	}

    Freq_List <- getFreq_BasePoln(cycle1GenoTable,MarkerEffects)
	Freq <- Freq_List[[1]]
	FavAllele <- Freq_List[[2]]

	if(Weighted==TRUE && WghtMethod=="JM"){
		genoValues  <- PredictBayesPhenoValues_Wght(newCycle1GenoTable,PredictionModel_Geno,Freq)
		phenoValues <- PredictBayesPhenoValues_Wght(newCycle1GenoTable,PredictionModel_Pheno,Freq)
	}else if(Weighted==TRUE && WghtMethod=="DW" ){
		genoValues_PredList <- PredictBayesPhenoValues_WGS_DWModel(newCycle1GenoTable,PredictionModel_Geno,Freq,alphaShape,betaShape,cycleNumber,timeHorizon)
		phenoValues_PredList <- PredictBayesPhenoValues_WGS_DWModel(newCycle1GenoTable,PredictionModel_Pheno,Freq,alphaShape,betaShape,cycleNumber,timeHorizon)
		
		genoValues <- genoValues_PredList[[1]]
		phenoValues <- phenoValues_PredList[[1]]
		alphaPar_Vector[nCyc] <- genoValues_PredList[[2]]
	}else if(Weighted==FALSE){
		
		genoValues <- PredictBayesPhenoValues(newCycle1GenoTable,PredictionModel_Geno)
		phenoValues <- PredictBayesPhenoValues(newCycle1GenoTable,PredictionModel_Pheno)
	}
	
	if(GIndStats == TRUE){	 
###  Get population vector

		populations <- rep(0,20*100)
		init <- 1
		for(pop in 1:20){

			final <- pop*100
			populations[init:final]<- rep(pop,100)
			init <- final+1
		}
		
      cycle1GenoTable_AlleleFormat <- getAlleleFormat(cycle1GenoTable,AlleleConversionTable_Combined)
 	  cycle1GenoTable_GInd <- df2genind(cycle1GenoTable_AlleleFormat,pop=populations,sep="")
		
	  cycle1GenoData_DF <- genind2df(cycle1GenoTable_GInd)
	  write.table(cycle1GenoData_DF,paste("PM_",modelType,"_",no_QTL,"_",h2,"_GenoDataInd_Rep",Rep,"_Cyc_",nCyc,sep=""),sep="\t")
	  	
	}
    
### Initiate Arrays and Variables

  GenoVal_NX_2k_3c <- matrix(rep(0,(nCrosses*nProgeny)*nCycles),nrow=(nCrosses*nProgeny),ncol=nCycles)
  GenoVal_NX_N_3c <- matrix(rep(0,no_selected*nCycles),nrow=no_selected,ncol=nCycles)
  GenoVal_Sim_NX_2k_3c <- matrix(rep(0,(nCrosses*nProgeny)*nCycles),nrow=(nCrosses*nProgeny),ncol=nCycles)

  PhenoVal_NX_2k_3c <- matrix(rep(0,(nCrosses*nProgeny)*nCycles),nrow=(nCrosses*nProgeny),ncol=nCycles)
  PhenoVal_NX_N_3c <- matrix(rep(0,no_selected*nCycles),nrow=no_selected,ncol=nCycles)
  PhenoVal_Sim_NX_2k_3c <- matrix(rep(0,(nCrosses*nProgeny)*nCycles),nrow=(nCrosses*nProgeny),ncol=nCycles)

  attainedGenoValues <- rep(0,nCycles)
  selectedGenoIndividualIndices_List <- list()
  selectedPhenoIndividualIndices_List <- list()
  
######## Functions on Cycle 1 Geno Data ################################################################################

	if(selectionOnSimulated ==FALSE){
		  sortedGenoValues <- sort(genoValues,decreasing=TRUE,index.return=TRUE)
		  topNGenoValues <- sortedGenoValues[[1]][1:no_selected]
		  GenoTableIndices_topN <- sortedGenoValues[[2]][1:no_selected]

		  sortedPhenoValues <- sort(phenoValues,decreasing=TRUE,index.return=TRUE)
		  topNPhenoValues <- sortedPhenoValues[[1]][1:no_selected]
		  PhenoTableIndices_topN <- sortedPhenoValues[[2]][1:no_selected]

### Sort genoSim values for  attainedGenoValues

		  sortedGenoSimValues <- sort(genoSimValues,decreasing=TRUE,index.return=TRUE)
		  topNGenoSimValues <- sortedGenoSimValues[[1]][1:no_selected]

	} else if(selectionOnSimulated ==TRUE){

		  sortedGenoValues <- sort(genoSimValues,decreasing=TRUE,index.return=TRUE)
		  topNGenoValues <- sortedGenoValues[[1]][1:no_selected]
		  GenoTableIndices_topN <- sortedGenoValues[[2]][1:no_selected]

		  sortedPhenoValues <- sort(phenoSimValues,decreasing=TRUE,index.return=TRUE)
		  topNPhenoValues <- sortedPhenoValues[[1]][1:no_selected]
		  PhenoTableIndices_topN <- sortedPhenoValues[[2]][1:no_selected]

### Sort genoSim values for  attainedGenoValues

		  sortedGenoSimValues <- sort(genoSimValues,decreasing=TRUE,index.return=TRUE)
		  topNGenoSimValues <- sortedGenoSimValues[[1]][1:no_selected]
	}

## Assign output variables for cycle 1

	  GenoVal_Sim_NX_2k_3c[,1] <- genoSimValues
	  GenoVal_NX_2k_3c[,1] <- genoValues
	  GenoVal_NX_N_3c[,1] <- topNGenoValues
	
	  PhenoVal_Sim_NX_2k_3c[,1] <- phenoSimValues
	  PhenoVal_NX_2k_3c[,1] <- phenoValues
	  PhenoVal_NX_N_3c[,1] <- topNPhenoValues

	  attainedGenoValues[1] <- max(topNGenoSimValues)
	  
	  selectedGenoIndividualIndices <- GenoTableIndices_topN
	  selectedPhenoIndividualIndices <- PhenoTableIndices_topN
	  
	  selectedGenoIndividualIndices_List[[nCyc]] <- selectedGenoIndividualIndices
	  selectedPhenoIndividualIndices_List[[nCyc]] <- selectedPhenoIndividualIndices
		

      rm(cycle1GenoTable)
	  rm(newCycle1GenoTable) 

    
	  
	  combinedTableCount <- 1

############### Cycles 2*29 ###################################################################################
### extract genotype data of selected individuals

  for(nCyc in 2:nCycles){
  
   	print(paste("Cycle_No",nCyc))

    if(nCyc==2){

		if(selectionOnGeno == TRUE){
			selectedGenoData <- extractGenoData(selectedGenoIndividualIndices,cycle1_Geno_Data,no_selected)
		}else if(selectionOnGeno == FALSE){
			selectedGenoData <- extractGenoData(selectedPhenoIndividualIndices,cycle1_Geno_Data,no_selected)
		}

    }
    if(nCyc>2){

		if(selectionOnGeno == TRUE){

			selectedGenoData <- extractGenoData(selectedGenoIndividualIndices,Cycle_Progeny_F5,no_selected)
		}else if(selectionOnGeno == FALSE){
			selectedGenoData <- extractGenoData(selectedPhenoIndividualIndices,Cycle_Progeny_F5,no_selected)
		}

    }


    Parent_Combn_indices <- getParentCombnIndices(no_selected)

    Cycle_Progeny_F1 <- array(0,c(no_selected,nMarkers,2,1))

    Cycle_Progeny_F2 <- array(0,c(no_selected,nMarkers,2,nProgeny))

    Cycle_Progeny_F3 <- array(0,c(no_selected,nMarkers,2,nProgeny))

    Cycle_Progeny_F4 <- array(0,c(no_selected,nMarkers,2,nProgeny))

    Cycle_Progeny_F5 <- array(0,c(no_selected,nMarkers,2,nProgeny))


###################################################################################

    for( j in 1:(nCrosses)){

      parents<- Parent_Combn_indices[,j]

      Parent1<-  selectedGenoData[parents[1],,]
      Parent2<-  selectedGenoData[parents[2],,]

      Cycle_Progeny_F1[j,,,] <- generateProgeny(Parent1,Parent2,1,NAM_LinkMap_New)

      Parent1<- Cycle_Progeny_F1[j,,,1]
      Parent2<- Cycle_Progeny_F1[j,,,1]

      progeny1<- generateProgeny(Parent1,Parent2,nProgeny,NAM_LinkMap_New)
      Cycle_Progeny_F2[j,,,]  <- progeny1


      for(m in 1:nProgeny){

        Parent1<- Cycle_Progeny_F2[j,,,m]
        Parent2<- Cycle_Progeny_F2[j,,,m]
        progeny1<- generateProgeny(Parent1,Parent2,1,NAM_LinkMap_New)
        Cycle_Progeny_F3[j,,,m] <- progeny1
      }


      for(m in 1:nProgeny){

        Parent1<- Cycle_Progeny_F3[j,,,m]
        Parent2<- Cycle_Progeny_F3[j,,,m]
        progeny1<- generateProgeny(Parent1,Parent2,1,NAM_LinkMap_New)
        Cycle_Progeny_F4[j,,,m] <- progeny1
      }

      for(m in 1:nProgeny){

        Parent1<- Cycle_Progeny_F4[j,,,m]
        Parent2<- Cycle_Progeny_F4[j,,,m]
        progeny1<- generateProgeny(Parent1,Parent2,1,NAM_LinkMap_New)
        Cycle_Progeny_F5[j,,,m] <- progeny1
      }

    }

#######################

    nextGenGenoTable <- generateMinus101GenoFormat(Cycle_Progeny_F5,no_selected,nProgeny)
    Freq <- getFreq(nextGenGenoTable,FavAllele)

### Simulate genotypic and phenotypic values 

    genoSimValues <- simulateGenotypicValues(nextGenGenoTable,no_QTL)
    phenoSimValues <- simulatePhenotypicValues(nextGenGenoTable,varE,no_QTL)
    newNextGenGenoTable <- generateWeightedGenoTable(nextGenGenoTable,no_QTL)


if(nCyc%%updateFrequency==0 && modelUpdate ==TRUE){
## Extract training and test data for model build
    IndividualNames <-rep(0,(nCrosses*nProgeny))
      
    IndividualNames <- paste("Ind",c(1:(nCrosses*nProgeny)),sep="")
 
##################################################################################################

    names(phenoSimValues)<-IndividualNames

    Mean_Fixed<- rep(1,nIndividuals)

    phenotypicValuesSimTable <- cbind(phenoSimValues,Mean_Fixed)

#### Table for Simulated Genotypic Values ####################################

    names(genoSimValues)<-IndividualNames

    Mean_Fixed<- rep(1,nIndividuals)

    genotypicValuesSimTable <- cbind(genoSimValues,Mean_Fixed)

####################################
### Model update

    set.seed(25+Rep+nCyc)

	if(modelRetrain==TRUE){ 
        PredDefined <- FALSE

        while(PredDefined ==FALSE){

		indices<-c(1:nIndividuals)

		trainIndices<-sample(c(1:nIndividuals),(0.8*nIndividuals))
		testIndices<- indices[which(!indices %in% trainIndices)]

#### Training Set table for Genotype and Simulated Phenotype ###################

		# trainGenoTable <- genoTable_Mod[trainIndices,]
		trainGenoNewTable <- newNextGenGenoTable[trainIndices,]
		trainSimPhenoTable <- phenotypicValuesSimTable[trainIndices,]
		trainSimGenoTable <-  genotypicValuesSimTable[trainIndices,]

#### Validation Set table for Genotype and Simulated Phenotype ###################

		# testGenoTable<- genoTable_Mod[testIndices,]
		testGenoNewTable <- newNextGenGenoTable[testIndices,]
		testSimPhenoTable <- phenotypicValuesSimTable[testIndices,]
		testSimGenoTable <-  genotypicValuesSimTable[testIndices,]

	    print(length(trainSimGenoTable[,1]))
		print(sd(trainSimGenoTable[,1]))	
		print(sd(trainSimPhenoTable[,1]))
        print(i%%updateFrequency) 
		print(nCyc%%updateFrequency)
        print(summary(trainSimPhenoTable[,1]))		
############### Change RRBLUP to Bayes
      if(!is.null(sd(trainSimGenoTable[,1])) && !is.null(sd(trainSimPhenoTable[,1])) && !is.na(sd(trainSimGenoTable[,1])) && !is.na(sd(trainSimPhenoTable[,1]))){
	    if(sd(trainSimGenoTable[,1])!=0 && sd(trainSimPhenoTable[,1])!=0 && nCyc%%retrainFrequency==0){

			if(modelType=="RRBLUP"){
				PredictionModel_Geno <- (buildRRBLUPModel(trainGenoNewTable,trainSimGenoTable,testGenoNewTable,testSimGenoTable))
				PredictionModel_Pheno <-(buildRRBLUPModel(trainGenoNewTable,trainSimPhenoTable,testGenoNewTable,testSimPhenoTable))
			}else if(modelType=="RRBLUP_REML"){
				PredictionModel_Geno <- (buildRRBLUPModel_REML(trainGenoNewTable,trainSimGenoTable,testGenoNewTable,testSimGenoTable))
				PredictionModel_Pheno <-(buildRRBLUPModel_REML(trainGenoNewTable,trainSimPhenoTable,testGenoNewTable,testSimPhenoTable))
			}
	    }
	  }
	
        if(length(levels(factor(PredictionModel_Pheno[[1]])))>1 && mean(PredictionModel_Pheno[[1]])>0) {
                         PredDefined <- TRUE }      
        }
   }

	gc()	
		
	if(modelUpdate ==TRUE && updateType=="FullSet"){

        indices<-c(1:nIndividuals)

        trainIndices<-sample(c(1:nIndividuals),(0.8*nIndividuals))

        testIndices<- indices[which(!indices %in% trainIndices)]

#### Training Set table for Genotype and Simulated Phenotype ###################

        trainGenoNewTable <- newNextGenGenoTable[trainIndices,]

        trainSimPhenoTable <- phenotypicValuesSimTable[trainIndices,]

        trainSimGenoValTable <-  genotypicValuesSimTable[trainIndices,]

#### Validation Set table for Genotype and Simulated Phenotype ###################

        testGenoNewTable <- newNextGenGenoTable[testIndices,]

        testSimPhenoTable <- phenotypicValuesSimTable[testIndices,]

        testSimGenoValTable <-  genotypicValuesSimTable[testIndices,]


##### Build RRBLUP prediction model every cycle


        trainGenoNewTableComb <- as.big.matrix(rbind(bigmemory::as.matrix(trainGenoNewTablePreCycle),as.matrix(trainGenoNewTable)))

        trainSimGenoValTableComb <- rbind((trainSimGenoValTablePreCycle),trainSimGenoValTable)

        trainSimPhenoValTableComb <- rbind((trainSimPhenoTablePreCycle),trainSimPhenoTable)

        print(dim(trainGenoNewTableComb))

        if(!is.null(sd(trainSimGenoValTableComb[,1])) && !is.null(sd(trainSimPhenoValTableComb[,1])) && !is.na(sd(trainSimGenoValTableComb[,1])) && !is.na(sd(trainSimPhenoValTableComb[,1]))){
            if(sd(trainSimGenoValTableComb[,1])!=0 && sd(trainSimPhenoValTableComb[,1])!=0 && nCyc%%updateFrequency==0){ 


                    
            if(modelType=="BayesB"){
					PredictionModel_Geno <- tryCatch(buildBayesBModel(bigmemory::as.matrix(trainGenoNewTableComb),trainSimGenoValTableComb[,1],testGenoNewTable,testSimGenoValTable[,1],h2),error=function(e) print(paste("BuildBB-Error -",nCyc,e)))
					PredictionModel_Pheno <- tryCatch(buildBayesBModel(bigmemory::as.matrix(trainGenoNewTableComb),trainSimPhenoValTableComb[,1],testGenoNewTable,testSimPhenoTable[,1],h2),error=function(e) print(paste("BuildBB-Error -",nCyc,e)))
			}else if(modelType=="BL"){
			
					PredictionModel_Geno <- tryCatch(buildBLModel(bigmemory::as.matrix(trainGenoNewTableComb),trainSimGenoValTableComb[,1],testGenoNewTable,testSimGenoValTable[,1],h2),error=function(e) print(paste("BuildBL-Error -",nCyc,e)))
					PredictionModel_Pheno <- tryCatch(buildBLModel(bigmemory::as.matrix(trainGenoNewTableComb),trainSimPhenoValTableComb[,1],testGenoNewTable,testSimPhenoTable[,1],h2),error=function(e) print(paste("BuildBL-Error -",nCyc,e)))
			}

	      } 
			   

                trainGenoNewTablePreCycle <-  trainGenoNewTableComb

                trainSimPhenoTablePreCycle <-   (trainSimPhenoValTableComb)

                trainSimGenoValTablePreCycle <- (trainSimGenoValTableComb)


                rm(trainGenoNewTableComb)

                rm(trainSimGenoValTableComb)

                rm(trainSimPhenoValTableComb)

                gc()

        }

    }
    

	if(modelUpdate ==TRUE && updateType=="TrainingCycleWindow"){

			
		PredictionModel_Geno_PreCycle <- PredictionModel_Geno
		PredictionModel_Pheno_PreCycle <- PredictionModel_Pheno
        
        indices<-c(1:nIndividuals)

        trainIndices<- sample(c(1:nIndividuals),(0.8*nIndividuals))

        testIndices<- indices[which(!indices %in% trainIndices)]
		
	    nTrainIndices <- length(trainIndices)

#### Training Set table for Genotype and Simulated Phenotype ###################

        trainGenoNewTable <- newNextGenGenoTable[trainIndices,]

        trainSimPhenoTable <- phenotypicValuesSimTable[trainIndices,]

        trainSimGenoValTable <-  genotypicValuesSimTable[trainIndices,]

#### Validation Set table for Genotype and Simulated Phenotype ###################
        testGenoNewTable <- newNextGenGenoTable[testIndices,]

        testSimPhenoTable <- phenotypicValuesSimTable[testIndices,]

        testSimGenoValTable <-  genotypicValuesSimTable[testIndices,]
	
		trainGenoNewTablePreCycle <- attach.big.matrix(trainGeno_descriptorFileName)
						
##### Build RRBLUP prediction model every cycle
         # bigmemory::as.matrix

        trainGenoNewTableComb <- (rbind(bigmemory::as.matrix(trainGenoNewTablePreCycle),as.matrix(trainGenoNewTable)))
        trainSimGenoValTableComb <- rbind((trainSimGenoValTablePreCycle),trainSimGenoValTable)
        trainSimPhenoValTableComb <- rbind((trainSimPhenoTablePreCycle),trainSimPhenoTable)
        print(dim(trainGenoNewTableComb))
	    Avg_GenoSim_Prev <- mean(trainSimGenoValTableComb[,1])
	    Avg_GenoSim_Current <- mean(trainSimGenoValTable[,1])

		change_GenoSim <- Avg_GenoSim_Prev - Avg_GenoSim_Current
		combinedTableCount <- combinedTableCount+1
        print(change_GenoSim)

        if(!is.null(sd(trainSimGenoValTableComb[,1])) && !is.null(sd(trainSimPhenoValTableComb[,1])) && !is.na(sd(trainSimGenoValTableComb[,1])) && !is.na(sd(trainSimPhenoValTableComb[,1]))){

       	if(sd(trainSimGenoValTableComb[,1])!=0 && sd(trainSimPhenoValTableComb[,1])!=0 && change_GenoSim !=0){

	        if(modelType=="BayesB"){
					PredictionModel_Geno <-  tryCatch(buildBayesBModel((trainGenoNewTableComb),trainSimGenoValTableComb[,1],testGenoNewTable,testSimGenoValTable[,1],h2),error=function(e) print(paste("BuildRR-Error -",nCyc,e)))
					PredictionModel_Pheno <- tryCatch(buildBayesBModel((trainGenoNewTableComb),trainSimPhenoValTableComb[,1],testGenoNewTable,testSimPhenoTable[,1],h2),error=function(e) print(paste("BuildRR-Error -",nCyc,e)))
			}else if(modelType=="BL"){
			
					PredictionModel_Geno <- tryCatch(buildBLModel((trainGenoNewTableComb),trainSimGenoValTableComb[,1],testGenoNewTable,testSimGenoValTable[,1],h2),error=function(e) print(paste("BuildRR-Error -",nCyc,e)))
					PredictionModel_Pheno <- tryCatch(buildBLModel((trainGenoNewTableComb),trainSimPhenoValTableComb[,1],testGenoNewTable,testSimPhenoTable[,1],h2),error=function(e) print(paste("BuildRR-Error -",nCyc,e)))
			}

	    }
		
			

           print(sd(PredictionModel_Pheno[[3]])) 
           print(sd(PredictionModel_Geno[[3]])) 
           print(length(levels(factor(PredictionModel_Pheno[[3]]))))
           print(length(levels(factor(PredictionModel_Geno[[3]]))))	 

           genoValues <- PredictBayesPhenoValues(newNextGenGenoTable,PredictionModel_Geno)
           phenoValues <- PredictBayesPhenoValues(newNextGenGenoTable,PredictionModel_Pheno)
           print(sd(phenoValues))
           print(sd(genoValues))

 
           if(length(levels(factor(PredictionModel_Pheno[[3]]))) ==1 || length(levels(factor(PredictionModel_Geno[[3]]))) ==1 || (is.na(sd(phenoValues)) || sd(phenoValues)==0) || (is.na(sd(genoValues)) || sd(genoValues)==0) || change_GenoSim==0){

                         PredictionModel_Pheno <- PredictionModel_Pheno_PreCycle
                         PredictionModel_Geno <- PredictionModel_Geno_PreCycle

           }
	}
	 
	 
	 
     trainGeno_backingFileName <- paste("trainTable_",no_QTL,"_",h2,"_",condition,"_",Rep,"_Cyc",nCyc,".bin",sep="")
     trainGeno_descriptorFileName <- paste("trainTable_",no_QTL,"_",h2,"_",condition,"_",Rep,"_Cyc",nCyc,".desc",sep="")
	     		
		if(combinedTableCount < trainTableWindowSize){
			
							
				trainGenoNewTablePreCycle <- deepcopy(as.big.matrix(trainGenoNewTableComb),backingfile=trainGeno_backingFileName,descriptorfile=trainGeno_descriptorFileName)

                trainSimPhenoTablePreCycle <-   (trainSimPhenoValTableComb)

                trainSimGenoValTablePreCycle <- (trainSimGenoValTableComb)
					
		}else if(combinedTableCount >= trainTableWindowSize){
			
			  				
				trainGenoNewTablePreCycle <- deepcopy(as.big.matrix(trainGenoNewTableComb[-c(1:nTrainIndices),]),backingfile=trainGeno_backingFileName,descriptorfile=trainGeno_descriptorFileName)

                trainSimPhenoTablePreCycle <-   (trainSimPhenoValTableComb[-c(1:nTrainIndices),])

                trainSimGenoValTablePreCycle <- (trainSimGenoValTableComb[-c(1:nTrainIndices),])
				
				combinedTableCount <- (trainTableWindowSize-1)
						
					
		}
	
    gc()

   }
 
 }  

gc()

#################get predicted geno and pheno values with JM and unweighted GP models

    if(Weighted==TRUE && WghtMethod =="JM"){
		genoValues <- PredictBayesPhenoValues_Wght(newNextGenGenoTable,PredictionModel_Geno,Freq)
		phenoValues <- PredictBayesPhenoValues_Wght(newNextGenGenoTable,PredictionModel_Pheno,Freq)
    } else if (Weighted==TRUE && WghtMethod=="DW"){
		genoValues_PredList <- PredictBayesPhenoValues_WGS_DWModel(newNextGenGenoTable,PredictionModel_Geno,Freq,alphaShape,betaShape,cycleNumber,timeHorizon)
		phenoValues_PredList <- PredictBayesPhenoValues_WGS_DWModel(newNextGenGenoTable,PredictionModel_Pheno,Freq,alphaShape,betaShape,cycleNumber,timeHorizon)

		genoValues <- genoValues_PredList[[1]]
		phenoValues <- phenoValues_PredList[[1]]

		alphaPar_Vector[nCyc] <- genoValues_PredList[[2]]
     }else if(Weighted ==FALSE){
		genoValues <- PredictBayesPhenoValues(newNextGenGenoTable,PredictionModel_Geno)
		phenoValues <- PredictBayesPhenoValues(newNextGenGenoTable,PredictionModel_Pheno)
    }

   	if(GIndStats == TRUE){ 
###  Get population vector

		populations <- rep(0,20*100)
		init <- 1
		for(pop in 1:20){

			final <- pop*100
			populations[init:final]<- rep(pop,100)
			init <- final+1
		}
		
### Get GInd objects 	

		print(length(populations))
		nextGenGenoTable_AlleleFormat <- getAlleleFormat(nextGenGenoTable,AlleleConversionTable_Combined)
		nextGenGenoTable_GInd <- df2genind(as.data.frame(nextGenGenoTable_AlleleFormat),pop=populations,sep="")
		nextGenGenoData_DF <- genind2df(nextGenGenoTable_GInd)
		write.table(nextGenGenoData_DF,paste("PM_",modelType,"trainTable_Cyc_",trainTableWindowSize,"_",no_QTL,"_",h2,"_GenoDataInd_Rep",Rep,"_Cyc_",nCyc,sep=""),sep="\t")	

    }
#############
    print(paste("GenoVal_Length",length(genoValues)))
    print(paste("PhenoVal_Length",length(phenoValues)))

    GenoVal_Sim_NX_2k_3c[,nCyc] <- genoSimValues
    GenoVal_NX_2k_3c[,nCyc] <- genoValues
   
    PhenoVal_Sim_NX_2k_3c[,nCyc]<- phenoSimValues
    PhenoVal_NX_2k_3c[,nCyc] <- phenoValues
   

### Selection based on simulated or predicted values

    if(selectionOnSimulated ==TRUE){

		genoSelectionTable <- ApplySelection(genoSimValues,no_selected)
		phenoSelectionTable <- ApplySelection(phenoSimValues,no_selected)


		if(selectionOnGeno==TRUE){

			selectedGenoIndividualIndices <- as.vector(genoSelectionTable[,2])
			genoSimSelectedValues <- genoSimValues[selectedGenoIndividualIndices]
			phenoSimSelectedValues <- phenoSimValues[selectedGenoIndividualIndices]

		}else if(selectionOnGeno==FALSE){

			selectedPhenoIndividualIndices <- as.vector(phenoSelectionTable[,2])
			genoSimSelectedValues <- genoSimValues[selectedPhenoIndividualIndices]
			phenoSimSelectedValues <- phenoSimValues[selectedPhenoIndividualIndices]
		}

		GenoVal_NX_N_3c[,nCyc] <- genoSimSelectedValues
	    PhenoVal_NX_N_3c[,nCyc] <- phenoSimSelectedValues
		attainedGenoValues[nCyc] <- max(genoSimSelectedValues)
		
		selectedGenoIndividualIndices_List[[nCyc]] <- selectedGenoIndividualIndices
		selectedPhenoIndividualIndices_List[[nCyc]] <- selectedPhenoIndividualIndices
		

	}
    
	if(selectionOnSimulated==FALSE){

		genoSelectionTable <- ApplySelection(genoValues,no_selected)
		phenoSelectionTable <- ApplySelection(phenoValues,no_selected)

        print(paste("genoSelectionTable-dim-",dim(genoSelectionTable)))
		print(paste("phenoSelectionTable-dim-",dim(phenoSelectionTable)))
		print(paste("selectedGenoIndividualIndices",is.na(phenoSelectionTable[,2])))

		if(selectionOnGeno==TRUE){

			selectedGenoIndividualIndices <- as.vector(genoSelectionTable[,2])
			genoSelectedValues <- genoValues[selectedGenoIndividualIndices]
			phenoSelectedValues <- phenoValues[selectedGenoIndividualIndices]
		    genoSimSelectedValues <- genoSimValues[selectedPhenoIndividualIndices]
		
		}else if(selectionOnGeno==FALSE){

			selectedPhenoIndividualIndices <- as.vector(phenoSelectionTable[,2])
			genoSelectedValues <- genoValues[selectedPhenoIndividualIndices]
			phenoSelectedValues <- phenoValues[selectedPhenoIndividualIndices]
		        genoSimSelectedValues <- genoSimValues[selectedPhenoIndividualIndices]
		}
	        GenoVal_NX_N_3c[,nCyc]<- genoSelectedValues
	        PhenoVal_NX_N_3c[,nCyc]<- phenoSelectedValues
        	attainedGenoValues[nCyc] <- max(genoSimSelectedValues)
			
			selectedGenoIndividualIndices_List[[nCyc]] <- selectedGenoIndividualIndices
			selectedPhenoIndividualIndices_List[[nCyc]] <- selectedPhenoIndividualIndices
    }

   	rm(nextGenGenoTable)
	rm(newNextGenGenoTable)
    gc()
  
   }
 
  simResults_List <- list(GenoVal_Sim_NX_2k_3c,GenoVal_NX_2k_3c,GenoVal_NX_N_3c,PhenoVal_Sim_NX_2k_3c,PhenoVal_NX_2k_3c,PhenoVal_NX_N_3c,attainedGenoValues,selectedGenoIndividualIndices_List,selectedPhenoIndividualIndices_List)

  return(simResults_List)

}  


### 

runSimulations40X_RRBLUP_WGS_V3 <- function(model_Sol_Pheno,model_Sol_Geno,F5_Progeny_List,genoValSimList,phenoValSimList,varEList,noQTLs,H2,numberSelected,noCrosses,no_Progeny,noCycles,selectionOnGenoValues,selectionOnSimulatedValues,ModelRetrain,RetrainFrequency,ModelUpdate,UpdateType,UpdateFrequency,i,k,ModelType,weighted,weightingMethod,AlphaShape,BetaShape,TimeHorizon,trainGenoNewTable,trainSimGenoValTable,trainSimPhenoValTable,TrainTableWindowSize,BreedingDesign,NAM_LinkMap,gIndStats,alleleConversionTable_Combined){


### Assign Variables

	  h2 <- H2
	  no_QTL <-noQTLs

	  no_selected <- numberSelected
	  nCrosses <- noCrosses
	  nProgeny <- no_Progeny
	  nCycles <- noCycles
	  selectionOnGeno <- selectionOnGenoValues
	  selectionOnSimulated <- selectionOnSimulatedValues

####################################################

	  PredictionModel_Pheno <- model_Sol_Pheno
	  PredictionModel_Geno <- model_Sol_Geno

	  cycle1_Geno_Data <- F5_Progeny_List
	  genoSimValues <- genoValSimList
	  phenoSimValues <- phenoValSimList

	  varE <- varEList

	  condition <- i
	  Rep <- k
	  modelType <- ModelType
	  modelRetrain <- ModelRetrain
	  updateFrequency <- UpdateFrequency
	  retrainFrequency <- RetrainFrequency
	  updateType <- UpdateType
    modelUpdate <- ModelUpdate

	  nFamilies <- 20
    cycle1_nProgeny <- 100
	  nMarkers <- 4289
    nIndividuals <- 2000

    Weighted <- weighted
	  WghtMethod <- weightingMethod  
	  alphaShape <- AlphaShape
	  betaShape <- BetaShape
	  timeHorizon <- TimeHorizon
	  alphaPar_Vector <- rep(0,nCycles)
    BD <- BreedingDesign
         
	  NAM_LinkMap_New <- NAM_LinkMap 
	  trainTableWindowSize <- TrainTableWindowSize
	  
	  nCyc <- 1
		        
	  trainGeno_backingFileName <- paste("trainTable_Cyc_",trainTableWindowSize,"_",no_QTL,"_",h2,"_",condition,"_",Rep,"_Cyc",nCyc,".bin",sep="")
      trainGeno_descriptorFileName <- paste("trainTable_Cyc_",trainTableWindowSize,"_",no_QTL,"_",h2,"_",condition,"_",Rep,"_Cyc",nCyc,".desc",sep="")
 						
	  trainGenoNewTablePreCycle <- deepcopy(as.big.matrix(trainGenoNewTable),backingfile=trainGeno_backingFileName,descriptorfile=trainGeno_descriptorFileName)
	  
	  #trainGenoNewTablePreCycle <- as.big.matrix(trainGenoNewTable)
      trainSimPhenoTablePreCycle <- trainSimPhenoValTable  
      trainSimGenoValTablePreCycle <- trainSimGenoValTable

    
    GIndStats <- gIndStats
	  AlleleConversionTable_Combined <- alleleConversionTable_Combined 
#########################################################################################################
	  nCyc <- 1
    print(paste("cycle_No",nCyc))
	  combinedTableCount <- 1
	  
####### Predict geno and pheno values ###################################################################

	  cycle1GenoTable <- generateMinus101GenoFormat(cycle1_Geno_Data,nFamilies,cycle1_nProgeny)

	  newCycle1GenoTable <- generateWeightedGenoTable(cycle1GenoTable,no_QTL)

	  if(selectionOnGeno==TRUE){
		  MarkerEffects <- unlist(PredictionModel_Geno[1])
	  }else if(selectionOnGeno==FALSE){
	    MarkerEffects <- unlist(PredictionModel_Pheno[1])
	  }

    Freq_List <- getFreq_BasePoln(cycle1GenoTable,MarkerEffects)
	  Freq <- Freq_List[[1]]
	  FavAllele <- Freq_List[[2]]
	  
	  if(GIndStats == TRUE){ 
###  Get population vector

	    populations <- rep(0,20*100)
		  init <- 1
		  for(pop in 1:20){

			  final <- pop*100
			  populations[init:final]<- rep(pop,100)
			  init <- final+1
		  }
		
      cycle1GenoTable_AlleleFormat <- getAlleleFormat(cycle1GenoTable,AlleleConversionTable_Combined)
 	  cycle1GenoTable_GInd <- df2genind(cycle1GenoTable_AlleleFormat,pop=populations,sep="")
		
	  cycle1GenoData_DF <- genind2df(cycle1GenoTable_GInd)
	  write.table(cycle1GenoData_DF,paste("PM_",modelType,"_",no_QTL,"_",h2,"_GenoDataInd_Rep",Rep,"_Cyc_",nCyc,sep=""),sep="\t")
	 }	

###### Predict geno and pheno values ###################################################################

    if(Weighted==TRUE && WghtMethod=="JM"){
			genoValues <- PredictRRBLUPPhenoValues_WGS_Jannink1(newCycle1GenoTable,PredictionModel_Geno,Freq)
			phenoValues <- PredictRRBLUPPhenoValues_WGS_Jannink1(newCycle1GenoTable,PredictionModel_Pheno,Freq)
        
		}else if (Weighted==TRUE && WghtMethod =="DW"){
		
		  cycleNumber <- nCyc
		  genoValues_PredList <- PredictRRBLUPPhenoValues_WGS_DWModel(newCycle1GenoTable,PredictionModel_Geno,Freq,alphaShape,betaShape,cycleNumber,timeHorizon)
			phenoValues_PredList <- PredictRRBLUPPhenoValues_WGS_DWModel(newCycle1GenoTable,PredictionModel_Pheno,Freq,alphaShape,betaShape,cycleNumber,timeHorizon)

			genoValues <- genoValues_PredList[[1]]
			phenoValues <- phenoValues_PredList[[1]]
			alphaPar_Vector[i] <- genoValues_PredList[[2]]
		
		}else if(Weighted==FALSE){
      genoValues  <- PredictRRBLUPPhenoValues(newCycle1GenoTable,PredictionModel_Geno)
      phenoValues <- PredictRRBLUPPhenoValues(newCycle1GenoTable,PredictionModel_Pheno)
    }

	
### Initiate Arrays and Variables  #######################################################################

	  GenoVal_NX_2k_3c <- matrix(rep(0,(nCrosses*nProgeny)*nCycles),nrow=(nCrosses*nProgeny),ncol=nCycles)
	  GenoVal_NX_N_3c <- matrix(rep(0,no_selected*nCycles),nrow=no_selected,ncol=nCycles)
	  GenoVal_Sim_NX_2k_3c <- matrix(rep(0,(nCrosses*nProgeny)*nCycles),nrow=(nCrosses*nProgeny),ncol=nCycles)

	  PhenoVal_NX_2k_3c <- matrix(rep(0,(nCrosses*nProgeny)*nCycles),nrow=(nCrosses*nProgeny),ncol=nCycles)
	  PhenoVal_NX_N_3c <- matrix(rep(0,no_selected*nCycles),nrow=no_selected,ncol=nCycles)
	  PhenoVal_Sim_NX_2k_3c <- matrix(rep(0,(nCrosses*nProgeny)*nCycles),nrow=(nCrosses*nProgeny),ncol=nCycles)

    attainedGenoValues <- rep(0,nCycles)
	 
	  selectedGenoIndividualIndices_List <- list()
    selectedPhenoIndividualIndices_List <- list()


######## Functions on Cycle 1 Geno Data ################################################################################

	if(selectionOnSimulated ==FALSE){
	
		if(selectionOnGeno==TRUE){
	
		  sortedGenoValues <- sort(genoValues,decreasing=TRUE,index.return=TRUE)
		  topNGenoValues <- sortedGenoValues[[1]][1:no_selected]
		  GenoTableIndices_topN <- sortedGenoValues[[2]][1:no_selected]
      topNPhenoValues <- phenoValues[GenoTableIndices_topN] 
		  topNGenoSimValues <- genoSimValues[GenoTableIndices_topN]
		  PhenoTableIndices_topN <- GenoTableIndices_topN
		}else if (selectionOnGeno==FALSE) {
		  	  
		  sortedPhenoValues <- sort(phenoValues,decreasing=TRUE,index.return=TRUE)
		  topNPhenoValues <- sortedPhenoValues[[1]][1:no_selected]
		  PhenoTableIndices_topN <- sortedPhenoValues[[2]][1:no_selected]
		  topNGenoValues <- genoValues[PhenoTableIndices_topN]
		  topNGenoSimValues <- genoSimValues[PhenoTableIndices_topN]
		  GenoTableIndices_topN <- PhenoTableIndices_topN
		} 
  
    }
	if(selectionOnSimulated ==TRUE){
	
		if(selectionOnGeno==TRUE){
		 
   		  sortedGenoValues <- sort(genoSimValues,decreasing=TRUE,index.return=TRUE)
		  topNGenoValues <- sortedGenoValues[[1]][1:no_selected]
		  GenoTableIndices_topN <- sortedGenoValues[[2]][1:no_selected]
		  topNPhenoValues <- phenoValues[GenoTableIndices_topN] 
		  topNGenoSimValues <- genoSimValues[GenoTableIndices_topN]
		  PhenoTableIndices_topN <- GenoTableIndices_topN
		
		}else if(selectionOnGeno==FALSE){

		  sortedPhenoValues <- sort(phenoSimValues,decreasing=TRUE,index.return=TRUE)
		  topNPhenoValues <- sortedPhenoValues[[1]][1:no_selected]
		  PhenoTableIndices_topN <- sortedPhenoValues[[2]][1:no_selected]
		  topNGenoValues <- genoValues[PhenoTableIndices_topN]
		  topNGenoSimValues <- genoSimValues[PhenoTableIndices_topN]
		  GenoTableIndices_topN <- PhenoTableIndices_topN
	    }
	}

##################################################################################################################
## Assign output variables for cycle 1

	  GenoVal_Sim_NX_2k_3c[,1] <- genoSimValues
	  GenoVal_NX_2k_3c[,1] <- genoValues
	  GenoVal_NX_N_3c[,1] <- topNGenoValues
    
	  attainedGenoValues[1] <- max(topNGenoSimValues)
	  
	  selectedGenoIndividualIndices <- GenoTableIndices_topN
      selectedPhenoIndividualIndices <- PhenoTableIndices_topN
	  
	  selectedGenoIndividualIndices_List[[nCyc]] <- selectedGenoIndividualIndices
	  selectedPhenoIndividualIndices_List[[nCyc]] <- selectedPhenoIndividualIndices
		

	  PhenoVal_Sim_NX_2k_3c[,1] <- phenoSimValues
	  PhenoVal_NX_2k_3c[,1] <- phenoValues
	  PhenoVal_NX_N_3c[,1] <- topNPhenoValues


      rm(cycle1GenoTable)
	  rm(newCycle1GenoTable)

############### Cycles 2*39 ###################################################################################
### extract genotype data of selected individuals
  
  for(nCyc in 2:nCycles){

   print(paste("cycleNo-",nCyc))
		
	 if(nCyc==2){

		if(selectionOnGeno == TRUE){
			selectedGenoData <- extractGenoData(selectedGenoIndividualIndices,cycle1_Geno_Data,no_selected)
		}else if(selectionOnGeno == FALSE){
			selectedGenoData <- extractGenoData(selectedPhenoIndividualIndices,cycle1_Geno_Data,no_selected)
		}

	}else if(nCyc>2){

		if(selectionOnGeno == TRUE){

			selectedGenoData <- extractGenoData(selectedGenoIndividualIndices,Cycle_Progeny_F5,no_selected)
		}else if(selectionOnGeno == FALSE){
			selectedGenoData <- extractGenoData(selectedPhenoIndividualIndices,Cycle_Progeny_F5,no_selected)
		}

	}
	
	Cycle_Progeny_F5 <- getF5RILs_BD(BD,selectedGenoData,no_selected,nProgeny,nMarkers,NAM_LinkMap_New)
	
	
########################
##  nextGenGenoTable <- generate012GenoFormat(Cycle_Progeny_F5,no_selected)

    nextGenGenoTable <- generateMinus101GenoFormat(Cycle_Progeny_F5,nCrosses,nProgeny)

    Freq <- getFreq(nextGenGenoTable,FavAllele)
    genoSimValues <- simulateGenotypicValues(nextGenGenoTable,no_QTL)
    phenoSimValues <- simulatePhenotypicValues(nextGenGenoTable,varE,no_QTL)

    newNextGenGenoTable <- generateWeightedGenoTable(nextGenGenoTable,no_QTL)

 if(nCyc%%updateFrequency==0 && modelUpdate ==TRUE){
 
    set.seed(25+Rep+nCyc)
## Extract training and test data for model build

 
    IndividualNames <-rep(0,(nCrosses*nProgeny))

   IndividualNames <- paste("Ind",c(1:(nCrosses*nProgeny)),sep="")
##################################################################################################

    names(phenoSimValues)<-IndividualNames

    Mean_Fixed<- rep(1,nIndividuals)

    phenotypicValuesSimTable<- cbind(phenoSimValues,Mean_Fixed)


#### Table for Simulated Genotypic Values ####################################

    names(genoSimValues)<-IndividualNames

    Mean_Fixed<- rep(1,nIndividuals)

    genotypicValuesSimTable <- cbind(genoSimValues,Mean_Fixed)


### Model Retrain 

	if(modelRetrain==TRUE){ 
        PredDefined <- FALSE

        while(PredDefined ==FALSE){

		indices<-c(1:nIndividuals)

		trainIndices<-sample(c(1:nIndividuals),(0.8*nIndividuals))
		testIndices<- indices[which(!indices %in% trainIndices)]

#### Training Set table for Genotype and Simulated Phenotype ###################

		# trainGenoTable <- genoTable_Mod[trainIndices,]
		trainGenoNewTable <- newNextGenGenoTable[trainIndices,]
		trainSimPhenoTable <- phenotypicValuesSimTable[trainIndices,]
		trainSimGenoTable <-  genotypicValuesSimTable[trainIndices,]

#### Validation Set table for Genotype and Simulated Phenotype ###################

		# testGenoTable<- genoTable_Mod[testIndices,]
		testGenoNewTable <- newNextGenGenoTable[testIndices,]
		testSimPhenoTable <- phenotypicValuesSimTable[testIndices,]
		testSimGenoTable <-  genotypicValuesSimTable[testIndices,]

	    print(length(trainSimGenoTable[,1]))
		print(sd(trainSimGenoTable[,1]))	
		print(sd(trainSimPhenoTable[,1]))
        print(i%%updateFrequency) 
		print(nCyc%%updateFrequency)
        print(summary(trainSimPhenoTable[,1]))		
############### Change RRBLUP to Bayes
      if(!is.null(sd(trainSimGenoTable[,1])) && !is.null(sd(trainSimPhenoTable[,1])) && !is.na(sd(trainSimGenoTable[,1])) && !is.na(sd(trainSimPhenoTable[,1]))){
	    if(sd(trainSimGenoTable[,1])!=0 && sd(trainSimPhenoTable[,1])!=0 && nCyc%%retrainFrequency==0){

			if(modelType=="RRBLUP"){
				PredictionModel_Geno <- (buildRRBLUPModel(trainGenoNewTable,trainSimGenoTable,testGenoNewTable,testSimGenoTable))
				PredictionModel_Pheno <-(buildRRBLUPModel(trainGenoNewTable,trainSimPhenoTable,testGenoNewTable,testSimPhenoTable))
			}else if(modelType=="RRBLUP_REML"){
				PredictionModel_Geno <- (buildRRBLUPModel_REML(trainGenoNewTable,trainSimGenoTable,testGenoNewTable,testSimGenoTable))
				PredictionModel_Pheno <-(buildRRBLUPModel_REML(trainGenoNewTable,trainSimPhenoTable,testGenoNewTable,testSimPhenoTable))
			}
	    }
	  }
	
        if(length(levels(factor(PredictionModel_Pheno[[1]])))>1 && mean(PredictionModel_Pheno[[1]])>0) {
                         PredDefined <- TRUE }      
        }
   }

	gc()	
		
	if(modelUpdate ==TRUE && updateType=="FullSet"){

        indices<-c(1:nIndividuals)

        trainIndices<-sample(c(1:nIndividuals),(0.8*nIndividuals))

        testIndices<- indices[which(!indices %in% trainIndices)]

#### Training Set table for Genotype and Simulated Phenotype ###################

        trainGenoNewTable <- newNextGenGenoTable[trainIndices,]

        trainSimPhenoTable <- phenotypicValuesSimTable[trainIndices,]

        trainSimGenoValTable <-  genotypicValuesSimTable[trainIndices,]

#### Validation Set table for Genotype and Simulated Phenotype ###################

        testGenoNewTable <- newNextGenGenoTable[testIndices,]

        testSimPhenoTable <- phenotypicValuesSimTable[testIndices,]

        testSimGenoValTable <-  genotypicValuesSimTable[testIndices,]


##### Build RRBLUP prediction model every cycle


        trainGenoNewTableComb <- as.big.matrix(rbind(bigmemory::as.matrix(trainGenoNewTablePreCycle),as.matrix(trainGenoNewTable)))

        trainSimGenoValTableComb <- rbind((trainSimGenoValTablePreCycle),trainSimGenoValTable)

        trainSimPhenoValTableComb <- rbind((trainSimPhenoTablePreCycle),trainSimPhenoTable)

        print(dim(trainGenoNewTableComb))

        if(!is.null(sd(trainSimGenoValTableComb[,1])) && !is.null(sd(trainSimPhenoValTableComb[,1])) && !is.na(sd(trainSimGenoValTableComb[,1])) && !is.na(sd(trainSimPhenoValTableComb[,1]))){
             if(sd(trainSimGenoValTableComb[,1])!=0 && sd(trainSimPhenoValTableComb[,1])!=0 && nCyc%%updateFrequency==0){ 


	
           if(modelType=="RRBLUP"){
                        PredictionModel_Geno <- (buildRRBLUPModel(bigmemory::as.matrix(trainGenoNewTableComb),trainSimGenoValTableComb,testGenoNewTable,testSimGenoValTable))
                        PredictionModel_Pheno <-(buildRRBLUPModel(bigmemory::as.matrix(trainGenoNewTableComb),trainSimPhenoTable,testGenoNewTable,testSimPhenoTable))
           }else if(modelType=="RRBLUP_REML"){

                        PredictionModel_Geno <- buildRRBLUPModel_REML(bigmemory::as.matrix(trainGenoNewTableComb), trainSimGenoValTableComb,as.matrix(testGenoNewTable),testSimGenoValTable)
                        PredictionModel_Pheno <- buildRRBLUPModel_REML(bigmemory::as.matrix(trainGenoNewTableComb), trainSimPhenoValTableComb,as.matrix(testGenoNewTable),testSimPhenoTable)
           }

                trainGenoNewTablePreCycle <-  trainGenoNewTableComb

                trainSimPhenoTablePreCycle <-   (trainSimPhenoValTableComb)

                trainSimGenoValTablePreCycle <- (trainSimGenoValTableComb)


                rm(trainGenoNewTableComb)

                rm(trainSimGenoValTableComb)

                rm(trainSimPhenoValTableComb)

                gc()

                }

             }
    }

	if(modelUpdate ==TRUE && updateType=="TrainingCycleWindow"){

			
		PredictionModel_Geno_PreCycle <- PredictionModel_Geno
		PredictionModel_Pheno_PreCycle <- PredictionModel_Pheno
        
        indices<-c(1:nIndividuals)

        trainIndices<-sample(c(1:nIndividuals),(0.8*nIndividuals))

        testIndices<- indices[which(!indices %in% trainIndices)]
		
	    nTrainIndices <- length(trainIndices)

#### Training Set table for Genotype and Simulated Phenotype ###################

        trainGenoNewTable <- newNextGenGenoTable[trainIndices,]

        trainSimPhenoTable <- phenotypicValuesSimTable[trainIndices,]

        trainSimGenoValTable <-  genotypicValuesSimTable[trainIndices,]

#### Validation Set table for Genotype and Simulated Phenotype ###################
        testGenoNewTable <- newNextGenGenoTable[testIndices,]

        testSimPhenoTable <- phenotypicValuesSimTable[testIndices,]

        testSimGenoValTable <-  genotypicValuesSimTable[testIndices,]

        trainGenoNewTablePreCycle <- attach.big.matrix(trainGeno_descriptorFileName)

##### Build RRBLUP prediction model every cycle
        

        trainGenoNewTableComb <- (rbind(bigmemory::as.matrix(trainGenoNewTablePreCycle),as.matrix(trainGenoNewTable)))
        trainSimGenoValTableComb <- rbind((trainSimGenoValTablePreCycle),trainSimGenoValTable)
        trainSimPhenoValTableComb <- rbind((trainSimPhenoTablePreCycle),trainSimPhenoTable)
        print(dim(trainGenoNewTableComb))
	      Avg_GenoSim_Prev <- mean(trainSimGenoValTableComb[,1])
	      Avg_GenoSim_Current <- mean(trainSimGenoValTable[,1])
	
	     change_GenoSim <- Avg_GenoSim_Prev - Avg_GenoSim_Current
	     combinedTableCount <- combinedTableCount+1
       print(change_GenoSim)
    	   
	### 
      
        if(!is.null(sd(trainSimGenoValTableComb[,1])) && !is.null(sd(trainSimPhenoValTableComb[,1])) && !is.na(sd(trainSimGenoValTableComb[,1])) && !is.na(sd(trainSimPhenoValTableComb[,1]))){
           if(sd(trainSimGenoValTableComb[,1])!=0 && sd(trainSimPhenoValTableComb[,1])!=0 && change_GenoSim !=0){ 

	### 
             if(modelType=="RRBLUP"){

                        PredictionModel_Geno <- (buildRRBLUPModel((trainGenoNewTableComb),trainSimGenoValTableComb,as.matrix(testGenoNewTable),testSimGenoValTable))

                        PredictionModel_Pheno <-(buildRRBLUPModel((trainGenoNewTableComb),trainSimPhenoValTableComb,as.matrix(testGenoNewTable),testSimPhenoTable))

           }else if(modelType=="RRBLUP_REML"){

                      					
						PredictionModel_Geno <- tryCatch(buildRRBLUPModel_REML((trainGenoNewTableComb),trainSimGenoValTableComb[,1],as.matrix(testGenoNewTable),testSimGenoValTable[,1]),error=function(e) print(paste("BuildRR-Error -",nCyc,e)))

                        PredictionModel_Pheno <- tryCatch(buildRRBLUPModel_REML((trainGenoNewTableComb),trainSimPhenoValTableComb[,1],as.matrix(testGenoNewTable),testSimPhenoTable[,1]),error=function(e) print(paste("BuildRR-Error -",nCyc,e)))

           }

	     }

           print(sd(PredictionModel_Pheno[[1]]))
           print(sd(PredictionModel_Geno[[1]])) 
           print(length(levels(factor(PredictionModel_Pheno[[1]]))))
           print(length(levels(factor(PredictionModel_Geno[[1]]))))	 

           genoValues <- PredictRRBLUPPhenoValues(newNextGenGenoTable,PredictionModel_Geno)
           phenoValues <- PredictRRBLUPPhenoValues(newNextGenGenoTable,PredictionModel_Pheno)
           print(sd(phenoValues))
           print(sd(genoValues))

 
           if(length(levels(factor(PredictionModel_Pheno[[1]]))) ==1 || length(levels(factor(PredictionModel_Geno[[1]]))) ==1 || (is.na(sd(phenoValues)) || sd(phenoValues)==0) || (is.na(sd(genoValues))) || sd(genoValues)==0 || change_GenoSim==0){

                         PredictionModel_Pheno <- PredictionModel_Pheno_PreCycle
                         PredictionModel_Geno <- PredictionModel_Geno_PreCycle

           }
	 }
		
		trainGeno_backingFileName <- paste("trainTable_Cyc_",trainTableWindowSize,"_",no_QTL,"_",h2,"_",condition,"_",Rep,"_Cyc",nCyc,".bin",sep="")
     
	    trainGeno_descriptorFileName <- paste("trainTable_Cyc_",trainTableWindowSize,"_",no_QTL,"_",h2,"_",condition,"_",Rep,"_Cyc",nCyc,".desc",sep="")
	 	 
		
		if(combinedTableCount < trainTableWindowSize){
			
			   	trainGenoNewTablePreCycle <- deepcopy(as.big.matrix(trainGenoNewTableComb),backingfile=trainGeno_backingFileName,descriptorfile=trainGeno_descriptorFileName)
	 
                trainSimPhenoTablePreCycle <-   (trainSimPhenoValTableComb)

                trainSimGenoValTablePreCycle <- (trainSimGenoValTableComb)
					
		 }else if(combinedTableCount >= trainTableWindowSize){
			
			  	trainGenoNewTablePreCycle <- deepcopy(as.big.matrix(trainGenoNewTableComb[-c(1:nTrainIndices),]),backingfile=trainGeno_backingFileName,descriptorfile=trainGeno_descriptorFileName)

                trainSimPhenoTablePreCycle <-   (trainSimPhenoValTableComb[-c(1:nTrainIndices),])

                trainSimGenoValTablePreCycle <- (trainSimGenoValTableComb[-c(1:nTrainIndices),])
				
				combinedTableCount <- (trainTableWindowSize-1)
					
		  }
		  
		   rm(trainGenoNewTableComb)

           rm(trainSimGenoValTableComb)

           rm(trainSimPhenoValTableComb)
	
    gc()

   }
 }
#################get predicted geno and pheno values with JM and unweighted GP models
  	
	
	if (Weighted==TRUE && WghtMethod =="JM"){
		genoValues <- PredictRRBLUPPhenoValues_WGS_Jannink1(newNextGenGenoTable,PredictionModel_Geno,Freq)
		phenoValues <- PredictRRBLUPPhenoValues_WGS_Jannink1(newNextGenGenoTable,PredictionModel_Pheno,Freq)
    } else if(Weighted ==TRUE && WghtMethod =="DW"){
	
		cycleNumber <- nCyc
		genoValues_PredList <- PredictRRBLUPPhenoValues_WGS_DWModel(newNextGenGenoTable,PredictionModel_Geno,Freq,alphaShape,betaShape,cycleNumber,timeHorizon)
		phenoValues_PredList <- PredictRRBLUPPhenoValues_WGS_DWModel(newNextGenGenoTable,PredictionModel_Pheno,Freq,alphaShape,betaShape,cycleNumber,timeHorizon)

		genoValues <- genoValues_PredList[[1]]
		phenoValues <- phenoValues_PredList[[1]]

		alphaPar_Vector[i] <- genoValues_PredList[[2]]
		
	} else if(Weighted==FALSE){
		genoValues <- PredictRRBLUPPhenoValues(newNextGenGenoTable,PredictionModel_Geno)
		phenoValues <- PredictRRBLUPPhenoValues(newNextGenGenoTable,PredictionModel_Pheno)
    }
	
############# 
  if(GIndStats == TRUE){
###  Get population vector

		populations <- rep(0,20*100)
		init <- 1
		for(pop in 1:20){

			final <- pop*100
			populations[init:final]<- rep(pop,100)
			init <- final+1
		}
		
### Get GInd objects 	
	
		print(length(populations))
		nextGenGenoTable_AlleleFormat <- getAlleleFormat(nextGenGenoTable,AlleleConversionTable_Combined)
		nextGenGenoTable_GInd <- df2genind(as.data.frame(nextGenGenoTable_AlleleFormat),pop=populations,sep="")
		nextGenGenoData_DF <- genind2df(nextGenGenoTable_GInd)
		write.table(nextGenGenoData_DF,paste("PM_",modelType,"trainTable_Cyc_",trainTableWindowSize,"_",no_QTL,"_",h2,"_GenoDataInd_Rep",Rep,"_Cyc_",nCyc,sep=""),sep="\t")
	
	}
#############
    print(paste("GenoVal_Length",length(genoValues)))
    print(paste("PhenoVal_Length",length(phenoValues)))

    GenoVal_Sim_NX_2k_3c[,nCyc] <- genoSimValues
    GenoVal_NX_2k_3c[,nCyc] <- genoValues
   
    PhenoVal_Sim_NX_2k_3c[,nCyc]<- phenoSimValues
    PhenoVal_NX_2k_3c[,nCyc] <- phenoValues
   

### Selection based on simulated or predicted values

    if(selectionOnSimulated ==TRUE){

		genoSelectionTable <- ApplySelection(genoSimValues,no_selected)
		phenoSelectionTable <- ApplySelection(phenoSimValues,no_selected)


		if(selectionOnGeno==TRUE){

			selectedGenoIndividualIndices <- as.vector(genoSelectionTable[,2])
			genoSimSelectedValues <- genoSimValues[selectedGenoIndividualIndices]
			phenoSimSelectedValues <- phenoSimValues[selectedGenoIndividualIndices]

		}else if(selectionOnGeno==FALSE){

			selectedPhenoIndividualIndices <- as.vector(phenoSelectionTable[,2])
			genoSimSelectedValues <- genoSimValues[selectedPhenoIndividualIndices]
			phenoSimSelectedValues <- phenoSimValues[selectedPhenoIndividualIndices]
		}

		GenoVal_NX_N_3c[,nCyc] <- genoSimSelectedValues
	    PhenoVal_NX_N_3c[,nCyc] <- phenoSimSelectedValues
		attainedGenoValues[nCyc] <- max(genoSimSelectedValues)
		selectedGenoIndividualIndices_List[[nCyc]] <- selectedGenoIndividualIndices
	    selectedPhenoIndividualIndices_List[[nCyc]] <- selectedPhenoIndividualIndices
		


	}
    if(selectionOnSimulated==FALSE){

		genoSelectionTable <- ApplySelection(genoValues,no_selected)
		phenoSelectionTable <- ApplySelection(phenoValues,no_selected)

    	print(paste("genoSelectionTable-dim-",dim(genoSelectionTable)))
		print(paste("phenoSelectionTable-dim-",dim(phenoSelectionTable)))
		print(paste("selectedGenoIndividualIndices",is.na(phenoSelectionTable[,2])))

		if(selectionOnGeno==TRUE){

			selectedGenoIndividualIndices <- as.vector(genoSelectionTable[,2])
			genoSelectedValues <- genoValues[selectedGenoIndividualIndices]
			phenoSelectedValues <- phenoValues[selectedGenoIndividualIndices]
		  genoSimSelectedValues <- genoSimValues[selectedPhenoIndividualIndices]
		
		}else if(selectionOnGeno==FALSE){

			selectedPhenoIndividualIndices <- as.vector(phenoSelectionTable[,2])
			genoSelectedValues <- genoValues[selectedPhenoIndividualIndices]
			phenoSelectedValues <- phenoValues[selectedPhenoIndividualIndices]
		    genoSimSelectedValues <- genoSimValues[selectedPhenoIndividualIndices]
		}
		    GenoVal_NX_N_3c[,nCyc]<- genoSelectedValues
	        PhenoVal_NX_N_3c[,nCyc]<- phenoSelectedValues
        	attainedGenoValues[nCyc] <- max(genoSimSelectedValues)
			
			selectedGenoIndividualIndices_List[[nCyc]] <- selectedGenoIndividualIndices
	        selectedPhenoIndividualIndices_List[[nCyc]] <- selectedPhenoIndividualIndices
		

	}

   	rm(nextGenGenoTable)
	rm(newNextGenGenoTable)
    gc()
  }

  simResults_List <- list(GenoVal_Sim_NX_2k_3c,GenoVal_NX_2k_3c,GenoVal_NX_N_3c,PhenoVal_Sim_NX_2k_3c,PhenoVal_NX_2k_3c,PhenoVal_NX_N_3c,attainedGenoValues,selectedGenoIndividualIndices_List,selectedPhenoIndividualIndices_List)

  return(simResults_List)

}
	
### 

runSimulations40X_Drift <- function(model_Sol_Pheno,model_Sol_Geno,F5_Progeny_List,genoValSimList,phenoValSimList,varEList,noQTLs,H2,numberSelected,noCrosses,no_Progeny,noCycles,selectionOnGenoValues,selectionOnSimulatedValues,ModelRetrain,RetrainFrequency,ModelUpdate,UpdateType,UpdateFrequency,i,k,ModelType,weighted,weightingMethod,AlphaShape,BetaShape,TimeHorizon,trainGenoNewTable,trainSimGenoValTable,trainSimPhenoValTable,TrainTableWindowSize,BreedingDesign,NAM_LinkMap,gIndStats,alleleConversionTable_Combined){


### Assign Variables

	  h2 <- H2
	  no_QTL <-noQTLs

	  no_selected <- numberSelected
	  nCrosses <- noCrosses
	  nProgeny <- no_Progeny
	  nCycles <- noCycles
	  selectionOnGeno <- selectionOnGenoValues
	  selectionOnSimulated <- selectionOnSimulatedValues

####################################################

	  PredictionModel_Pheno <- model_Sol_Pheno
	  PredictionModel_Geno <- model_Sol_Geno

	  cycle1_Geno_Data <- F5_Progeny_List
	  genoSimValues <- genoValSimList
	  phenoSimValues <- phenoValSimList

	  varE <- varEList

	  condition <- i
	  Rep <- k
	  modelType <- ModelType
	  modelRetrain <- ModelRetrain
	  updateFrequency <- UpdateFrequency
	  retrainFrequency <- RetrainFrequency
	  updateType <- UpdateType
      modelUpdate <- ModelUpdate

	  nFamilies <- 20
      cycle1_nProgeny <- 100
	  nMarkers <- 4289
      nIndividuals <- 2000

      Weighted <- weighted
	  WghtMethod <- weightingMethod  
	  alphaShape <- AlphaShape
	  betaShape <- BetaShape
	  timeHorizon <- TimeHorizon
	  alphaPar_Vector <- rep(0,nCycles)
      BD <- BreedingDesign
         
	  NAM_LinkMap_New <- NAM_LinkMap 
	  trainTableWindowSize <- TrainTableWindowSize
	  
	  nCyc <- 1
		        
	  trainGeno_backingFileName <- paste("trainTable_Cyc_",trainTableWindowSize,"_",no_QTL,"_",h2,"_",condition,"_",Rep,"_Cyc",nCyc,".bin",sep="")
      trainGeno_descriptorFileName <- paste("trainTable_Cyc_",trainTableWindowSize,"_",no_QTL,"_",h2,"_",condition,"_",Rep,"_Cyc",nCyc,".desc",sep="")
 							
	  trainGenoNewTablePreCycle <- deepcopy(as.big.matrix(trainGenoNewTable),backingfile=trainGeno_backingFileName,descriptorfile=trainGeno_descriptorFileName)
   
      trainSimPhenoTablePreCycle <- trainSimPhenoValTable  
      trainSimGenoValTablePreCycle <- trainSimGenoValTable

    
      GIndStats <- gIndStats
	  AlleleConversionTable_Combined <- alleleConversionTable_Combined 
#########################################################################################################
	  nCyc <- 1
      print(paste("cycle_No",nCyc))
	  combinedTableCount <- 1
	  
####### Predict geno and pheno values ###################################################################

	  cycle1GenoTable <- generateMinus101GenoFormat(cycle1_Geno_Data,nFamilies,cycle1_nProgeny)

	  newCycle1GenoTable <- generateWeightedGenoTable(cycle1GenoTable,no_QTL)

	  if(selectionOnGeno==TRUE){
		MarkerEffects <- unlist(PredictionModel_Geno[1])
	  }else if(selectionOnGeno==FALSE){
	    	MarkerEffects <- unlist(PredictionModel_Pheno[1])
	  }

      Freq_List <- getFreq_BasePoln(cycle1GenoTable,MarkerEffects)
	  Freq <- Freq_List[[1]]
	  FavAllele <- Freq_List[[2]]
	  
	  if(GIndStats == TRUE){ 
###  Get population vector

	    populations <- rep(0,20*100)
		init <- 1
		for(pop in 1:20){

			final <- pop*100
			populations[init:final]<- rep(pop,100)
			init <- final+1
		}
		
      cycle1GenoTable_AlleleFormat <- getAlleleFormat(cycle1GenoTable,AlleleConversionTable_Combined)
 	  cycle1GenoTable_GInd <- df2genind(cycle1GenoTable_AlleleFormat,pop=populations,sep="")
		
	  cycle1GenoData_DF <- genind2df(cycle1GenoTable_GInd)
	  write.table(cycle1GenoData_DF,paste("PM_",modelType,"_",no_QTL,"_",h2,"_GenoDataInd_Rep",Rep,"_Cyc_",nCyc,sep=""),sep="\t")
	}	

###### Predict geno and pheno values ###################################################################

        if(Weighted==TRUE && WghtMethod=="JM"){
			genoValues <- PredictRRBLUPPhenoValues_WGS_Jannink1(newCycle1GenoTable,PredictionModel_Geno,Freq)
			phenoValues <- PredictRRBLUPPhenoValues_WGS_Jannink1(newCycle1GenoTable,PredictionModel_Pheno,Freq)
        
		}else if (Weighted==TRUE && WghtMethod =="DW"){
		
		    cycleNumber <- nCyc
		    genoValues_PredList <- PredictRRBLUPPhenoValues_WGS_DWModel(newCycle1GenoTable,PredictionModel_Geno,Freq,alphaShape,betaShape,cycleNumber,timeHorizon)
			phenoValues_PredList <- PredictRRBLUPPhenoValues_WGS_DWModel(newCycle1GenoTable,PredictionModel_Pheno,Freq,alphaShape,betaShape,cycleNumber,timeHorizon)

			genoValues <- genoValues_PredList[[1]]
			phenoValues <- phenoValues_PredList[[1]]
			alphaPar_Vector[i] <- genoValues_PredList[[2]]
		
		}else if(Weighted==FALSE){
            genoValues  <- PredictRRBLUPPhenoValues(newCycle1GenoTable,PredictionModel_Geno)
            phenoValues <- PredictRRBLUPPhenoValues(newCycle1GenoTable,PredictionModel_Pheno)
        }

	
### Initiate Arrays and Variables  #######################################################################

	  GenoVal_NX_2k_3c <- matrix(rep(0,(nCrosses*nProgeny)*nCycles),nrow=(nCrosses*nProgeny),ncol=nCycles)
	  GenoVal_NX_N_3c <- matrix(rep(0,no_selected*nCycles),nrow=no_selected,ncol=nCycles)
	  GenoVal_Sim_NX_2k_3c <- matrix(rep(0,(nCrosses*nProgeny)*nCycles),nrow=(nCrosses*nProgeny),ncol=nCycles)

	  PhenoVal_NX_2k_3c <- matrix(rep(0,(nCrosses*nProgeny)*nCycles),nrow=(nCrosses*nProgeny),ncol=nCycles)
	  PhenoVal_NX_N_3c <- matrix(rep(0,no_selected*nCycles),nrow=no_selected,ncol=nCycles)
	  PhenoVal_Sim_NX_2k_3c <- matrix(rep(0,(nCrosses*nProgeny)*nCycles),nrow=(nCrosses*nProgeny),ncol=nCycles)

      attainedGenoValues <- rep(0,nCycles)
	 
	  selectedGenoIndividualIndices_List <- list()
      selectedPhenoIndividualIndices_List <- list()


######## Functions on Cycle 1 Geno Data ################################################################################

	if(selectionOnSimulated ==FALSE){
	
		if(selectionOnGeno==TRUE){
	
		  sortedGenoValues <- sort(genoValues,decreasing=TRUE,index.return=TRUE)
		  topNGenoValues <- sortedGenoValues[[1]][1:no_selected]
		  GenoTableIndices_topN <- sortedGenoValues[[2]][1:no_selected]
          topNPhenoValues <- phenoValues[GenoTableIndices_topN] 
		  topNGenoSimValues <- genoSimValues[GenoTableIndices_topN]
		  PhenoTableIndices_topN <- GenoTableIndices_topN
		}else if (selectionOnGeno==FALSE) {
		  	  
		  sortedPhenoValues <- sort(phenoValues,decreasing=TRUE,index.return=TRUE)
		  topNPhenoValues <- sortedPhenoValues[[1]][1:no_selected]
		  PhenoTableIndices_topN <- sortedPhenoValues[[2]][1:no_selected]
		  topNGenoValues <- genoValues[PhenoTableIndices_topN]
		  topNGenoSimValues <- genoSimValues[PhenoTableIndices_topN]
		  GenoTableIndices_topN <- PhenoTableIndices_topN
		} 
  
    }
	if(selectionOnSimulated ==TRUE){
	
		if(selectionOnGeno==TRUE){
		 
   		  sortedGenoValues <- sort(genoSimValues,decreasing=TRUE,index.return=TRUE)
		  topNGenoValues <- sortedGenoValues[[1]][1:no_selected]
		  GenoTableIndices_topN <- sortedGenoValues[[2]][1:no_selected]
		  topNPhenoValues <- phenoValues[GenoTableIndices_topN] 
		  topNGenoSimValues <- genoSimValues[GenoTableIndices_topN]
		  PhenoTableIndices_topN <- GenoTableIndices_topN
		
		}else if(selectionOnGeno==FALSE){

		  sortedPhenoValues <- sort(phenoSimValues,decreasing=TRUE,index.return=TRUE)
		  topNPhenoValues <- sortedPhenoValues[[1]][1:no_selected]
		  PhenoTableIndices_topN <- sortedPhenoValues[[2]][1:no_selected]
		  topNGenoValues <- genoValues[PhenoTableIndices_topN]
		  topNGenoSimValues <- genoSimValues[PhenoTableIndices_topN]
		  GenoTableIndices_topN <- PhenoTableIndices_topN
	    }
	}


##################################################################################################################
## Assign output variables for cycle 1

	  GenoVal_Sim_NX_2k_3c[,1] <- genoSimValues
	  GenoVal_NX_2k_3c[,1] <- genoValues
	  GenoVal_NX_N_3c[,1] <- topNGenoValues
    
	  attainedGenoValues[1] <- max(topNGenoSimValues)
	  
	  selectedGenoIndividualIndices <- GenoTableIndices_topN
      selectedPhenoIndividualIndices <- PhenoTableIndices_topN
	  
	  selectedGenoIndividualIndices_List[[nCyc]] <- selectedGenoIndividualIndices
	  selectedPhenoIndividualIndices_List[[nCyc]] <- selectedPhenoIndividualIndices
		

	  PhenoVal_Sim_NX_2k_3c[,1] <- phenoSimValues
	  PhenoVal_NX_2k_3c[,1] <- phenoValues
	  PhenoVal_NX_N_3c[,1] <- topNPhenoValues
      
	  initIndex <- 1
	  finalIndex <- cycle1_nProgeny
	  
	  cycle1_Geno_Data_PM <- array(0,c(nFamilies*cycle1_nProgeny,nMarkers,2))

      for(nfam in 1:nFamilies){ 
	  
	    
		cycle1_Geno_Data_PM[initIndex:finalIndex,,] <- cycle1_Geno_Data[nfam,,,1:100]
		initIndex <- finalIndex +1 
		finalIndex <- finalIndex + cycle1_nProgeny
		
	  }
	   
	
      rm(cycle1GenoTable)
	  rm(newCycle1GenoTable)

############### Cycles 2*39 ###################################################################################
### extract genotype data of selected individuals
   
  for(nCyc in 2:nCycles){
  
	

    print(paste("cycleNo-",nCyc))
	
	if(selectionOnGeno == TRUE){
			selectedGenoData <- cycle1_Geno_Data_PM[selectedGenoIndividualIndices,,]
	}else if(selectionOnGeno == FALSE){
			selectedGenoData <- cycle1_Geno_Data_PM[selectedPhenoIndividualIndices,,] 
	}
	

	Cycle_Progeny_F5 <- getF5RILs_BD(BD,selectedGenoData,no_selected,nProgeny,nMarkers,NAM_LinkMap_New)
	
	
########################
##  nextGenGenoTable <- generate012GenoFormat(Cycle_Progeny_F5,no_selected)

    nextGenGenoTable <- generateMinus101GenoFormat(Cycle_Progeny_F5,nCrosses,nProgeny)

    Freq <- getFreq(nextGenGenoTable,FavAllele)
    genoSimValues <- simulateGenotypicValues(nextGenGenoTable,no_QTL)
    phenoSimValues <- simulatePhenotypicValues(nextGenGenoTable,varE,no_QTL)

    newNextGenGenoTable <- generateWeightedGenoTable(nextGenGenoTable,no_QTL)

 if(nCyc%%updateFrequency==0 && modelUpdate ==TRUE){
 
    set.seed(25+Rep+nCyc)
## Extract training and test data for model build

 
    IndividualNames <-rep(0,(nCrosses*nProgeny))

   IndividualNames <- paste("Ind",c(1:(nCrosses*nProgeny)),sep="")
##################################################################################################

    names(phenoSimValues)<-IndividualNames

    Mean_Fixed<- rep(1,nIndividuals)

    phenotypicValuesSimTable<- cbind(phenoSimValues,Mean_Fixed)


#### Table for Simulated Genotypic Values ####################################

    names(genoSimValues)<-IndividualNames

    Mean_Fixed<- rep(1,nIndividuals)

    genotypicValuesSimTable <- cbind(genoSimValues,Mean_Fixed)


### Model Retrain 

	if(modelRetrain==TRUE){ 
        PredDefined <- FALSE

        while(PredDefined ==FALSE){

		indices<-c(1:nIndividuals)

		trainIndices<-sample(c(1:nIndividuals),(0.8*nIndividuals))
		testIndices<- indices[which(!indices %in% trainIndices)]

#### Training Set table for Genotype and Simulated Phenotype ###################

		# trainGenoTable <- genoTable_Mod[trainIndices,]
		trainGenoNewTable <- newNextGenGenoTable[trainIndices,]
		trainSimPhenoTable <- phenotypicValuesSimTable[trainIndices,]
		trainSimGenoTable <-  genotypicValuesSimTable[trainIndices,]

#### Validation Set table for Genotype and Simulated Phenotype ###################

		# testGenoTable<- genoTable_Mod[testIndices,]
		testGenoNewTable <- newNextGenGenoTable[testIndices,]
		testSimPhenoTable <- phenotypicValuesSimTable[testIndices,]
		testSimGenoTable <-  genotypicValuesSimTable[testIndices,]

	    print(length(trainSimGenoTable[,1]))
		print(sd(trainSimGenoTable[,1]))	
		print(sd(trainSimPhenoTable[,1]))
        print(i%%updateFrequency) 
		print(nCyc%%updateFrequency)
        print(summary(trainSimPhenoTable[,1]))		
############### Change RRBLUP to Bayes
      if(!is.null(sd(trainSimGenoTable[,1])) && !is.null(sd(trainSimPhenoTable[,1])) && !is.na(sd(trainSimGenoTable[,1])) && !is.na(sd(trainSimPhenoTable[,1]))){
	    if(sd(trainSimGenoTable[,1])!=0 && sd(trainSimPhenoTable[,1])!=0 && nCyc%%retrainFrequency==0){

			if(modelType=="RRBLUP"){
				PredictionModel_Geno <- (buildRRBLUPModel(trainGenoNewTable,trainSimGenoTable,testGenoNewTable,testSimGenoTable))
				PredictionModel_Pheno <-(buildRRBLUPModel(trainGenoNewTable,trainSimPhenoTable,testGenoNewTable,testSimPhenoTable))
			}else if(modelType=="RRBLUP_REML"){
				PredictionModel_Geno <- (buildRRBLUPModel_REML(trainGenoNewTable,trainSimGenoTable,testGenoNewTable,testSimGenoTable))
				PredictionModel_Pheno <-(buildRRBLUPModel_REML(trainGenoNewTable,trainSimPhenoTable,testGenoNewTable,testSimPhenoTable))
			}
	    }
	  }
	
        if(length(levels(factor(PredictionModel_Pheno[[1]])))>1 && mean(PredictionModel_Pheno[[1]])>0) {
                         PredDefined <- TRUE }      
        }
   }

	gc()	
		
	if(modelUpdate ==TRUE && updateType=="FullSet"){

        indices<-c(1:nIndividuals)

        trainIndices<-sample(c(1:nIndividuals),(0.8*nIndividuals))

        testIndices<- indices[which(!indices %in% trainIndices)]

#### Training Set table for Genotype and Simulated Phenotype ###################

        trainGenoNewTable <- newNextGenGenoTable[trainIndices,]

        trainSimPhenoTable <- phenotypicValuesSimTable[trainIndices,]

        trainSimGenoValTable <-  genotypicValuesSimTable[trainIndices,]

#### Validation Set table for Genotype and Simulated Phenotype ###################

        testGenoNewTable <- newNextGenGenoTable[testIndices,]

        testSimPhenoTable <- phenotypicValuesSimTable[testIndices,]

        testSimGenoValTable <-  genotypicValuesSimTable[testIndices,]


##### Build RRBLUP prediction model every cycle


        trainGenoNewTableComb <- as.big.matrix(rbind(bigmemory::as.matrix(trainGenoNewTablePreCycle),as.matrix(trainGenoNewTable)))

        trainSimGenoValTableComb <- rbind((trainSimGenoValTablePreCycle),trainSimGenoValTable)

        trainSimPhenoValTableComb <- rbind((trainSimPhenoTablePreCycle),trainSimPhenoTable)

        print(dim(trainGenoNewTableComb))

        if(!is.null(sd(trainSimGenoValTableComb[,1])) && !is.null(sd(trainSimPhenoValTableComb[,1])) && !is.na(sd(trainSimGenoValTableComb[,1])) && !is.na(sd(trainSimPhenoValTableComb[,1]))){
             if(sd(trainSimGenoValTableComb[,1])!=0 && sd(trainSimPhenoValTableComb[,1])!=0 && nCyc%%updateFrequency==0){ 


	
           if(modelType=="RRBLUP"){
                        PredictionModel_Geno <- (buildRRBLUPModel(bigmemory::as.matrix(trainGenoNewTableComb),trainSimGenoValTableComb,testGenoNewTable,testSimGenoValTable))
                        PredictionModel_Pheno <-(buildRRBLUPModel(bigmemory::as.matrix(trainGenoNewTableComb),trainSimPhenoTable,testGenoNewTable,testSimPhenoTable))
           }else if(modelType=="RRBLUP_REML"){

                        PredictionModel_Geno <- buildRRBLUPModel_REML(bigmemory::as.matrix(trainGenoNewTableComb), trainSimGenoValTableComb,as.matrix(testGenoNewTable),testSimGenoValTable)
                        PredictionModel_Pheno <- buildRRBLUPModel_REML(bigmemory::as.matrix(trainGenoNewTableComb), trainSimPhenoValTableComb,as.matrix(testGenoNewTable),testSimPhenoTable)
           }

                trainGenoNewTablePreCycle <-  trainGenoNewTableComb

                trainSimPhenoTablePreCycle <-   (trainSimPhenoValTableComb)

                trainSimGenoValTablePreCycle <- (trainSimGenoValTableComb)


                rm(trainGenoNewTableComb)

                rm(trainSimGenoValTableComb)

                rm(trainSimPhenoValTableComb)

                gc()

                }

             }
    }

	if(modelUpdate ==TRUE && updateType=="TrainingCycleWindow"){

			
		PredictionModel_Geno_PreCycle <- PredictionModel_Geno
		PredictionModel_Pheno_PreCycle <- PredictionModel_Pheno
        
        indices<-c(1:nIndividuals)

        trainIndices<-sample(c(1:nIndividuals),(0.8*nIndividuals))

        testIndices<- indices[which(!indices %in% trainIndices)]
		
	    nTrainIndices <- length(trainIndices)

#### Training Set table for Genotype and Simulated Phenotype ###################

        trainGenoNewTable <- newNextGenGenoTable[trainIndices,]

        trainSimPhenoTable <- phenotypicValuesSimTable[trainIndices,]

        trainSimGenoValTable <-  genotypicValuesSimTable[trainIndices,]

#### Validation Set table for Genotype and Simulated Phenotype ###################
        testGenoNewTable <- newNextGenGenoTable[testIndices,]

        testSimPhenoTable <- phenotypicValuesSimTable[testIndices,]

        testSimGenoValTable <-  genotypicValuesSimTable[testIndices,]

        trainGenoNewTablePreCycle <- attach.big.matrix(trainGeno_descriptorFileName)

##### Build RRBLUP prediction model every cycle
        

        trainGenoNewTableComb <- (rbind(bigmemory::as.matrix(trainGenoNewTablePreCycle),as.matrix(trainGenoNewTable)))
        trainSimGenoValTableComb <- rbind((trainSimGenoValTablePreCycle),trainSimGenoValTable)
        trainSimPhenoValTableComb <- rbind((trainSimPhenoTablePreCycle),trainSimPhenoTable)
        print(dim(trainGenoNewTableComb))
	    Avg_GenoSim_Prev <- mean(trainSimGenoValTableComb[,1])
	    Avg_GenoSim_Current <- mean(trainSimGenoValTable[,1])
		
		 

	    change_GenoSim <- Avg_GenoSim_Prev - Avg_GenoSim_Current
	    combinedTableCount <- combinedTableCount+1
        print(change_GenoSim)
    	   
	### 
      
        if(!is.null(sd(trainSimGenoValTableComb[,1])) && !is.null(sd(trainSimPhenoValTableComb[,1])) && !is.na(sd(trainSimGenoValTableComb[,1])) && !is.na(sd(trainSimPhenoValTableComb[,1]))){
           if(sd(trainSimGenoValTableComb[,1])!=0 && sd(trainSimPhenoValTableComb[,1])!=0 && change_GenoSim !=0){ 

	### 
             if(modelType=="RRBLUP"){

                        PredictionModel_Geno <- (buildRRBLUPModel((trainGenoNewTableComb),trainSimGenoValTableComb,as.matrix(testGenoNewTable),testSimGenoValTable))

                        PredictionModel_Pheno <-(buildRRBLUPModel((trainGenoNewTableComb),trainSimPhenoValTableComb,as.matrix(testGenoNewTable),testSimPhenoTable))

           }else if(modelType=="RRBLUP_REML"){

                      					
						PredictionModel_Geno <- tryCatch(buildRRBLUPModel_REML((trainGenoNewTableComb),trainSimGenoValTableComb[,1],as.matrix(testGenoNewTable),testSimGenoValTable[,1]),error=function(e) print(paste("BuildRR-Error -",nCyc,e)))

                        PredictionModel_Pheno <- tryCatch(buildRRBLUPModel_REML((trainGenoNewTableComb),trainSimPhenoValTableComb[,1],as.matrix(testGenoNewTable),testSimPhenoTable[,1]),error=function(e) print(paste("BuildRR-Error -",nCyc,e)))

           }

	     }

           print(sd(PredictionModel_Pheno[[1]]))
           print(sd(PredictionModel_Geno[[1]])) 
           print(length(levels(factor(PredictionModel_Pheno[[1]]))))
           print(length(levels(factor(PredictionModel_Geno[[1]]))))	 

           genoValues <- PredictRRBLUPPhenoValues(newNextGenGenoTable,PredictionModel_Geno)
           phenoValues <- PredictRRBLUPPhenoValues(newNextGenGenoTable,PredictionModel_Pheno)
           print(sd(phenoValues))
           print(sd(genoValues))

 
           if(length(levels(factor(PredictionModel_Pheno[[1]]))) ==1 || length(levels(factor(PredictionModel_Geno[[1]]))) ==1 || (is.na(sd(phenoValues)) || sd(phenoValues)==0) || (is.na(sd(genoValues))) || sd(genoValues)==0 || change_GenoSim==0){

                         PredictionModel_Pheno <- PredictionModel_Pheno_PreCycle
                         PredictionModel_Geno <- PredictionModel_Geno_PreCycle

           }
	 }
		
		trainGeno_backingFileName <- paste("trainTable_Cyc_",trainTableWindowSize,"_",no_QTL,"_",h2,"_",condition,"_",Rep,"_Cyc",nCyc,".bin",sep="")
     
	    trainGeno_descriptorFileName <- paste("trainTable_Cyc_",trainTableWindowSize,"_",no_QTL,"_",h2,"_",condition,"_",Rep,"_Cyc",nCyc,".desc",sep="")
	 	 
		
		if(combinedTableCount < trainTableWindowSize){
			
			   	trainGenoNewTablePreCycle <- deepcopy(as.big.matrix(trainGenoNewTableComb),backingfile=trainGeno_backingFileName,descriptorfile=trainGeno_descriptorFileName)
	 
                trainSimPhenoTablePreCycle <-   (trainSimPhenoValTableComb)

                trainSimGenoValTablePreCycle <- (trainSimGenoValTableComb)
					
		 }else if(combinedTableCount >= trainTableWindowSize){
			
			  	trainGenoNewTablePreCycle <- deepcopy(as.big.matrix(trainGenoNewTableComb[-c(1:nTrainIndices),]),backingfile=trainGeno_backingFileName,descriptorfile=trainGeno_descriptorFileName)

                trainSimPhenoTablePreCycle <-   (trainSimPhenoValTableComb[-c(1:nTrainIndices),])

                trainSimGenoValTablePreCycle <- (trainSimGenoValTableComb[-c(1:nTrainIndices),])
				
				combinedTableCount <- (trainTableWindowSize-1)
					
		  }
		  
		   rm(trainGenoNewTableComb)

           rm(trainSimGenoValTableComb)

           rm(trainSimPhenoValTableComb)
	
    gc()

   }
 }
#################get predicted geno and pheno values with JM and unweighted GP models
  	
	
	if (Weighted==TRUE && WghtMethod =="JM"){
		genoValues <- PredictRRBLUPPhenoValues_WGS_Jannink1(newNextGenGenoTable,PredictionModel_Geno,Freq)
		phenoValues <- PredictRRBLUPPhenoValues_WGS_Jannink1(newNextGenGenoTable,PredictionModel_Pheno,Freq)
    } else if(Weighted ==TRUE && WghtMethod =="DW"){
	
		cycleNumber <- nCyc
		genoValues_PredList <- PredictRRBLUPPhenoValues_WGS_DWModel(newNextGenGenoTable,PredictionModel_Geno,Freq,alphaShape,betaShape,cycleNumber,timeHorizon)
		phenoValues_PredList <- PredictRRBLUPPhenoValues_WGS_DWModel(newNextGenGenoTable,PredictionModel_Pheno,Freq,alphaShape,betaShape,cycleNumber,timeHorizon)

		genoValues <- genoValues_PredList[[1]]
		phenoValues <- phenoValues_PredList[[1]]

		alphaPar_Vector[i] <- genoValues_PredList[[2]]
		
	} else if(Weighted==FALSE){
		genoValues <- PredictRRBLUPPhenoValues(newNextGenGenoTable,PredictionModel_Geno)
		phenoValues <- PredictRRBLUPPhenoValues(newNextGenGenoTable,PredictionModel_Pheno)
    }
	
############# 
    if(GIndStats == TRUE){
###  Get population vector

		populations <- rep(0,20*100)
		init <- 1
		for(pop in 1:20){

			final <- pop*100
			populations[init:final]<- rep(pop,100)
			init <- final+1
		}
		
### Get GInd objects 	
	
		print(length(populations))
		nextGenGenoTable_AlleleFormat <- getAlleleFormat(nextGenGenoTable,AlleleConversionTable_Combined)
		nextGenGenoTable_GInd <- df2genind(as.data.frame(nextGenGenoTable_AlleleFormat),pop=populations,sep="")
		nextGenGenoData_DF <- genind2df(nextGenGenoTable_GInd)
		write.table(nextGenGenoData_DF,paste("PM_",modelType,"trainTable_Cyc_",trainTableWindowSize,"_",no_QTL,"_",h2,"_GenoDataInd_Rep",Rep,"_Cyc_",nCyc,sep=""),sep="\t")
	
	}
#############
    print(paste("GenoVal_Length",length(genoValues)))
    print(paste("PhenoVal_Length",length(phenoValues)))

    GenoVal_Sim_NX_2k_3c[,nCyc] <- genoSimValues
    GenoVal_NX_2k_3c[,nCyc] <- genoValues
   
    PhenoVal_Sim_NX_2k_3c[,nCyc]<- phenoSimValues
    PhenoVal_NX_2k_3c[,nCyc] <- phenoValues
   

### Selection based on simulated or predicted values

    if(selectionOnSimulated ==TRUE){

		genoSelectionTable <- ApplySelection(genoSimValues,no_selected)
		phenoSelectionTable <- ApplySelection(phenoSimValues,no_selected)


		if(selectionOnGeno==TRUE){

			selectedGenoIndividualIndices <- as.vector(genoSelectionTable[,2])
			genoSimSelectedValues <- genoSimValues[selectedGenoIndividualIndices]
			phenoSimSelectedValues <- phenoSimValues[selectedGenoIndividualIndices]

		}else if(selectionOnGeno==FALSE){

			selectedPhenoIndividualIndices <- as.vector(phenoSelectionTable[,2])
			genoSimSelectedValues <- genoSimValues[selectedPhenoIndividualIndices]
			phenoSimSelectedValues <- phenoSimValues[selectedPhenoIndividualIndices]
		}

		GenoVal_NX_N_3c[,nCyc] <- genoSimSelectedValues
	    PhenoVal_NX_N_3c[,nCyc] <- phenoSimSelectedValues
		attainedGenoValues[nCyc] <- max(genoSimSelectedValues)
		selectedGenoIndividualIndices_List[[nCyc]] <- selectedGenoIndividualIndices
	    selectedPhenoIndividualIndices_List[[nCyc]] <- selectedPhenoIndividualIndices
		


	}
      if(selectionOnSimulated==FALSE){

		genoSelectionTable <- ApplySelection(genoValues,no_selected)
		phenoSelectionTable <- ApplySelection(phenoValues,no_selected)

    print(paste("genoSelectionTable-dim-",dim(genoSelectionTable)))
		print(paste("phenoSelectionTable-dim-",dim(phenoSelectionTable)))
		print(paste("selectedGenoIndividualIndices",is.na(phenoSelectionTable[,2])))

		if(selectionOnGeno==TRUE){

			selectedGenoIndividualIndices <- as.vector(genoSelectionTable[,2])
			genoSelectedValues <- genoValues[selectedGenoIndividualIndices]
			phenoSelectedValues <- phenoValues[selectedGenoIndividualIndices]
		  genoSimSelectedValues <- genoSimValues[selectedPhenoIndividualIndices]
		
		}else if(selectionOnGeno==FALSE){

			selectedPhenoIndividualIndices <- as.vector(phenoSelectionTable[,2])
			genoSelectedValues <- genoValues[selectedPhenoIndividualIndices]
			phenoSelectedValues <- phenoValues[selectedPhenoIndividualIndices]
		  genoSimSelectedValues <- genoSimValues[selectedPhenoIndividualIndices]
		}
		  GenoVal_NX_N_3c[,nCyc]<- genoSelectedValues
	    PhenoVal_NX_N_3c[,nCyc]<- phenoSelectedValues
      attainedGenoValues[nCyc] <- max(genoSimSelectedValues)
			
			selectedGenoIndividualIndices_List[[nCyc]] <- selectedGenoIndividualIndices
	    selectedPhenoIndividualIndices_List[[nCyc]] <- selectedPhenoIndividualIndices
		

	}

   	rm(nextGenGenoTable)
	rm(newNextGenGenoTable)
    gc()
  }

  simResults_List <- list(GenoVal_Sim_NX_2k_3c,GenoVal_NX_2k_3c,GenoVal_NX_N_3c,PhenoVal_Sim_NX_2k_3c,PhenoVal_NX_2k_3c,PhenoVal_NX_N_3c,attainedGenoValues,selectedGenoIndividualIndices_List,selectedPhenoIndividualIndices_List)

  return(simResults_List)

}
	


runSimulations40X_Bayes_WGS_V2_Win <- function(model_Sol_Pheno,model_Sol_Geno,F5_Progeny_List,genoValSimList,phenoValSimList,varEList,noQTLs,H2,numberSelected,noCrosses,no_Progeny,noCycles,selectionOnGenoValues,selectionOnSimulatedValues,ModelRetrain,RetrainFrequency,ModelUpdate,UpdateType,UpdateFrequency,i,k,ModelType,weighted,weightingMethod,AlphaShape,BetaShape,TimeHorizon,trainGenoNewTable,trainSimGenoValTable,trainSimPhenoValTable,TrainTableWindowSize,NAM_LinkMap,gIndStats,alleleConversionTable_Combined){

 options(warn=-1)

### Assign Variables

  h2 <- H2
  no_QTL <-noQTLs

  no_selected <- numberSelected
  nCrosses <- noCrosses
  nProgeny <- no_Progeny
  nCycles <- noCycles
  selectionOnGeno <- selectionOnGenoValues
  selectionOnSimulated <- selectionOnSimulatedValues

####################################################################################################

  PredictionModel_Pheno <- model_Sol_Pheno
  PredictionModel_Geno <- model_Sol_Geno

  cycle1_Geno_Data <- F5_Progeny_List
  genoSimValues <- genoValSimList
  phenoSimValues <- phenoValSimList

  varE <- varEList

  nFamilies <- 20
  cycle1_nProgeny <- 100

  modelType <- ModelType
  modelRetrain <- ModelRetrain
  retrainFrequency <- RetrainFrequency 

  modelUpdate <- ModelUpdate
  updateFrequency <- UpdateFrequency
  updateType <- UpdateType

  nMarkers <- 4289
  nIndividuals <- 2000

  Weighted <- weighted 
  WghtMethod <- weightingMethod

  
  alphaShape <- AlphaShape
  betaShape <- BetaShape
  timeHorizon <- TimeHorizon
  alphaPar_Vector <- rep(0,nCycles)

  condition <- i
  Rep <- k
  
    
  trainTableWindowSize <- TrainTableWindowSize
#######################################################################################################
   NAM_LinkMap_New <- NAM_LinkMap 
   GIndStats <- gIndStats
   AlleleConversionTable_Combined <- alleleConversionTable_Combined
  
   nCyc <- 1
   print(paste("cycle_No",nCyc))
           
   trainGeno_backingFileName <- paste("trainTable_",no_QTL,"_",h2,"_",condition,"_",Rep,"_Cyc",nCyc,".bin",sep="")
   trainGeno_descriptorFileName <- paste("trainTable_",no_QTL,"_",h2,"_",condition,"_",Rep,"_Cyc",nCyc,".desc",sep="")
 							
  # trainGenoNewTablePreCycle <- deepcopy(as.big.matrix(trainGenoNewTable),backingfile=trainGeno_backingFileName,descriptorfile=trainGeno_descriptorFileName)
   
   trainGenoNewTablePreCycle <- as.big.matrix(trainGenoNewTable)
   trainSimPhenoTablePreCycle <- trainSimPhenoValTable  
   trainSimGenoValTablePreCycle <- trainSimGenoValTable

####### Predict geno and pheno values ###################################################################

	cycle1GenoTable <- generateMinus101GenoFormat(cycle1_Geno_Data,nFamilies,cycle1_nProgeny)
	newCycle1GenoTable <- generateWeightedGenoTable(cycle1GenoTable,no_QTL)

    if(selectionOnGeno==TRUE){
		MarkerEffects <- unlist(PredictionModel_Geno[3])
	}else if(selectionOnGeno==FALSE){
	    MarkerEffects <- unlist(PredictionModel_Pheno[3])
	}

    Freq_List <- getFreq_BasePoln(cycle1GenoTable,MarkerEffects)
	Freq <- Freq_List[[1]]
	FavAllele <- Freq_List[[2]]

	if(Weighted==TRUE && WghtMethod=="JM"){
		genoValues  <- PredictBayesPhenoValues_Wght(newCycle1GenoTable,PredictionModel_Geno,Freq)
		phenoValues <- PredictBayesPhenoValues_Wght(newCycle1GenoTable,PredictionModel_Pheno,Freq)
	}else if(Weighted==TRUE && WghtMethod=="DW" ){
		genoValues_PredList <- PredictBayesPhenoValues_WGS_DWModel(newCycle1GenoTable,PredictionModel_Geno,Freq,alphaShape,betaShape,cycleNumber,timeHorizon)
		phenoValues_PredList <- PredictBayesPhenoValues_WGS_DWModel(newCycle1GenoTable,PredictionModel_Pheno,Freq,alphaShape,betaShape,cycleNumber,timeHorizon)
		
		genoValues <- genoValues_PredList[[1]]
		phenoValues <- phenoValues_PredList[[1]]
		alphaPar_Vector[nCyc] <- genoValues_PredList[[2]]
	}else if(Weighted==FALSE){
		
		genoValues <- PredictBayesPhenoValues(newCycle1GenoTable,PredictionModel_Geno)
		phenoValues <- PredictBayesPhenoValues(newCycle1GenoTable,PredictionModel_Pheno)
	}
	
	if(GIndStats == TRUE){	 
###  Get population vector

		populations <- rep(0,20*100)
		init <- 1
		for(pop in 1:20){

			final <- pop*100
			populations[init:final]<- rep(pop,100)
			init <- final+1
		}
		
      cycle1GenoTable_AlleleFormat <- getAlleleFormat(cycle1GenoTable,AlleleConversionTable_Combined)
 	  cycle1GenoTable_GInd <- df2genind(cycle1GenoTable_AlleleFormat,pop=populations,sep="")
		
	  cycle1GenoData_DF <- genind2df(cycle1GenoTable_GInd)
	  write.table(cycle1GenoData_DF,paste("PM_",modelType,"_",no_QTL,"_",h2,"_GenoDataInd_Rep",Rep,"_Cyc_",nCyc,sep=""),sep="\t")
	  	
	}
    
### Initiate Arrays and Variables

  GenoVal_NX_2k_3c <- matrix(rep(0,(nCrosses*nProgeny)*nCycles),nrow=(nCrosses*nProgeny),ncol=nCycles)
  GenoVal_NX_N_3c <- matrix(rep(0,no_selected*nCycles),nrow=no_selected,ncol=nCycles)
  GenoVal_Sim_NX_2k_3c <- matrix(rep(0,(nCrosses*nProgeny)*nCycles),nrow=(nCrosses*nProgeny),ncol=nCycles)

  PhenoVal_NX_2k_3c <- matrix(rep(0,(nCrosses*nProgeny)*nCycles),nrow=(nCrosses*nProgeny),ncol=nCycles)
  PhenoVal_NX_N_3c <- matrix(rep(0,no_selected*nCycles),nrow=no_selected,ncol=nCycles)
  PhenoVal_Sim_NX_2k_3c <- matrix(rep(0,(nCrosses*nProgeny)*nCycles),nrow=(nCrosses*nProgeny),ncol=nCycles)

  attainedGenoValues <- rep(0,nCycles)
  selectedGenoIndividualIndices_List <- list()
  selectedPhenoIndividualIndices_List <- list()
  
######## Functions on Cycle 1 Geno Data ################################################################################

	if(selectionOnSimulated ==FALSE){
		  sortedGenoValues <- sort(genoValues,decreasing=TRUE,index.return=TRUE)
		  topNGenoValues <- sortedGenoValues[[1]][1:no_selected]
		  GenoTableIndices_topN <- sortedGenoValues[[2]][1:no_selected]

		  sortedPhenoValues <- sort(phenoValues,decreasing=TRUE,index.return=TRUE)
		  topNPhenoValues <- sortedPhenoValues[[1]][1:no_selected]
		  PhenoTableIndices_topN <- sortedPhenoValues[[2]][1:no_selected]

### Sort genoSim values for  attainedGenoValues

		  sortedGenoSimValues <- sort(genoSimValues,decreasing=TRUE,index.return=TRUE)
		  topNGenoSimValues <- sortedGenoSimValues[[1]][1:no_selected]

	} else if(selectionOnSimulated ==TRUE){

		  sortedGenoValues <- sort(genoSimValues,decreasing=TRUE,index.return=TRUE)
		  topNGenoValues <- sortedGenoValues[[1]][1:no_selected]
		  GenoTableIndices_topN <- sortedGenoValues[[2]][1:no_selected]

		  sortedPhenoValues <- sort(phenoSimValues,decreasing=TRUE,index.return=TRUE)
		  topNPhenoValues <- sortedPhenoValues[[1]][1:no_selected]
		  PhenoTableIndices_topN <- sortedPhenoValues[[2]][1:no_selected]

### Sort genoSim values for  attainedGenoValues

		  sortedGenoSimValues <- sort(genoSimValues,decreasing=TRUE,index.return=TRUE)
		  topNGenoSimValues <- sortedGenoSimValues[[1]][1:no_selected]
	}

## Assign output variables for cycle 1

	  GenoVal_Sim_NX_2k_3c[,1] <- genoSimValues
	  GenoVal_NX_2k_3c[,1] <- genoValues
	  GenoVal_NX_N_3c[,1] <- topNGenoValues
	
	  PhenoVal_Sim_NX_2k_3c[,1] <- phenoSimValues
	  PhenoVal_NX_2k_3c[,1] <- phenoValues
	  PhenoVal_NX_N_3c[,1] <- topNPhenoValues

	  attainedGenoValues[1] <- max(topNGenoSimValues)
	  
	  selectedGenoIndividualIndices <- GenoTableIndices_topN
	  selectedPhenoIndividualIndices <- PhenoTableIndices_topN
	  
	  selectedGenoIndividualIndices_List[[nCyc]] <- selectedGenoIndividualIndices
	  selectedPhenoIndividualIndices_List[[nCyc]] <- selectedPhenoIndividualIndices
		

      rm(cycle1GenoTable)
	  rm(newCycle1GenoTable) 

    
	  
	  combinedTableCount <- 1

############### Cycles 2*29 ###################################################################################
### extract genotype data of selected individuals

  for(nCyc in 2:nCycles){
  
   	print(paste("Cycle_No",nCyc))

    if(nCyc==2){

		if(selectionOnGeno == TRUE){
			selectedGenoData <- extractGenoData(selectedGenoIndividualIndices,cycle1_Geno_Data,no_selected)
		}else if(selectionOnGeno == FALSE){
			selectedGenoData <- extractGenoData(selectedPhenoIndividualIndices,cycle1_Geno_Data,no_selected)
		}

    }
    if(nCyc>2){

		if(selectionOnGeno == TRUE){

			selectedGenoData <- extractGenoData(selectedGenoIndividualIndices,Cycle_Progeny_F5,no_selected)
		}else if(selectionOnGeno == FALSE){
			selectedGenoData <- extractGenoData(selectedPhenoIndividualIndices,Cycle_Progeny_F5,no_selected)
		}

    }


    Parent_Combn_indices <- getParentCombnIndices(no_selected)

    Cycle_Progeny_F1 <- array(0,c(no_selected,nMarkers,2,1))

    Cycle_Progeny_F2 <- array(0,c(no_selected,nMarkers,2,nProgeny))

    Cycle_Progeny_F3 <- array(0,c(no_selected,nMarkers,2,nProgeny))

    Cycle_Progeny_F4 <- array(0,c(no_selected,nMarkers,2,nProgeny))

    Cycle_Progeny_F5 <- array(0,c(no_selected,nMarkers,2,nProgeny))


###################################################################################

    for( j in 1:(nCrosses)){

      parents<- Parent_Combn_indices[,j]

      Parent1<-  selectedGenoData[parents[1],,]
      Parent2<-  selectedGenoData[parents[2],,]

      Cycle_Progeny_F1[j,,,] <- generateProgeny(Parent1,Parent2,1,NAM_LinkMap_New)

      Parent1<- Cycle_Progeny_F1[j,,,1]
      Parent2<- Cycle_Progeny_F1[j,,,1]

      progeny1<- generateProgeny(Parent1,Parent2,nProgeny,NAM_LinkMap_New)
      Cycle_Progeny_F2[j,,,]  <- progeny1


      for(m in 1:nProgeny){

        Parent1<- Cycle_Progeny_F2[j,,,m]
        Parent2<- Cycle_Progeny_F2[j,,,m]
        progeny1<- generateProgeny(Parent1,Parent2,1,NAM_LinkMap_New)
        Cycle_Progeny_F3[j,,,m] <- progeny1
      }


      for(m in 1:nProgeny){

        Parent1<- Cycle_Progeny_F3[j,,,m]
        Parent2<- Cycle_Progeny_F3[j,,,m]
        progeny1<- generateProgeny(Parent1,Parent2,1,NAM_LinkMap_New)
        Cycle_Progeny_F4[j,,,m] <- progeny1
      }

      for(m in 1:nProgeny){

        Parent1<- Cycle_Progeny_F4[j,,,m]
        Parent2<- Cycle_Progeny_F4[j,,,m]
        progeny1<- generateProgeny(Parent1,Parent2,1,NAM_LinkMap_New)
        Cycle_Progeny_F5[j,,,m] <- progeny1
      }

    }

#######################

    nextGenGenoTable <- generateMinus101GenoFormat(Cycle_Progeny_F5,no_selected,nProgeny)
    Freq <- getFreq(nextGenGenoTable,FavAllele)

### Simulate genotypic and phenotypic values 

    genoSimValues <- simulateGenotypicValues(nextGenGenoTable,no_QTL)
    phenoSimValues <- simulatePhenotypicValues(nextGenGenoTable,varE,no_QTL)
    newNextGenGenoTable <- generateWeightedGenoTable(nextGenGenoTable,no_QTL)


if(nCyc%%updateFrequency==0 && modelUpdate ==TRUE){
## Extract training and test data for model build
    IndividualNames <-rep(0,(nCrosses*nProgeny))
      
    IndividualNames <- paste("Ind",c(1:(nCrosses*nProgeny)),sep="")
 
##################################################################################################

    names(phenoSimValues)<-IndividualNames

    Mean_Fixed<- rep(1,nIndividuals)

    phenotypicValuesSimTable <- cbind(phenoSimValues,Mean_Fixed)

#### Table for Simulated Genotypic Values ####################################

    names(genoSimValues)<-IndividualNames

    Mean_Fixed<- rep(1,nIndividuals)

    genotypicValuesSimTable <- cbind(genoSimValues,Mean_Fixed)

####################################
### Model update

    set.seed(25+Rep+nCyc)

	if(modelRetrain==TRUE){ 
        PredDefined <- FALSE

        while(PredDefined ==FALSE){

		indices<-c(1:nIndividuals)

		trainIndices<-sample(c(1:nIndividuals),(0.8*nIndividuals))
		testIndices<- indices[which(!indices %in% trainIndices)]

#### Training Set table for Genotype and Simulated Phenotype ###################

		# trainGenoTable <- genoTable_Mod[trainIndices,]
		trainGenoNewTable <- newNextGenGenoTable[trainIndices,]
		trainSimPhenoTable <- phenotypicValuesSimTable[trainIndices,]
		trainSimGenoTable <-  genotypicValuesSimTable[trainIndices,]

#### Validation Set table for Genotype and Simulated Phenotype ###################

		# testGenoTable<- genoTable_Mod[testIndices,]
		testGenoNewTable <- newNextGenGenoTable[testIndices,]
		testSimPhenoTable <- phenotypicValuesSimTable[testIndices,]
		testSimGenoTable <-  genotypicValuesSimTable[testIndices,]

	    print(length(trainSimGenoTable[,1]))
		print(sd(trainSimGenoTable[,1]))	
		print(sd(trainSimPhenoTable[,1]))
        print(i%%updateFrequency) 
		print(nCyc%%updateFrequency)
        print(summary(trainSimPhenoTable[,1]))		
############### Change RRBLUP to Bayes
      if(!is.null(sd(trainSimGenoTable[,1])) && !is.null(sd(trainSimPhenoTable[,1])) && !is.na(sd(trainSimGenoTable[,1])) && !is.na(sd(trainSimPhenoTable[,1]))){
	    if(sd(trainSimGenoTable[,1])!=0 && sd(trainSimPhenoTable[,1])!=0 && nCyc%%retrainFrequency==0){

			if(modelType=="RRBLUP"){
				PredictionModel_Geno <- (buildRRBLUPModel(trainGenoNewTable,trainSimGenoTable,testGenoNewTable,testSimGenoTable))
				PredictionModel_Pheno <-(buildRRBLUPModel(trainGenoNewTable,trainSimPhenoTable,testGenoNewTable,testSimPhenoTable))
			}else if(modelType=="RRBLUP_REML"){
				PredictionModel_Geno <- (buildRRBLUPModel_REML(trainGenoNewTable,trainSimGenoTable,testGenoNewTable,testSimGenoTable))
				PredictionModel_Pheno <-(buildRRBLUPModel_REML(trainGenoNewTable,trainSimPhenoTable,testGenoNewTable,testSimPhenoTable))
			}
	    }
	  }
	
        if(length(levels(factor(PredictionModel_Pheno[[1]])))>1 && mean(PredictionModel_Pheno[[1]])>0) {
                         PredDefined <- TRUE }      
        }
   }

	gc()	
		
	if(modelUpdate ==TRUE && updateType=="FullSet"){

        indices<-c(1:nIndividuals)

        trainIndices<-sample(c(1:nIndividuals),(0.8*nIndividuals))

        testIndices<- indices[which(!indices %in% trainIndices)]

#### Training Set table for Genotype and Simulated Phenotype ###################

        trainGenoNewTable <- newNextGenGenoTable[trainIndices,]

        trainSimPhenoTable <- phenotypicValuesSimTable[trainIndices,]

        trainSimGenoValTable <-  genotypicValuesSimTable[trainIndices,]

#### Validation Set table for Genotype and Simulated Phenotype ###################

        testGenoNewTable <- newNextGenGenoTable[testIndices,]

        testSimPhenoTable <- phenotypicValuesSimTable[testIndices,]

        testSimGenoValTable <-  genotypicValuesSimTable[testIndices,]


##### Build RRBLUP prediction model every cycle


        trainGenoNewTableComb <- as.big.matrix(rbind(bigmemory::as.matrix(trainGenoNewTablePreCycle),as.matrix(trainGenoNewTable)))

        trainSimGenoValTableComb <- rbind((trainSimGenoValTablePreCycle),trainSimGenoValTable)

        trainSimPhenoValTableComb <- rbind((trainSimPhenoTablePreCycle),trainSimPhenoTable)

        print(dim(trainGenoNewTableComb))

        if(!is.null(sd(trainSimGenoValTableComb[,1])) && !is.null(sd(trainSimPhenoValTableComb[,1])) && !is.na(sd(trainSimGenoValTableComb[,1])) && !is.na(sd(trainSimPhenoValTableComb[,1]))){
            if(sd(trainSimGenoValTableComb[,1])!=0 && sd(trainSimPhenoValTableComb[,1])!=0 && nCyc%%updateFrequency==0){ 


                    
            if(modelType=="BayesB"){
					PredictionModel_Geno <- tryCatch(buildBayesBModel(bigmemory::as.matrix(trainGenoNewTableComb),trainSimGenoValTableComb[,1],testGenoNewTable,testSimGenoValTable[,1],h2),error=function(e) print(paste("BuildBB-Error -",nCyc,e)))
					PredictionModel_Pheno <- tryCatch(buildBayesBModel(bigmemory::as.matrix(trainGenoNewTableComb),trainSimPhenoValTableComb[,1],testGenoNewTable,testSimPhenoTable[,1],h2),error=function(e) print(paste("BuildBB-Error -",nCyc,e)))
			}else if(modelType=="BL"){
			
					PredictionModel_Geno <- tryCatch(buildBLModel(bigmemory::as.matrix(trainGenoNewTableComb),trainSimGenoValTableComb[,1],testGenoNewTable,testSimGenoValTable[,1],h2),error=function(e) print(paste("BuildBL-Error -",nCyc,e)))
					PredictionModel_Pheno <- tryCatch(buildBLModel(bigmemory::as.matrix(trainGenoNewTableComb),trainSimPhenoValTableComb[,1],testGenoNewTable,testSimPhenoTable[,1],h2),error=function(e) print(paste("BuildBL-Error -",nCyc,e)))
			}

	      } 
			   

                trainGenoNewTablePreCycle <-  trainGenoNewTableComb

                trainSimPhenoTablePreCycle <-   (trainSimPhenoValTableComb)

                trainSimGenoValTablePreCycle <- (trainSimGenoValTableComb)


                rm(trainGenoNewTableComb)

                rm(trainSimGenoValTableComb)

                rm(trainSimPhenoValTableComb)

                gc()

        }

    }
    

	if(modelUpdate ==TRUE && updateType=="TrainingCycleWindow"){

			
		PredictionModel_Geno_PreCycle <- PredictionModel_Geno
		PredictionModel_Pheno_PreCycle <- PredictionModel_Pheno
        
        indices<-c(1:nIndividuals)

        trainIndices<- sample(c(1:nIndividuals),(0.8*nIndividuals))

        testIndices<- indices[which(!indices %in% trainIndices)]
		
	    nTrainIndices <- length(trainIndices)

#### Training Set table for Genotype and Simulated Phenotype ###################

        trainGenoNewTable <- newNextGenGenoTable[trainIndices,]

        trainSimPhenoTable <- phenotypicValuesSimTable[trainIndices,]

        trainSimGenoValTable <-  genotypicValuesSimTable[trainIndices,]

#### Validation Set table for Genotype and Simulated Phenotype ###################
        testGenoNewTable <- newNextGenGenoTable[testIndices,]

        testSimPhenoTable <- phenotypicValuesSimTable[testIndices,]

        testSimGenoValTable <-  genotypicValuesSimTable[testIndices,]
	
		trainGenoNewTablePreCycle <- attach.big.matrix(trainGeno_descriptorFileName)
						
##### Build RRBLUP prediction model every cycle
         # bigmemory::as.matrix

        trainGenoNewTableComb <- (rbind(bigmemory::as.matrix(trainGenoNewTablePreCycle),as.matrix(trainGenoNewTable)))
        trainSimGenoValTableComb <- rbind((trainSimGenoValTablePreCycle),trainSimGenoValTable)
        trainSimPhenoValTableComb <- rbind((trainSimPhenoTablePreCycle),trainSimPhenoTable)
        print(dim(trainGenoNewTableComb))
	    Avg_GenoSim_Prev <- mean(trainSimGenoValTableComb[,1])
	    Avg_GenoSim_Current <- mean(trainSimGenoValTable[,1])

		change_GenoSim <- Avg_GenoSim_Prev - Avg_GenoSim_Current
		combinedTableCount <- combinedTableCount+1
        print(change_GenoSim)

        if(!is.null(sd(trainSimGenoValTableComb[,1])) && !is.null(sd(trainSimPhenoValTableComb[,1])) && !is.na(sd(trainSimGenoValTableComb[,1])) && !is.na(sd(trainSimPhenoValTableComb[,1]))){

       	if(sd(trainSimGenoValTableComb[,1])!=0 && sd(trainSimPhenoValTableComb[,1])!=0 && change_GenoSim !=0){

	        if(modelType=="BayesB"){
					PredictionModel_Geno <-  tryCatch(buildBayesBModel((trainGenoNewTableComb),trainSimGenoValTableComb[,1],testGenoNewTable,testSimGenoValTable[,1],h2),error=function(e) print(paste("BuildRR-Error -",nCyc,e)))
					PredictionModel_Pheno <- tryCatch(buildBayesBModel((trainGenoNewTableComb),trainSimPhenoValTableComb[,1],testGenoNewTable,testSimPhenoTable[,1],h2),error=function(e) print(paste("BuildRR-Error -",nCyc,e)))
			}else if(modelType=="BL"){
			
					PredictionModel_Geno <- tryCatch(buildBLModel((trainGenoNewTableComb),trainSimGenoValTableComb[,1],testGenoNewTable,testSimGenoValTable[,1],h2),error=function(e) print(paste("BuildRR-Error -",nCyc,e)))
					PredictionModel_Pheno <- tryCatch(buildBLModel((trainGenoNewTableComb),trainSimPhenoValTableComb[,1],testGenoNewTable,testSimPhenoTable[,1],h2),error=function(e) print(paste("BuildRR-Error -",nCyc,e)))
			}

	    }
		
			

           print(sd(PredictionModel_Pheno[[3]])) 
           print(sd(PredictionModel_Geno[[3]])) 
           print(length(levels(factor(PredictionModel_Pheno[[3]]))))
           print(length(levels(factor(PredictionModel_Geno[[3]]))))	 

           genoValues <- PredictBayesPhenoValues(newNextGenGenoTable,PredictionModel_Geno)
           phenoValues <- PredictBayesPhenoValues(newNextGenGenoTable,PredictionModel_Pheno)
           print(sd(phenoValues))
           print(sd(genoValues))

 
           if(length(levels(factor(PredictionModel_Pheno[[3]]))) ==1 || length(levels(factor(PredictionModel_Geno[[3]]))) ==1 || (is.na(sd(phenoValues)) || sd(phenoValues)==0) || (is.na(sd(genoValues)) || sd(genoValues)==0) || change_GenoSim==0){

                         PredictionModel_Pheno <- PredictionModel_Pheno_PreCycle
                         PredictionModel_Geno <- PredictionModel_Geno_PreCycle

           }
	}
	 
	 
	 
     trainGeno_backingFileName <- paste("trainTable_",no_QTL,"_",h2,"_",condition,"_",Rep,"_Cyc",nCyc,".bin",sep="")
     trainGeno_descriptorFileName <- paste("trainTable_",no_QTL,"_",h2,"_",condition,"_",Rep,"_Cyc",nCyc,".desc",sep="")
	     		
		if(combinedTableCount < trainTableWindowSize){
			
							
				trainGenoNewTablePreCycle <- deepcopy(as.big.matrix(trainGenoNewTableComb),backingfile=trainGeno_backingFileName,descriptorfile=trainGeno_descriptorFileName)

                trainSimPhenoTablePreCycle <-   (trainSimPhenoValTableComb)

                trainSimGenoValTablePreCycle <- (trainSimGenoValTableComb)
					
		}else if(combinedTableCount >= trainTableWindowSize){
			
			  				
				trainGenoNewTablePreCycle <- deepcopy(as.big.matrix(trainGenoNewTableComb[-c(1:nTrainIndices),]),backingfile=trainGeno_backingFileName,descriptorfile=trainGeno_descriptorFileName)

                trainSimPhenoTablePreCycle <-   (trainSimPhenoValTableComb[-c(1:nTrainIndices),])

                trainSimGenoValTablePreCycle <- (trainSimGenoValTableComb[-c(1:nTrainIndices),])
				
				combinedTableCount <- (trainTableWindowSize-1)
						
					
		}
	
    gc()

   }
 
 }  

gc()

#################get predicted geno and pheno values with JM and unweighted GP models

    if(Weighted==TRUE && WghtMethod =="JM"){
		genoValues <- PredictBayesPhenoValues_Wght(newNextGenGenoTable,PredictionModel_Geno,Freq)
		phenoValues <- PredictBayesPhenoValues_Wght(newNextGenGenoTable,PredictionModel_Pheno,Freq)
    } else if (Weighted==TRUE && WghtMethod=="DW"){
		genoValues_PredList <- PredictBayesPhenoValues_WGS_DWModel(newNextGenGenoTable,PredictionModel_Geno,Freq,alphaShape,betaShape,cycleNumber,timeHorizon)
		phenoValues_PredList <- PredictBayesPhenoValues_WGS_DWModel(newNextGenGenoTable,PredictionModel_Pheno,Freq,alphaShape,betaShape,cycleNumber,timeHorizon)

		genoValues <- genoValues_PredList[[1]]
		phenoValues <- phenoValues_PredList[[1]]

		alphaPar_Vector[nCyc] <- genoValues_PredList[[2]]
     }else if(Weighted ==FALSE){
		genoValues <- PredictBayesPhenoValues(newNextGenGenoTable,PredictionModel_Geno)
		phenoValues <- PredictBayesPhenoValues(newNextGenGenoTable,PredictionModel_Pheno)
    }

   	if(GIndStats == TRUE){ 
###  Get population vector

		populations <- rep(0,20*100)
		init <- 1
		for(pop in 1:20){

			final <- pop*100
			populations[init:final]<- rep(pop,100)
			init <- final+1
		}
		
### Get GInd objects 	

		print(length(populations))
		nextGenGenoTable_AlleleFormat <- getAlleleFormat(nextGenGenoTable,AlleleConversionTable_Combined)
		nextGenGenoTable_GInd <- df2genind(as.data.frame(nextGenGenoTable_AlleleFormat),pop=populations,sep="")
		nextGenGenoData_DF <- genind2df(nextGenGenoTable_GInd)
		write.table(nextGenGenoData_DF,paste("PM_",modelType,"trainTable_Cyc_",trainTableWindowSize,"_",no_QTL,"_",h2,"_GenoDataInd_Rep",Rep,"_Cyc_",nCyc,sep=""),sep="\t")	

    }
#############
    print(paste("GenoVal_Length",length(genoValues)))
    print(paste("PhenoVal_Length",length(phenoValues)))

    GenoVal_Sim_NX_2k_3c[,nCyc] <- genoSimValues
    GenoVal_NX_2k_3c[,nCyc] <- genoValues
   
    PhenoVal_Sim_NX_2k_3c[,nCyc]<- phenoSimValues
    PhenoVal_NX_2k_3c[,nCyc] <- phenoValues
   

### Selection based on simulated or predicted values

    if(selectionOnSimulated ==TRUE){

		genoSelectionTable <- ApplySelection(genoSimValues,no_selected)
		phenoSelectionTable <- ApplySelection(phenoSimValues,no_selected)


		if(selectionOnGeno==TRUE){

			selectedGenoIndividualIndices <- as.vector(genoSelectionTable[,2])
			genoSimSelectedValues <- genoSimValues[selectedGenoIndividualIndices]
			phenoSimSelectedValues <- phenoSimValues[selectedGenoIndividualIndices]

		}else if(selectionOnGeno==FALSE){

			selectedPhenoIndividualIndices <- as.vector(phenoSelectionTable[,2])
			genoSimSelectedValues <- genoSimValues[selectedPhenoIndividualIndices]
			phenoSimSelectedValues <- phenoSimValues[selectedPhenoIndividualIndices]
		}

		GenoVal_NX_N_3c[,nCyc] <- genoSimSelectedValues
	    PhenoVal_NX_N_3c[,nCyc] <- phenoSimSelectedValues
		attainedGenoValues[nCyc] <- max(genoSimSelectedValues)
		
		selectedGenoIndividualIndices_List[[nCyc]] <- selectedGenoIndividualIndices
		selectedPhenoIndividualIndices_List[[nCyc]] <- selectedPhenoIndividualIndices
		

	}
    
	if(selectionOnSimulated==FALSE){

		genoSelectionTable <- ApplySelection(genoValues,no_selected)
		phenoSelectionTable <- ApplySelection(phenoValues,no_selected)

        print(paste("genoSelectionTable-dim-",dim(genoSelectionTable)))
		print(paste("phenoSelectionTable-dim-",dim(phenoSelectionTable)))
		print(paste("selectedGenoIndividualIndices",is.na(phenoSelectionTable[,2])))

		if(selectionOnGeno==TRUE){

			selectedGenoIndividualIndices <- as.vector(genoSelectionTable[,2])
			genoSelectedValues <- genoValues[selectedGenoIndividualIndices]
			phenoSelectedValues <- phenoValues[selectedGenoIndividualIndices]
		    genoSimSelectedValues <- genoSimValues[selectedPhenoIndividualIndices]
		
		}else if(selectionOnGeno==FALSE){

			selectedPhenoIndividualIndices <- as.vector(phenoSelectionTable[,2])
			genoSelectedValues <- genoValues[selectedPhenoIndividualIndices]
			phenoSelectedValues <- phenoValues[selectedPhenoIndividualIndices]
		        genoSimSelectedValues <- genoSimValues[selectedPhenoIndividualIndices]
		}
	        GenoVal_NX_N_3c[,nCyc]<- genoSelectedValues
	        PhenoVal_NX_N_3c[,nCyc]<- phenoSelectedValues
        	attainedGenoValues[nCyc] <- max(genoSimSelectedValues)
			
			selectedGenoIndividualIndices_List[[nCyc]] <- selectedGenoIndividualIndices
			selectedPhenoIndividualIndices_List[[nCyc]] <- selectedPhenoIndividualIndices
    }

   	rm(nextGenGenoTable)
	rm(newNextGenGenoTable)
    gc()
  
   }
 
  simResults_List <- list(GenoVal_Sim_NX_2k_3c,GenoVal_NX_2k_3c,GenoVal_NX_N_3c,PhenoVal_Sim_NX_2k_3c,PhenoVal_NX_2k_3c,PhenoVal_NX_N_3c,attainedGenoValues,selectedGenoIndividualIndices_List,selectedPhenoIndividualIndices_List)

  return(simResults_List)

}  


### 

runSimulations40X_RRBLUP_WGS_V3_Win <- function(model_Sol_Pheno,model_Sol_Geno,F5_Progeny_List,genoValSimList,phenoValSimList,varEList,noQTLs,H2,numberSelected,noCrosses,no_Progeny,noCycles,selectionOnGenoValues,selectionOnSimulatedValues,ModelRetrain,RetrainFrequency,ModelUpdate,UpdateType,UpdateFrequency,i,k,ModelType,weighted,weightingMethod,AlphaShape,BetaShape,TimeHorizon,trainGenoNewTable,trainSimGenoValTable,trainSimPhenoValTable,TrainTableWindowSize,BreedingDesign,NAM_LinkMap,gIndStats,alleleConversionTable_Combined){


### Assign Variables

	  h2 <- H2
	  no_QTL <-noQTLs

	  no_selected <- numberSelected
	  nCrosses <- noCrosses
	  nProgeny <- no_Progeny
	  nCycles <- noCycles
	  selectionOnGeno <- selectionOnGenoValues
	  selectionOnSimulated <- selectionOnSimulatedValues

####################################################

	  PredictionModel_Pheno <- model_Sol_Pheno
	  PredictionModel_Geno <- model_Sol_Geno

	  cycle1_Geno_Data <- F5_Progeny_List
	  genoSimValues <- genoValSimList
	  phenoSimValues <- phenoValSimList

	  varE <- varEList

	  condition <- i
	  Rep <- k
	  modelType <- ModelType
	  modelRetrain <- ModelRetrain
	  updateFrequency <- UpdateFrequency
	  retrainFrequency <- RetrainFrequency
	  updateType <- UpdateType
    modelUpdate <- ModelUpdate

	  nFamilies <- 20
    cycle1_nProgeny <- 100
	  nMarkers <- 4289
    nIndividuals <- 2000

    Weighted <- weighted
	  WghtMethod <- weightingMethod  
	  alphaShape <- AlphaShape
	  betaShape <- BetaShape
	  timeHorizon <- TimeHorizon
	  alphaPar_Vector <- rep(0,nCycles)
    BD <- BreedingDesign
         
	  NAM_LinkMap_New <- NAM_LinkMap 
	  trainTableWindowSize <- TrainTableWindowSize
	  
	  nCyc <- 1
		        
	  trainGeno_backingFileName <- paste("trainTable_Cyc_",trainTableWindowSize,"_",no_QTL,"_",h2,"_",condition,"_",Rep,"_Cyc",nCyc,".bin",sep="")
      trainGeno_descriptorFileName <- paste("trainTable_Cyc_",trainTableWindowSize,"_",no_QTL,"_",h2,"_",condition,"_",Rep,"_Cyc",nCyc,".desc",sep="")
 						
	  #trainGenoNewTablePreCycle <- deepcopy(as.big.matrix(trainGenoNewTable),backingfile=trainGeno_backingFileName,descriptorfile=trainGeno_descriptorFileName)
	  
	  trainGenoNewTablePreCycle <- as.big.matrix(trainGenoNewTable)
      trainSimPhenoTablePreCycle <- trainSimPhenoValTable  
      trainSimGenoValTablePreCycle <- trainSimGenoValTable

    
    GIndStats <- gIndStats
	  AlleleConversionTable_Combined <- alleleConversionTable_Combined 
#########################################################################################################
	  nCyc <- 1
      print(paste("cycle_No",nCyc))
	  combinedTableCount <- 1
	  
####### Predict geno and pheno values ###################################################################

	  cycle1GenoTable <- generateMinus101GenoFormat(cycle1_Geno_Data,nFamilies,cycle1_nProgeny)

	  newCycle1GenoTable <- generateWeightedGenoTable(cycle1GenoTable,no_QTL)

	  if(selectionOnGeno==TRUE){
		  MarkerEffects <- unlist(PredictionModel_Geno[1])
	  }else if(selectionOnGeno==FALSE){
	    MarkerEffects <- unlist(PredictionModel_Pheno[1])
	  }

      Freq_List <- getFreq_BasePoln(cycle1GenoTable,MarkerEffects)
	  Freq <- Freq_List[[1]]
	  FavAllele <- Freq_List[[2]]
	  
	  if(GIndStats == TRUE){ 
###  Get population vector

	    populations <- rep(0,20*100)
		  init <- 1
		  for(pop in 1:20){

			  final <- pop*100
			  populations[init:final]<- rep(pop,100)
			  init <- final+1
		  }
		
		cycle1GenoTable_AlleleFormat <- getAlleleFormat(cycle1GenoTable,AlleleConversionTable_Combined)
 	    cycle1GenoTable_GInd <- df2genind(cycle1GenoTable_AlleleFormat,pop=populations,sep="")
		
	  cycle1GenoData_DF <- genind2df(cycle1GenoTable_GInd)
	  write.table(cycle1GenoData_DF,paste("PM_",modelType,"_",no_QTL,"_",h2,"_GenoDataInd_Rep",Rep,"_Cyc_",nCyc,sep=""),sep="\t")
	 }	

###### Predict geno and pheno values ###################################################################

    if(Weighted==TRUE && WghtMethod=="JM"){
			genoValues <- PredictRRBLUPPhenoValues_WGS_Jannink1(newCycle1GenoTable,PredictionModel_Geno,Freq)
			phenoValues <- PredictRRBLUPPhenoValues_WGS_Jannink1(newCycle1GenoTable,PredictionModel_Pheno,Freq)
        
		}else if (Weighted==TRUE && WghtMethod =="DW"){
		
		  cycleNumber <- nCyc
		  genoValues_PredList <- PredictRRBLUPPhenoValues_WGS_DWModel(newCycle1GenoTable,PredictionModel_Geno,Freq,alphaShape,betaShape,cycleNumber,timeHorizon)
			phenoValues_PredList <- PredictRRBLUPPhenoValues_WGS_DWModel(newCycle1GenoTable,PredictionModel_Pheno,Freq,alphaShape,betaShape,cycleNumber,timeHorizon)

			genoValues <- genoValues_PredList[[1]]
			phenoValues <- phenoValues_PredList[[1]]
			alphaPar_Vector[i] <- genoValues_PredList[[2]]
		
		}else if(Weighted==FALSE){
      genoValues  <- PredictRRBLUPPhenoValues(newCycle1GenoTable,PredictionModel_Geno)
      phenoValues <- PredictRRBLUPPhenoValues(newCycle1GenoTable,PredictionModel_Pheno)
    }

	
### Initiate Arrays and Variables  #######################################################################

	  GenoVal_NX_2k_3c <- matrix(rep(0,(nCrosses*nProgeny)*nCycles),nrow=(nCrosses*nProgeny),ncol=nCycles)
	  GenoVal_NX_N_3c <- matrix(rep(0,no_selected*nCycles),nrow=no_selected,ncol=nCycles)
	  GenoVal_Sim_NX_2k_3c <- matrix(rep(0,(nCrosses*nProgeny)*nCycles),nrow=(nCrosses*nProgeny),ncol=nCycles)

	  PhenoVal_NX_2k_3c <- matrix(rep(0,(nCrosses*nProgeny)*nCycles),nrow=(nCrosses*nProgeny),ncol=nCycles)
	  PhenoVal_NX_N_3c <- matrix(rep(0,no_selected*nCycles),nrow=no_selected,ncol=nCycles)
	  PhenoVal_Sim_NX_2k_3c <- matrix(rep(0,(nCrosses*nProgeny)*nCycles),nrow=(nCrosses*nProgeny),ncol=nCycles)

    attainedGenoValues <- rep(0,nCycles)
	 
	  selectedGenoIndividualIndices_List <- list()
    selectedPhenoIndividualIndices_List <- list()


######## Functions on Cycle 1 Geno Data ################################################################################

	if(selectionOnSimulated ==FALSE){
	
		if(selectionOnGeno==TRUE){
	
		  sortedGenoValues <- sort(genoValues,decreasing=TRUE,index.return=TRUE)
		  topNGenoValues <- sortedGenoValues[[1]][1:no_selected]
		  GenoTableIndices_topN <- sortedGenoValues[[2]][1:no_selected]
      topNPhenoValues <- phenoValues[GenoTableIndices_topN] 
		  topNGenoSimValues <- genoSimValues[GenoTableIndices_topN]
		  PhenoTableIndices_topN <- GenoTableIndices_topN
		}else if (selectionOnGeno==FALSE) {
		  	  
		  sortedPhenoValues <- sort(phenoValues,decreasing=TRUE,index.return=TRUE)
		  topNPhenoValues <- sortedPhenoValues[[1]][1:no_selected]
		  PhenoTableIndices_topN <- sortedPhenoValues[[2]][1:no_selected]
		  topNGenoValues <- genoValues[PhenoTableIndices_topN]
		  topNGenoSimValues <- genoSimValues[PhenoTableIndices_topN]
		  GenoTableIndices_topN <- PhenoTableIndices_topN
		} 
  
    }
	if(selectionOnSimulated ==TRUE){
	
		if(selectionOnGeno==TRUE){
		 
   		  sortedGenoValues <- sort(genoSimValues,decreasing=TRUE,index.return=TRUE)
		  topNGenoValues <- sortedGenoValues[[1]][1:no_selected]
		  GenoTableIndices_topN <- sortedGenoValues[[2]][1:no_selected]
		  topNPhenoValues <- phenoValues[GenoTableIndices_topN] 
		  topNGenoSimValues <- genoSimValues[GenoTableIndices_topN]
		  PhenoTableIndices_topN <- GenoTableIndices_topN
		
		}else if(selectionOnGeno==FALSE){

		  sortedPhenoValues <- sort(phenoSimValues,decreasing=TRUE,index.return=TRUE)
		  topNPhenoValues <- sortedPhenoValues[[1]][1:no_selected]
		  PhenoTableIndices_topN <- sortedPhenoValues[[2]][1:no_selected]
		  topNGenoValues <- genoValues[PhenoTableIndices_topN]
		  topNGenoSimValues <- genoSimValues[PhenoTableIndices_topN]
		  GenoTableIndices_topN <- PhenoTableIndices_topN
	    }
	}

##################################################################################################################
## Assign output variables for cycle 1

	  GenoVal_Sim_NX_2k_3c[,1] <- genoSimValues
	  GenoVal_NX_2k_3c[,1] <- genoValues
	  GenoVal_NX_N_3c[,1] <- topNGenoValues
    
	  attainedGenoValues[1] <- max(topNGenoSimValues)
	  
	  selectedGenoIndividualIndices <- GenoTableIndices_topN
      selectedPhenoIndividualIndices <- PhenoTableIndices_topN
	  
	  selectedGenoIndividualIndices_List[[nCyc]] <- selectedGenoIndividualIndices
	  selectedPhenoIndividualIndices_List[[nCyc]] <- selectedPhenoIndividualIndices
		

	  PhenoVal_Sim_NX_2k_3c[,1] <- phenoSimValues
	  PhenoVal_NX_2k_3c[,1] <- phenoValues
	  PhenoVal_NX_N_3c[,1] <- topNPhenoValues


      rm(cycle1GenoTable)
	  rm(newCycle1GenoTable)

############### Cycles 2*39 ###################################################################################
### extract genotype data of selected individuals
  
  for(nCyc in 2:nCycles){

   print(paste("cycleNo-",nCyc))
		
	 if(nCyc==2){

		if(selectionOnGeno == TRUE){
			selectedGenoData <- extractGenoData(selectedGenoIndividualIndices,cycle1_Geno_Data,no_selected)
		}else if(selectionOnGeno == FALSE){
			selectedGenoData <- extractGenoData(selectedPhenoIndividualIndices,cycle1_Geno_Data,no_selected)
		}

	}else if(nCyc>2){

		if(selectionOnGeno == TRUE){

			selectedGenoData <- extractGenoData(selectedGenoIndividualIndices,Cycle_Progeny_F5,no_selected)
		}else if(selectionOnGeno == FALSE){
			selectedGenoData <- extractGenoData(selectedPhenoIndividualIndices,Cycle_Progeny_F5,no_selected)
		}

	}
	
	Cycle_Progeny_F5 <- getF5RILs_BD(BD,selectedGenoData,no_selected,nProgeny,nMarkers,NAM_LinkMap_New)
	
	
########################
##  nextGenGenoTable <- generate012GenoFormat(Cycle_Progeny_F5,no_selected)

    nextGenGenoTable <- generateMinus101GenoFormat(Cycle_Progeny_F5,nCrosses,nProgeny)

    Freq <- getFreq(nextGenGenoTable,FavAllele)
    genoSimValues <- simulateGenotypicValues(nextGenGenoTable,no_QTL)
    phenoSimValues <- simulatePhenotypicValues(nextGenGenoTable,varE,no_QTL)

    newNextGenGenoTable <- generateWeightedGenoTable(nextGenGenoTable,no_QTL)

 if(nCyc%%updateFrequency==0 && modelUpdate ==TRUE){
 
    set.seed(25+Rep+nCyc)
## Extract training and test data for model build

 
    IndividualNames <-rep(0,(nCrosses*nProgeny))

   IndividualNames <- paste("Ind",c(1:(nCrosses*nProgeny)),sep="")
##################################################################################################

    names(phenoSimValues)<-IndividualNames

    Mean_Fixed<- rep(1,nIndividuals)

    phenotypicValuesSimTable<- cbind(phenoSimValues,Mean_Fixed)


#### Table for Simulated Genotypic Values ####################################

    names(genoSimValues)<-IndividualNames

    Mean_Fixed<- rep(1,nIndividuals)

    genotypicValuesSimTable <- cbind(genoSimValues,Mean_Fixed)


### Model Retrain 

	if(modelRetrain==TRUE){ 
        PredDefined <- FALSE

        while(PredDefined ==FALSE){

		indices<-c(1:nIndividuals)

		trainIndices<-sample(c(1:nIndividuals),(0.8*nIndividuals))
		testIndices<- indices[which(!indices %in% trainIndices)]

#### Training Set table for Genotype and Simulated Phenotype ###################

		# trainGenoTable <- genoTable_Mod[trainIndices,]
		trainGenoNewTable <- newNextGenGenoTable[trainIndices,]
		trainSimPhenoTable <- phenotypicValuesSimTable[trainIndices,]
		trainSimGenoTable <-  genotypicValuesSimTable[trainIndices,]

#### Validation Set table for Genotype and Simulated Phenotype ###################

		# testGenoTable<- genoTable_Mod[testIndices,]
		testGenoNewTable <- newNextGenGenoTable[testIndices,]
		testSimPhenoTable <- phenotypicValuesSimTable[testIndices,]
		testSimGenoTable <-  genotypicValuesSimTable[testIndices,]

	    print(length(trainSimGenoTable[,1]))
		print(sd(trainSimGenoTable[,1]))	
		print(sd(trainSimPhenoTable[,1]))
        print(i%%updateFrequency) 
		print(nCyc%%updateFrequency)
        print(summary(trainSimPhenoTable[,1]))		
############### Change RRBLUP to Bayes
      if(!is.null(sd(trainSimGenoTable[,1])) && !is.null(sd(trainSimPhenoTable[,1])) && !is.na(sd(trainSimGenoTable[,1])) && !is.na(sd(trainSimPhenoTable[,1]))){
	    if(sd(trainSimGenoTable[,1])!=0 && sd(trainSimPhenoTable[,1])!=0 && nCyc%%retrainFrequency==0){

			if(modelType=="RRBLUP"){
				PredictionModel_Geno <- (buildRRBLUPModel(trainGenoNewTable,trainSimGenoTable,testGenoNewTable,testSimGenoTable))
				PredictionModel_Pheno <-(buildRRBLUPModel(trainGenoNewTable,trainSimPhenoTable,testGenoNewTable,testSimPhenoTable))
			}else if(modelType=="RRBLUP_REML"){
				PredictionModel_Geno <- (buildRRBLUPModel_REML(trainGenoNewTable,trainSimGenoTable,testGenoNewTable,testSimGenoTable))
				PredictionModel_Pheno <-(buildRRBLUPModel_REML(trainGenoNewTable,trainSimPhenoTable,testGenoNewTable,testSimPhenoTable))
			}
	    }
	  }
	
        if(length(levels(factor(PredictionModel_Pheno[[1]])))>1 && mean(PredictionModel_Pheno[[1]])>0) {
                         PredDefined <- TRUE }      
        }
   }

	gc()	
		
	if(modelUpdate ==TRUE && updateType=="FullSet"){

        indices<-c(1:nIndividuals)

        trainIndices<-sample(c(1:nIndividuals),(0.8*nIndividuals))

        testIndices<- indices[which(!indices %in% trainIndices)]

#### Training Set table for Genotype and Simulated Phenotype ###################

        trainGenoNewTable <- newNextGenGenoTable[trainIndices,]

        trainSimPhenoTable <- phenotypicValuesSimTable[trainIndices,]

        trainSimGenoValTable <-  genotypicValuesSimTable[trainIndices,]

#### Validation Set table for Genotype and Simulated Phenotype ###################

        testGenoNewTable <- newNextGenGenoTable[testIndices,]

        testSimPhenoTable <- phenotypicValuesSimTable[testIndices,]

        testSimGenoValTable <-  genotypicValuesSimTable[testIndices,]


##### Build RRBLUP prediction model every cycle


        trainGenoNewTableComb <- as.big.matrix(rbind(bigmemory::as.matrix(trainGenoNewTablePreCycle),as.matrix(trainGenoNewTable)))

        trainSimGenoValTableComb <- rbind((trainSimGenoValTablePreCycle),trainSimGenoValTable)

        trainSimPhenoValTableComb <- rbind((trainSimPhenoTablePreCycle),trainSimPhenoTable)

        print(dim(trainGenoNewTableComb))

        if(!is.null(sd(trainSimGenoValTableComb[,1])) && !is.null(sd(trainSimPhenoValTableComb[,1])) && !is.na(sd(trainSimGenoValTableComb[,1])) && !is.na(sd(trainSimPhenoValTableComb[,1]))){
             if(sd(trainSimGenoValTableComb[,1])!=0 && sd(trainSimPhenoValTableComb[,1])!=0 && nCyc%%updateFrequency==0){ 


	
           if(modelType=="RRBLUP"){
                        PredictionModel_Geno <- (buildRRBLUPModel(bigmemory::as.matrix(trainGenoNewTableComb),trainSimGenoValTableComb,testGenoNewTable,testSimGenoValTable))
                        PredictionModel_Pheno <-(buildRRBLUPModel(bigmemory::as.matrix(trainGenoNewTableComb),trainSimPhenoTable,testGenoNewTable,testSimPhenoTable))
           }else if(modelType=="RRBLUP_REML"){

                        PredictionModel_Geno <- buildRRBLUPModel_REML(bigmemory::as.matrix(trainGenoNewTableComb), trainSimGenoValTableComb,as.matrix(testGenoNewTable),testSimGenoValTable)
                        PredictionModel_Pheno <- buildRRBLUPModel_REML(bigmemory::as.matrix(trainGenoNewTableComb), trainSimPhenoValTableComb,as.matrix(testGenoNewTable),testSimPhenoTable)
           }

                trainGenoNewTablePreCycle <-  trainGenoNewTableComb

                trainSimPhenoTablePreCycle <-   (trainSimPhenoValTableComb)

                trainSimGenoValTablePreCycle <- (trainSimGenoValTableComb)


                rm(trainGenoNewTableComb)

                rm(trainSimGenoValTableComb)

                rm(trainSimPhenoValTableComb)

                gc()

                }

             }
    }

	if(modelUpdate ==TRUE && updateType=="TrainingCycleWindow"){

			
		PredictionModel_Geno_PreCycle <- PredictionModel_Geno
		PredictionModel_Pheno_PreCycle <- PredictionModel_Pheno
        
        indices<-c(1:nIndividuals)

        trainIndices<-sample(c(1:nIndividuals),(0.8*nIndividuals))

        testIndices<- indices[which(!indices %in% trainIndices)]
		
	    nTrainIndices <- length(trainIndices)

#### Training Set table for Genotype and Simulated Phenotype ###################

        trainGenoNewTable <- newNextGenGenoTable[trainIndices,]

        trainSimPhenoTable <- phenotypicValuesSimTable[trainIndices,]

        trainSimGenoValTable <-  genotypicValuesSimTable[trainIndices,]

#### Validation Set table for Genotype and Simulated Phenotype ###################
        testGenoNewTable <- newNextGenGenoTable[testIndices,]

        testSimPhenoTable <- phenotypicValuesSimTable[testIndices,]

        testSimGenoValTable <-  genotypicValuesSimTable[testIndices,]

        trainGenoNewTablePreCycle <- attach.big.matrix(trainGeno_descriptorFileName)

##### Build RRBLUP prediction model every cycle
        

        trainGenoNewTableComb <- (rbind(bigmemory::as.matrix(trainGenoNewTablePreCycle),as.matrix(trainGenoNewTable)))
        trainSimGenoValTableComb <- rbind((trainSimGenoValTablePreCycle),trainSimGenoValTable)
        trainSimPhenoValTableComb <- rbind((trainSimPhenoTablePreCycle),trainSimPhenoTable)
        print(dim(trainGenoNewTableComb))
	      Avg_GenoSim_Prev <- mean(trainSimGenoValTableComb[,1])
	      Avg_GenoSim_Current <- mean(trainSimGenoValTable[,1])
	
	     change_GenoSim <- Avg_GenoSim_Prev - Avg_GenoSim_Current
	     combinedTableCount <- combinedTableCount+1
       print(change_GenoSim)
    	   
	### 
      
        if(!is.null(sd(trainSimGenoValTableComb[,1])) && !is.null(sd(trainSimPhenoValTableComb[,1])) && !is.na(sd(trainSimGenoValTableComb[,1])) && !is.na(sd(trainSimPhenoValTableComb[,1]))){
           if(sd(trainSimGenoValTableComb[,1])!=0 && sd(trainSimPhenoValTableComb[,1])!=0 && change_GenoSim !=0){ 

	### 
             if(modelType=="RRBLUP"){

                        PredictionModel_Geno <- (buildRRBLUPModel((trainGenoNewTableComb),trainSimGenoValTableComb,as.matrix(testGenoNewTable),testSimGenoValTable))

                        PredictionModel_Pheno <-(buildRRBLUPModel((trainGenoNewTableComb),trainSimPhenoValTableComb,as.matrix(testGenoNewTable),testSimPhenoTable))

           }else if(modelType=="RRBLUP_REML"){

                      					
						PredictionModel_Geno <- tryCatch(buildRRBLUPModel_REML((trainGenoNewTableComb),trainSimGenoValTableComb[,1],as.matrix(testGenoNewTable),testSimGenoValTable[,1]),error=function(e) print(paste("BuildRR-Error -",nCyc,e)))

                        PredictionModel_Pheno <- tryCatch(buildRRBLUPModel_REML((trainGenoNewTableComb),trainSimPhenoValTableComb[,1],as.matrix(testGenoNewTable),testSimPhenoTable[,1]),error=function(e) print(paste("BuildRR-Error -",nCyc,e)))

           }

	     }

           print(sd(PredictionModel_Pheno[[1]]))
           print(sd(PredictionModel_Geno[[1]])) 
           print(length(levels(factor(PredictionModel_Pheno[[1]]))))
           print(length(levels(factor(PredictionModel_Geno[[1]]))))	 

           genoValues <- PredictRRBLUPPhenoValues(newNextGenGenoTable,PredictionModel_Geno)
           phenoValues <- PredictRRBLUPPhenoValues(newNextGenGenoTable,PredictionModel_Pheno)
           print(sd(phenoValues))
           print(sd(genoValues))

 
           if(length(levels(factor(PredictionModel_Pheno[[1]]))) ==1 || length(levels(factor(PredictionModel_Geno[[1]]))) ==1 || (is.na(sd(phenoValues)) || sd(phenoValues)==0) || (is.na(sd(genoValues))) || sd(genoValues)==0 || change_GenoSim==0){

                         PredictionModel_Pheno <- PredictionModel_Pheno_PreCycle
                         PredictionModel_Geno <- PredictionModel_Geno_PreCycle

           }
	 }
		
		trainGeno_backingFileName <- paste("trainTable_Cyc_",trainTableWindowSize,"_",no_QTL,"_",h2,"_",condition,"_",Rep,"_Cyc",nCyc,".bin",sep="")
     
	    trainGeno_descriptorFileName <- paste("trainTable_Cyc_",trainTableWindowSize,"_",no_QTL,"_",h2,"_",condition,"_",Rep,"_Cyc",nCyc,".desc",sep="")
	 	 
		
		if(combinedTableCount < trainTableWindowSize){
			
			   	trainGenoNewTablePreCycle <- deepcopy(as.big.matrix(trainGenoNewTableComb),backingfile=trainGeno_backingFileName,descriptorfile=trainGeno_descriptorFileName)
	 
                trainSimPhenoTablePreCycle <-   (trainSimPhenoValTableComb)

                trainSimGenoValTablePreCycle <- (trainSimGenoValTableComb)
					
		 }else if(combinedTableCount >= trainTableWindowSize){
			
			  	trainGenoNewTablePreCycle <- deepcopy(as.big.matrix(trainGenoNewTableComb[-c(1:nTrainIndices),]),backingfile=trainGeno_backingFileName,descriptorfile=trainGeno_descriptorFileName)

                trainSimPhenoTablePreCycle <-   (trainSimPhenoValTableComb[-c(1:nTrainIndices),])

                trainSimGenoValTablePreCycle <- (trainSimGenoValTableComb[-c(1:nTrainIndices),])
				
				combinedTableCount <- (trainTableWindowSize-1)
					
		  }
		  
		   rm(trainGenoNewTableComb)

           rm(trainSimGenoValTableComb)

           rm(trainSimPhenoValTableComb)
	
    gc()

   }
 }
#################get predicted geno and pheno values with JM and unweighted GP models
  	
	
  if (Weighted==TRUE && WghtMethod =="JM"){
		genoValues <- PredictRRBLUPPhenoValues_WGS_Jannink1(newNextGenGenoTable,PredictionModel_Geno,Freq)
		phenoValues <- PredictRRBLUPPhenoValues_WGS_Jannink1(newNextGenGenoTable,PredictionModel_Pheno,Freq)
    } else if(Weighted ==TRUE && WghtMethod =="DW"){
	
		cycleNumber <- nCyc
		genoValues_PredList <- PredictRRBLUPPhenoValues_WGS_DWModel(newNextGenGenoTable,PredictionModel_Geno,Freq,alphaShape,betaShape,cycleNumber,timeHorizon)
		phenoValues_PredList <- PredictRRBLUPPhenoValues_WGS_DWModel(newNextGenGenoTable,PredictionModel_Pheno,Freq,alphaShape,betaShape,cycleNumber,timeHorizon)

		genoValues <- genoValues_PredList[[1]]
		phenoValues <- phenoValues_PredList[[1]]

		alphaPar_Vector[i] <- genoValues_PredList[[2]]
		
	} else if(Weighted==FALSE){
		genoValues <- PredictRRBLUPPhenoValues(newNextGenGenoTable,PredictionModel_Geno)
		phenoValues <- PredictRRBLUPPhenoValues(newNextGenGenoTable,PredictionModel_Pheno)
    }
	
############# 
  if(GIndStats == TRUE){
###  Get population vector

		populations <- rep(0,20*100)
		init <- 1
		for(pop in 1:20){

			final <- pop*100
			populations[init:final]<- rep(pop,100)
			init <- final+1
		}
		
### Get GInd objects 	
	
		print(length(populations))
		nextGenGenoTable_AlleleFormat <- getAlleleFormat(nextGenGenoTable,AlleleConversionTable_Combined)
		nextGenGenoTable_GInd <- df2genind(as.data.frame(nextGenGenoTable_AlleleFormat),pop=populations,sep="")
		nextGenGenoData_DF <- genind2df(nextGenGenoTable_GInd)
		write.table(nextGenGenoData_DF,paste("PM_",modelType,"trainTable_Cyc_",trainTableWindowSize,"_",no_QTL,"_",h2,"_GenoDataInd_Rep",Rep,"_Cyc_",nCyc,sep=""),sep="\t")
	
	}
#############
    print(paste("GenoVal_Length",length(genoValues)))
    print(paste("PhenoVal_Length",length(phenoValues)))

    GenoVal_Sim_NX_2k_3c[,nCyc] <- genoSimValues
    GenoVal_NX_2k_3c[,nCyc] <- genoValues
   
    PhenoVal_Sim_NX_2k_3c[,nCyc]<- phenoSimValues
    PhenoVal_NX_2k_3c[,nCyc] <- phenoValues
   

### Selection based on simulated or predicted values

    if(selectionOnSimulated ==TRUE){

		genoSelectionTable <- ApplySelection(genoSimValues,no_selected)
		phenoSelectionTable <- ApplySelection(phenoSimValues,no_selected)


		if(selectionOnGeno==TRUE){

			selectedGenoIndividualIndices <- as.vector(genoSelectionTable[,2])
			genoSimSelectedValues <- genoSimValues[selectedGenoIndividualIndices]
			phenoSimSelectedValues <- phenoSimValues[selectedGenoIndividualIndices]

		}else if(selectionOnGeno==FALSE){

			selectedPhenoIndividualIndices <- as.vector(phenoSelectionTable[,2])
			genoSimSelectedValues <- genoSimValues[selectedPhenoIndividualIndices]
			phenoSimSelectedValues <- phenoSimValues[selectedPhenoIndividualIndices]
		}

		GenoVal_NX_N_3c[,nCyc] <- genoSimSelectedValues
	    PhenoVal_NX_N_3c[,nCyc] <- phenoSimSelectedValues
		attainedGenoValues[nCyc] <- max(genoSimSelectedValues)
		selectedGenoIndividualIndices_List[[nCyc]] <- selectedGenoIndividualIndices
	    selectedPhenoIndividualIndices_List[[nCyc]] <- selectedPhenoIndividualIndices
		


	}
    if(selectionOnSimulated==FALSE){

		genoSelectionTable <- ApplySelection(genoValues,no_selected)
		phenoSelectionTable <- ApplySelection(phenoValues,no_selected)

    	print(paste("genoSelectionTable-dim-",dim(genoSelectionTable)))
		print(paste("phenoSelectionTable-dim-",dim(phenoSelectionTable)))
		print(paste("selectedGenoIndividualIndices",is.na(phenoSelectionTable[,2])))

		if(selectionOnGeno==TRUE){

			selectedGenoIndividualIndices <- as.vector(genoSelectionTable[,2])
			genoSelectedValues <- genoValues[selectedGenoIndividualIndices]
			phenoSelectedValues <- phenoValues[selectedGenoIndividualIndices]
		  genoSimSelectedValues <- genoSimValues[selectedPhenoIndividualIndices]
		
		}else if(selectionOnGeno==FALSE){

			selectedPhenoIndividualIndices <- as.vector(phenoSelectionTable[,2])
			genoSelectedValues <- genoValues[selectedPhenoIndividualIndices]
			phenoSelectedValues <- phenoValues[selectedPhenoIndividualIndices]
		    genoSimSelectedValues <- genoSimValues[selectedPhenoIndividualIndices]
		}
		    GenoVal_NX_N_3c[,nCyc]<- genoSelectedValues
	        PhenoVal_NX_N_3c[,nCyc]<- phenoSelectedValues
        	attainedGenoValues[nCyc] <- max(genoSimSelectedValues)
			
			selectedGenoIndividualIndices_List[[nCyc]] <- selectedGenoIndividualIndices
	        selectedPhenoIndividualIndices_List[[nCyc]] <- selectedPhenoIndividualIndices
		

	}

   	rm(nextGenGenoTable)
	rm(newNextGenGenoTable)
    gc()
  }

  simResults_List <- list(GenoVal_Sim_NX_2k_3c,GenoVal_NX_2k_3c,GenoVal_NX_N_3c,PhenoVal_Sim_NX_2k_3c,PhenoVal_NX_2k_3c,PhenoVal_NX_N_3c,attainedGenoValues,selectedGenoIndividualIndices_List,selectedPhenoIndividualIndices_List)

  return(simResults_List)

}
	
### 
```
