## Supplementary material for "Factors Affecting Response to Recurrent Genomic Selection in Soybeans": File S1: functionSVMLinearModel.r

```
#### Function for SVM Linear genomic prediction model with caret package

buildSVMLinearModel <- function(trainGenoTable,trainPhenoTable,testGenoTable,testPhenoTable){
  
  library(caret)
  library(kernlab)  
  trainPheno <- trainPhenoTable[,1]
  trainGeno <- trainGenoTable
  nIndividuals <- 2000
  
  folds=5
  repeats=1
  myControl <- trainControl(method='cv', number=folds, repeats=repeats, 
                          returnResamp='none', classProbs=FALSE,
                          returnData=FALSE, savePredictions=TRUE, 
                          verboseIter=TRUE, allowParallel=TRUE,
                          index=createMultiFolds(trainPheno, k=folds, times=repeats)) 
						  
  
  model1<- train(trainGeno,trainPheno, method='svmLinear',trControl=myControl)
  
 
## Predict with test data set ###########################################################################s
  testPheno <- testPhenoTable[,1] 
  testGeno <- testGenoTable 
 
### Predictions with probability values
#preds_prob <- data.frame(sapply(all.models.2, function(x){predict(x, testGeno,type='prob')}))
 
  pred.pheno.train <- predict(model1,trainGeno)
  pred.pheno.valid <- predict(model1,testGeno)

# cor(Pred.pheno.train,trainPheno)
   u.effects <- coef(model1$finalModel)
#################################################################################################################
      
  Pred.Accuracy <- cor(pred.pheno.valid,testPheno)
  
  
  train_MSE <- (sum((pred.pheno.train-trainPheno))^2)/(0.8*nIndividuals)
  test_MSE <- (sum((pred.pheno.valid-testPheno))^2)/(0.2*nIndividuals) 
  
  sol <- list(Pred.Accuracy,model1,u.effects,train_MSE,test_MSE)
  
  return(sol)
  
}  


################################################################


# model2 <- train(Data1625OR,Y, method='svmRadial', trControl=myControl,metric='Accuracy')
```
