## Supplementary material for "Factors Affecting Response to Recurrent Genomic Selection in Soybeans": File S1: functionSVMRBFModel.r

```
#### Function for SVM RBF genomic prediction model with caret package

buildSVMRBFModel <- function(trainGenoTable,trainPhenoTable,testGenoTable,testPhenoTable){
  
  
  trainPheno <- trainPhenoTable
  trainGeno <- trainGenoTable
  nIndividuals_train <- nrow(trainGenoTable)
  nIndividuals_test <- nrow(testGenoTable)
  
  folds=5
  repeats=5
  myControl <- trainControl(method='cv', number=folds, repeats=repeats, 
                          returnResamp='none', classProbs=FALSE,
                          returnData=FALSE, savePredictions=TRUE, 
                          verboseIter=TRUE, allowParallel=TRUE,
                          index=createMultiFolds(trainPheno, k=folds, times=repeats)) 
						  
  
   model1<- (train(trainGeno,trainPheno, method='svmRadial',trControl=myControl))
  
   # gridRF <- expand.grid(mfinal=c(50,100,150,175,200))
   
   # model1<- train(trainGeno,trainPheno, method='rf',trControl=myControl,tuneGrid=gridRF)
	
## Predict with test data set ###########################################################################s
  testPheno <- testPhenoTable 
  testGeno <- testGenoTable 
 
### Predictions with probability values

  #preds_prob <- data.frame(sapply(all.models.2, function(x){predict(x, testGeno,type='prob')}))
 
  pred.pheno.train <- (kernlab::predict(model1,trainGeno))
  pred.pheno.valid <- (kernlab::predict(model1,testGeno))

# cor(Pred.pheno.train,trainPheno)
   u.effects <- coef(model1$finalModel)
#################################################################################################################
      
  Pred.Accuracy <- cor(pred.pheno.valid,testPheno)
  
  train_MSE <- (sum((pred.pheno.train-trainPheno))^2)/(nIndividuals_train)
  test_MSE <- (sum((pred.pheno.valid-testPheno))^2)/(nIndividuals_test) 
  
  sol <- list(Pred.Accuracy,model1,u.effects,train_MSE,test_MSE)
  
  return(sol)
  
} 

################################################################
```
