## Supplementary material for "Factors Affecting Response to Recurrent Genomic Selection in Soybeans": File S1: getBasePopulation_List.R

```
### Generate founder population similar to SoyNAM populations with the specified number of markers, QTLs, genetic architecture, h^2 

## Number of markers - 4289 high quality marker set with linkage information

################################################################################
## Generate parent genotypes (with same no of markers in a 6K genotypic dataset), 20 parents X 1 common parent 
################################################################################
# 50% of QTL with positive effect and 50% with negative effect
# Effect size is  a= 1/No_QTL * maximum_genotypic value with equal effect size for all QTL 
# QTL can have three possible genotypes (-1,0,1) 

## Function to generate F5 RILs includes function to simulate meiosis and generate progeny genotype by selfing F1 for four generations #########################################

##### Initiate Lists ###########################################################

getBasePopulation_FamilyWise_List <- function(nReps){

	nreps <- nReps
##################################### 
 
####### Conditions ###################################################################
## 1) QTL_No: 40 - h2=0.7 ; 2) QTL_No: 40 - h2=0.3 ; 3) QTL_No: 400 - h2=0.7 ; 
## 4) QTL_No: 400 - h2=0.3; 5)  QTL_No: nMarkers- h2=0.7; 6)QTL_No: nMarkers- h2=0.7; 

######################################################################################

## Get Recombination frequency vector from NAM linkage data 6K chip.

  NAM_LinkMap_New <- getNAMLinkageMapTable()

	RF_Vector <- as.vector(NAM_LinkMap_New[,3])
	nMarkers <- length(RF_Vector)

### Set Base population PG variables 

	no_QTL <- c(40,400,nMarkers)
	nCrosses <-20 
	nProgeny <- 100 
	no_selected <- 20
	nIndividuals <- 2000
    h2 <- c(0.7,0.3)
	
  plotHClust <- FALSE
	founderSimilarity <- FALSE
##Initiate output variables

  F_Progeny_List <- list()

    
	phenotypicValuesSimTableList_Reps <- rep(list(list()),nreps)
	genotypicValuesSimTableList_Reps <- rep(list(list()),nreps)

	phenotypicValuesSimList_Reps <- rep(list(list()),nreps)
	genotypicValuesSimList_Reps <-rep(list(list()),nreps)

	trainIndicesList_Reps <- rep(list(list()),nreps)
	testIndicesList_Reps<- rep(list(list()),nreps)

	trainGenoTableList_Reps <- rep(list(list()),nreps)
	trainGenoNewTableList_Reps <-rep(list(list()),nreps)
	trainGenoNewTableList_w_o_QTL_Reps <-rep(list(list()),nreps)
	trainSimGenoValuesTableList_Reps <- rep(list(list()),nreps)
	trainSimPhenoValuesTableList_Reps <- rep(list(list()),nreps)

	
	testGenoTableList_Reps <- rep(list(list()),nreps)
	testGenoNewTableList_Reps <- rep(list(list()),nreps)
	testGenoNewTableList_w_o_QTL_Reps <- rep(list(list()),nreps)
	testSimGenoValuesTableList_Reps <- rep(list(list()),nreps)
	testSimPhenoValuesTableList_Reps <- rep(list(list()),nreps)
	
	F_Progeny_List_Reps <- rep(list(list()),nreps)
	errorVarList_Reps <- rep(list(list()),nreps)

### Generate base population data for n reps
   
	
    for(nrep in 1:nreps){
  
   		nCondition <- 1
		
### Initiate variables 
	 
   	    genotypicValuesSimList <- list()
		genotypicValuesSimTableList <- list()

		phenotypicValuesSimList <- list()
		phenotypicValuesSimTableList <- list()

		trainIndicesList <- list() 
		testIndicesList <- list() 


		trainGenoTableList <- list()
		trainGenoNewTableList <- list()
		trainGenoNewTableList_w_o_QTL <- list()
		
		testGenoTableList <- list()
		testGenoNewTableList <- list()
		testGenoNewTableList_w_o_QTL <- list()
		
		trainSimPhenoValuesTableList <- list()
		testSimPhenoValuesTableList <- list()
			
		trainSimGenoValuesTableList <- list()
		testSimGenoValuesTableList <- list() 
		
		
		F2_Progeny_List <- list() 
		F3_Progeny_List <- list() 
		F4_Progeny_List <- list() 
		F5_Progeny_List <- list() 
		

		errorVarList <- list() 
		percent_heterozygous_List <- list()
	  
### Loop through QTL and heritability conditions 	
    
	for(numQTL in 1:length(no_QTL)){
  
     for (numH in 1:2){
 #######################################################################
        
		nLoci <- no_QTL[numQTL]
### Define parent genotype ###################################
      
		cPC1 <- sample(c(0,1),nMarkers,replace=TRUE,prob=c(0.1,0.9))
		cPC2 <- cPC1
		
		commonParent <- as.array(cbind(cPC1,cPC2))
		Parent1 <- commonParent

		parent2 <- array(0,c(nCrosses,nMarkers,2)) 
		
		for(i in 1:20){ 
		
			P2C1 <- sample(c(0,1),nMarkers,replace=TRUE,prob=c(0.1,0.9))
			P2C2 <- P2C1
			parent2C <- as.array(cbind(P2C1,P2C2)) 
			parent2[i,,] <- parent2C
		
		}

	  parent2_table <- matrix(rep(0,4289*20),nrow=4289,ncol=20,byrow=TRUE)
	  
	  for(i in 1:20){
	  
		parent2_table[,i] <- parent2[i,,1]+parent2[i,,2]
	  
	  
	  }
	  
	  parent1_table <- Parent1[,1]+Parent1[,2]
	  Parent_table <- cbind(parent1_table,parent2_table)
      colnames(Parent_table) <- c(1:21)


	 if(plotHClust== TRUE){
	  
	    d <- as.matrix(dist(t(Parent_table)))
		dim(d)
		
		png(plotFilename,width=1024,height=768,pointsize=20)
		plotFilename <- paste("Hierarchical Clustering of NAM Families.png")
		Parent_clust <- hclust(as.dist(d))
		plot(Parent_clust,xlab="NAM Families")
		dev.off()
	  }
#### Optional part where you can 
#### Similarity Table

	 if(founderSimilarity==TRUE){

		similarityTable3C<-c()
		intXTable3C <-c()

		fractionVec <- rep(0,20)

		commonParentC <- commonParent[,1] + commonParent[,2]

		for (i in 1:20) { 
		 
		   Parent2C <- parent2[i,,1]+parent2[i,,2]
		   
		   fractionVec[i] <- length(which(Parent2C == commonParentC))/ length(commonParentC)
		 
		}
### Similarity Table among founder parents
		similarityTable3C<-c()
		fractionVec <- rep(0,20)
		for (i in 1:20){ 
		  Parent2C <- parent2[i,,1]+parent2[i,,2]
		  for(j in 1:20){ 
		  
			Parent2IC <- parent2[j,,1]+parent2[j,,2]
			
			fractionVec[j] <- length(which(Parent2C == Parent2IC))/ length(Parent2C)
			
		  }
		 similarityTable3C <- cbind(similarityTable3C,fractionVec)

		}
		dim(similarityTable3C)

}

### Generate 1 Progeny for the first four cycles #############
        
      Cycle_1_Progeny<- array(0,c(nCrosses,nMarkers,2,1))
      Cycle_2_Progeny<- array(0,c(nCrosses,nMarkers,2,nProgeny))
      Cycle_3_Progeny<- array(0,c(nCrosses,nMarkers,2,nProgeny))
      Cycle_4_Progeny<- array(0,c(nCrosses,nMarkers,2,nProgeny))
	    Cycle_5_Progeny<- array(0,c(nCrosses,nMarkers,2,nProgeny))
     
        
      for(i in 1:nCrosses){
     
                
            Parent1 <- commonParent
            Parent2 <- parent2[i,,]
            progeny1<- generateProgeny(Parent1,Parent2,1,NAM_LinkMap_New) 
       
            Cycle_1_Progeny[i,,,] <- progeny1 
            
           
			 
 ######## Generate 100 RILS for each of the twenty families by selfing F1 hybrids ########################### 		       
           
             Parent1<- Cycle_1_Progeny[i,,,1]  
             Parent2<- Cycle_1_Progeny[i,,,1] 
             progeny1<- generateProgeny(Parent1,Parent2,nProgeny,NAM_LinkMap_New) 
             Cycle_2_Progeny[i,,,] <- progeny1 
                 
             F2_Progeny_List[[nrep]] <-  Cycle_2_Progeny
			 
 ######## Generate 100 RILS for each of the twenty families by selfing generation 2 ########################### 
				for(m in 1:nProgeny){ 
				
				  Parent1<- Cycle_2_Progeny[i,,,m]  
				  Parent2<- Cycle_2_Progeny[i,,,m] 
				  progeny1<- generateProgeny(Parent1,Parent2,1,NAM_LinkMap_New) 
				  Cycle_3_Progeny[i,,,m] <- progeny1 
				}
				F3_Progeny_List[[nrep]] <-  Cycle_3_Progeny
						
######## Generate 100 RILS for each of the twenty families by selfing generation 3 ########################### 
				for(m in 1:nProgeny){ 
				
				  Parent1 <- Cycle_3_Progeny[i,,,m]  
				  Parent2 <- Cycle_3_Progeny[i,,,m] 
				  progeny1<- generateProgeny(Parent1,Parent2,1,NAM_LinkMap_New) 
				  Cycle_4_Progeny[i,,,m] <- progeny1 
				}
				 F4_Progeny_List[[nrep]] <-  Cycle_4_Progeny
			
######## Generate 100 RILS for each of the twenty families by selfing generation 4 ########################### 
			
				for(m in 1:nProgeny){ 
		  
				  Parent1<- Cycle_4_Progeny[i,,,m]
				  Parent2<- Cycle_4_Progeny[i,,,m]
			
				  Cycle_5_Progeny[i,,,m] <- generateProgeny(Parent1,Parent2,1,NAM_LinkMap_New) 
			
				}
				
			}
				
				F5_Progeny_List[[nrep]] <-   Cycle_5_Progeny
		
				F_Progeny <-  F5_Progeny_List[[nrep]]

############################################################################################
### Genotype Table in GenSel and rrBLUP format (genotypes coded as -1,0,1)
     
	  genoTable <- generateMinus101GenoFormat(F_Progeny,no_selected,nProgeny)
	  dim(genoTable)
	       
	  genoTable_Mod <- apply(genoTable,2,as.numeric)
	  genoTable_New <- generateWeightedGenoTable(genoTable_Mod,nLoci)

	  genoSimValues <- simulateGenotypicValues(genoTable_Mod,nLoci)
	  
#########################################################################
      genoVar <- var(genoSimValues)
	    heritability <- h2[numH]
      errorVar <- ((1-heritability) * genoVar)/(heritability)
      errorSD <- sqrt(errorVar)
      varE <-  errorVar
##########################################################################
	    phenoSimValues <- simulatePhenotypicValues(genoTable_Mod,varE,nLoci)
           
   
############################# ColNames  #######################################################       
    nLines <- nCrosses*nProgeny 
    LineId <-rep(0,(nCrosses*nProgeny))
    for(i in 1:(nCrosses*nProgeny)){ 
        
        LineId[i]<- paste("Line",i,sep="")
        
    }
#### RowNames # IndividualNames <- c("Animal_ID",IndividualNames)

    markerId <-rep(0,nMarkers) 
    for(i in 1:(nMarkers)){  
        
        markerId[i]<- paste("m",i,sep="")
        
    }

#### Table for Simulated Phenotype ###########################################
    names(phenoSimValues)<- LineId
    Mean_Fixed<- rep(1,nLines)
    phenotypicValuesSimTable <-cbind(phenoSimValues,Mean_Fixed) 
      
#### Table for Simulated Genotypic Values ####################################
      
    names(genoSimValues)<- LineId
    Mean_Fixed<- rep(1,nLines)
    genotypicValuesSimTable <- cbind(genoSimValues,Mean_Fixed)
    
#### Indices for geno and pheno tables 
	
##### Get family-wise genotable list 

     genoTable_Mod_List <- list()
	 genoTable_New_List <- list()
	 phenoSimValues_List <- list() 
	 genoSimValues_List <- list()
	 phenotypicValuesSimTable_List <- list()
	 genotypicValuesSimTable_List <- list()
	 
	 
     nFamilies <- 20
     nProgeny_per_Family <- 100
     initIndex <- 1
	 finalIndex <- nProgeny_per_Family
	 
    for(nFamily in 1:nFamilies){
	
	    genoTable_Mod_List[[nFamily]] <- genoTable_Mod[initIndex:finalIndex,]
		genoTable_New_List[[nFamily]] <- genoTable_New[initIndex:finalIndex,]
		phenotypicValuesSimTable_List[[nFamily]] <- phenotypicValuesSimTable[initIndex:finalIndex,]
		genotypicValuesSimTable_List[[nFamily]] <- genotypicValuesSimTable[initIndex:finalIndex,]
		phenoSimValues_List[[nFamily]] <- phenoSimValues[initIndex:finalIndex]
	    genoSimValues_List[[nFamily]] <- genoSimValues[initIndex:finalIndex]
		
		initIndex <- finalIndex +1 
		finalIndex <- finalIndex + nProgeny_per_Family
	}


##
    trainIndices_List <- list()
	testIndices_List <- list()

    trainGenoTable_List <- list()
    trainGenoNewTable_List <- list()
	trainGenoNewTable_List_w_o_QTL <- list()
    trainSimPhenoValuesTable_List <- list()
    trainSimGenoValuesTable_List <- list()
	
	testGenoTable_List <- list()
    testGenoNewTable_List <- list()
	testGenoNewTable_List_w_o_QTL <- list()
    testSimPhenoValuesTable_List <- list()
    testSimGenoValuesTable_List <- list()
	
	
	QTL_Indices <- getQTLIndices(nLoci)

	
## (80-20) data split for training and test data set for training GS models   
   
   for(nFamily in 1:nFamilies){
    nLines_per_Family <- 100
    initseed <- 25+nrep 
	set.seed(initseed)
	indices<-c(1:nLines_per_Family)
    trainIndices <-sample(c(1:nLines_per_Family),(0.8*nLines_per_Family)) 
    testIndices <- indices[which(!indices %in% trainIndices)] 
     
    trainIndices_List[[nFamily]] <- trainIndices
    testIndices_List[[nFamily]]	<- testIndices
#### Training Set table for Genotype and Simulated Phenotype ################### 
      
    trainGenoTable_List[[nFamily]] <- genoTable_Mod_List[[nFamily]][trainIndices,] 
    trainGenoNewTable_List[[nFamily]] <- genoTable_New_List[[nFamily]][trainIndices,] 
	trainGenoNewTable_List_w_o_QTL[[nFamily]] <- genoTable_New_List[[nFamily]][trainIndices,-c(QTL_Indices)] 
	
    trainSimPhenoValuesTable_List[[nFamily]] <- phenotypicValuesSimTable_List[[nFamily]][trainIndices,] 
    trainSimGenoValuesTable_List[[nFamily]] <-  genotypicValuesSimTable_List[[nFamily]][trainIndices,]
	
	     
#### Validation Set table for Genotype and Simulated Phenotype ###################
      
    testGenoTable_List[[nFamily]]<- genoTable_Mod_List[[nFamily]][testIndices,] 
    testGenoNewTable_List[[nFamily]] <- genoTable_New_List[[nFamily]][testIndices,]
    testGenoNewTable_List_w_o_QTL[[nFamily]] <- genoTable_New_List[[nFamily]][testIndices,-c(QTL_Indices)]
	testSimPhenoValuesTable_List[[nFamily]]  <- phenotypicValuesSimTable_List[[nFamily]][testIndices,]  
    testSimGenoValuesTable_List[[nFamily]] <-  genotypicValuesSimTable_List[[nFamily]][testIndices,]
 
  }
#### List of Tables for nCondition (combination of QTL & h2 condition) #######################################     
    
	phenotypicValuesSimTableList[[nCondition]] <- phenotypicValuesSimTable_List
    genotypicValuesSimTableList[[nCondition]] <- genotypicValuesSimTable_List
	
    phenotypicValuesSimList[[nCondition]] <- phenoSimValues_List
    genotypicValuesSimList[[nCondition]] <- genoSimValues_List
    errorVarList[[nCondition]] <- errorVar
      
    trainIndicesList[[nCondition]] <- trainIndices_List
    testIndicesList[[nCondition]] <- testIndices_List
      
      
    trainGenoTableList[[nCondition]] <- trainGenoTable_List
    trainGenoNewTableList[[nCondition]] <- trainGenoNewTable_List
	trainGenoNewTableList_w_o_QTL[[nCondition]] <- trainGenoNewTable_List_w_o_QTL
	
    trainSimPhenoValuesTableList[[nCondition]] <- trainSimPhenoValuesTable_List
    trainSimGenoValuesTableList[[nCondition]] <- trainSimGenoValuesTable_List
      
    testGenoTableList[[nCondition]] <- testGenoTable_List
    testGenoNewTableList[[nCondition]] <- testGenoNewTable_List
	testGenoNewTableList_w_o_QTL[[nCondition]] <- testGenoNewTable_List_w_o_QTL
	
	
    testSimPhenoValuesTableList[[nCondition]] <- testSimPhenoValuesTable_List
    testSimGenoValuesTableList[[nCondition]] <- testSimGenoValuesTable_List
      
    F_Progeny_List[[nCondition]] <- F_Progeny

      
    nCondition<-nCondition+1
   
    }
   }
	
	
	phenotypicValuesSimTableList_Reps[[nrep]] <- phenotypicValuesSimTableList
	genotypicValuesSimTableList_Reps[[nrep]] <-  genotypicValuesSimTableList
	
	phenotypicValuesSimList_Reps[[nrep]]<- phenotypicValuesSimList
	genotypicValuesSimList_Reps[[nrep]] <- genotypicValuesSimList
	
	
	trainIndicesList_Reps[[nrep]] <- trainIndicesList 
	testIndicesList_Reps[[nrep]]<- testIndicesList

	trainGenoTableList_Reps[[nrep]] <- trainGenoTableList
	trainGenoNewTableList_Reps[[nrep]] <- trainGenoNewTableList
	trainGenoNewTableList_w_o_QTL_Reps[[nrep]] <- trainGenoNewTableList_w_o_QTL
	trainSimPhenoValuesTableList_Reps[[nrep]] <- trainSimPhenoValuesTableList
	trainSimGenoValuesTableList_Reps[[nrep]] <- trainSimGenoValuesTableList

	
	testGenoTableList_Reps[[nrep]] <- testGenoTableList
	testGenoNewTableList_Reps[[nrep]] <- testGenoNewTableList
	testGenoNewTableList_w_o_QTL_Reps[[nrep]] <- testGenoNewTableList_w_o_QTL
	testSimGenoValuesTableList_Reps[[nrep]] <- testSimGenoValuesTableList
	testSimPhenoValuesTableList_Reps[[nrep]] <- testSimPhenoValuesTableList
    
	F_Progeny_List_Reps[[nrep]] <- F_Progeny_List
	errorVarList_Reps[[nrep]] <- errorVarList

  }
 
 

	return(list(genotypicValuesSimTableList_Reps,phenotypicValuesSimTableList_Reps,genotypicValuesSimList_Reps,phenotypicValuesSimList_Reps,trainIndicesList_Reps,testIndicesList_Reps,trainGenoTableList_Reps,trainGenoNewTableList_Reps,trainGenoNewTableList_w_o_QTL_Reps, testGenoTableList_Reps,testGenoNewTableList_Reps,testGenoNewTableList_w_o_QTL_Reps,trainSimGenoValuesTableList_Reps,trainSimPhenoValuesTableList_Reps,testSimGenoValuesTableList_Reps,testSimPhenoValuesTableList_Reps,F_Progeny_List_Reps,errorVarList_Reps))

}


###################################################################################################################
# nReps <- 1
# NAMBasePopulationData_List <- getBasePopulation_FamilyWise_List(nReps)

  
```
