## Supplementary material for "Factors Affecting Response to Recurrent Genomic Selection in Soybeans": File S1: getGInd_PGStatsOutput.R

```
getGIndData <- function(ncon,nReps,ModelType_List){
  
  nMarkers <- 4289
  h2 <- c(0.7,0.3,0.7,0.3,0.7,0.3)
  no_QTL <- c(40,40,400,400,nMarkers,nMarkers)
  nCycles <- 40
 
  Default <- TRUE
  NAM_LinkMap_New <- getNAMLinkageMapTable()
 
  nMethods <- length(ModelType_List)
  
  nRep <- nReps
  nCon <- ncon
  trainWindowSize <- 14
  
  
  for(nMet in 1:nMethods){
    
    ModelType <- ModelType_List[[nMet]]
    
    GenTable_AF_List_Cyc <- list()
    
    for(nCyc in 1:nCycles){
      
      GenTable_AF_List <- list()
      
      for(nrep in 1:nRep){
        
        ## Read Geno Table
        
        if(nCyc ==1){
          filename <- paste("PM_",ModelType,"_",no_QTL[nCon],"_",h2[nCon],"_GenoDataInd_Rep",nrep,"_Cyc_",nCyc,sep="")
          GenTable_AF_List[[nrep]] <- read.table(filename,sep="\t")
        }else if(nCyc >1){
          filename <- paste("PM_",ModelType,"trainTable_Cyc_",trainWindowSize,"_",no_QTL[nCon],"_",h2[nCon],"_GenoDataInd_Rep",nrep,"_Cyc_",nCyc,sep="")
          GenTable_AF_List[[nrep]] <- read.table(filename,sep="\t")
        }
        
      }
      
      GenTable_AF_List_Cyc[[nCyc]] <- GenTable_AF_List
    }
    
  
##########
    GS_GenTable_AFList_Cyc[[nMet]] <- GenTable_AF_List_Cyc
   
 }
  
  return(GS_GenTable_AFList_Cyc)
}


get_PG_LD_StatsGSMethods<- function(GS_GenTable_AFList_Cyc,ncon,nReps,ncycles){
  
  nMarkers <- 4289
  h2 <- c(0.7,0.3,0.7,0.3,0.7,0.3)
  no_QTL <- c(40,40,400,400,nMarkers,nMarkers)
  nCycles <- ncycles
  MarkerEffects_Reps <- list()
  nMethods <- length(GS_GenTable_AFList_Cyc)
  nCon <- ncon
  i <- nCon
  nRep <- nReps
  nreps <- nReps
  
  nCycles <- ncycles
  selectionOnGeno <- FALSE
  noQTLs <- no_QTL[nCon]
  
  RRREML_Model <- GSModels_List[[1]]
  
  alleleConversionTable_Combined <- AlleleConversionTable_Combined
  nextGenGenoTable <- list()
  nextGenGenoTable_List <- list()
  
  for(nMet in 1:nMethods){
    
    ModelType <- ModelType_List[[nMet]]
    
    for(nrep in 1:nRep){
      
      PredictionModel_Pheno <- RRREML_Model[[2]][[i]][[nrep]]
      PredictionModel_Geno <- RRREML_Model[[1]][[i]][[nrep]]
      
      if(ModelType =="RRREML" || ModelType =="PS" || ModelType=="RRBLUP_REML"){
        if(selectionOnGeno==TRUE){
          MarkerEffects <- unlist(PredictionModel_Geno[1])
        }else if(selectionOnGeno==FALSE){
          MarkerEffects <- unlist(PredictionModel_Pheno[1])
        }
      }
      
      MarkerEffects_Reps[[nrep]] <- MarkerEffects
    }
    
    GenTable_AF_List_Cyc <- GS_GenTable_AFList_Cyc[[nMet]]
    
    nextGenGenoTable <- foreach(nCyc=1:nCycles) %:% foreach(k=1:nRep) %dopar% (SoyNAMPredictionMethods::getNumericFormat(GenTable_AF_List_Cyc[[nCyc]][[k]][,-1],alleleConversionTable_Combined))
   
   
    noQTLs <- no_QTL[nCon]
    nCyc <-1
    GenTable_AF_List <- GenTable_AF_List_Cyc[[nCyc]]
     
    nextGenGenoTable <- foreach(k=1:nReps) %dopar% (SoyNAMPredictionMethods::getNumericFormat(GenTable_AF_List_Cyc[[nCyc]][[k]][,-1],alleleConversionTable_Combined))
  
    PGStats <- foreach(k=1:nReps) %dopar% (SoyNAMPredictionMethods::getPGStatsGSMethods_Cyc1(GenTable_AF_List_Cyc[[nCyc]][[k]][,-1],nextGenGenoTable[[k]],populations,MarkerEffects_Reps[[k]],noQTLs,gIndObjects,ModelType,nCon,k,nCyc))
    PGStats_List[[nCyc]] <- PGStats
  
    rm(PGStats)
  
  
  
    nextGenGenoTable <- foreach(nCyc=2:nCycles) %:% foreach(k=1:nReps) %dopar% (SoyNAMPredictionMethods::getNumericFormat(GenTable_AF_List_Cyc[[nCyc]][[k]][,-1],alleleConversionTable_Combined))
    #nextGenGenoTable <- lapply(nextGenGenoTable_Numeric,function(x) apply(x,2,as.numeric))
  
    PGStats <- foreach(nCyc=2:nCycles) %:% foreach(k=1:nReps) %dopar% (SoyNAMPredictionMethods::getPGStatsGSMethods(GenTable_AF_List_Cyc[[nCyc]][[k]][,-1],nextGenGenoTable[[nCyc]][[k]],populations,MarkerEffects_Reps[[k]],PGStats_List[[1]][[k]][[2]],noQTLs,gIndObjects,ModelType,nCon,k,nCyc))
    
    LDStats <- foreach(nCyc=1:nCycles) %:% foreach(nrep=1:nReps) %dopar% tryCatch(SoyNAMPredictionMethods::getLDTable_GSMethods(nextGenGenoTable[[nCyc]][[nrep]],noQTLs,ModelType,nCon,nrep,nCyc),error=function(e) print(paste("Error LDStats Cycle No",nCyc,e)))       
    
    
    PGStats_List[[nCyc]] <- PGStats
    LDStats_List[[nCyc]] <- LDStats
   
    
    rm(PGStats)
    
    
    GS_PGStats_List_SC[[nMet]] <- PGStats_List
    GS_LDStats_List_SC[[nMet]] <- PGStats_List
   
  }

   return(list(GS_PGStats_List_SC,GS_LDStats_List_SC))
}
 

getPGStatsOutput <- function(GS_PGStats_List_SC,ncon,nreps,ncycles){ 
  
  nCycles <- ncycles
  nRep <- nreps
  
  nSelCriteria <- length(GS_PGStats_List_SC)
  
  Diff_Stats_List_Local <- list()
  Diff_Stats_List_Global <- list()
  
  Diff_Stats_List_Local_SC <- list()
  Diff_Stats_List_Global_SC <- list()
  
  F <-  rep(list(rep(list(list()),nRep)),nCycles)
  ExpHet <-  rep(list(rep(list(list()),nRep)),nCycles)
  Freq_GInd <- rep(list(rep(list(list()),nRep)),nCycles)
  MAF <- rep(list(rep(list(list()),nRep)),nCycles)
  
  F_List <-  rep(list(rep(list(rep(list(list()),nRep)),nCycles)),nSelCriteria)
  ExpHet_List <-  rep(list(rep(list(rep(list(list()),nRep)),nCycles)),nSelCriteria)
  Freq_GInd_List <- rep(list(rep(list(rep(list(list()),nRep)),nCycles)),nSelCriteria)
  MAF_List <- rep(list(rep(list(rep(list(list()),nRep)),nCycles)),nSelCriteria)
  
 
  
  for(nSC in 1:nSelCriteria){	
    
    PGStats_List <- GS_PGStats_List_SC[[nSC]] 
    
    for(nCyc in 1:nCycles){ 
      
      for(nrep in 1:nRep){
        
        F[[nCyc]][[nrep]] <-  PGStats_List[[nCyc]][[nrep]][[8]]
        ExpHet[[nCyc]][[nrep]] <-  PGStats_List[[nCyc]][[nrep]][[9]] 
        Freq_GInd[[nCyc]][[nrep]] <- PGStats_List[[nCyc]][[nrep]][[10]]
        MAF[[nCyc]][[nrep]] <- PGStats_List[[nCyc]][[nrep]][[11]] 
        
        
      }
    } 
    
    F_List[[nSC]] <- F
    ExpHet_List[[nSC]] <- ExpHet
    Freq_GInd_List[[nSC]] <- Freq_GInd
    MAF_List[[nSC]] <- MAF
    
  }
  
  
  return(list(F_List,ExpHet_List,Freq_GInd_List,MAF_List))
}


# # PGStats_List_14Cyc <- list(PS_PGStats_List,RR1X_14_PGStats_List,BB1X_14_PGStats_List,BL1X_14_PGStats_List,SVMRBF1X_14_PGStats_List) 

# # ncon <- 3
# # reps <- 10 
# # ncycles <- 40
# # PGStats_Out <- getPGStatsOutput(PGStats_List_14Cyc,ncon,nreps,ncycles)                                       
# # ###########  


getLDStats_Mat_Avg <- function(LDStats_List,nCyc,nReps){

  
  
  LDStats <- LDStats_List[[nCyc]]
  LDStats_Mat_D <- melt(LDStats[[1]][[1]][[1]])
  LDStats_Mat_D1 <- melt(LDStats[[1]][[1]][[2]])
  LDStats_Mat_R2 <- melt(LDStats[[1]][[1]][[3]])
  
  
  LDStats_List_Comb_D <- split(LDStats_Mat_D,LDStats_Mat_D$Var2) 
  LDStats_List_Comb_D1 <- split(LDStats_Mat_D1,LDStats_Mat_D1$Var2) 
  LDStats_List_Comb_R2 <- split(LDStats_Mat_R2,LDStats_Mat_R2$Var2)
  
  for(nrep in 2:nReps){
    
    
    LDStats_Mat_Melt_D <- melt(LDStats[[nrep]][[1]][[1]]) 
    LDStats_Mat_Melt_D1 <- melt(LDStats[[nrep]][[1]][[2]]) 
    LDStats_Mat_Melt_R2 <- melt(LDStats[[nrep]][[1]][[3]]) 
    
    LDStats_List_D <- split(LDStats_Mat_Melt_D,LDStats_Mat_Melt_D$Var2) 
    LDStats_List_D1 <- split(LDStats_Mat_Melt_D1,LDStats_Mat_Melt_D1$Var2) 
    LDStats_List_R2 <- split(LDStats_Mat_Melt_R2,LDStats_Mat_Melt_R2$Var2) 
    
    
    if(nrep>=2){
      b_D <- mapply(function(X,Y){(X[,3])+(Y[,3])},X=LDStats_List_Comb_D,Y=LDStats_List_D)
      b_D1 <- mapply(function(X,Y){(X[,3])+(Y[,3])},X=LDStats_List_Comb_D1,Y=LDStats_List_D1)
      b_R2 <- mapply(function(X,Y){(X[,3])+(Y[,3])},X=LDStats_List_Comb_R2,Y=LDStats_List_R2)
    }
    
    LDStats_Mat_Total_D <-  b_D
    LDStats_Mat_Total_D1 <-  b_D1
    LDStats_Mat_Total_R2 <-  b_R2
    
    LDStats_Mat_Total_Melt_D <- melt(LDStats_Mat_Total_D)
    LDStats_Mat_Total_Melt_D1 <- melt(LDStats_Mat_Total_D1)
    LDStats_Mat_Total_Melt_R2 <- melt(LDStats_Mat_Total_R2) 
    
    LDStats_List_Comb_D <- split(LDStats_Mat_Total_Melt_D,LDStats_Mat_Total_Melt_D[,2]) 
    LDStats_List_Comb_D1 <- split(LDStats_Mat_Total_Melt_D1,LDStats_Mat_Total_Melt_D1[,2])
    LDStats_List_Comb_R2 <- split(LDStats_Mat_Total_Melt_R2,LDStats_Mat_Total_Melt_R2[,2])
    
    
    if(nrep==nReps){
      LDStats_Mat_Avg_D <- apply(LDStats_Mat_Total_D,2,function(x) x/nReps)
      LDStats_Mat_Avg_D1 <- apply(LDStats_Mat_Total_D1,2,function(x) x/nReps)
      LDStats_Mat_Avg_R2 <- apply(LDStats_Mat_Total_R2,2,function(x) x/nReps)
    }
    
  }
  return(list(LDStats_Mat_Avg_D,LDStats_Mat_Avg_D1,LDStats_Mat_Avg_R2))
}

#getLDStats_Mat_Avg(LDStats,nCyc,nReps)
#######################################################################


getLDPlotStats <- function(GS_LDStats_List_SC,ncon,nReps,ncycles,Methods,nsel,NAM_LINKAGE_MAP){

  
  nRep <- nReps
  nreps <- nReps
  nCon <- ncon
  nCycles <- ncycles
  
  Cycles <- c(1:nCycles)
  nCycles <- length(Cycles)
  nMarkers <- 4289
    
  NAM_Table <- NAM_LINKAGE_MAP
  dim(NAM_Table)
  
  Marker_Marker_cM_List_Cyc_Met <- list()
  Marker_Marker_BP_List_Cyc_Met <- list()
  Marker_Marker_LD_List_D_Cyc_Met <- list()
  Marker_Marker_LD_List_D1_Cyc_Met <- list()
  Marker_Marker_LD_List_R2_Cyc_Met <- list()
  
  
  Methods <- methods
  nMethods <- length(Methods)
  nSel <- nsel
  
  for(nMet in 1:nMethods){
     Met_LDStats <- GS_LDStats_List_SC[[nMet]]
    
     LDStats_List_Cyc <- list()
  
     for(nCyc in 1:(length(Cycles))){ 
    
        LDStats_List_Cyc[[nCyc]] <- (getLDStats_Mat_Avg(Met_LDStats,nCyc,nReps))
    
     }
    
   ###### Marker-marker LD Table ###########
  
   Marker_Marker_cM_List_Cyc <- list()
   Marker_Marker_BP_List_Cyc <- list()
  
   Marker_Marker_LD_List_Cyc <- list()
   Marker_Marker_LD_List_D_Cyc <- list()
   Marker_Marker_LD_List_D1_Cyc <- list()
   Marker_Marker_LD_List_R2_Cyc <- list()
  
   Marker_Marker_cM_Table <- matrix(rep(0,nMarkers*nMarkers),nrow=nMarkers,ncol=nMarkers)
  
   Marker_Indices_Chr <- NAM_Table[,3]  
  
   for(nCyc in 1:(length(Cycles))){
    
    
    Marker_Marker_LD_List_D <- list() 
    Marker_Marker_LD_List_D1 <- list() 
    Marker_Marker_LD_List_R2 <- list() 
    Marker_Marker_cM_List <- list()
    Marker_Marker_BP_List <- list()
    
    for(nMarker in 1:nMarkers){ 
      
      Marker_ID <- NAM_Table[nMarker,2]
      chr <- Marker_Indices_Chr[nMarker]
      
      LD_Table_Chr_D <- LDStats_List_Cyc[[nCyc]][[1]][nMarker,which(as.character(NAM_Table[,3]) %in% as.character(chr))]
      LD_Table_Chr_D1 <- LDStats_List_Cyc[[nCyc]][[2]][nMarker,which(as.character(NAM_Table[,3]) %in% as.character(chr))]
      LD_Table_Chr_R2 <- LDStats_List_Cyc[[nCyc]][[3]][nMarker,which(as.character(NAM_Table[,3]) %in% as.character(chr))]
      
      NAM_Table_Chr <- NAM_Table[which(as.character(NAM_Table[,3]) %in% as.character(chr)),] 
      Marker_Marker_Chr_Index <- which(NAM_Table_Chr[,2] %in% Marker_ID)
      nMarkers_in_Chr <- nrow(NAM_Table_Chr)
      
      Marker_marker_cM <- rep(0,nMarkers_in_Chr)
      Marker_marker_BP <- rep(0,nMarkers_in_Chr) 
      
      Marker_Marker_LD_D <- rep(0,nMarkers_in_Chr)
      Marker_Marker_LD_D1 <- rep(0,nMarkers_in_Chr)
      Marker_Marker_LD_R2 <- rep(0,nMarkers_in_Chr)
      
      
      if(Marker_Marker_Chr_Index != nMarkers_in_Chr){ 
        
        for(nMarker_pos in Marker_Marker_Chr_Index:nMarkers_in_Chr){ 
          
          Marker_pos <- NAM_Table_Chr[Marker_Marker_Chr_Index,6]
          marker_cM <- NAM_Table_Chr[nMarker_pos,6]
          
          Marker_Dist_pos <-  NAM_Table_Chr[Marker_Marker_Chr_Index,14]
          marker_BP <- NAM_Table_Chr[nMarker_pos,14]
          
          Marker_marker_cM[nMarker_pos] <- marker_cM - Marker_pos  
          Marker_marker_BP[nMarker_pos] <- marker_BP- Marker_Dist_pos
          
          
          Marker_Marker_LD_D[nMarker_pos] <- LD_Table_Chr_D[nMarker_pos]
          Marker_Marker_LD_D1[nMarker_pos] <- LD_Table_Chr_D1[nMarker_pos]
          Marker_Marker_LD_R2[nMarker_pos] <- LD_Table_Chr_R2[nMarker_pos]
          
        } 
      }else if(Marker_Marker_Chr_Index != 1){ 
        
        for(nMarker_neg in (Marker_Marker_Chr_Index-1):1){ 
          
          Marker_neg <- NAM_Table_Chr[Marker_Marker_Chr_Index,6]
          marker_cM <- NAM_Table_Chr[nMarker_neg,6]
          
          Marker_Dist_neg <-  NAM_Table_Chr[Marker_Marker_Chr_Index,14]
          marker_BP <- NAM_Table_Chr[nMarker_neg,14]
          
          Marker_marker_cM[nMarker_neg] <- Marker_neg - marker_cM 
          Marker_marker_BP[nMarker_neg] <- Marker_Dist_neg - marker_BP
          
          Marker_Marker_LD_D[nMarker_neg] <- LD_Table_Chr_D[nMarker_neg]
          Marker_Marker_LD_D1[nMarker_neg] <- LD_Table_Chr_D1[nMarker_neg]
          Marker_Marker_LD_R2[nMarker_neg] <- LD_Table_Chr_R2[nMarker_neg]
        }  
      } 
      
      Marker_Marker_cM_List[[nMarker]] <- Marker_marker_cM
      Marker_Marker_BP_List[[nMarker]] <- Marker_marker_BP
      Marker_Marker_LD_List_D[[nMarker]] <- Marker_Marker_LD_D
      # Marker_Marker_LD_List_D1[[nMarker]] <- Marker_Marker_LD_D1
      # Marker_Marker_LD_List_R2[[nMarker]] <- Marker_Marker_LD_R2 
      
    }
    
    Marker_Marker_cM_List_Cyc[[nCyc]] <-  Marker_Marker_cM_List
    Marker_Marker_BP_List_Cyc[[nCyc]] <-  Marker_Marker_BP_List
    Marker_Marker_LD_List_D_Cyc[[nCyc]] <- Marker_Marker_LD_List_D
    # Marker_Marker_LD_List_D1_Cyc[[nCyc]] <- Marker_Marker_LD_List_D1
    # Marker_Marker_LD_List_R2_Cyc[[nCyc]] <- Marker_Marker_LD_List_R2
    
  }
  
  Marker_Marker_cM_List_Cyc_Met[[nMet]] <-  Marker_Marker_cM_List_Cyc
  Marker_Marker_BP_List_Cyc_Met[[nMet]] <-  Marker_Marker_BP_List_Cyc
  Marker_Marker_LD_List_D_Cyc_Met[[nMet]] <- Marker_Marker_LD_List_D_Cyc
  # Marker_Marker_LD_List_D1_Cyc_Met[[nMet]] <- Marker_Marker_LD_List_D1_Cyc
  # Marker_Marker_LD_List_R2_Cyc_Met[[nMet]] <- Marker_Marker_LD_List_R2_Cyc
  
}

return(list(Marker_Marker_cM_List_Cyc_Met,Marker_Marker_BP_List_Cyc_Met,Marker_Marker_LD_List_D_Cyc_Met))

}


#########


    
    
  
  
```
