## Supplementary material for "Factors Affecting Response to Recurrent Genomic Selection in Soybeans": File S1: getLDStats_GSMethods.R

```
 
 getNumericFormat_012_LD <- function(GenoTable_AlleleFormat,AlleleConversionTable_Combined){

        options(warn=-1)
        genoTable <- apply(GenoTable_AlleleFormat,2,as.character)
        nIndividuals <- nrow(genoTable)

        IA3023_alleles <- as.character(AlleleConversionTable_Combined[,8])
        Alt_Homozygous_alleles <- as.character(AlleleConversionTable_Combined[,10])
        Het_alleles <- as.character(AlleleConversionTable_Combined[,22])

        IA3023_alleles_Indices <- apply(genoTable,1,function(x) which(as.character(x) == (IA3023_alleles)))

        Het_alleles_Indices <-  apply(genoTable,1,function(x) if(length(which( as.character(x)==(Het_alleles)))!=0){which(as.character(x) == (Het_alleles))} else{0})
         
        Alt_Homozygous_alleles_Indices <- apply(genoTable,1,function(x) which(as.character(x) == (Alt_Homozygous_alleles)))
		
		
				
		for(i in 1:nIndividuals){

           if(length(IA3023_alleles_Indices[[i]]) >1){
                genoTable[i,(IA3023_alleles_Indices[[i]])] <- "2"
           }

           if(length(Het_alleles_Indices[[i]]) >1 & Het_alleles_Indices[[i]] !=0){
                genoTable[i,(Het_alleles_Indices[[i]])] <- "1"
           }
           if(length(Alt_Homozygous_alleles_Indices[[i]]) >1){

                genoTable[i,(Alt_Homozygous_alleles_Indices[[i]])] <- "0"
           }
        }
		
	    genoTable_Num <- apply(genoTable,2,as.numeric)
        return(genoTable_Num)
 }


getLDTable_GSMethods <- function(nextGenGenoTable_NumF,no_QTL,ModelType,nCon,nrep,nCyc){ 

    nextGenGenoTable <- (nextGenGenoTable_NumF)
	noQTL <- no_QTL
	LD_List <-  getLD(nextGenGenoTable,noQTL)
		
    LD_GenoTable <-  LD_List[[1]]
    LD_QTL_GenoTable <- LD_List[[2]]
	rm(LD_List)
	  	
	return(list(LD_GenoTable,LD_QTL_GenoTable))
		

}


getFreq_Alleles_LD <- function(GenoTable){


	nMarkers <- dim(GenoTable)[2]
	nIndividuals <- dim(GenoTable)[1]
	Freq_1 <- rep(0,nMarkers)
	Freq_0 <- rep(0,nMarkers)
	count_1 <-rep(0,nMarkers)
	count_0 <-rep(0,nMarkers)
	n_alleles <- 2*nIndividuals

	for(j in 1:nMarkers){

		p_1_P <- length(which(GenoTable[,j]==1))
		p_0_H <- length(which(GenoTable[,j]==0))
		p_minus1_Q <- length(which(GenoTable[,j]==-1))

		count_1[j] <- ((2*p_1_P)+p_0_H )
		count_0[j] <- ((2*p_minus1_Q)+p_0_H )

		Freq_1[j] <- count_1[j]/n_alleles
		Freq_0[j] <- count_0[j]/n_alleles
	 }
 
	return(list(Freq_1,Freq_0))

}


getLD <- function(nextGenGenoTable,noQTL){ 

  nLines <- nrow(nextGenGenoTable)
  nMarkers <- ncol(nextGenGenoTable) 
 
 
  Allele_Freq <- getFreq_Alleles_LD(nextGenGenoTable)
  OneAllele_Freq  <- Allele_Freq[[1]]
  ZeroAllele_Freq  <- Allele_Freq[[2]]

  nQTL <- noQTL
  
  p1 <- OneAllele_Freq 
  q1 <- ZeroAllele_Freq
  a <- c(1:length(p1)) 
  Pro <- nextGenGenoTable
  D <- matrix(rep(0,nMarkers*nMarkers),nrow=nMarkers,ncol=nMarkers) 
  R2 <- matrix(rep(0,nMarkers*nMarkers),nrow=nMarkers,ncol=nMarkers)
  D1 <- matrix(rep(0,nMarkers*nMarkers),nrow=nMarkers,ncol=nMarkers)

  QTLIndices <- getQTLIndices(nQTL)
 
### 
 
  D_Geno <- LD_V3(p1,q1,a,Pro)
  #D_Geno <- LD(p1,q1,a,Pro,D,D1,R2)
  D_QTL_Geno <- list(D_Geno[[1]][QTLIndices,],D_Geno[[2]][QTLIndices,],D_Geno[[3]][QTLIndices,])
   
  
  return(list(D_Geno,D_QTL_Geno))
}


 
LD_V3 <- function(p1,q1,a,pro){

 D <- matrix(0,nrow=4289,ncol=4289)
 D1 <- matrix(0,nrow=4289,ncol=4289)
 R2 <- matrix(0,nrow=4289,ncol=4289)
 
#  pa1a1b1b1 = 0
#  pa1a1b1b2 = 0
#  pa1a2b1b2 = 0
#  pa1a2b1b1 = 0

 for(i in 1:4289){ 
 
  for(j in i:4289){
     
	b <- length(pro[,i])
	pa1a1b1b1 <- length(which(pro[,i]==1 & pro[,j]==1))
	pa1a1b1b2 <- length(which(pro[,i]==1 & pro[,j]==0))
	pa1a2b1b2 <- length(which(pro[,i]==0 & pro[,j]==0))
	pa1a2b1b1 <- length(which(pro[,i]==0 & pro[,j]==1))
	
    D[i,j] <- (((2*pa1a1b1b1)+pa1a1b1b2+(pa1a2b1b2/2)+pa1a2b1b1)/(2*b)) - (p1[i]*p1[j])
    
	if(D[i,j] > 0){ 
       PQ1 <- p1[i]*q1[j]
       PQ2 <- q1[i]*p1[j] 
       Dmax <- min(PQ1,PQ2)
	   if(Dmax !=0){
          D1[i,j] <- D[i,j]/Dmax 
		} else if(Dmax ==0){ 
		  D1[i,j] <- 999 
		}
	}
	if(D[i,j] < 0){
         PQ1 <- p1[i]*p1[j]
         PQ2 <- q1[i]*q1[j]
         Dmax <- min(PQ1,PQ2)
		if(Dmax !=0){
          D1[i,j] <- D[i,j]/Dmax 
		} else if(Dmax ==0){ 
		  D1[i,j] <- 999 
		}
    } 
	if(D[i,j]==0){ 
		D1[i,j] <- D[i,j]
	}
	
	R2_denom <- (p1[j]*p1[i]*q1[j]*q1[i])
	if(R2_denom !=0){
		R2[i,j] <- (D[i,j]*D[i,j])/ R2_denom 
    }else if(R2_denom ==0){
		R2[i,j] <- 999
	}
	
  }
 }
 
  return(list(D,D1,R2)) 
}
 
######### 

	
```
