## Supplementary material for "Factors Affecting Response to Recurrent Genomic Selection in Soybeans": File S1: getPGPlots_SoyNAMPredMethods.R

```
###########

 
getInbreedingPlot <- function(F_List,nCon,nreps){ 
   
	cycleNo <- c(3:40)
	nCycles <- 40
  nSelCriteria <- length(F_List)
  	
	for(nSC in 1:nSelCriteria){

        F <-  F_List[[nSC]]
     
		F_Cyc <- list()
		F_Rate <- c()

		for(nCyc in 1:nCycles){ 
  		 
			F_Cyc[[nCyc]] <- mean(unlist(lapply(F[[nCyc]],function(x) length(x)/sum(1/x))))
	  
			if(nCyc==1){
				F_Rate[nCyc] <- 0
			}
	  
			if(nCyc >1){
	  
				F_CycVec <- unlist(F_Cyc)
				F_Rate[nCyc] <-  (F_CycVec[nCyc] - F_CycVec[nCyc-1])/(1-F_CycVec[nCyc-1])
			}
	  
		} 
  
	   
		leg_col <- c("dark green","blue","black","red","purple","gold3","dark orange")
		pchar <- c(1,3,5,7,9,11,13)
		nQTL <- 2
		
		initConVector <- c(1,3,5)
		initCon <- initConVector[nQTL]
		nCon <- initCon
		plotTitle <- "Average Rate of Inbreeding"
		par(oma=c(0,0,0,4),mar=c(4,4,4,12))
	    
		if(nSC==1){
				
			plot(cycleNo,(F_Rate[3:40]),type="o",xlim=c(1,40),ylim=c(-2.5,1.00),xlab="Cycle Number", ylab="Average Rate of Inbreeding ",main=plotTitle,lwd=3,lty=pchar[nSC],pch=pchar[nSC],col=leg_col[nSC])
			
		}else if(nSC >1){
			
			lines(cycleNo,(F_Rate[3:40]),type="o",pch=pchar[nSC],lwd=3,lty=pchar[nSC],col=leg_col[nSC])
			
		}
				
	}
		
		
        index <- 1
		label <- c("PS","RRREML","BayesB","BL","SVMRBF")
       # label <- c("PS","RRREML_1X_14Cyc","BayesB_1X_14Cyc","BL_1X_14Cyc","SVMRBF_1X_14Cyc")
	    sel <- c("QTL_40_H0.7","QTL_40_H0.3","QTL_400_H0.7","QTL_400_H0.3","QTL_4289_H0.7","QTL_4289_H0.3")
        Sellabel <- c("1.0%","2.5%","10.0%")
        nSelSch <- 3
  		
	    par(font=2)
	    ymax <- 1.0
        yoffset1 <- 0.90 
        yoffset2 <- 1.35
        par(font=2)
	   legend(41.5,ymax,legend=label[1:nSelCriteria],xpd=TRUE,lty=c(1,1),lwd=c(3,3),col=leg_col[1:nSelCriteria],cex=0.7,title="Method")
       legend(41.5,ymax-yoffset1,legend=Sellabel[nSelSch],xpd=TRUE,col="black",pch=c(1,3,5),title="Selected Top Fraction",text.width=strwidth("Selected Top Fraction")+0.05,cex=0.7) 
	   legend(41.5,ymax-yoffset2,legend=sel[nCon],xpd=TRUE,col="black",pch=c(1,3,5),title="Genetic Architecture",text.width=strwidth("Genetic Architecture")+0.05,cex=0.7) 
	
	}
	
# dev.off() 

getQTLIndices <- function(nLoci){

   nMarkers <- 4289
   nCycles <- 40

  if(nLoci== 40){
      QTL_indices<- seq(50,(nMarkers/2),by=53)
    }


   if(nLoci == 400){
      QTL_indices_1<- seq(8,nMarkers,by=12)
      QTL_indices_2 <- seq(50,nMarkers,by=107)
      n_3 <- 400-(length(QTL_indices_1))-(length(QTL_indices_2))
      QTL_indices_3<- sample(c(1:nMarkers),n_3)

      QTL_indices <- c(QTL_indices_1,QTL_indices_2,QTL_indices_3)
    }

    if(nLoci == nMarkers){
      QTL_indices <- c(1:nMarkers)
    }

    return(QTL_indices) 

 }
 


getExpHetPlot <- function(ExpHet_List,nCon,nReps){
 
  #plotFilename <- paste("ExpHet_GSMethods_14Cyc.png",sep="")
  #png(plotFilename,width=1024,height=768,pointsize=20)
  par(oma=c(0,0,0,4),mar=c(4,4,4,12))
  nCycles <- 40
  
  nSelCriteria <- length(ExpHet_List)
  cycleNo <- c(1:40)
  ExpHet_Cyc <- list()
  
  for(nSC in 1:nSelCriteria){	

  
    ExpHet <- ExpHet_List[[nSC]] 
 	ExpHet_Cyc <- list()
	ExpHet_Rate <- c()

    for(nCyc in 1:nCycles){ 
  
		 
      ExpHet_Cyc[[nCyc]] <- mean(unlist(lapply(ExpHet[[nCyc]],function(x) mean(x))))
	 	  	  
    } 
  
	
        
	leg_col <- c("dark green","blue","black","red","purple","gold3","dark orange")
	pchar <- c(1,3,5,7,9,11,13)
	nQTL <- 2
		
		#nConVector <- c(2,4,6)
		#nConditions <-nConVector[nQTL]
		
	initConVector <- c(1,3,5)
	initCon <- initConVector[nQTL]
	nCon <- initCon
	plotTitle <- "Average Expected Heterozygosity"
			
	if(nSC==1){
				
			plot(cycleNo,unlist(ExpHet_Cyc),type="o",ylim=c(0,0.12),xlab="Cycle Number", ylab="Average Expected Heterozygosity",main=plotTitle,lwd=3,lty=pchar[nSC],pch=pchar[nSC],col=leg_col[nSC])
			
	}else if(nSC >1){
			
			lines(cycleNo,unlist(ExpHet_Cyc),type="o",pch=pchar[nSC],lwd=3,lty=pchar[nSC],col=leg_col[nSC])
			
	}
	
				
	}
		
		
    index <- 1
	label <- c("PS","RRREML","BayesB","BL","SVMRBF")
    #label <- c("PS","RRREML_1X_14Cyc","BayesB_1X_14Cyc","BL_1X_14Cyc","SVMRBF_1X_14Cyc")
    sel <- c("QTL_40_H0.7","QTL_40_H0.3","QTL_400_H0.7","QTL_400_H0.3","QTL_4289_H0.7","QTL_4289_H0.3")
    Sellabel <- c("1.0%","2.5%","10.0%")
    nSelSch <- 3
    ymax <- 0.12
    yoffset1 <- 0.042
    yoffset2 <- 0.062
  		
	par(font=2)
	legend(41.5,ymax,legend=label[1:nSelCriteria],xpd=TRUE,lty=c(1,1),lwd=c(3,3),col=leg_col[1:nSelCriteria],cex=0.7,title="Method")
    legend(41.5,ymax-yoffset1,legend=Sellabel[nSelSch],xpd=TRUE,col="black",pch=c(1,3,5),title="Selected Top Fraction",text.width=strwidth("Selected Top Fraction")+0.10,cex=0.7) 
	legend(41.5,ymax-yoffset2,legend=sel[nCon],xpd=TRUE,col="black",pch=c(1,3,5),title="Genetic Architecture",text.width=strwidth("Genetic Architecture")+0.10,cex=0.7)
    #dev.off()	
			
}


getFavAlleleOutput <- function(MAF_List,FavAlleleList,QTL_Indices_400){

 nSelCriteria <- length(MAF_List)
 nRep <- 2
 nCycles <- 40 
 FavAllele_List <- list()
 FavAll_Freq_List_Cyc <- rep(list(rep(list(list()),nRep)),nCycles)
 FavAll_Freq_List_Met <- rep(list(rep(list(rep(list(list()),nRep)),nCycles)),nSelCriteria)

 for(nSC in 1:nSelCriteria){

  for(nCyc in 1:nCycles){

   for(nrep in 1:nRep){ 

    FavAllele_List[[nrep]] <- FavAlleleList[[nSC]][[1]][[nrep]][[2]] 
 
    FavAllele <-  FavAllele_List[[nrep]]

    OneIndices <-  which(FavAllele ==1)
    ZeroIndices <- which(FavAllele ==0)
    FavAll_Freq_List <- rep(0,4289)

    FavAll_Freq_List[OneIndices]  <- 1-MAF_List[[nSC]][[nCyc]][[nrep]][OneIndices] 
    FavAll_Freq_List[ZeroIndices] <- MAF_List[[nSC]][[nCyc]][[nrep]][ZeroIndices] 

    FavAll_Freq_List_Cyc[[nCyc]][[nrep]]<- FavAll_Freq_List


   }
  }
    FavAll_Freq_List_Met[[nSC]] <- FavAll_Freq_List_Cyc

}
 
 QTL_indices <- QTL_Indices_400
 nRep <- 2
 nCycles <- 40
 
 FavAll_Table <- matrix(rep(0,nRep*4289),nrow=nRep,ncol=4289)
 FavAll_List <- rep(list(rep(list(list()),nRep)),nCycles)

 Lost_FavAll_Table <- rep(0,nRep)
 Lost_FavQTLAll_Table <- rep(0,nRep)

 Avg_FavAll_List <- rep(list(rep(list(list()),nCycles)),nSelCriteria)
 Tot_FavAll_List <- rep(list(rep(list(list()),nCycles)),nSelCriteria)
 Tot_FavQTL_List <- rep(list(rep(list(list()),nCycles)),nSelCriteria)

 Tot_LostFavAll_List <- rep(list(rep(list(list()),nCycles)),nSelCriteria)
 Tot_LostFavQTLAll_List <- rep(list(rep(list(list()),nCycles)),nSelCriteria)


 for(nSC in 1:nSelCriteria){
  #nRep <- 1
  for(nCyc in 1:40){
   for(nrep in 1:nRep){

		FavAll_Table[nrep,] <- FavAll_Freq_List_Met[[nSC]][[nCyc]][[nRep]]

		Lost_FavAll_Table[nrep] <- length(which(FavAll_Freq_List_Met[[nSC]][[nCyc]][[nRep]] %in% 0))
		Lost_FavQTLAll_Table[nrep] <- length(which(FavAll_Freq_List_Met[[nSC]][[nCyc]][[nRep]][QTL_indices] %in% 0))
   
    }
		Avg_FavAll_List[[nSC]][[nCyc]] <- apply(FavAll_Table,2,mean)
  
		Tot_FavAll_List[[nSC]][[nCyc]] <- sum(Avg_FavAll_List[[nSC]][[nCyc]]*4000)

		Tot_FavQTL_List[[nSC]][[nCyc]] <- sum(Avg_FavAll_List[[nSC]][[nCyc]][QTL_indices]*4000)

		Tot_LostFavAll_List[[nSC]][[nCyc]] <- mean(Lost_FavAll_Table)
		Tot_LostFavQTLAll_List[[nSC]][[nCyc]] <- mean(Lost_FavQTLAll_Table)
  }
 }
 
	return(list(Tot_LostFavAll_List,Tot_LostFavQTLAll_List))
}


 
getLostGenoValue <- function(MAF_List_nTS,FavAlleleList,QTL_Indices_400){

 MAF_List <- MAF_List_nTS
 nSelCriteria <- length(MAF_List)
 nRep <- 10
 nCycles <- 40 
 FavAllele_List <- list()
 FavAll_Freq_List_Cyc <- rep(list(rep(list(list()),nRep)),nCycles)
 FavAll_Freq_List_Met <- rep(list(rep(list(rep(list(list()),nRep)),nCycles)),nSelCriteria)
 QTL_Freq_List_Cyc <- rep(list(rep(list(list()),nRep)),nCycles)
 QTL_Freq_List_Met <- rep(list(rep(list(rep(list(list()),nRep)),nCycles)),nSelCriteria) 
 
 

 for(nSC in 1:nSelCriteria){

  for(nCyc in 1:nCycles){

   for(nrep in 1:nRep){ 

    FavAllele_List[[nrep]] <- FavAlleleList[[nrep]][[2]] 
 
    FavAllele <-  FavAllele_List[[nrep]]
    
    FavAll_Freq_List <- rep(0,4289)
    QTL_Freq_List <- rep(0,400)
    
	evenIndices<- seq(2,400,by=2)
	oddIndices <- seq(1,399,by=2)

   
	QTL_Freq_List[evenIndices] <- 1-MAF_List[[nSC]][[nCyc]][[nrep]][QTL_Indices_400[evenIndices]]
	QTL_Freq_List[oddIndices] <- MAF_List[[nSC]][[nCyc]][[nrep]][QTL_Indices_400[oddIndices]]
	
    FavAll_Freq_List_Cyc[[nCyc]][[nrep]]<- FavAll_Freq_List
    QTL_Freq_List_Cyc[[nCyc]][[nrep]] <- QTL_Freq_List

   }
  }
    FavAll_Freq_List_Met[[nSC]] <- FavAll_Freq_List_Cyc
    QTL_Freq_List_Met[[nSC]] <- QTL_Freq_List_Cyc
}

############## 
 
 QTL_indices <- QTL_Indices_400
 nRep <- 10
 nCycles <- 40
 
 FavAll_Table <- matrix(rep(0,nRep*4289),nrow=nRep,ncol=4289)
 FavAll_List <- rep(list(rep(list(list()),nRep)),nCycles)

 Lost_FavAll_Table <- rep(0,nRep)
 Lost_FavQTLAll_Table <- rep(0,nRep)

 Avg_FavAll_List <- rep(list(rep(list(list()),nCycles)),nSelCriteria)
 Tot_FavAll_List <- rep(list(rep(list(list()),nCycles)),nSelCriteria)
 Tot_FavQTL_List <- rep(list(rep(list(list()),nCycles)),nSelCriteria)

 Tot_LostFavAll_List <- rep(list(rep(list(list()),nCycles)),nSelCriteria)
 Tot_LostFavQTLAll_List <- rep(list(rep(list(list()),nCycles)),nSelCriteria)

Lost_FavQTLAll_Indices <- rep(list(list()),nRep)
Lost_FavQTLAll_Indices_List <- rep(list(rep(list(list()),nCycles)),nSelCriteria)

 Lost_Genetic_Value <- rep(list(list()),nRep)
 Lost_Genetic_Value_List <-rep(list(rep(list(list()),nCycles)),nSelCriteria)
 Lost_FavQTLAll_Freq <- rep(0,nRep)
 Lost_FavQTLAll_Freq_List <- rep(list(rep(list(list()),nCycles)),nSelCriteria)
 
 for(nSC in 1:nSelCriteria){
  for(nCyc in 1:40){
    for(nrep in 1:nRep){

				
		Lost_FavQTLAll_Freq[nrep] <- length(which(QTL_Freq_List_Met[[nSC]][[nCyc]][[nRep]] %in% 0))
		
		Lost_Genetic_Value[[nrep]] <- Lost_FavQTLAll_Freq * 0.5
		
   
    }
	    Lost_FavQTLAll_Freq_List[[nSC]][[nCyc]] <- mean(Lost_FavQTLAll_Freq)
					
		Lost_Genetic_Value_List[[nSC]][[nCyc]] <- mean(unlist(Lost_Genetic_Value))

  }
 }
 
	return(list(Lost_FavQTLAll_Freq_List,Lost_Genetic_Value_List))
	
}


getLostFavorableAllelePlot <- function(Tot_LostFavQTLAll_List,ncon,nrep,TrainCycle,ncycles){

 cycleNo <- c(1:ncycles)
 ymin <- min(unlist(Tot_LostFavQTLAll_List))-5
 ymax <- max(unlist(Tot_LostFavQTLAll_List))+25
 nSelCriteria <- length(Tot_LostFavQTLAll_List)
 leg_col <- c("dark green","blue","black","red","purple")
 nCycles <- ncycles
 trainCycle<- TrainCycle
 
 nCon <- ncon
 plotTitle <- "Average Number of Lost Favorable QTL in Recurrent GS"


for(nSC in 1:nSelCriteria){
   
 Tot_LostFavQTLAll <- unlist(Tot_LostFavQTLAll_List[[nSC]])

 if(nSC==1){
 	plotFilename <- paste("GSMethods_Lost_QTL_Sel_400_QTL_",TrainCycle,"_",nCycles,"_.png",sep="")
 	
    png(plotFilename,width=1024,height=768,pointsize=20)
	par(oma=c(0,0,0,2),mar=c(4,4,4,12))
 	plot(cycleNo,Tot_LostFavQTLAll[1:nCycles],type="o",xlim=c(0,nCycles),ylim=c(ymin,ymax),lty=3,lwd=3,col=leg_col[nSC],xlab="",ylab="",main="") 
 }
 else if(nSC >1){ 
 
	lines(cycleNo,Tot_LostFavQTLAll[1:nCycles],type="o",lty=3,lwd=3,col=leg_col[nSC])
 }
}

 title(xlab="Cycle",ylab="Average Lost Favorable QTL Alleles",main=plotTitle,font=2)
 
 
 if(trainCycle ==1){
		SelMethod <- c("PS","RRREML","BayesB","BL","SVMRBF")
		
		}
 if(trainCycle ==4){
		SelMethod <- c("PS","RRREML_1X_14Cyc","BayesB_1X_14Cyc","BL_1X_14Cyc","SVMRBF_1X_14Cyc")
 }	

 par(font=2)
 sel <- c("QTL_40_H0.7","QTL_40_H0.3","QTL_400_H0.7","QTL_400_H0.3","QTL_4289_H0.7","QTL_4289_H0.3")
 Sellabel <- c("1.0%","2.5%","10.0%")

 yoffset1 <- 70 
 yoffset2 <- 100
 nSelSch <- 3
 x_c <- nCycles +1.5
 legend(x_c,ymax,legend=SelMethod[1:nSelCriteria],xpd=TRUE,lty=c(1,1),lwd=c(3,3),col=leg_col[1:nSelCriteria],cex=0.7,title="Method") 
 legend(x_c,ymax-yoffset1,legend=Sellabel[nSelSch],xpd=TRUE,col="black",pch=c(1,3,5),title="Selected Top Fraction",text.width=strwidth("Selected Top Fraction"),cex=0.70) 
 legend(x_c,ymax-yoffset2,legend=sel[nCon],xpd=TRUE,col="black",pch=c(1,3,5),title="Genetic Model",text.width=strwidth(
 "Genetic Model")+0.01,cex=0.7) 
 
 dev.off()
 
 }


#### 
# Tot_LostFavQTLAll_List <- FavAllele_Out[[2]]
# 
# getLostFavorableAllelePlot(Tot_LostFavQTLAll_List,ncon,nrep)

getLostFavorableAllelePlot <- function(Tot_LostFavQTLAll_List,ncon,nrep){

 cycleNo <- c(1:40)
 ymin <- min(unlist(Tot_LostFavQTLAll_List))-5
 ymax <- max(unlist(Tot_LostFavQTLAll_List))
 nSelCriteria <- length(Tot_LostFavQTLAll_List)
 leg_col <- c("dark green","blue","black","red","purple")
 nCycles <- 40 
 
 nCon <- ncon
 plotTitle <- "Average Number of Lost Favorable QTL in Recurrent GS"

for(nSC in 1:nSelCriteria){
   
 Tot_LostFavQTLAll <- unlist(Tot_LostFavQTLAll_List[[nSC]])

 if(nSC==1){
 	plotFilename <- paste("GSMethods_Lost_QTL_Sel_400_QTL.png")
 	#png(plotFilename,width=1024,height=768,pointsize=20)
	par(oma=c(0,0,0,4),mar=c(4,4,4,8))

 	plot(cycleNo,Tot_LostFavQTLAll,type="o",xlim=c(0,40),ylim=c(ymin,ymax),lty=3,lwd=3,col=leg_col[nSC],xlab="Cycle",ylab="Average Lost Favorable QTL Alleles",main=plotTitle) 
 }
 else if(nSC >1){ 
 
	lines(cycleNo,Tot_LostFavQTLAll,type="o",lty=3,lwd=3,col=leg_col[nSC])
 }
}

 #label <- c("PS","RRREML","BayesB","BL","SVMRBF")
 par(font=2)
 sel <- c("QTL_40_H0.7","QTL_40_H0.3","QTL_400_H0.7","QTL_400_H0.3","QTL_4289_H0.7","QTL_4289_H0.3")
 label <- c("PS","RRREML","BayesB","BL","SVMRBF")
 Sellabel <- c("1.0%","2.5%","10.0%")

 yoffset1 <- 50 
 yoffset2 <- 100
 nSelSch <- 3
 
 legend(41.5,ymax,legend=label[1:nSelCriteria],xpd=TRUE,lty=c(1,1),lwd=c(3,3),col=leg_col[1:nSelCriteria],cex=0.7,title="Method") 
 legend(41.5,ymax-yoffset1,legend=Sellabel[nSelSch],xpd=TRUE,col="black",pch=c(1,3,5),title="Selected Top Fraction",text.width=strwidth("Selected Top Fraction")+0.10,cex=0.7) 
 legend(41.5,ymax-yoffset2,legend=sel[nCon],xpd=TRUE,col="black",pch=c(1,3,5),title="Genetic Architecture",text.width=strwidth(
 "Genetic Architecture")+0.10,cex=0.7) 
 
 }
 
 
 
 


getLDPlots <- function(Marker_Marker_cM_List_Cyc,Marker_Marker_LD_List_Cyc){
  
  Cycles <- c(1,5,10,15,20,25,30,35)[1:5]
  
  for(nCyc in 1:(length(Cycles))){
    
    # nCyc <- 1
    
    LDPlotTableMarker <- c() 
    
    cM_Marker_Code_List <- list()
    cM_Marker_cMCode_List <- list()
    nMarkers <- length(Marker_Marker_cM_List_Cyc[[nCyc]])
    for(nMarker in 1:nMarkers){
      
      a <- unlist(Marker_Marker_cM_List_Cyc[[nCyc]][[nMarker]])
      aCM <- a
      
      a[which(a>=0 & a<=20)] <- 1 
      a[which(a>20 & a<=40)] <- 2 
      a[which(a>40 & a<=60)] <- 3 
      a[which(a>60 & a<=80)] <- 4 
      a[which(a>80 & a<=100)] <- 5 
      a[which(a>100 & a<=130)] <- 6  
      
      aCM[which(a>=0 & a<=20.0)] <- "0-20" 
      aCM[which(a>20.0 & a<=40.0)] <- "20-40" 
      aCM[which(a>40.0 & a<=60.0)] <- "40-60"
      aCM[which(a>60.0 & a<=80.0)] <- "60-80"
      aCM[which(a>80.0 & a<=100.0)] <- "80-100"
      aCM[which(a>100.0 & a<=130.0)] <- "100-130"
      
      #,cM_Marker_cMCode_List[[nMarker]]
      
      cM_Marker_Code_List[[nMarker]] <- a
      cM_Marker_cMCode_List[[nMarker]] <- aCM
      LDPlotTable <- cbind(Marker_Marker_LD_List_Cyc[[nCyc]][[nMarker]],Marker_Marker_cM_List_Cyc[[nCyc]][[nMarker]],cM_Marker_Code_List[[nMarker]])
      LDPlotTableMarker <- rbind(LDPlotTableMarker,LDPlotTable)
    }
    
    #(D_Table[,1])
    
    D_Table <- LDPlotTableMarker
 
    par(oma=c(0,0,0,6),mar=c(4,8,4,12))
  
    boxplot(D_Table[,1] ~ (D_Table[,3]),data=D_Table,ylab="D",xlab="Intervals cM",main=paste("LD Decay over Distance cM- Selection Cycle -",Cycles[nCyc],sep=""),outline=TRUE)
    
    label <- c("1 -> 0-20", "2 -> 20-40", "3 -> 40-60", "4 -> 60-80", "5 -> 80-100", "6 -> 100-130")
    legend(7.0,0.7,label,xpd=TRUE,title="Intervals in cM")
  }
}
################    
    


	 
# ################ 	 Requires another check   
#getSelGenoVariancePlot <- function(GenoVal_NX_N_List_3c_PhenoPred_GP_List2,ncon,nReps){

    
 	

    # nCycles <-40
    # nreps <- 10
    # cycleNo <- c(1:nCycles)

# for(nSelCon in 1:2){ 

 # for(nTrnCyc in 1:4){
	
    # for(nQTL in 1:3){
	
	# GenoVal_NX_N_List_3c_Pheno_GP_Update_List <- GenoVal_NX_N_List_3c_Pheno200_GP_Update_List2_TrainCyc[[nTrnCyc]]
	
      	# nSelInt <- length(GenoVal_NX_N_List_3c_Pheno_GP_Update_List)
        # nSelInt_Series <- 1:nSelInt
 
        
	# leg_col <- c("dark green","blue","black","red","purple","gold3","coral3","lightgoldenrod3","sienna4","grey30","steelblue3","indianred2","wheat4","tomato","dark orange")
		
	# pchar <- c(1,3,5,7,9,11,13)
		
		
		# nConVector <- c(2,4,6)
		# initConVector <- c(1,3,5)
		# finalCon <-nConVector[nQTL]
		# initCon <- initConVector[nQTL]
		# selCon <- c(initCon,finalCon)[nSelCon]
                # H2_Levels <- c("H2-0.7","H2-0.3")
	   	
			# if(nQTL==1){
				# ymin<- 0
				# ymax <- 80
				# yoffset1 <- 50
				# yoffset2 <- 60
		    # }else if(nQTL==2){
				# ymin <- 0
				# ymax <- 8.5
				# yoffset1 <- 3.5
				# yoffset2 <- 4.5
		    # }else if(nQTL==3){ 
				# ymin <- 0
				# ymax<- 0.85
				# yoffset1 <- 0.35
				# yoffset2 <- 0.45
		    # }
		
	
		
	# Rs_List <- list()
	
	
	# for(nSelSeries in (nSelInt_Series)){
	  
	   	  
	   # for(nSel in 1:nSelSeries){
	
	   	
	      # for(nCon in selCon){
		    
		  
        		
		    # GenoVal_NX_N_List_3c_Pheno_GP_Update <- GenoVal_NX_N_List_3c_Pheno_GP_Update_List[[nSel]]
		
					
			# Rt <- rep(0,40)
			# Rs <- rep(0,40)
			# pValue.t <- rep(0,40)

			# Gt_Rep <- matrix(rep(0,nreps*nCycles),nrow=nreps,ncol=nCycles)
			# GVart_Rep <- matrix(rep(0,nreps*nCycles),nrow=nreps,ncol=nCycles)
			
			
			# ## mean genotypic value of a population of 2000 individuals every generation for 10 replicates 
			# ## genotypic variance - variance of genetic values of a ppoulation every generation for 10 replicates 
			# ## Average of mean genotypic value of 10 reps every generation 
            # ## Average of genotypic variance of 10 reps every generation 			
			
			
			# for(nReps in 1:nreps){ 
			 
			  # for (t in 1:nCycles){
				
				# Gt_Rep[nReps,t] <- mean(GenoVal_NX_N_List_3c_Pheno_GP_Update[[nCon]][[nReps]][,t])
				# GVart_Rep[nReps,t] <- var(GenoVal_NX_N_List_3c_Pheno_GP_Update[[nCon]][[nReps]][,t]) 
			
			  # }
			# }
										
			# R0 <- mean(Gt_Rep[,1])
			# sd0 <- mean(GVart_Rep[,1])
			# Std0 <- abs(R0)/sd0
			# Rm <-  200

			# Gt <- apply(Gt_Rep,2,mean)
			
		    # GVart <-apply(GVart_Rep,2,mean)
					
	   
			
			# if(nCon==selCon && nSel ==1){ 

				  
	  			 # plotFilename <- paste("Avg_Sel_GenVar_GS_TrainCycSet_Sel200","_QTL_",nQTL,"TrainCycSet_",nTrnCyc,"_h2_",H2_Levels[nSelCon],nSelSeries,".png")
	   			 # png(plotFilename,width=1024,height=768,pointsize=20)

	  			 # par(oma=c(0,0,0,6),mar=c(4,8,4,12))
				 # plotTitle <- paste("Genotypic Variance in Selected Population",nSelSch,sep="")

				 # plot(cycleNo,(GVart),type="o",ylim=c(ymin,ymax),xlab="Cycle Number", ylab="Average Genenotypic Variance",main=plotTitle,lwd=3,lty=pchar[nCon],pch=pchar[nCon],col=leg_col[nSel])
			# }else if(((nSel==1)&&(nCon>selCon)) || ((nSel!=1)&&(nCon>=selCon))){
			
			    # lines(cycleNo,(GVart),type="o",pch=pchar[nCon],lwd=3,lty=pchar[nCon],col=leg_col[nSel])
			
			# }
			
			
		# }
	
	# }
		
    
	
	# label <- list(c("PS","RRREML","BayesB","BL","SVMRBF"),c("PS","RRREML_1X_10Cyc","BayesB_1X_10Cyc","BL_1X_10Cyc","SVMRBF_1X_10Cyc"),c("PS","RRREML_1X_12Cyc","BayesB_1X_12Cyc","BL_1X_12Cyc","SVMRBF_1X_12Cyc"),c("PS","RRREML_1X_14Cyc","BayesB_1X_14Cyc","BL_1X_14Cyc","SVMRBF_1X_14Cyc"))[[nTrnCyc]]
        # sel<- c("QTL_40_H0.7","QTL_40_H0.3","QTL_400_H0.7","QTL_400_H0.3","QTL_4289_H0.7","QTL_4289_H0.3")
  			
	# par(font=2)
	# legend(41.5,ymax,legend=label,xpd=TRUE,lty=c(1,1),lwd=c(3,3),col=leg_col,cex=0.7,title="GS Model") 
        # legend(41.5,ymax-yoffset1,legend=Sellabel[nSelSch],xpd=TRUE,col="black",pch=c(1,3,5),title="Selected Top Fraction",text.width=strwidth("Selected Top Fraction")+0.05,cex=0.85) 
	# legend(41.5,ymax-yoffset2,legend=sel[nCon],xpd=TRUE,col="black",pch=c(1,3,5),title="Genetic Model",text.width=strwidth("Genetic Model")+0.10,cex=0.85) 
	 
   	 # dev.off() 
  # } 
 # }
# }
# }
#}
# #########################
 
```
