## Supplementary material for "Factors Affecting Response to Recurrent Genomic Selection in Soybeans": File S1: getPGStats_GSMethods.R

```
getPGStatsGSMethods_Cyc1 <- function(CycleGenoTable_AF,nextGenGenoTable_NumF,Populations,MarkerEffects,no_QTL,gIndObjects,ModelType,nCon,nrep,nCyc){ 

        GenTable_AF <- CycleGenoTable_AF
		nextGenGenoTable <- nextGenGenoTable_NumF
		populations <- Populations
		noQTL <- no_QTL
		
		QTL_GenoTable <- getQTLTable(nextGenGenoTable,no_QTL)
					
### get PG stats from genotable
		
		Freq_Allele <-	getFreq_BasePoln(nextGenGenoTable,MarkerEffects)
		FavorableAlleleFreq <- Freq_Allele[[1]]
		FavorableAllele <- Freq_Allele[[2]]
		percentHeterozygous_List <- getPercentHeterozygous(nextGenGenoTable)

		Freq_QTLAllele <- getFreq_BasePoln(QTL_GenoTable,MarkerEffects)
		FavorableQTLAlleleFreq <- Freq_QTLAllele[[1]]
		FavorableQTLAllele <- Freq_QTLAllele[[2]]
		percentHeterozygous_QTL <- getPercentHeterozygous(QTL_GenoTable)

  
		rm(nextGenGenoTable)
	
### get PG parameters from GenInd

    if(gIndObjects==TRUE){

        CycleGenoTable_GInd <- df2genind(GenTable_AF,pop=populations,sep="")
 
		Diff_Stats <- diff_stats(CycleGenoTable_GInd)
	    		
		F <- inbreeding(CycleGenoTable_GInd,res.type="estimate")
        ExpHet <- Hs(CycleGenoTable_GInd)
		Freq_GInd <- makefreq(CycleGenoTable_GInd)
		MAF <- minorAllele(CycleGenoTable_GInd)
	}
	
	
	
	if(gIndObjects==TRUE){
		return(list(FavorableAlleleFreq,FavorableAllele,percentHeterozygous_List,FavorableQTLAlleleFreq,FavorableQTLAllele,percentHeterozygous_QTL,Diff_Stats,F,ExpHet,Freq_GInd,MAF))
	}else if(gIndObjects==FALSE){ 
		return(list(FavorableAlleleFreq,FavorableAllele,percentHeterozygous_List,FavorableQTLAlleleFreq,FavorableQTLAllele,percentHeterozygous_QTL))
	}

}


 
getPGStatsGSMethods <- function(CycleGenoTable_AF,nextGenGenoTable_NumF,Populations,MarkerEffects,FavAllele,no_QTL,gIndObjects,ModelType,nCon,nrep,nCyc){ 

        GenTable_AF <- CycleGenoTable_AF
		populations <- Populations
		noQTL <- no_QTL
		nextGenGenoTable <- nextGenGenoTable_NumF
### get PG stats from genotable
		
		QTL_GenoTable <- getQTLTable(nextGenGenoTable,noQTL)
		
		Freq_Allele <-	getFreq(nextGenGenoTable,FavAllele)
		FavorableAlleleFreq <- Freq_Allele[[1]]
		FavorableAllele  <- Freq_Allele[[2]]
		percentHeterozygous_List <- getPercentHeterozygous(nextGenGenoTable)

		Freq_QTLAllele <- getFreq(QTL_GenoTable,FavAllele)
		FavorableQTLAlleleFreq <- Freq_QTLAllele[[1]]
		FavorableQTLAllele  <- Freq_QTLAllele[[2]]
		percentHeterozygous_QTL <- getPercentHeterozygous(QTL_GenoTable)
		
		rm(nextGenGenoTable)

    		
    if(gIndObjects==TRUE){		
		
### get PG parameters from GenInd

        CycleGenoTable_GInd <- df2genind(GenTable_AF,pop=populations,sep="")
 		Diff_Stats <- diff_stats(CycleGenoTable_GInd)
	    
				
		F <- inbreeding(CycleGenoTable_GInd,res.type="estimate")
        ExpHet <- Hs(CycleGenoTable_GInd)
		Freq_GInd <- makefreq(CycleGenoTable_GInd)
		MAF <- minorAllele(CycleGenoTable_GInd)
		
		return(list(FavorableAlleleFreq,FavorableAllele,percentHeterozygous_List,FavorableQTLAlleleFreq,FavorableQTLAllele,percentHeterozygous_QTL,Diff_Stats,F,ExpHet,Freq_GInd,MAF))
	} 
	else if(gIndObjects==FALSE){ 
		return(list(FavorableAlleleFreq,FavorableAllele,percentHeterozygous_List,FavorableQTLAlleleFreq,FavorableQTLAllele,percentHeterozygous_QTL))
	}

}


###########################################
 
 getNumericFormat <- function(GenoTable_AlleleFormat,alleleConversionTable_Combined){

        options(warn=-1)
        genoTable <- apply(GenoTable_AlleleFormat,2,as.character)
        nIndividuals <- nrow(genoTable)
		nMarkers <- ncol(genoTable)
        #AlleleConversionTable_Combined <- alleleConversionTable_Combined[,-1]
		AlleleConversionTable_Combined <- apply(alleleConversionTable_Combined,2,as.character)
        IA3023_alleles <- as.character(AlleleConversionTable_Combined[,8])
        Alt_Homozygous_alleles <- as.character(AlleleConversionTable_Combined[,10])
        # Het_alleles1 <- as.character(AlleleConversionTable_Combined[,22])
		# Het_alleles2 <- as.character(AlleleConversionTable_Combined[,23])
		Het_alleles1 <-paste(as.character(AlleleConversionTable_Combined[,7]),as.character(AlleleConversionTable_Combined[,9]),sep="")
		Het_alleles2 <- paste(as.character(AlleleConversionTable_Combined[,9]),as.character(AlleleConversionTable_Combined[,7]),sep="")
		
		Recombinant_Hom1_alleles <- c()
		Recombinant_Hom2_alleles <- c()
		Recombinant_Het1_alleles <- c()
		Recombinant_Het2_alleles <- c()
		
		NR1_Recombinant_Het1_alleles <- c()
		NR1_Recombinant_Het2_alleles <- c() 
		NR1_Recombinant_Het3_alleles <- c() 
		NR1_Recombinant_Het4_alleles <- c()
		
		NR2_Recombinant_Het1_alleles <- c()
		NR2_Recombinant_Het2_alleles <- c() 
		NR2_Recombinant_Het3_alleles <- c() 
		NR2_Recombinant_Het4_alleles <- c()
		
		for(i in 1:nMarkers){
		   completeSet <- c("A","T","G","C") 
		   NR1 <- as.character(AlleleConversionTable_Combined[i,7])
		   NR2 <- as.character(AlleleConversionTable_Combined[i,9])
		   d2 <- setdiff(completeSet,c(NR1,NR2))
		   #d2 <- setdiff(d1,as.character(AlleleConversionTable_Combined[i,9])) 
		
		   Recombinant_Hom1_alleles <- c(Recombinant_Hom1_alleles,paste(d2[1],d2[1],sep=""))
		   Recombinant_Hom2_alleles <- c(Recombinant_Hom2_alleles,paste(d2[2],d2[2],sep=""))
		   
		   Recombinant_Het1_alleles <- c(Recombinant_Het1_alleles,paste(d2[1],d2[2],sep=""))
		   Recombinant_Het2_alleles <- c(Recombinant_Het2_alleles,paste(d2[2],d2[1],sep=""))
		  
		   NR1_Recombinant_Het1_alleles <- c(NR1_Recombinant_Het1_alleles,paste(d2[1],NR1,sep=""))
		   NR1_Recombinant_Het2_alleles <- c(NR1_Recombinant_Het2_alleles,paste(NR1,d2[1],sep=""))
		 
		   NR1_Recombinant_Het3_alleles <- c(NR1_Recombinant_Het3_alleles,paste(d2[2],NR1,sep=""))
		   NR1_Recombinant_Het4_alleles <- c(NR1_Recombinant_Het4_alleles,paste(NR1,d2[2],sep=""))
				 
		   NR2_Recombinant_Het1_alleles <- c(NR2_Recombinant_Het1_alleles,paste(d2[1],NR2,sep=""))
		   NR2_Recombinant_Het2_alleles <- c(NR2_Recombinant_Het2_alleles,paste(NR2,d2[1],sep=""))
		   
		   NR2_Recombinant_Het3_alleles <- c(NR2_Recombinant_Het3_alleles,paste(d2[2],NR2,sep=""))
		   NR2_Recombinant_Het4_alleles <- c(NR2_Recombinant_Het4_alleles,paste(NR2,d2[2],sep=""))
		
		}
		
		
#####   NR

        IA3023_alleles_Indices <- apply(genoTable,1,function(x) which(as.character(x) == (IA3023_alleles)))

        Het_alleles1_Indices <-  apply(genoTable,1,function(x) if(length(which(as.character(x)==(Het_alleles1)))!=0){which(as.character(x)==(Het_alleles1))} else{0})
		
		Het_alleles2_Indices <-  apply(genoTable,1,function(x) if(length(which(as.character(x)==(Het_alleles2)))!=0){which(as.character(x)==(Het_alleles2))} else{0})
         
        Alt_Homozygous_alleles_Indices <- apply(genoTable,1,function(x) which(as.character(x) == (Alt_Homozygous_alleles)))
		
####### Recombinant 

        Recombinant_Hom1_alleles_Indices <- apply(genoTable,1,function(x) if(length(which(as.character(x)==(Recombinant_Hom1_alleles)))!=0){which(as.character(x)==(Recombinant_Hom1_alleles))} else{0})
       
	    Recombinant_Hom2_alleles_Indices <- apply(genoTable,1,function(x) if(length(which(as.character(x)==(Recombinant_Hom2_alleles)))!=0){which(as.character(x)==(Recombinant_Hom2_alleles))} else{0})
		
		Recombinant_Het1_alleles_Indices <- apply(genoTable,1,function(x) if(length(which(as.character(x)==(Recombinant_Het1_alleles)))!=0){which(as.character(x)==(Recombinant_Het1_alleles))} else{0})
		
		Recombinant_Het2_alleles_Indices <- apply(genoTable,1,function(x) if(length(which(as.character(x)==(Recombinant_Het2_alleles)))!=0){which(as.character(x)==(Recombinant_Het2_alleles))} else{0})

####### NR-Recombinant
	
       	NR1_Recombinant_Het1_alleles_Indices <- apply(genoTable,1,function(x) if(length(which(as.character(x)==(NR1_Recombinant_Het1_alleles)))!=0){which(as.character(x)==(NR1_Recombinant_Het1_alleles))} else{0})
       
	    NR1_Recombinant_Het2_alleles_Indices <- apply(genoTable,1,function(x) if(length(which(as.character(x)==(NR1_Recombinant_Het2_alleles)))!=0){which(as.character(x)==(NR1_Recombinant_Het2_alleles))} else{0})
	
	    NR1_Recombinant_Het3_alleles_Indices <- apply(genoTable,1,function(x) if(length(which(as.character(x)==(NR1_Recombinant_Het3_alleles)))!=0){which(as.character(x)==(NR1_Recombinant_Het3_alleles))} else{0})
	
		NR1_Recombinant_Het4_alleles_Indices <- apply(genoTable,1,function(x) if(length(which(as.character(x)==(NR1_Recombinant_Het4_alleles)))!=0){which(as.character(x)==(NR1_Recombinant_Het4_alleles))} else{0})
	
		
		NR2_Recombinant_Het1_alleles_Indices <- apply(genoTable,1,function(x) if(length(which(as.character(x)==(NR2_Recombinant_Het1_alleles)))!=0){which(as.character(x)==(NR2_Recombinant_Het1_alleles))} else{0})
       
	    NR2_Recombinant_Het2_alleles_Indices <- apply(genoTable,1,function(x) if(length(which(as.character(x)==(NR2_Recombinant_Het2_alleles)))!=0){which(as.character(x)==(NR2_Recombinant_Het2_alleles))} else{0})
	
	    NR2_Recombinant_Het3_alleles_Indices <- apply(genoTable,1,function(x) if(length(which(as.character(x)==(NR2_Recombinant_Het3_alleles)))!=0){which(as.character(x)==(NR2_Recombinant_Het3_alleles))} else{0})
	
		NR2_Recombinant_Het4_alleles_Indices <- apply(genoTable,1,function(x) if(length(which(as.character(x)==(NR2_Recombinant_Het4_alleles)))!=0){which(as.character(x)==(NR2_Recombinant_Het4_alleles))} else{0})
		
	
			
		Total_Alleles <- rep(0,nIndividuals)
					
		for(i in 1:nIndividuals){

           if(length(IA3023_alleles_Indices[[i]]) >1){
		          genoTable[i,(IA3023_alleles_Indices[[i]])] <- "1"
			}
           if(length(Het_alleles1_Indices[[i]])>=1 & Het_alleles1_Indices[[i]] !=0){
                genoTable[i,(Het_alleles1_Indices[[i]])] <- "0"
			}
		   if(length(Het_alleles2_Indices[[i]])>=1 & Het_alleles2_Indices[[i]] !=0){
                genoTable[i,(Het_alleles2_Indices[[i]])] <- "0"
			}
		    if(length(Alt_Homozygous_alleles_Indices[[i]]) >1){
                genoTable[i,(Alt_Homozygous_alleles_Indices[[i]])] <- "-1"
			}
		   
		  #  & Recombinant_Hom2_alleles_Indices[[i]] !=0 ; & Recombinant_Het1_alleles_Indices[[i]] !=0 & Recombinant_Het2_alleles_Indices[[i]] !=0
		  
		   if(length(Recombinant_Hom1_alleles_Indices[[i]])>1){
                 genoTable[i,(Recombinant_Hom1_alleles_Indices [[i]])] <- "1"
		   }
		   if(length(Recombinant_Hom2_alleles_Indices[[i]])>1){
                 genoTable[i,(Recombinant_Hom2_alleles_Indices[[i]])] <- "1"
		   }
		   
		   if(length(Recombinant_Het1_alleles_Indices[[i]])>1){
                 genoTable[i,(Recombinant_Het1_alleles_Indices[[i]])] <- "0"
		   }
		   if(length(Recombinant_Het2_alleles_Indices[[i]])>1){
                 genoTable[i,(Recombinant_Het2_alleles_Indices[[i]])] <- "0"
		   }
		   
		   Allele_Indices <- c((IA3023_alleles_Indices[[i]]),(Het_alleles1_Indices[[i]]),(Het_alleles2_Indices[[i]]),(Alt_Homozygous_alleles_Indices[[i]]),(Recombinant_Hom1_alleles_Indices[[i]]),(Recombinant_Hom2_alleles_Indices[[i]]),(Recombinant_Het1_alleles_Indices[[i]]),(Recombinant_Het2_alleles_Indices[[i]]))

		   NR_Allele_Indices <- c(NR1_Recombinant_Het1_alleles_Indices[[i]],NR1_Recombinant_Het2_alleles_Indices[[i]],NR1_Recombinant_Het3_alleles_Indices[[i]],NR1_Recombinant_Het4_alleles_Indices[[i]],NR2_Recombinant_Het1_alleles_Indices[[i]],NR2_Recombinant_Het2_alleles_Indices[[i]],NR2_Recombinant_Het3_alleles_Indices[[i]],NR2_Recombinant_Het4_alleles_Indices[[i]])

		   
		   Rest_of_Indices <- setdiff(c(1:nMarkers),c(Allele_Indices,NR_Allele_Indices))
		   
		   Total_Alleles[i] <- length(unique(c(Allele_Indices,NR_Allele_Indices,Rest_of_Indices)))
		   
		   if(length(NR_Allele_Indices)>=1 & NR_Allele_Indices[1] !=0){
                 genoTable[i,(NR_Allele_Indices)] <- "0"
		   } 
		    if(length(Rest_of_Indices)>=1 & Rest_of_Indices[1] !=0){
                 genoTable[i,(Rest_of_Indices)] <- "0"
		   } 
		   
		   
		}
		
	      genoTable_Num <- apply(genoTable,2,as.numeric)
          return(genoTable_Num)
 }


 
###


getQTLIndices <- function(noQTL){  

   nLoci <- noQTL
   nMarkers <- 4289
##### Extract QTL table ######

    if(nLoci== 40){
      QTL_indices<- seq(50,(nMarkers/2),by=53)
    }


    if(nLoci == 400){
      QTL_indices_1<- seq(8,nMarkers,by=12)
      QTL_indices_2 <- seq(50,nMarkers,by=107)
      n_3 <- 400-(length(QTL_indices_1))-(length(QTL_indices_2))
      QTL_indices_3<- sample(c(1:nMarkers),n_3)

      QTL_indices <- c(QTL_indices_1,QTL_indices_2,QTL_indices_3)
    }


    if(nLoci == nMarkers){
      QTL_indices<- c(1:nMarkers)
    }
   
    return(QTL_indices)   

}

####
	 
# LD_Table <- getLD(nextGenGenoTable,400) 


#########################################################################
getQTLTable <- function(GenoTable,noLoci){


	genoTable <- GenoTable
    nIndividuals <-nrow(genoTable)
    genotypicValuesSim <-rep(0,nIndividuals)
    nLoci <- noLoci
	nMarkers <- ncol(genoTable)
  
##### Extract QTL table ######

    if(nLoci== 40){
      QTL_indices<- seq(50,(nMarkers/2),by=53)
    }


    if(nLoci == 400){
      QTL_indices_1<- seq(8,nMarkers,by=12)
      QTL_indices_2 <- seq(50,nMarkers,by=107)
      n_3 <- 400-(length(QTL_indices_1))-(length(QTL_indices_2))
      QTL_indices_3<- sample(c(1:nMarkers),n_3)

      QTL_indices <- c(QTL_indices_1,QTL_indices_2,QTL_indices_3)
    }


    if(nLoci == nMarkers){
      QTL_indices<- c(1:nMarkers)
    }


    QTL_table <- genoTable[,c(QTL_indices)]


	return(QTL_table)

}

###
	


 # missingIndices <- setdiff(c(1:4289),Allele_Indices)
 # Allele_Indices <- c((IA3023_alleles_Indices[[i]]),(Het_alleles1_Indices[[i]]),(Het_alleles2_Indices[[i]]),(Alt_Homozygous_alleles_Indices[[i]]),(Recombinant_Hom1_alleles_Indices[[i]]),(Recombinant_Hom2_alleles_Indices[[i]]),(Recombinant_Het1_alleles_Indices[[i]]),(Recombinant_Het2_alleles_Indices[[i]]))

 # NR_Allele_Indices <- c(NR1_Recombinant_Het1_alleles_Indices[[i]],NR1_Recombinant_Het2_alleles_Indices[[i]],NR1_Recombinant_Het3_alleles_Indices[[i]],NR1_Recombinant_Het4_alleles_Indices[[i]],NR2_Recombinant_Het1_alleles_Indices[[i]],NR2_Recombinant_Het2_alleles_Indices[[i]],NR2_Recombinant_Het3_alleles_Indices[[i]],NR2_Recombinant_Het4_alleles_Indices[[i]])
 
 # Total_Alleles[i] <- length(IA3023_alleles_Indices[[i]])+length(Het_alleles1_Indices[[i]])+length(Het_alleles2_Indices[[i]])+length(Alt_Homozygous_alleles_Indices[[i]])+length(Recombinant_Hom1_alleles_Indices[[i]])+length(Recombinant_Hom2_alleles_Indices[[i]])+length(Recombinant_Het1_alleles_Indices[[i]])+ length(Recombinant_Het2_alleles_Indices[[i]])+ length(NR1_Recombinant_Het1_alleles_Indices[[i]])+ length(NR1_Recombinant_Het2_alleles_Indices[[i]]) + length(NR1_Recombinant_Het3_alleles_Indices[[i]])+ length(NR1_Recombinant_Het4_alleles_Indices[[i]])+ length(NR2_Recombinant_Het1_alleles_Indices[[i]])+ length(NR2_Recombinant_Het2_alleles_Indices[[i]]) + length(NR2_Recombinant_Het3_alleles_Indices[[i]])+ length(NR2_Recombinant_Het4_alleles_Indices[[i]])
 
```
