## Supplementary material for "Factors Affecting Response to Recurrent Genomic Selection in Soybeans": File S1: getPlots_SoyNAM_PredMethods.R

```
#### Function to get plots for dynamics of PG 

 collectOutputVariables <- function(simResults_List,NConditions,NReps,SelFraction,noCycles){ 
	
	    simResults_ListPheno200_GP_Update_List2 <- simResults_List
	
	
	    GenoVal_Sim_NX_2k_List_3c_Pheno200_GP_Update_List2 <- list()
		GenoVal_NX_2k_List_3c_Pheno200_GP_Update_List2 <- list()
		GenoVal_NX_N_List_3c_Pheno200_GP_Update_List2 <- list()
		PhenoVal_Sim_NX_2k_List_3c_Pheno200_GP_Update_List2 <- list()
		PhenoVal_NX_2k_List_3c_Pheno200_GP_Update_List2 <- list()
		PhenoVal_NX_N_List_3c_Pheno200_GP_Update_List2 <- list() 
		attained_GenoValues_List_3c_Pheno200_GP_Update_List2 <- list()
	
        for(j in 1:length(simResults_ListPheno200_GP_Update_List2)){  

			simResults_List2Pheno200_GP_Update <- simResults_ListPheno200_GP_Update_List2[[j]]

			nIndividuals <- 2000
			nCycles <- noCycles
			nConditions <- NConditions
			nreps <- NReps
			nSelected <- SelFraction
	 
			GenoVal_Sim_NX_2k_List_3c_Pheno200_GP_Update <- rep(list(rep(list(matrix(nrow=nIndividuals,ncol=nCycles)),nreps)),nConditions)  
			GenoVal_NX_2k_List_3c_Pheno200_GP_Update <- rep(list(rep(list(matrix(nrow=nIndividuals,ncol=nCycles)),nreps)),nConditions) 
			GenoVal_NX_N_List_3c_Pheno200_GP_Update <- rep(list(rep(list(matrix(nrow=nSelected,ncol=nCycles)),nreps)),nConditions) 

			PhenoVal_Sim_NX_2k_List_3c_Pheno200_GP_Update <- rep(list(rep(list(matrix(nrow=nIndividuals,ncol=nCycles)),nreps)),nConditions)  
			PhenoVal_NX_2k_List_3c_Pheno200_GP_Update <- rep(list(rep(list(matrix(nrow=nIndividuals,ncol=nCycles)),nreps)),nConditions) 
			PhenoVal_NX_N_List_3c_Pheno200_GP_Update <-  rep(list(rep(list(matrix(nrow=nSelected,ncol=nCycles)),nreps)),nConditions) 

			attained_GenoValues_List_3c_Pheno200_GP_Update <- rep(list(rep(list(list()),nreps)),nConditions)
		  
			#for(i in 1:nConditions){ 
			
			  i <- 3
          				
				simResults <-  simResults_List2Pheno200_GP_Update[[i]]

				for (k in 1:length(simResults)){
			
					GenoVal_Sim_NX_2k_List_3c_Pheno200_GP_Update[[i]][[k]] <- simResults[[k]][[1]]
					GenoVal_NX_2k_List_3c_Pheno200_GP_Update[[i]][[k]] <-  simResults[[k]][[2]]
					GenoVal_NX_N_List_3c_Pheno200_GP_Update[[i]][[k]] <-   simResults[[k]][[3]]
			
			  
					PhenoVal_Sim_NX_2k_List_3c_Pheno200_GP_Update[[i]][[k]] <-  simResults[[k]][[4]] 
					PhenoVal_NX_2k_List_3c_Pheno200_GP_Update[[i]][[k]] <-  simResults[[k]][[5]]
					PhenoVal_NX_N_List_3c_Pheno200_GP_Update[[i]][[k]] <-  simResults[[k]][[6]]
		  		
					attained_GenoValues_List_3c_Pheno200_GP_Update[[i]][[k]] <- simResults[[k]][[7]]
						
				}
			
			
	
	
			GenoVal_Sim_NX_2k_List_3c_Pheno200_GP_Update_List2[[j]] <- GenoVal_Sim_NX_2k_List_3c_Pheno200_GP_Update
			GenoVal_NX_2k_List_3c_Pheno200_GP_Update_List2[[j]] <- GenoVal_NX_2k_List_3c_Pheno200_GP_Update
			GenoVal_NX_N_List_3c_Pheno200_GP_Update_List2[[j]] <- GenoVal_NX_N_List_3c_Pheno200_GP_Update 

			PhenoVal_Sim_NX_2k_List_3c_Pheno200_GP_Update_List2[[j]] <- PhenoVal_Sim_NX_2k_List_3c_Pheno200_GP_Update 
			PhenoVal_NX_2k_List_3c_Pheno200_GP_Update_List2[[j]] <- PhenoVal_NX_2k_List_3c_Pheno200_GP_Update 
			PhenoVal_NX_N_List_3c_Pheno200_GP_Update_List2[[j]] <- PhenoVal_NX_N_List_3c_Pheno200_GP_Update 
		
			attained_GenoValues_List_3c_Pheno200_GP_Update_List2[[j]] <- attained_GenoValues_List_3c_Pheno200_GP_Update 
		
		}
		
		
		return(list(GenoVal_Sim_NX_2k_List_3c_Pheno200_GP_Update_List2,GenoVal_NX_2k_List_3c_Pheno200_GP_Update_List2,
		GenoVal_NX_N_List_3c_Pheno200_GP_Update_List2,PhenoVal_Sim_NX_2k_List_3c_Pheno200_GP_Update_List2,	PhenoVal_NX_2k_List_3c_Pheno200_GP_Update_List2,PhenoVal_NX_N_List_3c_Pheno200_GP_Update_List2,		attained_GenoValues_List_3c_Pheno200_GP_Update_List2))                                                   
		
	}

#### 


 getPlots <- function(OutputList){ 

		getResponseMaxPlot(OutputList[[1]])
		getStdGenoVariancePlot(OutputList[[1]]) 
		getPredictionAccuracyPlot(OutputList[[4]],OutputList[[5]]) 
		getAttainedGenoValuePlot(OutputList[[1]])
	 
    }
   

# ResponseSDPlot(OutputList[[1]]) 	

##### a) RsMax
## GenoVal_Sim_NX_2k_List_3c_PhenoPred_GP_List2  <- output_List[[1]]
 
 getResponseMaxPlot <- function(GenoVal_Sim_NX_2k_List_3c_PhenoPred_GP_List2,ncon,nReps){
 
	
	GenoVal_Sim_NX_2k_List_3c_PhenoPred_GP_List <- GenoVal_Sim_NX_2k_List_3c_PhenoPred_GP_List2 
	
	# <-  list(GenoVal_Sim_NX_2k_List_3c_PhenoPred20_GP_List2,GenoVal_Sim_NX_2k_List_3c_PhenoPred50_GP_List2,GenoVal_Sim_NX_2k_List_3c_PhenoPred200_GP_List2)
  
	nSelMethod <- length(GenoVal_Sim_NX_2k_List_3c_PhenoPred_GP_List)
	  
	for(nSel in 1:nSelMethod){

		GenoVal_Sim_NX_2k_List_3c_PhenoPred_GP <- GenoVal_Sim_NX_2k_List_3c_PhenoPred_GP_List[[nSel]]
     		
	  # for(nQTL in 1:3){
		#  for(nSelSch in 1:nSelScheme){
   		#	selMethods <- list(SelMethodPairs2,SelMethodPairs3,SelMethodPairs4)
		#	GenoVal_Sim_NX_2k_List_3c_PhenoPred_GP_List <- GenoVal_Sim_NX_2k_List_3c_PhenoPred_GP_List2[[nSelSch]] 
					
		nCycles <-40
		nreps <- nReps
		cycleNo <- c(1:nCycles)
		nCon <- ncon
				
		leg_col <- c("dark green","blue","black","red","purple","dark orange","gold3")
			
		nSelSch <- 3
			
		lineType <- 1
		pchar <- 1
						
		initCon <- 3
		nConditions <- 6
				
			#for (nCon in initCon:nConditions){ 
					
				nCycles <-40
				Rt <- rep(0,40)
				Rs <- rep(0,40)
				pValue.t <- rep(0,40)

				Gt_Rep <- matrix(rep(0,nreps*nCycles),nrow=nreps,ncol=nCycles)
				GVart_Rep <- matrix(rep(0,nreps*nCycles),nrow=nreps,ncol=nCycles)
				
				
				## mean genotypic value of a population of 2000 individuals every generation for 10 replicates 
				## genotypic variance - variance of genetic values of a ppoulation every generation for 10 replicates 
				## Average of mean genotypic value of 10 reps every generation 
				## Average of genotypic variance of 10 reps every generation 			
				
				
				for(nReps in 1:nreps){ 
				 
				  for (t in 1:nCycles){
					
					Gt_Rep[nReps,t] <- mean(GenoVal_Sim_NX_2k_List_3c_PhenoPred_GP[[nCon]][[nReps]][,t])
					GVart_Rep[nReps,t] <- sd(GenoVal_Sim_NX_2k_List_3c_PhenoPred_GP[[nCon]][[nReps]][,t]) 
				
				  }
				}
											
				R0 <- mean(Gt_Rep[,1])
				sd0 <- mean(GVart_Rep[,1])
							
				Std0 <- abs(R0)/sd0
				Rm <- 200 

				Gt <- apply(Gt_Rep,2,mean)
				
				Rt <- Gt -R0

				Rs = Rt /(Rm - R0)
						
				# if(label[nSel] == "PS"){
					# plotFilename <- paste("Rs_Max_",label[nSel],".png") 
					# plotTitle <- "Standardized Response in Recurrent PS"
				# }else if(label[nSel] != "PS"){ }
					plotFilename <- paste("Rs_Max_GS_Methods.png")
					plotTitle <- "Standardized Response in Recurrent GS"
				
			# nSelSch==3 && nSelSch>1 && 		   png(plotFilename,width=1024,height=768,pointsize=20)
			    
				if( nSel==1){ 
				   par(oma=c(0,0,0,4),mar=c(4,4,4,12))
				
				   plot(cycleNo,(Rs),type="o",ylim=c(0,1),xlab="Cycle Number", ylab="Standardized Response (Rs)",main=plotTitle,lwd=3,lty=lineType,pch=pchar,col=leg_col[nSel])
				}else if((nSel>1)){
				
					lines(cycleNo,(Rs),type="o",pch=pchar,lwd=3,lty=lineType,col=leg_col[nSel])
				
				}
		
	}
		
		#}
		#label <- c("PS","RR-BLUP","BayesB","BL","SVMRBF")
		# legend(41.5,ymax-yoffset1,legend=label,xpd=TRUE,lty=c(1,1),lwd=c(3,3),col=leg_col,cex=0.7,title="Method")
		# dev.off() 
		nTS <- 1
		index <- 1
		Sellabel <- c("1.0%","2.5%","10.0%")
		SelCondition <- c("QTL_40_H0.7","QTL_40_H0.3","QTL_400_H0.7","QTL_400_H0.3","QTL_4289_H0.7","QTL_4289_H0.3")
		SelMethod <- list(c("PS","RRREML","BayesB","BL","SVMRBF"),c("PS","RRREML_1X_14Cyc","BayesB_1X_14Cyc","BL_1X_14Cyc","SVMRBF_1X_14Cyc"))[[nTS]]
			
		
		par(font=2)
		legend(41.5,1.0,legend=SelMethod[1:nSelMethod],xpd=TRUE,lty=c(1,1),lwd=c(3,3),col=leg_col,cex=0.7,title="Method") 
		legend(41.5,0.65,legend=SelCondition[nCon],xpd=TRUE,col="black",pch=c(1,3,5),title="Genetic Architecture",text.width=strwidth("Genetic Architecture")+0.05,cex=0.7)
		legend(41.5,0.45,legend=Sellabel[nSelSch],xpd=TRUE,col="black",pch=c(1,3,5),title=" Selected Top Fraction",text.width=strwidth("Selected Top Fraction"),cex=0.7) 
	 	
 }
 
 
 


####################

####################
#2 ### Standardized Variance 

 getStdGenoVariancePlot <- function(GenoVal_Sim_NX_2k_List_3c_PhenoPred_GP_List2,ncon,nReps){
    
	#nSelScheme <- 3  
	#GenoVal_Sim_NX_2k_List_3c_PhenoPred_GP_List2 <- 	#list(GenoVal_Sim_NX_2k_List_3c_PhenoPred20_GP_List2,GenoVal_Sim_NX_2k_List_3c_PhenoPred50_GP_List2,GenoVal_Sim_NX_2k_List_3c_PhenoPred200_GP_List2)
  
	nSelMethod <- length(GenoVal_Sim_NX_2k_List_3c_PhenoPred_GP_List2)
	label <- c("PS","RR-BLUP","BayesB","BL","SVMRBF")  
	  
	for(nSel in 1:nSelMethod){
      	
		initCon <- 3
		nConditions <- 6
		
		#for(nSelSch in 1:nSelScheme){
  		#GenoVal_Sim_NX_2k_List_3c_PhenoPred_GP_List <- GenoVal_Sim_NX_2k_List_3c_PhenoPred_GP_List2[[nSel]] 
		
		GenoVal_Sim_NX_2k_List <- GenoVal_Sim_NX_2k_List_3c_PhenoPred_GP_List2[[nSel]]
		
		nCycles <- 40
		nreps <- nReps
		nCon <- ncon
		cycleNo <- c(1:nCycles)
	
		leg_col <- c("dark green","blue","black","red","purple","gray")
		
       	lineType <- 1
		pchar <-1 
		nSelSch <- 3
	  	  	  
		#for(nCon in initCon:nConditions) {	
		

		   Var_SimGenoVal_2k <- lapply(GenoVal_Sim_NX_2k_List[[nCon]],function(x) apply(x,2,var))
		
		   Var_SimGeno_2k_plot <- c()
	  
		   for(nReps in 1:nreps) { 
		
				Var_SimGeno_2k_plot <- rbind(Var_SimGeno_2k_plot,unlist(Var_SimGenoVal_2k[[nReps]]))
		
		   }
			
		   MeanVar_SimGeno_2k_Plot <- rep(0,nCycles)
		
		   for(i in 1:nCycles) { 
			  
			  MeanVar_SimGeno_2k_Plot[i] <- mean(Var_SimGeno_2k_plot[,i])
			  
		   }
			
			MeanVar_Max <- max(MeanVar_SimGeno_2k_Plot)
			
			MeanVar_Adjusted <- rep(0,nCycles)
			
			for(i in 2:nCycles){
				MeanVar_Adjusted[i] <- (MeanVar_SimGeno_2k_Plot[1]-MeanVar_SimGeno_2k_Plot[i])/(MeanVar_Max)
			}

			# if(label[nSel] == "PS"){
				# plotFilename <- paste("Standardized_Variance_GS_",label[nSel],"_SelSch_",nSelSch,".png",sep="") 
  				# plotTitle <- paste("Genotypic Variance in Recurrent PS","_SelSch_",nSelSch,".png",sep="")
			# }else if(label[nSel] != "PS"){ 
				plotFilename <- paste("Standardized_Variance_GS_Methods.png",sep="") 
  				plotTitle <- "Genotypic Variance in Recurrent GS"
						
			# if(nCon==initCon && nSelSch ==1){ 
	    
			if(nSel==1){
				par(oma=c(0,0,0,4),mar=c(4,4,4,12))
				
				plot(cycleNo,(1-MeanVar_Adjusted),ylim=c(0,1),xlim=c(1,nCycles),type="o",lty=lineType,pch=pchar,lwd=3,cex.lab=1.25,cex.main=1.25,font.lab=2,col=leg_col[nSel],xlab="Cycle Number"
				     ,ylab="Standardized Genotypic Variance",main=plotTitle)
			
			}else if(nSel>1){
	
			    lines(cycleNo,(1-MeanVar_Adjusted),type="o",lwd=3,lty=lineType,pch=pchar,col=leg_col[nSel])
			}
		
	}
 
    
		  
    index <- 1
	
	
	index <- 1
	Sellabel <- c("1.0%","2.5%","10.0%")
	SelCondition <- c("QTL_40_H0.7","QTL_40_H0.3","QTL_400_H0.7","QTL_400_H0.3","QTL_4289_H0.7","QTL_4289_H0.3")
	SelMethod <- c("PS","RRREML","BayesB","BL","SVMRBF")
			
		
	par(font=2)
	legend(41.5,1.0,legend=SelMethod[1:nSelMethod],xpd=TRUE,lty=c(1,1),lwd=c(3,3),col=leg_col,cex=0.7,title="Method") 
	legend(41.5,0.65,legend=SelCondition[nCon],xpd=TRUE,col="black",pch=c(1,3,5),title="Genetic Architecture",text.width=strwidth("Genetic Architecture")+0.045,cex=0.7)
	legend(41.5,0.45,legend=Sellabel[nSelSch],xpd=TRUE,col="black",pch=c(1,3,5),title=" Selected Top Fraction",text.width=strwidth("Selected Top Fraction")+0.02,cex=0.7) 
	
	# dev.off()
	
   }
  
 
#####################################################################################
## 4 ### Attained Genotypic Values
  
getAttainedGenoValuePlot <- function(GenoVal_Sim_NX_2k_List_3c_PhenoPred_GP_List2,ncon,nReps){
  
	  
	nSelMethod <- length(GenoVal_Sim_NX_2k_List_3c_PhenoPred_GP_List2)
	label <- c("PS","RR-REML","BayesB","BL","SVMRBF")  
	label <- "RR-BLUP" 
	nSelSch <- 3
	nCon <- ncon
	nrep <- nReps
	
	for(nSel in 1:nSelMethod){
     	
		initCon <- 3
		nConditions <- 6
		
	    GenoVal_Sim_NX_2k_List_3c_PhenoPred_GP_List <- GenoVal_Sim_NX_2k_List_3c_PhenoPred_GP_List2
		
		GenoVal_Sim_NX_2k_List<- GenoVal_Sim_NX_2k_List_3c_PhenoPred_GP_List[[nSel]]
		
		nCycles <-40
		nreps <- 1
		cycleNo <- c(1:nCycles)

	 	leg_col <- c("dark green","blue","black","red","purple","gray")
			
		lineType <- 1
		pchar <-1 
		
		#for(nCon in initCon:nConditions){  
		
		    Avg_Sim_Attained_Geno_2k_Plot_RepTable <- matrix(rep(0,nCycles*nreps),nrow=nreps,ncol=nCycles)
			Avg_Sim_Attained_Geno_2k_Plot <- rep(0,nCycles)
	  
			SimGenoVal_NX_2k_Plot<- GenoVal_Sim_NX_2k_List[[nCon]]
	  
	 
			for(j in 1:nreps){	 
		  
				for(i in 1:nCycles) { 
		 		
			
					Avg_Sim_Attained_Geno_2k_Plot_RepTable[j,i] <- max(SimGenoVal_NX_2k_Plot[[j]][,i])
					
				}
			}
	  
	
			Avg_Sim_Attained_Geno_2k_Plot <- apply(Avg_Sim_Attained_Geno_2k_Plot_RepTable,2,mean)

	   
			ylim_min <- 0
			ylim_max <- 200
	   
			# if(label[nSel] == "PS"){
				# plotFilename <- paste("Attained Genotypic Value_",label[nSel],".png") 
  				# plotTitle <- "Attained Genotypic Value in Recurrent PS"
			# }else if(label[nSel] != "PS"){ 
				
			plotFilename <- paste("Attained Genotypic Value_GS Methods.png")
  			plotTitle <- "Attained Genotypic Value in Recurrent GS"
			
			
			if(nSel==1){ 
   			#	png(plotFilename,width=1352,height=748,pointsize=20)
				par(oma=c(0,0,0,4),mar=c(4,4,4,12))
			
				plot(cycleNo,Avg_Sim_Attained_Geno_2k_Plot,ylim=c(0,ylim_max),xlim=c(0,nCycles),xaxp=c(0,40,4),type="o",lwd=3,col=leg_col[nSel],cex.lab=1.25,cex.main=1.5,font.lab=2,xlab="Cycle Number",ylab="Attained Genotypic Value",main=plotTitle,pch=pchar,lty=lineType)
			}else if(((nSel>1))){ 
			#|| ((nSelSch !=1)&&(nCon>=initCon))){

			    lines(cycleNo,Avg_Sim_Attained_Geno_2k_Plot,ylim=c(ylim_min,ylim_max),type="o",lwd=3,pch=pchar,lty=lineType,col=leg_col[nSel])
			}
	   
	  }
    
      abline(h= 200,lty=2,col=leg_col[2])
	  abline(h= 80,lty=2,col=leg_col[2])
	  abline(h= 25,lty=2,col=leg_col[2])
	  
	  		  
	  index <- 1
		Sellabel <- c("1.0%","2.5%","10.0%")
		SelCondition <- c("QTL_40_H0.7","QTL_40_H0.3","QTL_400_H0.7","QTL_400_H0.3","QTL_4289_H0.7","QTL_4289_H0.3")
		SelMethod <- c("PS","RRREML","BayesB","BL","SVMRBF")
			
		
		par(font=2)
		legend(41.5,150,legend=SelMethod[1:nSelMethod],xpd=TRUE,lty=c(1,1),lwd=c(3,3),col=leg_col,cex=0.7,title="Method") 
		legend(41.5,100,legend=SelCondition[nCon],xpd=TRUE,col="black",pch=c(1,3,5),title="Genetic Architecture",text.width=strwidth("Genetic Architecture")+0.05,cex=0.7)
		legend(41.5,70,legend=Sellabel[nSelSch],xpd=TRUE,col="black",pch=c(1,3,5),title=" Selected Top Fraction",text.width=strwidth("Selected Top Fraction")+0.02,cex=0.7) 
	  	  
	    #dev.off()
    
	}
 

 
##########################################################
##5 ## Prediction accuracy 

# nSelScheme <- 3 , OutputList[[4]],OutputList[[5]]

 getPredictionAccuracyPlot <- function(PhenoVal_Sim_NX_2k_List_3c_PhenoPred_GP_List2,PhenoVal_NX_2k_List_3c_PhenoPred_GP_List2,ncon,nReps){ 
 
	
	nSelMethod <- length(PhenoVal_Sim_NX_2k_List_3c_PhenoPred_GP_List2)
 	nSelSch <- 3
	nCon <- ncon
	
	for(nSel in 1:nSelMethod){

      	
		initCon <- 3
		nConditions <- 6
	
		nCycles <-40
		nreps <- nReps
		cycleNo <- c(1:nCycles)
		Cycle_No <- c(1:nCycles)
			   	
		PhenoVal_Sim_NX_2k_List <- PhenoVal_Sim_NX_2k_List_3c_PhenoPred_GP_List2[[nSel]]
		
		PhenoVal_NX_2k_List <- PhenoVal_NX_2k_List_3c_PhenoPred_GP_List2[[nSel]]
       
	   	
		leg_col <- c("dark green","blue","black","red","purple","dark orange","gray")

		lineType <- 1
		pchar <-1 
		
						
		dataR <- c()
		dataR2 <- c()
		       
           	   
		rho1_List_3c <-rep(list(rep(list(list()),10,)),6)
		  
		rho1 <-rep(0,nCycles)
				
		for(nReps in 1:nreps){
				
				PhenoVal_Sim <- PhenoVal_Sim_NX_2k_List[[nCon]][[nReps]]
				PhenoVal <- PhenoVal_NX_2k_List[[nCon]][[nReps]]
								
				for(i in 1:nCycles){
				  
					rho1[i] <- cor(PhenoVal_Sim[,i],PhenoVal[,i]) 
				  
				}
				rho1_List_3c[[nCon]][[nReps]] <- rho1
		} 
		
		   
			rho_List_3c_Table <- matrix(rep(0,nCycles*nreps),nrow=nreps,ncol=nCycles)
			rho1_List <- list()
			  
		#	for(nCon in initCon:nConditions) {
			
			for(nReps in 1:nreps){ 
				for(i in 1:nCycles){ 
						
						   rho_List_3c_Table[nReps,i] <- rho1_List_3c[[nCon]][[nreps]][i]
							
				}
			}		
					
				rho1_List[[nCon]] <- rho_List_3c_Table
			#}
			
			rho1_List_3c_final <- list()
			rho1_List_3c_SE_final <- list()
			rho_final <- rep(0,40)
			rho_final_se <- rep(0,40)
			
			# for(nCon in initCon:nConditions){ 
			   
			   		
				rho_final<- apply(rho1_List[[nCon]],2,mean)
				rho_final_sd <-apply(rho1_List[[nCon]],2,sd)
			  	rho_final_se <- rho_final_sd/sqrt(nreps)
				 
				rho1_List_3c_final[[nCon]] <- rho_final
				
				rho1_List_3c_SE_final[[nCon]] <-rho_final_se
						    			   
				Pred_accuracy <- rho1_List_3c_final[[nCon]]
				
				Pred_accuracy_se <- rho1_List_3c_SE_final[[nCon]]
				
				NAIndices<- which(is.na(Pred_accuracy))
				
				plotFilename <- paste("Prediction Accuracy_GS_Method.png") 
				plotTitle <- "Prediction Accuracy in Recurrent GS"
				
						
				 if(nSel==1){
	#			   png(plotFilename,width=1532,height=748,pointsize=20)
				   par(oma=c(0,0,0,4),mar=c(4,4,4,12))
			  	   plot(Cycle_No,Pred_accuracy,type="o",col=leg_col[nSel],lwd=3,xlim=c(1,nCycles),lty=lineType,pch=pchar,ylim=c(-0.2,1),xlab="Cycle Number",ylab="Prediction Accuracy",main=plotTitle)
			  
				   #}else if(((nSelSch==1)&&(nCon>initCon)) || ((nSelSch!=1)&&(nCon>=initCon))){
		         }else if((nSel>1)){
		
					lines(Cycle_No,Pred_accuracy,type="o",col=leg_col[nSel],lwd=3,lty=lineType,pch=pchar) 
					abline(h=0,col="black",lwd=2,lty=1)
				}
		}	
		
		index <- 1
				
				  
	    Sellabel <- c("1.0%","2.5%","10.0%")
		SelCondition <- c("QTL_40_H0.7","QTL_40_H0.3","QTL_400_H0.7","QTL_400_H0.3","QTL_4289_H0.7","QTL_4289_H0.3")
		SelMethod <- c("PS","RRREML","BayesB","BL","SVMRBF")
			
		
		par(font=2)
		legend(41.5,1.0,legend=SelMethod[1:nSelMethod],xpd=TRUE,lty=c(1,1),lwd=c(3,3),col=leg_col[1:nSelMethod],cex=0.8,title="Method") 
		legend(41.5,0.65,legend=SelCondition[nCon],xpd=TRUE,col="black",pch=c(1,3,5),title="Genetic Architecture",text.width=strwidth("Genetic Architecture")+0.10,cex=0.7)
		legend(41.5,0.45,legend=Sellabel[nSelSch],xpd=TRUE,col="black",pch=c(1,3,5),title=" Selected Top Fraction",text.width=strwidth("Selected Top Fraction")+0.10,cex=0.7) 
		
		#dev.off()
		
    }		
	
###################################################

## 1b) Response standardized to sd of base population genotypic values  
 
 getResponseSDPlot <- function(GenoVal_Sim_NX_2k_List_3c_PhenoPred_GP_List2){

    nSelScheme <- 3
 
    GenoVal_Sim_NX_2k_List_3c_PhenoPred_GP_List <- GenoVal_Sim_NX_2k_List_3c_PhenoPred_GP_List2
	
	# GenoVal_Sim_NX_2k_List_3c_PhenoPred_GP_List2 <- 
	# list(GenoVal_Sim_NX_2k_List_3c_PhenoPred20_GP_List2,GenoVal_Sim_NX_2k_List_3c_PhenoPred50_GP_List2,GenoVal_Sim_NX_2k_List_3c_PhenoPred200_GP_List2)
 
 
	nSelMethod <- length(GenoVal_Sim_NX_2k_List_3c_PhenoPred_GP_List)
	  
	for(nSel in 1:nSelMethod){

		GenoVal_Sim_NX_2k_List_3c_PhenoPred_GP <- GenoVal_Sim_NX_2k_List_3c_PhenoPred_GP_List[[nSel]]
     
		label <- c("PS","RR-BLUP","BayesB","BL","SVMRBF")
    
		# for(nQTL in 1:3){
		
		#  for(nSelSch in 1:nSelScheme){

   		
			selMethods <- list(SelMethodPairs2,SelMethodPairs3,SelMethodPairs4)

		#	GenoVal_Sim_NX_2k_List_3c_PhenoPred_GP_List <- GenoVal_Sim_NX_2k_List_3c_PhenoPred_GP_List2[[nSelSch]] 
			
			GenoVal_Sim_NX_2k_List_3c_PhenoPred_GP <- GenoVal_Sim_NX_2k_List_3c_PhenoPred_GP_List[[nSel]]
			
			nCycles <-40
			nreps <- 1
			cycleNo <- c(1:nCycles)

					
			leg_col <- c("dark green","blue","black","red","purple","dark orange","gold3")
			
			
			lineType <- 1
			pchar <-1 
			  
									
			initCon <- 3
			nConditions <- 6
			ymin <- 0
			ymax <- 0.25
			nSelSch <- 3				
			
			# for (nCon in initCon:nConditions
			    nCon <- 3
				
				nCycles <-40
				nreps <- 10
			
				Rt <- rep(0,40)
				Rs <- rep(0,40)
				pValue.t <- rep(0,40)

				Gt_Rep <- matrix(rep(0,nreps*nCycles),nrow=nreps,ncol=nCycles)
				GVart_Rep <- matrix(rep(0,nreps*nCycles),nrow=nreps,ncol=nCycles)
				
				
				## mean genotypic value of a population of 2000 individuals every generation for 10 replicates 
				## genotypic variance - variance of genetic values of a ppoulation every generation for 10 replicates 
				## Average of mean genotypic value of 10 reps every generation 
				## Average of genotypic variance of 10 reps every generation 			
				
				
				for(nReps in 1:nreps){ 
				 
				  for (t in 1:nCycles){
					
					Gt_Rep[nReps,t] <- mean(GenoVal_Sim_NX_2k_List_3c_PhenoPred_GP[[nCon]][[nReps]][,t])
					GVart_Rep[nReps,t] <- sd(GenoVal_Sim_NX_2k_List_3c_PhenoPred_GP[[nCon]][[nReps]][,t]) 
				
				  }
				}
											
				R0 <- mean(Gt_Rep[,1])
				sd0 <- mean(GVart_Rep[,1])
							
				Std0 <- abs(R0)/sd0
				Rm <- 200 

				Gt <- apply(Gt_Rep,2,mean)
				
				Rt <- Gt -R0

				Rs = Rt /sd0
				Rs_Norm <- Rs/norm(Rs,"2")
						
				if(label[nSel] == "PS"){
					plotFilename <- paste("Rs_SD_",label[nSel],"_SelSch_",nSelSch,".png") 
					plotTitle <- paste("Standardized_Response in Recurrent ","PS",sep="")
				}else if(label[nSel] != "PS"){ 
					plotFilename <- paste("Rs_SD_",label[nSel],"_SelSch_",nSelSch,".png") 
					plotTitle <- paste("Standardized Response_in Recurrent_GS_Model-",label[nSel],sep="")
				}
							
				if(nCon==initCon){
				   #png(plotFilename,width=1024,height=768,pointsize=20)

			       par(oma=c(0,0,0,4),mar=c(4,4,4,8))
					
				   plot(cycleNo,(Rs_Norm),type="o",ylim=c(ymin,ymax),xlab="Cycle Number", ylab="Standardized Long-term Genetic Gain (Rs)",main=plotTitle,lwd=3,lty=lineType,pch=pchar,col=leg_col[nSel])
				}else if((nCon >initCon )){
				
					lines(cycleNo,(Rs_Norm),type="o",pch=pchar,lwd=3,lty=lineType,col=leg_col[nSel])
				
				}
		
		}	
				
		index <- 1
		Sellabel <- c("1.0%","2.5%","10.0%")
		sel<- c("QTL_40_H0.7","QTL_40_H0.3","QTL_400_H0.7","QTL_400_H0.3","QTL_4289_H0.7","QTL_4289_H0.3")
		
		par(font=2)
		# legend(41.5,ymax-yoffset1,legend=label,xpd=TRUE,lty=c(1,1),lwd=c(3,3),col=leg_col,cex=0.7,title="Method")
		
		legend(41.5,0.16,legend=Sellabel[nSelSch],xpd=TRUE,col="black",pch=c(1,3,5)[nSelSch],title=" Selected Top Fraction",text.width=strwidth("Selected Top Fraction")+0.05,cex=0.8) 
		
		legend(41.5,0.24,legend=sel[nSel],xpd=TRUE,lty=c(1,1),lwd=c(3,3),col=leg_col,cex=0.7,title="Condition") 
	 
	    dev.off() 
	}

  

## 1c) ### Rs_SD across QTL 
 getResponseSDPlot_QTL_V2 <- function(GenoVal_Sim_NX_2k_List_3c_PhenoPred_GP_List2){
  
 nSelScheme <- 3
 
 GenoVal_Sim_NX_2k_List_3c_PhenoPred_GP_List <- GenoVal_Sim_NX_2k_List_3c_PhenoPred_GP_List2
 nSelMethod <- length(GenoVal_Sim_NX_2k_List_3c_PhenoPred_GP_List)
	  
 for(nSel in 1:nSelMethod){

    # GenoVal_Sim_NX_2k_List_3c_PhenoPred_GP <- GenoVal_Sim_NX_2k_List_3c_PhenoPred_GP_List[[nSel]]
     
    label <- c("PS","RR-BLUP","BayesB","BL","SVMRBF")
    label <- "RR-BLUP"
	
    for(nQTL in 1:3){
		
		
		nConVector <- c(2,4,6)
		initConVector <- c(1,3,5)
		
		nConditions <-nConVector[nQTL]
		initCon <- initConVector[nQTL]
		pchar <- c(1,3,5,7,9,11,13)
		
		#for (nCon in initCon:nConditions){		
        	
			#for(nSelSch in 1:nSelScheme){
			# GenoVal_Sim_NX_2k_List_3c_PhenoPred_GP_List <- GenoVal_Sim_NX_2k_List_3c_PhenoPred_GP_List2[[nSelSch]] 
			
			GenoVal_Sim_NX_2k_List_3c_PhenoPred_GP <- GenoVal_Sim_NX_2k_List_3c_PhenoPred_GP_List[[nSel]]
			
			nCycles <-40
			nreps <- 10
			cycleNo <- c(1:nCycles)

					
			leg_col <- c("dark orange","blue","black","red","purple","dark green","gold3")
			
			lineType <- 1
			pchar <-1 
			
			ymin <- 0
			ymax <- 0.25
									
			nCycles <-40
			nreps <- 1
			
			Rt <- rep(0,40)
			Rs <- rep(0,40)
			pValue.t <- rep(0,40)

			Gt_Rep <- matrix(rep(0,nreps*nCycles),nrow=nreps,ncol=nCycles)
			GVart_Rep <- matrix(rep(0,nreps*nCycles),nrow=nreps,ncol=nCycles)
				
				
			## mean genotypic value of a population of 2000 individuals every generation for 10 replicates 
			## genotypic variance - variance of genetic values of a ppoulation every generation for 10 replicates 
			## Average of mean genotypic value of 10 reps every generation 
			## Average of genotypic variance of 10 reps every generation 			
				
				
			for(nReps in 1:nreps){ 
				 
				  for (t in 1:nCycles){
					
					Gt_Rep[nReps,t] <- mean(GenoVal_Sim_NX_2k_List_3c_PhenoPred_GP[[nCon]][[nReps]][,t])
					GVart_Rep[nReps,t] <- sd(GenoVal_Sim_NX_2k_List_3c_PhenoPred_GP[[nCon]][[nReps]][,t]) 
				
				  }
			}
											
				R0 <- mean(Gt_Rep[,1])
				sd0 <- mean(GVart_Rep[,1])
							
				Std0 <- abs(R0)/sd0
				Rm <- 200 

				Gt <- apply(Gt_Rep,2,mean)
				
				Rt <- Gt -R0

				Rs = Rt /sd0
				Rs_Norm <- Rs/norm(Rs,"2")
						
				if(label[nSel] == "PS"){
					plotFilename <- paste("Rs_SD_",label[nSel],"_QTL_",nQTL,".png") 
					plotTitle <- paste("Standardized_Response in Recurrent ","PS",sep="")
				}else if(label[nSel] != "PS"){ 
					plotFilename <- paste("Rs_SD_",label[nSel],"_QTL_",nQTL,".png") 
					plotTitle <- paste("Standardized Response_in Recurrent_GS_Model-",label[nSel],sep="")
				}
							
				if(nSelSch==1 && nCon==initCon){
				  # png(plotFilename,width=1024,height=768,pointsize=20)

			       par(oma=c(0,0,0,4),mar=c(4,4,4,8))
					
				   plot(cycleNo,(Rs_Norm),type="o",ylim=c(ymin,ymax),xlab="Cycle Number", ylab="Standardized Long-term Genetic Gain (Rs)",main=plotTitle,lwd=3,lty=lineType,pch=pchar,col=leg_col[nCon])
				}else if((nSelSch==1 &&(nCon >initCon )) || ((nSelSch>1) && (nCon>=initCon))){
				
					lines(cycleNo,(Rs_Norm),type="o",pch=pchar,lwd=3,lty=lineType,col=leg_col[nCon])
				
				}
		
		}
	
				
		index <- 1
		Sellabel <- c("1.0%","2.5%","10.0%")
		sel<- c("QTL_40_H0.7","QTL_40_H0.3","QTL_400_H0.7","QTL_400_H0.3","QTL_4289_H0.7","QTL_4289_H0.3")
		
		par(font=2)
		# legend(41.5,ymax-yoffset1,legend=label,xpd=TRUE,lty=c(1,1),lwd=c(3,3),col=leg_col,cex=0.7,title="Method")
		
		legend(41.5,0.16,legend=Sellabel[nSelSch],xpd=TRUE,col="black",pch=c(1,3,5),title=" Selected Top Fraction",text.width=strwidth("Selected Top Fraction")+0.05,cex=0.7) 
		
		legend(41.5,0.24,legend=sel[nCon],xpd=TRUE,lty=c(1,1),lwd=c(3,3),col=leg_col[nCon],cex=0.7,title="Condition") 
	 
	    dev.off() 
	}
  }


    
	
```
