## Supplementary material for "Factors Affecting Response to Recurrent Genomic Selection in Soybeans": File S1: GP_LMap.R

```
### Create RF Table 
## 

getNAMLinkageMapTable<- function(){

#data(package="SoyNAMPredictionMethods")
  
NAM_Link_Map <- NAM_LINKAGE_MAP

#levels(factor(NAM_Link_Map[,"LGroup"]))

RF_Vec_New <- c()
LG_Vec_New <- c() 
MarkerName_Vec_New <- c() 
MapOrder_Vec_New <- c() 
Position_Vec_New <- c()

##no_LG - number of linkage groups

no_LG <- 20

a<-rep(0,no_LG)
b<-rep(0,no_LG)

for(i in 1:no_LG){   

	trial1 <- NAM_Link_Map[which(NAM_Link_Map[,"LGroup"] %in% i),"RecombinationF"]
	trial1_LG <- NAM_Link_Map[which(NAM_Link_Map[,"LGroup"] %in% i),"LGroup"]
	trial1_MarkerName<- as.character(NAM_Link_Map[which(NAM_Link_Map[,"LGroup"] %in% i),"Marker_label"])
	trial1_MapOrder <- NAM_Link_Map[which(NAM_Link_Map[,"LGroup"] %in% i),"Map_Order"] 
	trial1_Position <- NAM_Link_Map[which(NAM_Link_Map[,"LGroup"] %in% i),"Position_Wma2.v1"] 
	  
	trial1[1]<-0.5
	trial1_new <- trial1


	RF_Vec_New <-c(RF_Vec_New,trial1_new) 
	LG_Vec_New <-c(LG_Vec_New,trial1_LG) 
	MarkerName_Vec_New <-c(MarkerName_Vec_New,trial1_MarkerName) 
	MapOrder_Vec_New <-c(MapOrder_Vec_New,trial1_MapOrder) 
	Position_Vec_New <-c(Position_Vec_New,trial1_Position) 

}
 
 NAM_LinkMap_New <- cbind(as.numeric(LG_Vec_New),as.numeric(MapOrder_Vec_New),as.numeric(RF_Vec_New),as.numeric(Position_Vec_New))
 colnames(NAM_LinkMap_New) <- c("Linkage_Group","Map_Order","Recombination_Frequency","Position")
 rownames(NAM_LinkMap_New) <- as.character(MarkerName_Vec_New) 

 return(NAM_LinkMap_New)
}


###################
```
