## Supplementary material for "Factors Affecting Response to Recurrent Genomic Selection in Soybeans": File S1: RcppExports.R

```
# Generated by using Rcpp::compileAttributes() -> do not edit by hand
# Generator token: 10BE3573-1514-4C36-9D1C-5A225CD40393

LD <- function(p1, q1, a, pro, D, D1, R2) {
    .Call('_SoyNAMPredictionMethods_LD', PACKAGE = 'SoyNAMPredictionMethods', p1, q1, a, pro, D, D1, R2)
}

LD_V3 <- function(p1, q1, a, pro, D, D1, R2) {
    .Call('_SoyNAMPredictionMethods_LD_V3', PACKAGE = 'SoyNAMPredictionMethods', p1, q1, a, pro, D, D1, R2)
}

rrReml <- function(y, gen, maxit = 500L, tol = 10e-6) {
    .Call('_SoyNAMPredictionMethods_rrReml', PACKAGE = 'SoyNAMPredictionMethods', y, gen, maxit, tol)
}
```
