## Supplementary material for "Factors Affecting Response to Recurrent Genomic Selection in Soybeans": File S1: RunSimulations_GS.R

```
runSimulations_GS_40X <- function(NAMBasePopulationData,GSModels,ModelType,no_QTL=400,h2=0.7,SelFraction=200,nRep=1,nReps,parameter_List=list(),default,PlatForm,NAM_LinkMap_New,alleleConversionTable_Combined){
    
   
  if(PlatForm =="Windows" || PlatForm =="windows"){
    platForm <- "Windows"
  }
  
  noQTLs <- no_QTL
  H2 <- h2
  numberSelected <- SelFraction
  modelType <- ModelType
  nrep <- nReps
  
  if(no_QTL==400 & h2==0.7){
      nCon <- 3
  }
  
  AlleleConversionTable_Combined <- alleleConversionTable_Combined
  Default <- default
  ## GS Models 
  
  model_Sol_Pheno_List <- GSModels[[2]]
  model_Sol_Geno_List <- GSModels[[1]] 
  
  ## NAMBase population data
  
  genotypicValuesSimTableList_Reps <-  NAMBasePopulationData[[1]]
  phenotypicValuesSimTableList_Reps <- NAMBasePopulationData[[2]]
  genotypicValuesSimList_Reps  <- NAMBasePopulationData[[3]]
  phenotypicValuesSimList_Reps <- NAMBasePopulationData[[4]]
  trainIndicesList_Reps <- NAMBasePopulationData[[5]]
  testIndicesList_Reps <- NAMBasePopulationData[[6]]
  trainGenoTableList_Reps <- NAMBasePopulationData[[7]]
  trainGenoNewTableList_Reps <- NAMBasePopulationData[[8]]
  testGenoTableList_Reps <- NAMBasePopulationData[[9]]
  testGenoNewTableList_Reps <-  NAMBasePopulationData[[10]]
  trainSimGenoValuesTableList_Reps <- NAMBasePopulationData[[11]]
  trainSimPhenoValuesTableList_Reps <- NAMBasePopulationData[[12]]
  testSimGenoValuesTableList_Reps <- NAMBasePopulationData[[13]]
  testSimPhenoValuesTableList_Reps <- NAMBasePopulationData[[14]]
  F_Progeny_List_Reps <-NAMBasePopulationData[[15]]
  errorVarList_Reps <- NAMBasePopulationData[[16]]
  
  
  
  model_Sol_Pheno <-  model_Sol_Pheno_List[[nCon]][[nrep]]
  model_Sol_Geno <-  model_Sol_Geno_List[[nCon]][[nrep]]
  F5_Progeny_List <- F_Progeny_List_Reps[[nrep]][[nCon]]
  genoValSimList <- genotypicValuesSimList_Reps[[nrep]][[nCon]]
  phenoValSimList <- phenotypicValuesSimList_Reps[[nrep]][[nCon]]
  varEList <- errorVarList_Reps[[nrep]][[nCon]]
  
  trainGenoNewTable <- trainGenoNewTableList_Reps[[nrep]][[nCon]]
  trainSimGenoValTable <-  trainSimGenoValuesTableList_Reps[[nrep]][[nCon]]
  trainSimPhenoValTable <- trainSimPhenoValuesTableList_Reps[[nrep]][[nCon]]
  
  
  ### Parameter List
  
  if(Default==TRUE){
    if(modelType!="PS"){
      Parameter_List <- list(numberSelected=200,noCrosses=200,no_Progeny=10,noCycles=40,selectionOnGenoValues=FALSE,selectionOnSimulatedValues=FALSE,ModelRetrain=FALSE,RetrainFrequency=1,ModelUpdate=FALSE,UpdateType="FULLSET",UpdateFrequency=1,weighted=FALSE,weightingMethod="JM",AlphaShape=0.025,BetaShape=0.1,TimeHorizon=40,TrainTableWindowSize=14,NAM_LinkMap=NAM_LinkMap_New,gIndStats=TRUE,AlleleConversionTable_Combined)
    }else if(modelType=="PS"){ 
      Parameter_List <- list(numberSelected=200,noCrosses=200,no_Progeny=10,noCycles=40,selectionOnGenoValues=FALSE,selectionOnSimulatedValues=TRUE,ModelRetrain=FALSE,RetrainFrequency=1,ModelUpdate=FALSE,UpdateType="FULLSET",UpdateFrequency=1,weighted=FALSE,weightingMethod="JM",AlphaShape=0.025,BetaShape=0.1,TimeHorizon=40,TrainTableWindowSize=14,NAM_LinkMap=NAM_LinkMap_New,gIndStats=TRUE,AlleleConversionTable_Combined)
    }
   }
   if(Default==FALSE){ 
    
    Parameter_List <- parameter_List
   }
  
  i <- nCon
  k<- nrep
  BreedingDesign <- "BD2"
  
  
  numberSelected <- Parameter_List[[1]]
  noCrosses <- Parameter_List[[2]]
  no_Progeny <- Parameter_List[[3]]
  noCycles <- Parameter_List[[4]]
  selectionOnGenoValues <- Parameter_List[[5]]
  selectionOnSimulatedValues <- Parameter_List[[6]]
  ModelRetrain <- Parameter_List[[7]]
  RetrainFrequency <- Parameter_List[[8]]
  ModelUpdate <- Parameter_List[[9]]
  UpdateType <- Parameter_List[[10]]
  UpdateFrequency <- Parameter_List[[11]]
  
  weighted <- Parameter_List[[12]]
  weightingMethod <- Parameter_List[[13]]
  AlphaShape <- Parameter_List[[14]]
  BetaShape <- Parameter_List[[15]]
  TimeHorizon <- Parameter_List[[16]]
  TrainTableWindowSize <- Parameter_List[[17]]
  NAM_LinkMap <- Parameter_List[[18]]
  gIndStats <- Parameter_List[[19]]
  alleleConversionTable_Combined <- apply(Parameter_List[[20]],2,as.character)
  
  
####
if(platForm != "Windows"){
  if(modelType=="SVMRBF"){
    simResults <- runSimulations40X_ML_Update(model_Sol_Pheno,model_Sol_Geno,F5_Progeny_List,genoValSimList,phenoValSimList,varEList,noQTLs,H2,numberSelected,noCrosses,no_Progeny,noCycles,selectionOnGenoValues,selectionOnSimulatedValues,ModelRetrain,RetrainFrequency,ModelUpdate,UpdateType,UpdateFrequency,i,k,ModelType,weighted,weightingMethod,AlphaShape,BetaShape,TimeHorizon,trainGenoNewTable,trainSimGenoValTable,trainSimPhenoValTable,TrainTableWindowSize,NAM_LinkMap,gIndStats,alleleConversionTable_Combined)
  }else if(modelType=="BayesB" | modelType=="BL"){
    simResults <- runSimulations40X_Bayes_WGS_V2(model_Sol_Pheno,model_Sol_Geno,F5_Progeny_List,genoValSimList,phenoValSimList,varEList,noQTLs,H2,numberSelected,noCrosses,no_Progeny,noCycles,selectionOnGenoValues,selectionOnSimulatedValues,ModelRetrain,RetrainFrequency,ModelUpdate,UpdateType,UpdateFrequency,i,k,ModelType,weighted,weightingMethod,AlphaShape,BetaShape,TimeHorizon,trainGenoNewTable,trainSimGenoValTable,trainSimPhenoValTable,TrainTableWindowSize,NAM_LinkMap,gIndStats,alleleConversionTable_Combined) 
  }else if(modelType=="RRBLUP" | modelType=="RRBLUP_REML" | modelType=="PS"){
    simResults <- runSimulations40X_RRBLUP_WGS_V3(model_Sol_Pheno,model_Sol_Geno,F5_Progeny_List,genoValSimList,phenoValSimList,varEList,noQTLs,H2,numberSelected,noCrosses,no_Progeny,noCycles,selectionOnGenoValues,selectionOnSimulatedValues,ModelRetrain,RetrainFrequency,ModelUpdate,UpdateType,UpdateFrequency,i,k,ModelType,weighted,weightingMethod,AlphaShape,BetaShape,TimeHorizon,trainGenoNewTable,trainSimGenoValTable,trainSimPhenoValTable,TrainTableWindowSize,BreedingDesign,NAM_LinkMap,gIndStats,alleleConversionTable_Combined)
  } 
}
if(platForm == "Windows"){
  
  if(modelType=="SVMRBF"){
    simResults <- runSimulations40X_ML_Update(model_Sol_Pheno,model_Sol_Geno,F5_Progeny_List,genoValSimList,phenoValSimList,varEList,noQTLs,H2,numberSelected,noCrosses,no_Progeny,noCycles,selectionOnGenoValues,selectionOnSimulatedValues,ModelRetrain,RetrainFrequency,ModelUpdate,UpdateType,UpdateFrequency,i,k,ModelType,weighted,weightingMethod,AlphaShape,BetaShape,TimeHorizon,trainGenoNewTable,trainSimGenoValTable,trainSimPhenoValTable,TrainTableWindowSize,NAM_LinkMap,gIndStats,alleleConversionTable_Combined)
  }else if(modelType=="BayesB" | modelType=="BL"){
    simResults <- runSimulations40X_Bayes_WGS_V2_Win(model_Sol_Pheno,model_Sol_Geno,F5_Progeny_List,genoValSimList,phenoValSimList,varEList,noQTLs,H2,numberSelected,noCrosses,no_Progeny,noCycles,selectionOnGenoValues,selectionOnSimulatedValues,ModelRetrain,RetrainFrequency,ModelUpdate,UpdateType,UpdateFrequency,i,k,ModelType,weighted,weightingMethod,AlphaShape,BetaShape,TimeHorizon,trainGenoNewTable,trainSimGenoValTable,trainSimPhenoValTable,TrainTableWindowSize,NAM_LinkMap,gIndStats,alleleConversionTable_Combined) 
  }else if(modelType=="RRBLUP" | modelType=="RRBLUP_REML" | modelType=="PS"){
    simResults <- runSimulations40X_RRBLUP_WGS_V3_Win(model_Sol_Pheno,model_Sol_Geno,F5_Progeny_List,genoValSimList,phenoValSimList,varEList,noQTLs,H2,numberSelected,noCrosses,no_Progeny,noCycles,selectionOnGenoValues,selectionOnSimulatedValues,ModelRetrain,RetrainFrequency,ModelUpdate,UpdateType,UpdateFrequency,i,k,ModelType,weighted,weightingMethod,AlphaShape,BetaShape,TimeHorizon,trainGenoNewTable,trainSimGenoValTable,trainSimPhenoValTable,TrainTableWindowSize,BreedingDesign,NAM_LinkMap,gIndStats,alleleConversionTable_Combined)
  } 
} 
  
  
  return(simResults)
}
```
