## Supplementary material for "Factors Affecting Response to Recurrent Genomic Selection in Soybeans": File S1: trainGSModel.R

```
## function to train GS model given training and test geno and pheno set


trainGSModel <- function(NAMBasePopulationData, ModelType, nCores, MakeCluster,RegisterParallel){
  
 trainGenoNewTableListReps <- NAMBasePopulationData[[8]]
 trainSimGenoValuesTableListReps <- NAMBasePopulationData[[11]]
 trainSimPhenoValuesTableListReps <- NAMBasePopulationData[[12]]
 testGenoNewTableListReps <- NAMBasePopulationData[[10]]
 testSimGenoValuesTableListReps <- NAMBasePopulationData[[13]]
 testSimPhenoValuesTableListReps <- NAMBasePopulationData[[14]]
 
  
  modelType <- ModelType
  nreps <- length(trainGenoNewTableListReps)
  nCon <- length(trainGenoNewTableListReps[[1]])
  h2 <- c(0.7,0.3,0.7,0.3,0.7,0.3)
  no_Cores <- nCores
  
  ###
  
  if(MakeCluster ==TRUE && RegisterParallel==TRUE){
    cl <- makeCluster(no_Cores)
    registerDoParallel(cl)
  }else if(MakeCluster == FALSE && RegisterParallel==TRUE){
    registerDoParallel(no_Cores)
  }
  
  ####
  
  if(modelType == "BL"){
    
    
    model_Sol_Geno_List <- foreach(nConditions=1:nCon) %:% foreach(i=1:nreps, .export=c("trainGenoNewTableListReps","trainSimGenoValuesTableListReps","testGenoNewTableListReps","testSimGenoValuesTableListReps","buildBLModel")) %dopar% (buildBLModel(trainGenoNewTableListReps[[i]][[nConditions]],trainSimGenoValuesTableListReps[[i]][[nConditions]],testGenoNewTableListReps[[i]][[nConditions]],testSimGenoValuesTableListReps[[i]][[nConditions]],h2[nConditions]))
    
    
    model_Sol_Pheno_List <- foreach(nConditions=1:nCon) %:% foreach(i=1:nreps,.export=c("trainGenoNewTableListReps","trainSimPhenoValuesTableListReps","testGenoNewTableListReps","testSimPhenoValuesTableListReps","buildBLModel")) %dopar% (buildBLModel(trainGenoNewTableListReps[[i]][[nConditions]],trainSimPhenoValuesTableListReps[[i]][[nConditions]],testGenoNewTableListReps[[i]][[nConditions]],testSimPhenoValuesTableListReps[[i]][[nConditions]],h2[nConditions]))
    
  }
  
  if(modelType=="BayesB"){ 
    
    
    model_Sol_Geno_List <- foreach(nConditions=1:nCon) %:% foreach(i=1:nreps, .export=c("trainGenoNewTableListReps","trainSimGenoValuesTableListReps","testGenoNewTableListReps","testSimGenoValuesTableListReps","buildBayesBModel")) %dopar% (buildBayesBModel(trainGenoNewTableListReps[[i]][[nConditions]], trainSimGenoValuesTableListReps[[i]][[nConditions]],testGenoNewTableListReps[[i]][[nConditions]],testSimGenoValuesTableListReps[[i]][[nConditions]],h2[nConditions]))
    
    
    
    model_Sol_Pheno_List <- foreach(nConditions=1:nCon) %:% foreach(i=1:nreps,.export=c("trainGenoNewTableListReps","trainSimPhenoValuesTableListReps","testGenoNewTableListReps","testSimPhenoValuesTableListReps","buildBayesBModel")) %dopar% (buildBayesBModel(trainGenoNewTableListReps[[i]][[nConditions]],trainSimPhenoValuesTableListReps[[i]][[nConditions]],testGenoNewTableListReps[[i]][[nConditions]],testSimPhenoValuesTableListReps[[i]][[nConditions]],h2[nConditions]))
    
  }
  
  if(modelType=="RRBLUP"){ 
       
    
    
    model_Sol_Geno_List <- foreach(nConditions=1:nCon) %:% foreach(i=1:nreps, .export=c("trainGenoNewTableListReps","trainSimGenoValuesTableListReps","testGenoNewTableListReps","testSimGenoValuesTableListReps","buildRRBLUPModel"), .packages="rrBLUP") %dopar% (buildRRBLUPModel(trainGenoNewTableListReps[[i]][[nConditions]],trainSimGenoValuesTableListReps[[i]][[nConditions]][,1],testGenoNewTableListReps[[i]][[nConditions]],testSimGenoValuesTableListReps[[i]][[nConditions]][,1]))
    
    
    model_Sol_Pheno_List <- foreach(nConditions=1:nCon) %:% foreach(i=1:nreps, .export=c("trainGenoNewTableListReps","trainSimPhenoValuesTableListReps","testGenoNewTableListReps","testSimPhenoValuesTableListReps","buildRRBLUPModel"), .packages="rrBLUP") %dopar% (buildRRBLUPModel(trainGenoNewTableListReps[[i]][[nConditions]],trainSimPhenoValuesTableListReps[[i]][[nConditions]][,1],testGenoNewTableListReps[[i]][[nConditions]],testSimPhenoValuesTableListReps[[i]][[nConditions]][,1]))
    
  } 
  
  if(modelType=="RRBLUP_REML"){
    
    
    model_Sol_Geno_List <- foreach(nConditions=1:nCon) %:% foreach(i=1:nreps,.export=c("trainGenoNewTableListReps","trainSimGenoValuesTableListReps","testGenoNewTableListReps","testSimGenoValuesTableListReps","buildRRBLUPModel_REML")) %dopar% (buildRRBLUPModel_REML(trainGenoNewTableListReps[[i]][[nConditions]], trainSimGenoValuesTableListReps[[i]][[nConditions]][,1],testGenoNewTableListReps[[i]][[nConditions]],testSimGenoValuesTableListReps[[i]][[nConditions]][,1]))
    
    
    
    model_Sol_Pheno_List <- foreach(nConditions=1:nCon) %:% foreach(i=1:nreps,.export=c("trainGenoNewTableListReps","trainSimPhenoValuesTableListReps","testGenoNewTableListReps","testSimPhenoValuesTableListReps","buildRRBLUPModel_REML")) %dopar% (buildRRBLUPModel_REML(trainGenoNewTableListReps[[i]][[nConditions]],trainSimPhenoValuesTableListReps[[i]][[nConditions]][,1],testGenoNewTableListReps[[i]][[nConditions]],testSimPhenoValuesTableListReps[[i]][[nConditions]][,1]))
    
    
  }
  
  if(modelType=="SVMRBF"){
    
    
    model_Sol_Geno_List <- foreach(nConditions=1:nCon) %:% foreach(i=1:nreps, .export=c("trainGenoNewTableListReps","trainSimGenoValuesTableListReps","testGenoNewTableListReps","testSimGenoValuesTableListReps","buildBayesBModel")) %dopar% (buildSVMRBFModel(trainGenoNewTableListReps[[i]][[nConditions]], trainSimGenoValuesTableListReps[[i]][[nConditions]],testGenoNewTableListReps[[i]][[nConditions]],testSimGenoValuesTableListReps[[i]][[nConditions]]))
    
    #########################################################################################################################
    model_Sol_Pheno_List <- foreach(nConditions=1:nCon) %:% foreach(i=1:nreps,.export=c("trainGenoNewTableListReps","trainSimPhenoValuesTableListReps","testGenoNewTableListReps","testSimPhenoValuesTableListReps","buildSVMRBFModel")) %dopar% (buildSVMRBFModel(trainGenoNewTableListReps[[i]][[nConditions]],trainSimPhenoValuesTableListReps[[i]][[nConditions]],testGenoNewTableListReps[[i]][[nConditions]],testSimPhenoValuesTableListReps[[i]][[nConditions]]))
    
  }
  
  
  return(list(model_Sol_Geno_List,model_Sol_Pheno_List))
  
}


  
trainGSModel_Old <- function(trainGenoNewTableListReps,trainSimGenoValuesTableListReps,trainSimPhenoValuesTableListReps,testGenoNewTableListReps,testSimGenoValuesTableListReps,testSimPhenoValuesTableListReps,H2,ModelType,nCores,MakeCluster,RegisterParallel){

    modelType <- ModelType
	nreps <- length(trainGenoNewTableListReps)
	nCon <- length(trainGenoNewTableListReps[[1]])
	h2 <- H2
	no_Cores <- nCores
	
###

  if(MakeCluster ==TRUE && RegisterParallel==TRUE){
	cl <- makeCluster(no_Cores)
	registerDoParallel(cl)
  }else if(MakeCluster == FALSE && RegisterParallel==TRUE){
	registerDoParallel(no_Cores)
  }
  
####

    if(modelType == "BL"){
	
	  
		model_Sol_Geno_List <- foreach(nConditions=1:nCon) %:% foreach(i=1:nreps, .export=c("trainGenoNewTableListReps","trainSimGenoValuesTableListReps","testGenoNewTableListReps","testSimGenoValuesTableListReps","buildBLModel")) %dopar% (buildBLModel(trainGenoNewTableListReps[[i]][[nConditions]],trainSimGenoValuesTableListReps[[i]][[nConditions]],testGenoNewTableListReps[[i]][[nConditions]],testSimGenoValuesTableListReps[[i]][[nConditions]],h2[nConditions]))
     
		 
		model_Sol_Pheno_List <- foreach(nConditions=1:nCon) %:% foreach(i=1:nreps,.export=c("trainGenoNewTableListReps","trainSimPhenoValuesTableListReps","testGenoNewTableListReps","testSimPhenoValuesTableListReps","buildBLModel")) %dopar% (buildBLModel(trainGenoNewTableListReps[[i]][[nConditions]],trainSimPhenoValuesTableListReps[[i]][[nConditions]],testGenoNewTableListReps[[i]][[nConditions]],testSimPhenoValuesTableListReps[[i]][[nConditions]],h2[nConditions]))

	}

    if(modelType=="BayesB"){ 
	
  
			model_Sol_Geno_List <- foreach(nConditions=1:nCon) %:% foreach(i=1:nreps, .export=c("trainGenoNewTableListReps","trainSimGenoValuesTableListReps","testGenoNewTableListReps","testSimGenoValuesTableListReps","buildBayesBModel")) %dopar% (buildBayesBModel(trainGenoNewTableListReps[[i]][[nConditions]], trainSimGenoValuesTableListReps[[i]][[nConditions]],testGenoNewTableListReps[[i]][[nConditions]],testSimGenoValuesTableListReps[[i]][[nConditions]],h2[nConditions]))
     
	
		
			model_Sol_Pheno_List <- foreach(nConditions=1:nCon) %:% foreach(i=1:nreps,.export=c("trainGenoNewTableListReps","trainSimPhenoValuesTableListReps","testGenoNewTableListReps","testSimPhenoValuesTableListReps","buildBayesBModel")) %dopar% (buildBayesBModel(trainGenoNewTableListReps[[i]][[nConditions]],trainSimPhenoValuesTableListReps[[i]][[nConditions]],testGenoNewTableListReps[[i]][[nConditions]],testSimPhenoValuesTableListReps[[i]][[nConditions]],h2[nConditions]))

	}
	
	if(modelType=="RRBLUP"){ 
	
	

	
			model_Sol_Geno_List <- foreach(nConditions=1:nCon) %:% foreach(i=1:nreps, .export=c("trainGenoNewTableListReps","trainSimGenoValuesTableListReps","testGenoNewTableListReps","testSimGenoValuesTableListReps","buildRRBLUPModel"), .packages="rrBLUP") %dopar% (buildRRBLUPModel(trainGenoNewTableListReps[[i]][[nConditions]],trainSimGenoValuesTableListReps[[i]][[nConditions]],testGenoNewTableListReps[[i]][[nConditions]],testSimGenoValuesTableListReps[[i]][[nConditions]]))
     
				
			model_Sol_Pheno_List <- foreach(nConditions=1:nCon) %:% foreach(i=1:nreps, .export=c("trainGenoNewTableListReps","trainSimPhenoValuesTableListReps","testGenoNewTableListReps","testSimPhenoValuesTableListReps","buildRRBLUPModel"), .packages="rrBLUP") %dopar% (buildRRBLUPModel(trainGenoNewTableListReps[[i]][[nConditions]],trainSimPhenoValuesTableListReps[[i]][[nConditions]],testGenoNewTableListReps[[i]][[nConditions]],testSimPhenoValuesTableListReps[[i]][[nConditions]]))

	} 
	
	if(modelType=="RRBLUP_REML"){
	
	
	    model_Sol_Geno_List <- foreach(nConditions=1:nCon) %:% foreach(i=1:nreps, .export=c("trainGenoNewTableListReps","trainSimGenoValuesTableListReps","testGenoNewTableListReps","testSimGenoValuesTableListReps","buildRRBLUPModel_REML")) %dopar% (buildRRBLUPModel_REML(trainGenoNewTableListReps[[i]][[nConditions]], trainSimGenoValuesTableListReps[[i]][[nConditions]],testGenoNewTableListReps[[i]][[nConditions]],testSimGenoValuesTableListReps[[i]][[nConditions]]))
     
 

		model_Sol_Pheno_List <- foreach(nConditions=1:nCon) %:% foreach(i=1:nreps,.export=c("trainGenoNewTableListReps","trainSimPhenoValuesTableListReps","testGenoNewTableListReps","testSimPhenoValuesTableListReps","buildRRBLUPModel_REML")) %dopar% (buildRRBLUPModel_REML(trainGenoNewTableListReps[[i]][[nConditions]],trainSimPhenoValuesTableListReps[[i]][[nConditions]],testGenoNewTableListReps[[i]][[nConditions]],testSimPhenoValuesTableListReps[[i]][[nConditions]]))

		
	}

	if(modelType=="SVMRBF"){ 
 

		model_Sol_Geno_List <- foreach(nConditions=1:nCon) %:% foreach(i=1:nreps, .export=c("trainGenoNewTableListReps","trainSimGenoValuesTableListReps","testGenoNewTableListReps","testSimGenoValuesTableListReps","buildBayesBModel")) %dopar% (buildSVMRBFModel(trainGenoNewTableListReps[[i]][[nConditions]], trainSimGenoValuesTableListReps[[i]][[nConditions]],testGenoNewTableListReps[[i]][[nConditions]],testSimGenoValuesTableListReps[[i]][[nConditions]]))
    
#########################################################################################################################
		model_Sol_Pheno_List <- foreach(nConditions=1:nCon) %:% foreach(i=1:nreps,.export=c("trainGenoNewTableListReps","trainSimPhenoValuesTableListReps","testGenoNewTableListReps","testSimPhenoValuesTableListReps","buildSVMRBFModel")) %dopar% (buildSVMRBFModel(trainGenoNewTableListReps[[i]][[nConditions]],trainSimPhenoValuesTableListReps[[i]][[nConditions]],testGenoNewTableListReps[[i]][[nConditions]],testSimPhenoValuesTableListReps[[i]][[nConditions]]))

	}


    return(list(model_Sol_Geno_List,model_Sol_Pheno_List))
	
 }
	
	
```
