## Supplementary material for "Factors Affecting Response to Recurrent Genomic Selection in Soybeans": File S1: trainGSModel_w_o_QTL.R

```
# library(SoyNAMPredictionMethods)
#library(SoyNAMPredictionMethodsGPU)

## function to train GS model given training and test geno and pheno set

# QTL_in_Model <- c(TRUE,FALSE)
# Model_Type_List <- c("BayesB","BL","RRBLUP_REML")

# Model_Sol_List_Q <- list() 

# ModelType <- Model_Type_List[1]
# nCores <- 10 
# MakeCluster <- TRUE 
# RegisterParallel <- TRUE

# for(QM in QTL_in_Model){


# # trainGSModel <- function(NAMBasePopulationData, ModelType, nCores, MakeCluster,RegisterParallel){

  
  # if(QM==TRUE){
  
	 # trainGenoNewTableListReps <- NAMBasePopulationData[[8]]
	 # testGenoNewTableListReps <- NAMBasePopulationData[[11]]
	 
  # } 
  
  # if(QM==FALSE){
	
	# trainGenoNewTableListReps <- NAMBasePopulationData[[9]]
 	# testGenoNewTableListReps <- NAMBasePopulationData[[12]]
	
   # }
 
 # trainSimGenoValuesTableListReps <- NAMBasePopulationData[[13]]
 # trainSimPhenoValuesTableListReps <- NAMBasePopulationData[[14]]
 
 # testSimGenoValuesTableListReps <- NAMBasePopulationData[[15]]
 # testSimPhenoValuesTableListReps <- NAMBasePopulationData[[16]]
 
  
  # modelType <- ModelType
  # nreps <- length(trainGenoNewTableListReps)
  # nCon <- length(trainGenoNewTableListReps[[1]])
  # h2 <- c(0.7,0.3,0.7,0.3,0.7,0.3)
  # no_Cores <- nCores
  
  # ###
  
  # if(MakeCluster ==TRUE && RegisterParallel==TRUE){
    # cl <- makeCluster(no_Cores)
    # registerDoParallel(cl)
  # }else if(MakeCluster == FALSE && RegisterParallel==TRUE){
    # registerDoParallel(no_Cores)
  # }
  
  # ####
  
  # if(modelType == "BL"){
    
    
    # model_Sol_Geno_List <- foreach(nConditions=1:nCon,.packages="SoyNAMPredictionMethods") %:% foreach(nRep=1:nreps, .export=c("trainGenoNewTableListReps","trainSimGenoValuesTableListReps","testGenoNewTableListReps","testSimGenoValuesTableListReps","buildBLModel")) %dopar% (SoyNAMPredictionMethods::buildBLModel(trainGenoNewTableListReps[[nRep]][[nConditions]],trainSimGenoValuesTableListReps[[nRep]][[nConditions]][,1],testGenoNewTableListReps[[nRep]][[nConditions]],testSimGenoValuesTableListReps[[nRep]][[nConditions]][,1],h2[nConditions]))
    
    
    # model_Sol_Pheno_List <- foreach(nConditions=1:nCon,.packages="SoyNAMPredictionMethods") %:% foreach(nRep=1:nreps,.export=c("trainGenoNewTableListReps","trainSimPhenoValuesTableListReps","testGenoNewTableListReps","testSimPhenoValuesTableListReps","buildBLModel")) %dopar% (SoyNAMPredictionMethods::buildBLModel(trainGenoNewTableListReps[[nRep]][[nConditions]],trainSimPhenoValuesTableListReps[[nRep]][[nConditions]][,1],testGenoNewTableListReps[[nRep]][[nConditions]],testSimPhenoValuesTableListReps[[nRep]][[nConditions]][,1],h2[nConditions]))
    
  # }
  
  # if(modelType=="BayesB"){ 
    
    
    # model_Sol_Geno_List <- foreach(nConditions=1:nCon,.packages="SoyNAMPredictionMethods") %:% foreach(nRep=1:nreps, .export=c("trainGenoNewTableListReps","trainSimGenoValuesTableListReps","testGenoNewTableListReps","testSimGenoValuesTableListReps","buildBayesBModel")) %dopar% (SoyNAMPredictionMethods::buildBayesBModel(trainGenoNewTableListReps[[nRep]][[nConditions]], trainSimGenoValuesTableListReps[[nRep]][[nConditions]][,1],testGenoNewTableListReps[[nRep]][[nConditions]],testSimGenoValuesTableListReps[[nRep]][[nConditions]][,1],h2[nConditions]))
    
    
    
    # model_Sol_Pheno_List <- foreach(nConditions=1:nCon,packages="SoyNAMPredictionMethods") %:% foreach(nRep=1:nreps,.export=c("trainGenoNewTableListReps","trainSimPhenoValuesTableListReps","testGenoNewTableListReps","testSimPhenoValuesTableListReps","buildBayesBModel")) %dopar% (SoyNAMPredictionMethods::buildBayesBModel(trainGenoNewTableListReps[[nRep]][[nConditions]],trainSimPhenoValuesTableListReps[[nRep]][[nConditions]][,1],testGenoNewTableListReps[[nRep]][[nConditions]],testSimPhenoValuesTableListReps[[nRep]][[nConditions]][,1],h2[nConditions]))
    
  # }
  
  # if(modelType=="RRBLUP"){ 
       
    
    
    # model_Sol_Geno_List <- foreach(nConditions=1:nCon) %:% foreach(nRep=1:nreps, .export=c("trainGenoNewTableListReps","trainSimGenoValuesTableListReps","testGenoNewTableListReps","testSimGenoValuesTableListReps","buildRRBLUPModel"), .packages="rrBLUP") %dopar% (SoyNAMPredictionMethods::buildRRBLUPModel(trainGenoNewTableListReps[[nRep]][[nConditions]],trainSimGenoValuesTableListReps[[nRep]][[nConditions]][,1],testGenoNewTableListReps[[nRep]][[nConditions]],testSimGenoValuesTableListReps[[nRep]][[nConditions]][,1]))
    
    
    # model_Sol_Pheno_List <- foreach(nConditions=1:nCon) %:% foreach(nRep=1:nreps, .export=c("trainGenoNewTableListReps","trainSimPhenoValuesTableListReps","testGenoNewTableListReps","testSimPhenoValuesTableListReps","buildRRBLUPModel"), .packages="rrBLUP") %dopar% (SoyNAMPredictionMethods::buildRRBLUPModel(trainGenoNewTableListReps[[nRep]][[nConditions]],trainSimPhenoValuesTableListReps[[nRep]][[nConditions]][,1],testGenoNewTableListReps[[nRep]][[nConditions]],testSimPhenoValuesTableListReps[[nRep]][[nConditions]][,1]))
    
  # } 
  
  # if(modelType=="RRBLUP_REML"){
    
    
    # model_Sol_Geno_List <- foreach(nConditions=1:nCon,.packages="SoyNAMPredictionMethods") %:% foreach(nRep=1:nreps,.export=c("trainGenoNewTableListReps","trainSimGenoValuesTableListReps","testGenoNewTableListReps","testSimGenoValuesTableListReps","buildRRBLUPModel_REML")) %dopar% (SoyNAMPredictionMethods::buildRRBLUPModel_REML(trainGenoNewTableListReps[[nRep]][[nConditions]], trainSimGenoValuesTableListReps[[nRep]][[nConditions]][,1],testGenoNewTableListReps[[nRep]][[nConditions]],testSimGenoValuesTableListReps[[nRep]][[nConditions]][,1]))
    
    
    
    # model_Sol_Pheno_List <- foreach(nConditions=1:nCon,.packages="SoyNAMPredictionMethods") %:% foreach(nRep=1:nreps,.export=c("trainGenoNewTableListReps","trainSimPhenoValuesTableListReps","testGenoNewTableListReps","testSimPhenoValuesTableListReps","buildRRBLUPModel_REML")) %dopar% (SoyNAMPredictionMethods::buildRRBLUPModel_REML(trainGenoNewTableListReps[[nRep]][[nConditions]],trainSimPhenoValuesTableListReps[[nRep]][[nConditions]][,1],testGenoNewTableListReps[[nRep]][[nConditions]],testSimPhenoValuesTableListReps[[nRep]][[nConditions]][,1]))
    
    
  # }
  
  # if(modelType=="SVMRBF"){
    
    
    # model_Sol_Geno_List <- foreach(nConditions=1:nCon) %:% foreach(nRep=1:nreps, .export=c("trainGenoNewTableListReps","trainSimGenoValuesTableListReps","testGenoNewTableListReps","testSimGenoValuesTableListReps","buildBayesBModel")) %dopar% (buildSVMRBFModel(trainGenoNewTableListReps[[nRep]][[nConditions]], trainSimGenoValuesTableListReps[[nRep]][[nConditions]],testGenoNewTableListReps[[nRep]][[nConditions]],testSimGenoValuesTableListReps[[nRep]][[nConditions]]))
    
    # #########################################################################################################################
    # model_Sol_Pheno_List <- foreach(nConditions=1:nCon) %:% foreach(nRep=1:nreps,.export=c("trainGenoNewTableListReps","trainSimPhenoValuesTableListReps","testGenoNewTableListReps","testSimPhenoValuesTableListReps","buildSVMRBFModel")) %dopar% (buildSVMRBFModel(trainGenoNewTableListReps[[nRep]][[nConditions]],trainSimPhenoValuesTableListReps[[nRep]][[nConditions]],testGenoNewTableListReps[[nRep]][[nConditions]],testSimPhenoValuesTableListReps[[nRep]][[nConditions]]))
    
  # }
  
 
 
# buildSVMRBFModel_GPU <- function(trainGenoTable,trainPhenoTable,testGenoTable,testPhenoTable){
  
  # nIndividuals <- 2000
  # trainPheno <- trainPhenoTable
  # #trainGeno <- trainGenoTable
  
  # folds=5
   
  # model1<- (svm(x=trainGenoTable,y=trainPheno,kernel="radial",type="eps-regression",cross=folds,probability=FALSE,verbose=FALSE,no.change.x=FALSE,scale=FALSE))
  # #myControl <- trainControl(method='cv', number=folds, repeats=repeats, 
  # #                       returnResamp='none', classProbs=FALSE,
  # #                        returnData=FALSE, savePredictions=TRUE, 
  # #                        verboseIter=TRUE, allowParallel=TRUE,
  # #                        index=createMultiFolds(trainPheno, k=folds, times=repeats)) 
						  
  # # gridRF <- expand.grid(mfinal=c(50,100,150,175,200))
  # # model1<- train(trainGeno,trainPheno, method='rf',trControl=myControl,tuneGrid=gridRF)	
# ## Predict with test data set ###########################################################################s
  # testPheno <- testPhenoTable
  # testGeno <- testGenoTable 
 
# ### Predictions with probability values
# #preds_prob <- data.frame(sapply(all.models.2, function(x){predict(x, testGeno,type='prob')}))
 
  # pred.pheno.train <- (predict(model1,trainGenoTable))
  # pred.pheno.valid <- (predict(model1,testGeno))

# # cor(Pred.pheno.train,trainPheno)
   # u.effects <- (model1$coefs)
# #################################################################################################################
      
  # Pred.Accuracy <- cor(pred.pheno.valid,testPheno)
  
  # train_MSE <- (sum((pred.pheno.train-trainPheno))^2)/(0.8*nIndividuals)
  # test_MSE <- (sum((pred.pheno.valid-testPheno))^2)/(0.2*nIndividuals) 
  
  # sol <- list(Pred.Accuracy,model1,u.effects,train_MSE,test_MSE)
  
  # return(sol)
  
# } 
 
  
   # if(modelType=="SVMRBFGPU"){
   
    
  
      # model_Sol_Geno_List <- foreach(nConditions=1:nCon) %:% foreach(nRep=1:nreps,.export=c("trainGenoNewTableListReps","trainSimPhenoValuesTableListReps","testGenoNewTableListReps","testSimPhenoValuesTableListReps","buildSVMRBFModel")) %dopar% (tryCatch(SoyNAMPredictionMethodsGPU::buildSVMRBFModel_GPU(trainGenoNewTableListReps[[nRep]][[nConditions]],trainSimGenoValuesTableListReps[[nRep]][[nConditions]][,1],testGenoNewTableListReps[[nRep]][[nConditions]],testSimPhenoValuesTableListReps[[nRep]][[nConditions]][,1]),error=function(e) print(paste("BuildSVMRBF-Error -",e))))
	 
	
			
	  # model_Sol_Pheno_List <- foreach(nConditions=1:nCon) %:% foreach(nRep=1:nreps,.export=c("trainGenoNewTableListReps","trainSimPhenoValuesTableListReps","testGenoNewTableListReps","testSimPhenoValuesTableListReps","buildSVMRBFModel")) %dopar% (tryCatch(SoyNAMPredictionMethodsGPU::buildSVMRBFModel_GPU(trainGenoNewTableListReps[[nRep]][[nConditions]],trainSimPhenoValuesTableListReps[[nRep]][[nConditions]][,1],testGenoNewTableListReps[[nRep]][[nConditions]],testSimPhenoValuesTableListReps[[nRep]][[nConditions]][,1]),error=function(e) print(paste("BuildSVMRBF-Error -",e))))
			
			
  
    # }
  
  
  
  
 # # return
 
  # Model_Sol_List <- (list(model_Sol_Geno_List,model_Sol_Pheno_List)) 
  
  # Model_Sol_List_Q[[QM]] <- Model_Sol_List
# }


###
# 
# for(nConditions in 1:nCon){
# 
#        model_Sol_Geno_List[[nConditions]] <- foreach(nRep=1:nreps,.export=c("trainGenoNewTableListReps","trainSimGenoValuesTableListReps","testGenoNewTableListReps","testSimGenoValuesTableListReps","buildRRBLUPModel_REML")) %dopar% (SoyNAMPredictionMethods::buildRRBLUPModel_REML(trainGenoNewTableListReps[[nRep]][[nConditions]], trainSimGenoValuesTableListReps[[nRep]][[nConditions]][,1],testGenoNewTableListReps[[nRep]][[nConditions]],testSimGenoValuesTableListReps[[nRep]][[nConditions]][,1]))
# 
# }


# 
```
