## Supplementary material for "Factors Affecting Response to Recurrent Genomic Selection in Soybeans": File S1: trainGSModel_within_Families.R

```
# ## function to train GS model given training and test geno and pheno set

# #load("BasePopulation_Accuracy_Families.RData")

# # model_Sol_Geno_List <- list()
# # model_Sol_Pheno_List <- list()

# # NAMBasePopulationData <- NAMBasePopulationData_List
# # ModelType <- "RRBLUP_REML"
# # nCores <- 10
# # MakeCluster <- TRUE
# # RegisterParallel <- TRUE


# trainGSModel_within_Families <- function(NAMBasePopulationData, ModelType, nCores, MakeCluster,RegisterParallel){
  
 # trainGenoNewTableListReps <- NAMBasePopulationData[[8]]
 # trainSimGenoValuesTableListReps <- NAMBasePopulationData[[11]]
 # trainSimPhenoValuesTableListReps <- NAMBasePopulationData[[12]]
 # testGenoNewTableListReps <- NAMBasePopulationData[[10]]
 # testSimGenoValuesTableListReps <- NAMBasePopulationData[[13]]
 # testSimPhenoValuesTableListReps <- NAMBasePopulationData[[14]]
 
  
  # modelType <- ModelType
  # nreps <- length(trainGenoNewTableListReps)
  # nCon <- length(trainGenoNewTableListReps[[1]])
  # h2 <- c(0.7,0.3,0.7,0.3,0.7,0.3)
  # no_Cores <- nCores
  
  # ###
  
  # if(MakeCluster ==TRUE && RegisterParallel==TRUE){
    # cl <- makeCluster(no_Cores)
    # registerDoParallel(cl)
  # }else if(MakeCluster == FALSE && RegisterParallel==TRUE){
    # registerDoParallel(no_Cores)
  # }
  
  # ####
  
  # if(modelType == "BL"){
    
    
    # model_Sol_Geno_List <- foreach(i=1:nreps,.export=c("trainGenoNewTableListReps","trainSimGenoValuesTableListReps","testGenoNewTableListReps","testSimGenoValuesTableListReps","buildBayesBModel")) %:% foreach(k=1:nFamilies) %dopar% 
	# (buildBLModel(trainGenoNewTableListReps[[i]][[nConditions]][[k]], trainSimGenoValuesTableListReps[[i]][[nConditions]][[k]][,1],testGenoNewTableListReps[[i]][[nConditions]][[k]],testSimGenoValuesTableListReps[[i]][[nConditions]][[k]][,1],h2[nConditions]))
    
    
    # model_Sol_Pheno_List <- foreach(i=1:nreps,.export=c("trainGenoNewTableListReps","trainSimGenoValuesTableListReps","testGenoNewTableListReps","testSimGenoValuesTableListReps","buildBayesBModel")) %:% foreach(k=1:nFamilies) %dopar% 
	# (buildBLModel(trainGenoNewTableListReps[[i]][[nConditions]][[k]],trainSimPhenoValuesTableListReps[[i]][[nConditions]][[k]][,1],testGenoNewTableListReps[[i]][[nConditions]][[k]],testSimPhenoValuesTableListReps[[i]][[nConditions]][[k]][,1],h2[nConditions]))
    
  # }
  
  # if(modelType=="BayesB"){ 
    
	# for(nConditions in 1:6){
	
    
		# model_Sol_Geno_List <- foreach(i=1:nreps,.export=c("trainGenoNewTableListReps","trainSimGenoValuesTableListReps","testGenoNewTableListReps","testSimGenoValuesTableListReps","buildBayesBModel")) %:% foreach(k=1:nFamilies) %dopar%
		# (buildBayesBModel(trainGenoNewTableListReps[[i]][[nConditions]][[k]], trainSimGenoValuesTableListReps[[i]][[nConditions]][[k]][,1],testGenoNewTableListReps[[i]][[nConditions]][[k]],testSimGenoValuesTableListReps[[i]][[nConditions]][[k]][,1],h2[nConditions]))
		
		  
		# model_Sol_Pheno_List <- foreach(i=1:nreps,.export=c("trainGenoNewTableListReps","trainSimGenoValuesTableListReps","testGenoNewTableListReps","testSimGenoValuesTableListReps","buildBayesBModel")) %:% foreach(k=1:nFamilies) %dopar% (buildBayesBModel(trainGenoNewTableListReps[[i]][[nConditions]][[k]],trainSimPhenoValuesTableListReps[[i]][[nConditions]][[k]][,1],testGenoNewTableListReps[[i]][[nConditions]][[k]],testSimPhenoValuesTableListReps[[i]][[nConditions]][[k]][,1],h2[nConditions]))
    
    # }
	
  # }
  
  # if(modelType=="RRBLUP"){ 
       
    
    
    # model_Sol_Geno_List <- foreach(nConditions=1:nCon) %:% foreach(i=1:nreps, .export=c("trainGenoNewTableListReps","trainSimGenoValuesTableListReps","testGenoNewTableListReps","testSimGenoValuesTableListReps","buildRRBLUPModel"), .packages="rrBLUP") %dopar% (buildRRBLUPModel(trainGenoNewTableListReps[[i]][[nConditions]],trainSimGenoValuesTableListReps[[i]][[nConditions]][,1],testGenoNewTableListReps[[i]][[nConditions]],testSimGenoValuesTableListReps[[i]][[nConditions]][,1]))
    
    
    # model_Sol_Pheno_List <- foreach(nConditions=1:nCon) %:% foreach(i=1:nreps, .export=c("trainGenoNewTableListReps","trainSimPhenoValuesTableListReps","testGenoNewTableListReps","testSimPhenoValuesTableListReps","buildRRBLUPModel"), .packages="rrBLUP") %dopar% (buildRRBLUPModel(trainGenoNewTableListReps[[i]][[nConditions]],trainSimPhenoValuesTableListReps[[i]][[nConditions]][,1],testGenoNewTableListReps[[i]][[nConditions]],testSimPhenoValuesTableListReps[[i]][[nConditions]][,1]))
    
  # } 
  
  # if(modelType=="RRBLUP_REML"){
    
  
  
	# # model_Sol_Geno_List <- foreach(nConditions=1:nCon,.export=c("trainGenoNewTableListReps","trainSimGenoValuesTableListReps","testGenoNewTableListReps","testSimGenoValuesTableListReps","buildRRBLUPModel_REML")) %:% 
	
	# for(nConditions in 1:6){
	
	   # model_Sol_Geno_List[[nConditions]] <- foreach(i=1:nreps,.export=c("trainGenoNewTableListReps","trainSimGenoValuesTableListReps","testGenoNewTableListReps","testSimGenoValuesTableListReps","buildRRBLUPModel_REML")) %:% foreach(k=1:nFamilies) %dopar% (SoyNAMPredictionMethods::buildRRBLUPModel_REML(trainGenoNewTableListReps[[i]][[nConditions]][[k]], trainSimGenoValuesTableListReps[[i]][[nConditions]][[k]][,1],testGenoNewTableListReps[[i]][[nConditions]][[k]],testSimGenoValuesTableListReps[[i]][[nConditions]][[k]][,1]))
    
    
    
	    # model_Sol_Pheno_List[[nConditions]] <- foreach(i=1:nreps,.export=c("trainGenoNewTableListReps","trainSimPhenoValuesTableListReps","testGenoNewTableListReps","testSimPhenoValuesTableListReps","buildRRBLUPModel_REML")) %:% foreach(k=1:nFamilies) %dopar% (SoyNAMPredictionMethods::buildRRBLUPModel_REML(trainGenoNewTableListReps[[i]][[nConditions]][[k]],trainSimPhenoValuesTableListReps[[i]][[nConditions]][[k]][,1],testGenoNewTableListReps[[i]][[nConditions]][[k]],testSimPhenoValuesTableListReps[[i]][[nConditions]][[k]][,1]))
    
    
    # }
  
  # }
  
  # if(modelType=="SVMRBF"){
    
    
    # model_Sol_Geno_List <- foreach(nConditions=1:nCon) %:% foreach(i=1:nreps, .export=c("trainGenoNewTableListReps","trainSimGenoValuesTableListReps","testGenoNewTableListReps","testSimGenoValuesTableListReps","buildBayesBModel")) %dopar% (buildSVMRBFModel(trainGenoNewTableListReps[[i]][[nConditions]], trainSimGenoValuesTableListReps[[i]][[nConditions]],testGenoNewTableListReps[[i]][[nConditions]],testSimGenoValuesTableListReps[[i]][[nConditions]]))
    
    # #########################################################################################################################
    # model_Sol_Pheno_List <- foreach(nConditions=1:nCon) %:% foreach(i=1:nreps,.export=c("trainGenoNewTableListReps","trainSimPhenoValuesTableListReps","testGenoNewTableListReps","testSimPhenoValuesTableListReps","buildSVMRBFModel")) %dopar% (buildSVMRBFModel(trainGenoNewTableListReps[[i]][[nConditions]],trainSimPhenoValuesTableListReps[[i]][[nConditions]],testGenoNewTableListReps[[i]][[nConditions]],testSimPhenoValuesTableListReps[[i]][[nConditions]]))
    
  # }
  
  
  
  
  # return(list(model_Sol_Geno_List,model_Sol_Pheno_List))
  
# }

# # nFamilies <- 20 
# # nReps <- 10
# # nConditions <- 6

# # PredAccuracy <- rep(list(rep(list(rep(list(list()),nFamilies)),nReps)),nConditions)


# # for(nCon in 1:6){

  # # for(nrep in 1:10){ 
  
    # # for(nFamily in 1:20){
		
		
		# # PredAccuracy[[nCon]][[nrep]][[nFamily]] <- (model_Sol_Pheno_List[[nCon]][[nrep]][[nFamily]][[3]])
		
	# # }
 
  # # }
# # }

# # nCon <- 3
# # PredAccuracy_Condition <- list()

# # for(nCon in 1:6){

 # # PredAccuracy_Table <- matrix(rep(0,10*20),nrow=10,ncol=20)

 # # for(nrep in 1:10){
  
  # # for(nFamily in 1:20){ 

   # # PredAccuracy_Table[nrep,nFamily] <- PredAccuracy[[nCon]][[nrep]][[nFamily]]

  # # }
 # # } 
 
   # # PredAccuracy_Condition[[nCon]] <- apply(PredAccuracy_Table,2,mean) 
  
 # # } 
 
 
 
 
 # # ## Complete population with same population size as within family training
 
 # # trainGenoNewTableListReps <- NAMBasePopulationData[[8]]
 # # trainSimGenoValuesTableListReps <- NAMBasePopulationData[[11]]
 # # trainSimPhenoValuesTableListReps <- NAMBasePopulationData[[12]]
 # # testGenoNewTableListReps <- NAMBasePopulationData[[10]]
 # # testSimGenoValuesTableListReps <- NAMBasePopulationData[[13]]
 # # testSimPhenoValuesTableListReps <- NAMBasePopulationData[[14]]
 
 # # trainGenoNewTableListReps_Combined <- rep(list(rep(list(list()),nConditions)),nReps)
 # # trainSimGenoValuesTableListReps_Combined <- rep(list(rep(list(list()),nConditions)),nReps)
 # # trainSimPhenoValuesTableListReps_Combined <- rep(list(rep(list(list()),nConditions)),nReps)
 # # testGenoNewTableListReps_Combined <- rep(list(rep(list(list()),nConditions)),nReps)
 # # testSimGenoValuesTableListReps_Combined <- rep(list(rep(list(list()),nConditions)),nReps)
 # # testSimPhenoValuesTableListReps_Combined <- rep(list(rep(list(list()),nConditions)),nReps)
 
# # for(nCon in 1:6){

  # # for(nrep in 1:10){ 
    # # trainGenoNewTableReps_Combined <- c()
	# # trainSimGenoValuesTableReps_Combined <- c() 
	# # trainSimPhenoValuesTableReps_Combined <- c() 
	# # testGenoNewTableReps_Combined <- c() 
	# # testSimGenoValuesTableReps_Combined <- c()
	# # testSimPhenoValuesTableReps_Combined <- c()
	
 
	# # for(nFamily in 1:nFamilies){ 
  
    # # trainGenoNewTableReps_Combined <- rbind(trainGenoNewTableReps_Combined,trainGenoNewTableListReps[[nrep]][[nCon]][[nFamily]])
	# # trainGenoNewTableListReps_Combined[[nrep]][[nCon]] <- trainGenoNewTableReps_Combined 
	    
	# # trainSimGenoValuesTableReps_Combined <- rbind(trainSimGenoValuesTableReps_Combined,trainSimGenoValuesTableListReps[[nrep]][[nCon]][[nFamily]])
	# # trainSimGenoValuesTableListReps_Combined[[nrep]][[nCon]] <- trainSimGenoValuesTableReps_Combined
	
	# # trainSimPhenoValuesTableReps_Combined <- rbind(trainSimPhenoValuesTableReps_Combined,trainSimPhenoValuesTableListReps[[nrep]][[nCon]][[nFamily]])
	# # trainSimPhenoValuesTableListReps_Combined[[nrep]][[nCon]] <- trainSimPhenoValuesTableReps_Combined
	
	# # testGenoNewTableReps_Combined <- rbind(testGenoNewTableReps_Combined,testGenoNewTableListReps[[nrep]][[nCon]][[nFamily]])
	# # testGenoNewTableListReps_Combined[[nrep]][[nCon]] <- testGenoNewTableReps_Combined 
	
	
	# # testSimGenoValuesTableReps_Combined <-  rbind(testSimGenoValuesTableReps_Combined,testSimGenoValuesTableListReps[[nrep]][[nCon]][[nFamily]])
    # # testSimGenoValuesTableListReps_Combined[[nrep]][[nCon]] <- 	testSimGenoValuesTableReps_Combined
	
	
	# # testSimPhenoValuesTableReps_Combined <- rbind(testSimPhenoValuesTableReps_Combined,testSimPhenoValuesTableListReps[[nrep]][[nCon]][[nFamily]])
	# # testSimPhenoValuesTableListReps_Combined[[nrep]][[nCon]] <-  testSimPhenoValuesTableReps_Combined 
	
	# # } 
	
   # # }
   
  
  # # }
 
 
 
 
 # # trainGenoNewTableListReps_Combined_Sampled <- rep(list(rep(list(list()),nConditions)),nReps)
 # # trainSimGenoValuesTableListReps_Combined_Sampled <- rep(list(rep(list(list()),nConditions)),nReps)
 # # trainSimPhenoValuesTableListReps_Combined_Sampled <- rep(list(rep(list(list()),nConditions)),nReps)
 # # testGenoNewTableListReps_Combined_Sampled <- rep(list(rep(list(list()),nConditions)),nReps)
 # # testSimGenoValuesTableListReps_Combined_Sampled <- rep(list(rep(list(list()),nConditions)),nReps)
 # # testSimPhenoValuesTableListReps_Combined_Sampled <- rep(list(rep(list(list()),nConditions)),nReps)
 
# # for(nCon in 1:6){

  # # for(nrep in 1:10){ 
  
        
		# # nLines <- nrow(trainGenoNewTableListReps_Combined[[nrep]][[nCon]])
		# # nLines_per_Family <- 100
		# # initseed <- 25+nrep 
		# # set.seed(initseed)
		# # indices<-c(1:nLines)
		# # trainIndices <-sample(c(1:nLines),(0.8*nLines_per_Family)) 
		# # nLines_Test <- nrow(testGenoNewTableListReps_Combined[[nrep]][[nCon]])
        # # indices<-c(1:nLines_Test)
		# # testIndices <-sample(c(1:nLines_Test),(0.2*nLines_per_Family)) 
 
    
		# # trainGenoNewTableListReps_Combined_Sampled[[nrep]][[nCon]]  <- trainGenoNewTableListReps_Combined[[nrep]][[nCon]][trainIndices,]
	    
	
		# # trainSimGenoValuesTableListReps_Combined_Sampled[[nrep]][[nCon]]  <- trainSimGenoValuesTableListReps_Combined[[nrep]][[nCon]][trainIndices,]
	
	
		# # trainSimPhenoValuesTableListReps_Combined_Sampled[[nrep]][[nCon]] <- trainSimPhenoValuesTableListReps_Combined[[nrep]][[nCon]][trainIndices,]
	
	
		# # testGenoNewTableListReps_Combined_Sampled[[nrep]][[nCon]]  <- testGenoNewTableListReps_Combined[[nrep]][[nCon]][testIndices,]
	

		# # testSimGenoValuesTableListReps_Combined_Sampled[[nrep]][[nCon]] <- 	testSimGenoValuesTableListReps_Combined[[nrep]][[nCon]][testIndices,]
	
	
	
		# # testSimPhenoValuesTableListReps_Combined_Sampled[[nrep]][[nCon]] <- testSimPhenoValuesTableListReps_Combined[[nrep]][[nCon]][testIndices,]
	
    # # }
 # # } 
  
 
 
 
  
  # # modelType <- ModelType
  # # nreps <- length(trainGenoNewTableListReps)
  # # nCon <- length(trainGenoNewTableListReps[[1]])
  # # h2 <- c(0.7,0.3,0.7,0.3,0.7,0.3)
  # # no_Cores <- nCores
  
 
 
 
 
 
  # # ####
  
  # # if(modelType == "BL"){
    
    
    # # model_Sol_Geno_List_Combined_Sampled[[nConditions]] <-	foreach(i=1:nreps,.export=c("trainGenoNewTableListReps_Combined_Sampled","trainSimGenoValuesTableListReps_Combined_Sampled","testGenoNewTableListReps_Combined_Sampled","testSimGenoValuesTableListReps_Combined_Sampled","buildRRBLUPModel_REML")) %dopar% (SoyNAMPredictionMethods::buildBLModel(trainGenoNewTableListReps_Combined_Sampled[[i]][[nConditions]], trainSimGenoValuesTableListReps_Combined_Sampled[[i]][[nConditions]][,1],testGenoNewTableListReps_Combined_Sampled[[i]][[nConditions]],testSimGenoValuesTableListReps_Combined_Sampled[[i]][[nConditions]][,1],h2[nConditions]))
    
    
    
	# # model_Sol_Pheno_List_Combined_Sampled[[nConditions]] <- foreach(i=1:nreps,.export=c("trainGenoNewTableListReps_Combined_Sampled","trainSimPhenoValuesTableListReps_Combined_Sampled","testGenoNewTableListReps_Combined_Sampled","testSimPhenoValuesTableListReps_Combined_Sampled","buildRRBLUPModel_REML")) %dopar% (SoyNAMPredictionMethods::buildBLModel(trainGenoNewTableListReps_Combined_Sampled[[i]][[nConditions]],trainSimPhenoValuesTableListReps_Combined_Sampled[[i]][[nConditions]][,1],testGenoNewTableListReps_Combined_Sampled[[i]][[nConditions]],testSimPhenoValuesTableListReps_Combined_Sampled[[i]][[nConditions]][,1],h2[nConditions]))
    
  # # }
  
  # # if(modelType=="BayesB"){ 
    
	# # for(nConditions in 1:6){
	
    	
	
	
	    # # model_Sol_Geno_List_Combined_Sampled[[nConditions]] <-	foreach(i=1:nreps,.export=c("trainGenoNewTableListReps_Combined_Sampled","trainSimGenoValuesTableListReps_Combined_Sampled","testGenoNewTableListReps_Combined_Sampled","testSimGenoValuesTableListReps_Combined_Sampled","buildRRBLUPModel_REML")) %dopar% (SoyNAMPredictionMethods::buildBayesBModel(trainGenoNewTableListReps_Combined_Sampled[[i]][[nConditions]], trainSimGenoValuesTableListReps_Combined_Sampled[[i]][[nConditions]][,1],testGenoNewTableListReps_Combined_Sampled[[i]][[nConditions]],testSimGenoValuesTableListReps_Combined_Sampled[[i]][[nConditions]][,1],h2[nConditions]))
    
    
    
		 # # model_Sol_Pheno_List_Combined_Sampled[[nConditions]] <- foreach(i=1:nreps,.export=c("trainGenoNewTableListReps_Combined_Sampled","trainSimPhenoValuesTableListReps_Combined_Sampled","testGenoNewTableListReps_Combined_Sampled","testSimPhenoValuesTableListReps_Combined_Sampled","buildRRBLUPModel_REML")) %dopar% (SoyNAMPredictionMethods::buildBayesBModel(trainGenoNewTableListReps_Combined_Sampled[[i]][[nConditions]],trainSimPhenoValuesTableListReps_Combined_Sampled[[i]][[nConditions]][,1],testGenoNewTableListReps_Combined_Sampled[[i]][[nConditions]],testSimPhenoValuesTableListReps_Combined_Sampled[[i]][[nConditions]][,1],h2[nConditions]))
    
    
      # # }
    # # }
	 
 
 
   # # if(modelType=="RRBLUP_REML"){
 
    # # # model_Sol_Geno_List_Combined_Sampled <- list()
	# # # model_Sol_Pheno_List_Combined_Sampled <- list()
  
	
	 # # for(nConditions in 1:6){
	
	    # # model_Sol_Geno_List_Combined_Sampled[[nConditions]] <-	foreach(i=1:nreps,.export=c("trainGenoNewTableListReps_Combined_Sampled","trainSimGenoValuesTableListReps_Combined_Sampled","testGenoNewTableListReps_Combined_Sampled","testSimGenoValuesTableListReps_Combined_Sampled","buildRRBLUPModel_REML")) %dopar% (SoyNAMPredictionMethods::buildRRBLUPModel_REML(trainGenoNewTableListReps_Combined_Sampled[[i]][[nConditions]], trainSimGenoValuesTableListReps_Combined_Sampled[[i]][[nConditions]][,1],testGenoNewTableListReps_Combined_Sampled[[i]][[nConditions]],testSimGenoValuesTableListReps_Combined_Sampled[[i]][[nConditions]][,1]))
    
    
    
		 # # model_Sol_Pheno_List_Combined_Sampled[[nConditions]] <- foreach(i=1:nreps,.export=c("trainGenoNewTableListReps_Combined_Sampled","trainSimPhenoValuesTableListReps_Combined_Sampled","testGenoNewTableListReps_Combined_Sampled","testSimPhenoValuesTableListReps_Combined_Sampled","buildRRBLUPModel_REML")) %dopar% (SoyNAMPredictionMethods::buildRRBLUPModel_REML(trainGenoNewTableListReps_Combined_Sampled[[i]][[nConditions]],trainSimPhenoValuesTableListReps_Combined_Sampled[[i]][[nConditions]][,1],testGenoNewTableListReps_Combined_Sampled[[i]][[nConditions]],testSimPhenoValuesTableListReps_Combined_Sampled[[i]][[nConditions]][,1]))
    
    
    # # }
   
   # # }

# # nReps <- 10
# # nConditions <- 6

# # PredAccuracy_Combined_Sampled <- rep(list(rep(list(list()),nReps)),nConditions)


# # for(nCon in 1:6){

  # # for(nrep in 1:10){ 
  
    
		
		
		# # PredAccuracy_Combined_Sampled[[nCon]][[nrep]] <- (model_Sol_Pheno_List_Combined_Sampled[[nCon]][[nrep]][[3]])
		
	 
  # # }
# # }


# # PredAccuracy_Condition_Combined_Sampled <- list()

# # for(nCon in 1:6){

 
    # # Avg_PredAccuracy_Combined_Sampled <- mean(unlist(PredAccuracy_Combined[[nCon]]))

    # # PredAccuracy_Condition_Combined_Sampled[[nCon]] <- Avg_PredAccuracy_Combined_Sampled
 # # } 
 
   
 # # # [[1]]
# # # [1] 0.5759605

# # # [[2]]
# # # [1] 0.2309474

# # # [[3]]
# # # [1] 0.4700512

# # # [[4]]
# # # [1] 0.1847925

# # # [[5]]
# # # [1] 0.5261836

# # # [[6]]
# # # [1] 0.3240252


# # ### Prediction Accuracy for Combined data : same as trainGS with/with out QTL


  
  # # modelType <- ModelType
  # # nreps <- length(trainGenoNewTableListReps)
  # # nCon <- length(trainGenoNewTableListReps[[1]])
  # # h2 <- c(0.7,0.3,0.7,0.3,0.7,0.3)
  # # no_Cores <- nCores
  
 
 
    # # model_Sol_Geno_List_Combined <- list()
	# # model_Sol_Pheno_List_Combined <- list()
  
	
	# # for(nConditions in 1:6){
	
	   # # model_Sol_Geno_List_Combined[[nConditions]] <-	foreach(i=1:nreps,.export=c("trainGenoNewTableListReps_Combined","trainSimGenoValuesTableListReps_Combined","testGenoNewTableListReps_Combined","testSimGenoValuesTableListReps_Combined","buildRRBLUPModel_REML")) %dopar% (SoyNAMPredictionMethods::buildRRBLUPModel_REML(trainGenoNewTableListReps_Combined[[i]][[nConditions]], trainSimGenoValuesTableListReps_Combined[[i]][[nConditions]][,1],testGenoNewTableListReps_Combined[[i]][[nConditions]],testSimGenoValuesTableListReps_Combined[[i]][[nConditions]][,1]))
    
    
    
		# # model_Sol_Pheno_List_Combined[[nConditions]] <- foreach(i=1:nreps,.export=c("trainGenoNewTableListReps_Combined","trainSimPhenoValuesTableListReps_Combined","testGenoNewTableListReps_Combined","testSimPhenoValuesTableListReps_Combined","buildRRBLUPModel_REML")) %dopar% (SoyNAMPredictionMethods::buildRRBLUPModel_REML(trainGenoNewTableListReps_Combined[[i]][[nConditions]],trainSimPhenoValuesTableListReps_Combined[[i]][[nConditions]][,1],testGenoNewTableListReps_Combined[[i]][[nConditions]],testSimPhenoValuesTableListReps_Combined[[i]][[nConditions]][,1]))
    
    
    # # }
   
 
 
   
# # nReps <- 10
# # nConditions <- 6

# # PredAccuracy_Combined <- rep(list(rep(list(list()),nReps)),nConditions)


# # for(nCon in 1:6){

  # # for(nrep in 1:10){ 
  
    
		
		
		# # PredAccuracy_Combined[[nCon]][[nrep]] <- (model_Sol_Pheno_List_Combined[[nCon]][[nrep]][[3]])
		
	 
  # # }
# # }


# # PredAccuracy_Condition_Combined <- list()

# # for(nCon in 1:6){

 
    # # Avg_PredAccuracy_Combined <- mean(unlist(PredAccuracy_Combined[[nCon]]))

    # # PredAccuracy_Condition_Combined[[nCon]] <- Avg_PredAccuracy_Combined
 # # } 
 
 
  # # PredAccuracy_Condition_Combined
# # [[1]]
# # [1] 0.7721484

# # [[2]]
# # [1] 0.457694

# # [[3]]
# # [1] 0.7694313

# # [[4]]
# # [1] 0.451491

# # [[5]]
# # [1] 0.7601086

# # [[6]]
# # [1] 0.4524069


# ## nQTL400 / h20.7

# # Average within Family TS - 0.586 
# # Combined TS_Constant TS Size - 0.47
# # Combined TS - 0.77 
```
