## Supplementary material for "Factors Affecting Response to Recurrent Genomic Selection in Soybeans": File S1: Rcpp_LDV3.cpp

```
#include <Rcpp.h>
using namespace Rcpp;
//[[Rcpp::export]]
SEXP LD_V3(NumericVector p1,NumericVector q1,NumericVector a, NumericMatrix pro,NumericMatrix D,NumericMatrix D1,NumericMatrix R2){
	
int m = D.ncol();
int n = a.size();
int s = pro.ncol();
double pab = 0;
double pa1a1b1b1 = 0;
double pa1a1b1b2 = 0;
double pa1a2b1b2 = 0;
double pa1a2b1b1 = 0;
int R2Denom = 0;

for (int j = 0; j < m-1; j++) {
 		
 for (int i = a(j); i < n; i++) { 
	
  for (int k = 0; k< s; k++) {     	
	if(pro(j,k)==1 && pro(i,k)==1){
		pa1a1b1b1 += 1;
	}
	if(pro(j,k)==1 && pro(i,k)==0){
		pa1a1b1b2 += 1;
	}
	if(pro(j,k)==0 && pro(i,k)==0){
		pa1a2b1b2 += 1;
	}
	if(pro(j,k)==0 && pro(i,k)==1){
		pa1a2b1b1 += 1;
	}
  }
  pab= ((2*pa1a1b1b1)+pa1a1b1b2+(0.5*pa1a2b1b2)+pa1a2b1b1)/(2*s);
  
  D(i,j)=pab-(p1(j)*p1(i));

  if(D(i,j) >0){ 

      double PQ1 = p1(i)*q1(j);
      double PQ2 = q1(i)*p1(j); 
      double Dmax = std::min(PQ1,PQ2);
      D1(i,j) = D(i,j)/Dmax ;
	  if(Dmax !=0 ){
		D1(i,j) = D(i,j)/Dmax ;
	   }
	  if(Dmax ==0 ){
        D1(i,j) = 999 ;
	  }
  } 
  if(D(i,j) <0){
     double PQ1 = p1(i)*p1(j);
     double PQ2 = q1(i)*q1(j); 
     double Dmax = std::min(PQ1,PQ2); 
	 if(Dmax !=0 ){
     D1(i,j) = D(i,j)/Dmax ;
	 }
	 if(Dmax ==0 ){
      D1(i,j) = 999 ;
	 }
  } 
  if(D(i,j)==0){ 
    D1(i,j) = D(i,j);
  }
     
  R2Denom = p1[j]*p1[i]*q1[j]*q1[i];
  
  if(R2Denom > 0){
	R2(i,j)= (D(i,j)*D(i,j))/R2Denom;
  }
  if(R2Denom ==0){
	R2(i,j)= 999; 
  }
  
   pab = 0;
  }
 }
  return List::create(Named("D")=D,Named("D1")=D1,Named("R2")=R2);
}
```
