## Supplementary material for "Factors Affecting Response to Recurrent Genomic Selection in Soybeans": File S1: RcppExports.cpp

```
// Generated by using Rcpp::compileAttributes() -> do not edit by hand
// Generator token: 10BE3573-1514-4C36-9D1C-5A225CD40393

#include <Rcpp.h>

using namespace Rcpp;

// LD
SEXP LD(NumericVector p1, NumericVector q1, NumericVector a, NumericMatrix pro, NumericMatrix D, NumericMatrix D1, NumericMatrix R2);
RcppExport SEXP _SoyNAMPredictionMethods_LD(SEXP p1SEXP, SEXP q1SEXP, SEXP aSEXP, SEXP proSEXP, SEXP DSEXP, SEXP D1SEXP, SEXP R2SEXP) {
BEGIN_RCPP
    Rcpp::RObject rcpp_result_gen;
    Rcpp::RNGScope rcpp_rngScope_gen;
    Rcpp::traits::input_parameter< NumericVector >::type p1(p1SEXP);
    Rcpp::traits::input_parameter< NumericVector >::type q1(q1SEXP);
    Rcpp::traits::input_parameter< NumericVector >::type a(aSEXP);
    Rcpp::traits::input_parameter< NumericMatrix >::type pro(proSEXP);
    Rcpp::traits::input_parameter< NumericMatrix >::type D(DSEXP);
    Rcpp::traits::input_parameter< NumericMatrix >::type D1(D1SEXP);
    Rcpp::traits::input_parameter< NumericMatrix >::type R2(R2SEXP);
    rcpp_result_gen = Rcpp::wrap(LD(p1, q1, a, pro, D, D1, R2));
    return rcpp_result_gen;
END_RCPP
}
// LD_V3
SEXP LD_V3(NumericVector p1, NumericVector q1, NumericVector a, NumericMatrix pro, NumericMatrix D, NumericMatrix D1, NumericMatrix R2);
RcppExport SEXP _SoyNAMPredictionMethods_LD_V3(SEXP p1SEXP, SEXP q1SEXP, SEXP aSEXP, SEXP proSEXP, SEXP DSEXP, SEXP D1SEXP, SEXP R2SEXP) {
BEGIN_RCPP
    Rcpp::RObject rcpp_result_gen;
    Rcpp::RNGScope rcpp_rngScope_gen;
    Rcpp::traits::input_parameter< NumericVector >::type p1(p1SEXP);
    Rcpp::traits::input_parameter< NumericVector >::type q1(q1SEXP);
    Rcpp::traits::input_parameter< NumericVector >::type a(aSEXP);
    Rcpp::traits::input_parameter< NumericMatrix >::type pro(proSEXP);
    Rcpp::traits::input_parameter< NumericMatrix >::type D(DSEXP);
    Rcpp::traits::input_parameter< NumericMatrix >::type D1(D1SEXP);
    Rcpp::traits::input_parameter< NumericMatrix >::type R2(R2SEXP);
    rcpp_result_gen = Rcpp::wrap(LD_V3(p1, q1, a, pro, D, D1, R2));
    return rcpp_result_gen;
END_RCPP
}
// rrReml
SEXP rrReml(NumericVector y, NumericMatrix gen, int maxit, double tol);
RcppExport SEXP _SoyNAMPredictionMethods_rrReml(SEXP ySEXP, SEXP genSEXP, SEXP maxitSEXP, SEXP tolSEXP) {
BEGIN_RCPP
    Rcpp::RObject rcpp_result_gen;
    Rcpp::RNGScope rcpp_rngScope_gen;
    Rcpp::traits::input_parameter< NumericVector >::type y(ySEXP);
    Rcpp::traits::input_parameter< NumericMatrix >::type gen(genSEXP);
    Rcpp::traits::input_parameter< int >::type maxit(maxitSEXP);
    Rcpp::traits::input_parameter< double >::type tol(tolSEXP);
    rcpp_result_gen = Rcpp::wrap(rrReml(y, gen, maxit, tol));
    return rcpp_result_gen;
END_RCPP
}

static const R_CallMethodDef CallEntries[] = {
    {"_SoyNAMPredictionMethods_LD", (DL_FUNC) &_SoyNAMPredictionMethods_LD, 7},
    {"_SoyNAMPredictionMethods_LD_V3", (DL_FUNC) &_SoyNAMPredictionMethods_LD_V3, 7},
    {"_SoyNAMPredictionMethods_rrReml", (DL_FUNC) &_SoyNAMPredictionMethods_rrReml, 4},
    {NULL, NULL, 0}
};

RcppExport void R_init_SoyNAMPredictionMethods(DllInfo *dll) {
    R_registerRoutines(dll, NULL, CallEntries, NULL, NULL);
    R_useDynamicSymbols(dll, FALSE);
}
```
