## Supplementary material for "Factors Affecting Response to Recurrent Genomic Selection in Soybeans": File S1: WGR_rrREML.cpp

```
#include <Rcpp.h>
using namespace Rcpp;
// [[Rcpp::export]]
SEXP rrReml(NumericVector y, NumericMatrix gen, int maxit = 500, double tol = 10e-6){
  // Functions starts here
  int p = gen.ncol();
  int n = gen.nrow();
  // Beta, mu and epsilon
  double b0, eM, ve, vb, h2, mu = mean(y), vy = var(y);
  NumericVector b(p), e = y-mu;
  // Marker variance
  NumericVector xx(p), vx(p);
  for(int i=0; i<p; i++){
    xx[i] = sum(gen(_,i)*gen(_,i));
    vx[i] = var(gen(_,i));}
  double MSx = sum(vx), Lmb=MSx;
  // Convergence control
  NumericVector bc(p);
  int numit = 0;
  double cnv = 1;
  // Loop
  while(numit<maxit){
    // Regression coefficients loop
    bc = b+0;
    for(int j=0; j<p; j++){
      b0 = b[j];
      b[j] = (sum(gen(_,j)*e)+xx[j]*b0)/(xx[j]+Lmb);
      e = e-gen(_,j)*(b[j]-b0);}
    // Variance components update
    ve = sum(e*y)/(n-1);
    vb = (vy-ve)/MSx;
    Lmb = ve/vb;
    // Intercept update
    eM = mean(e);
    mu = mu+eM;
    e = e-eM;
    // Convergence
    ++numit;
    cnv = sum(abs(bc-b));
    if( cnv<tol ){break;}}
  // Fitting the model
  NumericVector fit(n);
  for(int k=0; k<n; k++){ fit[k] = sum(gen(k,_)*b)+mu; }
  h2 = vb*MSx/(vb*MSx+ve);
  // Output
  return List::create(Named("mu")=mu, Named("b")=b,
                      Named("h2")=h2, Named("hat")=fit,
                      Named("Vb")=vb, Named("Ve")=ve);}
```
