## Supplemental Figures for "Factors Affecting Response to Recurrent Genomic Selection in Soybeans"

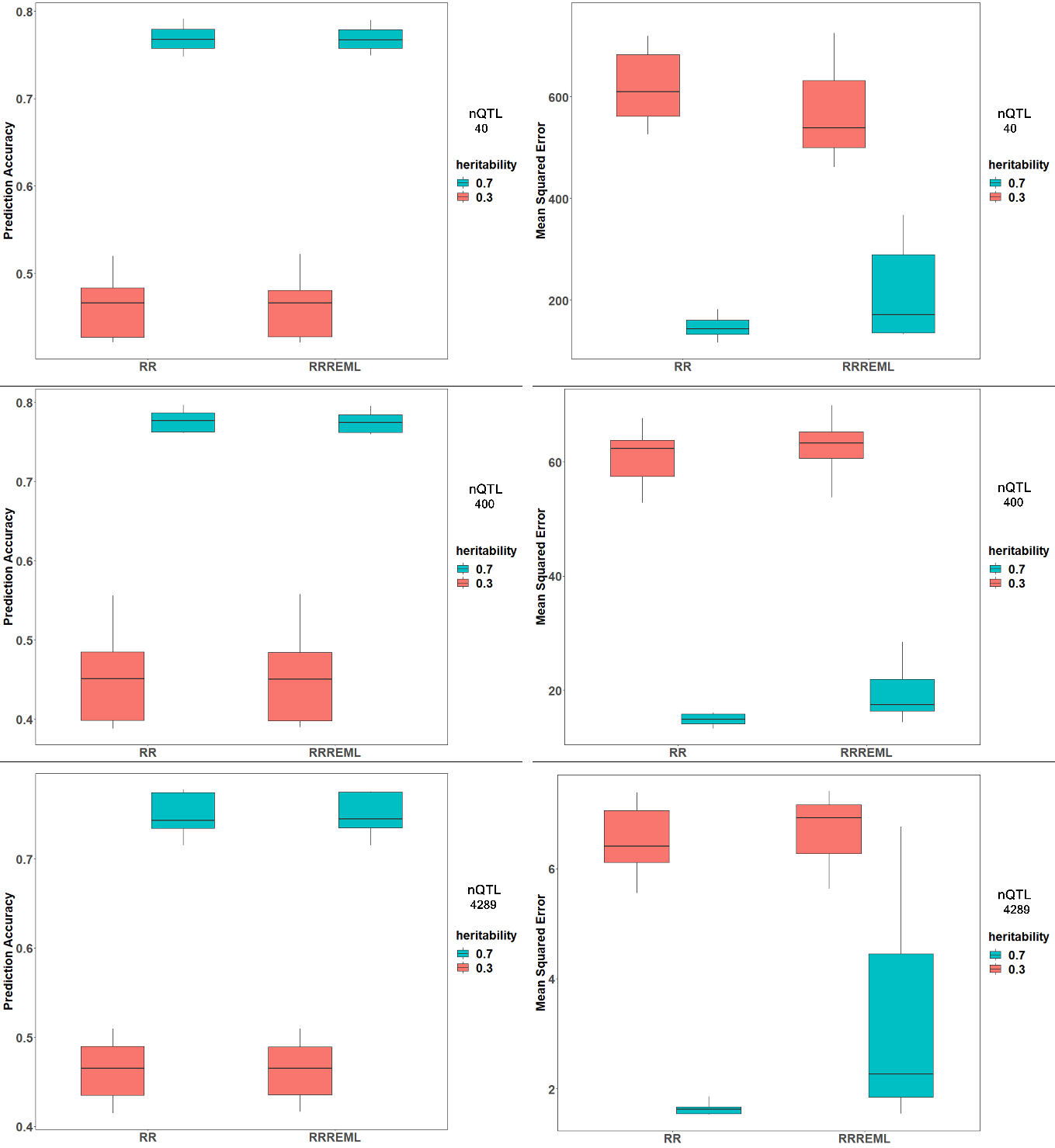

**Figure S1** **Evaluation of Two Ridge Regression GS Model Implementations** Prediction accuracy and Mean Squared Error are plotted for Ridge regression (RR) implemented in rrBLUP package and Ridge regression-REML for 40, 400 and 4289 QTL for two levels of heritability. Ridge regression GS model implemented in rrBLUP package uses mixed model equation solver that involves matrix inversion. This step is often time consuming and cannot be implemented in cases with singular matrices. Expectation Maximization algorithm to obtain Restricted Maximum Likelihood Estimates of marker effects, which doesn’t require the matrix inversion step (Xavier 2019). Code for implementing both the methods can be found in ‘SoyNAMPredictionMethods’ R package. Prediction accuracy and MSE of both the implementations of Ridge Regression are comparable, but EM-REML method is much faster than rrBLUP implementation. We used RR-REML training algorithm in our simulations that involve model updating with prior cycle training data set.

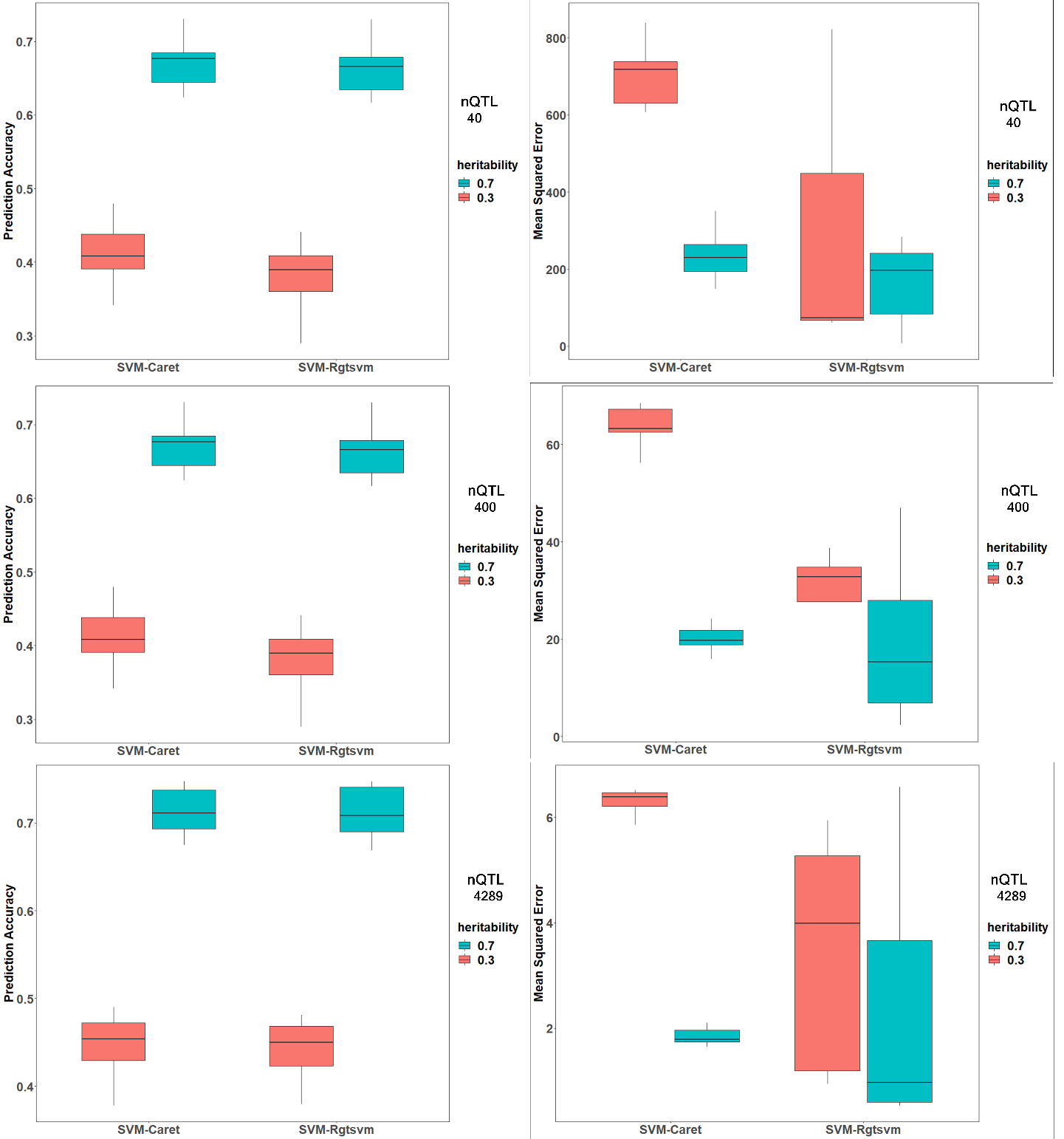

**Figure S2 Evaluation of Two Support Vector Machine Implementations**

SVM Model with radial basis function implemented in ‘caret’ and ‘Rgtsvm’ packages in R. ‘Rgtsvm’ implementation based on ‘e1071’ allows parallel model training in GPUs, which reduces computing time by several hundred fold. SVM with Radial Basis Kernel function implemented in ‘caret’ package is compared with SVM with RBF implemented in ‘Rgtsvm’. Prediction accuracy is plotted in the left panel and Mean Squared Error in the right panel for 40, 400 and 4289 QTL for two levels of heritability.

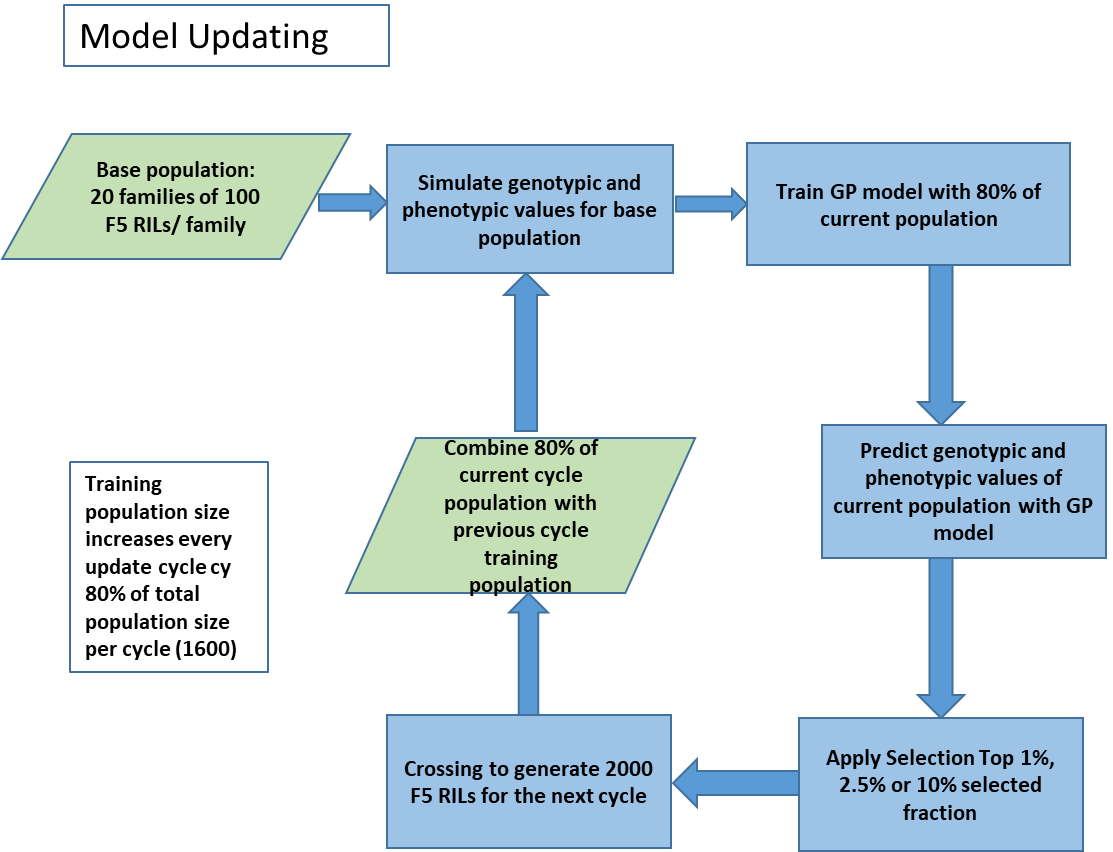

**Figure S3 Flow Chart for Model Updating in Simulations of Recurrent Genomic Selection**. Model updating involves combining training data from ‘n-1’ previous cycles (t-n-1...t-1) with training data from the t^th^ cycle to retrain genomic prediction models. ‘t’ refers to the selection cycle and ranges from 1 to 40 and ‘n’ refers to number of prior cycles that are included in the training set. The treatment design has four levels for ‘n’ ranging from 0 - 14. ‘n=0’ refers to no updating, whereas ‘n= 10’, ‘n=12’ and ‘n=14’ refer to inclusion of training data from 9, 11 and 13 prior cycles along with data from 10^th^, 12^th^, and 14^th^ cycle of selection.

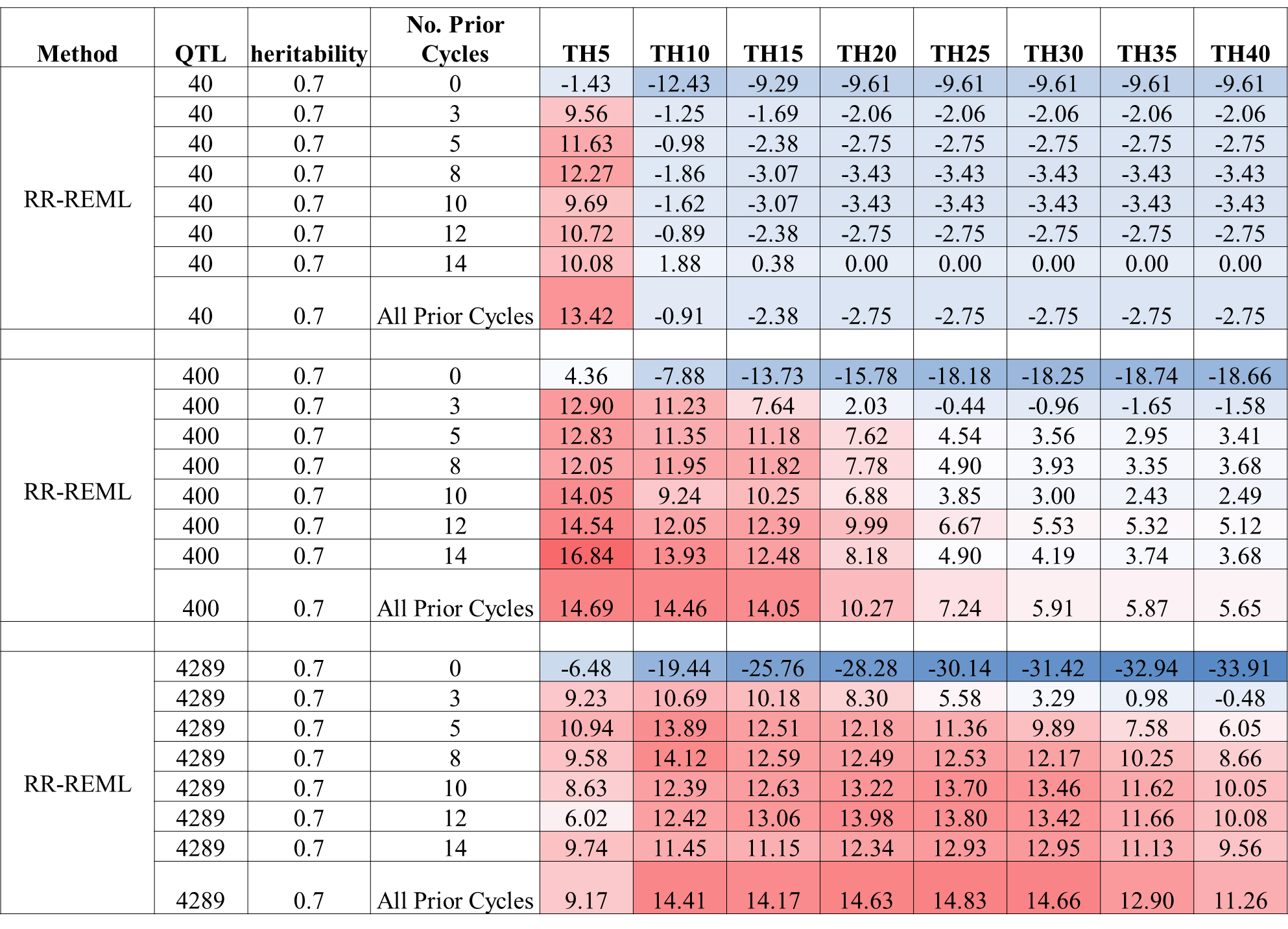

**Figure S4 Heat map for percent gain in Rs relative to PS for 0.7 H:** Heat map representing percentage gain in Rs_Max_ at the 40^th^ cycle with respect to PS with Ridge Regression genomic prediction model with no update and model updated every cycle with training data from prior cycles. Percent gain in response is provided for 40, 400 and 4289 simulated QTL responsible for 70% of phenotypic variability in the initial population with top 10% selected fraction. Blue to red shaded cells represent increasing gain in response relative to PS.

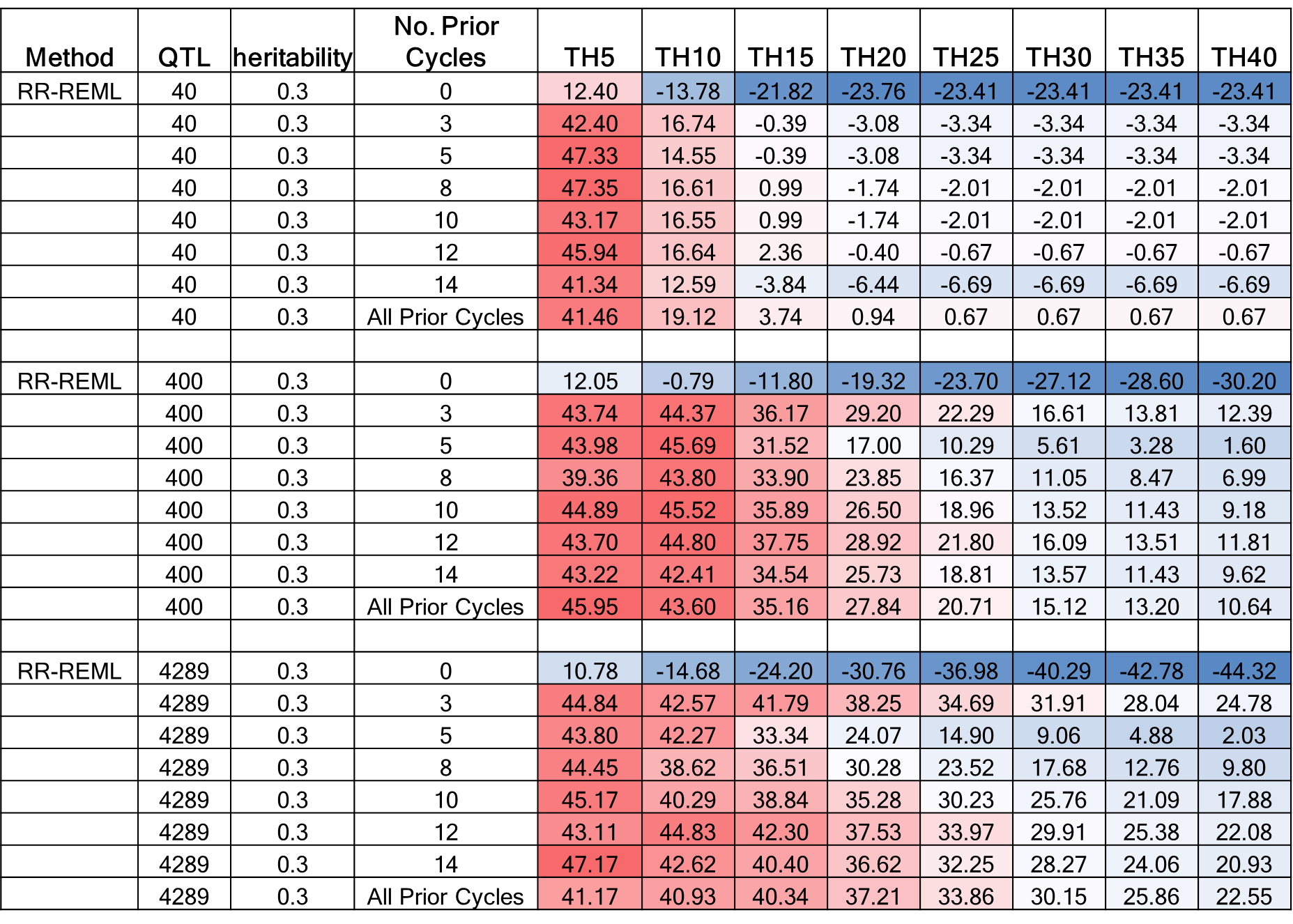

**Figure S5 Heat Map for Percent Gain in Responses for RR GS Relative to PS for 0.3 H:** Heat map representing percentage gain in Rs_Max_ with respect to PS with Ridge Regression genomic prediction model with no update and model updated every cycle with training data from prior cycles. Percent gain in response is provided for 40, 400 and 4289 simulated QTL responsible for 30% of phenotypic variability in the initial population for top 10% selected fraction. Blue to red shaded cells represent increasing gain in response relative to PS in the respective cycles of selection.

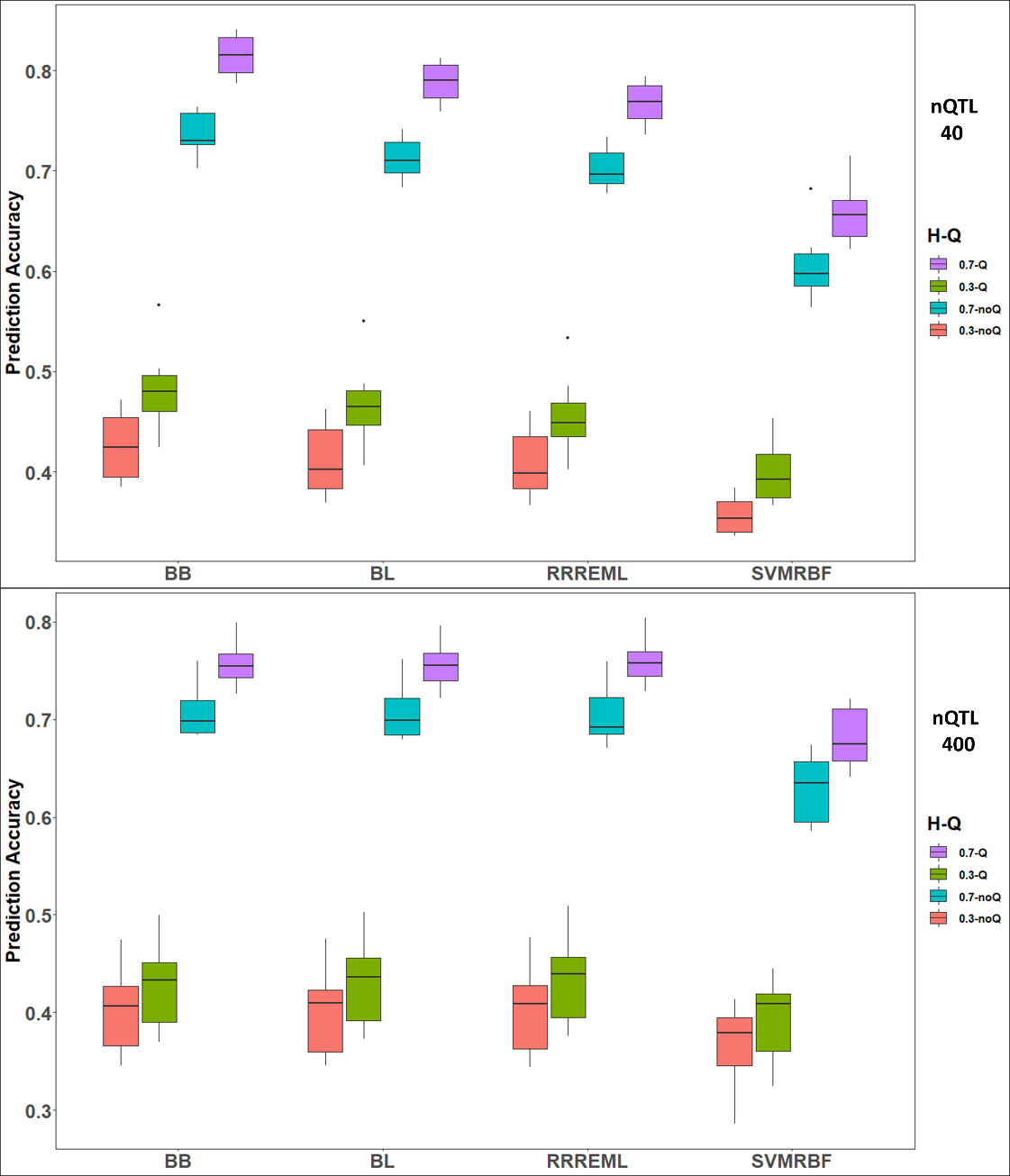

**Figure S6 Comparison of Estimated Prediction Accuracies of GP models in Founding Set of RILs using Training Sets with or without QTL:** Estimated prediction accuracies for four genomic prediction (GP) models. When QTL are included in the training set, estimated prediction accuracies (green and purple) are larger than that for models trained without QTL (red and blue) for all four models. Boxes represent distribution of estimates over 10 replicates for BB, BL and RRREML models, whereas SVMRBF distribution use estimates from less than five replicates. BB (BayesB), BL (Bayes LASSO), RRREML (Ridge Regression with REML) and SVMRBF (Support Vector Machines with Radial Basis Function Kernel) trained with F_5_ RILs derived from crosses of 20 homozygous founder lines with IA3023. Phenotypes used to train the GP models consisted of genetic architectures comprised of 40 and 400 simulated QTL (top and bottom) that were responsible for 70% (blue and purple) and 30% (green and red) of phenotypic variability in the initial populations.

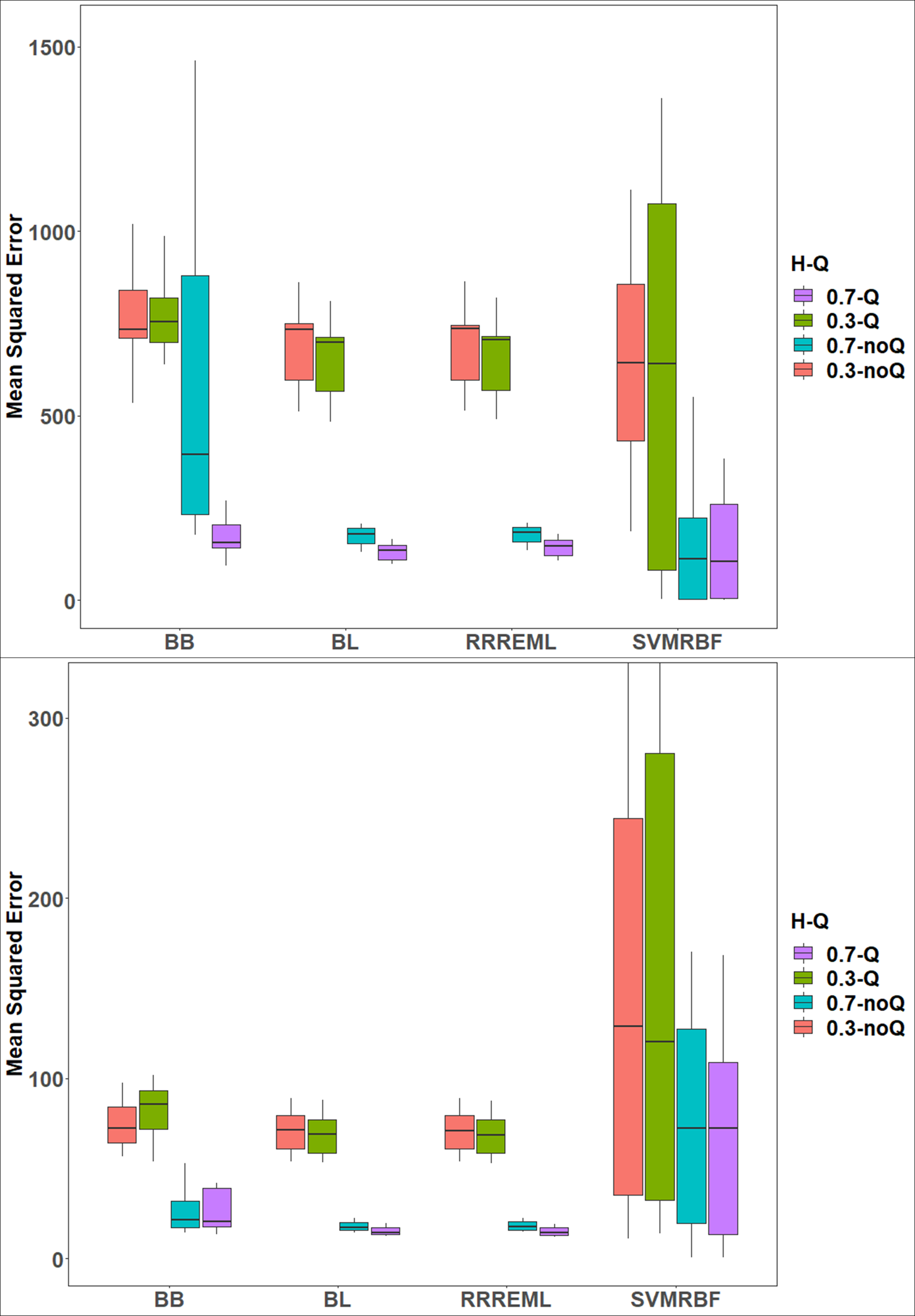

**Figure S7 Comparison of Estimated Mean Squared Error (MSE) of GP Models Trained Using Founding Set of RILs as Training Sets with or without QTL:** When QTL are included in the training set, MSE (green and purple) are larger or comparable to that of models trained without QTL (red and blue) for all four models. Boxes represent distribution of estimates over 10 replicates for BB, BL and RRREML models, whereas SVMRBF distribution use estimates from less than five replicates. BB (BayesB), BL (Bayes LASSO), RRREML (Ridge Regression with REML) and SVMRBF (Support Vector Machines with Radial Basis Function Kernel) trained with F_5_ RILs derived from crosses of 20 homozygous founder lines with IA3023. Phenotypes used to train the GP models consisted of genetic architectures comprised of 40 and 400 simulated QTL (top and bottom) that were responsible for 70% (blue and purple) and 30% (green and red) of phenotypic variability in the initial populations.

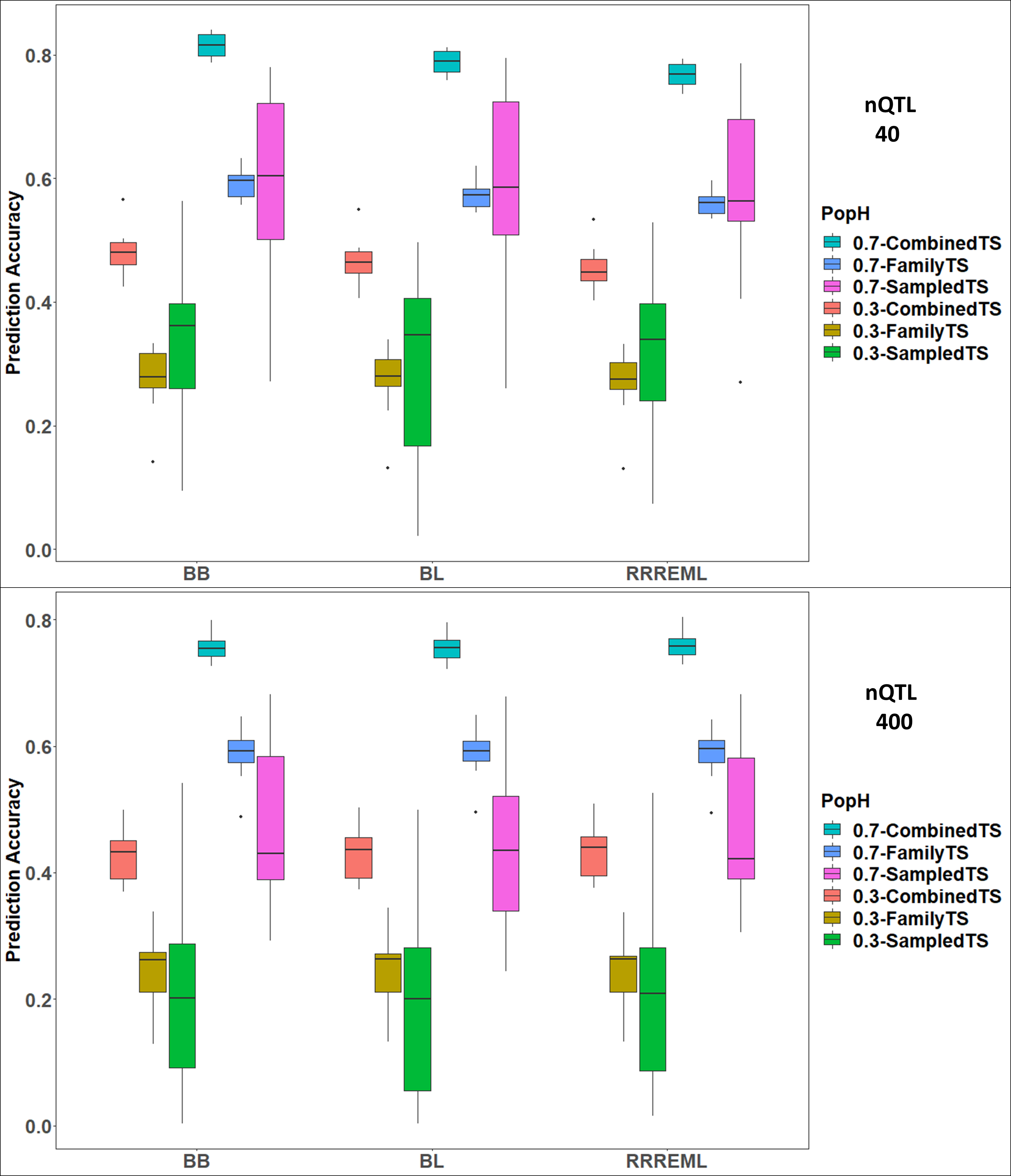

**Figure S8 Comparison of Estimated Prediction Accuracies of GP Models Trained Using Founding Set of RILs with Combined, Family and Sampled Training Sets (TS). i)** Combined TS are comprised of a combined population of RILs, ii) Family TS are comprised of RILs within families and iii) Sampled TS are comprised of RILs sampled from the combined population of RILs to keep the training set size same as family size. Prediction accuracies (sky blue and red) are larger when combined populations of RILs (1600 RILs) are used as TS than for models trained with RILS within families (80 RILs per family) (blue and brown) for the three parametric models. Accuracies are comparable to or lesser than average within family models when 80 RILs are sampled from a combined pool (green and magenta). Accuracies are lesser for 0.3 heritabilities than 0.7 heritabilities. Boxes represent distribution of estimates over 10 replicates for BB, BL and RRREML models, BB (BayesB), BL (Bayes LASSO), RRREML (Ridge Regression with REML) trained with F_5_ RILs derived from crosses of 20 homozygous founder lines with IA3023. Phenotypes used to train the GP models consisted of genetic architectures comprised of 40 and 400 simulated QTL (top and bottom) that were responsible for 70% (blue and purple) and 30% (green and red) of phenotypic variability in the initial populations. Accuracies of SVMRBF (Support Vector Machines with Radial Basis Function Kernel) are not provided as good fits were not obtained for most of the families.

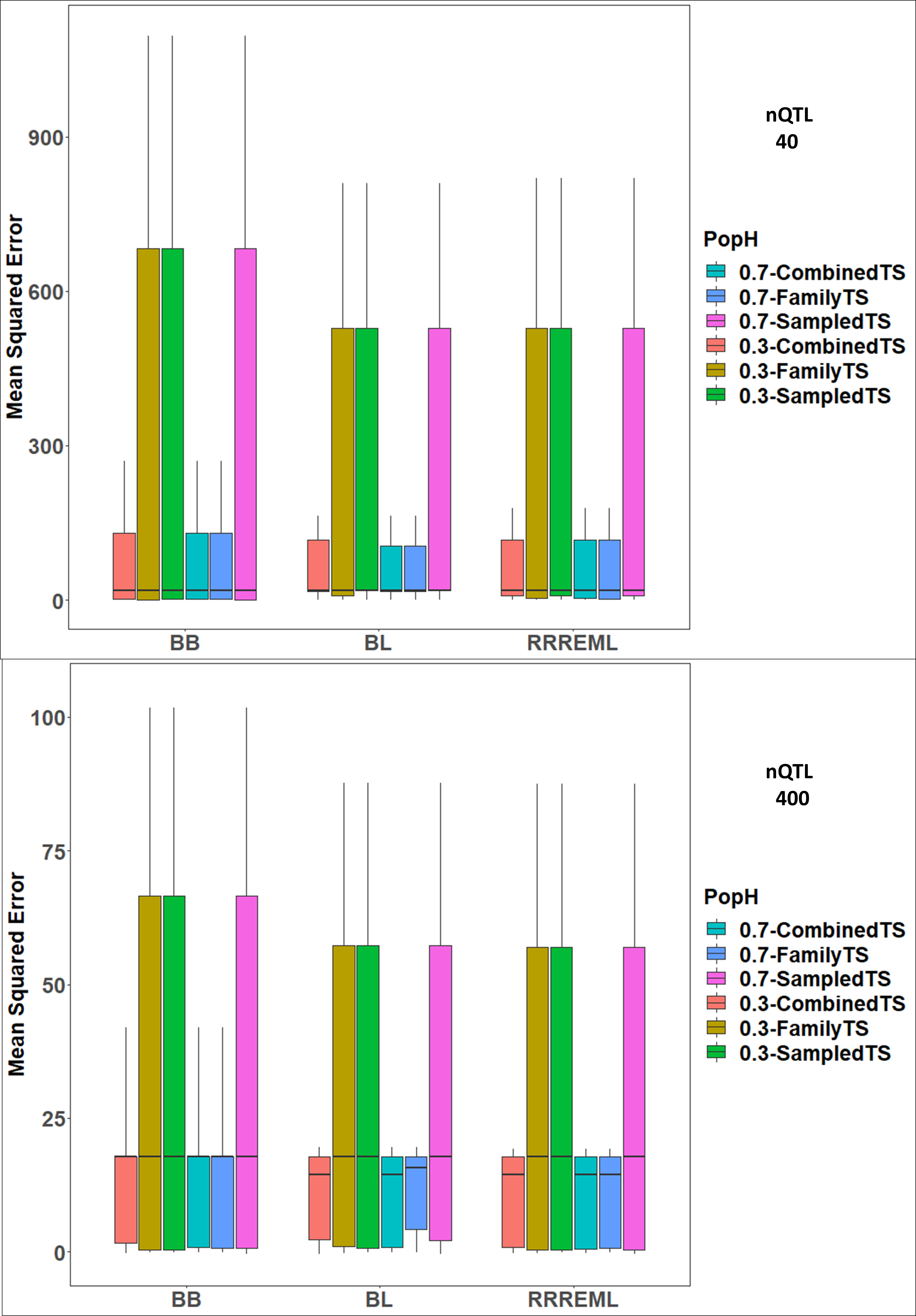

**Figure S9 Comparison of Estimated Mean Squared Error (MSE) of GP Models Trained Using Founding Set of RILs with Combined, Family and Sampled Training Sets.** Combined TS are comprised of a combined population of RILs, Family TS are comprised of RILs within families and Sampled TS are comprised of RILs sampled from the combined population of RILs to keep the training set size same as family size. MSE (sky blue and red) are lesser when combined populations of RILs (1600 RILs) RILS within families (80 RILs per family) (blue and brown) are used as TS for the three parametric models compared to models trained with 80 RILs that are sampled from a combined pool of 2000 RILs. MSE for sampled TS are larger due to sampling (green and magenta). MSE are lesser for 0.7 heritabilities compared to 0.3 heritabilities. Boxes represent distribution of estimates over 10 replicates for BB, BL and RRREML models, whereas SVMRBF distribution use estimates from less than five replicates. BB (BayesB), BL (Bayes LASSO), RRREML (Ridge Regression with REML) and SVMRBF (Support Vector Machines with Radial Basis Function Kernel) trained with F_5_ RILs derived from crosses of 20 homozygous founder lines with IA3023. Phenotypes used to train the GP models consisted of genetic architectures comprised of 40 and 400 simulated QTL (top and bottom) that were responsible for 70% (sky blue, blue and magenta) and 30% (red, brown and green) of phenotypic variability in the initial populations.

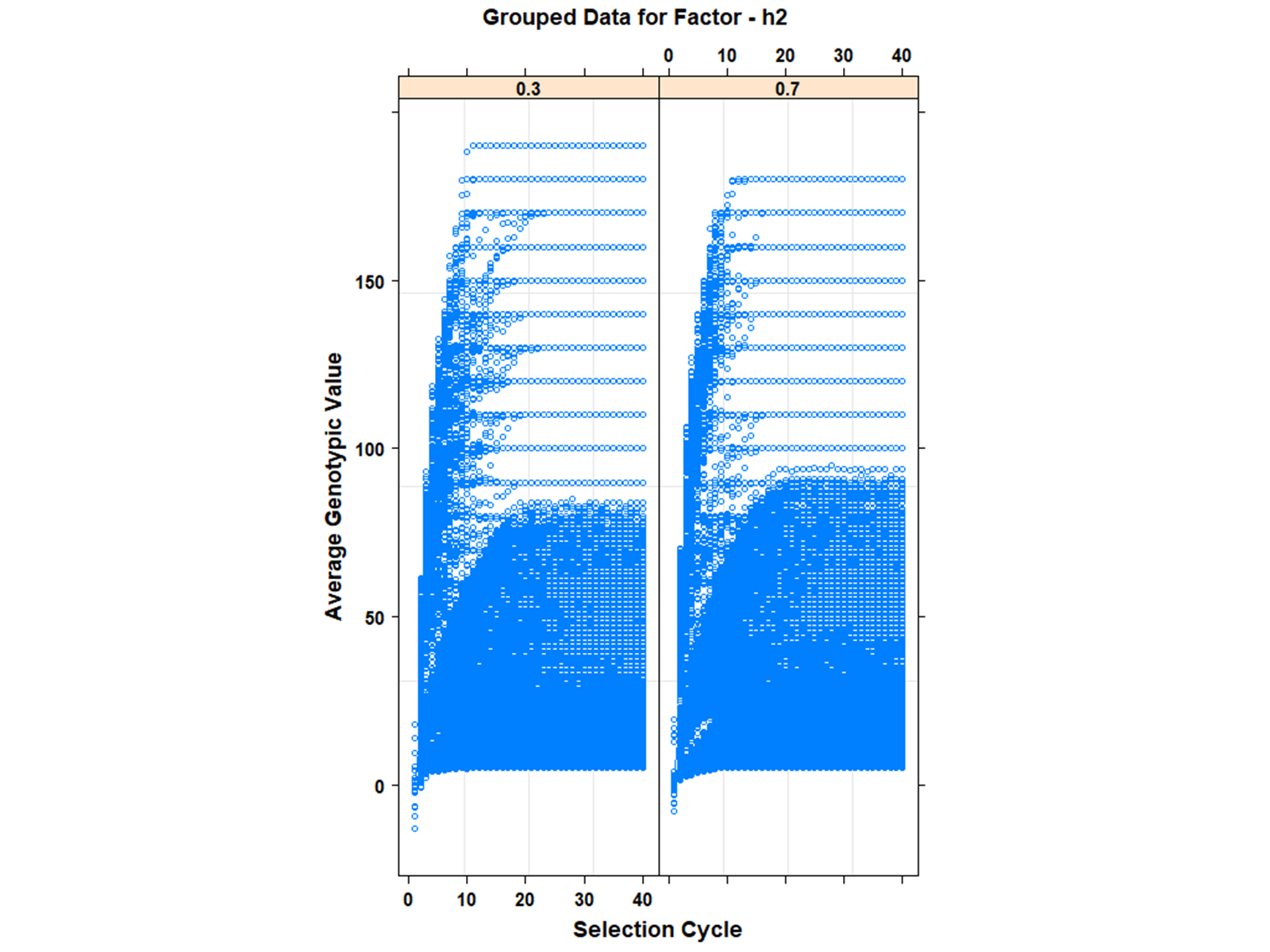

**Figure S7 Average Genotypic Values Grouped on Two H Levels:** Grouped data plot for selection intensity factor used in M1 ‘nlme’ fit with 2 curves for heritability levels 0.3 and 0.7 from left to right. Cycle number is plotted on x-axis and average genotypic values on y-axis.

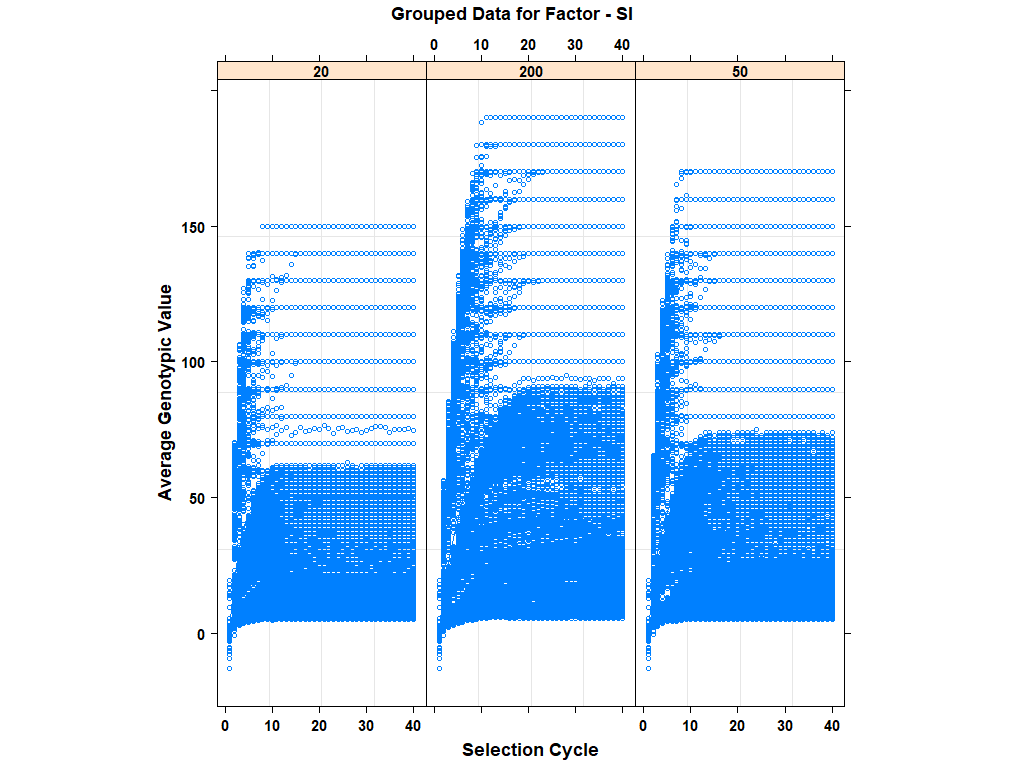

**Figure S11** **Average Genotypic Values Grouped on Three Selection Intensity Levels:** Grouped data plot for selection intensity factor used in M2 ‘nlme’ fit with three curves. Cycle number is plotted on x-axis and average genotypic value on y-axis. The three panels from left to right represent selection intensities of top 1% (20 RILs), 10% (200 RILs) and 2.5% (50 RILs)

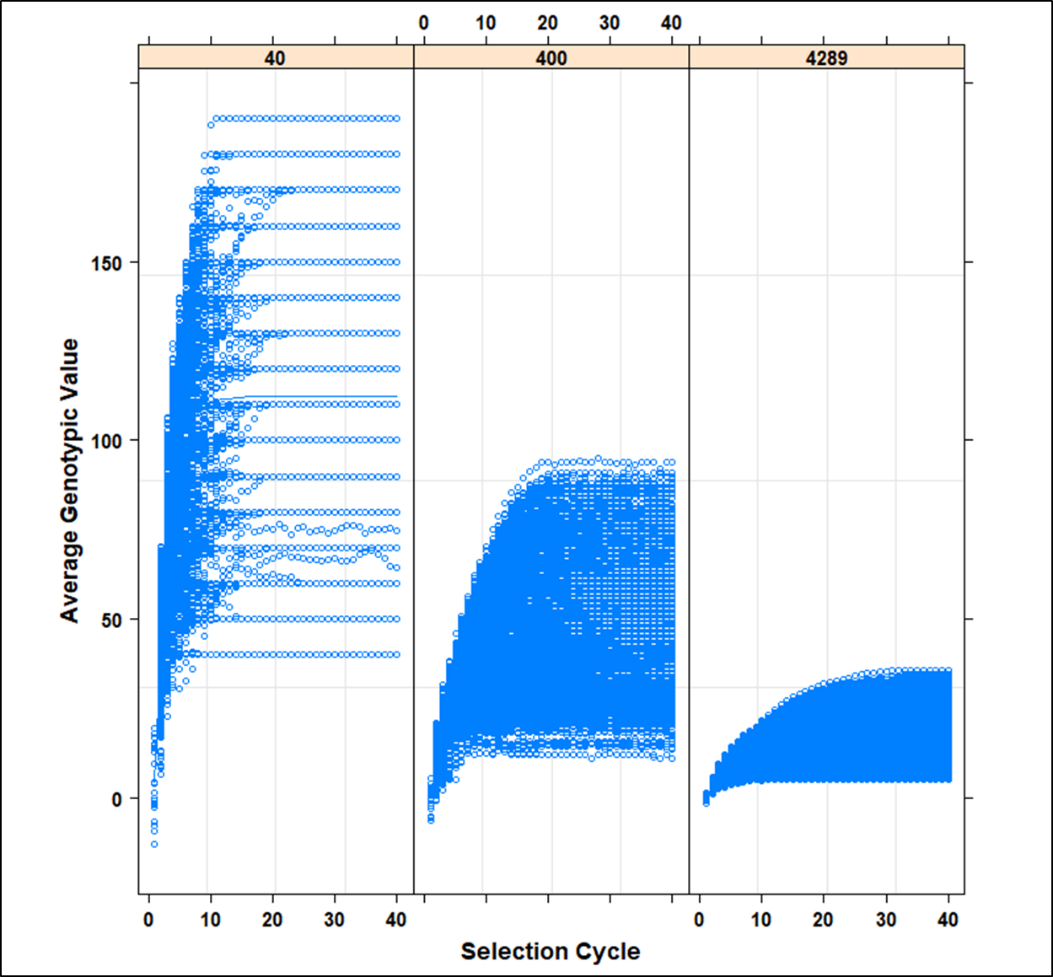

**Figure S12** **Average Genotypic Values Grouped on Three nQTL Levels:** Grouped data plot for nQTL factor used in M3 ‘nlme’ fit with three curves. Cycle number is plotted on x-axis and average genotypic value on y-axis. The three panels from left to right represent 40, 400 and 4289 simulated QTL.

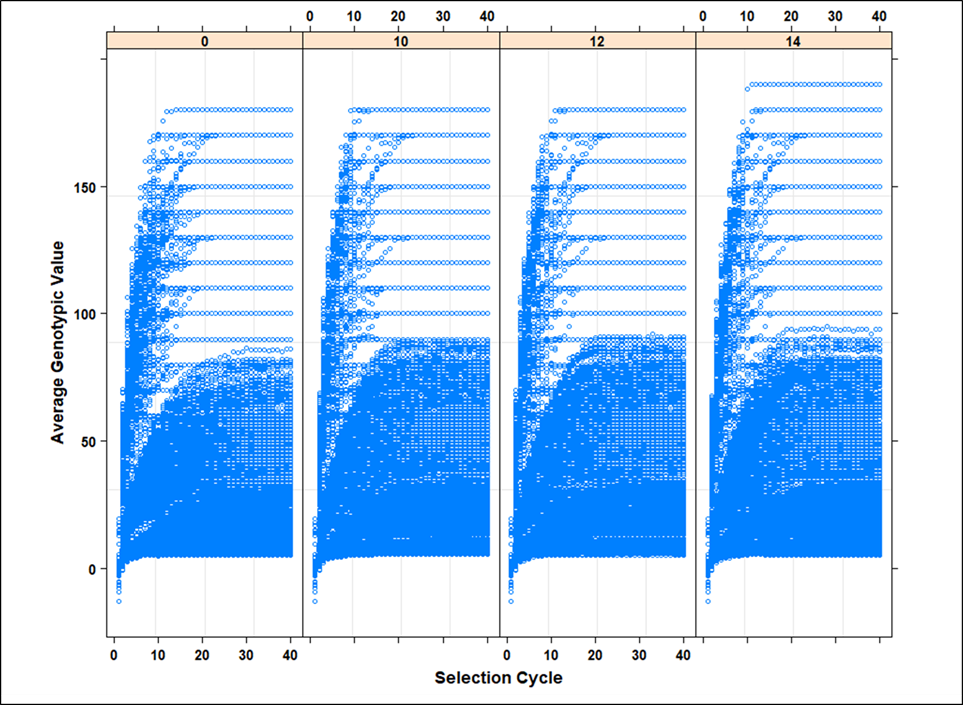

**Figure S13** **Average Genotypic Values Grouped on Four Training Set Levels:** Grouped data plot for number of prior cycles in GS model update training data used in M4 ‘nlme’ fit with four curves. Cycle number is plotted on x-axis and average genotypic value on y-axis. The four panels from left to right represent number of prior cycles in training sets with level 0, 10, 12 and 14. Level ‘0’ represent recurrent GS using GP models with no updating.

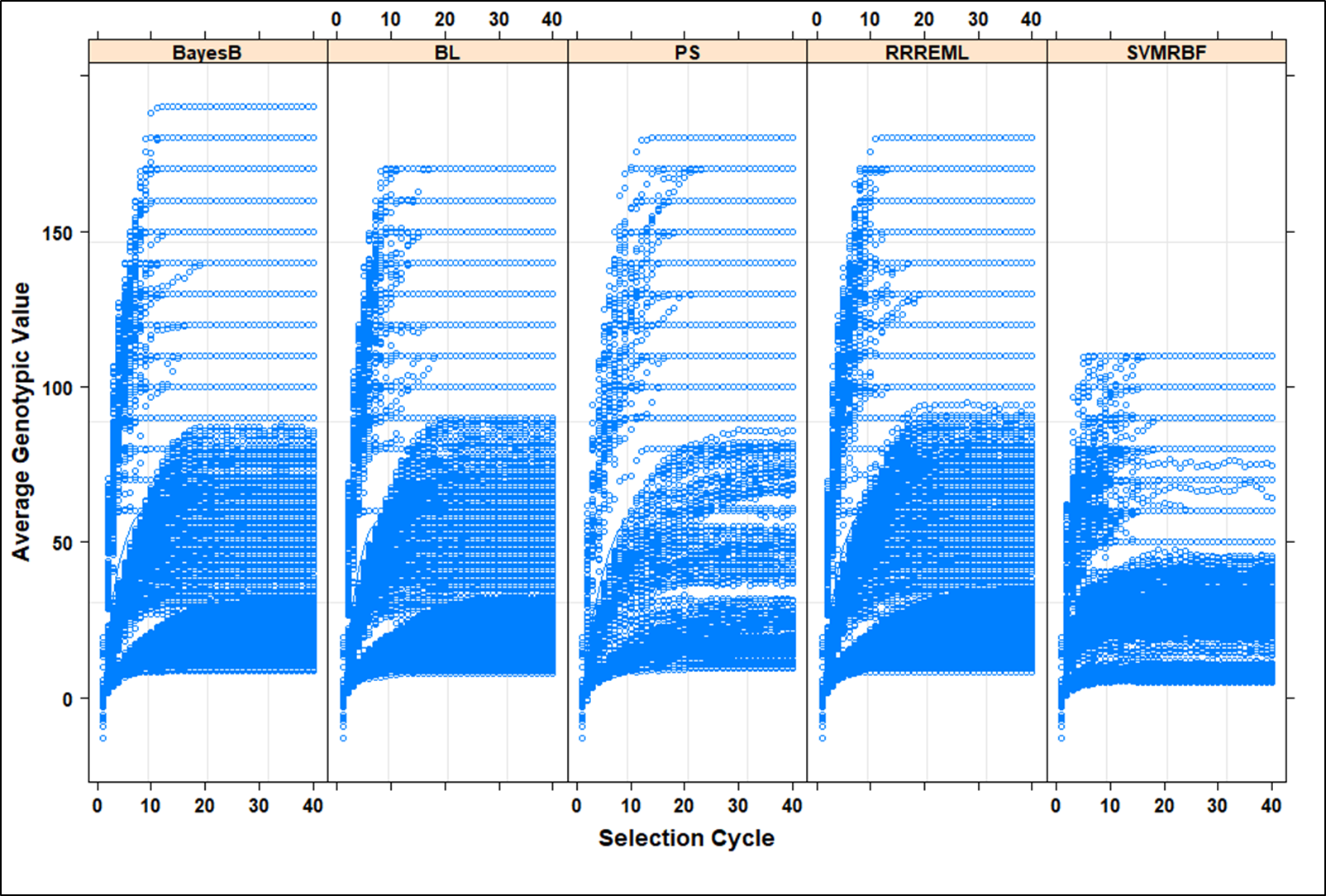

**Figure S14** **Average Genotypic Values Grouped on Five Selection Method Levels:** Grouped data plot for selection method that includes PS and GS with four GP models used in M5 ‘nlme’ fit with five curves. Cycle number is plotted on x-axis and average genotypic values on y-axis. The five panels from left to right represent selection methods that include BayesB, BL, PS, RR-REML and SVM-RBF. PS – Phenotypic Selection, RR-REML- Ridge Regression with Restricted Maximum Likelihood, BL – Bayes LASSO, and SVMRBF- Support Vector Machine with Radial Basis Kernel.

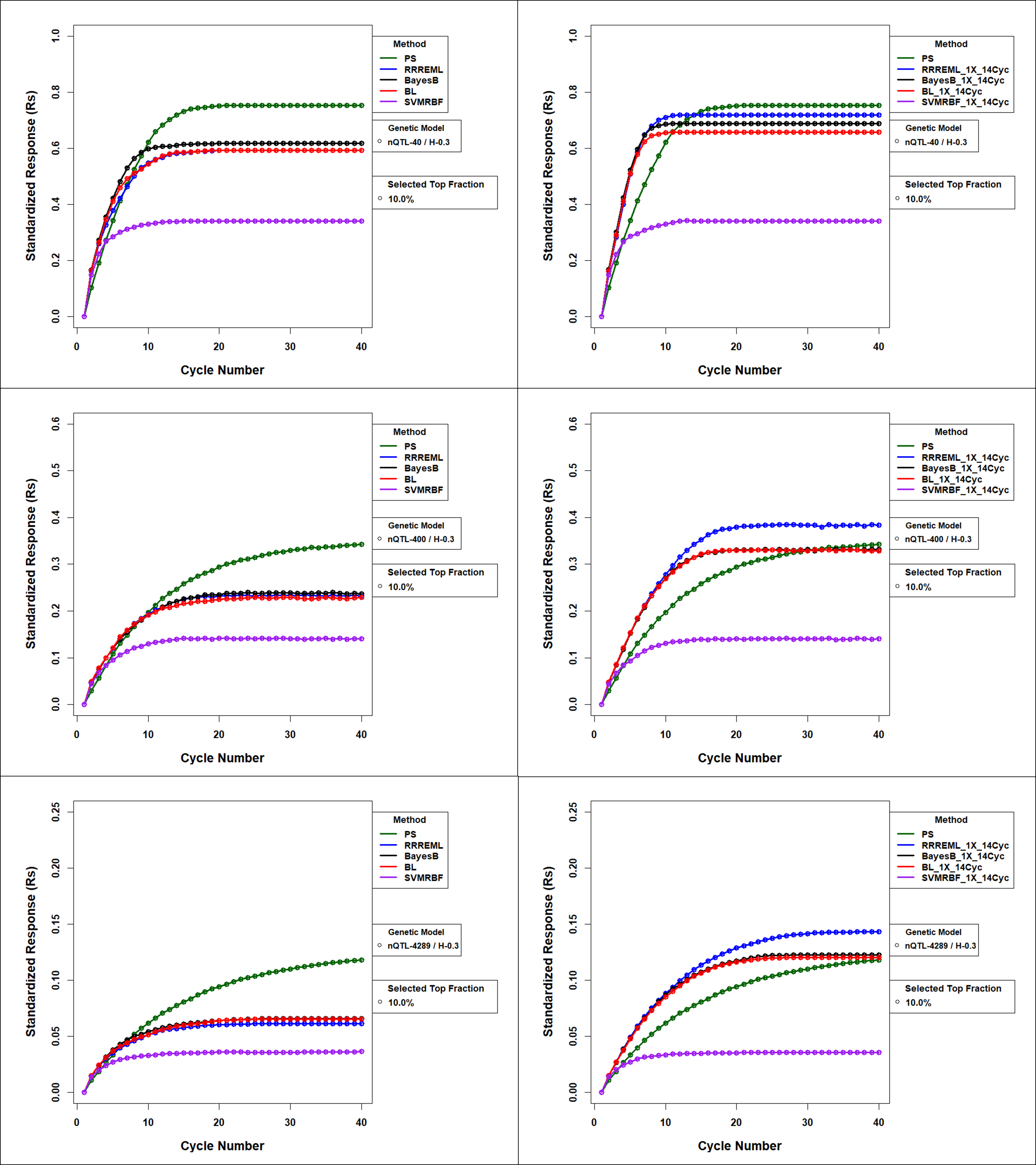

**Figure S15 Standardized Responses for Comparison of GS methods with and without Updating for 0.3 H and Top 10% Selected Fraction:** Forty cycles of response to selection of the top 10% of soybean RILs derived from SoyNAM founders. Responses are plotted by selection methods as genotypic values standardized to maximum possible value without model updating (left panels) and with model updating (right panels) using 14 prior cycles as training sets for the four GP models. Phenotypic selection (PS) is not updated and hence does not change between the left and right panels. Top panels consist of responses for genetic architectures consisting of 40 simulated QTL. Middle panels consist of responses for genetic architectures consisting of 400 simulated QTL and the bottom panels consist of responses for genetic architectures consisting of 4289 simulated QTL. All 40, 400 and 4289 are responsible for 30% of phenotypic variability in the initial population. PS – Phenotypic Selection, RR-REML- Ridge Regression with Restricted Maximum Likelihood, BL – Bayes LASSO, and SVMRBF- Support Vector Machine with Radial Basis Kernel.

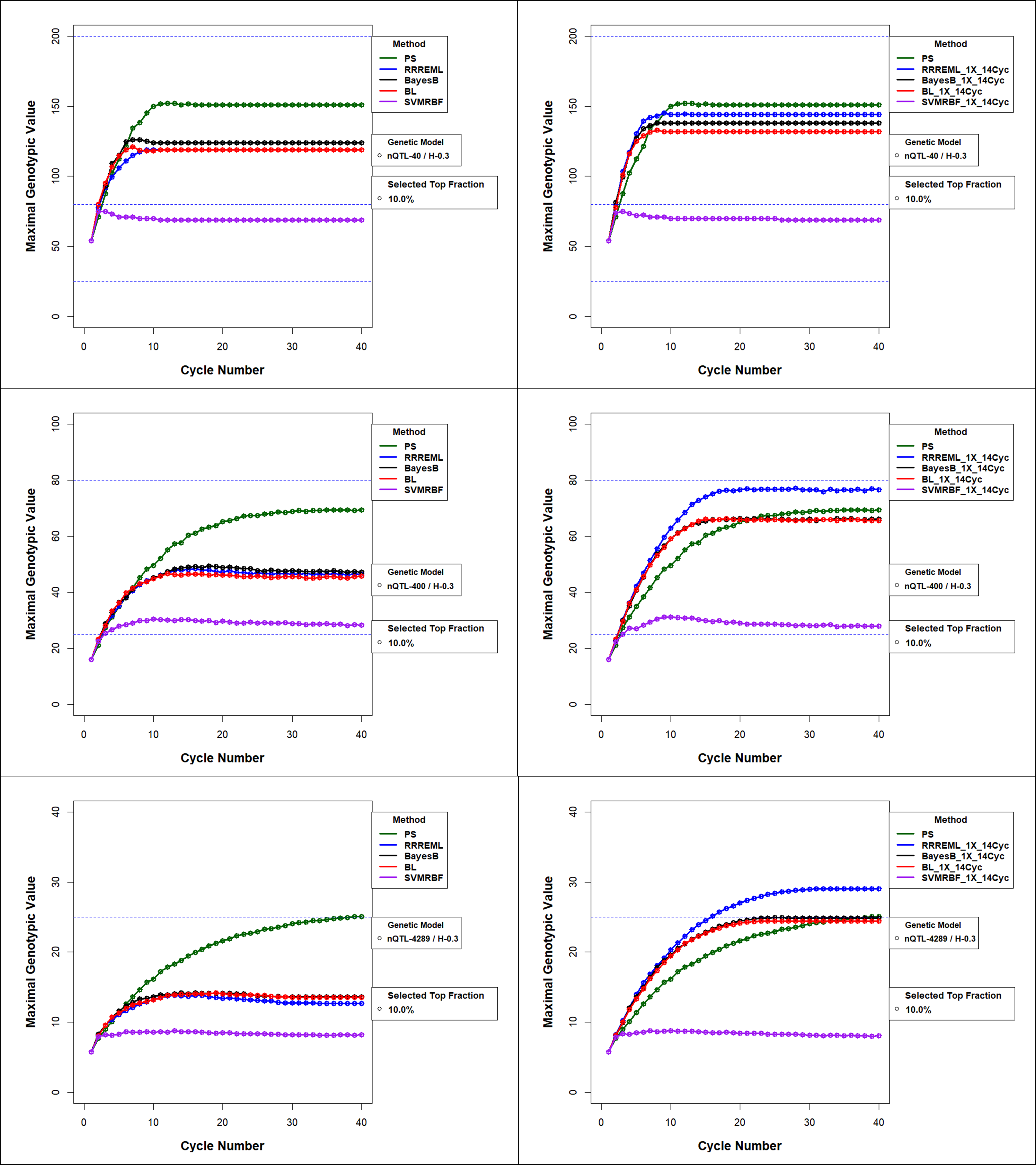

**Figure S16 Maximal Genotypic Value for Comparison of GS methods with and without Updating for 0.3 H and Top 10% Selected Fraction** Maximal genotypic values (Mgvs) in recurrent GS and PS without (left panels) and with model updating (right panels) using training data from up to 14 prior cycles for the four GP models. PS has no updating and hence does not change between the left and right panels. All treatments have top 10% selected fraction with 40 simulated QTL (top panels), 400 simulated QTL (middle panels) and 4289 simulated QTL (bottom panels) responsible for 30% of phenotypic variability in the initial population. PS – Phenotypic Selection, RR-REML- Ridge Regression with Restricted Maximum Likelihood, BL – Bayes LASSO, and SVMRBF- Support Vector Machine with Radial Basis Kernel.

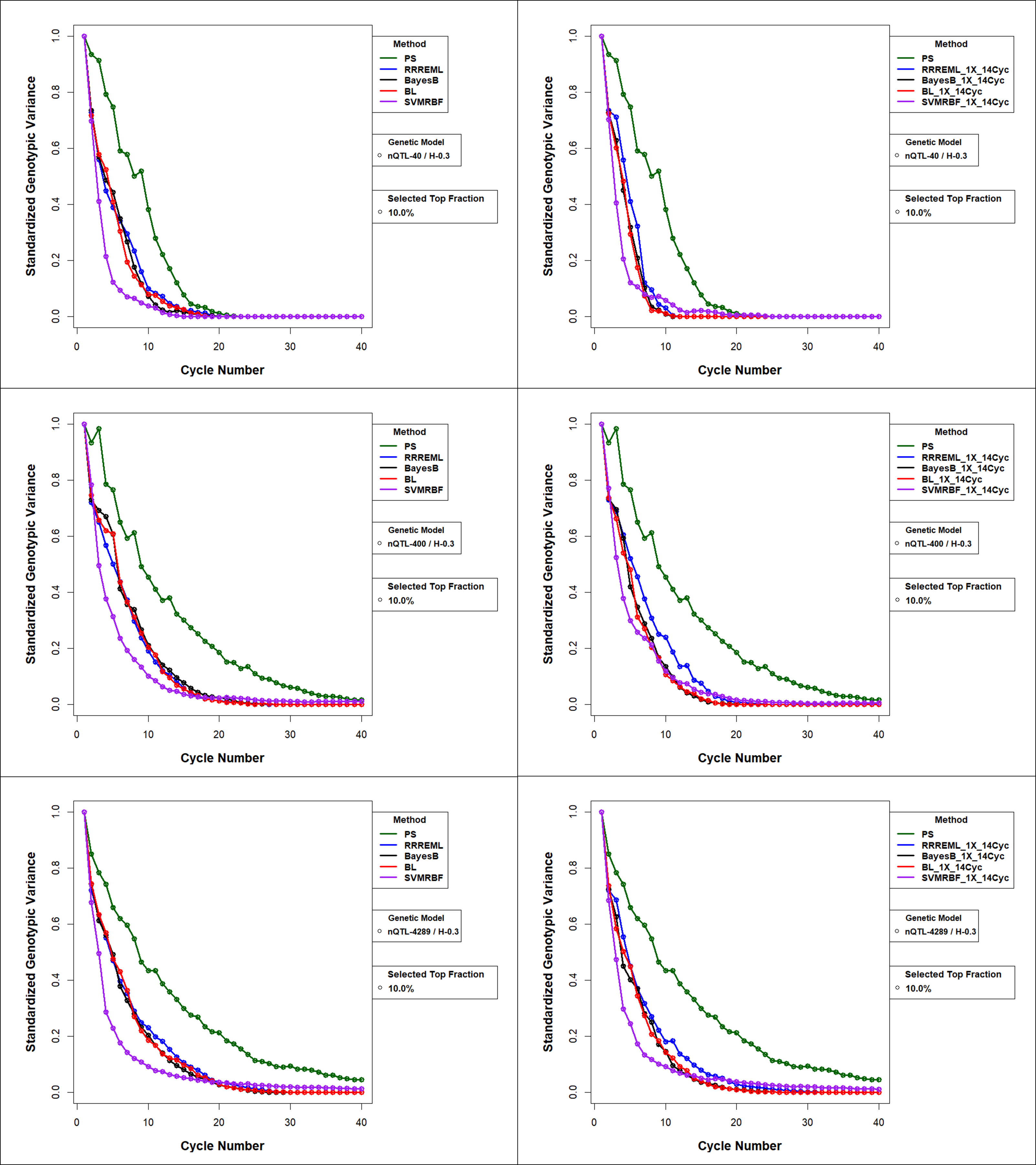

**Figure S17 Standardized Genotypic Variance (Sgv) for Comparison of GS methods with and without Updating for 0.3 H and Top 10% Selected Fraction** Standardized genotypic variance with model updating with training data from up to 14 prior cycles for the four GP models. PS has no updating and hence is the same in the left and right panels. All treatments have top 10% selected fraction with 40 simulated QTL (top panels), 400 simulated QTL (middle panels) and 4289 simulated QTL (bottom panels) responsible for 30% of phenotypic variability in the initial population. PS – Phenotypic Selection, RR-REML- Ridge Regression with Restricted Maximum Likelihood, BL – Bayes LASSO, and SVMRBF- Support Vector Machine with Radial Basis Kernel.

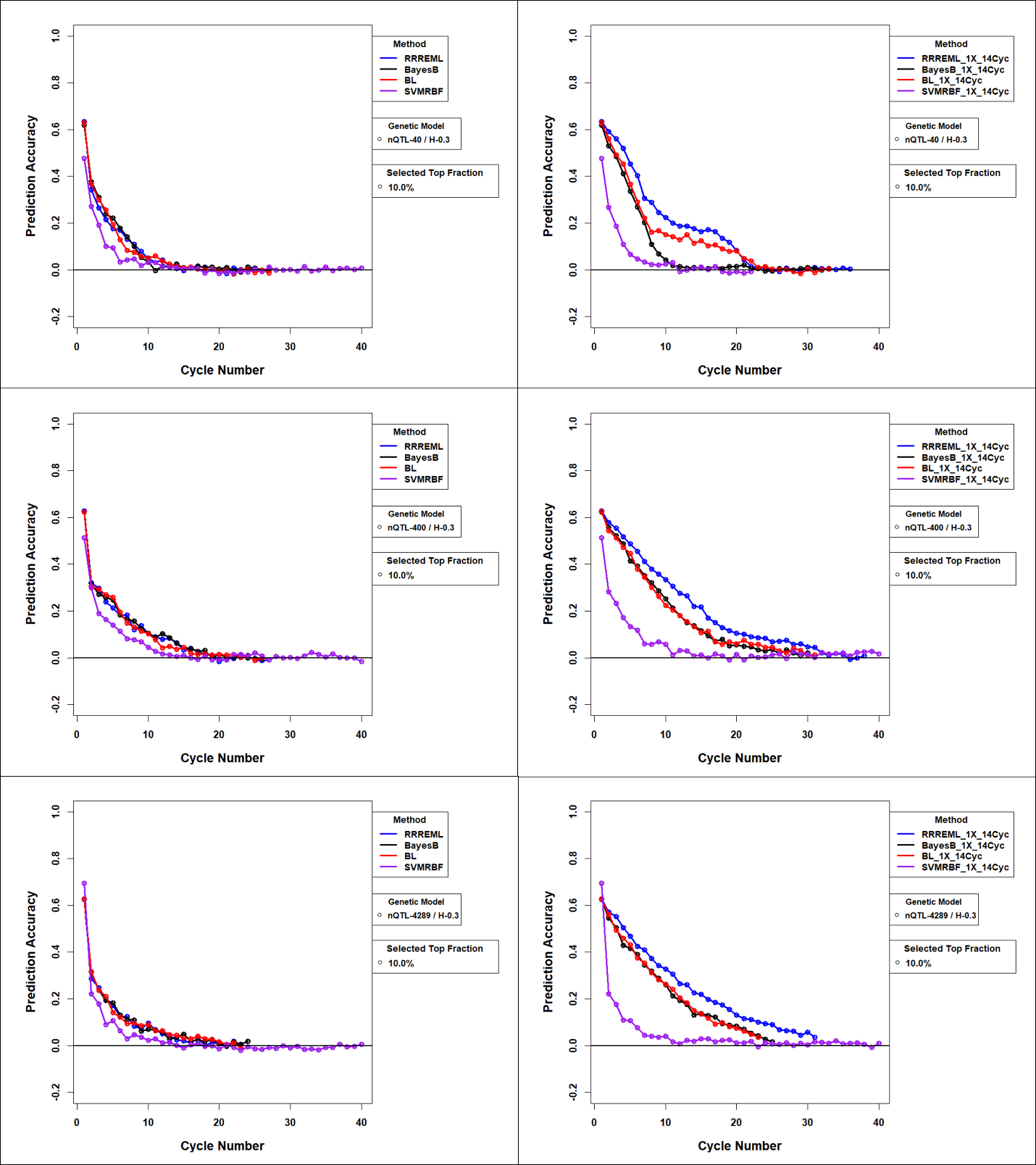

**Figure S18 Estimated Prediction Accuracies** **for Comparison of GS methods with and without Updating for 0.3 H and Top 10% Selected Fraction:** Estimated prediction accuracies with model updating with training data from up to 14 prior cycles for the four GP models. PS is not trained with 14 prior cycles of data and hence does not change between the left and right panels. All conditions have top 10% selected fraction with 14 prior cycles for 40 simulated QTL (top panels), 400 simulated QTL (middle panels), and 4289 simulated QTL (bottom panels). All 40, 400 and 4289 simulated QTL are responsible for 30% of phenotypic variability in the initial population. PS – Phenotypic Selection, RR-REML- Ridge Regression with Restricted Maximum Likelihood, BL – Bayes LASSO, and SVMRBF- Support Vector Machine with Radial Basis Kernel.

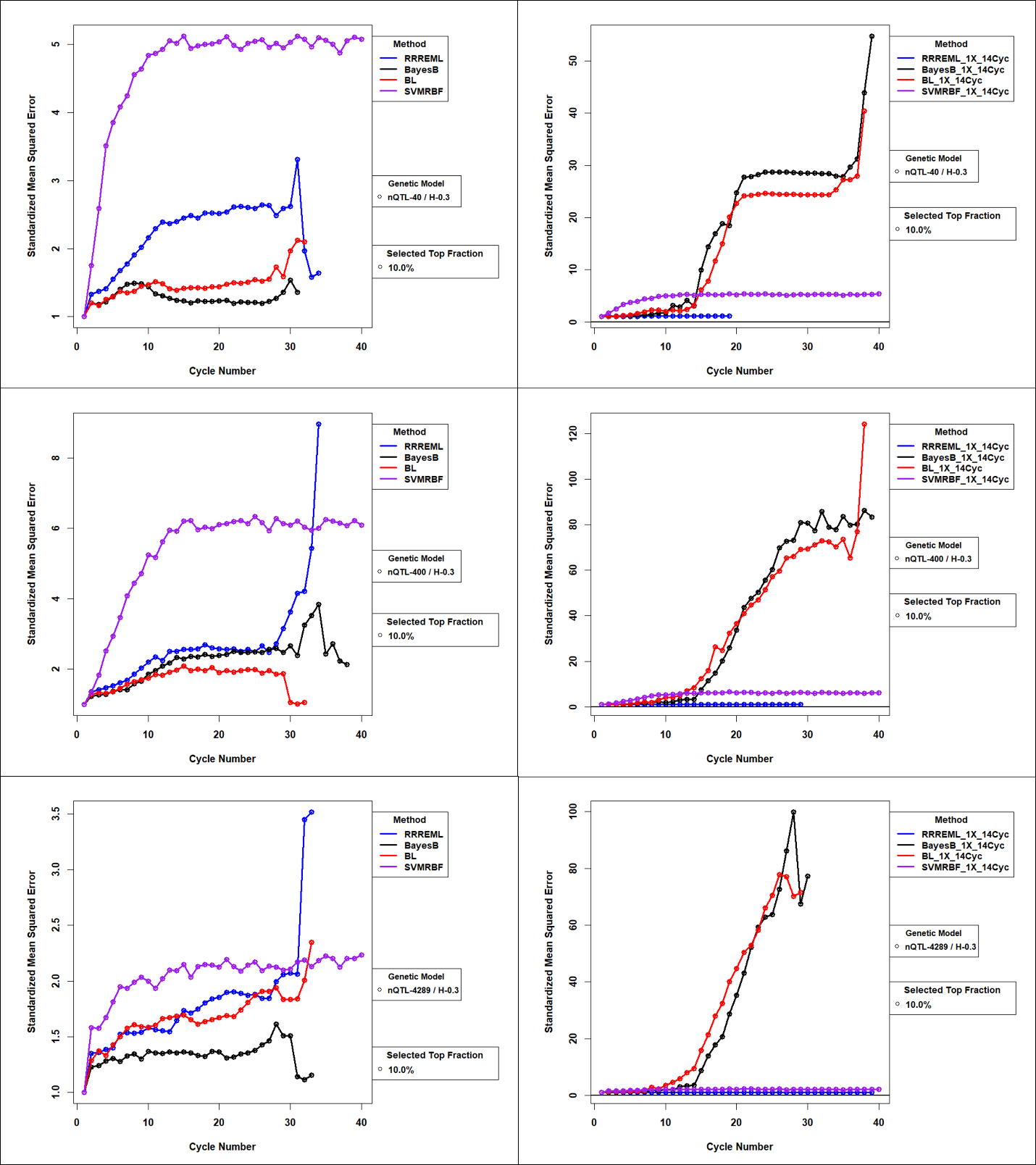

**Figure 19 Standardized Mean Squared Error** **for Comparison of GS methods with and without Updating for 0.7 H and Top 10% Selected Fraction:** Mean Squared Error of GP models with updates to the training sets used in genomic prediction (GP) models. Standardized MSE (>=1) is estimated as the ratio of MSE for GP models in cycle ‘c’ to MSE for GP models trained with founder population of RILs. While MSE for RRREML model were lesser with model updating, MSE for bayesian methods increased orders of magnitude in late cycles of selection with model updating. SVMEBF showed relatively constant MSE with and without updating. Training data from up to 14 prior selection cycles were used to update all four GP models for 40 QTL (top), 400 QTL (middle) and 4289 QTL (bottom) responsible for 70% of phenotypic variability in the initial population and top 10% of RILs with the greatest predicted values. RR-REML- Ridge Regression with Restricted Maximum Likelihood, BL – Bayes LASSO, and SVMRBF- Support Vector Machine with Radial Basis Kernel.

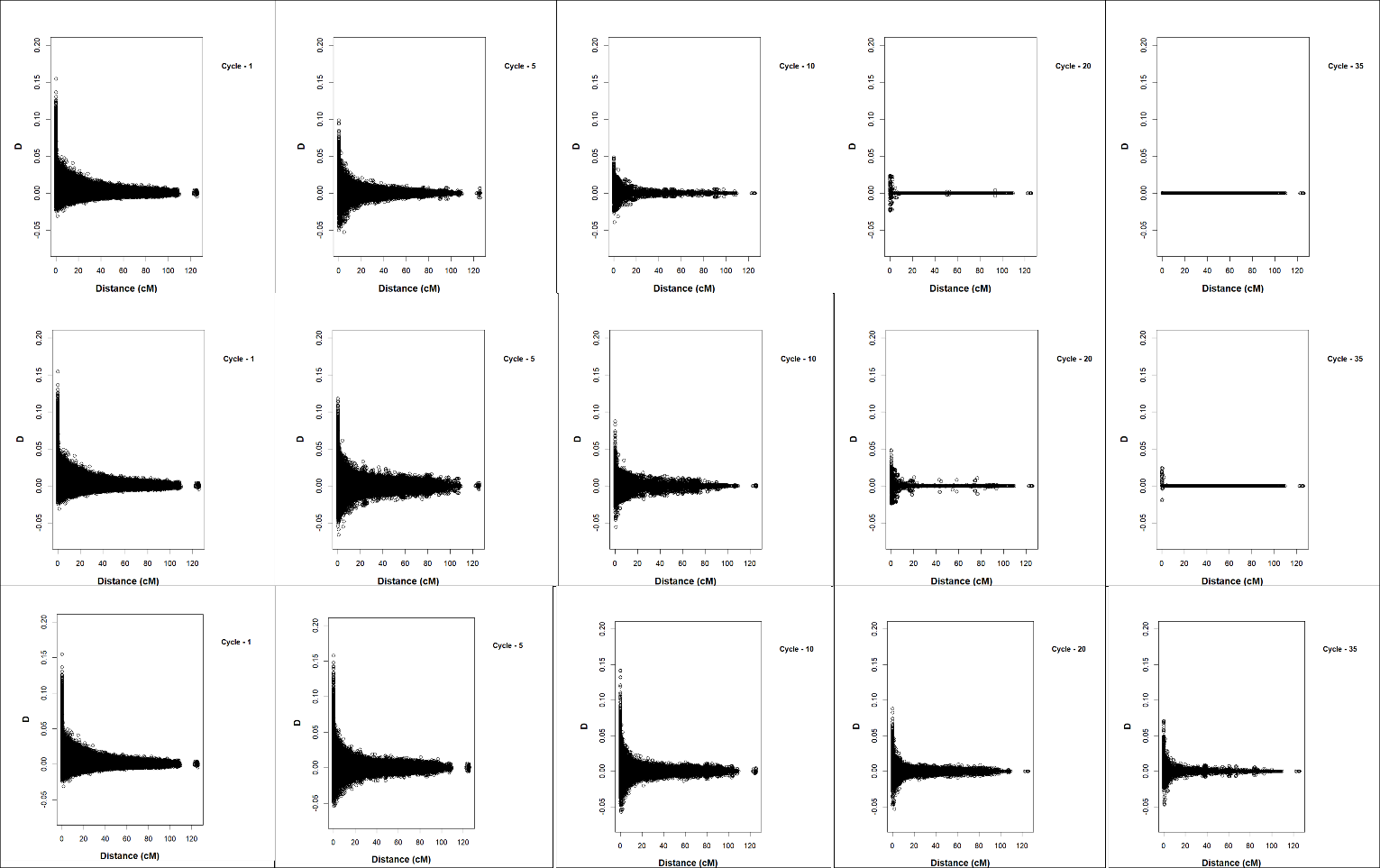

**Figure S20 Marker-marker LD decay with Distance between Marker Pairs within a Chromosome** **in cM for PS**. The figures represent distribution of LD estimated with ‘D’ statistic in selection cycles 1, 5, 10, 20 and 35 from left to right. ‘D’ is plotted on y-axis and distance between marker pairs within a chromosome in cM. Top panel represents population under PS with top 1% selected fraction. Middle panel represents population under PS with top 2.5% selected fraction. Bottom panel represents population under PS with top 10% selected fraction. All treatment combinations have 400 simulated QTL responsible for 70% of phenotypic variability in the initial population.

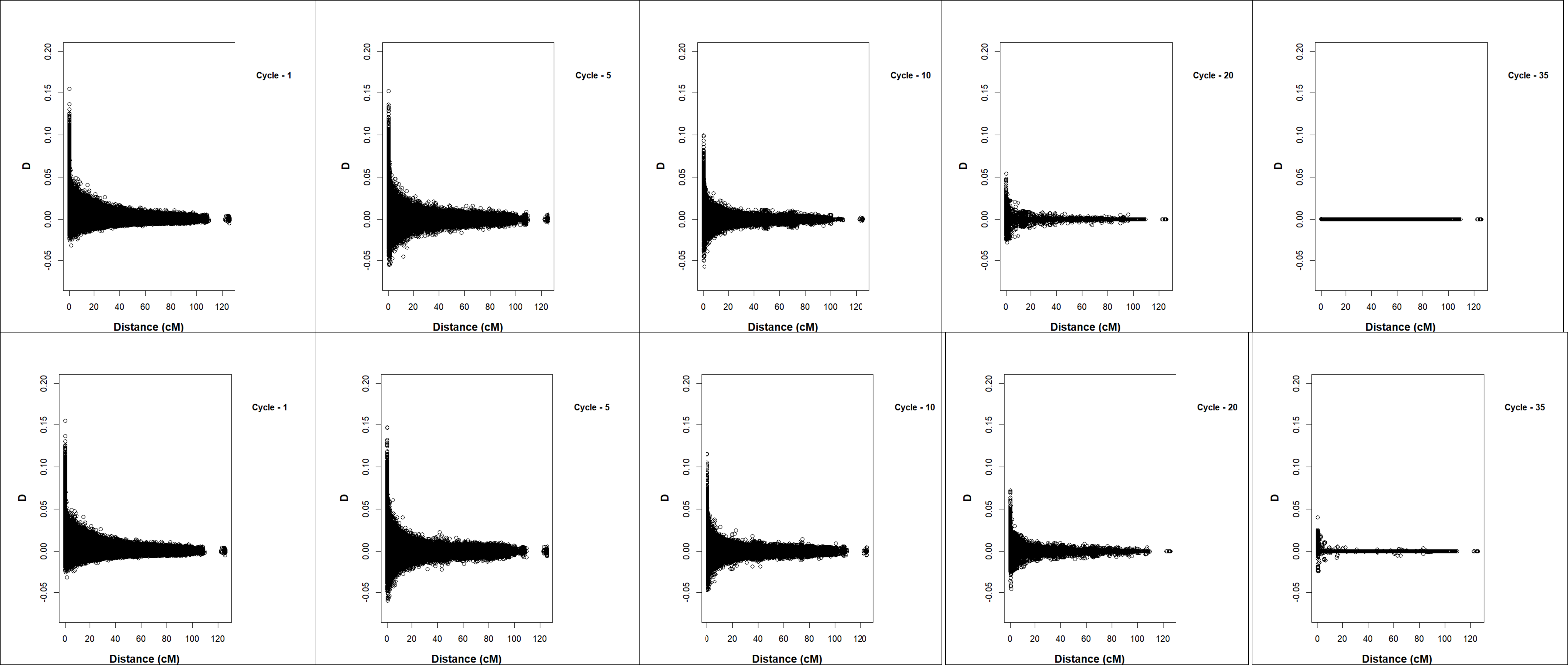

**Figure S21 Marker-marker LD decay with Distance between Marker Pairs within a Chromosome** **in cM for Ridge Regression**. Marker-marker LD decay with distance between marker pairs with in a chromosome in cM. The figures represent distribution of LD estimated with ‘D’ statistic in selection cycles 1, 5, 10, 20 and 35 from left to right. ‘D’ is plotted on y-axis and distance between marker pairs within a chromosome in cM. Top panel represents LD for populations under GS with Ridge Regression without updating and bottom panel represents GS with RR updated with training set from14 prior cycles. All treatment combinations have 400 simulated QTL responsible for 70% of phenotypic variability in the initial population and top 10% selected fraction.

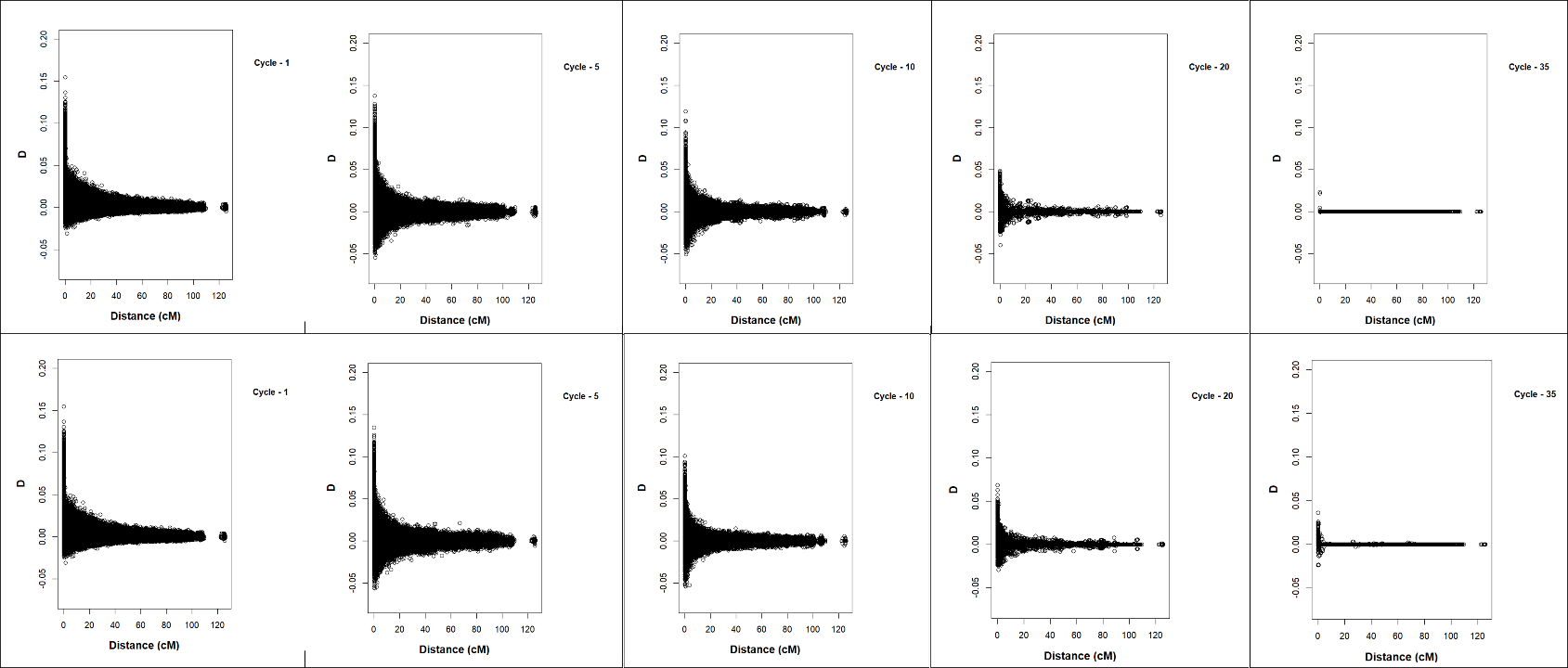

**Figure S22** **Marker-marker LD decay with Distance between Marker Pairs within a Chromosome** **in cM for BayesB**: Marker-marker LD decay with distance between marker pairs with in a chromosome in cM. The figures represent distribution of LD estimated with ‘D’ statistic in selection cycles 1, 5, 10, 20 and 35 from left to right. ‘D’ is plotted on y-axis and distance between marker pairs within a chromosome in cM. Top panel represents LD for populations under GS with BayesB without updating and bottom panel represents GS with BayesB updated with training set from14 prior cycles. All treatment combinations have 400 simulated QTL responsible for 70% of phenotypic variability in the initial population and top 10% selected fraction.

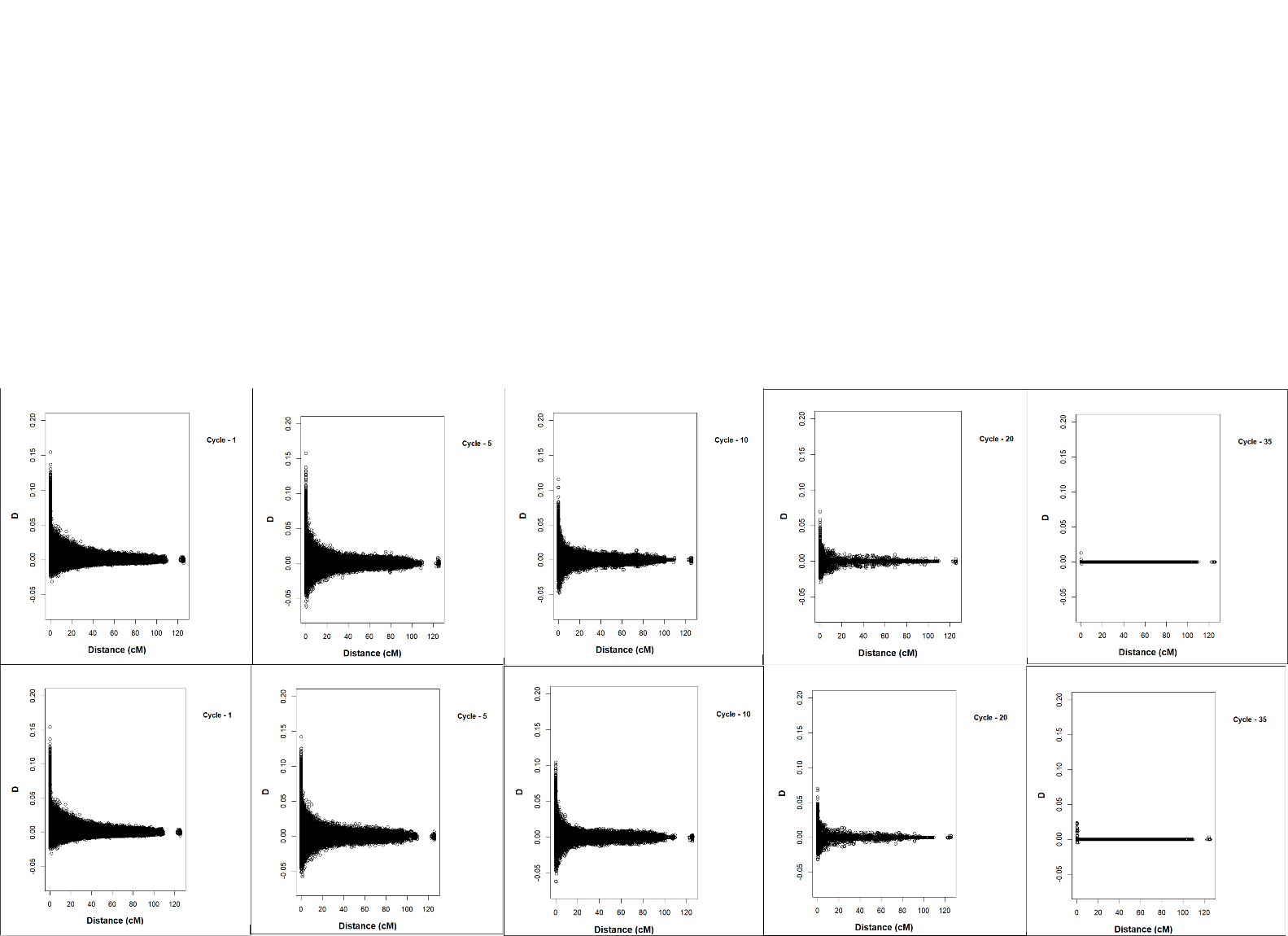

**Figure S23 Marker-marker LD decay with Distance between Marker Pairs within a Chromosome** **in cM for Bayes LASSO:** Marker-marker LD decay with distance between marker pairs with in a chromosome in cM. The figures represent distribution of LD estimated with ‘D’ statistic in selection cycles 1, 5, 10, 20 and 35 from left to right. ‘D’ is plotted on y-axis and distance between marker pairs within a chromosome in cM. Top panel represents LD for population under GS with Bayes LASSO without updating and bottom panel represents GS with Bayes LASSO updated with training data from 14 prior cycles. All treatment combinations have 400 simulated QTL responsible for 70% of phenotypic variability in the initial population and top 10% selected fraction.

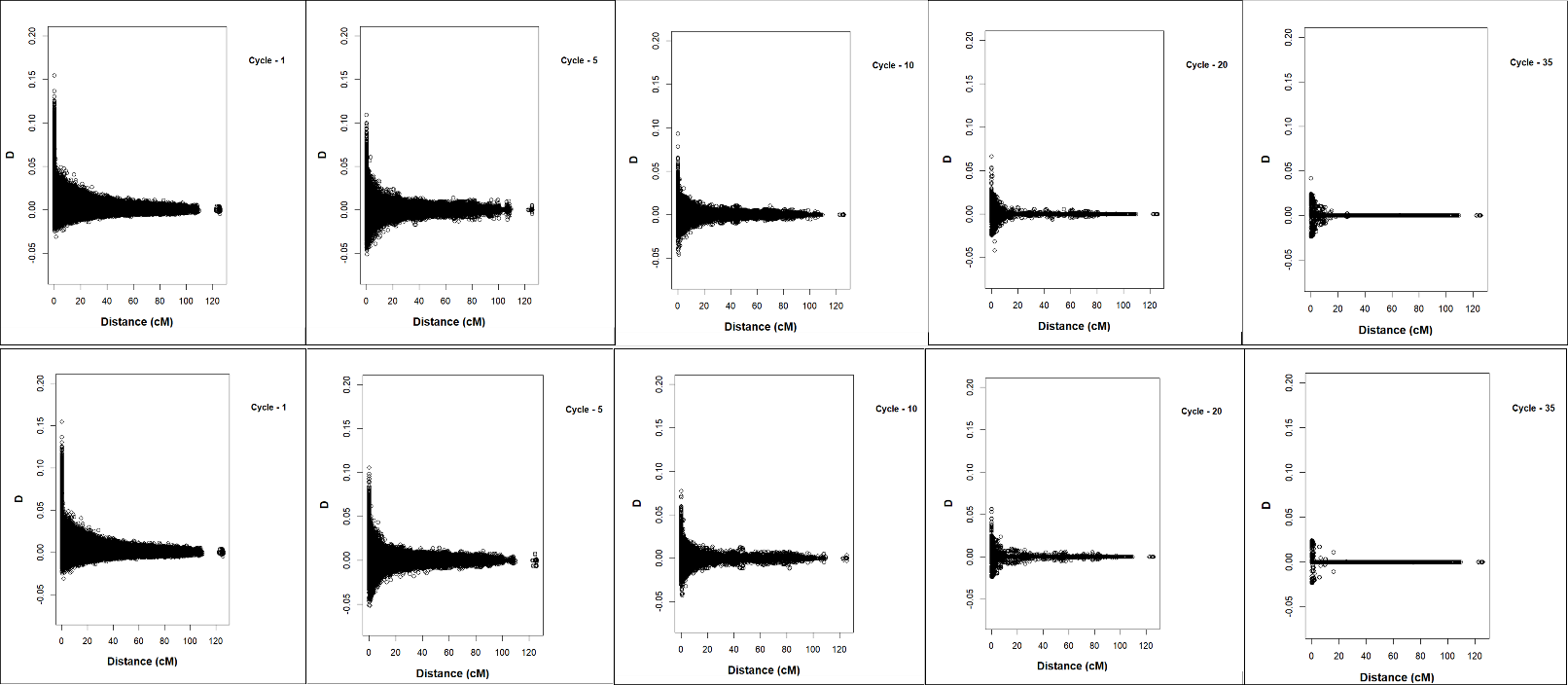

**Figure S24** **Marker-marker LD decay with Distance between Marker Pairs within a Chromosome** **in cM for SVMRBF:** The figures represent distribution of LD estimated with ‘D’ statistic in selection cycles 1, 5, 10, 20 and 35 from left to right. ‘D’ is plotted on y-axis and distance between marker pairs within a chromosome in cM. Top panel represents GS with SVMRBF without updating and bottom panel represents SVMRBF GS updated with training set from14 prior cycles. All treatment combinations have 400 simulated QTL responsible for 70% of phenotypic variability in the initial population and top 10% selected fraction.

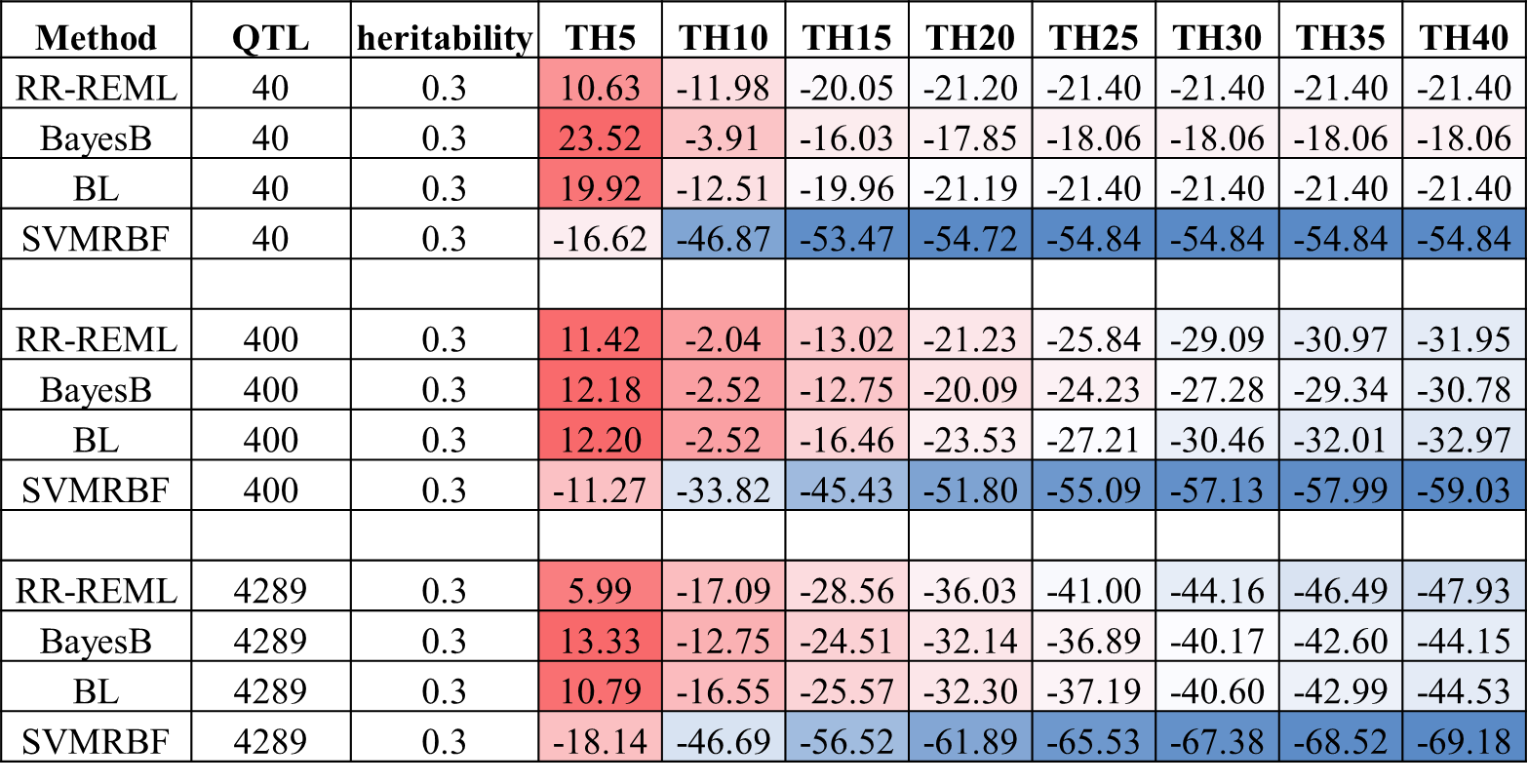

**Figure S25 Heat Map for Percent Gain in Rs Relative to PS for 0.3 H for GP Models without Updating** Heat map representing percent gain in response in recurrent GS for GS Models with no updating in cycles one to forty in steps of five cycles. Percent gain in responses are provided for genetic architectures consisting of 40 simulated QTL, 400 simulated QTL and 4289 simulated QTL responsible for 30% of phenotypic variability in the initial population (h2 -0.3). Blue to red shaded cells represent increasing gain in response relative to PS. PS – Phenotypic Selection, RR-REML- Ridge Regression with Restricted Maximum Likelihood, BL – Bayes LASSO, and SVMRBF- Support Vector Machine with Radial Basis Kernel.

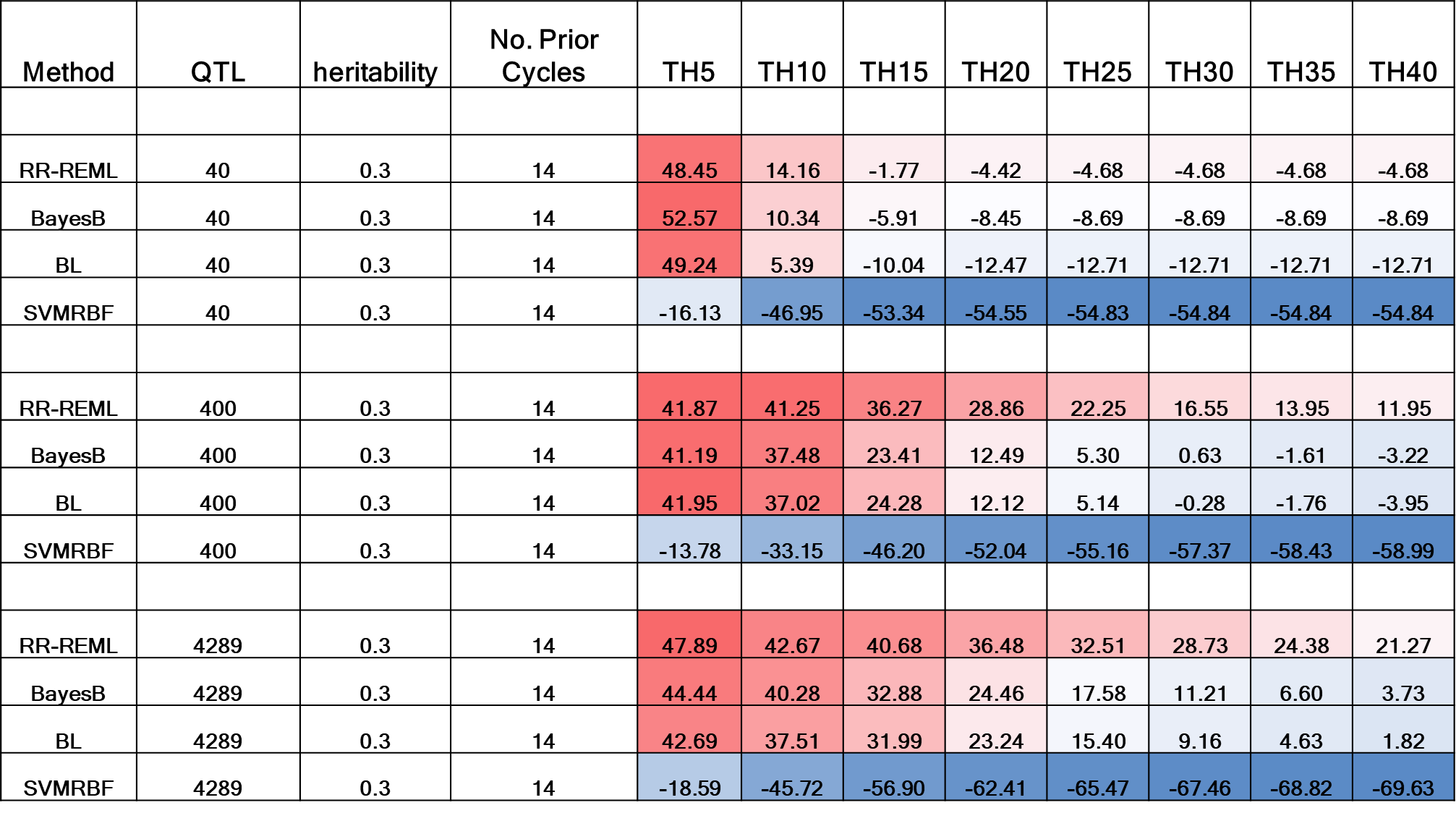

**Figure S26 Heat Map for Percent Gain in Responses Relative to PS for 0.3 H for GP Models with Updating** Heat map table representing percent gain in response with respect to response in PS in Recurrent GS for GP models updated with training data from 14 prior cycles in cycles one to forty in steps of five cycles. Percent gain in responses are provided for genetic architectures consisting of 40 simulated QTL, 400 simulated QTL and 4289 simulated QTL responsible for 30% of phenotypic variability in the initial population. Blue to red shaded cells represent increasing gain in response relative to PS. PS – Phenotypic Selection, RR-REML- Ridge Regression with Restricted Maximum Likelihood, BL – Bayes LASSO, and SVMRBF- Support Vector Machine with Radial Basis Kernel.

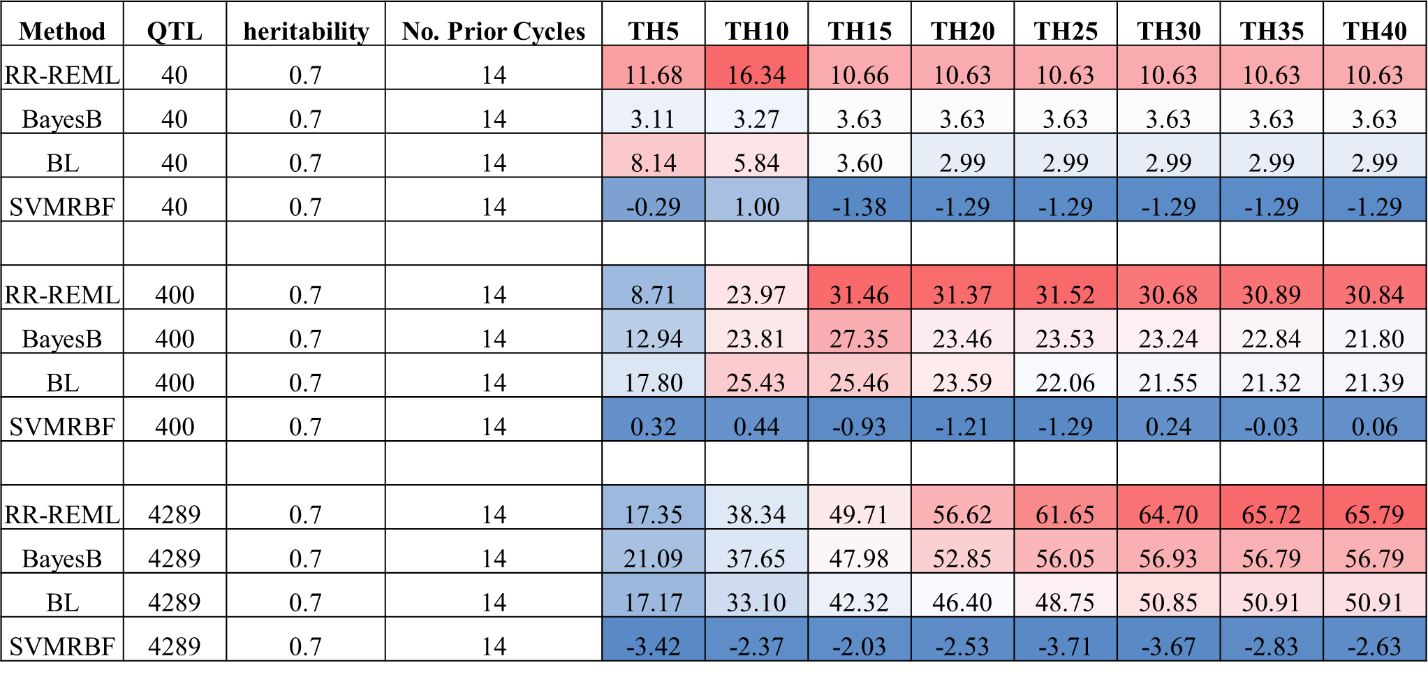

**Figure S27 Heat Map for Percent Gain in Responses Relative to GS without Updating for 0.7 H** Heat map representing percent gain in response with respect to same type of GP model without updating in Recurrent GS for GP models updated with training data from 14 prior cycles in cycles one to forty in steps of five cycles. Percent gain in responses are provided for genetic architectures consisting of 40 simulated QTL, 400 simulated QTL and 4289 simulated QTL responsible for 70% of phenotypic variability in the initial population (h2 -0.7). Blue to red shaded cells represent increasing gain in response relative to GS without updating. PS – Phenotypic Selection, RR-REML- Ridge Regression with Restricted Maximum Likelihood, BL – Bayes LASSO, and SVMRBF- Support Vector Machine with Radial Basis Kernel.

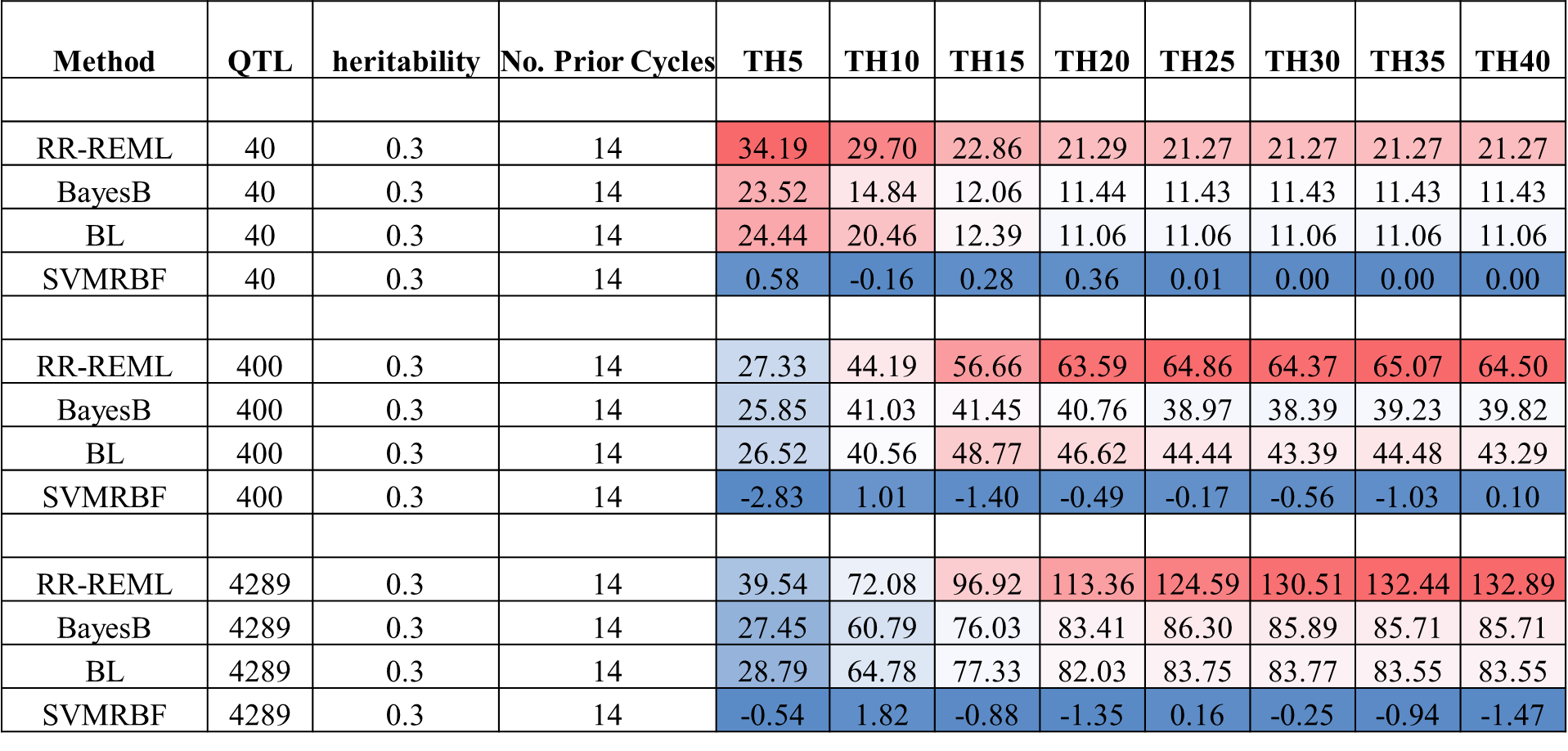

**Figure S28 Heat Map for Percent Gain in Responses Relative to GS without Updating for 0.3 H** Heat map table representing percent gain in response with respect to same type of GP model without updating in Recurrent GS for GP models updated with training data from 14 prior cycles in cycles one to forty in steps of five cycles. Percent gain in responses are provided for genetic architectures consisting of 40 simulated QTL, 400 simulated QTL and 4289 simulated QTL responsible for 30% of phenotypic variability in the initial population (h2 -0.3). Blue to red shaded cells represent increasing gain in response relative to GS without updating. PS – Phenotypic Selection, RR-REML- Ridge Regression with Restricted Maximum Likelihood, BL – Bayes LASSO, and SVMRBF- Support Vector Machine with Radial Basis Kernel.

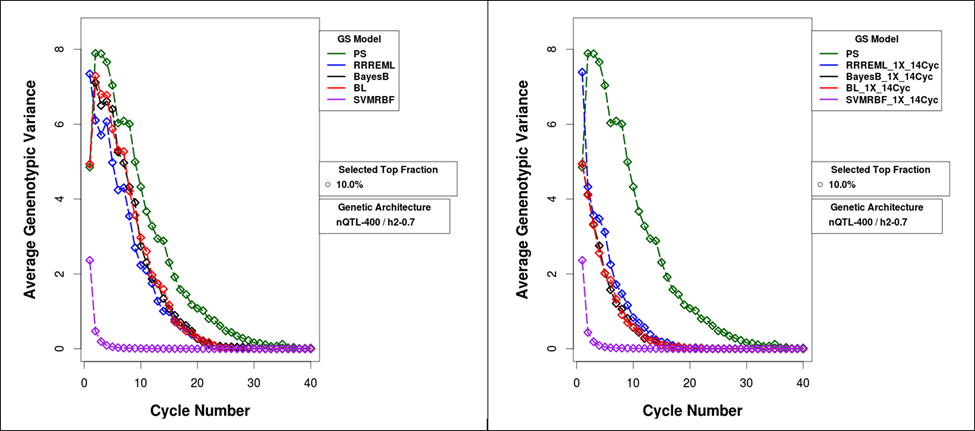

**Figure S29** **Average Genotypic Variance** **in Selected Populations for Comparison of GS methods with and without Updating for 400 QTL, 0.7 H and Top 10% Selected Fraction:** Average genotypic variance for 40 cycles of recurrent selection of 10% of RILs using five selection methods without updated training sets (left panel) and with updated training sets (right panel). Training sets consisted of genotypic and phenotypic data from up to14 prior cycles of recurrent selection. Simulated phenotypic values of the RILs consisted of 400 simulated QTL responsible for 70% of phenotypic variability in the initial population. PS – Phenotypic Selection, RR-REML- Ridge Regression with Restricted Maximum Likelihood, BL – Bayes LASSO, and SVMRBF- Support Vector Machine with Radial Basis Kernel.

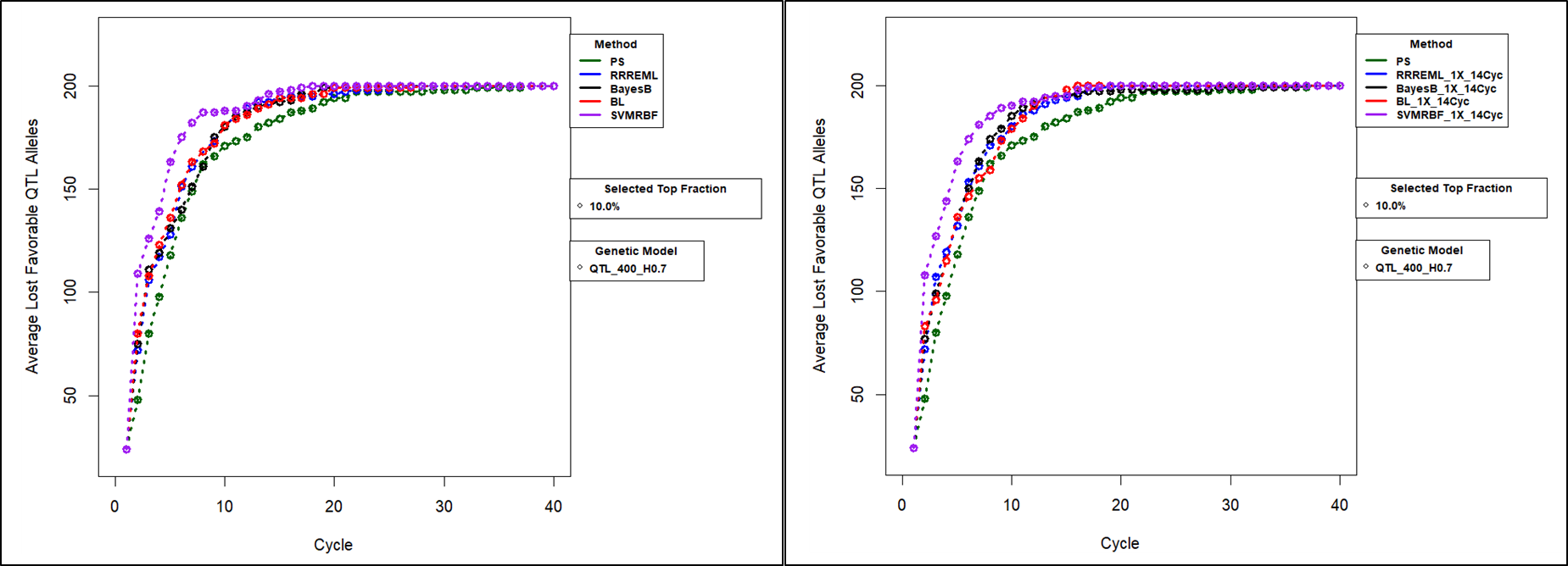

**Figure S30:** **Number of Lost Favorable Alleles** **for Comparison of GS methods with and without Updating for 400 QTL, 0.7 H and Top 10% Selected Fraction:** Number of favorable alleles that are cumulatively discarded across 40 cycles of recurrent selection of 10% of the RILs created every generation using five selection methods without updated training sets (left panel) and with updated training sets (right panel). Training sets consisted of genotypic and phenotypic data from up to14 prior cycles of recurrent selection. Simulated phenotypic values of the RILs consisted of 400 simulated QTL responsible for 70% of phenotypic variability in the initial population. PS – Phenotypic Selection, RR-REML- Ridge Regression with Restricted Maximum Likelihood, BL – Bayes LASSO, and SVMRBF- Support Vector Machine with Radial Basis Kernel.

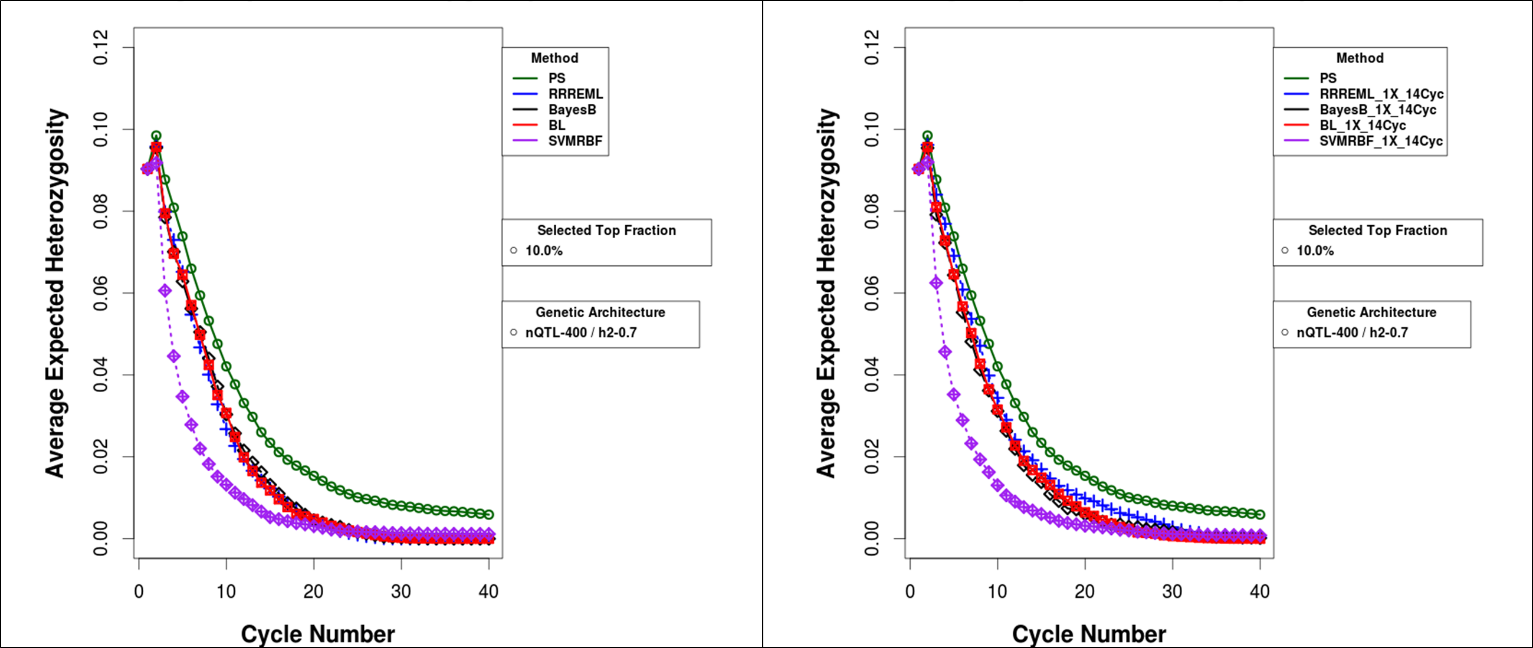

**Figure S31 Average Expected Heterozygosity** **for Comparison of GS methods with and without Updating for 400 QTL, 0.7 H and Top 10% Selected Fraction:** Average Expected Heterozygosity in recurrent GS without GP model updating (left panel) and GP models updated every cycle with training data from upto14 prior cycles (right panel). All treatment combinations have 400 simulated QTL responsible for 70% of phenotypic variability in the initial population and 10% top selected fraction. PS – Phenotypic Selection, RR-REML- Ridge Regression with Restricted Maximum Likelihood, BL – Bayes LASSO, and SVMRBF- Support Vector Machine with Radial Basis Kernel.

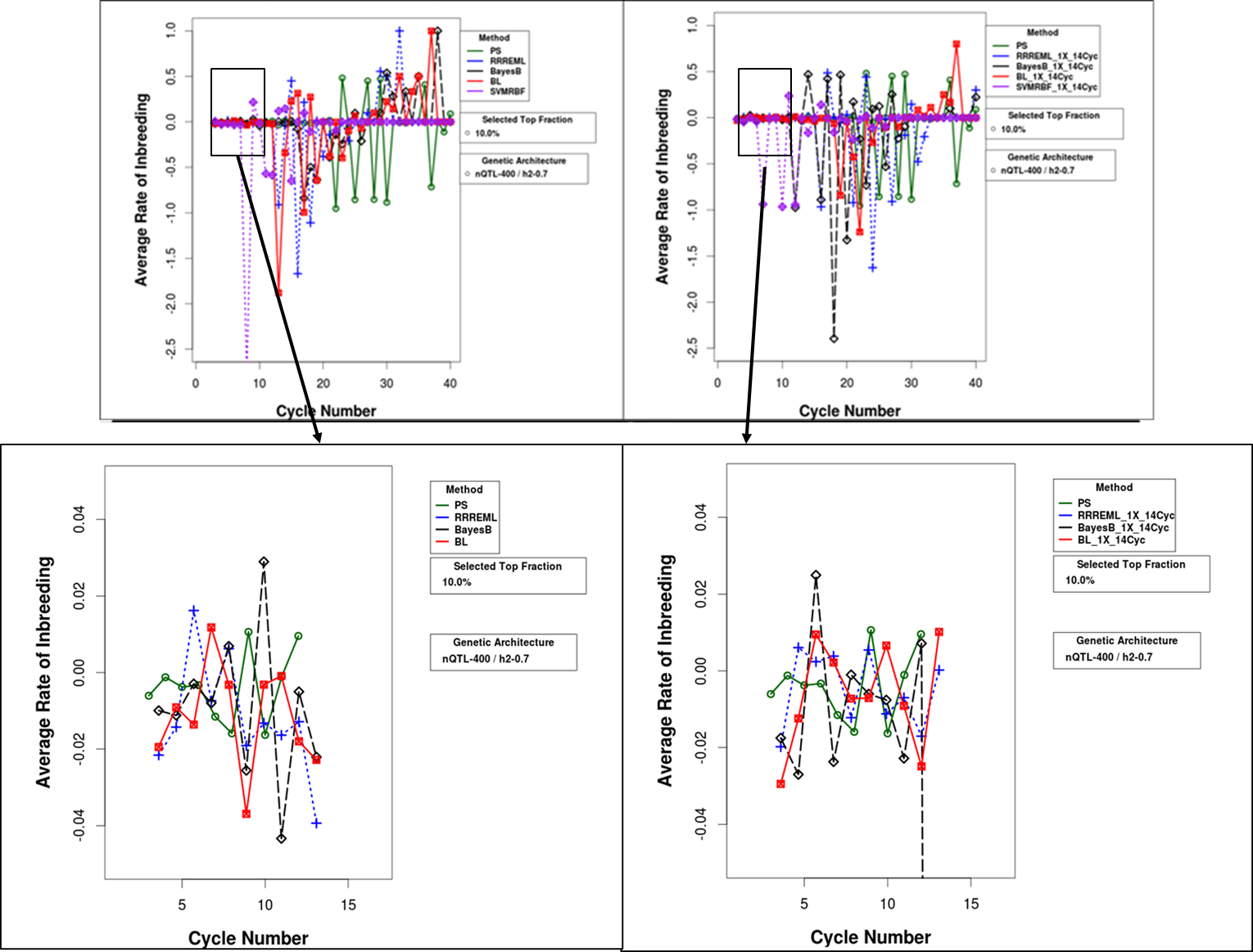

**Figure S32 Average Rate of Inbreeding** **for Comparison of GS methods with and without Updating for 400 QTL, 0.7 H and Top 10% Selected Fraction:** Average rate of inbreeding in recurrent GS without GP model updating (left panel) and GP models updated every cycle with training data from upto14 prior cycles. Inbreeding co-efficient is estimated as the harmonic mean of all individuals in the population for 40 cycles (top panel). Inset plot (bottom panel) shows magnified region from 3-12 cycles in average rate of inbreeding for PS, RR, BayesB, and BL GS methods. All treatment combinations have 400 simulated QTL responsible for 70% of phenotypic variability in the initial population and 10% top selected fraction. PS – Phenotypic Selection, RR-REML- Ridge Regression with Restricted Maximum Likelihood, BL – Bayes LASSO, and SVMRBF- Support Vector Machine with Radial Basis Kernel.

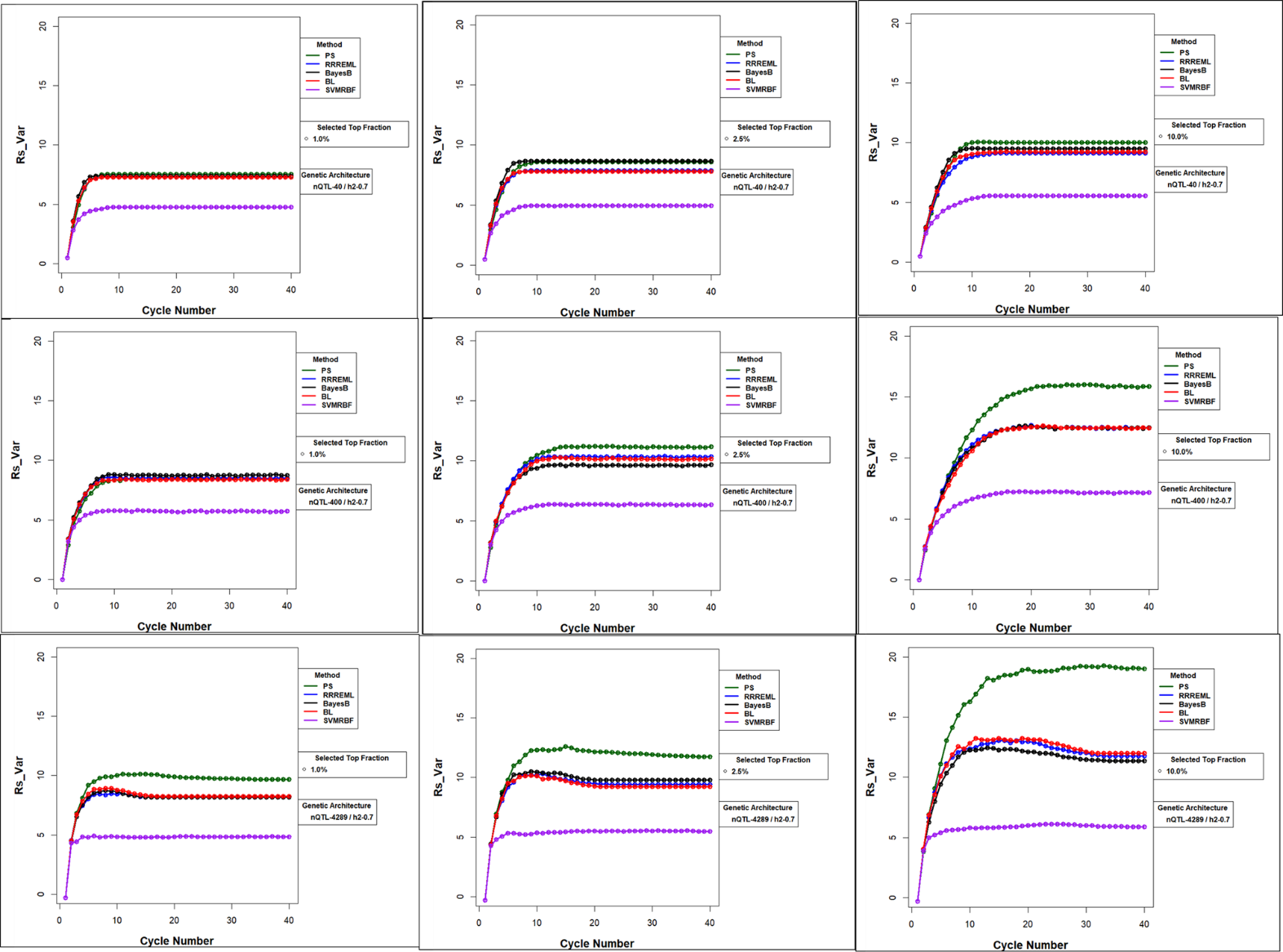

**Figure S33 Response Standardized to Change in Genotypic Variance (‘Rs_Var’)** **for Comparison of GS methods Without Updating for 0.7 H:** Response standardized to change in genotypic variance (‘Rs_Var’) across three levels of QTL and three levels of selection intensity for PS and four GS Methods that are not updated. ‘Rs_Var’ is plotted on the y-axis and selection cycle is plotted on x-axis. Response standardized to change in genotypic variance captures the difference in rate at which response increases with loss of genetic variance. Relaxed selection intensity results in greater limits (comparison of plots from left to right in the three panels). Greater number of QTL also result in greater limits (comparison of top, middle and bottom panels across three selection intensities).

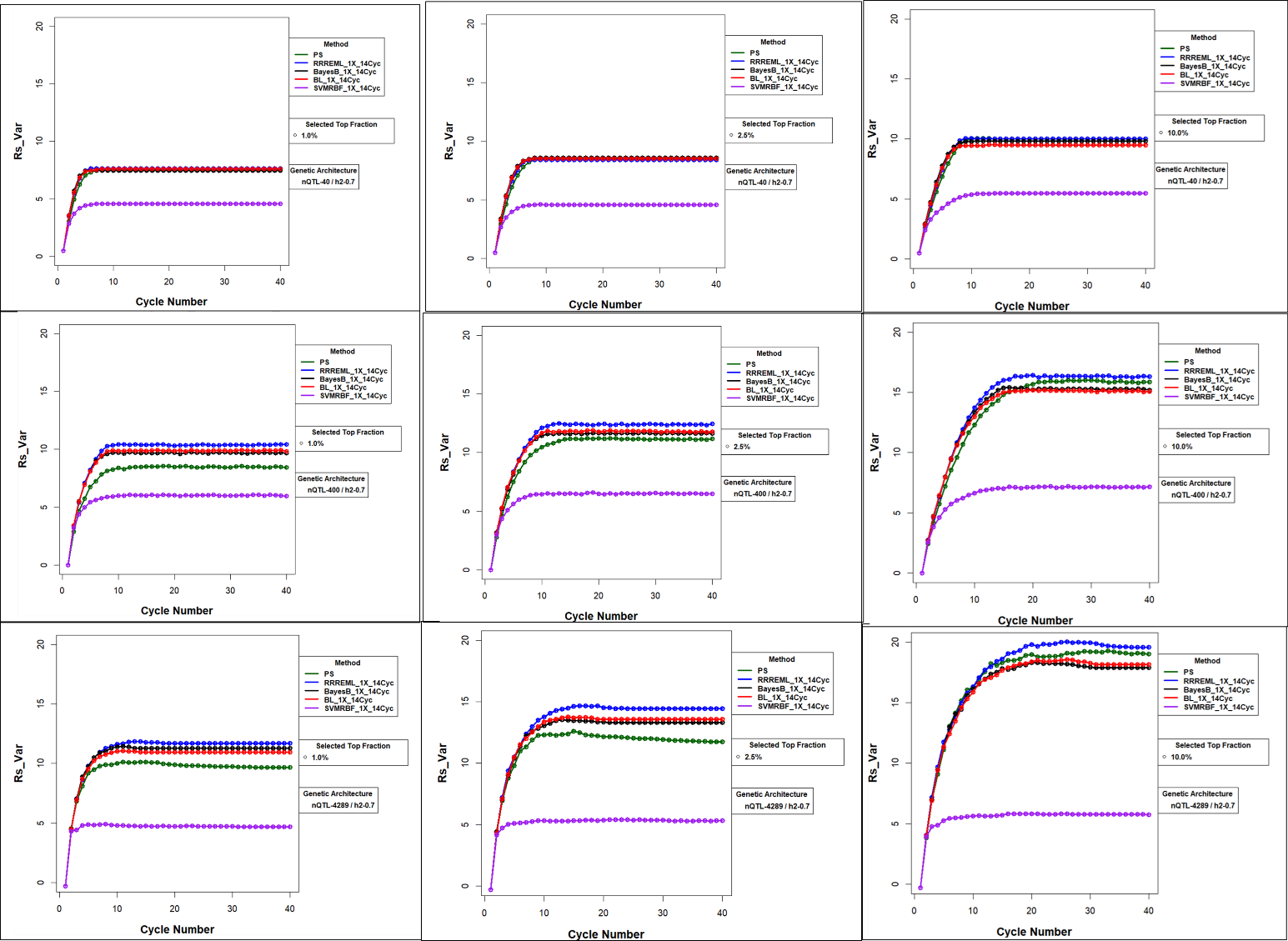

**Figure S34** **Response Standardized to Change in Genotypic Variance (‘Rs_Var’)** **for Comparison of GS methods with updating for 0.7 H:** Response standardized to change in genotypic variance (‘Rs_Var’) across three levels of QTL and three levels of selection intensity for PS and four GS Methods that are updated with training data from up to14 prior cycles. ‘Rs_Var’ is plotted on the y-axis and selection cycle is plotted on x-axis. Response standardized to change in genotypic variance captures the difference in rate at which response increases with loss of genetic variance. Relaxed selection intensity results in greater limits (comparison of plots from left to right in the three panels). Larger number of QTL also result in greater limits (comparison of top, middle and bottom panels across three selection intensities).

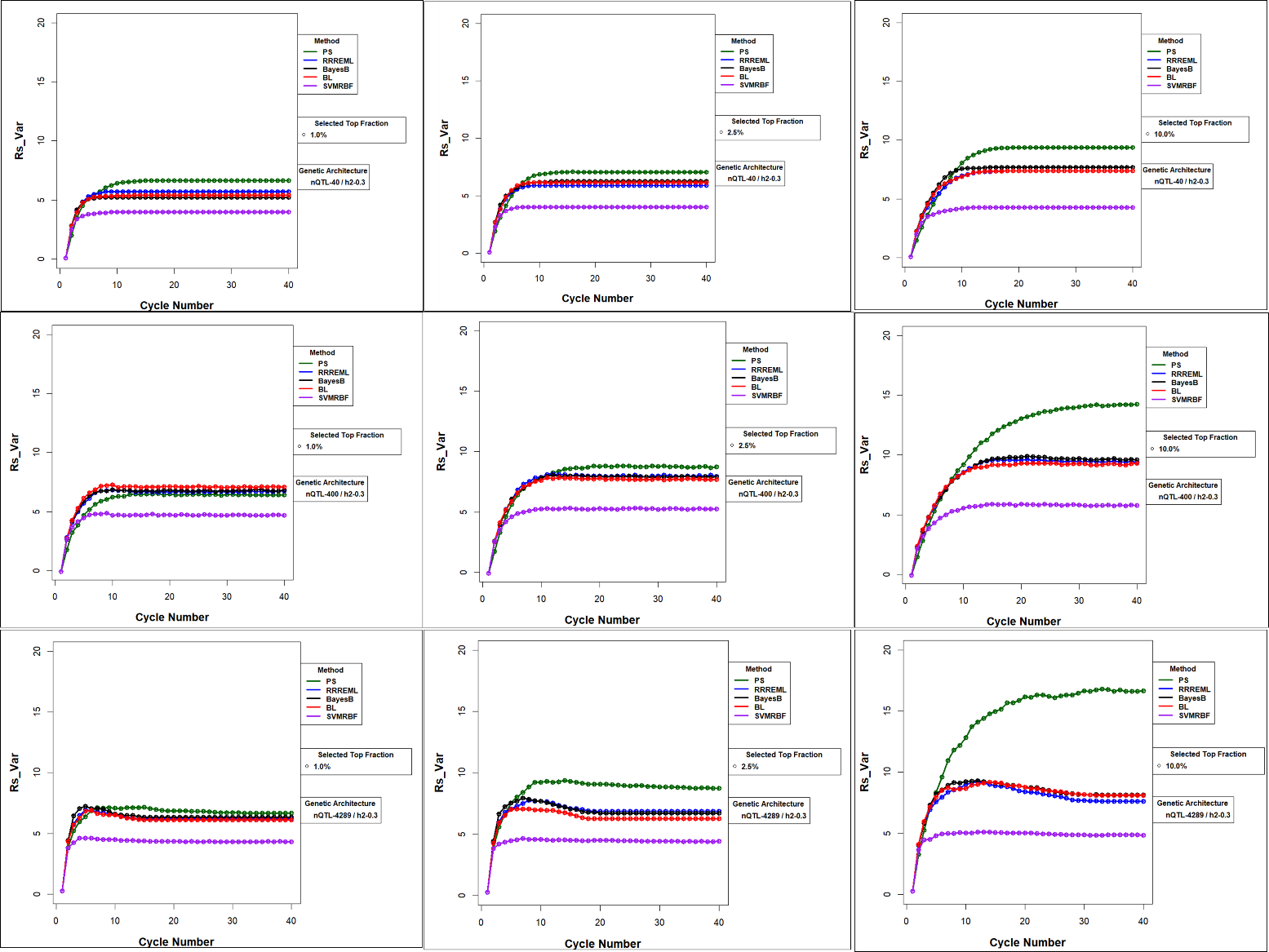

**Figure S35 Response Standardized to Change in Genotypic Variance (‘Rs_Var’)** **for Comparison of GS Methods without Updating for 0.3 H:** Response standardized to change in genotypic variance (‘Rs_Var’) across three levels of QTL and three levels of selection intensity for PS and four GS Methods that are not updated. ‘Rs_Var’ is plotted on the y-axis and selection cycle is plotted on x-axis. Response standardized to change in genotypic variance captures the difference in rate at which response increases with loss of genetic variance. Relaxed selection intensity results in greater limits (comparison of plots from left to right in the three panels). Larger number of QTL also result in greater limits (comparison of top, middle and bottom panels across three selection intensities).

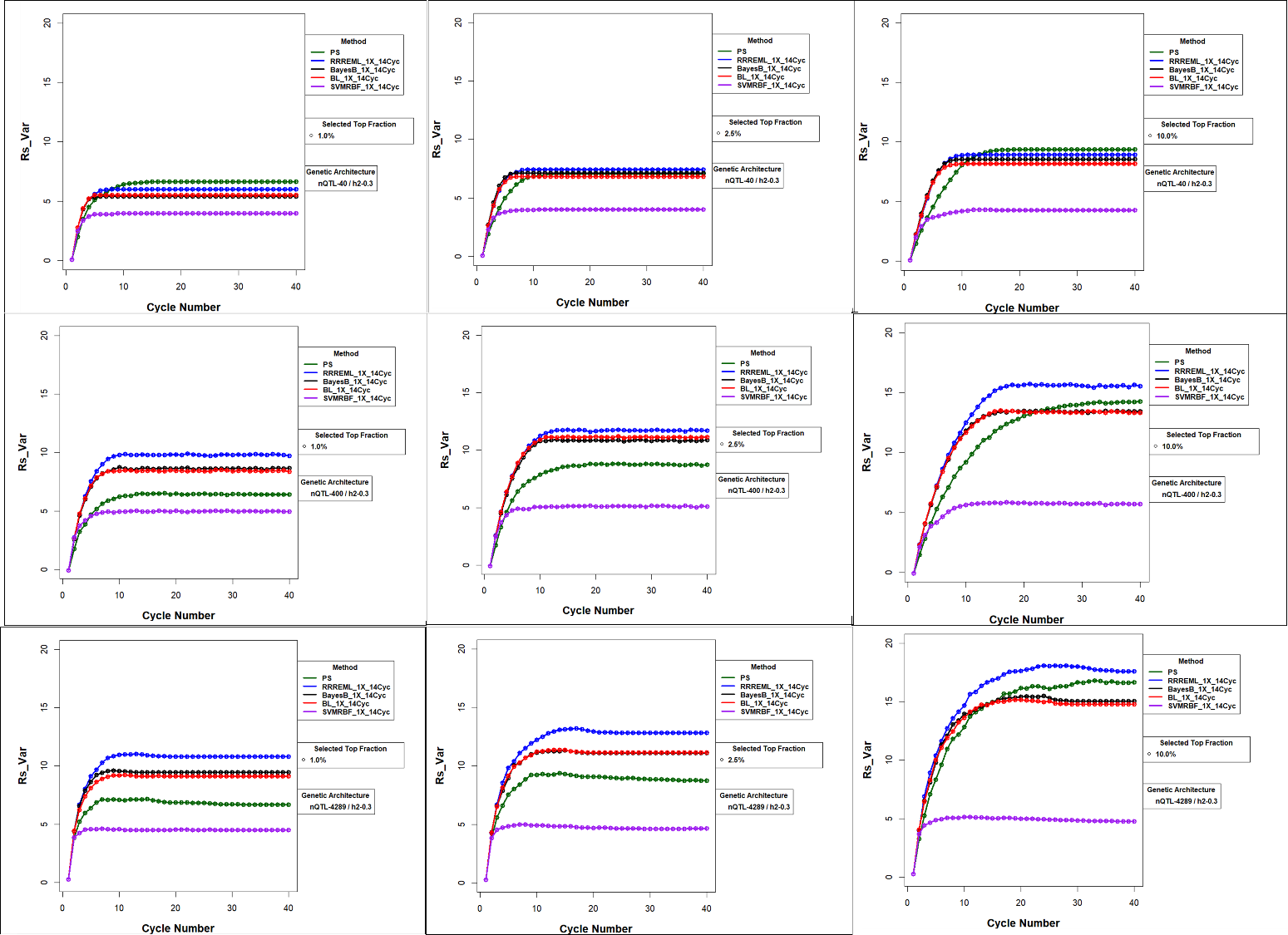

**Figure S36 Response Standardized to Change in Genotypic Variance (‘Rs_Var’)** **for Comparison of GS methods with updating for 0.3 H:** Response standardized to change in genotypic variance (‘Rs_Var’) across three levels of QTL and three levels of selection intensity for PS and four GS Methods that are not updated. ‘Rs_Var’ is plotted on the y-axis and selection cycle is plotted on x-axis. GP models are updated with training data from up to 14 prior cycles. Response standardized to change in genotypic variance captures the difference in rate at which response increases with loss of genetic variance. Relaxed selection intensity results in greater limits (comparison of plots from left to right in the three panels). Greater number of QTL also result in greater limits (comparison of top, middle and bottom panels across three selection intensities).

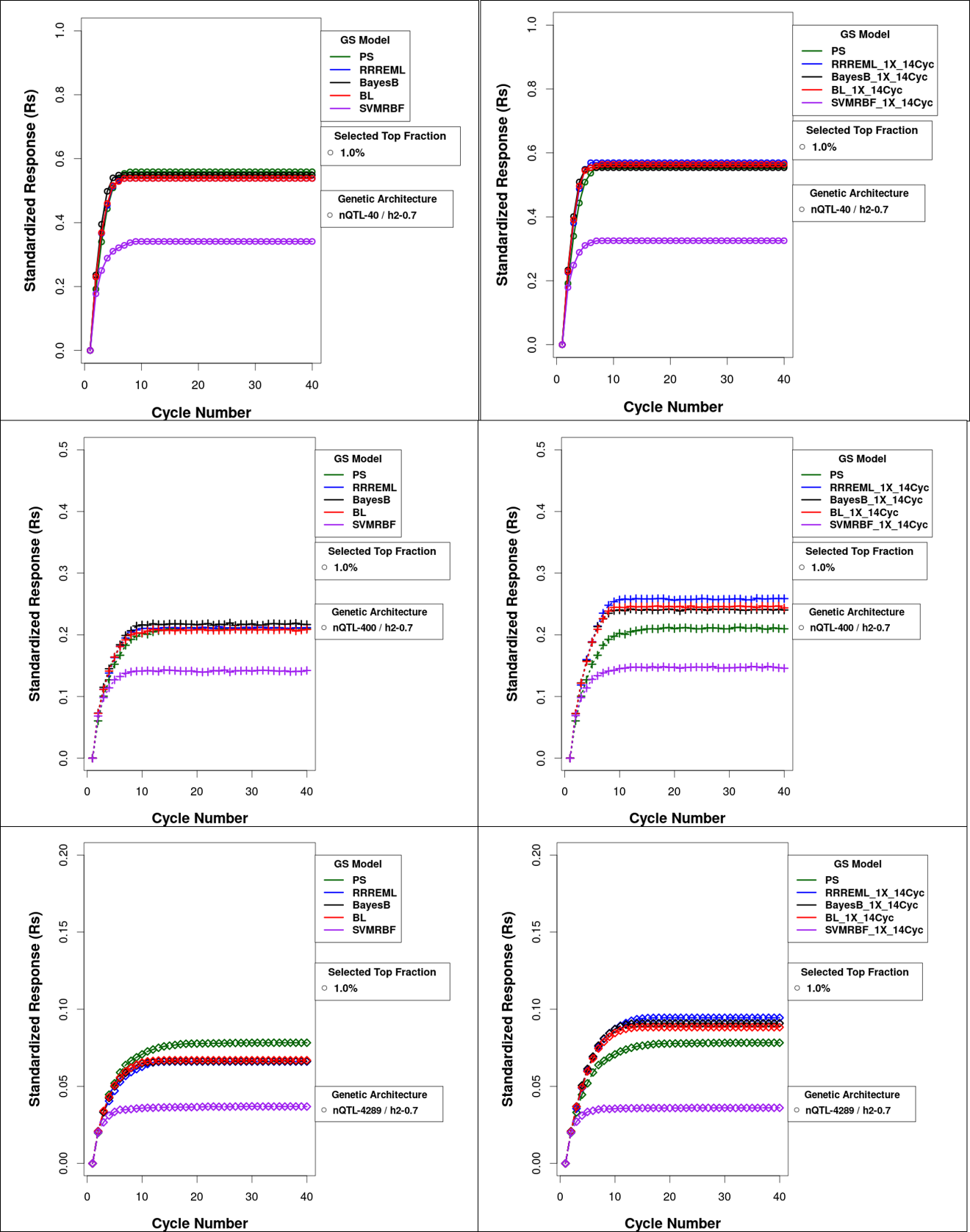

**Figure S37** **Standardized Responses for Comparison of GS methods with and without Updating for 0.7 H and Top 1% Selected Fraction:** Forty cycles of response to selection of the top 1% of soybean RILs derived from SoyNAM founders. Responses are plotted by selection methods as genotypic values standardized to maximum possible value without model updating (left panels) and with model updating (right panels) using 14 prior cycles as training sets for the four GP models. Phenotypic selection (PS) is not updated and hence does not change between the left and right panels. Top panels consist of responses for genetic architectures consisting of 40 simulated QTL. Middle panels consist of responses for genetic architectures consisting of 400 simulated QTL and the bottom panels consist of responses for genetic architectures consisting of 4289 simulated QTL. All 40, 400 and 4289 are responsible for 70% of phenotypic variability in the initial population. PS – Phenotypic Selection, RR-REML- Ridge Regression with Restricted Maximum Likelihood, BL – Bayes LASSO, and SVMRBF- Support Vector Machine with Radial Basis Kernel.

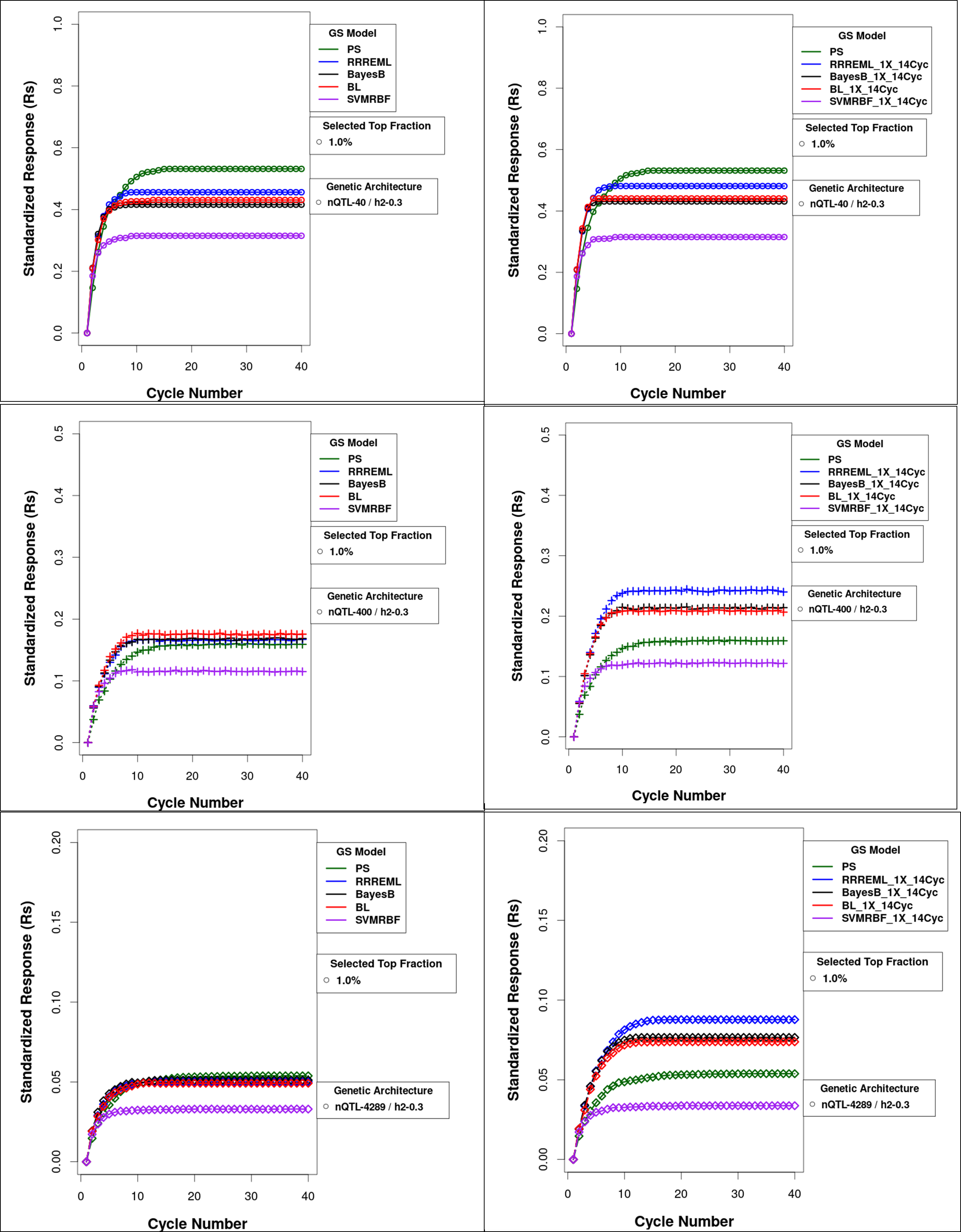

**Figure S38 Standardized Responses for Comparison of GS methods with and without Updating for 0.3 H and Top 1% Selected Fraction:** Forty cycles of response to selection of the top 1% of soybean RILs derived from SoyNAM founders. Responses are plotted by selection method as genotypic values standardized to maximum possible value without (left panels) and with (right panels) model updating using prior cycles as training sets for the four GP models. Phenotypic selection (PS) is not updated and hence does not change between the left and right panels. Top panels consist of responses for genetic architectures consisting of 40 simulated QTL. Middle panels consist of responses for genetic architectures consisting of 400 simulated QTL and the bottom panels consist of responses for genetic architectures consisting of 4289 simulated QTL. All 40, 400 and 4289 are responsible for 30% of phenotypic variability in the initial population. PS – Phenotypic Selection, RR-REML- Ridge Regression with Restricted Maximum Likelihood, BL – Bayes LASSO, and SVMRBF- Support Vector Machine with Radial Basis Kernel.

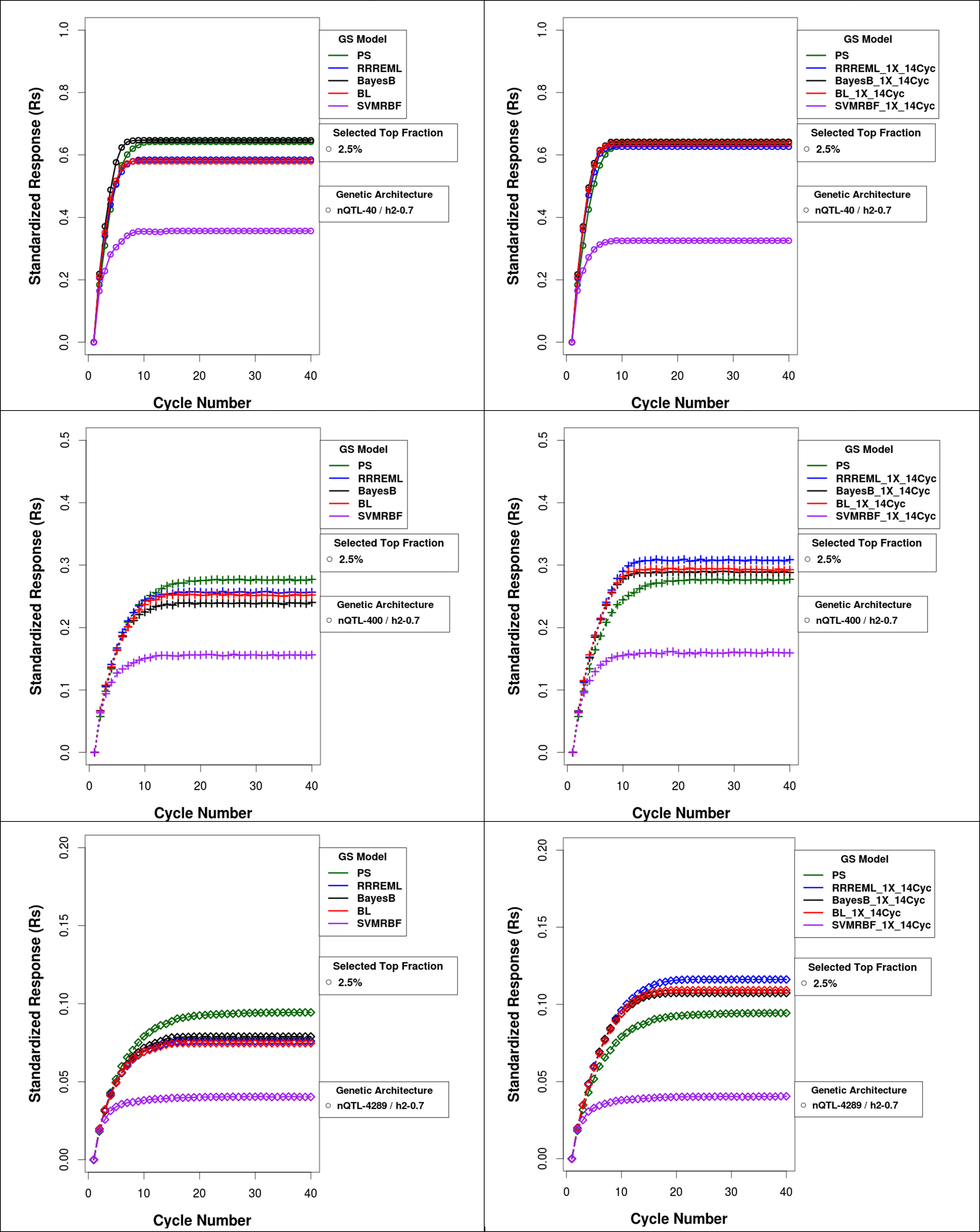

**Figure S39** **Standardized Responses for Comparison of GS methods with and without Updating for 0.7 H and Top 2.5% Selected Fraction:** Forty cycles of response to selection of the top 2.5% of soybean RILs derived from SoyNAM founders. Responses are plotted by selection methods as genotypic values standardized to maximum possible value without (left panels) and with (right panels) model updating using prior cycles as training sets for the four GP models. Phenotypic selection (PS) is not updated and hence does not change between the left and right panels. Top panels consist of responses for genetic architectures consisting of 40 simulated QTL. Middle panels consist of responses for genetic architectures consisting of 400 simulated QTL and the bottom panels consist of responses for genetic architectures consisting of 4289 simulated QTL. All 40, 400 and 4289 simulated QTL are responsible for 70% of phenotypic variability in the initial population.PS – Phenotypic Selection, RR-REML- Ridge Regression with Restricted Maximum Likelihood, BL – Bayes LASSO, and SVMRBF- Support Vector Machine with Radial Basis Kernel.

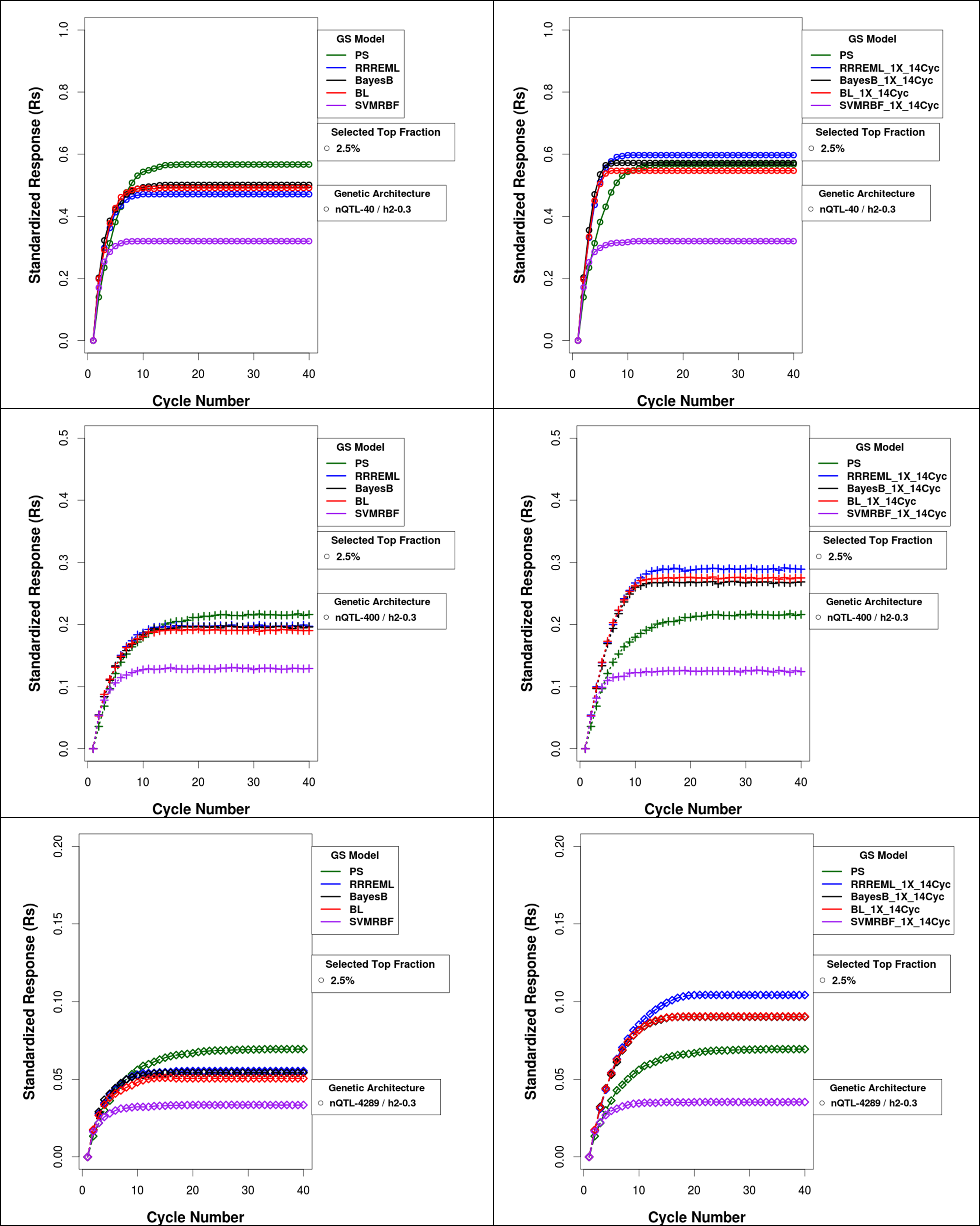

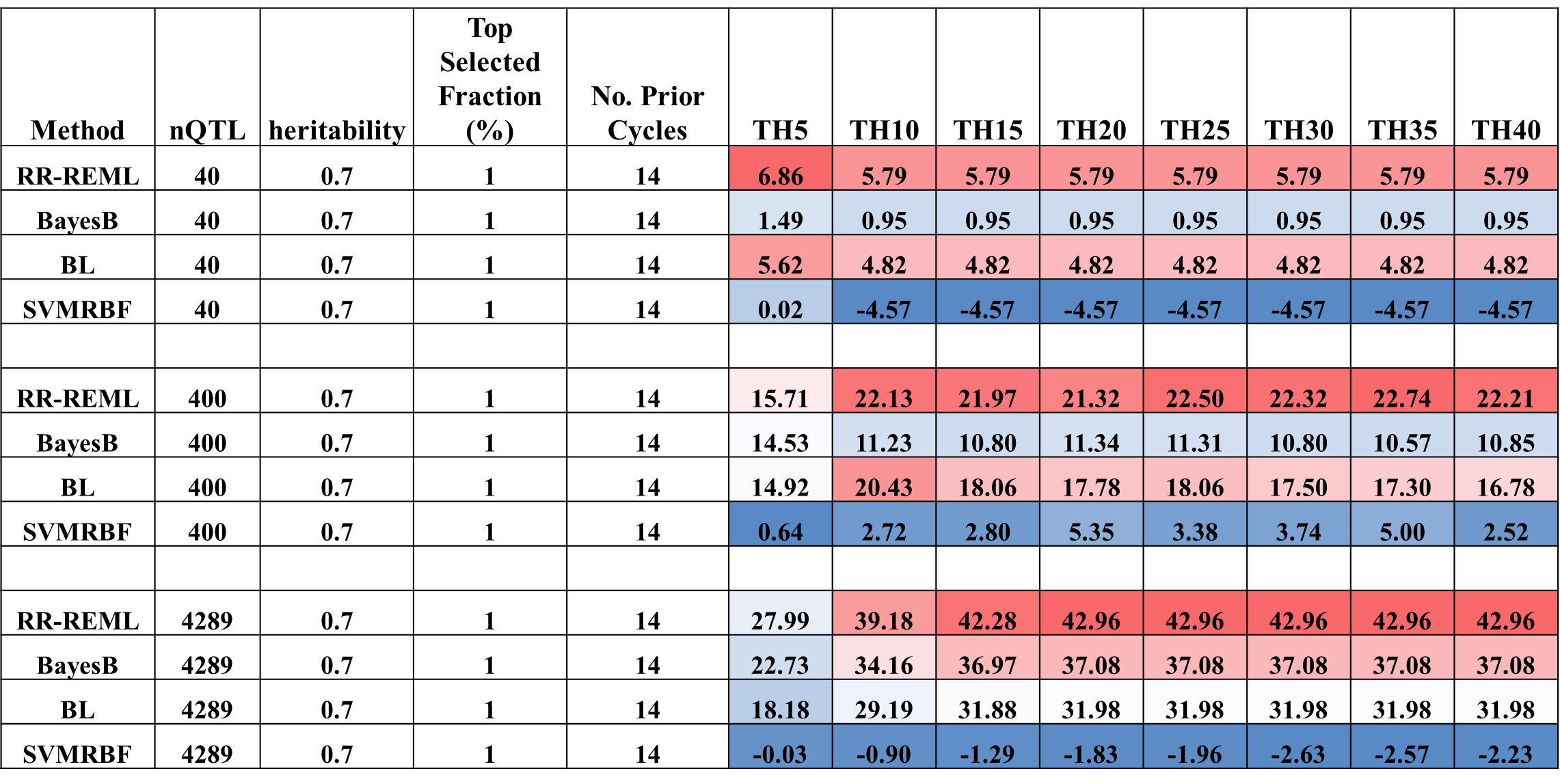

**Figure S40** **Standardized Responses for Comparison of GS methods with and without Updating for 0.3 H and Top 2.5% Selected Fraction:** Forty cycles of response to selection of the top 2.5% of soybean RILs derived from SoyNAM founders. Responses are plotted by selection method as genotypic values standardized to maximum possible value without model updating (left panels) and with model updating (right panels) using prior cycles as training sets for the four GP models. Phenotypic selection (PS) is not updated and hence does not change between the left and right panels. Top panels consist of responses for genetic architectures consisting of 40 simulated QTL. Middle panels consist of responses for genetic architectures consisting of 400 simulated QTL and the bottom panels consist of responses for genetic architectures consisting of 4289 simulated QTL. All 40, 400 and 4289 are responsible for 30% of phenotypic variability in the initial population. PS – Phenotypic Selection, RR-REML- Ridge Regression with Restricted Maximum Likelihood, BL – Bayes LASSO, and SVMRBF- Support Vector Machine with Radial Basis Kernel.

**Figure S41 Heat Map for Percent Gain in Rs Relative to GS without Updating for 0.7 H and Top 1% Selected Fraction** Heat map table representing percent gain in response with respect to same type of GP model without updating in Recurrent GS for GP models updated with training data from 14 prior cycles in cycles one to forty in steps of five cycles for 40, 400 and 4289 simulated QTL responsible for 70% of phenotypic variability in the initial population. Blue to red shaded cells represent increasing gain in response relative to GS without updating. PS – Phenotypic Selection, RR-REML- Ridge Regression with Restricted Maximum Likelihood, BL – Bayes LASSO, and SVMRBF- Support Vector Machine with Radial Basis Kernel.

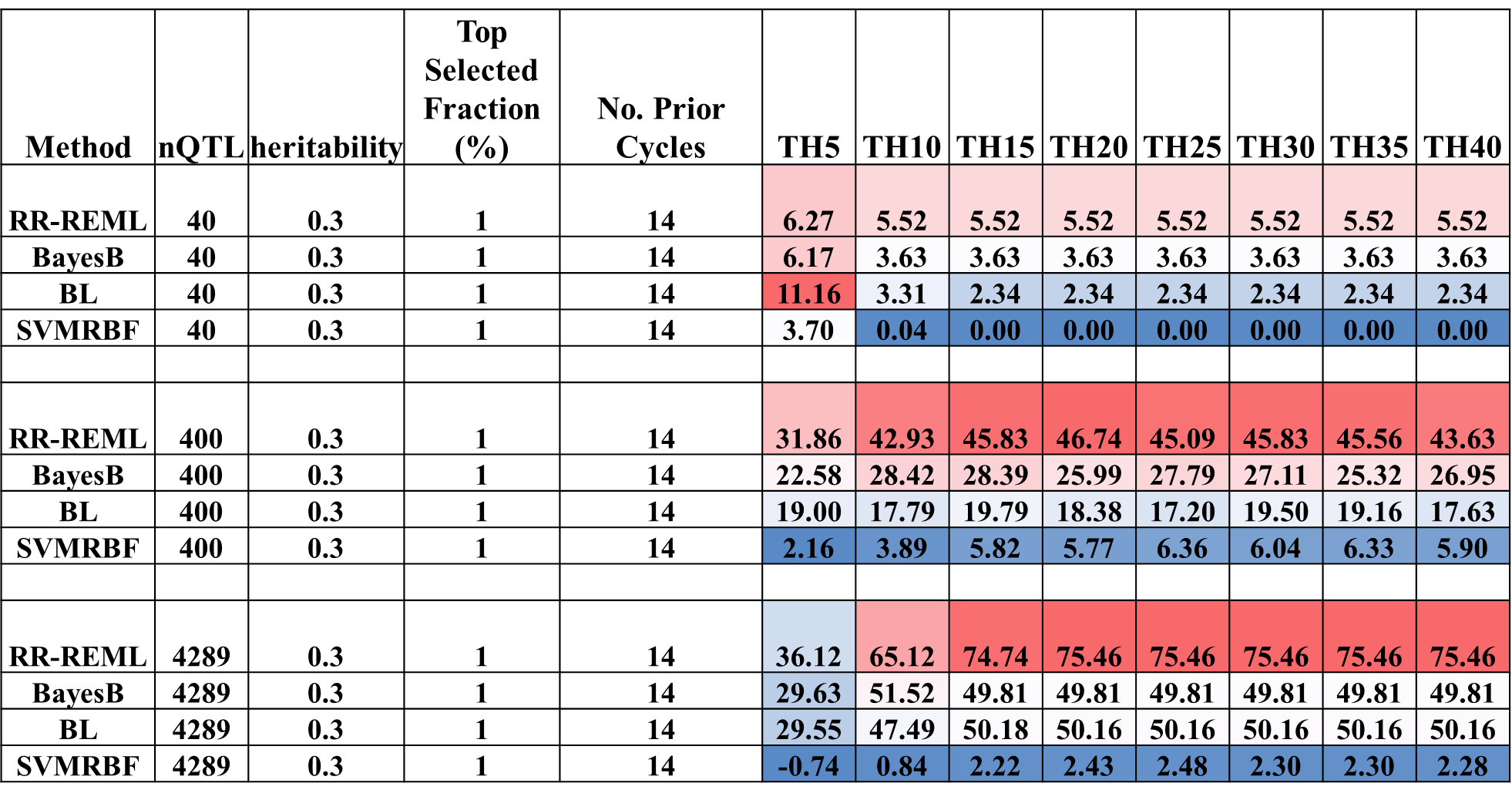

**Figure S42 Heat Map for Percent Gain in Responses Relative to GS without Updating for 0.3 H and Top 1% Selected Fraction** Heat map representing percent gain in response with respect to same type of GP model without updating in Recurrent GS for GP models updated with training data from 14 prior cycles in cycles one to forty in steps of five cycles for 40, 400 and 4289 simulated QTL responsible for 30% of phenotypic variability in the initial population. Blue to red shaded cells represent increasing gain in response relative to GS without updating. PS – Phenotypic Selection, RR-REML- Ridge Regression with Restricted Maximum Likelihood, BL – Bayes LASSO, and SVMRBF- Support Vector Machine with Radial Basis Kernel.

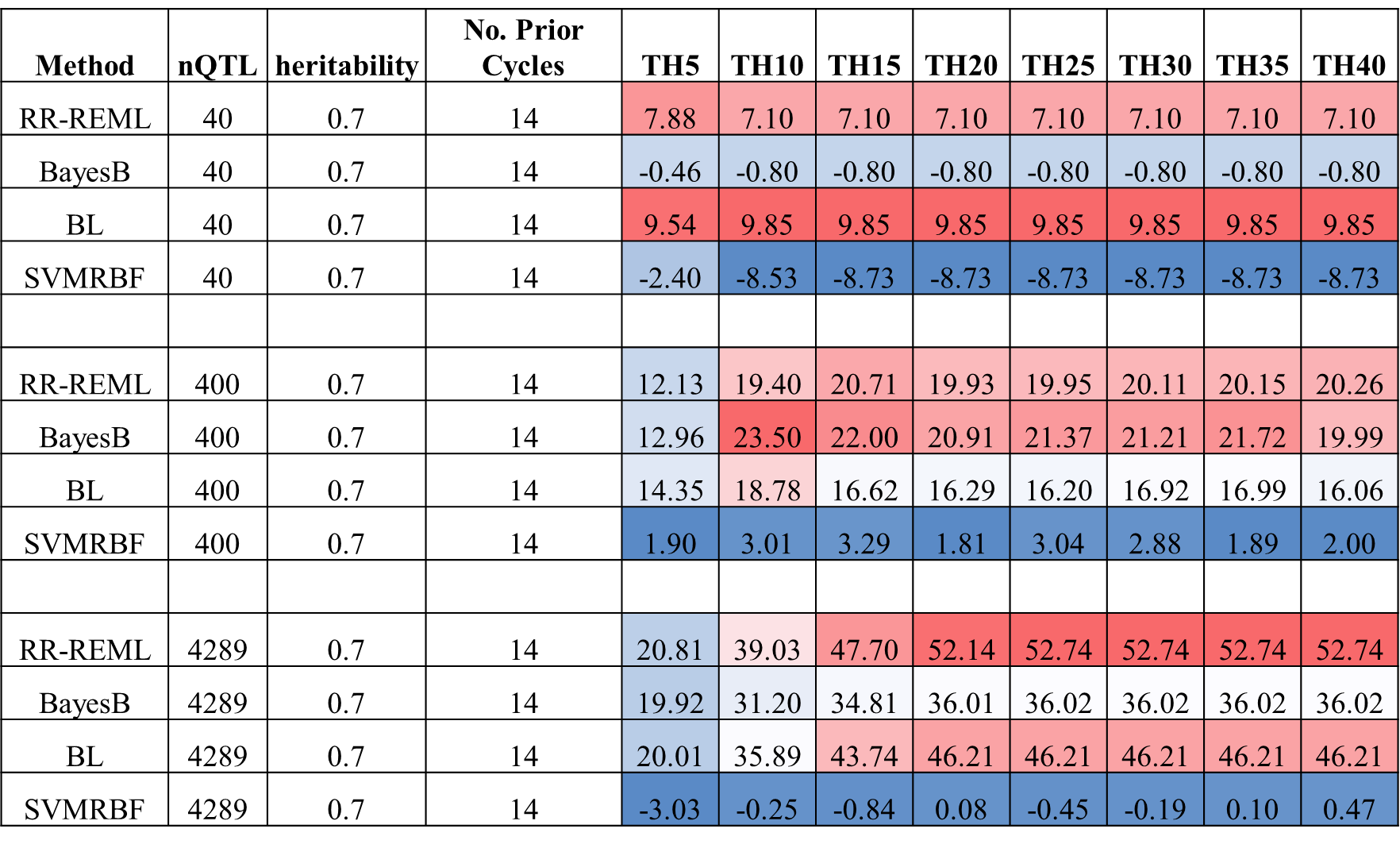

**Figure S43 Heat Map for Percent Gain in Responses Relative to GS without Updating for 0.7 H and Top 2.5% Selected Fraction** Heat map table for percent gain in response with respect to same type of GP model without updating in Recurrent GS for GP models updated with training data from 14 prior cycles in cycles one to forty in steps of five cycles for 40, 400 and 4289 simulated QTL responsible for 70% of phenotypic variability in the initial population. Blue to red shaded cells represent increasing gain in response relative to GS without updating. PS – Phenotypic Selection, RR-REML- Ridge Regression with Restricted Maximum Likelihood, BL – Bayes LASSO, and SVMRBF- Support Vector Machine with Radial Basis Kernel.

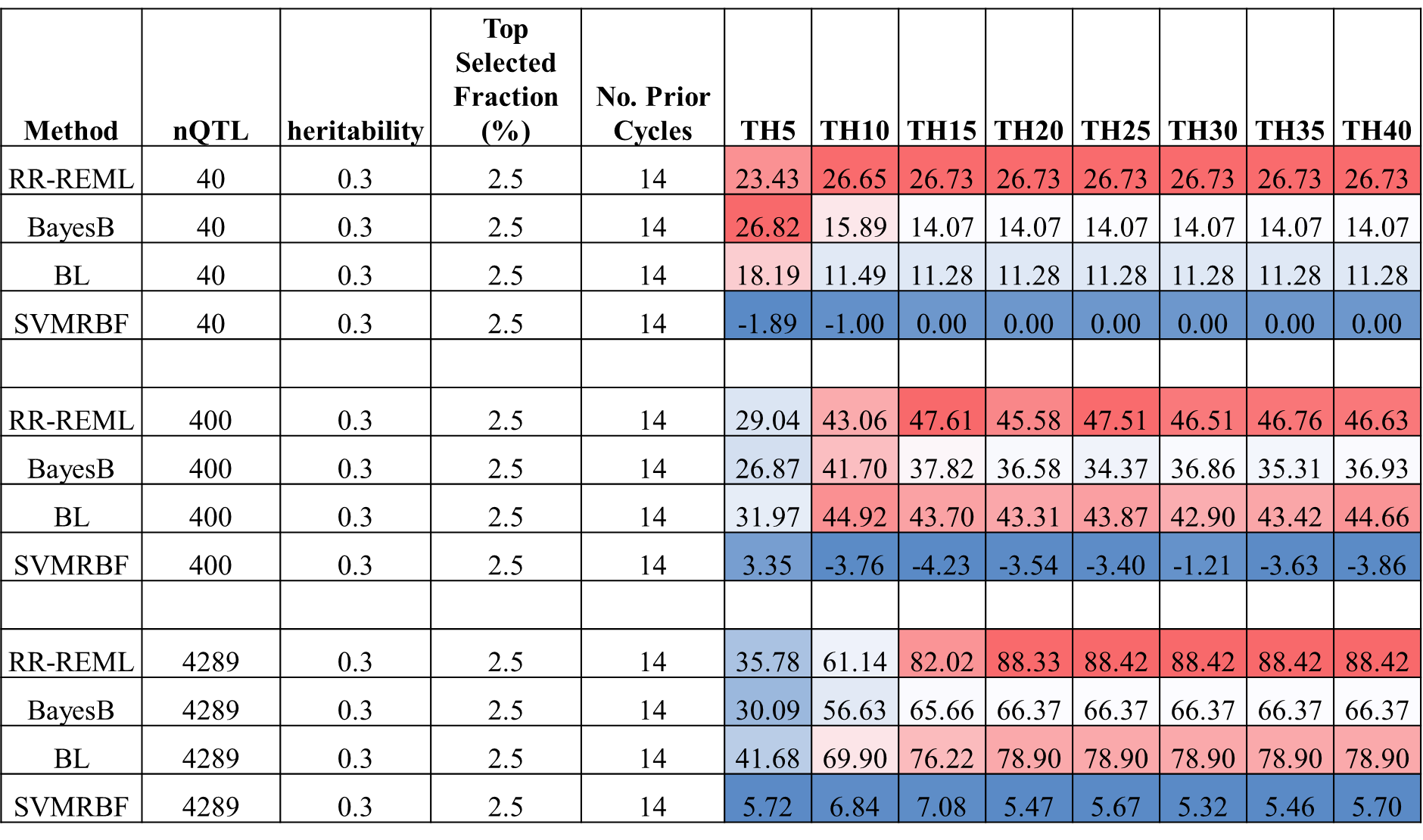

**Figure S44 Heat Map for Percent Gain in Responses Relative to GS without Updating for 0.3 H and Top 2.5% Selected Fraction:** Heat map table representing percent gain in response with respect to same type of GP model without updating in Recurrent GS for GP models updated with training data from up to 14 prior cycles in cycles one to forty in steps of five cycles for 40, 400 and 4289 simulated QTL responsible for 30% of phenotypic variability in the initial population. Blue to red shaded cells represent increasing gain in response relative to GS without updating. PS – Phenotypic Selection, RR-REML- Ridge Regression with Restricted Maximum Likelihood, BL – Bayes LASSO, and SVMRBF- Support Vector Machine with Radial Basis Kernel.

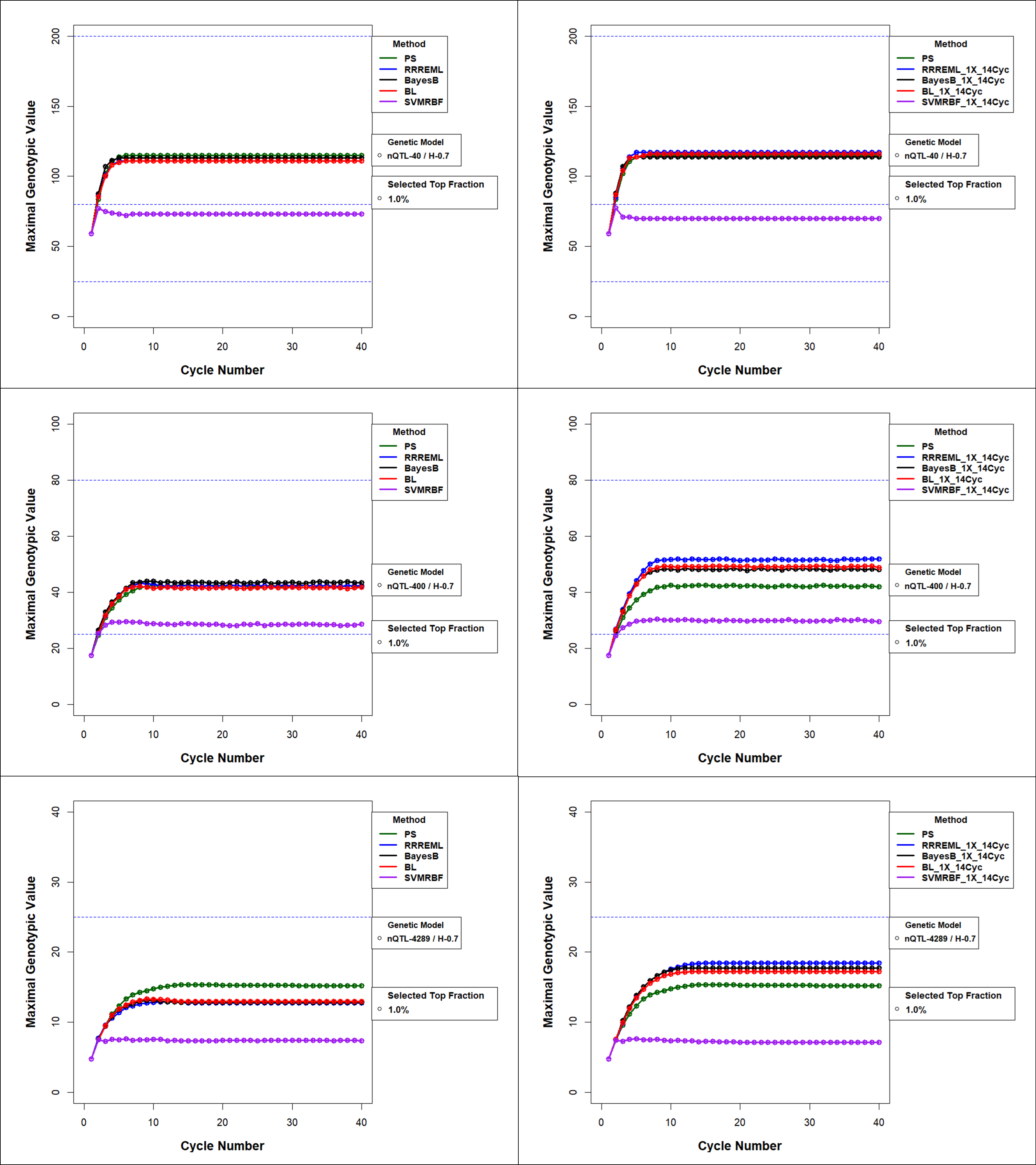

**Figure S45 Maximal Genotypic Value for Comparison of GS methods with and without Updating for 0.7 H and Top 1% Selected Fraction** Maximum Genotypic Value (Mgv) in recurrent GS and PS without model updating (left panels) and with model updating (right panels) using training data from up to 14 prior cycles for the four GP models. PS has no updating and hence does not change between the left and right panels for a) 40 simulated QTL (top), b) 400 simulated QTL (middle), and c) 4289 simulated QTL (bottom) responsible for 70% of phenotypic variability in the initial population and top 1% selected fraction. PS – Phenotypic Selection, RR-REML- Ridge Regression with Restricted Maximum Likelihood, BL – Bayes LASSO, and SVMRBF- Support Vector Machine with Radial Basis Kernel.

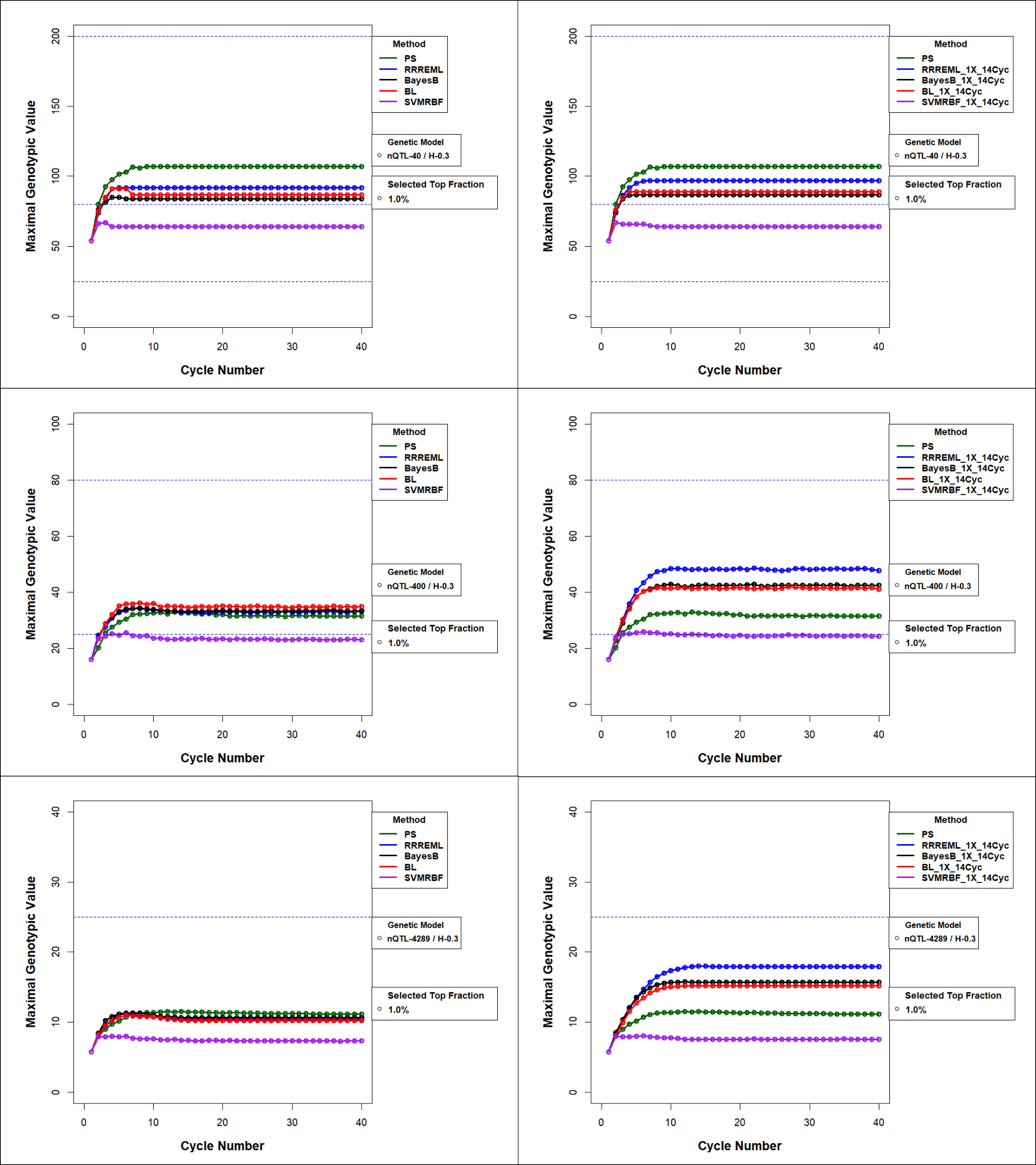

**Figure S46 Maximal Genotypic Value for Comparison of GS methods with and without Updating for 0.3 H and Top 1% Selected Fraction** Maximal genotypic value (Mgv) in recurrent GS and PS without model updating (left panels) and with model updating (right panels) using training data from up to 14 prior cycles for the four GP models. PS has no updating and hence does not change between the left and right panels for a) 40 simulated QTL (top), b) 400 simulated QTL (middle), and c) 4289 simulated QTL (bottom) responsible for 30% of phenotypic variability in the initial population and top 1% selected fraction. PS – Phenotypic Selection, RR-REML- Ridge Regression with Restricted Maximum Likelihood, BL – Bayes LASSO, and SVMRBF- Support Vector Machine with Radial Basis Kernel.

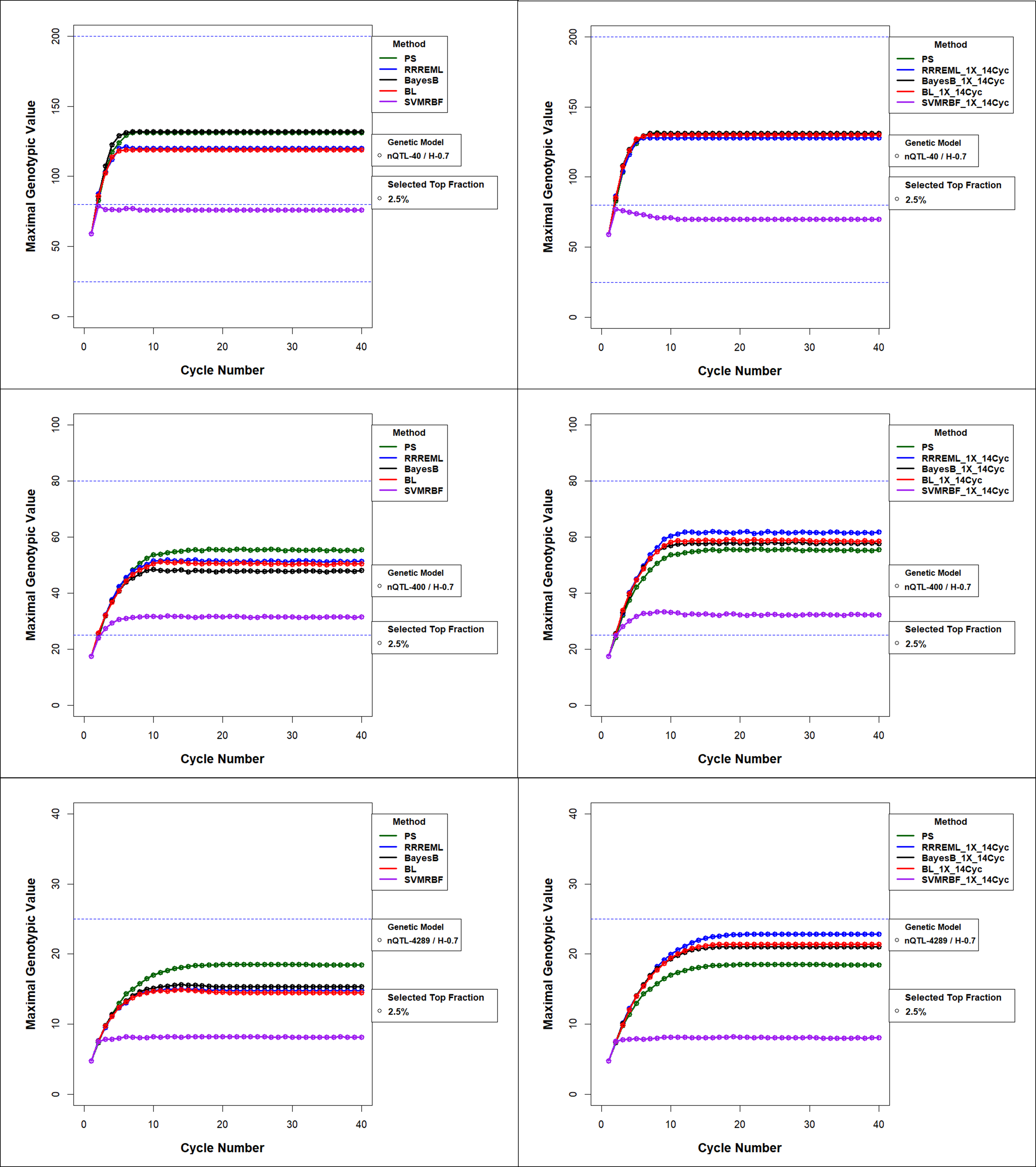

**Figure S47 Maximal Genotypic Value for Comparison of GS methods with and without Updating for 0.7 H and Top 2.5% Selected Fraction** Maximal genotypic value in recurrent GS and PS without model updating (left panels) and with model updating (right panels) using training data from up to 14 prior cycles for the four GP models. PS has no updating and hence does not change between the left and right panels. All treatment combinations have top 2.5% selected fraction with 40 simulated QTL (top), 400 simulated QTL (middle), and 4289 simulated QTL (bottom) responsible for 70% of phenotypic variability in the initial population. PS – Phenotypic Selection, RR-REML- Ridge Regression with Restricted Maximum Likelihood, BL – Bayes LASSO, and SVMRBF- Support Vector Machine with Radial Basis Kernel.

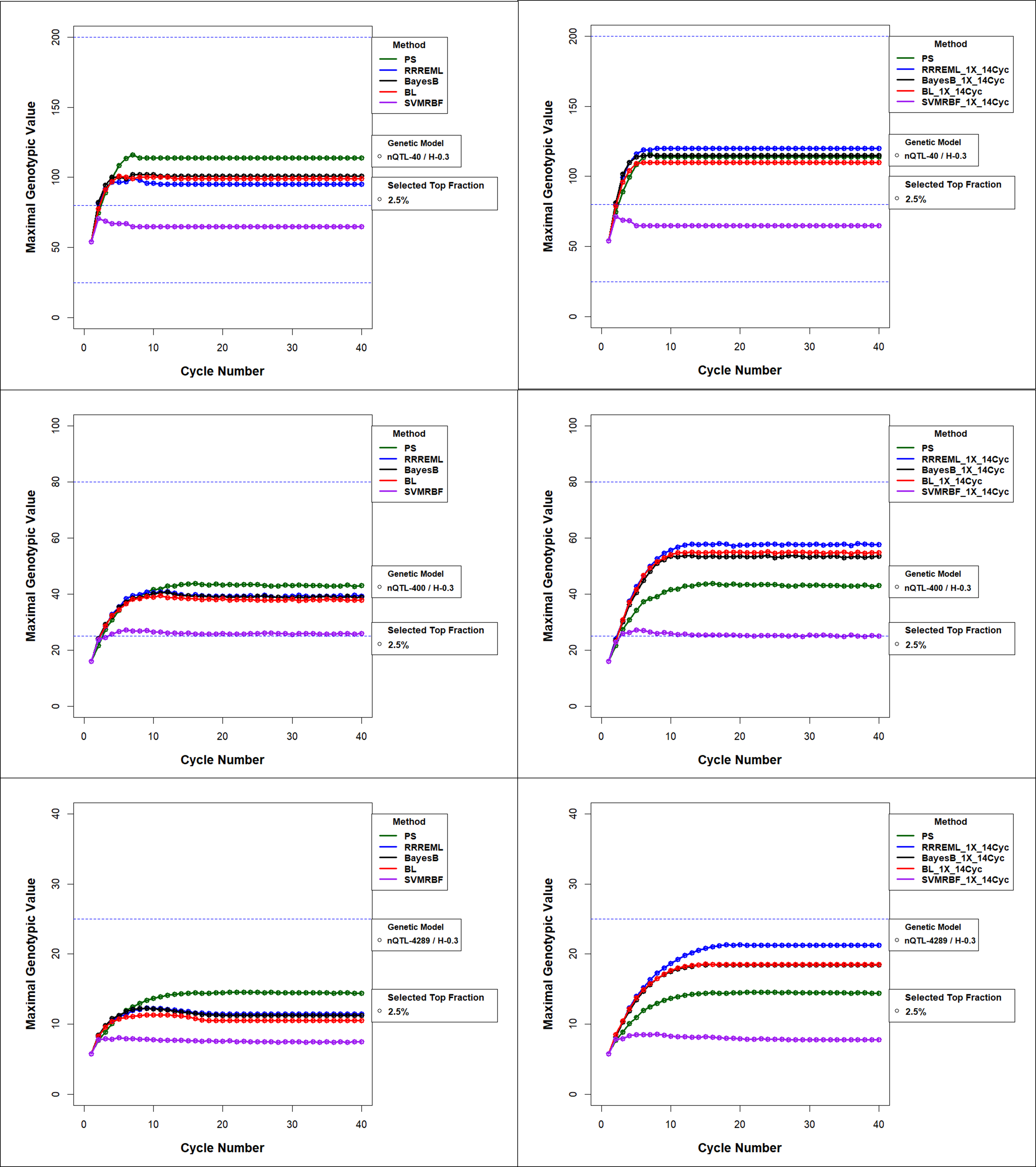

**Figure S48 Maximal Genotypic Value for Comparison of GS methods with and without Updating for 0.3 H and Top 2.5% Selected Fraction** Maximal genotypic value in recurrent GS and PS without (left panels) and with model updating (right panels) using training data from up to 14 prior cycles for the four GP models. PS has no updating and hence does not change between the left and right panels. All treatments have top 2.5% selected fraction with 40 simulated QTL (top), 400 simulated QTL (middle), and 4289 simulated QTL (bottom) responsible for 30% of phenotypic variability in the initial population, PS – Phenotypic Selection, RR-REML- Ridge Regression with Restricted Maximum Likelihood, BL – Bayes LASSO, and SVMRBF- Support Vector Machine with Radial Basis Kernel.

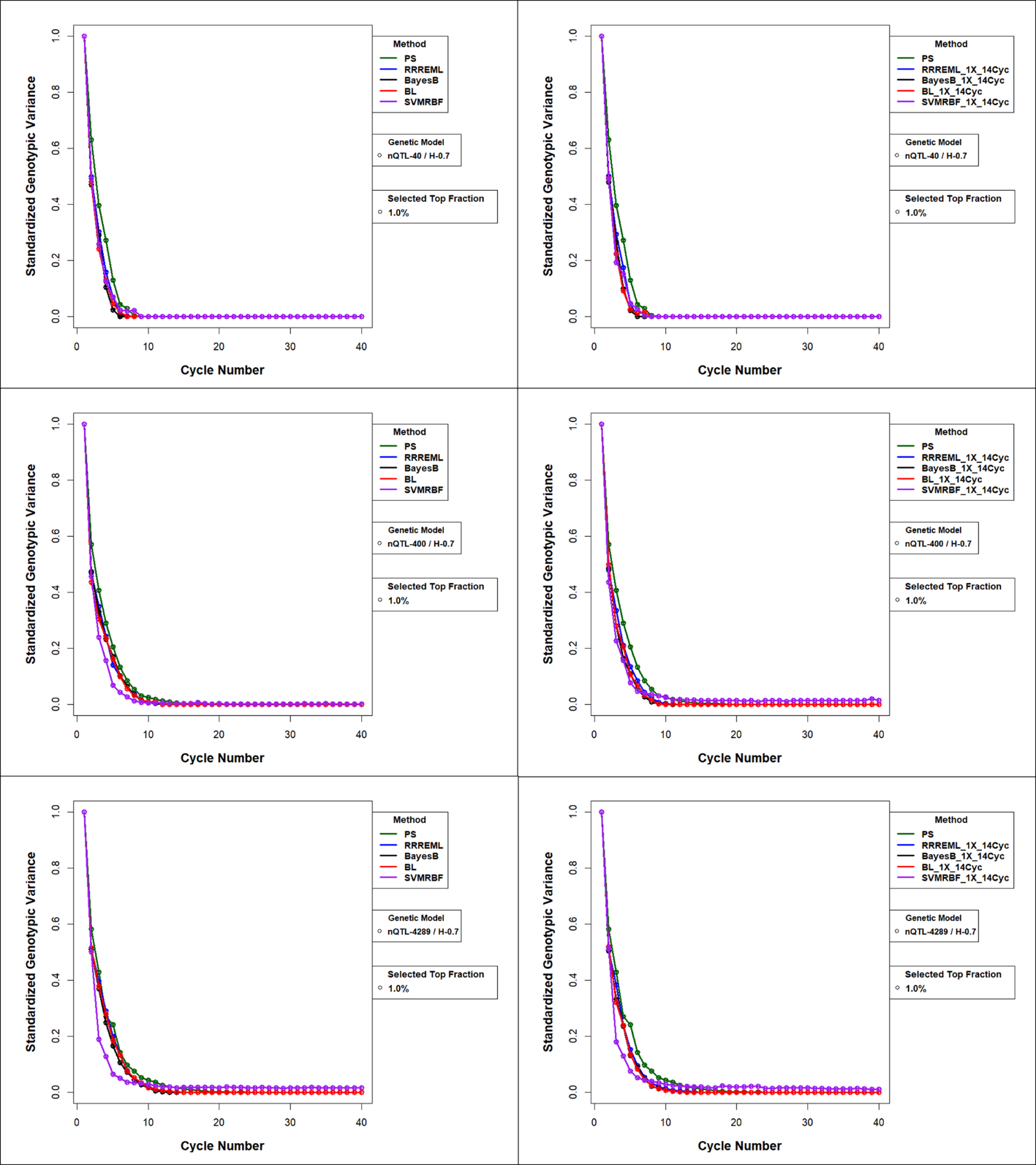

**Figure S49 Standardized Genotypic Variance (Sgv) for Comparison of GS methods with and without Updating for 0.7 H and Top 1% Selected Fraction** Standardized genotypic variance with model updating with training data from up to 14 prior cycles for the four GP models. PS has no updating and hence is the same in the left and right panels. All treatment factors top 1% selected fraction with 40 simulated QTL (top), 400 QTL simulated (middle), and 4289 simulated QTL (bottom) responsible for 70% of phenotypic variability in the initial population. PS – Phenotypic Selection, RR-REML- Ridge Regression with Restricted Maximum Likelihood, BL – Bayes LASSO, and SVMRBF- Support Vector Machine with Radial Basis Kernel.

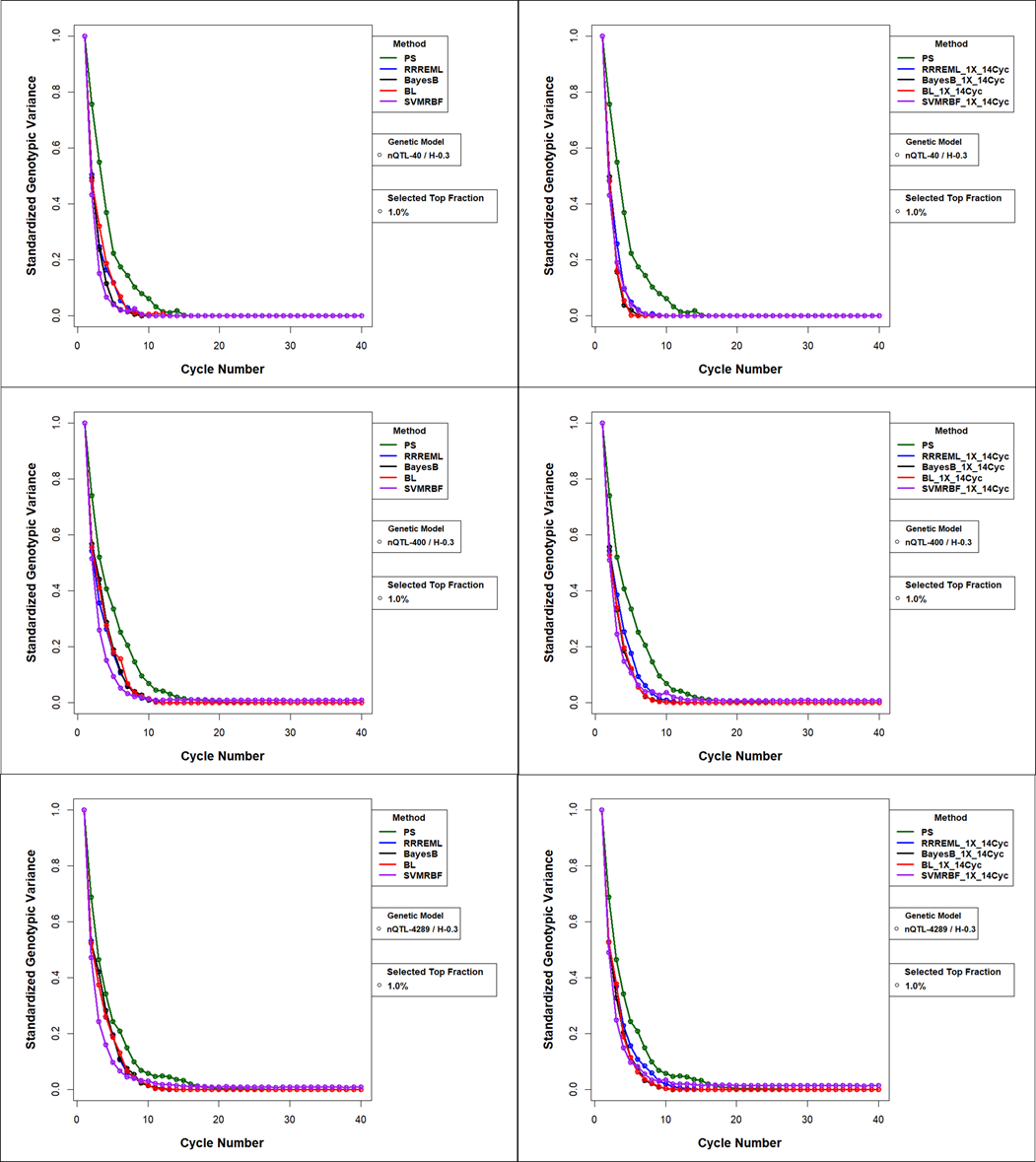

**Figure S50 Standardized Genotypic Variance (Sgv) for Comparison of GS methods with and without Updating for 0.3 H and Top 1% Selected Fraction** Standardized genotypic variance with model updating with training data from up to 14 prior cycles for the four GP models. PS has no updating and hence is the same in the left and right panels. All treatment factors have top 1% selected fraction with 40 simulated QTL (top), 400 simulated QTL (middle), and 4289 simulated QTL (bottom) responsible for 30% of phenotypic variability in the initial population. PS – Phenotypic Selection, RR-REML- Ridge Regression with Restricted Maximum Likelihood, BL – Bayes LASSO, and SVMRBF- Support Vector Machine with Radial Basis Kernel.

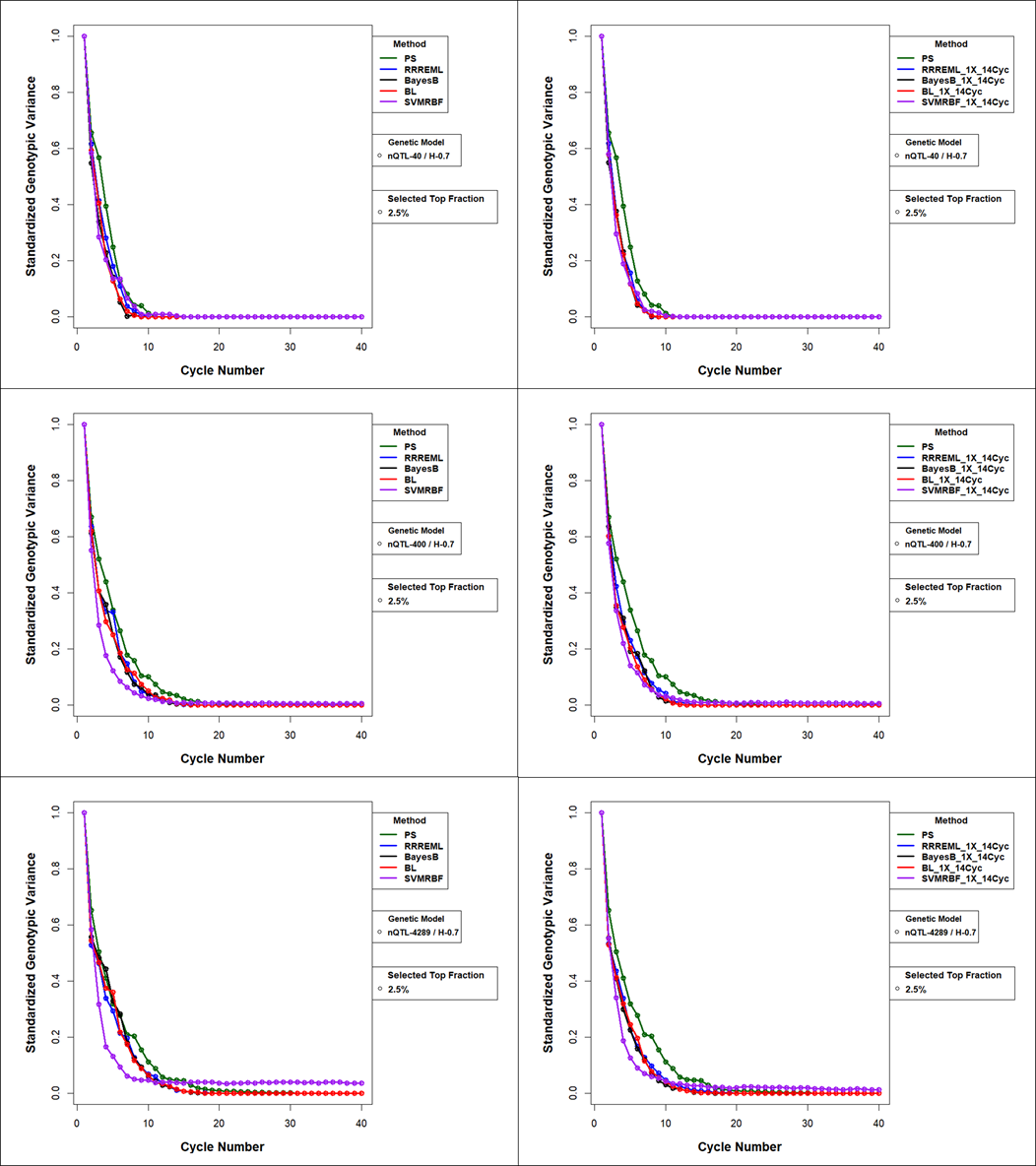

**Figure S51 Standardized Genotypic Variance (Sgv) for Comparison of GS methods with and without Updating for 0.7 H and Top 2.5% Selected Fraction** Standardized genotypic variance with model updating with training data from up to 14 prior cycles for the four GP models. PS has no updating and hence is the same in the left and right panels. All treatment factors have top 2.5% selected fraction with 40 simulated QTL (top), 400 simulated QTL (middle), and 4289 simulated QTL (bottom) responsible for 70% of phenotypic variability in the initial population. PS – Phenotypic Selection, RR-REML- Ridge Regression with Restricted Maximum Likelihood, BL – Bayes LASSO, and SVMRBF- Support Vector Machine with Radial Basis Kernel.

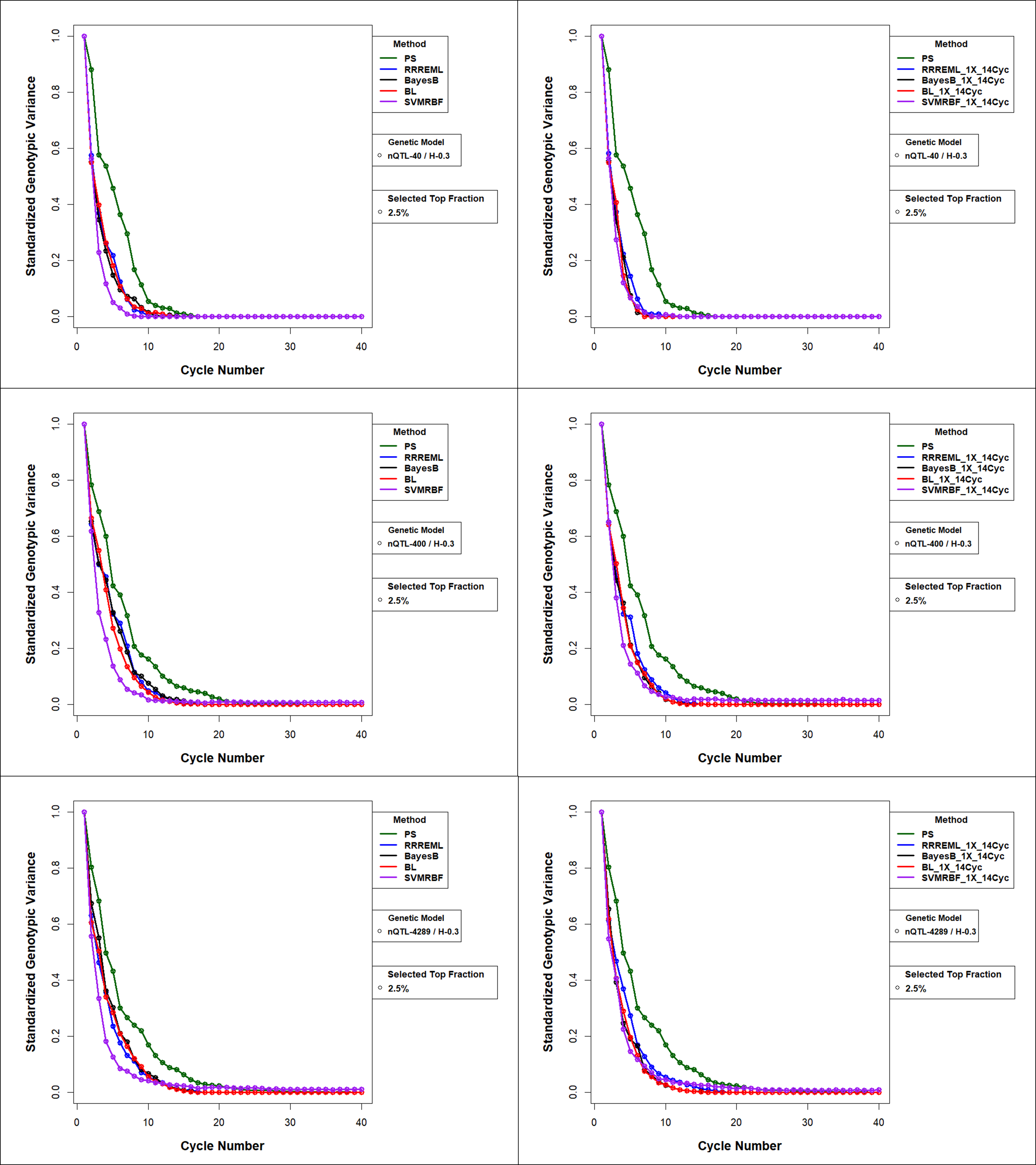

**Figure S52 Standardized Genotypic Variance (Sgv) for Comparison of GS methods with and without Updating for 0.3 H and Top 2.5% Selected Fraction** Standardized genotypic variance with model updating with training data from up to 14 prior cycles for the four GP models. All treatment factors have top 2.5% selected fraction with 40 QTL (top), 400 QTL (middle), and 4289 QTL (bottom) responsible for 30% of phenotypic variability in the initial population. PS – Phenotypic Selection, RR-REML- Ridge Regression with Restricted Maximum Likelihood, BL – Bayes LASSO, and SVMRBF- Support Vector Machine with Radial Basis Kernel.

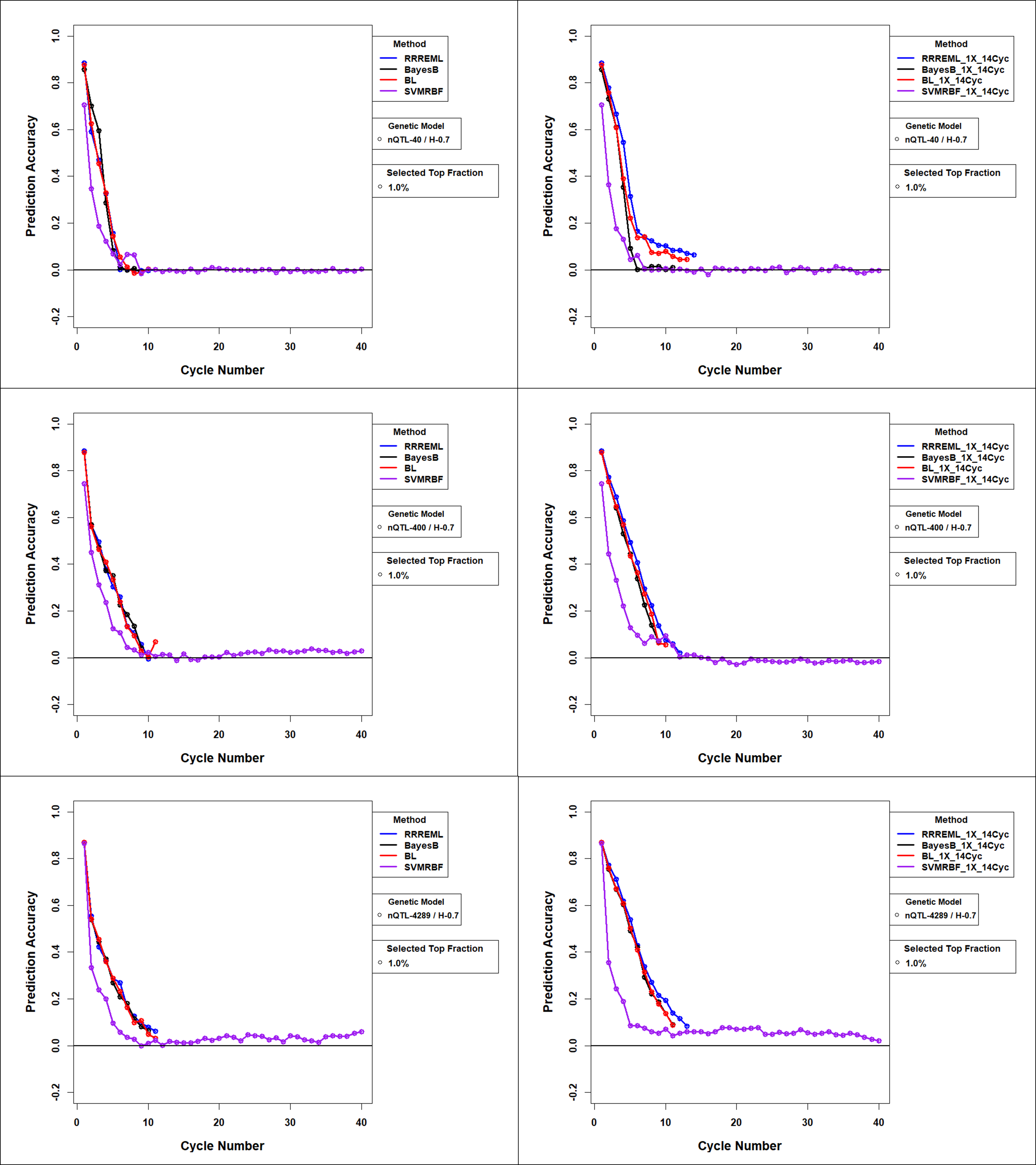

**Figure S53 Estimated Prediction Accuracies** **for Comparison of GS methods with and without Updating for 0.7 H and Top 1% Selected Fraction:** Estimated prediction accuracies with model updating with training data from up to 14 prior cycles for the four GP models. All treatment factors have top 1.0% selected fraction with 40 simulated QTL (top), 400 simulated QTL (middle), and 4289 simulated QTL (bottom) responsible for 70% of phenotypic variability in the initial population. RR-REML- Ridge Regression with Restricted Maximum Likelihood, BL – Bayes LASSO, and SVMRBF- Support Vector Machine with Radial Basis Kernel.

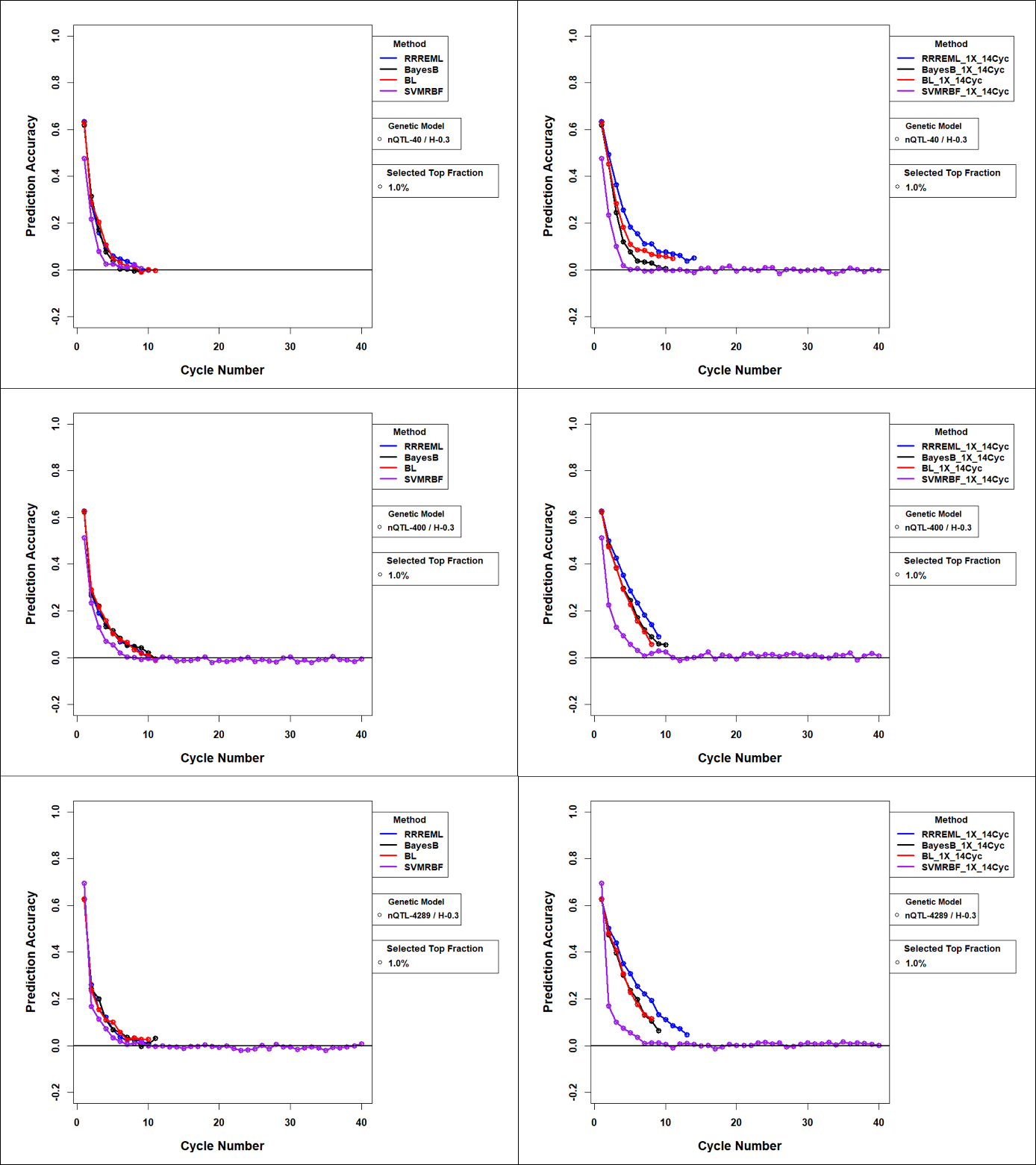

**Figure S54 Estimated Prediction Accuracies** **for Comparison of GS methods with and without Updating for 0.3 H and Top 1% Selected Fraction:** Estimated prediction accuracies with model updating with training data from up to 14 prior cycles for the four GP models. All conditions have top 1.0% selected fraction with 40 simulated QTL (top), 400 simulated QTL (middle), and 4289 simulated QTL (bottom) responsible for 30% of phenotypic variability in the initial population. RR-REML- Ridge Regression with Restricted Maximum Likelihood, BL – Bayes LASSO, and SVMRBF- Support Vector Machine with Radial Basis Kernel.

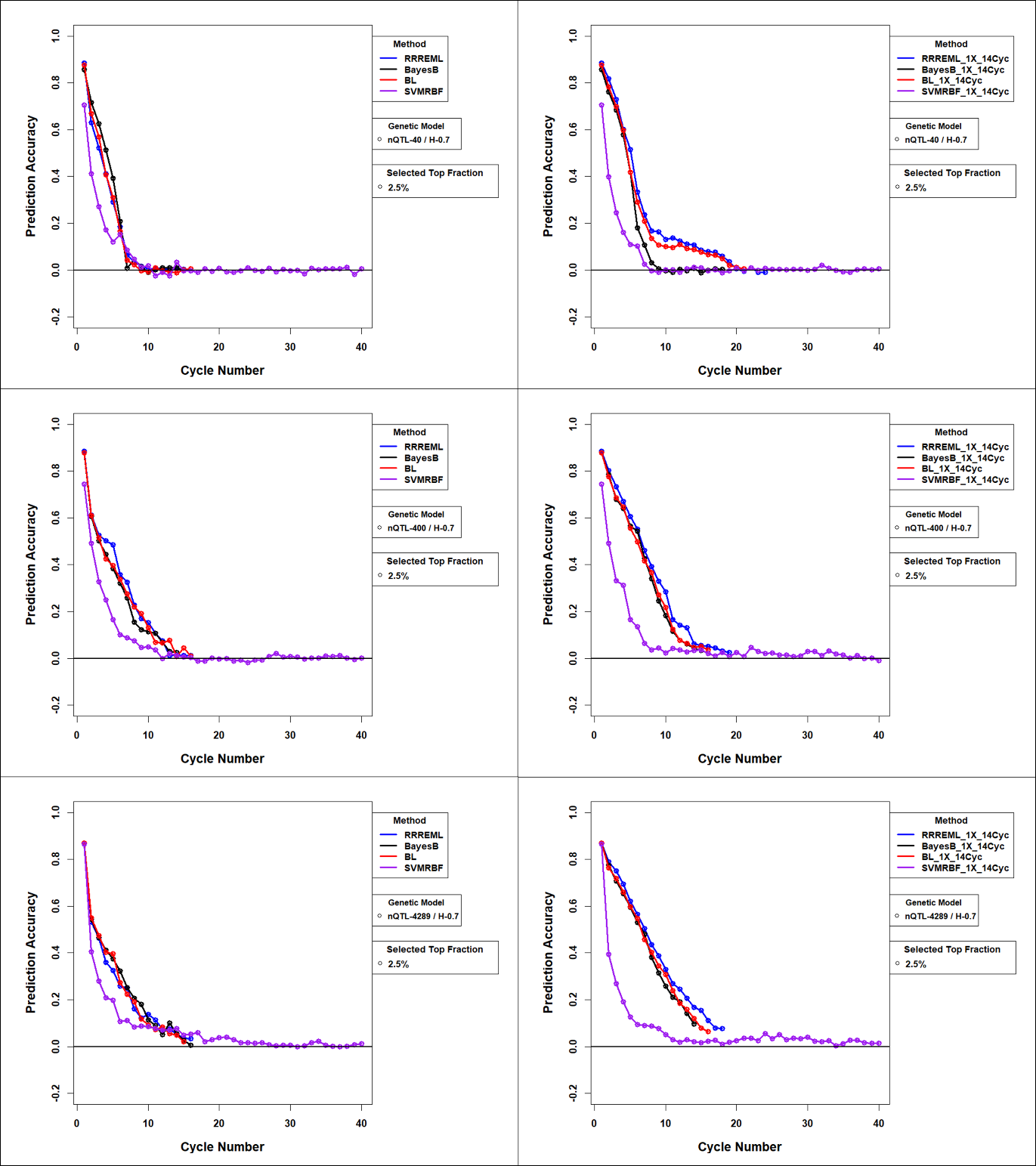

**Figure S55 Estimated Prediction Accuracies** **for Comparison of GS methods with and without Updating for 0.7 H and Top 2.5% Selected Fraction:** Estimated prediction accuracies with model updating with training data from up to 14 prior cycles for the four GP models. All conditions have narrow sense heritability 0.7 and top 2.5 % selected fraction with 40 simulated QTL (top left), 400 simulated QTL (top right), and 4289 simulated QTL (bottom) responsible for 70% of phenotypic variability in the initial population. PS – Phenotypic Selection, RR-REML- Ridge Regression with Restricted Maximum Likelihood, BL – Bayes LASSO, and SVMRBF- Support Vector Machine with Radial Basis Kernel.

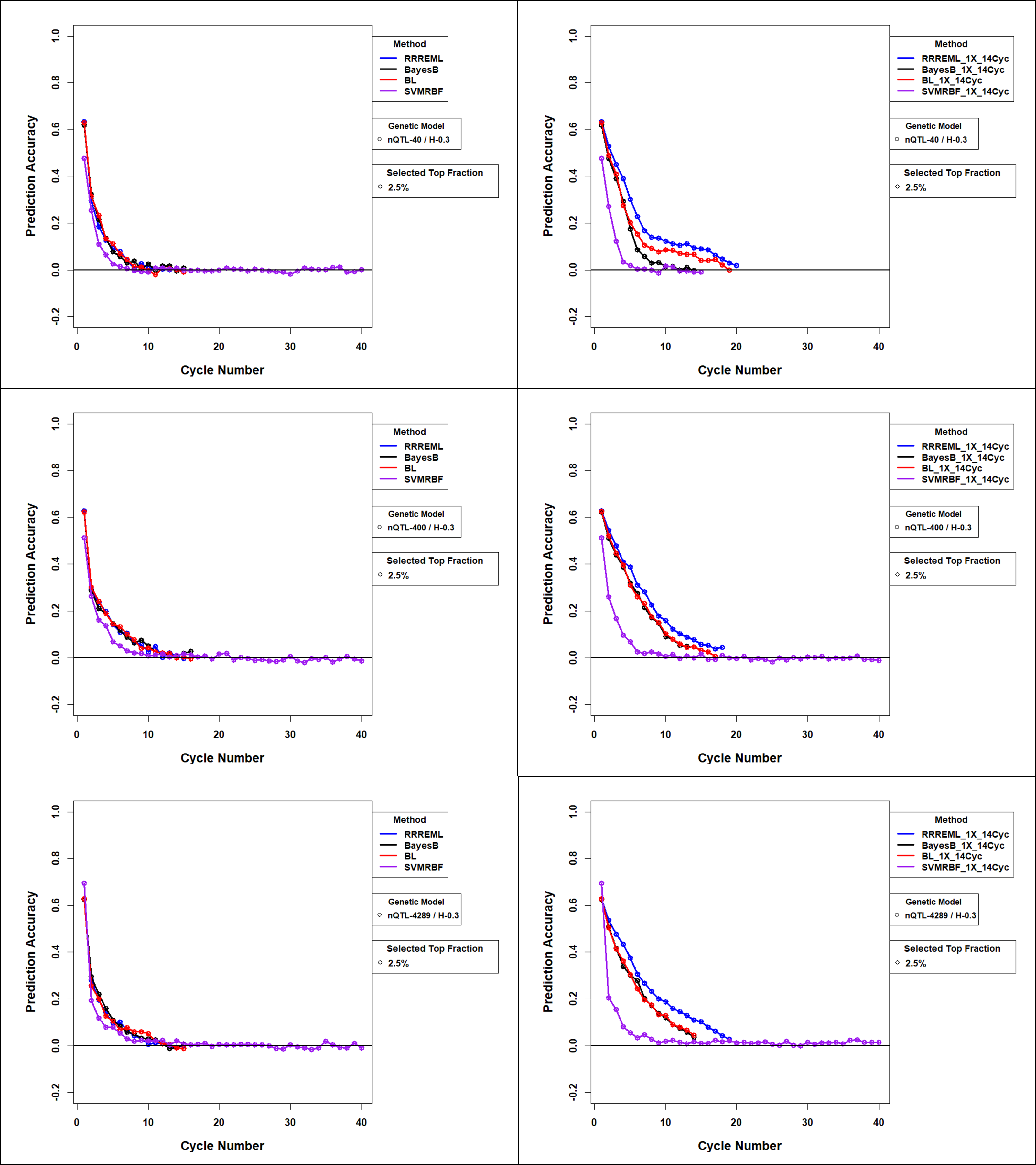

**Figure S56 Estimated Prediction Accuracies** **for Comparison of GS methods with and without Updating for 0.3 H and Top 2.5% Selected Fraction:** Estimated prediction accuracies with model updating with training data from up to 14 prior cycles for the four GP models. All conditions have top 2.5% selected fraction with 40 simulated QTL (top), 400 simulated QTL (middle), and 4289 simulated QTL (bottom) responsible for 30% of phenotypic variability in the initial population. PS – Phenotypic Selection, RR-REML- Ridge Regression with Restricted Maximum Likelihood, BL – Bayes LASSO, and SVMRBF- Support Vector Machine with Radial Basis Kernel.

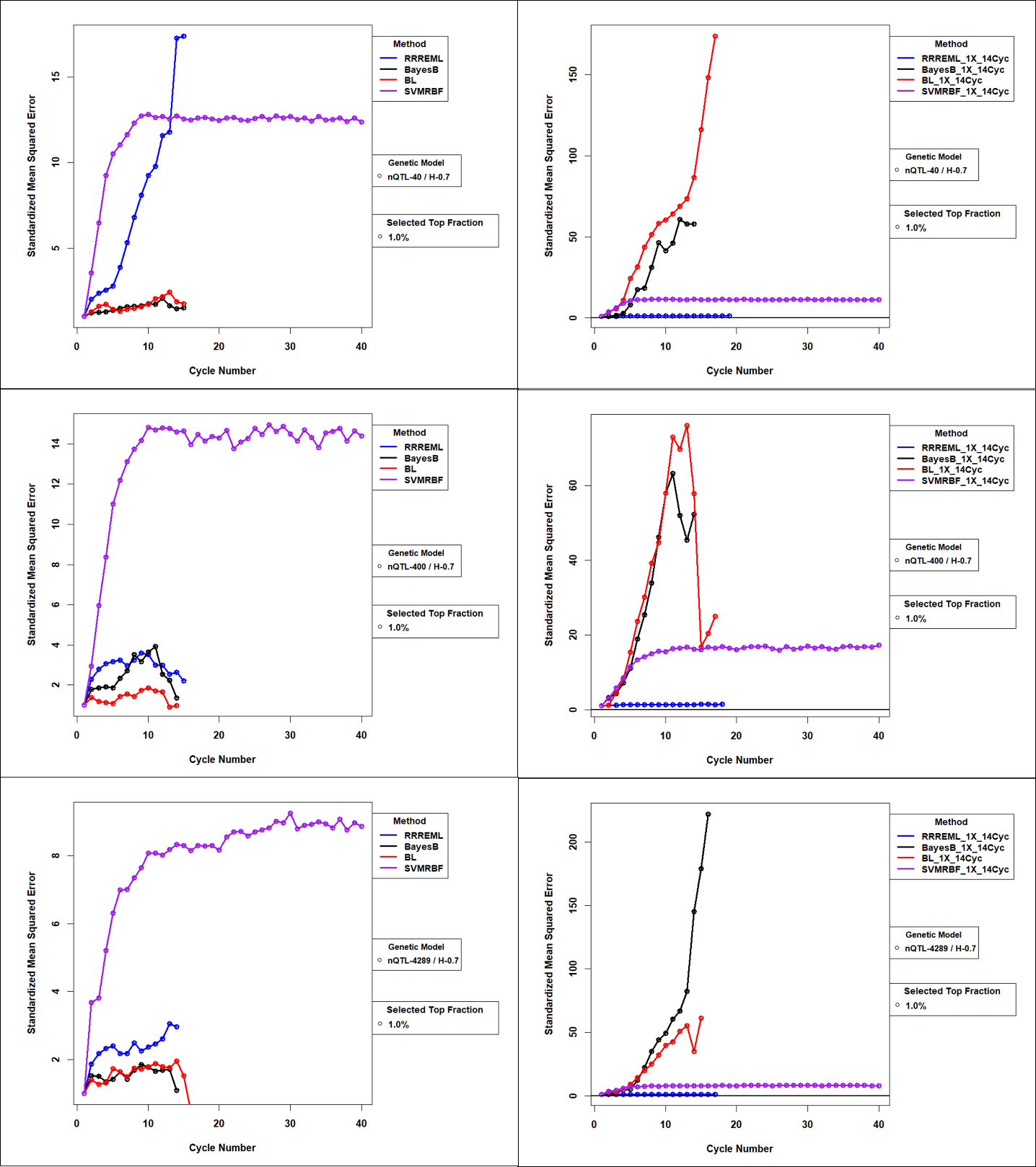

**Figure 57 Standardized Mean Squared Error** **for Comparison of GS methods with and without Updating for 0.7 H and Top 1% Selected Fraction:** Mean Squared Error of GP models with updates to the training sets used in genomic prediction (GP) models. Standardized MSE (>=1) is estimated as the ratio of MSE for GP models in cycle ‘c’ to MSE for GP models trained with founder population of RILs. While MSE for RRREML model were lesser with model updating, MSE for bayesian methods increased orders of magnitude in late cycles of selection with model updating. SVMEBF showed relatively constant MSE with and without updating. Training data from up to 14 prior selection cycles were used to update all four GP models for 40 QTL (top), 400 QTL (middle) and 4289 QTL (bottom) responsible for 70% of phenotypic variability in the initial population and top 1% of RILs with the greatest predicted values. RR-REML- Ridge Regression with Restricted Maximum Likelihood, BL – Bayes LASSO, and SVMRBF- Support Vector Machine with Radial Basis Kernel.

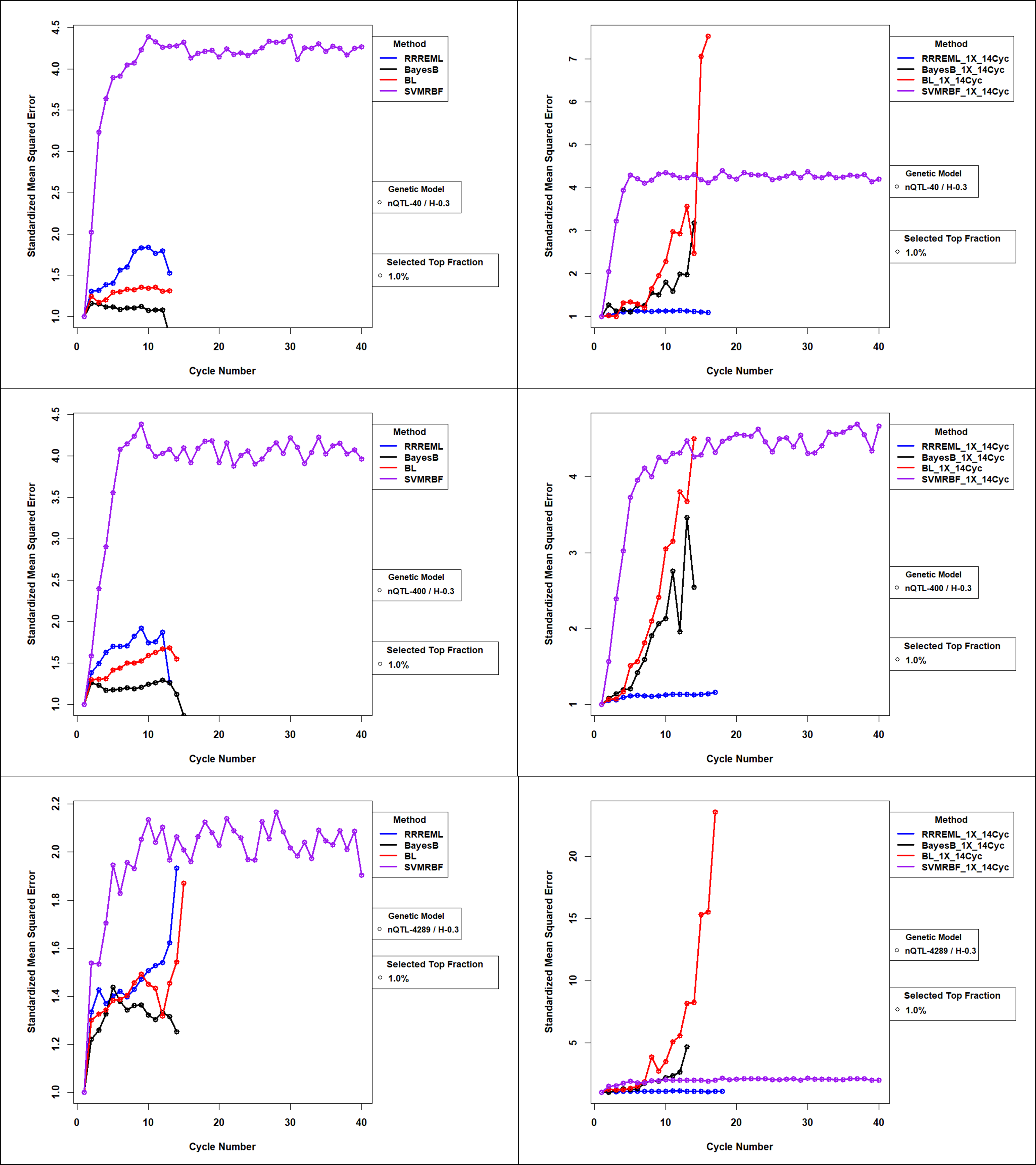

**Figure 58 Standardized Mean Squared Error** **for Comparison of GS methods with and without Updating for 0.3 H and Top 1.0 % Selected Fraction:** Mean Squared Error of GP models with updates to the training sets used in genomic prediction (GP) models. Standardized MSE (>=1) is estimated as the ratio of MSE for GP models in cycle ‘c’ to MSE for GP models trained with founder population of RILs. While MSE for RRREML model were lesser with model updating, MSE for bayesian methods increased orders of magnitude in late cycles of selection with model updating. SVMEBF showed relatively constant MSE with and without updating. Training data from up to 14 prior selection cycles were used to update all four GP models for 40 QTL (top), 400 QTL (middle) and 4289 QTL (bottom) responsible for 30% of phenotypic variability in the initial population and top 1.0% of RILs with the greatest predicted values. RR-REML- Ridge Regression with Restricted Maximum Likelihood, BL – Bayes LASSO, and SVMRBF- Support Vector Machine with Radial Basis Kernel.

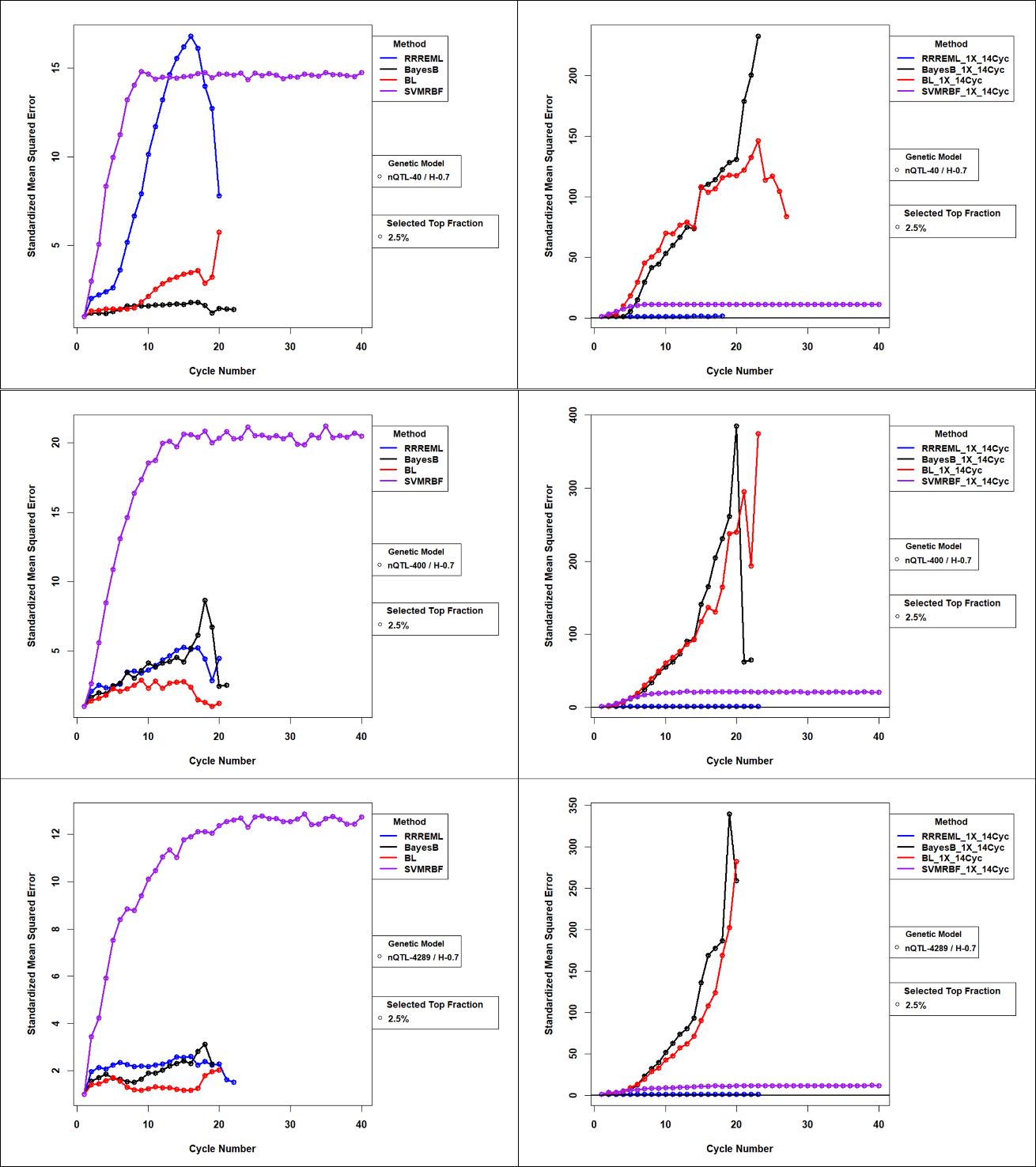

**Figure 59 Standardized Mean Squared Error** **for Comparison of GS methods with and without Updating for 0.7 H and Top 2.5% Selected Fraction:** Mean Squared Error of GP models with updates to the training sets used in genomic prediction (GP) models. Standardized MSE (>=1) is estimated as the ratio of MSE for GP models in cycle ‘c’ to MSE for GP models trained with founder population of RILs. While MSE for RRREML model were lesser with model updating, MSE for bayesian methods increased orders of magnitude in late cycles of selection with model updating. SVMEBF showed relatively constant MSE with and without updating. Training data from up to 14 prior selection cycles were used to update all four GP models for 40 QTL (top), 400 QTL (middle) and 4289 QTL (bottom) responsible for 70% of phenotypic variability in the initial population and top 2.5% of RILs with the greatest predicted values. RR-REML- Ridge Regression with Restricted Maximum Likelihood, BL – Bayes LASSO, and SVMRBF- Support Vector Machine with Radial Basis Kernel.

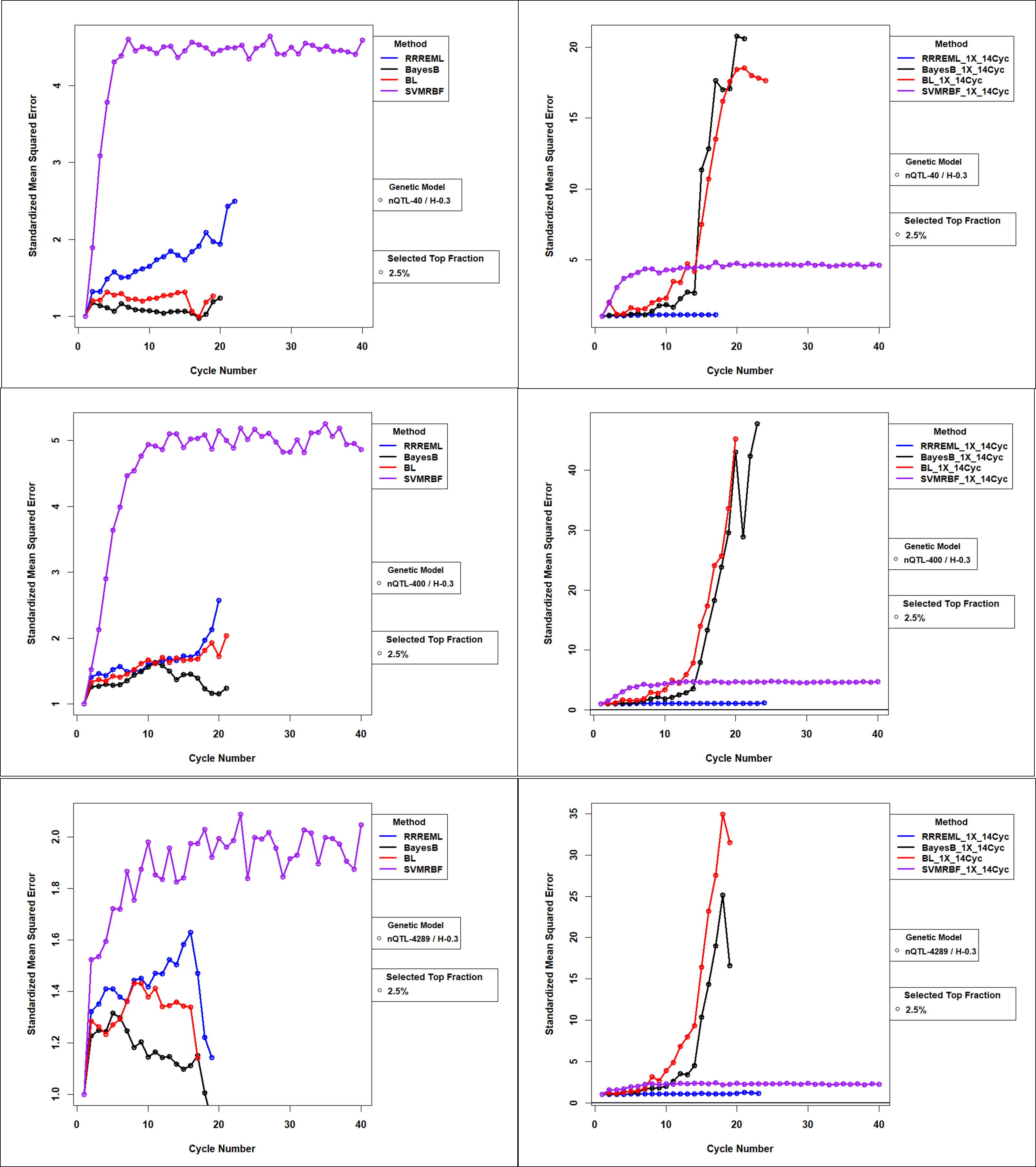

**Figure 60 Standardized Mean Squared Error** **for Comparison of GS methods with and without Updating for 0.3 H and Top 2.5% Selected Fraction:** Mean Squared Error of GP models with updates to the training sets used in genomic prediction (GP) models. Standardized MSE (>=1) is estimated as the ratio of MSE for GP models in cycle ‘c’ to MSE for GP models trained with founder population of RILs. While MSE for RRREML model were lesser with model updating, MSE for bayesian methods increased orders of magnitude in late cycles of selection with model updating. SVMEBF showed relatively constant MSE with and without updating. Training data from up to 14 prior selection cycles were used to update all four GP models for 40 QTL (top), 400 QTL (middle) and 4289 QTL (bottom) responsible for 30% of phenotypic variability in the initial population and top 2.5 % of RILs with the greatest predicted values. RR-REML- Ridge Regression with Restricted Maximum Likelihood, BL – Bayes LASSO, and SVMRBF- Support Vector Machine with Radial Basis Kernel.
